# Dendritic excitations govern back-propagation via a spike-rate accelerometer

**DOI:** 10.1101/2023.06.02.543490

**Authors:** Pojeong Park, David Wong-Campos, Daniel G. Itkis, Byung Hun Lee, Yitong Qi, Hunter Davis, Benjamin Antin, Amol Pasarkar, Jonathan B. Grimm, Sarah E. Plutkis, Katie L. Holland, Liam Paninski, Luke D. Lavis, Adam E. Cohen

## Abstract

Dendrites on neurons support nonlinear electrical excitations, but the computational significance of these events is not well understood. We developed molecular, optical, and analytical tools to map sub-millisecond voltage dynamics throughout the dendritic trees of CA1 pyramidal neurons under diverse optogenetic and synaptic stimulus patterns, in acute brain slices. We observed history-dependent spike back-propagation in distal dendrites, driven by locally generated Na^+^ spikes (dSpikes). Dendritic depolarization created a transient window for dSpike propagation, opened by A-type K_V_ channel inactivation, and closed by slow Na_V_ inactivation. Collisions of dSpikes with synaptic inputs triggered calcium channel and N-methyl-D-aspartate receptor (NMDAR)-dependent plateau potentials, with accompanying complex spikes at the soma. This hierarchical ion channel network acts as a spike-rate accelerometer, providing an intuitive picture of how dendritic excitations shape associative plasticity rules.

## Introduction

Dendrites carry information in two directions: they carry synaptic inputs to the soma, and they carry back-propagating action potentials (bAPs) from the soma into the dendritic tree. Dendritic integration determines when a neuron spikes, and is the basis of rapid single-cell information processing. Dendritic back-propagation conveys information about spike times to the synapses, leading to synaptic plasticity and slow changes in neuronal input-output functions.^1–3^ Electrical signals propagating in either direction encounter a diverse set of ion channels which give the dendrites nonlinear and history-dependent excitability.^4–6^ Simulation studies have proposed diverse computational roles of local excitations in dendritic integration.^7–9^ Despite many experimental studies showing that bAPs also activate dendritic excitations, there has not been a clear picture of the computational significance of these processes.

Spruston and colleagues first used dendritic patch clamp measurements to characterize history-dependent bAP propagation into distal dendrites of CA1 pyramidal cells.^10^ A-type potassium channels, localized in these distal dendrites, suppressed bAP propagation and prevented initiation of action potentials within the dendrites.^11^ Distal depolarization inactivated these channels, permitting bAPs to propagate further into the distal dendrites,^12,13^ and these bAPs could then engage dendritic Na_V_ channels leading to dichotomous success or failure of bAPs to trigger dendritic sodium spikes (dSpikes).^14,15^ Repetitive trains of bAPs drove slow inactivation of the dendritic Na_V_ channels and caused a gradual decrease in bAP penetration into distal dendrites.^16–18^ Thus the success or failure of a bAP to penetrate into the dendritic tree depends in a complex way on the history of neuronal spiking and dendritic voltage.

Engagement of NMDA receptors by excitatory synaptic inputs qualitatively changes the excitability properties of the dendritic tree. Upon overlapping glutamate exposure and membrane depolarization, cooperative activation of glutamate-bound NMDA receptors and voltage-gated calcium channels (VGCCs) can drive a large and self-sustaining inward current, leading to plateau potentials in the dendrites and complex spikes in the soma.^19^ Strong distal synaptic inputs alone evoked plateau potentials,^20^ but pairing with a burst of bAPs was a far more potent driver,^19^ while pairing with a single bAP was not effective at inducing plateau potentials, in either CA1^21–23^, CA3^24^ or cortical Layer 5 pyramidal^25^ neurons. NMDAR-mediated plateau potentials are effective triggers for synaptic plasticity in acute brain slices, and can trigger synaptic plasticity^26^ and formation of CA1 place cells *in vivo*, a phenomenon called behavioral timescale synaptic plasticity (BTSP).^27^ It has been controversial whether bAPs are necessary for triggering dendritic plateau potentials and synaptic plasticity, because bAPs are often too brief and too attenuated to engage NMDA receptors at distal dendrites (for a review see ^28,29^). The underlying logic of dendritic excitability remains unknown.

Here we show that under conditions where integration is dominated by passive cable properties, back-propagation is still strongly modulated by nonlinear dendritic excitations. Specifically, back-propagating action potentials successively inactivate A-type potassium channels, then activate dendritic sodium channels, calcium channels, and, in the presence of synaptic inputs, NMDA receptors. The net effect is that conjunction of distal synaptic inputs with an acceleration in the somatic spike rate (e.g. a period of silence followed by a pair or trio of closely spaced somatic spikes) leads to dendritic plateau potentials. These observations provide a biophysical basis for the “triplet learning rule” ^30^ which has been shown to tame instabilities of simple Hebbian plasticity, and to capture many aspects of plasticity experiments.^31^

## Results

We probed dendritic excitability by combining targeted optogenetic stimulation and high-speed structured illumination voltage imaging (Fig. 1). We modified a blue-shifted channelrhodopsin, CheRiff^32^, and a chemigenetic voltage indicator, Voltron2^33^, to improve dendritic expression by attaching an N-terminal Lucy-Rho tag^34^ and C-terminal ER export and TS trafficking motifs^35,36^. We expressed both constructs from a bicistronic vector at very low density in mouse hippocampal CA1 pyramidal neurons, and then prepared acute brain slices (Methods).

**Fig. 1.**
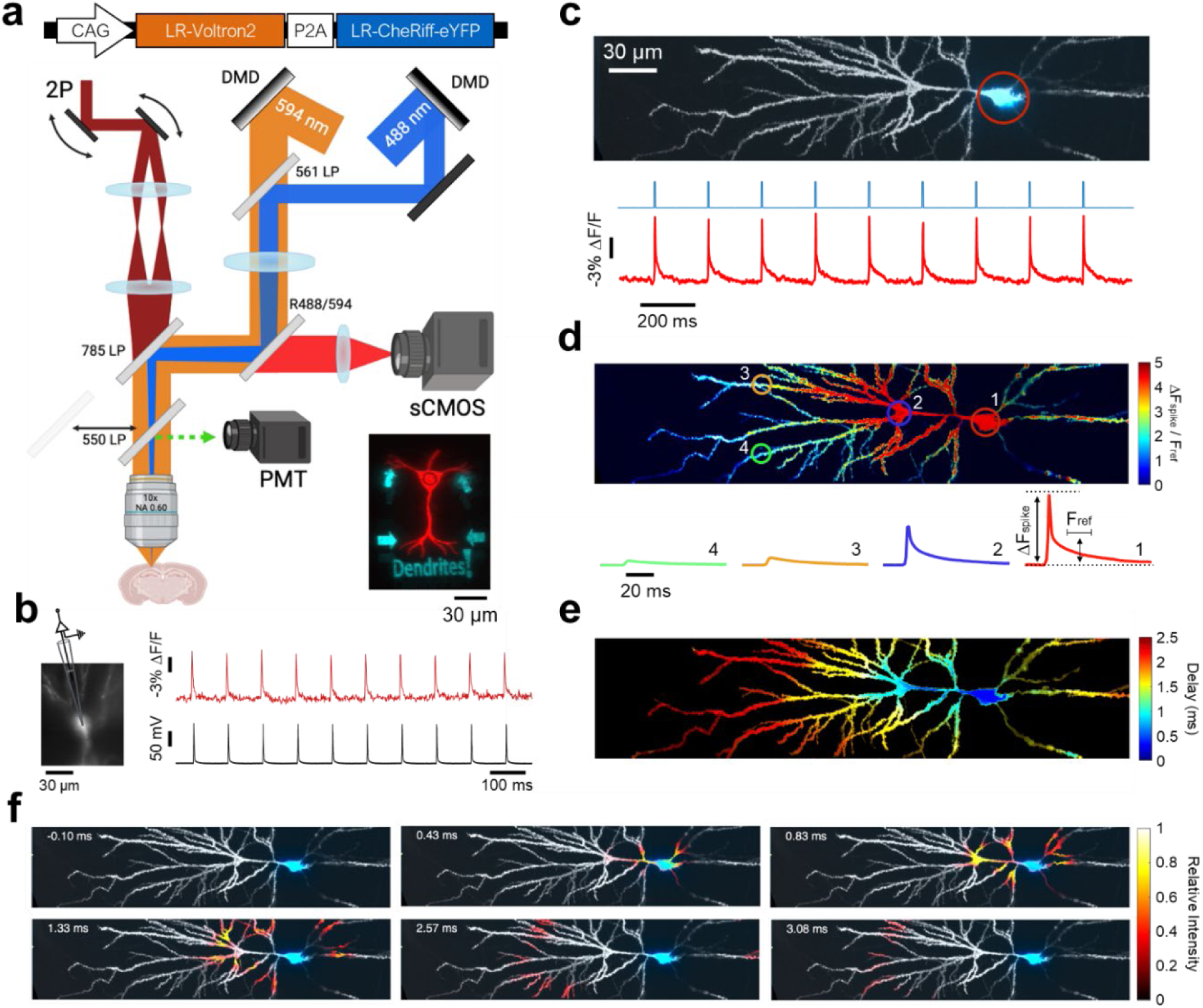
Mapping dendritic voltages with all-optical electrophysiology. **a,** Top: genetic construct for co-expression of LR-Voltron2 and LR-CheRiff-eYFP. Bottom: optical system combining two-photon (2P) static structural imaging (dark red), micromirror-patterned dynamic voltage imaging (orange), and micromirror-patterned optogenetic stimulation (blue). DMD, digital micromirror device. Inset: micromirror-patterned red and blue illumination on a test slide. **b,** Concurrent voltage imaging and whole-cell patch clamp recording at the soma. Sample rates: 1 kHz and 100 kHz, respectively. Spikes evoked by current injection (2 nA for 2 ms at 10 Hz). **c,** Top: 2P structural image of a CA1 neuron (gray), overlayed with eYFP epifluorescence indicating optogenetic stimulus region (blue; 10 ms duration, 5 Hz). Bottom: optogenetic stimulation (blue) and voltage-dependent fluorescence at the soma (red). **d,** Top: spike amplitude map from spike-triggered average of 59 well-isolated spikes. Bottom: spike-triggered average voltage traces in the correspondingly numbered circled regions. ΔF_spike_, peak spike amplitude. F_ref_, mean amplitude during the reference time (from t = 10-20 ms after spike, Methods). **e,** Spike delay map. **f,** Sub-frame interpolation showing details of spike back-propagation (see also Movie 3).

Using patch clamp recordings, we verified that these constructs did not perturb neuronal resting properties or excitability, in comparison to neighboring non-expressing neurons (Fig. S1). We then performed simultaneous patch clamp and fluorescence recordings and validated that the recordings measured voltage changes with sub-millisecond kinetics, ∼1 mV precision, and perfect spike-detection fidelity over thousands of spikes at spike rates up to the maximum tested, 100 Hz (Fig. S2).

The blue light used for optogenetic stimulation also excited fluorescence of the eYFP expression tag in CheRiff-eYFP, so we developed procedures to map the strength and distribution of the optogenetic stimuli via the eYFP fluorescence (Fig. S3). We tested the specificity of the patterned 1-photon optogenetic stimulation by comparing evoked responses for stimuli targeted directly to an in-focus dendrite vs. the same stimuli laterally offset by differing amounts (Fig. S4). Dendritic optogenetic stimuli could be localized within a ∼30 μm region.

Finally, we quantified how well the structured illumination voltage imaging rejected out-of-focus signals. Maps of -ΔF_spike_, the voltage-induced action potential amplitude, clearly showed the structure of the in-focus components of the dendritic tree, confirming that voltage signals from in-focus dendrites were readily distinguishable from background (Fig. S5).

After functional recordings, we made high-resolution, three-dimensional structural images via 2-photon (2P) microscopy. We mapped the functional recordings onto the independently measured cell morphology via a fit to a forward model of the microscope blurring function (Methods, Fig. S6). We further used PCA-based denoising similar to Ref. ^37^ (Methods, Fig. S7) to reduce the effect of pixel-wise shot noise. Together, these techniques produced simultaneously high spatial and temporal resolution voltage maps (Movies 1, 2, Fig. 1c – e) which accounted for > 93% of the variance in the raw data, confirming that the procedure captured most of the underlying dynamics. We then applied the Sub-Nyquist action potential timing (SNAPT)^32^ technique to map bAP wavefront motion with 25 μs time resolution (Movie 3, Fig. 1f, Methods). While the above image-processing steps were helpful for visualizing the voltage response maps, the biophysical analyses below were performed on the raw fluorescence to ensure fidelity to the underlying dynamics.

### Distal dendritic depolarization favors dendritic spikes (dSpikes)

We optogenetically stimulated a small cluster of proximal oblique dendrites (20 ms duration, 5 Hz, 54 repeats) and mapped the bioelectrical responses (Fig. 2). Each stimulus evoked two spikes at the soma, and dichotomous responses in the distal apical dendrites: in most trials the dendrites showed a low-pass filtered and strongly attenuated copy of the somatic activity; but in some cases (8 of 54 trials), the dendrites produced a single rapid spike (Fig. 2a). A histogram of event amplitudes in the soma and dendrites showed a unimodal distribution at the soma and a clearly bimodal distribution in the distal dendrites (Fig. 2b). A histogram of event times showed that in this experiment the dendritic excitations always emerged from the second somatic spike, never the first (Fig. 2c). We refer to these dendritic excitations as dSpikes.

**Fig. 2.**
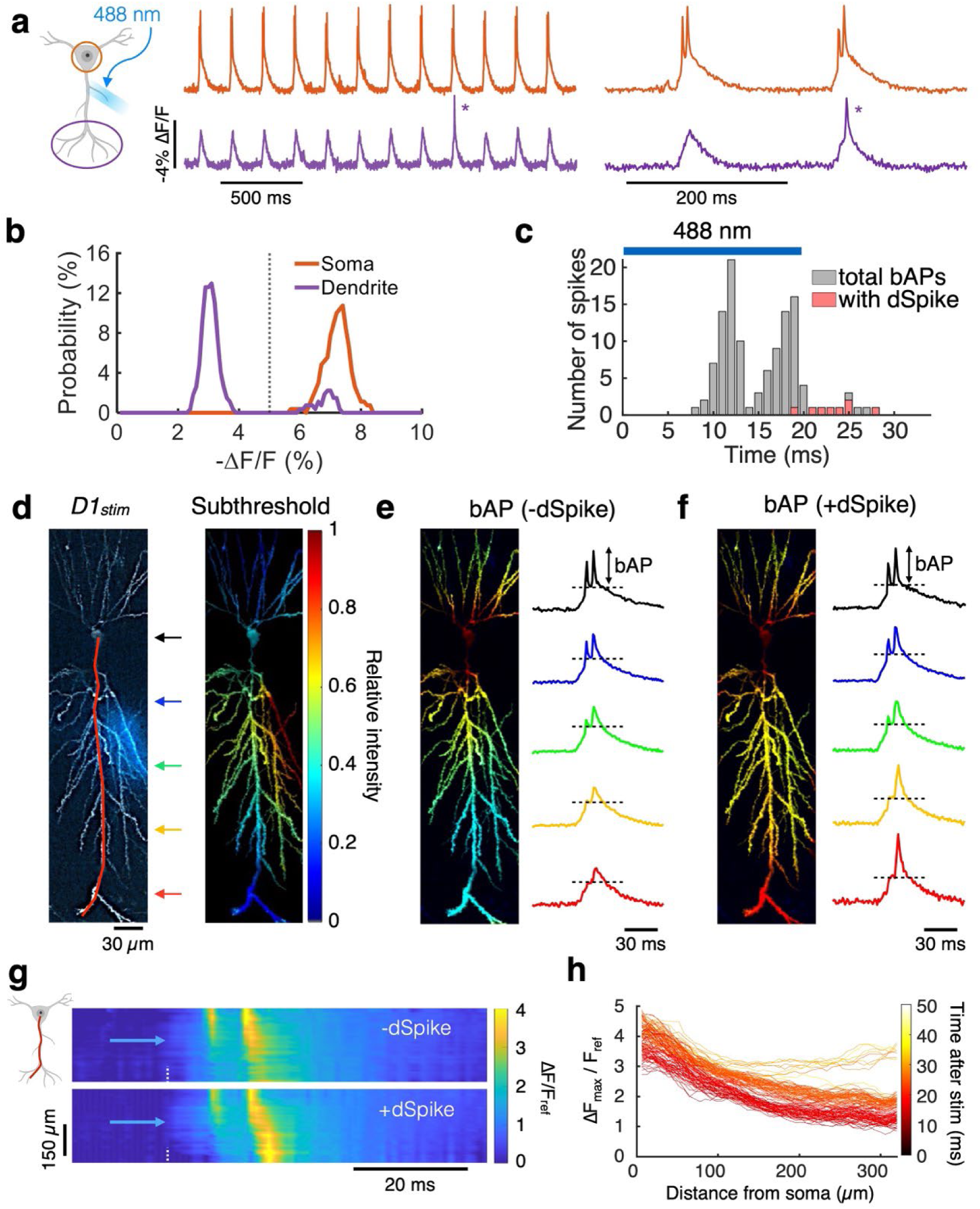
Spatial and temporal maps of dendritic spikes (dSpikes). **a,** Example traces at the soma (orange) and a distal dendrite (> 300 µm; purple) showing trials without and with a dSpike (*) in response to stimulation at a proximal dendritic branch (*D1_stim_*; 20 ms duration at 5 Hz). **b,** Distribution of bAP amplitudes at the soma (orange) and at a distal dendrite (> 300 µm; purple). The amplitude in the soma had a unimodal distribution and the amplitude in the dendrites had a bimodal distribution. Black dotted lines indicate threshold between -dSpike vs. +dSpike. **c,** Counting all bAPs (gray), and bAPs with dSpikes from the same cell (red), as a function of time after optogenetic stimulus onset. Blue bar shows the stimulus timing. **d,** Left: 2P structural image (gray) overlayed with fluorescence of eYFP (blue) indicating spatial distributions of the optogenetic stimulus. Right: normalized amplitude (ΔF/F_ref_) map for subthreshold depolarization (stimulus-triggered average from 54 trials). Amplitude heatmaps for the following panels share the same color scale. **e,** Normalized amplitude (ΔF/F_ref_) map for back-propagating action potentials (bAPs) without dSpikes (-dSpikes; spike-triggered average from 56 spikes). Right: single-trial example of bAPs without a dSpike, sampled along the red line in (**d**) at the locations indicated by colored arrows. **f,** Corresponding plots for bAPs with dSpikes (+dSpikes). Normalized amplitude map, spike-triggered average from 8 spikes. **g,** Kymographs comparing single-trial instances of bAPs with (+dSpike) and without (- dSpike) a dendritic spike along the red line in (**d**). The soma is at the top and the distal apical dendrites are at the bottom. Blue arrowhead indicates place and time of stimulus onset. White dotted line indicates time of stimulus onset. **h,** Amplitude profiles for each bAP along the red line in (**d**). Events that rise in the distal region indicate dSpikes. Plots were color-coded by the time after stimulus onset.

We then examined the spatial structures of the neuronal voltages in detail. The eYFP fluorescence in CheRiff-eYFP showed the stimulus profile (Fig. 2d). We calculated a stimulus-triggered average of the subthreshold depolarization prior to the onset of the first somatic spike. This voltage had greatest amplitude in the directly stimulated dendrites, and decayed smoothly away from the stimulus region (Fig. 2d), suggestive of passive dendritic integration.

We then divided the trials into those that did or did not evoke a dSpike. Spike-triggered averages of the peak depolarization showed that in the -dSpike trials the bAP was primarily confined to the soma and proximal apical and basal dendrites (Fig. 2e), while in the +dSpike trials, a second zone of amplification emerged in the distal dendrites (Fig. 2f).

To study the spatiotemporal structure of the voltage dynamics, we plotted a kymograph of the voltage along the main apical trunk (i.e., a heatmap of voltage with one space axis and one time axis; space axis shown by red line in Fig. 2d). The kymograph showed that upon stimulus onset, the depolarization rose first where the stimulated dendrites joined the trunk (Fig. 2g). This subthreshold depolarization propagated diffusively along the trunk in both the apical and basal directions. When the subthreshold depolarization reached the soma, it triggered a spike (we did not resolve the axon so we could not determine the precise initiation zone). The kymograph showed the finite velocity of spike back-propagation into the dendrites. Plots of the peak spike amplitude along the apical trunk showed exponential decay of the -dSpike events (length constant 255 ± 10 μm, mean ± s.e.m., *n* = 93 spikes) and distal amplification of the +dSpike events (Fig. 2h), very similar to theoretical predictions.^14^

We then repeated these experiments and analyses with optogenetic stimuli delivered either to one of several dendritic branches (Fig. 3a, S8-11), the soma (Fig. 3b), or the soma and a distal dendrite simultaneously (Fig. 3c). Experiments were repeated in *n* = 17 neurons from 15 animals. We noted several consistent features of the dendritic voltage responses:

**Fig. 3.**
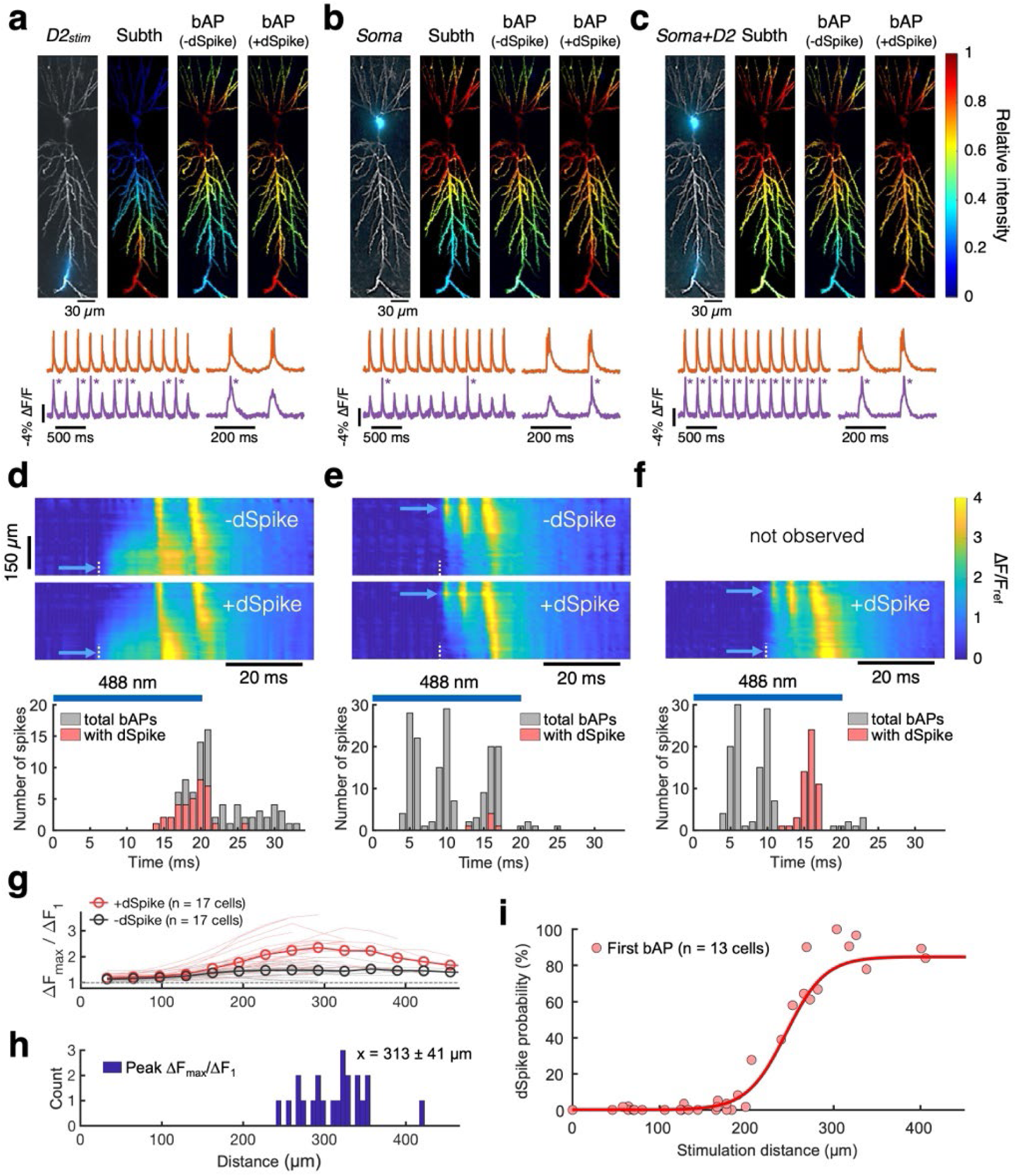
Distal dendritic depolarization favors dendritic spikes (dSpikes). **a-f,** Corresponding plots to those illustrated in Fig. 2 for stimuli at a distal dendrite (**a** and **d**, *D2_stim_*; 54 trials, 19 -dSpikes, and 37 +dSpikes), soma (**b** and **e**, *Soma*; 54 trials, 161 - dSpikes, and 6 +dSpikes), and both soma and distal dendrite simultaneously (**c** and **f,** *Soma+D2*; 54 trials, 116 -dSpikes, and 54 +dSpikes; all trials evoked at least 1 dSpike in this condition). **g,** Amplitude ratio of later bAPs to the first bAP (ΔF_max_/ΔF_1_). Data sorted by the presence (+dSpike) vs. absence (-dSpike) of dSpike. Soma or proximal dendritic branch (< 200 µm) was stimulated (*n* = 22 branches, 17 cells, 15 animals). Mean (open circles), individual stimuli (thin lines). **h,** Distance from soma to the area showing peak ΔF_max_/ΔF_1_ (x = mean ± s.d.), where ΔF_1_ is the amplitude of the first spike after stimulus onset and ΔF_max_ is the amplitude of a subsequent dSpike. **i,** Probability for the first bAP after stimulus onset to trigger a dSpike, as a function of stimulus distance from soma (*n* = 35 dendrites, 13 cells, 11 animals). Red line, sigmoidal fit.

First, the subthreshold voltage profile moved with the stimulus location and decayed smoothly outside the stimulated zone (Fig. 3d-f, S10-11). We only observed diffusive-like propagation of the subthreshold voltages, never dSpikes that initiated in the dendrites. Some prior dendritic patch clamp studies reported that strong current injection (e.g., 2 nA for 5 ms) triggered local dendritic sodium spikes,^38^ while others did not observe these events.^10,39^ Highly synchronous and spatially clustered synaptic inputs have also been reported to evoke local dSpikes in radial oblique dendrites^40^ and the apical trunk and tuft^41^ of CA1 pyramidal cells, though more gradual depolarization via asynchronous inputs did not evoke localized dSpikes. We performed numerical simulations of a conductance-based CA1 model (details below) and found that the absence of dendrite-initiated dSpikes in our experiments was likely due to the more gradual and distributed nature of optogenetic vs. patch clamp stimuli (Fig. S19a,b). Dendritic integration under our conditions appeared to be dominated by passive cable properties.

Second, the neurons produced precisely two kinds of excitations: perisomatic bAPs and bAP-driven distal dSpikes—and, surprisingly, neither spatial profile depended on the location of the driving stimulus. All spikes had similar waveforms at the soma, and then either decayed exponentially along the apical trunk or were amplified distally (Fig. S9), leading to a bimodal distribution of amplitudes in distal apical dendrites (Fig. S10). bAPs reliably engaged all perisomatic and proximal dendrites and failed along the distal trunk, while dSpikes reliably engaged distal apical dendrites too (Fig. S11). Neither bAPs nor dSpikes showed a preference for (or exclusion from) the stimulated dendritic branch. The amplitude of dSpikes relative to the first bAP (i.e., ΔF_max_/ΔF_1_) was maximum in the distal dendritic trunk (313 ± 41 µm from the soma, mean ± s.d., *n* = 22 stimulated regions (soma or proximal branch), 17 neurons, 15 animals; Fig. 3g,h).

Third, the probability that a bAP became a dSpike depended on the stimulus location and spike timing. Proximal stimuli (< 200 μm from the soma) almost never evoked dSpikes on the first bAP, but sometimes evoked dSpikes on the second or third bAP (Fig. 2c, 3e,f, S8). In contrast, distal stimuli (> 250 μm from the soma) reliably evoked dSpikes on the first bAP (Fig. 3d). Pooled data from 35 branch stimulations (*n* = 13 cells from 11 animals) revealed that the probability of evoking a dSpike with the first bAP followed a sigmoidal distance dependence, with a plateau of ∼80% success rate for stimuli > 300 µm from the soma (Fig. 3i).

These seemingly complex dSpike dynamics could be described with a simple rule. dSpikes arose if and only if the distal dendrites had been depolarized for at least 15 ms prior to arrival of a bAP. Distal stimuli took approximately 15 ms to depolarize the soma enough to elicit a spike. As a result, stimulation at a distal dendrite (D2) usually produced dSpikes on the first bAP (Fig. 3d). In contrast, stimuli at the soma or proximal dendrites evoked rapid somatic spikes before the distal dendrites were depolarized. These bAPs incrementally depolarized the distal dendrites, opening a window for dSpikes triggered by subsequent bAPs.

This rule is illustrated by comparing Figs. 3d, e, and f. Stimulation of the soma alone never evoked dSpikes on the first two bAPs, and occasionally evoked a dSpike on the third bAP (6 of 54 trials), which came 13 – 17 ms after stimulus onset (Fig. 3e). Stimulation of distal dendrite D2 alone evoked dSpikes on the first bAP, with 65% probability (35 of 54 trials, Fig. 3d). These bAPs came 14 – 21 ms after stimulus onset. Simultaneous stimulation at soma and D2 produced two quick bAPs which failed to evoke dSpikes, and then the third bAP (12 – 17 ms after stimulus onset) triggered a dSpike with 100% probability (54 of 54 trials, Fig. 3f).

Prior work had suggested that a critical somatic spike rate needed to be exceeded to evoke dendritic spikes in cortical neurons.^42,43^ Our results show that in CA1 pyramidal cells the somatic spike rate is not the key variable, but rather the dendritic depolarization. An isolated bAP can become a dSpike if the distal dendrites are pre-depolarized. Consistent with this notion, under a gradual optogenetic ramp stimulus delivered to the soma or proximal dendrites (< 150 µm from the soma), the subthreshold depolarization permeated the dendritic tree before the first spike. For this stimulus waveform, the first somatic spike almost always evoked a dSpike (92%, 188 ramps, 56 cells, 44 animals; Fig. S12). bAP amplitude in the distal dendrites then decreased, sometimes gradually and sometimes abruptly, a phenomenon previously attributed to dendritic sodium channel inactivation.^10^

### Dendritic spike-rate accelerometer, period-doubling, and stochastic back-propagation

Step-wise optogenetic stimuli to the soma or proximal dendrites (> 200 ms duration) evoked sustained spiking at the soma and a biphasic pattern of bAP propagation. For example, a step stimulus to a proximal apical dendrite evoked a bAP without a dSpike, then two bAPs with dSpikes, then a series of bAPs without dSpikes (Fig. 4a). Pooled data (*n* = 43 stimulated branches, 16 cells, 13 animals) showed that upon onset of step-wise optogenetic stimulation, dSpikes failed for the first 18 ± 7 ms (mean ± s.d.), then there was a period where dSpikes succeeded which lasted until 83 ± 47 ms after stimulus onset, followed by subsequent dSpike failures (Fig. 4b). Sustained spiking at the soma led to very few dSpikes later than 100 ms after stimulus onset, regardless of stimulus strength or location. The net effect of this window for dSpike formation was that the dendrites acted as a spike-rate accelerometer: a step-wise increase in the spike rate at the soma led to a transient increase in the dSpike rate in the distal apical dendrites.

**Fig. 4.**
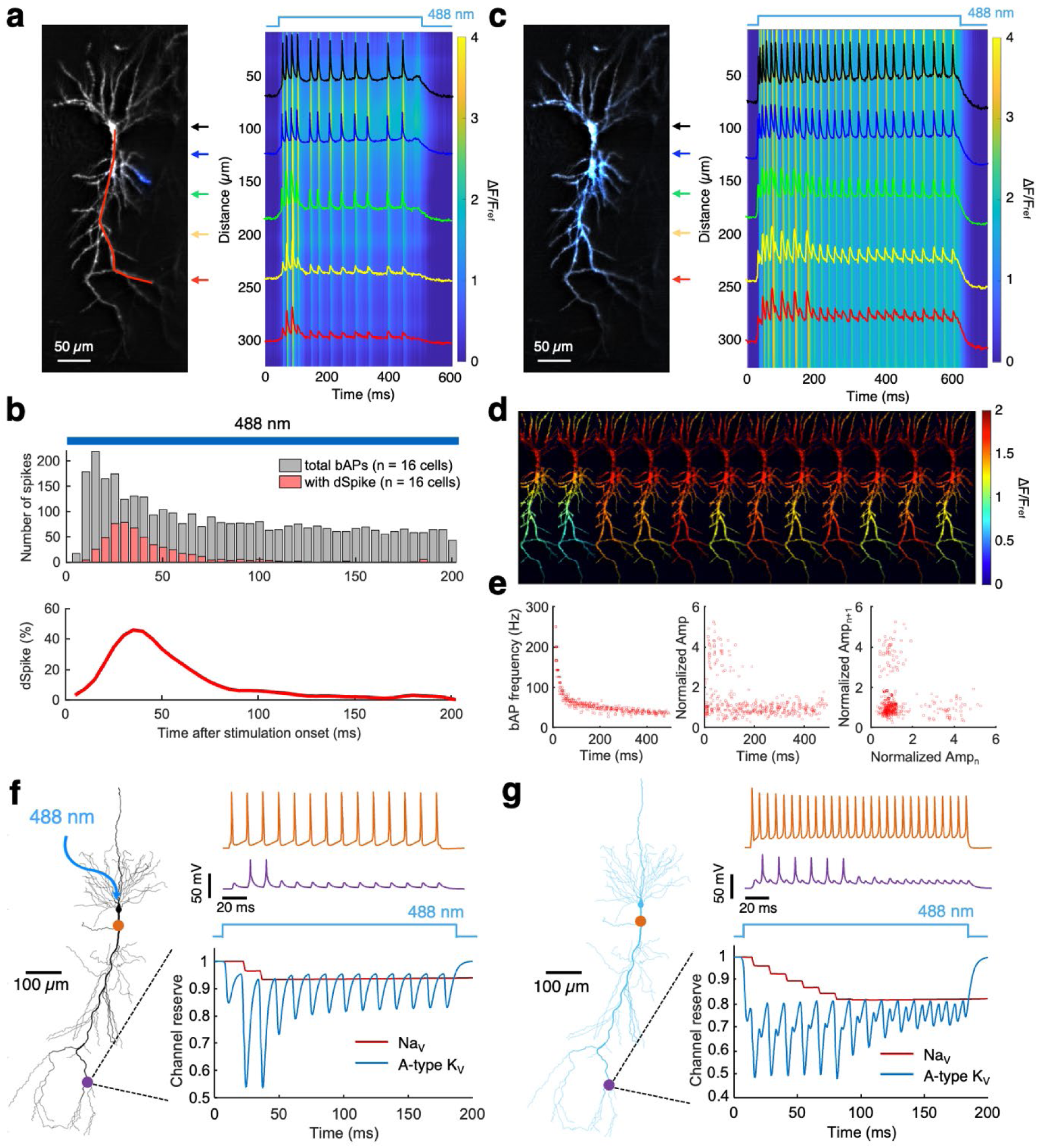
Dendritic excitations implement a spike-rate accelerometer. **a,** Structural image (gray) overlayed with fluorescence of eYFP (blue) showing the optogenetic stimulus. Kymograph along the red line. Example traces taken from the regions indicated by colored arrows. **b,** Top: number of bAPs (gray) and bAPs with dSpikes (red), as a function of time after stimulus onset, showing how acceleration of the somatic spike rate evokes a transient burst of dSpikes (*n* = 43 dendrite stimulus locations, 16 cells, and 13 animals). Bottom: percent of dSpikes among all bAPs. **c,** Equivalent experiment to (**a**) using wide-field illumination which covered the soma and apical trunk (blue). **d,** Amplitude maps of the first 12 bAPs showing two bAP failures, followed by alternating dSpikes and bAP failures. **e,** Plots showing period-doubling bifurcations (*n* = 9 cells from 9 animals). Left: bAP frequency as a function of time after stimulation onset. Middle: Normalized bAP amplitude relative to average of the final 5 bAPs. Right: Relationship between amplitudes of successive bAPs, bAP_n+1_ vs. bAP_n_ showing an alternating motif. **f-g,** Simulations showing spiking at the soma (orange) and distal dendrites (purple) and the dynamics of A-type K_V_ channels (blue) and Na_V_ channels (red). **f,** Soma-targeted stimulation opens a transient window for dSpike excitation. **g,** Wide-field stimulation evokes transient period-doubling bifurcation. Na_V_ channel reserve defined by the slow inactivation gate (fast inactivation and recovery not shown).

We observed a similar accelerometer-like pattern in bAP propagation when using patch clamp injection of current pulses at the soma. At low frequencies (e.g., 3 Hz), membrane depolarization did not accumulate sufficiently to trigger dSpikes; instead, we observed a progressive attenuation of bAP amplitudes in the distal dendrites, consistent with the slow Na_V_ inactivation as previously reported (Fig. S13d).^10,16,18^ At frequencies ≥ 80 Hz, however, we observed a characteristic motif of bAP failure, then a series of dSpikes, and then failures again (Fig. S13), similar to previous reports in L5 pyramidal cells.^42^ The fluorescence and patch clamp recordings at the soma showed excellent correspondence (Fig. S13b,c), further confirming the validity of our voltage imaging data.

We then used a wide-field optogenetic stimulus to test whether stronger depolarization could extend the dSpike time-window. To our surprise, the wide-area stimulus reliably evoked a period-doubling bifurcation: bAPs alternately succeeded and failed to evoke dSpikes (*n* = 9 cells from 9 animals, Fig. 4c, d). Despite the stronger stimulus, the dSpike window only persisted to ∼180 ms after stimulus onset, after which all bAPs failed to evoke dSpikes (Fig. 4e). Even stronger stimuli evoked an epoch of seemingly random interplay of bAPs and dSpikes (Fig. S14), similar to previously reported stochastic backpropagation in L5 cortical pyramidal cells,^44^ and reminiscent of the transition from alternans to arrhythmia in overdriven heart tissue,^45^ and the onset of chaos in linear chains of excitable HEK cells.^46^

We used pharmacology and numerical simulations to determine the biophysical origins of the dendritic spike-rate accelerometer function and of the period-doubling bifurcation. Motivated by prior studies of dendritic excitability,^11,12^ we hypothesized that opening of the dSpike window was driven by inactivation of A-type K_V_ channels. These transient K^+^ channels are expressed at a ∼6-fold higher level in distal dendrites than in soma.^11^ Subthreshold depolarization closes these channels with a voltage-dependent time constant between 6 ms (at -25 mV) and 27 ms (at +55 mV).^11^

To test the involvement of A-type K_V_ channels, we applied the potassium channel blockers, 4-AP (5 mM) or BaCl_2_ (150 µM), and applied brief optogenetic stimuli at the soma (20-30 ms duration, 5 Hz, 59 repeats). Both blockers significantly increased dSpike probability compared to the baseline (BaCl_2_: 11 ± 2% before to 64 ± 10% after, mean ± s.e.m., *n* = 9 cells from 6 animals, *p* < 0.001, paired t-test; 4-AP: 19 ± 4% before to 92 ± 4% after, mean ± s.e.m., *n* = 4 cells from 3 animals, *p* < 0.001, paired t-test; Fig. S15). These results are consistent with previous literature suggesting that A-type K_V_ channel inactivation opens the dendritic spike window at distal dendrites.^11,47^

We hypothesized that the dSpikes were primarily driven by fast Na^+^ currents. To test the involvement of Na_V_ channels, we applied the Na_V_ channel blocker TTX at very low concentration (20 nM).^48^ We stimulated a relatively large area to increase the baseline dSpike probability (∼100 µm diameter across the soma and proximal dendrites, 20 ms duration, 5 Hz, 59-118 repeats). This low TTX dose did not affect spiking at the soma, but the mean dSpike probability was significantly decreased (control: 44 ± 15%, TTX: 11 ± 8%, mean ± s.e.m., *n* = 4 cells from 3 animals, *p* = 0.008, paired t-test; Fig. S16a-d). In some experiments, a higher concentration of TTX (100 nM) eliminated all dSpikes at distal dendrites, while preserving somatic spiking (Fig. S16e).

We further tested the role of VGCC-dependent Ca^2+^ currents in mediating the dSpikes. The dSpikes were not affected by a range of Ni^2+^ concentrations (100-500 µM; Fig. S17a-e), suggesting that they are not mediated by VGCCs. Under strong optogenetic drive, dSpikes occasionally triggered broader (> 20 ms duration) dendritic spikes which we hypothesized were Ca^2+^ spikes (Fig. S17f). Addition of Ni^2+^ (500 μM) eliminated these broad spikes, confirming their dependence on VGCC activation. Addition of Ba^2+^ (150 µM) to otherwise untreated neurons increased the prevalence of putative Ca^2+^ spikes, and subsequent addition of Ni^2+^ (500 µM) eliminated these events while preserving dendritic Na^+^ spikes (Fig. S17g,h). Together, these results imply that dendritic Na^+^ spikes could trigger dendritic Ca^2+^ spikes, and that dendritic potassium channels typically supressed Ca^2+^ spikes.

These data confirmed that dSpikes were primarily mediated by Na^+^ currents, and suggested that the closing of the dSpike window was driven by dendritic Na_V_ channel slow inactivation.^10^ Dendritic Na_V_ slow inactivation time constants have been reported to range from 216 ms at 10 Hz spike rate to 58 ms at 50 Hz,^17^ broadly consistent with our measurement of a 100 - 180 ms dSpike window across a range of spike rates.

We simulated a multi-compartment conductance-based model of a CA1 pyramidal neuron, using biophysically realistic ion channels, modified from Ref. ^49^ (Methods, Fig. S18). To reproduce our data, we added a slow inactivation gate to the dendritic Na_V_ channels; and we adjusted the spatial distributions of dendritic A-type and Na_V_ channels. After tuning the model, a simulated step-wise optogenetic stimulation at the soma led to a dSpike motif of failure, success, success, failures (Fig. 4f), which closely matched our observations of a dendritic spike-rate accelerometer (Fig. 4a). Simulated wide-field optogenetic stimulation with the same model parameters evoked a period-doubling bifurcation with alternating dSpike successes and failures, followed by repeated failures (Fig. 4g), matching our observations with wide-field stimulation (Fig. 4c). We conclude that the numerical model accurately captured the dynamics of a CA1 pyramidal dendritic tree.

The simulations reported the contributions of each channel type to the dynamics, confirming that the dSpike window was plausibly opened by A-type K_V_ inactivation and closed by slow Na_V_ inactivation. Together, these two channels converted a step-wise increase in spike rate at the soma into a transient burst of dSpikes, i.e. implementing a spike-rate accelerometer. The simulations also explained the period-doubling: Under simultaneous distal and proximal stimulation, the absolute refractory period of the distal dendrites slightly exceeded the refractory period of the soma. Consequently, after a successful dSpike, the dendrites were still recovering when the next bAP arrived.

To characterize the robustness of the simulations, we systematically varied the dendritic Na_V_ density, amount of Na_V_ slow inactivation, A-type K_V_ density, and stimulus strength. While the precise number of bAP failures and bAP-evoked dSpikes depended on these parameters (consistent with the variable number of dSpikes in our experiments), the spike-rate accelerometer motif of bAP failure, dSpikes, and then failure persisted over a wide range of parameters (Fig. S19). This robustness highlights the biological plausibility of the mechanisms identified in our simulations.

Motivated by the parsimonious explanation of the seemingly complex dendritic phenomenology, we also developed a coarse-grained two-compartment Izhikevich-type model which also captured the opening and closing of the dSpike window (i.e. spike-rate accelerometer) and the period-doubling under simultaneous distal and proximal stimulation (Fig. S20). Such a model may be useful for computationally efficient large-scale simulations of neural dynamics with semi-realistic dendrites.

### Plateau potentials are evoked by collision of synaptic inputs and dSpikes

The experiments with patterned optogenetic stimulation raised a perplexing question: Since bAP and dSpike spatial footprints were each largely insensitive to the stimulus location, then where is the eligibility trace which determines the specific synapses to potentiate during long-term potentiation? Our voltage imaging experiments ruled out membrane voltage as a primary carrier of the eligibility trace.

Optogenetic stimulation provides a pure depolarization to the dendrites, bypassing the ligand-gated channels which are engaged during synaptic transmission. In particular, NMDA receptors show voltage-dependent gating only if first bound to glutamate.^50^ To determine how glutamatergic inputs affect dendritic electrophysiology, we performed experiments combining synaptic stimulation, optogenetic stimulation, and dendritic voltage imaging.

We used electric field stimulation (EFS; 0.1 ms, single pulses, 10-40 V) to activate presynaptic axon terminals in the temporoammonic pathway that synapses onto the distal apical dendrites. We sequentially applied optogenetic stimulation to the soma alone (30 ms), EFS alone, or EFS and optogenetic stimulation. As before, the optogenetic stimulation alone evoked bAPs. EFS alone evoked a distal depolarization which propagated diffusively toward the soma. Remarkably, combining optogenetic and electrical stimuli evoked a long-lasting (∼120 ms) plateau potential in the dendrites and a burst of spikes on top of a strong subthreshold depolarization at the soma, resembling a complex spike (Fig. 5a).^27,51^ This behavior was qualitatively different from the combined soma + distal dendrite optogenetic stimulation (Fig. 3c,f), pointing to a critical role of glutamate-gated channels in the process. Fig. S21 shows a gallery of kymographs of responses to optogenetic, EFS, and combined stimulation in *n* = 9 neurons (7 animals).

**Fig. 5.**
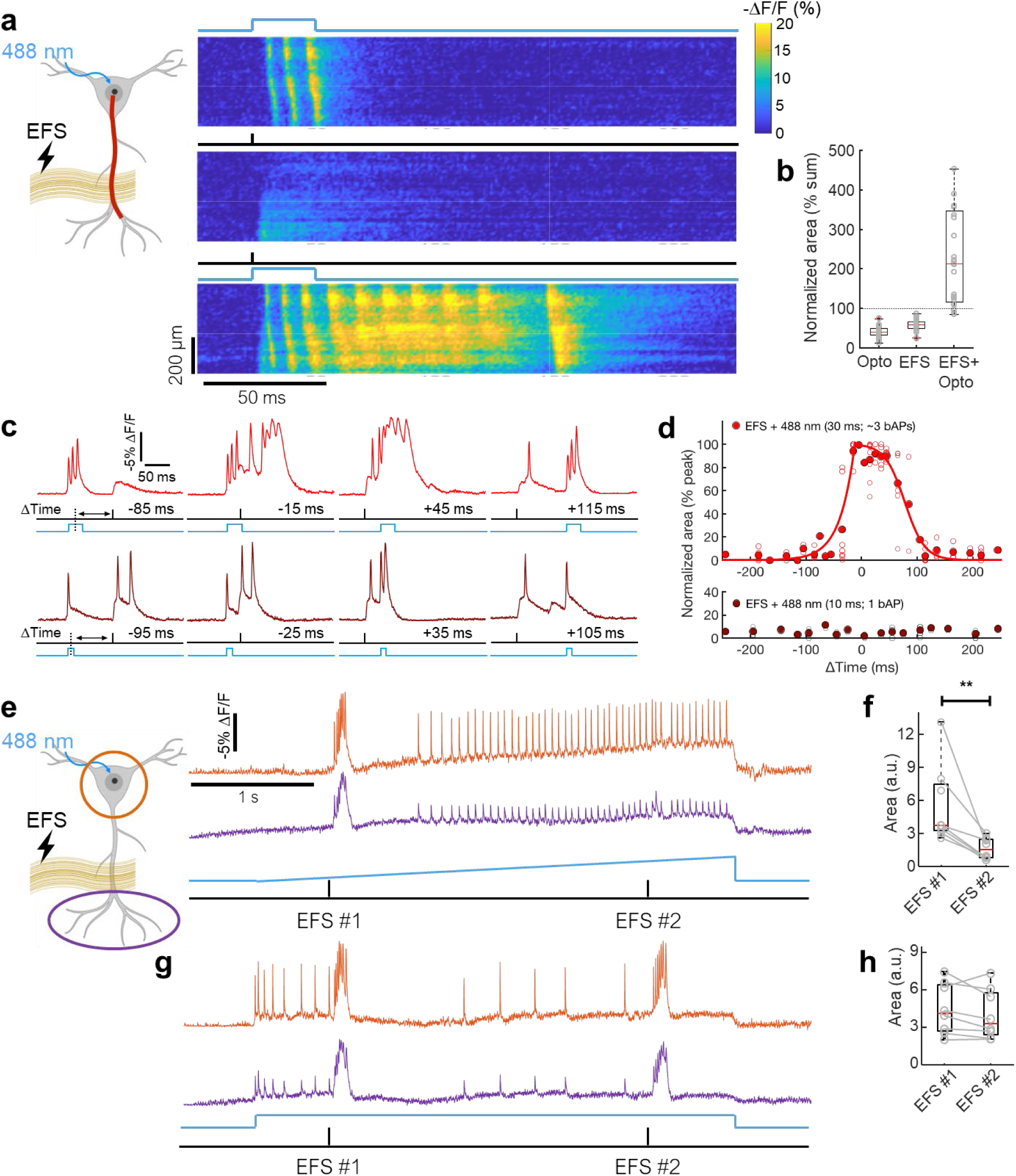
Collision of synaptic inputs and bAP-induced dSpikes triggers plateau potentials. **a,** Kymographs (ΔF/F) along the apical trunk (red line) showed the effects of (top) optogenetic stimuli targeted to the soma (30 ms duration), (middle) EFS-triggered synaptic inputs (0.1 ms duration), and (bottom) combined optical and electrical stimuli. **b,** Area under the curve (AUC) for combined stimulus normalized to the sum of AUC for optical and EFS stimuli alone (*n* = 28 cells from 12 animals). **c,** Top: Fluorescence in a distal dendrite (> 200 µm from the soma) in response to combinations of optogenetic (30 ms, 3 bAPs) and EFS stimulation at various time offsets (ΔTime). Bottom: Corresponding data using 10 ms optogenetic stimulation (1 bAP). **d,** AUC for combined stimulus as a function of ΔTime. Data for each cell scaled to the range [0, 1] (*n* = 8 cells from 7 animals for 30 ms stimulus; *n* = 5 cells from 4 animals for 10 ms stimulus). Open symbols represent individual data and filled symbols represent mean at each ΔTime. Red lines: exponential fit from -245 to 0 ms; sigmoidal fit from 0 to +245 ms. **e,** EFS and ramped soma-targeted optogenetic stimulation. Under weak optogenetic stimulus, EFS #1 evoked a plateau potential in the dendrites (purple) with an accompanying complex spike in the soma (orange). Under strong optogenetic stimulus, EFS #2 did not evoke a complex spike or plateau potential. **f,** AUC for the EFS-evoked event was significantly lower when paired with strong optogenetic stimuli vs. weak stimuli (*p* = 0.01, *n* = 7 neurons from 5 animals, paired t-test). **g,** Repeated EFS under tonic weak optogenetic stimulus. Both EFS stimuli evoked plateau potentials. **h,** AUC for the first and second EFS-evoked events was similar when paired with tonic weak optogenetic stimuli (*p* = 0.19, *n* = 7 neurons from 5 animals, paired t-test). Box plots show median, 25^th^ and 75^th^ percentiles, and extrema.

We compared the area under the curve (AUC) for the waveforms at distal dendrites (> 200 µm from the soma) induced by optical, electrical, and combined stimulation. The average AUC_combined_ was 3.1 ± 0.7-fold greater than AUC_optical_ + AUC_electrical_ (mean ± s.e.m.) and for more than half of the cells studied (15 of 28 cells from 12 animals), AUC_combined_ was more than twice AUC_optical_ + AUC_electrical_ (Fig. 5b). We characterized in detail the response properties of the cells that showed > 200% nonlinearity. The nonlinear amplification was greatest when the optical and electrical stimuli overlapped in time (*n* = 8 cells from 7 animals; Fig. 5c, d, S22). The amplification was an asymmetric function of the time offset: For ‘EFS before bAPs’, the amplification followed a sigmoidal profile, decaying by half in 87 ms. For ‘EFS after bAPs’, the amplification decayed exponentially with a time constant of 35 ms. These findings are consistent with prior results showing that when presynaptic inputs are paired with postsynaptic bursts, hippocampal LTP can occur for either relative timing of pre- and post-synaptic activity.^22,27^

We then tested how the number of optogenetically evoked bAPs affected the nonlinear amplification. Triggering a single bAP by a 10 ms optogenetic stimulus targeted to the soma did not evoke a nonlinear dendritic response, regardless of timing relative to the EFS (*n* = 5 cells from 4 animals; Fig. 5c, d). We conclude that at least one bAP-evoked sodium dSpike was necessary to create a plateau potential.

In our model, Na_V_-mediated dSpikes are a necessary trigger for plateau potentials. This model makes a counterintuitive prediction: if there is a sustained train of somatic spikes, and a distal synaptic input arrives after the dSpike window has closed (> 180 ms after spike-train onset), then this synaptic input will not be able to trigger a plateau potential despite coincidence with somatic spikes. We tested this prediction by stimulating the soma with gradual (3 s) optogenetic stimulus ramps, and then delivering EFS at the beginning (0.5 s after ramp start) and end (2.5 s after ramp start) of the ramps (Fig. 5e). The early EFS reliably evoked plateau potentials (AUC 5.6 ± 3.6, mean ± s.d.), whereas the late EFS, which occurred when the average spike rate was 24 ± 4 Hz (mean ± s.d.), did not (AUC 1.6 ± 1.0, mean ± s.d., n = 7 cells from 5 animals, *p* = 0.01, paired t-test, Fig. 5f). These experiments illustrate the important contribution of the dendritic spike-rate accelerometer to formation of plateau potentials: coincidence of distal synaptic inputs with somatic spikes alone was not sufficient to induce a plateau potential; rather, distal synaptic inputs had to coincide with an acceleration in the somatic spike rate.

As a control experiment, we delivered a weak 3 s optogenetic step stimulus to the soma, too small to drive dendritic Na_V_ inactivation on its own (Fig. 5g). In this case EFS at 0.5 s and at 2.5 s were equally effective at inducing plateau potentials (0.5 s: AUC 4.5 ± 2.0, mean ± s.d., 2.5 s: AUC 4.0 ± 2.0, mean ± s.d., *p* = 0.19, n = 7 cells from 5 animals, paired t-test; Fig. 5h).

We applied channel blockers to investigate the molecular mechanisms underlying dendritic plateau potentials. Block of NMDARs by D-AP5 (50 µM) reduced the plateau area to 42 ± 7% of baseline (*n* = 6 cells from 5 animals, mean ± s.e.m.) compared to vehicle control (107 ± 5% of baseline, *n* = 11 cells from 8 animals; *p* < 0.001, one-way ANOVA with Bonferroni’s *post hoc* test; Fig. 6a,b). Block of voltage-gated Ca^2+^ channels (VGCCs) by Ni^2+^ (100 µM) largely eliminated the plateau potential (AUC 18 ± 5% of baseline, *n* = 4 cells from 4 animals, mean ± s.e.m.; Fig. 6a,b), implying that VGCC currents were necessary for the initiation of the plateau potential. Block of dendritic Na_V_ channels by TTX (20 nM) also eliminated plateau potentials (AUC 26 ± 8% of baseline, *n* = 4 cells from 3 animals, mean ± s.e.m.; Fig. 6a,b), implying that Na_V_ currents were also necessary to trigger plateau potentials. None of these drugs substantially affected somatic spiking at the concentrations used. The drug effects on the plateau were much larger than their effects on the AUC for EFS or optogenetic stimulation alone (Fig. S23).

**Fig. 6.**
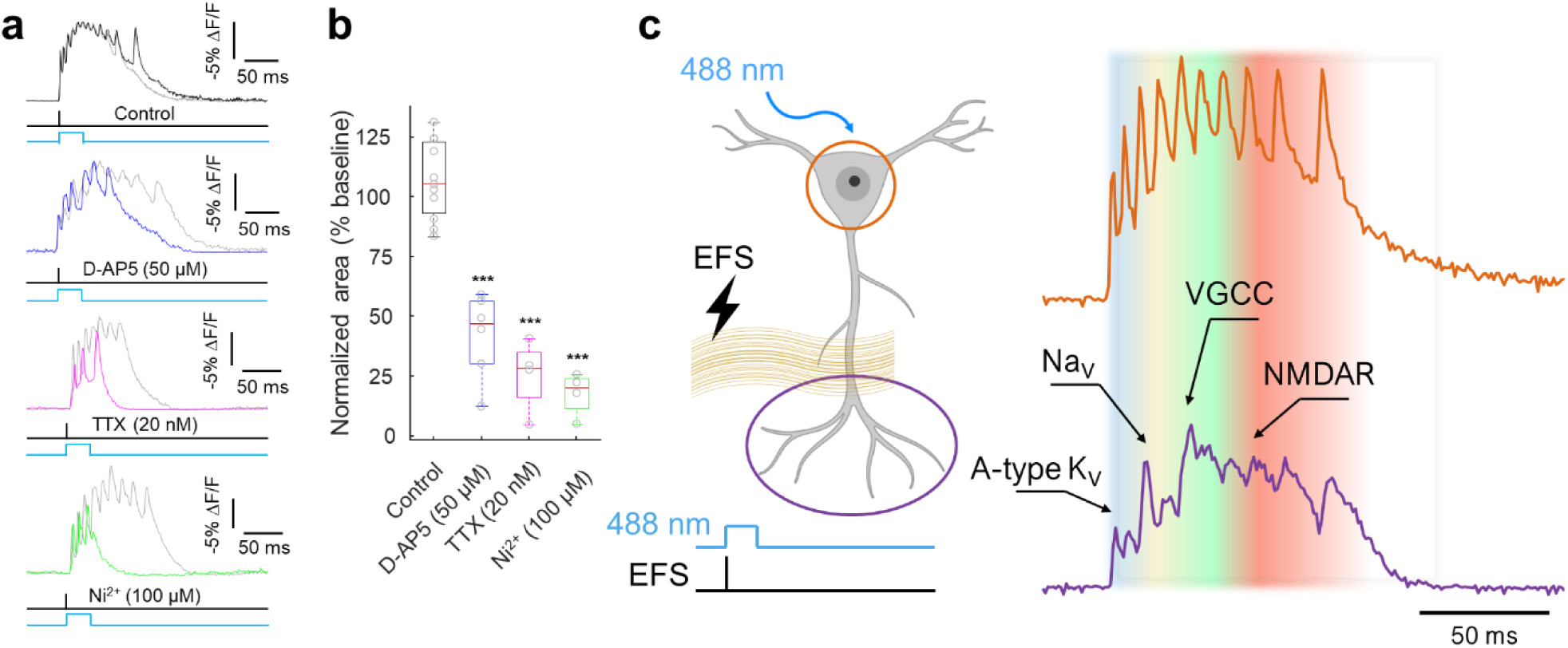
Contribution of dendritic ion channels to plateau potentials. **a,** Plateau potentials were evoked by concurrent optogenetic stimulation to the soma (30 ms) and EFS-triggered synaptic inputs (0.1 ms). Channel blockers: D-AP5 (50 µM, *n* = 6 cells from 5 animals), TTX (20 nM, *n* = 4 cells from 3 animals), and Ni^2+^ (100 µM, *n* = 4 cells from 4 animals) were compared to the vehicle control (*n* = 11 cells from 8 animals). Sample traces overlaid with the paired baseline trace measured from the same cell (gray). Fluorescence measured in a distal dendrite (> 200 µm from the soma). **b,** Quantification of effects in (**a**). Box plots show median, 25^th^ and 75^th^ percentiles, and extrema. \*\*\**p* < 0.001 vs. control, one-way ANOVA with Bonferroni’s *post hoc* test. **c,** Example traces of a complex spike at the soma (orange) and simultaneously recorded plateau potential in the distal dendrites (purple). Overlaid shading qualitatively indicates dominant contributions of distinct dendritic ion channels. bAP propagation within dendrites is initially limited by A-type K_V_ channel activation. The initial bAPs combine with synaptic depolarization to inactivate A-type K_V_ channels, allowing subsequent bAPs to evoke Na_V_-based dSpikes. These dSpikes lead to VGCC-dependent calcium spikes, causing prolonged dendritic membrane depolarization (> 20 ms). In the presence of glutamate from presynaptic inputs, this prolonged depolarization efficiently engages NMDARs, resulting in a global plateau potential.

## Discussion

High-resolution voltage imaging and optogenetics revealed the spatial structure and biophysical origin of plateau potentials (Fig. 6c). In a polarized dendrite, A-type K_V_ channels shunt the voltage, suppressing bAP propagation. The 6-27 ms time-constant for A-type channels to inactivate^11^ is slow enough that distal inputs usually trigger a spike at the soma before the dendrites become directly excitable. One or two closely spaced bAPs, direct distal depolarization, or gradual subthreshold somatic depolarization, transiently inactivates this shunt, opening a path for subsequent bAPs to activate Na_V_ channels, evoking dSpikes. The dSpikes then activate VGCC channels, leading to distal Ca^2+^ spikes. Together, the Na_V_ and VGCC currents activate the NMDAR receptors, leading to plateau potentials. Under sustained spiking, Na_V_ inactivation returns the dendrites to a non-excitable state.

These biophysical dynamics implement a spike-rate accelerometer: a period of silence followed by a burst of bAPs triggers dSpikes, whereas neither isolated low-frequency nor sustained high-frequency bAPs evoke these events. This property leads to the surprising observation that during sustained high frequency firing synaptic inputs cannot evoke plateau potentials (Fig. 5e-h). The precise parameters governing whether a sequence of bAPs triggers a dSpike depends on the recent history of dendritic subthreshold depolarization (e.g. Fig. 3), and on the channel expression levels (e.g. Fig. S19), suggesting modes of fast and slow regulation, respectively, of spike timing-dependent plasticity rules.

Under targeted optogenetic stimulation, bAP spatial profiles were highly stereotyped, comprising only two motifs: bAPs alone, or bAPs with broadly distributed dSpikes. Thus dSpikes appeared as a broadcast signal, covering the dendritic tree and carrying precise spike timing information but little spatial information. This idea had been proposed previously, but not directly observed.^52^ In contrast, the pattern of glutamate-bound NMDA receptors carried precise spatial information on the identities of the activated synapses, but little temporal information. Conjunction of these electrical and chemical signals drove spatiotemporally precise NMDA receptor activation and plateau potentials.^53^

The dynamics leading to plateau potentials closely resemble a triplet plasticity rule, which has been shown theoretically to rectify the inconsistencies between Hebbian plasticity and observed properties of LTP.^30^ The loss of dendritic excitability at sustained high spike rates suggests a mechanism for Bienenstock-Cooper-Munroe (BCM)-style metaplasticity, i.e. suppression of LTP at sustained high spike rates to avoid runaway plasticity.^31^ The dendritic spike-rate accelerometer provides an intuitive biophysical mechanism for a multiplet-based plasticity rule in CA1 pyramidal cells.

Our findings also suggest that neurons may have two distinct plasticity modes. If synaptic activation is sparse, then conjunction of dSpikes with synaptic activation may drive local NMDAR activation, calcium influx, and plasticity, without triggering a plateau potential.^20,28,48^ If synaptic activation in apical dendrites is sufficiently dense, then cooperative NMDAR and VGCC activation drives broadly distributed plateau potentials and somatic complex spikes, which may potentiate all synapses that are active within a broad time window surrounding the plateau potential. This second mode of plasticity resembles BTSP.^27^ This second plasticity mode implies an associative plasticity rule in which inputs from entorhinal cortex on distal dendrites gate, via plateau potentials, plasticity of inputs from CA3, which primarily synapse onto basal and proximal apical dendrites.^54^

*In vivo* calcium imaging experiments have reported localized dendritic signals, which are thought to arise from subthreshold calcium influx through VGCCs and NMDARs.^55–57^ It is unsurprising that the spatial structures of dendritic calcium and voltage signals differ. Whereas Ca^2+^ ions diffuse ∼2 μm during a typical 100 ms subthreshold event (effective buffered diffusion coefficient of calcium in dendrites is *D* < 20 μm^2^/s),^58,59^ electrical length constants are typically > 100 μm and so electrical events are much more homogeneous across space. Simultaneous voltage and calcium imaging experiments will be crucial for determining the quantitative relations between these two modalities.

Dynamics *in vivo* might differ from our observations in acute slices, but some *in vivo* experiments suggest that the overall picture is similar. Early extracellular recordings from CA1 pyramidal neurons of behaving rats found that spike bursts (indicative of complex spikes) were most likely to occur following 0.1 – 1 s of silence and were suppressed during epochs of sustained fast spiking.^60^ Furthermore, bursts of dendritic activity and putative Ca^2+^ spikes were always preceded by a large-amplitude fast dendritic spike, which we associate with a dSpike.^51^ We recently performed simultaneous voltage imaging in soma and apical dendrites of Layer 2/3 pyramidal cells *in vivo* and observed highly correlated voltages across the dendritic tree, and a biphasic trend in bAP propagation amplitude during sustained somatic spiking, similar to our observations here (e.g. Fig. 4b).^61^

Neurons *in vivo* receive inhibitory and neuromodulatory inputs that could modify the simple picture presented here. Branch-specific inhibition, local NMDAR activation, or modulation of other ion channels might lead to more branch-to-branch variability in voltage dynamics. It remains to be determined whether excitatory inputs *in vivo* are sufficiently strong and clustered to overwhelm the A-type suppression mechanism and to directly drive dendrite-initiated excitations without first initiating a bAP. Further study is required to map dendritic integration and back-propagation in CA1 pyramidal cells in live animals.

## Supporting information

Supplementary Izhikevich model

Supplementary NEURON models

## Acknowledgments

We thank A. Preecha and S. Begum for technical assistance, F. P. Brooks for assistance with software, and B. Sabatini, M. Triplett, D. Peterka, and N. Altunkeser for helpful discussions. This work was supported by Chan Zuckerberg Initiative Dynamic Imaging grant 2023-321177, a Vannevar Bush Faculty Fellowship, a Brain Research Foundation Scientific Innovations Award BRF-SIA-2022-02, the Harvard Brain Science Initiative, and NIH grants R01-NS126043 and R01-MH117042. J.D.W.-C. is a Merck Awardee of the Life Sciences Research Foundation. J. B. G., S. E. P. and L. D. L. are supported by the Howard Hughes Medical Institute.

## Author contributions

PP developed the genetic constructs, performed *IUE* surgeries, and performed all experiments and data analysis. JDWC developed the optical system. DGI performed numerical simulations. BHL helped with the experimental design. YQ identified and characterized the dye and opsin combination for *in vivo* Optopatch. HCD developed the instrument control software. JBG, SEP, KLJ and LDL provided HaloTag dyes and guidance on their use. BA, AP and LP developed methods to fuse functional and structural recordings. AEC supervised the project, helped with data analysis and wrote the manuscript along with PP.

## Competing interests

The authors declare no competing interests.

## Data availability

Data are available from the corresponding author upon reasonable request.

## Code availability

Code for numerical simulations of dendritic excitability is included with this submission.

## Methods

### Genetic constructs

We used Voltron2, an improved chemigenetic voltage indicator^33^, and co-expressed it with a blue-shifted channelrhodopsin, CheRiff by a self-cleaving p2a linker. To optimize expression and dendritic membrane trafficking we designed the construct CAG::LR-Voltron2-TS-ER2-p2a-LR-CheRiff-TS-eYFP-ER2 (Addgene: #203228). In this construct, LR is the membrane localization signal from Lucy-Rho^34^, TS is the trafficking sequence from K_ir_2.1^35^, and ER2 is the endoplasmic reticulum export signal FCYENEV^35^. In some experiments, we used a Cre recombinase-dependent DIO (double-floxed inverse open reading frame) construct, CAG::DIO-LR-Voltron2-TS-ER2-p2a-LR-CheRiff-TS-eYFP-ER2 (Addgene: #203229), and co-expressed it with CAG::Cre (Addgene: #13775) plasmid wt/wt = 30:1 for *in utero* electroporation. As both approaches yielded sparse hippocampal expression and similar data, the data were pooled.

The genes were cloned into an adeno-associated virus (AAV) backbone with a synthetic CAG promoter using standard Gibson Assembly. Briefly, the vector was linearized by double digestion using restriction enzymes (New England Biolabs) and purified by the GeneJET gel extraction kit (ThermoFisher). DNA fragments were generated by PCR amplification and then fused with the backbones using NEBuilder HiFi DNA assembly kit (New England Biolabs). All plasmids were verified by sequencing (GeneWiz).

### In utero electroporation (IUE)

All animal procedures adhered to the National Institutes of Health Guide for the care and use of laboratory animals and were approved by the Harvard University Institutional Animal Care and Use Committee (IACUC). The IUE surgery was performed as described previously.^62^ Timed-pregnant female CD1 mice (embryonic day 15.5, E15.5; Charles River) were deeply anesthetized and maintained with 2% isoflurane. The animal body temperature was maintained at 37 °C. Uterine horns were exposed and periodically rinsed with warm phosphate-buffered saline (PBS). Plasmid DNA was diluted in PBS (2 μg/μL; 0.05% fast green), and 1 µL of the mixture was injected into the left lateral ventricle of the embryos. Electrical pulses (40 V, 50 ms duration) targeting the hippocampus were delivered five times at 1 Hz using tweezers electroporation electrodes (CUY650P5; Nepa Gene). Injected embryos were returned to the abdominal cavity, and the surgical incision was closed with absorbable PGCL25 sutures (Patterson).

### Slice preparation

Coronal slices (300 µm) were prepared from CD1 mice of either sex between 2-4 postnatal weeks. Animals were anesthetized with isoflurane and euthanized by decapitation. The brain was then removed and placed in ice-chilled slicing solution containing (in mM): 210 sucrose, 3 KCl, 26 NaHCO_3_, 1.25 NaH_2_PO_4_, 5 MgCl_2_, 10 D-glucose, 3 sodium ascorbate, and 0.5 CaCl_2_ (saturated with 95% O_2_ and 5% CO_2_). Acute slices were made using a Vibratome (VT1200S, Leica) while maintained in the slicing solution. Slices were recovered at 34 °C for 10 min in the imaging solution (artificial cerebrospinal fluid, ACSF) containing (in mM): 124 NaCl, 3 KCl, 26 NaHCO_3_, 1.25 NaH_2_PO_4_, 2 MgCl_2_, 15 D-glucose, and 2 CaCl_2_ (saturated with 95% O_2_ and 5% CO_2_). Slices were then incubated in ACSF containing JFX_608_-HaloTag ligand^63^ (0.5-1 µM) for 30 min at room temperature, and moved to a fresh ACSF for another 30 min to wash out excess dye. Slices were maintained at room temperature until recordings were made.

Functional recordings were performed at 34 °C. We found that at 24 °C the dendritic excitability dynamics were significantly different (Fig. S24). After a sample had been recorded at 34 °C, the in-line solution heater was switched off for at least 10 min, with the bath temperature continuously monitored using a thermistor probe. We observed a significant increase in dSpike probability at 24 °C (0% vs. 29 ± 16% by the first bAP, *p* = 0.09; 16 ± 3% vs. 50 ± 12% by any bAP, *p* = 0.02; mean ± s.e.m., *n* = 6 cells from 4 animals, paired t-test). This temperature effect could explain the difference between our results and prior measurements at room temperature of bAP propagation into distal dendrites.^10^ Previously published simulations that accounted for the different temperature sensitivities of A-type K_V_ channels and Na_V_ channels^64,65^ predicted temperature-sensitive back-propagation dynamics.^44^

### Electrophysiology

Somatic whole-cell recordings were acquired from hippocampal CA1 pyramidal neurons using a custom upright microscope. All experiments were performed at 34 °C, and continuously perfused at 2 mL/min with ACSF. Patch pipettes (2–4 MΩ) were filled with an internal solution containing (in mM): 8 NaCl, 130 KmeSO_3_, 10 HEPES, 5 KCl, 0.5 EGTA, 4 Mg-ATP, and 0.3 Na_3_-GTP. The pH was adjusted to 7.3 using KOH and osmolality was adjusted to 285–295 mOsm/L with water. Signals were amplified using a Multiclamp 700B (Molecular Devices), filtered at 10 kHz with the internal Bessel filter, and digitized at 100 kHz using a PCIe-6323 (National Instruments) A/D board. After entering the whole-cell configuration, membrane capacitance and membrane resistance were measured under voltage-clamp mode (Fig. S1). Resting membrane potential, rheobase and spike rates were measured under current-clamp mode. Rheobase was defined as the minimum amplitude of a current step (500 ms duration) to evoke at least one spike. To induce a rapid, isolated single spike, 2 nA current (2 ms duration) was injected into the soma in current-clamp mode (Fig. 1b and Fig. S13).

For the experiments in Fig. 5-6, we applied electric field stimulation (EFS) to the temporoammonic (TA) pathway to evoke synaptic responses. We used a concentric bipolar electrode (CBAPB50, FHC) with stimuli of 10-40 V (0.1 ms duration). Stimulus intensity was adjusted to be high enough to obtain plateau potentials when combined with optogenetic stimulation at the soma (30 ms duration). ACSF contained picrotoxin (50 µM) to prevent GABA_A_ receptor-mediated currents, and MgCl_2_ concentration was lowered to 1 mM from 2 mM to enhance NMDA receptor-mediated currents.

### Voltage imaging in custom upright microscope

Voltage imaging experiments were conducted on a previously described home-built epifluorescence microscope.^36^ Briefly, blue (488 nm) light was patterned by a digital micromirror device (DMD) and used for targeted channelrhodopsin stimulation. Stimulated regions were confirmed by fluorescence of the eYFP marker in LR-CheRiff-eYFP. Orange (594 nm) illumination was also patterned by a separate DMD and used for structured illumination voltage imaging and *post hoc* HiLo reconstruction of dendritic morphology.^66^ Typical laser intensity for 594 nm was 10-20 mW/mm^2^. Intensity for 488 was up to 1 mW/mm^2^.

Laser lines from a blue laser (488 nm, 150 mW, Obis LS) and orange laser (594 nm, 100 mW, Cobolt Mambo) were combined by a dichroic (IDEX, FF506-Di03-25×36) and sent through an acousto-optic modulator (TF525-250-6-3-GH18A, Gooch and Housego) for amplitude control. After the modulator, blue and orange lines were split with a dichroic mirror (IDEX, FF506-Di03-25×36), expanded, and sent to two independent DMDs for spatial modulation; one for the blue (Lightcrafter DLP3000, Texas Instruments) and the other for the orange (V-7000 VIS, ViALUX). The DMD planes were recombined via a dichroic mirror and re-imaged onto the sample via a tube lens (U-TLU, Olympus) and a 10x water-immersion objective, NA 0.60 (Olympus XLPLN10XSVMP). Fluorescence was collected by the objective and separated from the excitation by a multi-band dichroic mirror (IDEX, Di01-R405/488/594-25×36, three bandpasses). The fluorescence was then imaged onto a sCMOS camera (Hamamatsu Orca Flash 4.0) with the appropriate emission filter for the orange (Chroma, ET645/75m, bandpass) and blue (Chroma, ET525/50m, bandpass). Voltage-imaging recordings were acquired at a 1 kHz frame rate unless stated otherwise.

To enhance the spatial resolution of the high temporal resolution movies, we mapped the timing data onto a static 2P structural image of either eYFP (from the LR-CheRiff-eYFP fusion, *λ*_exc_ = 920 nm) or JFX_608_ HaloTag ligand (*λ*_exc_ = 820 nm, targeting the ^2^S transition). Sometimes we made a structural image using 1P HiLo imaging (e.g., Fig. 4a).^66^ A 25x water-immersion objective, NA 1.05 (Olympus XLPLN25XSVMP2) was used to increase the spatial resolution. Maximum intensity projections of z-stacks were used to form images of the dendritic arbor. The distance for each recording site is a slight underestimate of the true on-path distance from the soma because we ignored changes along the z-axis.

For the experiments in Fig. S4, we created the DMD pattern by outlining a region of interest (ROI) around the soma or a single dendritic branch. This pattern was then sequentially shifted by 10 µm increments, up to a total of 100 µm from the original ROI. The shifting of the DMD pattern was triggered with digital clock pulses, cycling through the pre-defined set of patterns three times. The responses from these three cycles were averaged.

### Pharmacology

Drugs were prepared as frozen stock solutions (stored at -20 °C). Drugs were: picrotoxin (Abcam), D-(-)-2-Amino-5-phosphonopentanoic acid (D-AP5, Tocris), tetrodotoxin (TTX, Abcam), NiCl_2_ (Sigma), BaCl_2_ (Sigma), and 4-Aminopyridine (4-AP, Abcam). Ni^2+^ is considered a non-selective VGCC blocker, but T-type (Ca_V_3.x) and R-type (Ca_V_2.3) are sensitive at the used concentration of Ni^2+^ (100 µM; Fig. 6a,b).^67^ Drugs were mixed with ACSF and perfused over the slice for at least 15 min prior to measurements.

All treatment groups were interleaved with control experiments. Statistical significance was assessed using (two-tailed) paired or unpaired Student’s *t*-tests or one-way ANOVA with Bonferroni’s *post hoc* test as appropriate; the level of significance is denoted on the figures as follows: \**p* < 0.05, \*\**p* < 0.01 and \*\*\**p* < 0.001. The experiments were not randomized, and the investigators were not blinded to the experimental condition. Sample size was based on reports in related literature and was not predetermined by calculation.

**Image analysis**

All analysis was performed in MATLAB, as described below.

### Mapping functional data onto structural data

We reconstructed voltage in three dimensions by combining 2P anatomy scans and 1P voltage imaging functional datasets.

#### Image registration

A high-resolution two-photon z-stack of the cell was acquired with z-spacing of 2 μm (Fig. S6). A wide-field epifluorescence image was also acquired and used to define a DMD mask to restrict the 594 nm illumination to the cell and its immediate neighborhood. Patterned illumination substantially decreased background autofluorescence and increased the contrast of the dendrites. We then registered the 2P z-stack to the 1P image. First we performed registration in the x-y plane. We manually defined control points on in-focus parts of the 1P image, and on corresponding structures on a maximum-intensity projection of the 2P z-stack. We used a second-order polynomial fit to map the 2P z-stack onto the 1P image. We then used a cross-correlation approach to find the z-planes in the 2P z-stack which best matched the region around each in-focus control point on the 1P image. We used these coordinates to define rotations and translations out of the image plane, to register the 2P z-stack to the in-focus parts of the 1P image.

#### Determining the microscope model, *M*

We then built a forward model to simulate the 1P image from the 2P z-stack. We assumed a Gaussian beam point-spread function (PSF) with width at focus *w_0_* and confocal parameter *b* (while the Gaussian beam approximation is not strictly correct for the high NA objective, we found that in the presence of tissue light scattering this parameterization of the PSF was adequate). We convolved each plane of the registered 2P z-stack with the corresponding PSF to create a stack 2P_blur_, where each slice was blurred in accordance with its distance from the focal plane. Summing these blurred 2P-images with a depth-dependent weight (to account for signal attenuation) reproduced the 1P image. The depth-dependent PSF and weighting comprised the microscope model *M*.

#### Mapping 2D images to 3D voltages

To enforce the biophysical constraints of continuity and smoothness in the dendritic voltage, we approximate the voltage along the entire tree by linearly interpolating between a small number (a few hundred) of nodes. For the results in Fig. S6 and Movie 2, we used 106 nodes, with a maximum spacing between nodes of 50 μm. Nodes were placed at each branch point and end point of the segmented dendritic tree, to ensure that the model could capture voltage differences between branches.

The assumption of low-dimensional voltage dynamics (e.g. 106 dimensions in Fig. S6) is analogous to the assumptions made when using matrix factorization methods ^37,68,69^ to denoise calcium or voltage imaging movies, and allows us to solve the challenging inverse problem of inferring 3D voltages given 2D measurements (Fig S6a). Below, we describe a simplified version of the method and refer the reader to Ref. 70 for details.

We preprocessed the voltage movie to remove changing baselines by fitting a b-spline to each pixel, and subtracting this baseline estimate.^37^ We then segmented the anatomy stack using Ilastik^71^ to yield a 3D voxelized model of the apical dendrite and soma (our segmentation did not resolve the axon or basal dendrites).

We then skeletonized this 3D model of the dendritic tree into a set of nodes and edges using the NeuTube tracing software.^72^ Edges exist only between nodes which are connected on the dendritic tree. Combined, the nodes and edges form a tree-structured graph that will be used when solving the voltage inference problem. Crucially, the number of nodes is far fewer than the number of voxels in the tree, thus implementing our assumption of low dimensionality.

Let *Q* be the number of voxels in the segmented neuron and *P* be the number of nodes in the skeletonized graph. We define the *Q* × *P* basis matrix *B* to map from node weights to voxel voltages. The columns of *B* have a simple interpretation: they are smooth basis functions used to reconstruct the voltage. The node weights in a given frame, *w_t_*, control the relative intensities of these basis functions (Fig. S6b). Using *W* to refer to the *P* ×*T* matrix of node weights across all frames, we have *v^* = *BW*, where the *Q* ×*T* matrix *v^* represents our three-dimensional voltage estimate across all frames.

To account for the blurring and attenuation created by the 1P microscope, we pass the three-dimensional voltage estimate through the microscope model *M*. The microscope model performs a depth-dependent blur and summation (Fig S6e). A simplified version of the model is therefore:

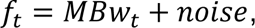

where *f_t_* is a (preprocessed) voltage imaging frame (*f* standing for fluorescence), and *w_t_* are the (unknown) node weights.

To infer the node weights *w_t_*, we use the graph of nodes to solve a regularized inverse problem (Fig S6a). The regularizer is given by:

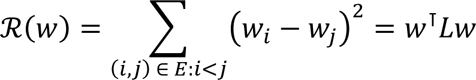

where *L* is the graph-Laplacian. This regularizer encourages adjacent nodes to have similar weights. With this regularizer, the (simplified) optimization problem used to infer node weights is:

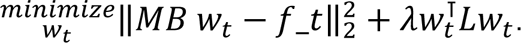

We use generalized cross validation (GCV^73^) to select the penalty parameter *λ* which best describes the data.

For brevity, we have omitted some complexities of the model in this summary. The full details of our voltage reconstruction can be found in Ref. 70. In practice, we found it helpful to include gain and offset terms to account for non-uniform indicator expression and baseline fluorescence. The full model jointly estimates these parameters along with the node voltages. We fit the model iteratively, alternating between estimating voltages and optimizing gains and offsets. This approach yields 3D voltage estimates *v^* for each frame (Fig S6f,h). We then passed these estimates through the microscope model $M$ to create the reconstructed frame shown in Fig. S6g.

### PCA-based filtering

Electrical length constants extended over many pixels, so we used Principal Component Analysis (PCA)-based filtering to remove pixel-wise shot noise, similar to the approach in Ref. ^37^. We found that > 96% of the variance in the recordings was contained within the first three principal components, whose spatial footprints broadly corresponded to the soma-proximal region, the dendritic trunk, and the distal dendrites, supporting the use of PCA to remove high spatial-frequency noise (Fig. S7). The first five temporal components contained signals related to neural activity and were used to resynthesize a denoised movie; the remaining components represented uncorrelated shot noise. To verify that the PCA filtering did not distort the underlying AP waveforms, we compared mean AP waveforms in subcellular compartments before and after the filtering steps. We observed no systematic deviations in the AP waveforms in the soma or dendrites (Fig. S7).

### Extracting fluorescence from movies

Fluorescence values were extracted from raw movies in one of two ways. One approach used the maximum-likelihood pixel-weighting algorithm described previously.^32^ Briefly, the fluorescence at each pixel was correlated with the whole-field average fluorescence. Pixels that showed stronger correlation to the mean were preferentially weighted. This algorithm automatically found the pixels carrying the most information, and de-emphasized background pixels. Alternatively, a user defined a region comprising the soma and dendrites and calculated fluorescence from the unweighted mean of pixel values within this region. These two approaches gave similar results. Photobleaching was corrected by dividing the frames by a regression fit to the mean fluorescence.

### Spike-triggered average (STA) movies

A simple threshold-and-maximum procedure was applied for spike detection. Fluorescence traces were first high-pass-filtered, and a threshold was manually selected. Once spike times were determined, movie segments surrounding each spike were averaged together. To calculate stimulus-triggered averages, traces or movies were aligned to the optogenetic stimulus onset.

### Sub-Nyquist Action Potential Timing (SNAPT)

Spike propagation delay was calculated using the SNAPT subframe interpolation algorithm as described previously.^32^ In brief, spike-triggered average (STA) movies were used as a template and fit with a quadratic spline interpolation of the spike waveform, pixel-by-pixel. For each pixel, we calculated the sub-frame interpolated time that the spike reached 50% of local maximum, to create a map of the spike delay. The fits were then converted into movies. Spike timing at each pixel was represented by a brief flash, which followed a Gaussian time-course with duration equal to the cell-average time resolution, *σ*.

The pixel matrix of the subframe interpolated movie was expanded to match the dimensions of the high-resolution image, and the amplitude at each pixel was then set equal to the mean brightness at that pixel. For assembly of the color movies, the timing signal was assigned to a color map that was overlaid on a grayscale image of mean fluorescence. The optically stimulated region of the cell was highlighted in blue.

### Normalization to reference signal (ΔF/F_ref_)

To account for sub-cellular variations in voltage sensitivity (likely due to variations in Voltron2 trafficking), we normalized the fluorescence signal associated with a spike, ΔF, by the amplitude of the change in fluorescence during the passive return to baseline after a stimulus (i.e., ΔF/F_ref_; Fig. 1d). ΔF/F_ref_ was used for all spike amplitude heatmaps and kymographs unless stated otherwise. By comparing two voltage-sensitive signals, this measure was insensitive to background and protein trafficking.

### Kymographs

We manually drew a line along the apical dendrite and then determined the mean fluorescence time-course in equal-length segments along the line. Typical segment size was 10 pixels (6.5 µm with 10x objective). The fluorescence waveforms were assembled into a kymograph matrix showing signal amplitude as a function of linear position and time. Example waveforms were calculated by averaging responses from 5 segments (∼33 μm contour length) and plotted on top of the kymographs.

### Counting bAPs and dSpikes

All spikes at the soma were counted as back-propagating action potentials (bAPs). The timing of each bAP was estimated relative to the optogenetic stimulation onset. Dendritic spikes (dSpikes) were detected by high-pass-filtering and simple threshold-and-maximum in a user-defined region (typically > 300 µm from soma). We defined dSpikes as a large and narrow discharge (typically < 5 ms in full width at half maximum). Dspike successes and failures were clearly distinguished in the fluorescence traces (e.g., Fig. 2a and Fig. S10).

### Period doubling bifurcation

We typically observed period doubling during wide-field optogenetic stimulation (Fig. 4c-e). Overly strong stimulation frequently led to the failure of all spikes, likely due to incomplete recovery of Na_V_ channels (i.e. depolarization block). In preliminary experiments, we determined a stimulation intensity approximately halfway between rheobase and depolarization block. The frequency of bAPs was estimated by measuring the time interval between bAP peaks, while the amplitude was normalized to the average of the final 5 bAPs in a stimulus epoch (Fig. 4e).

### Normalization of plateau potential area

The cumulative area under the curve (AUC) was determined by integrating fluorescence changes, ΔF, with respect to time (Fig. S22). In Fig. 5b, the normalized area (% sum) was calculated as AUC for combined stimulation (AUC_combined_), divided by the sum of the AUC for optogenetic stimulation alone (AUC_optical_) and the AUC for EFS alone (AUC_electrical_). In Fig. 5d, the normalized area (% peak) was calculated by mapping the AUC for combined stimulus for each cell vs. ΔTime to the range [0, 1] (Fig. S22b-c). In Fig. 6b, the normalized area (% baseline) was determined as the ratio of AUC to baseline AUC prior to any vehicle or drug treatment.

## Biophysical modeling

### Simulating a CA1 pyramidal cell in NEURON

Morphologically realistic simulations were carried out on an AMD64-Windows computer using NEURON^74^ through its Python interface (Python 3.11, NEURON 8.2) and exported into MATLAB to analyze and compare with experimental data. Model specifications and simulation code are available as Supplementary Files.

### Model properties

We adapted an existing CA1 pyramidal cell model (ModelDB accession number: 116084)^49^. Our model uses the same morphology as Ref. ^49^. We added a slow inactivation gate to the Na_V_ channels, varied the distributions of Na_V_ channels and A-type K_V_ channels, added a gradient in the maximal Na_V_ inactivation from soma to the distal dendrites, and introduced channelrhodopsin in the somatic and dendritic compartments. The distribution of delayed rectifier K_DR_ channels was homogeneous throughout the cell, as in Ref. ^49^. We sought to replicate the voltage profiles under pure optogenetic stimulation, so our model did not contain VGCC or NMDAR conductances. Table 1 shows values of the model parameters.

**Table 1:**
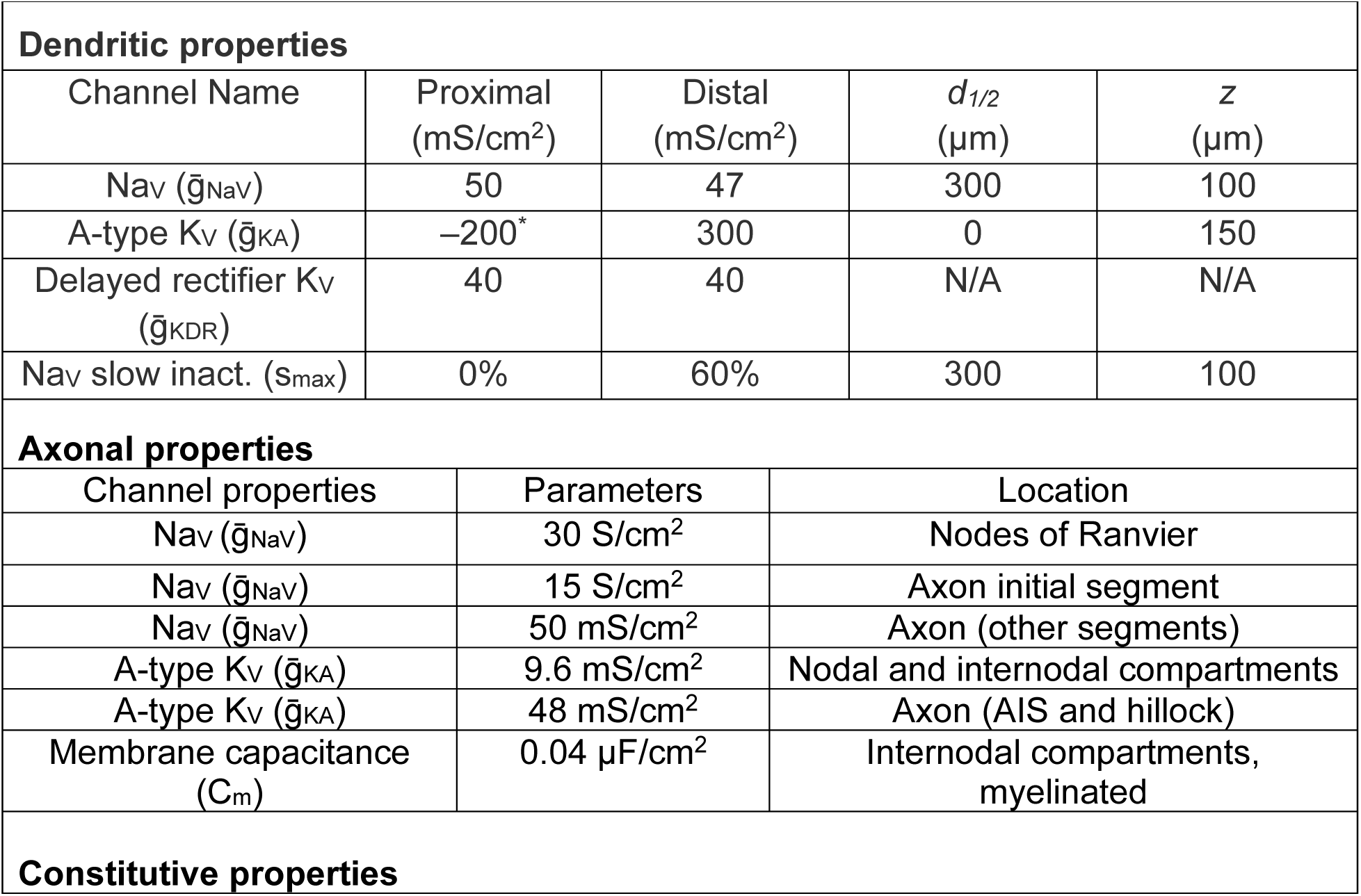
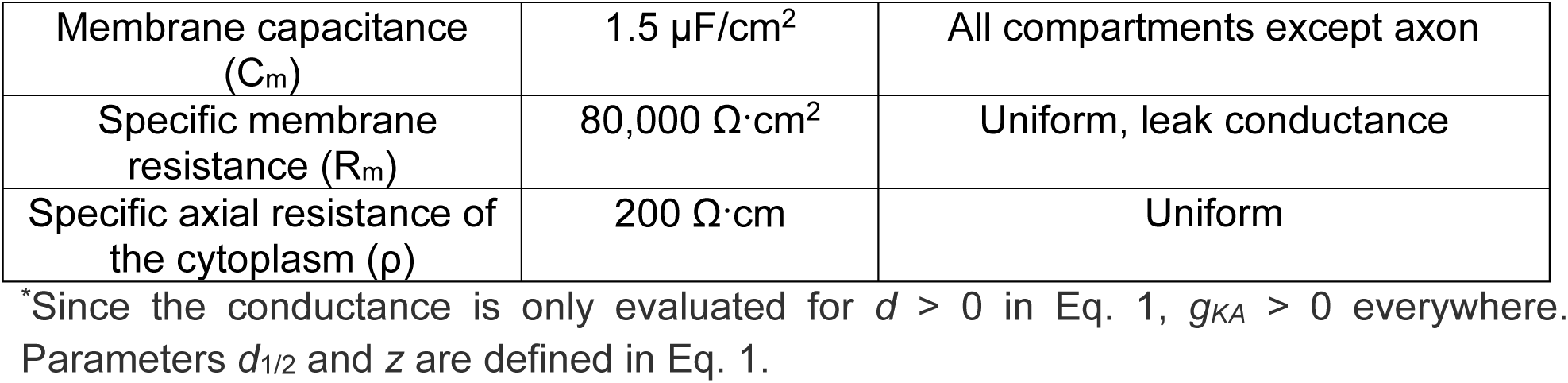
Ion channel densities in a model CA1 pyramidal cell. Channel models were adapted from Ref. ^49^.

| Dendritic properties |  |  |  |  |
| --- | --- | --- | --- | --- |
| Channel Name | Proximal<br>(mS/cm<sup>2</sup>) | Distal<br>(mS/cm<sup>2</sup>) | $d_{1/2}$<br>(μm) | $z$<br>(μm) |
| Nav ($\bar{g}_{Nav}$) | 50 | 47 | 300 | 100 |
| A-type Kv ($\bar{g}_{KA}$) | −200* | 300 | 0 | 150 |
| Delayed rectifier Kv<br>($\bar{g}_{KDR}$) | 40 | 40 | N/A | N/A |
| Nav slow inact. ($s_{max}$) | 0% | 60% | 300 | 100 |
| Axonal properties |  |  |  |  |
| Channel properties | Parameters |  | Location |  |
| Nav ($\bar{g}_{Nav}$) | 30 S/cm<sup>2</sup> | | Nodes of Ranvier | |
| Nav ($\bar{g}_{Nav}$) | 15 S/cm<sup>2</sup> | | Axon initial segment | |
| Nav ($\bar{g}_{Nav}$) | 50 mS/cm<sup>2</sup> | | Axon (other segments) | |
| A-type Kv ($\bar{g}_{KA}$) | 9.6 mS/cm<sup>2</sup> | | Nodal and internodal compartments | |
| A-type Kv ($\bar{g}_{KA}$) | 48 mS/cm<sup>2</sup> | | Axon (AIS and hillock) | |
| Membrane capacitance<br>($C_m$) | 0.04 μF/cm<sup>2</sup> | | Internodal compartments,<br>myelinated | |
| Constitutive properties |  |  |  |  |

|  |  |  |
| --- | --- | --- |
| Membrane capacitance ($C_m$) | 1.5 $\mu\text{F}/\text{cm}^2$ | All compartments except axon |
| Specific membrane resistance ($R_m$) | 80,000 $\Omega\cdot\text{cm}^2$ | Uniform, leak conductance |
| Specific axial resistance of the cytoplasm ($\rho$) | 200 $\Omega\cdot\text{cm}$ | Uniform |

\*Since the conductance is only evaluated for $d > 0$ in Eq. 1, $g_{KA} > 0$ everywhere. Parameters $d_{1/2}$ and $z$ are defined in Eq. 1.

For voltage-gated channels, the maximum channel conductance densities were modeled as sigmoidal functions of the contour distance, *d*, from the soma. These sigmoid distributions take the general form shown in Equation 1, with asymptotic densities *g_Prox_* and *g_Distal_*, a half-way distance *d_1/2_* (where the density is (*g_Prox_* + *g_Distal_*)/2) and a steepness parameter *z*.

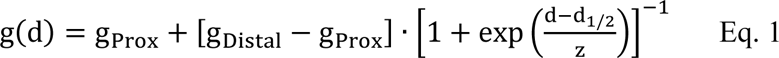

The *g* parameters were selected to agree with literature values,^11^ while the other parameters were adjusted to match our data. Fig. S18a-c show channel densities of Na_V_ and A-type K_V_ across the neuron morphology after tuning to match our data. For temperature-dependent channel gating parameters, we used a temperature of 35 °C.

In the axonal compartments channel densities and membrane capacitance were set separately to account for differences in channel targeting, myelination, and clustering of Na_V_ channels at the axon initial segment (AIS) and nodes of Ranvier. Table 1 summarizes the model parameters.

### A-type K_V_

A-type potassium channels are present in the soma and dendrites, with approximately six-fold higher density in the distal apical dendrites than the soma.^75^ Proximal (*d* < 100 μm) and distal (*d* > 100 μm) channels differed slightly in the kinetics and voltage dependence of the activation variable, to reproduce measured channel properties in

CA1 pyramidal dendrites.^11,62^ The detailed kinetic properties of the two types of A-type K channels are given below, where *v* is in mV, *I_KA_* is in µA/cm^2^ and time constants *τ* in ms.

Kinetic scheme for proximal A-type K_V_ channels (d < 100 μm):

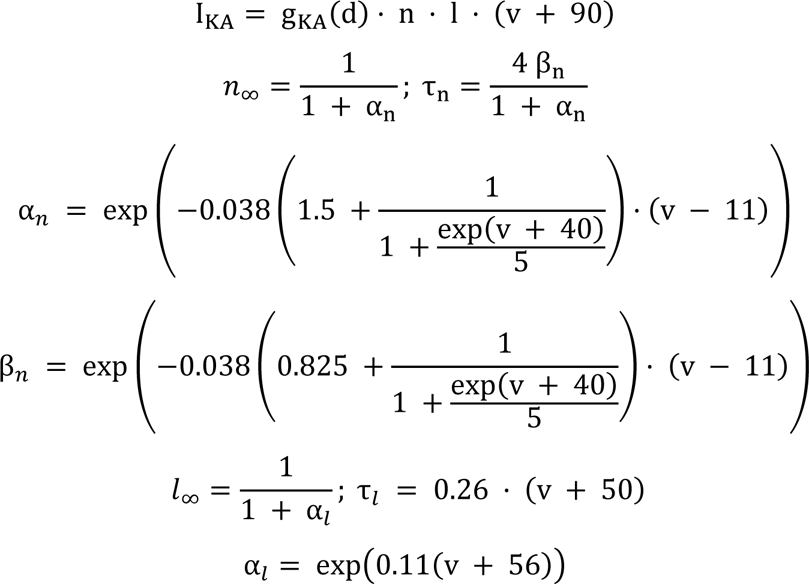

Time constants are constrained to *τ_n_* ≥ 0.1 ms and *τ_l_* ≥ 2 ms.

For distal A-type K_V_ channels (*d* > 100 μm) the kinetic scheme is similar, with the replacements:

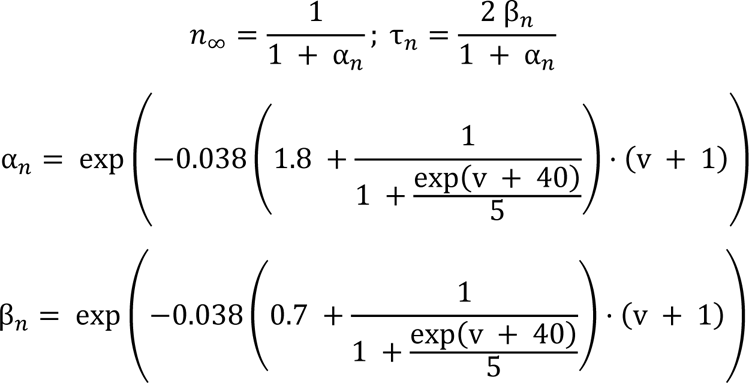

### Voltage-gated sodium channel, Na_V_

To capture the loss of dendritic excitability after several dSpikes, we used a Hodgkin-Huxley-type model with a slow inactivation variable, *s*, and assumed that a fraction, *a_r2_*, of Na_V_ channels could undergo slow inactivation (e.g. *a_r2_* = 70% means that at most 70% of Na_V_ channels can undergo slow inactivation). We further assumed that *a_r2_* followed a sigmoid function of *d* of the form of Eq. 1, with *a_r2_* = 0 (i.e. no slow inactivation) at the soma. Fig. S18d shows the steady state value of *s*, as a function of membrane voltage and *a_r2_*. Steady-state Na_V_ inactivation is thus modeled as a mixture of inactivating and persistent currents as s_∞_ (V, a_r2_) = a_r2_ ⋅ s_∞_ (V, 100%) + (1 – a_r2_), where s_∞_ (V, 100%) is the voltage dependent equilibrium value of the fully inactivating channels.

Kinetic scheme for Na_V_ channels with slow inactivation:

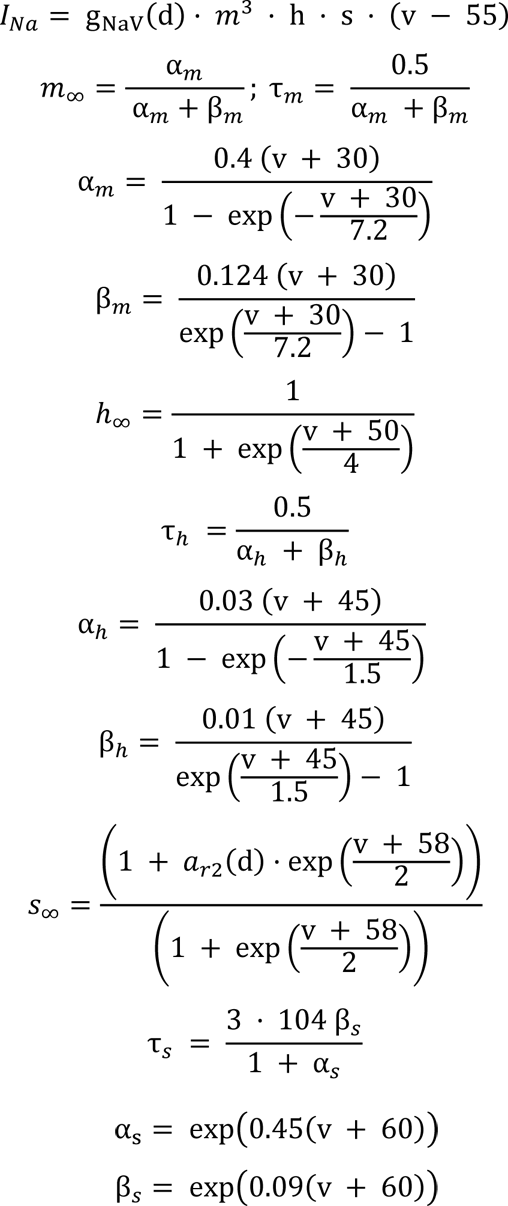

Na_V_ time constants are constrained to *τ_m_* ≥ 0.02 ms, *τ_h_* ≥ 0.5 ms and *τ_s_* ≥ 10 ms.

### Delayed rectifier potassium, K_DR_

Delayed rectifier K_V_ channels are uniformly distributed across the neuron as in ref. 41.

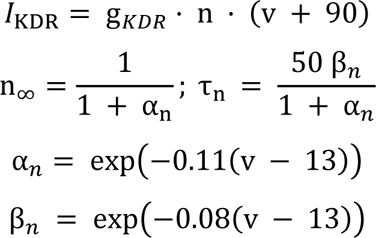

The K_DR_ time constant is constrained to τ_n_ ≥ 2 ms.

### Passive properties

The passive leak conductance (*V_rev_* = –66 mV) was uniform across the soma and dendritic arbor. Dendrites were modeled as smooth cylinders. To account for the excess surface area due to dendritic spines, the effective membrane capacitance was set to 1.50 μF/cm^2^, larger than the geometrical membrane capacitance of ∼1 μF/cm^2^.^76^

### Channelrhodopsin

Optogenetic stimulation was implemented using distributed time-dependent conductances with reversal potential 0 mV. The conductance was proportional to the simulated blue light illumination profile.

### Parameter fitting

The nonlinear interactions of Na_V_, K_DR_ and K_A_ channels produce a rich variety of subthreshold and spiking patterns, even in models with one or a few compartments. Also, a given electrical response pattern can often arise from multiple combinations of ion channel parameters. As our model uses realistic neuronal morphology, the parameter space grows exponentially as each compartment can have different channel parameters.

To reduce the size of the parameter search space we introduced constraints from literature data wherever possible, and reduced the number of spatially dependent parameters by imposing the parameter distributions of Equation 1.

Parameter space was mapped using sweep searches, varying optogenetic stimulus intensity, somatic and distal Na_V_ densities and the level of Na_V_ slow inactivation. The voltage traces were then classified based on the patterns of bAP successes and failures and arranged as phase diagrams (Fig. S19c-e). The threshold to distinguish bAP successes from failures was whether the voltage at a compartment at 500 μm from the soma reached a peak voltage above or below -40 mV.

### Model validation

The model was originally used to simulate responses to patch clamp and synaptic stimulation.^49^ To validate our version of the model, we first compared a localized current-clamp stimulus at the soma vs. distributed channelrhodopsin conductance at the soma (g_Distal_ = 0, *d_1/2_* = 40 μm, *z* = 10 μm). For similar net currents, these two stimulation methods yield similar spiking and bAP patterns, which also matched our experimental results (Fig. 4f-g). Under strong somatic or weaker widefield stimulation, the model also recreated the dendritic period doubling behavior observed experimentally.

To further validate the model, we compared simulation results to data on patch clamp current injection in oblique dendrites.^38^ We simulated 2 nA current injections and observed dendrite-localized dSpikes similar to experimental observations (Fig. S19a). Turning off the dendritic Na_V_ channels mimicked the effect of adding TTX in the experiments. We then simulated the same dendrite under optogenetic stimulation (*d_1/2_* = 50 μm, *z* = 10 μm, g_distal_ = 0, g_oblique_ = 1 mS/cm^2^ centered on an oblique dendrite as shown in Fig. S19b). The optogenetic stimulation resulted in AP starting at the soma and backpropagating into the dendritic arbor, with a bAP amplitude pattern consistent with our experimental results for stimulation of an oblique dendrite.

### Defining channel reserve

To determine the biophysical basis of our experimentally observed dSpike time window, we defined channel reserves for Na_V_ and A-type K_V_ channels. The channel reserve is the fraction of ion channels that could transition to the open state due to a bAP. For Na_V_ channels, the slow inactivation variable *s* measures the channel reserve. For example, at *s* = 0.6, only 60% of Na_V_ channels could participate in amplifying a bAP. In A-type K_V_ channels, the inactivation variable *l* plays a similar role. Tracking the channel reserves for these two channels indicated that a depleted A-type reserve and un-depleted Na_V_ reserve were necessary for dSpikes to occur.

### Simplifications and omissions

CA1 pyramidal cells contain diverse Na_V_ channels due to different genes, subunit compositions, and post-translational modifications,^77^ whereas we used a single Na_V_ with a gradient in slow inactivation. The mechanism of slow inactivation and the factors driving a spatial gradient in this parameter have not been fully elucidated but post-translational modification by protein kinase C has been implicated.^78^ The sigmoidal channel distribution of Eq. 1 may miss effects from finer-grained subcellular variations in channel density. Our channelrhodopsin model does not include the light-dependent opening kinetics of CheRiff, the conductance sag under continuous illumination, or finite closing kinetics.

Several classes of ion channels were omitted from our model, including calcium-activated potassium channels (K_CA_), VGCCs, NMDARs and hyperpolarization-activated cyclic nucleotide-gated (HCN) channels. The omission of these channels is justified because we focused on the short-time dynamics of bAP filtering under pure optogenetic drive (Fig. 2-4), and the model accurately captured the observed dynamics under these conditions. VGCCs and NMDARs drive apical calcium plateaus and initiate complex spikes under synaptic inputs. HCN-channels are preferentially targeted to the dendritic tuft and could play a role similar to the A-type K_V_ channels in gating back-propagation.^79^ These channels may also be important in mediating neuromodulatory effects, a variable not explored in this work.

### Coupled two-compartment Izhikevich model

As a complement to the morphologically and physiologically detailed CA1 model, we introduced a two-stage resistively coupled Izhikevich model (Fig. S20). Despite its simplicity, this model broadly reproduced the bAP filtering characteristics of the CA1 dendrites. The classical Izhikevich model is a computationally efficient spiking neuron model consisting of a voltage variable *v* and a slow adaptation variable *u*, and is capable of reproducing a wide variety of neuronal behaviors observed in mammalian brains.^80^ These characteristics made it an attractive starting point for a simplified CA1 model.

In our adaptation of the Izhikevich model, the soma was tuned to exhibit regular spiking without adaptation. The soma was resistively coupled to the dendrite compartment that featured an adjusting spiking threshold. Optogenetic-like stimulation was implemented as variable conductances with 0 mV reversal potential at the soma or dendrites. When the model was driven by optogenetic stimulation at the soma only, it replicated the behavior seen in Fig. 4f. A single spike from the soma did not depolarize the dendrites enough to trigger a spike, but a spike train from the soma was able to evoke dendritic spikes. The spike-threshold adaptation in the dendrites mimicked the effect of Na_V_ inactivation, causing the dendrites to lose excitability after several successful spikes. The distal dendrites thereby acted as a high-pass filter or accelerometer, selectively responding to a small number of somatic spikes following a step increase in somatic firing rate. Simultaneous stimulation of soma and dendrites recreated the experimentally observed period-doubling bifurcation shown in Fig. 4g.

Model specifications and simulation code are available as Supplementary Files.

---

**Fig. S1.**
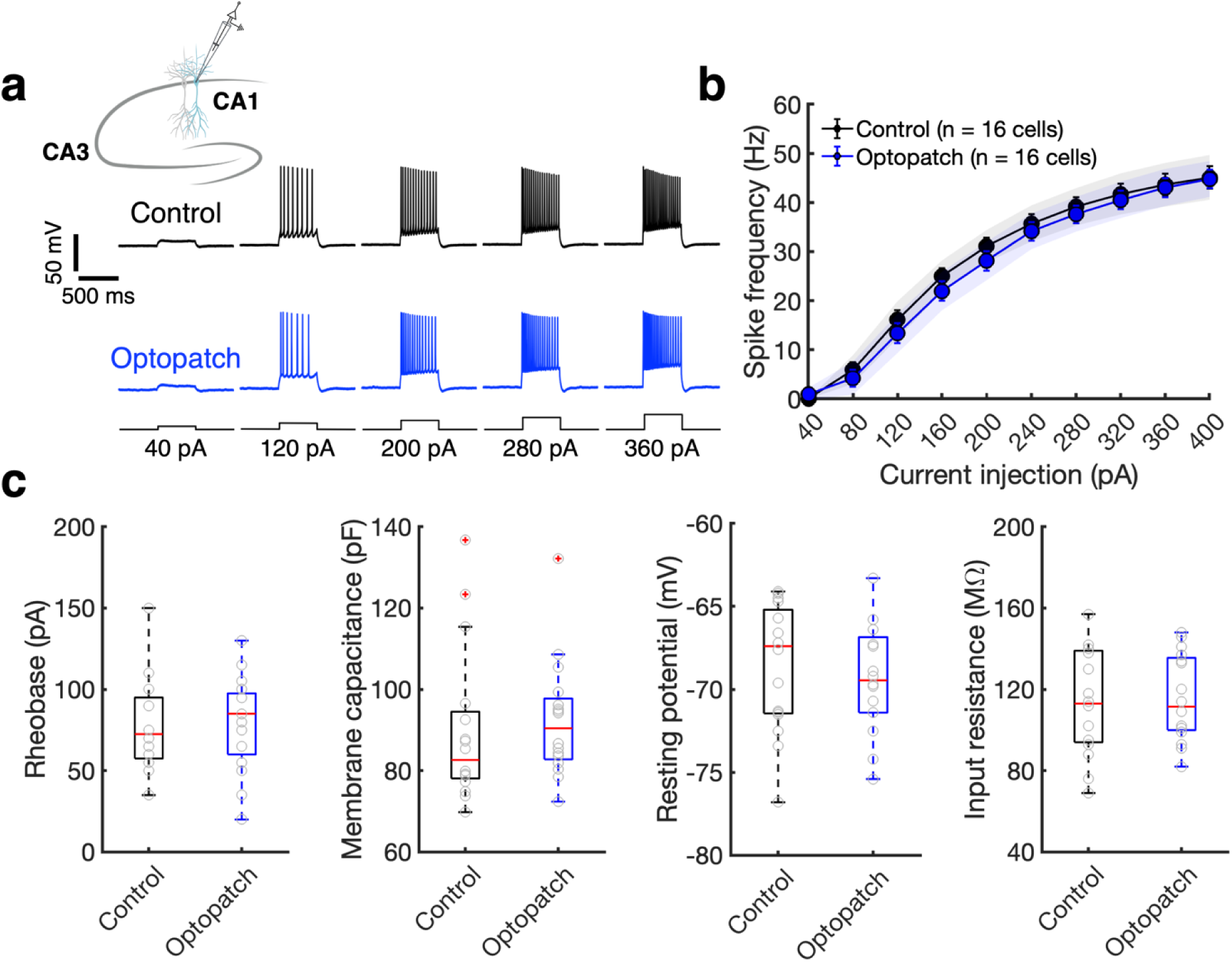
Effects of Optopatch expression on intrinsic electrical properties. **a,** Representative patch clamp recordings comparing the voltage responses of Optopatch-expressing neurons to their neighboring non-expressing neurons, elicited by 500 ms current injections. These observations were made in acute hippocampal brain slices. **b-c,** Pooled data on intrinsic membrane properties and excitability (n = 16 cells from 6 animals for each group). In **(b)**, error bars show the SEM, while shaded areas indicate the 95% confidence intervals for the mean. The box plots show the median, 25^th^ and 75^th^ percentile, with outliers (indicated by ‘+’ symbols) excluded. No significant difference was found between the groups (*p* > 0.05, two-sided paired Student’s *t*-test). Related to Fig. 1.

**Fig. S2.**
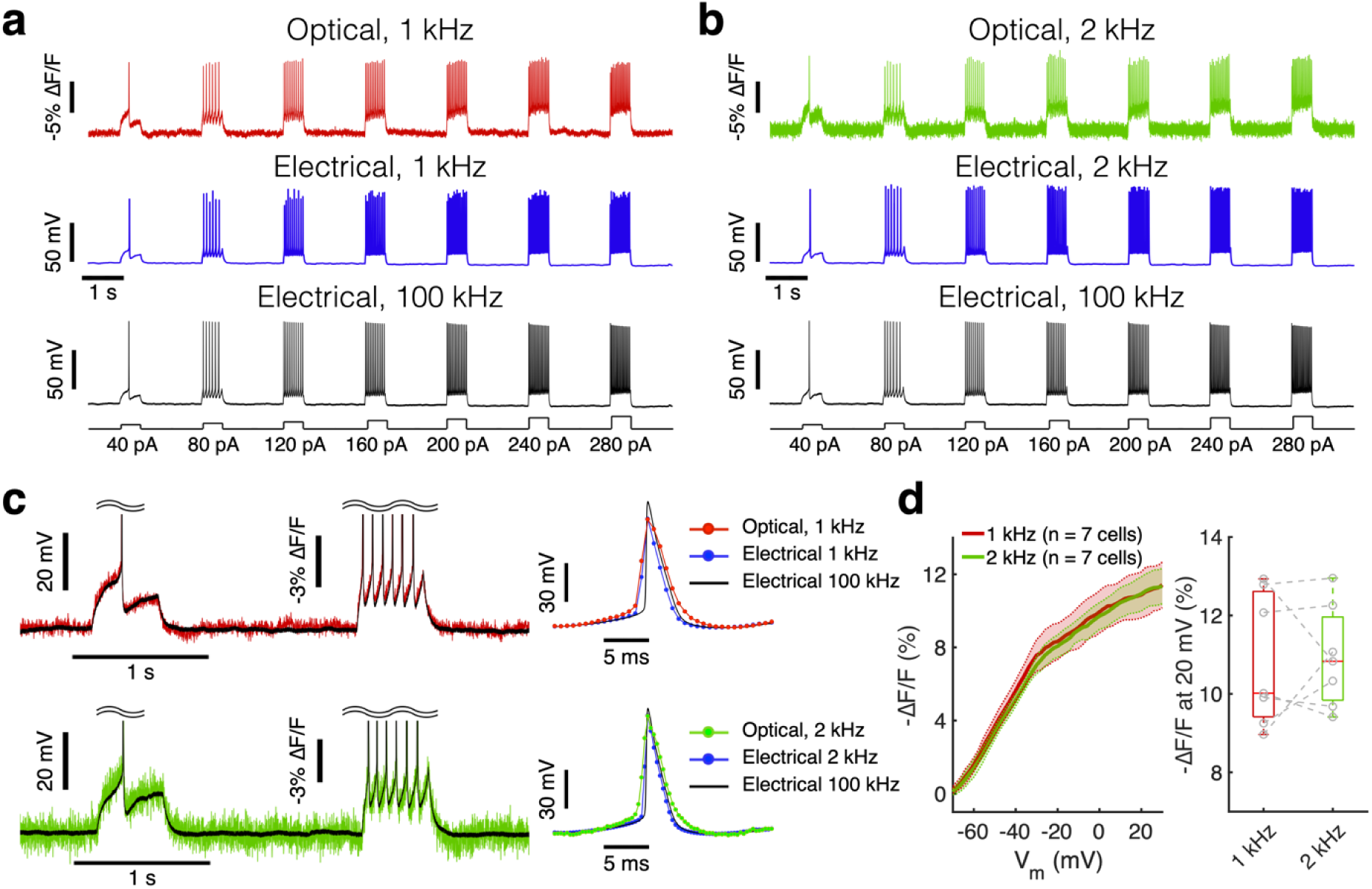
Characterization of LR-Voltron2. **a,** Concurrent fluorescence (red; frame rate 1 kHz) and current clamp recordings (black; sample rate 100 kHz) in CA1 neurons expressing the Voltron2-CheRiff Optopatch construct. Spikes were evoked by a 500 ms current injection. The 100 kHz electrical trace was down-sampled (blue) to illustrate the low-pass filtering effect of the 1 kHz camera frame rate. **b,** Similar experiments performed on the same cell, but with a 2 kHz camera frame rate (green), and the 100 kHz electrical trace down-sampled to 2 kHz. **c,** Overlay of raw (left) and spike-triggered average waveforms (right; 136-139 spikes) for the optical and electrical traces from **(a)** and **(b)**. **d,** Left: Relation between fluorescence and membrane voltage (n = 7 cells from 4 animals). The shaded areas indicate the 95% confidence intervals for the mean. Right: Fluorescence changes upon a voltage step from -70 to +20 mV were the same for 1 kHz vs. 2 kHz camera frame rates (*p* > 0.05, two-sided paired Student’s *t*-test). Box plots show the median, 25^th^ and 75^th^ percentile. The single-trial optical noise-equivalent voltage was 1.2 ± 0.5 mV at 1 kHz and 2.5 ± 0.8 mV at 2 kHz (mean ± s.d., n = 7 neurons, 4 animals). Optically recorded spikes were slightly broadened at 1 kHz compared to 2 kHz (full-width at half-maximum 1 kHz: 3.9 ms, 2 kHz: 3.4 ms, patch clamp digitized at 100 kHz: 2.5 ms). Recordings at both frame-rates clearly resolved every spike, with zero false-positives or false-negatives up to the maximum observed spike rate of 90 Hz (*n* = 7 cells, 2212 spikes). Related to Fig. 1.

**Fig. S3.**
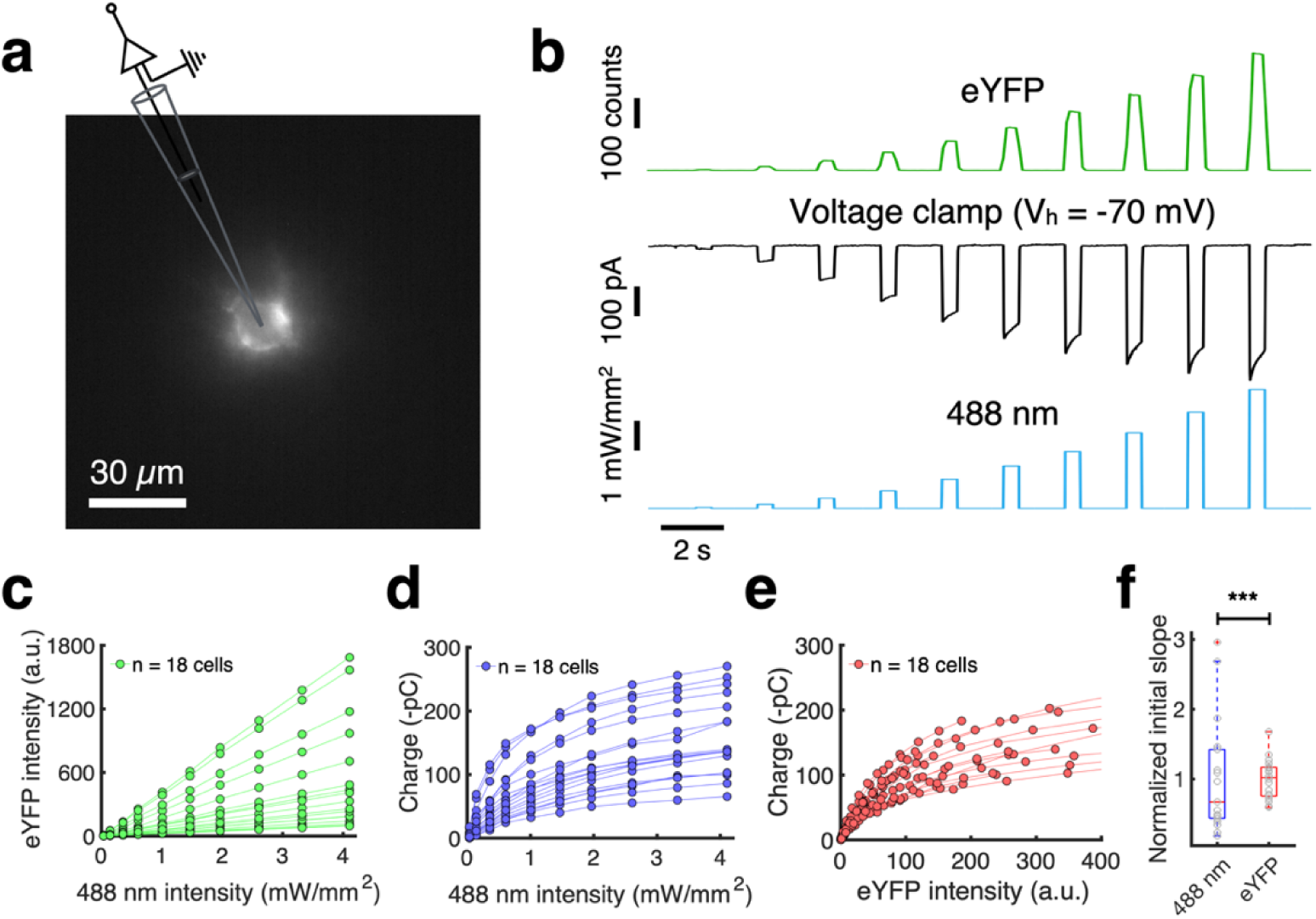
Characterization of LR-CheRiff-eYFP. We sought to calibrate optogenetic stimulus strengths across cells and sub-cellular regions which possibly differed in membrane areas or CheRiff expression levels. We performed optical dosimetry via the eYFP fluorescence in CheRiff-eYFP, reasoning that cumulative eYFP fluorescence would be proportional to photon flux at the locations of the CheRiff molecules. **a,** Epifluorescence image showing a CA1 neuron expressing the Voltron2-CheRiff Optopatch construct. Targeted illumination (488 nm) was directed to the soma using a digital micromirror device (DMD), revealing LR-CheRiff-eYFP fluorescence. **b,** Sample traces from concurrent fluorescence (green; frame rate 10 Hz) and voltage clamp (black; acquisition rate 100 kHz) recordings, showing responses to increasing laser power (blue; 0.5 s duration). Responses from three consecutive trials were averaged. **c,** Relationship between 488 nm illumination intensity and mean eYFP fluorescence intensity. **d,** Relationship between the 488 nm intensity and accumulated electric charge during the illumination pulse. Photocurrent showed saturation behavior, reaching 50% of saturation at *I* = 0.6 ± 0.2 mW/mm^2^ (mean ± s.d., *n* = 18 cells, 6 animals), consistent with prior results.^81^ **e,** Relationship between the cumulative eYFP intensity and the charge. **f,** Pooled data comparing the slope in the linear response regime intensity (*I* < 1 mW/mm^2^) of charge vs. blue light intensity (data from **(d)**) and charge vs. eYFP intensity (data from **(e)**). In both cases, the slopes were normalized to the mean of each group. Dosimetry via eYFP fluorescence had 7.8-fold lower variance than did dosimetry via raw blue laser power (0.084 vs. 0.66, n = 18 cells and 6 animals; *p* < 0.001, F-test). Box plots show the median, 25^th^ and 75^th^ percentile. Related to Fig. 1.

**Fig. S4.**
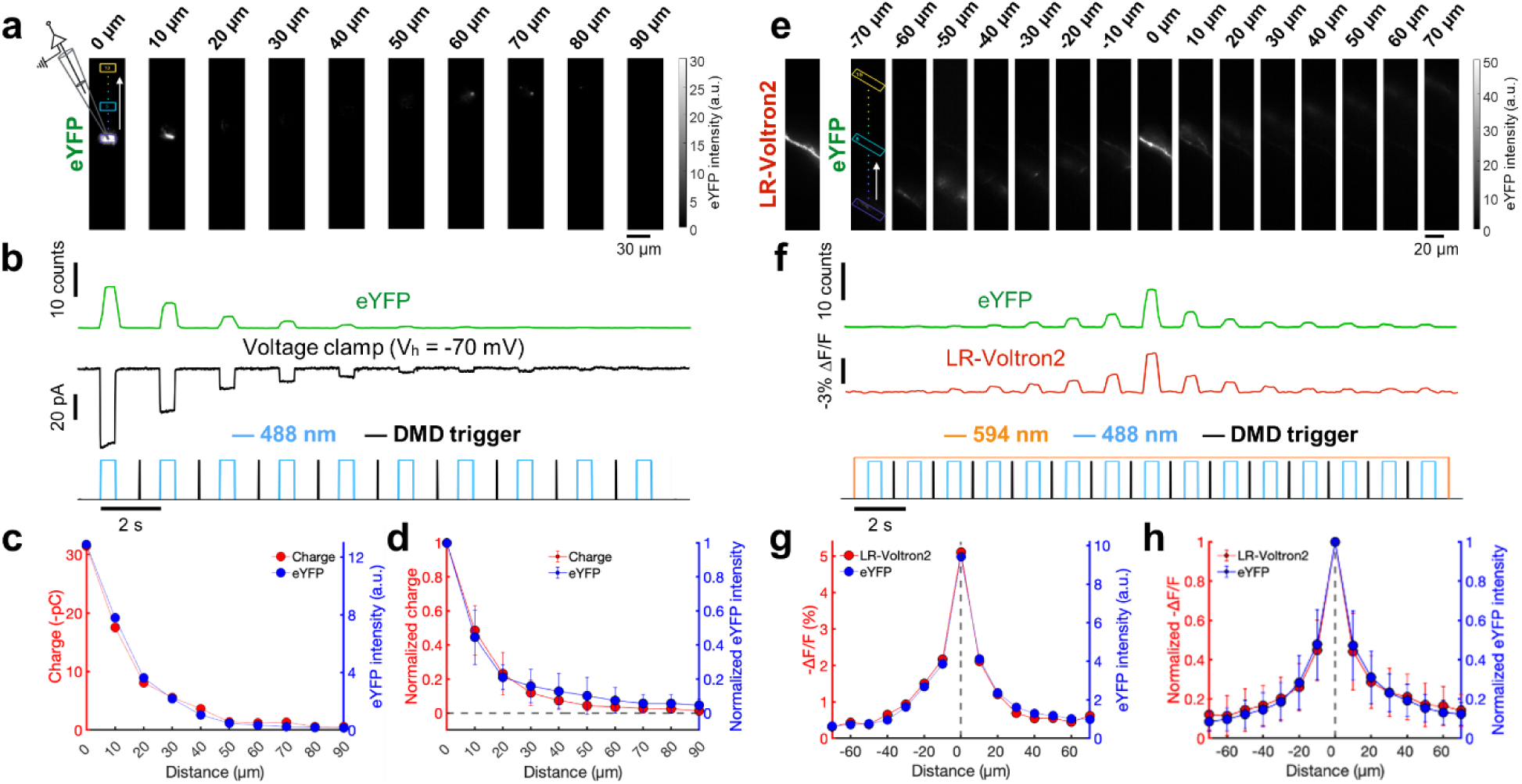
Target-selectivity of patterned 1-photon optogenetic stimulation. To calibrate the targeting specificity of the patterned 1-photon optogenetic stimulation, we projected a series of blue light spots which tiled strips crossing different sub-cellular structures and recorded the electrical responses. **a,** A series of epifluorescence images (all within the same field of view and contrast settings) show the effects of micromirror-patterned 488 nm illumination targeted to the soma of a neuron expressing the Optopatch construct. The illumination pattern was sequentially shifted upward (indicated by a white arrow) in 10 µm increments, up to 90 µm. **b,** Sample traces from concurrent fluorescence (green; frame rate 10 Hz) and voltage clamp (black; acquisition rate 100 kHz) recordings, showing responses to increasing distances of the illumination spot from the soma. The same 488 nm laser power (∼1 mW/mm^2^ for 0.5 s) was used across the measurements. Responses from three consecutive trials were averaged. **c,** Accumulated charge (red) and eYFP fluorescence (blue) as a function of the distance of the blue spot from the center of the soma, for the cell shown in **(a, b)**. **d,** Same as **(c)**, with pooled data (*n* = 10 cells, 4 animals). Data were normalized to the responses observed at 0 µm offset. Error bars represent s.d.. The induced photocurrent fell to half maximum when stimuli were offset by 11 ± 4 μm and to 20% of maximum at 23 ± 7 μm (mean ± s.d.). The eYFP fluorescence was an excellent predictor of photocurrent. **e,** For stimuli crossing distal dendrites, we used the change in Voltron2 fluorescence in the targeted dendrite (indicating local membrane depolarization) as a proxy for stimulus strength. Left: Fluorescence of LR-Voltron2 with targeted 594 nm illumination on a single dendritic branch. Right: Fluorescence of LR-CheRiff-eYFP (all within the same field of view and contrast settings) for a series of 488 nm illumination spots with different transverse displacements from -70 µm to +70 µm relative to the center of the branch, in 10 μm steps. **f,** Representative fluorescence traces from LR-CheRiff-eYFP (green; frame rate 10 Hz) and LR-Voltron2 (red; frame rate 100 Hz), showing responses as a function of blue light offset. The same 488 nm laser power (1 mW/mm^2^ for 0.5 s) was used across the measurements. Averages of three consecutive responses are shown. **g,** Mean fluorescence of LR-Voltron2 (red) and LR-CheRiff-eYFP (blue) as a function of the blue offset for the cell in **(f)**. **h,** Same as **(g)**, with pooled data from 29 branches across 20 cells and 4 animals. Data were normalized to the responses observed at 0 µm offset. Error bars represent s.d.. Evoked responses, |ΔF|, fell to half maximum for stimuli at 11 ± 5 μm offset and to 20% of maximum at 34 ± 14 μm offset (mean ± s.d., *n* = 29 branches, 20 cells, 4 animals). Related to Fig. 1.

**Fig. S5.**
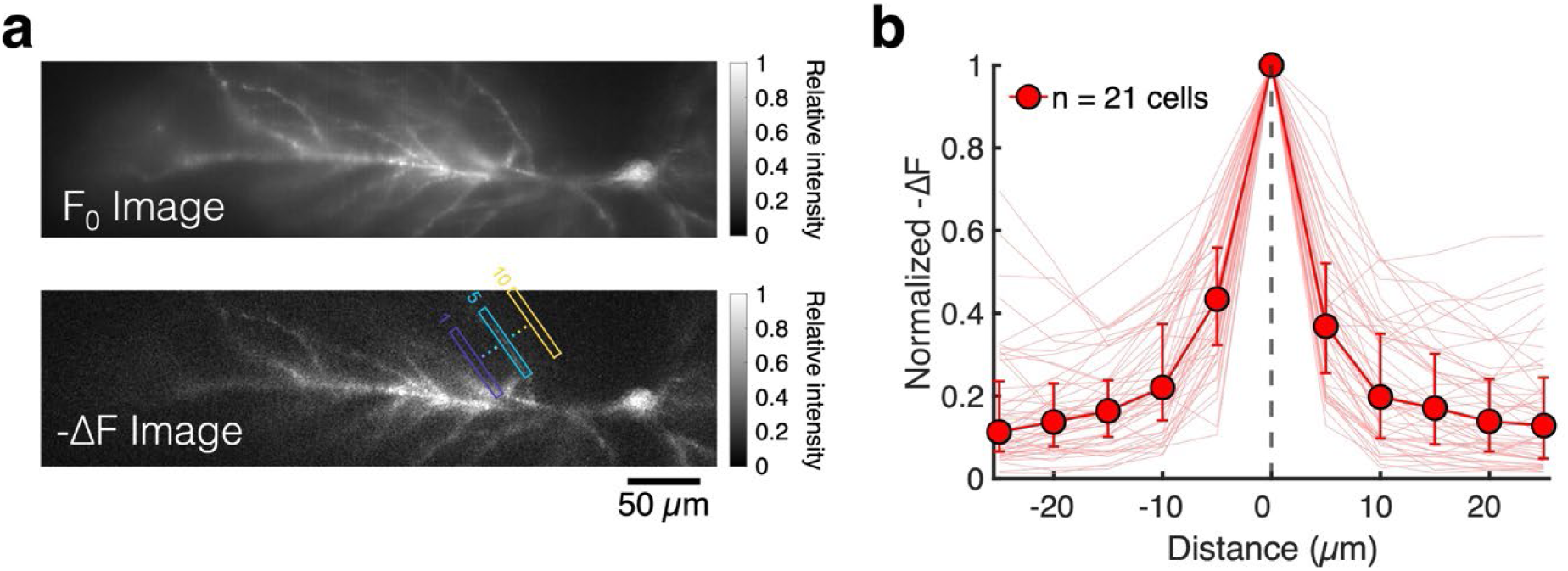
Focal-plane selectivity of 1P voltage recordings. **a,** Frames showing basal fluorescence levels (F_0_) and mean fluorescence changes (-ΔF) during spike back-propagation (frame rate 1 kHz). A rectangular ROI (rectangles) was unidirectionally shifted in 5 µm increments, up to 25 µm from the center of a branch. **b,** Quantitative analysis for the mean fluorescence changes of LR-Voltron2 as a function of ROI distance from the center of a branch. Data were normalized to the responses observed at 0 µm (median and interquantile range; n= 49 branches, 21 cells, 20 animals). The |ΔF| signal fell to half maximum at transverse offsets of 4 μm (*n* = 49 branches, 21 cells, median). At offsets > 20 μm, the fluorescence signal plateaued at 11% (median, IQR: 7 - 24%) of maximum, reflecting the contribution of out-of-focus dendrites. Related to Fig. 1.

**Fig. S6.**
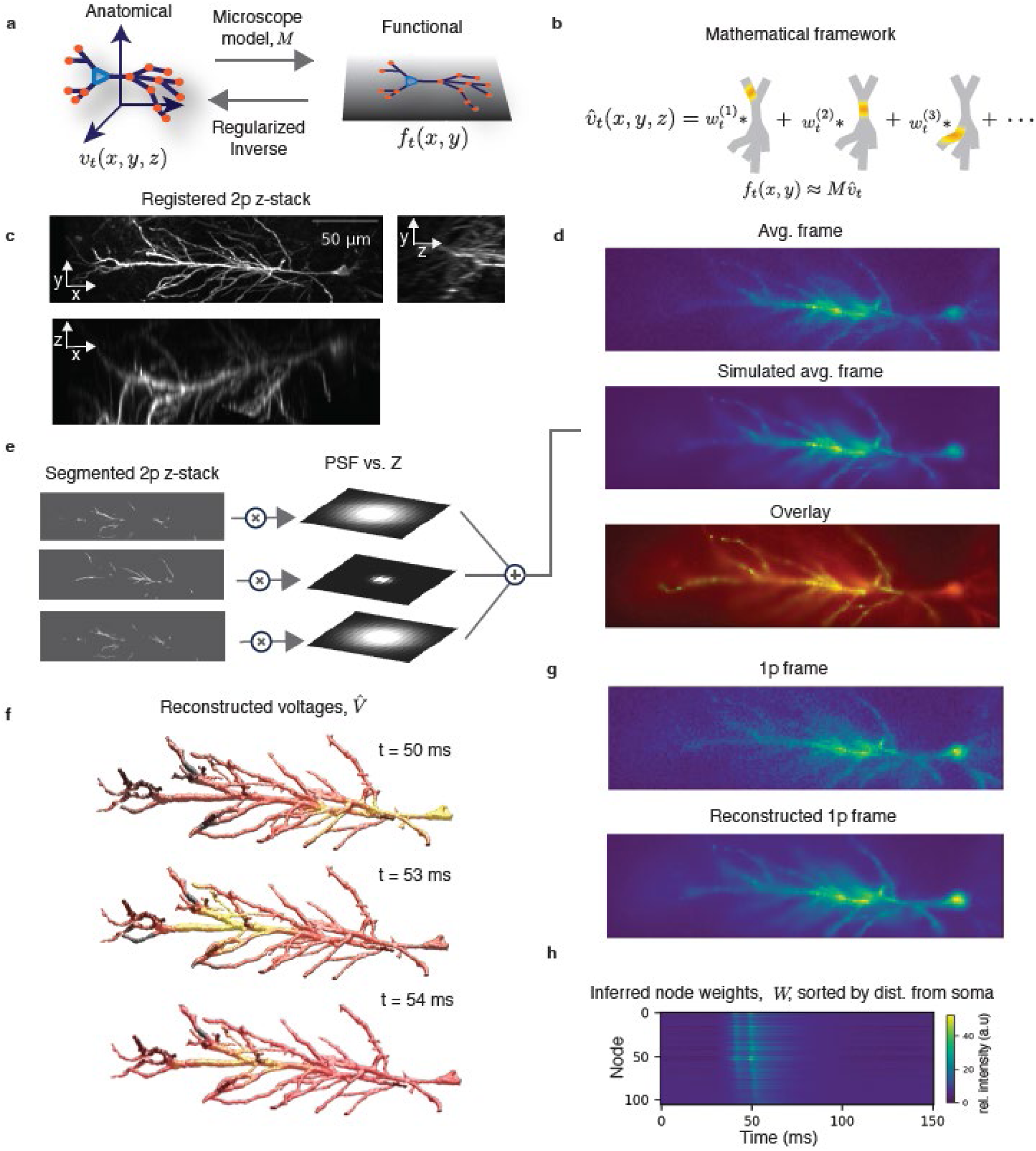
Fusing functional and structural data. **a**, The voltage inference problem is cast as an inverse problem. Projecting from 3D to 2D is modeled as blurring and projection, the parameters of which are determined by a microscope model which includes the microscope point spread function (PSF) and depth-dependent signal attenuation. The estimated three-dimensional voltage profile, *v^*_*t*_(*x*, *y*, *z*), is inferred from a two-dimensional fluorescence image *f*_*t*_(*x*, *y*) by solving a regularized inverse problem which incorporates biophysical constraints (continuity, smoothness) on the voltage. **b**, The voltage estimate, *v^*_*t*_(*x*, *y*, *z*), is reconstructed from a weighted sum of smooth basis functions (colored patches) which enforce continuity and smoothness. The estimated voltage is passed through the microscope model (given by the matrix *M*) to create an estimated movie frame. **c**, Three projections of a 2P z-stack of a CA1 pyramidal cell. **d**, Top: average 1P fluorescence frame in the functional recording. Middle: simulated average frame computed by applying the microscope model to the 2P z-stack. Bottom: overlay. **e**, The 2P z-stack was segmented into neuron and background, and aligned to the 1p coordinate frame via a polynomial transform. The basal dendrites were not included in the segmentation. To create a simulated 1p image, each plane of the segmented 2p stack was blurred with a gaussian filter whose width determined by the parameters of the PSF. The blurred planes were weighted and summed to create a simulated 1p frame (**d,** middle panel). **f**, Three snapshots of a spike-triggered-average voltage estimate during a bAP with a dSpike. **g**, Top: single frame from the STA movie. Bottom: Model estimated frame. **h**, Inferred voltages at points along the dendritic tree, sorted by increasing distance from the soma. Spike doublets are clearly visible, with the second spike penetrating further along the apical dendritic branch (a dSpike). See also Movie 2.

**Fig. S7.**
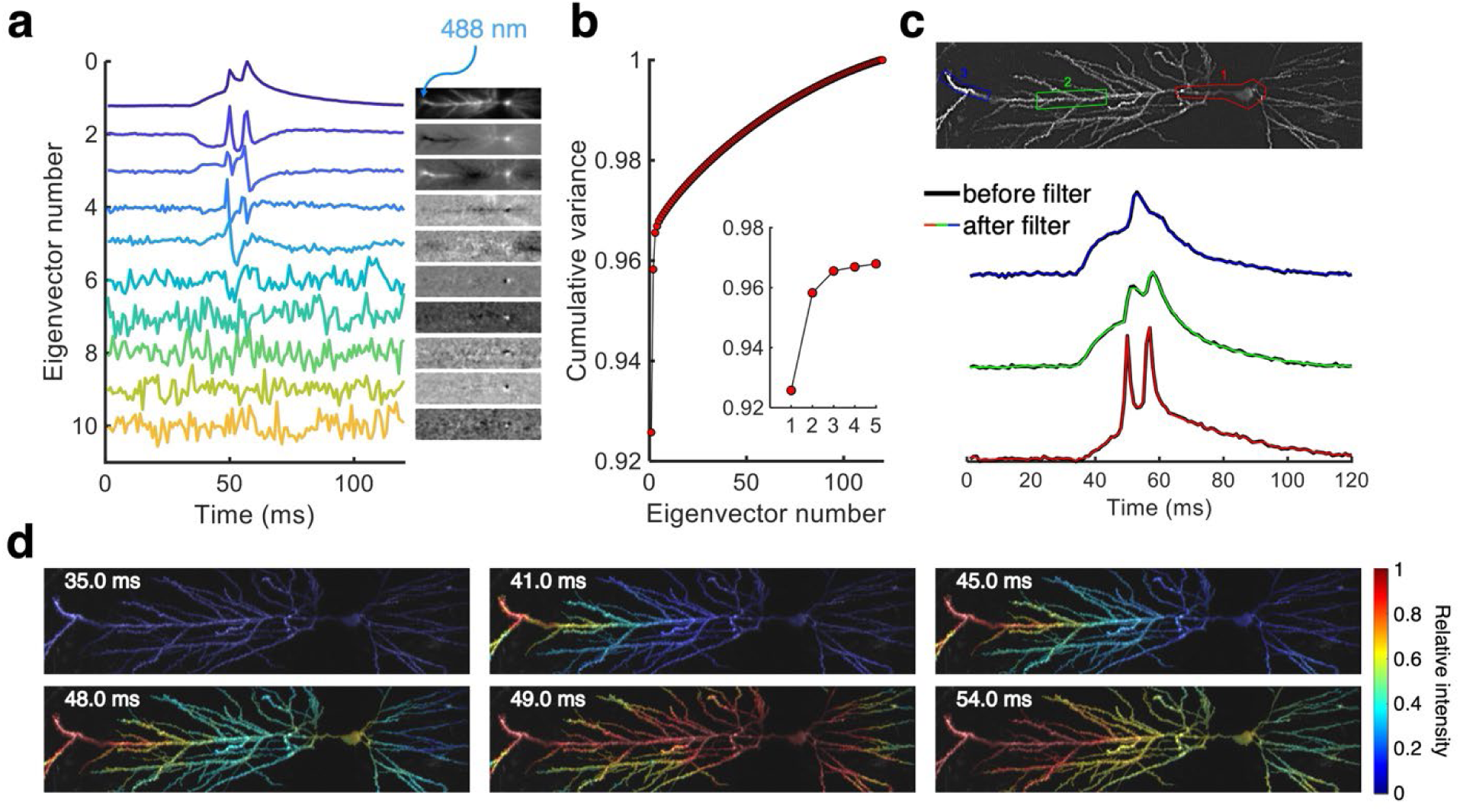
Principal component analysis (PCA)-based data smoothing. **a,** Patterned optogenetic stimulation at distal dendrite for 20 ms (blue arrow), and a stimulus-triggered average fluorescence movie (frame rate, 1 kHz, 11 trials). PCA was applied to the waveforms at individual pixels. The inset shows projection of the movie onto each of the first ten eigenvectors. **b,** The first 5 PCA eigenvectors account for > 96% of the pixel-to-pixel waveform variance, with the remaining eigenvectors largely corresponding to shot noise. The inset highlights the first 5 PCA eigenvectors. **c,** Comparison of the waveforms before and after projecting the movie into the space spanned by the first 5 principal components. Original waveforms are shown in black lines, and the PCA-filtered data in each corresponding region are shown in color. **d,** Frames after the PCA filtering. Data from Fig. 3a. Related to Fig. 1 (see also Movie 1).

**Fig. S8.**
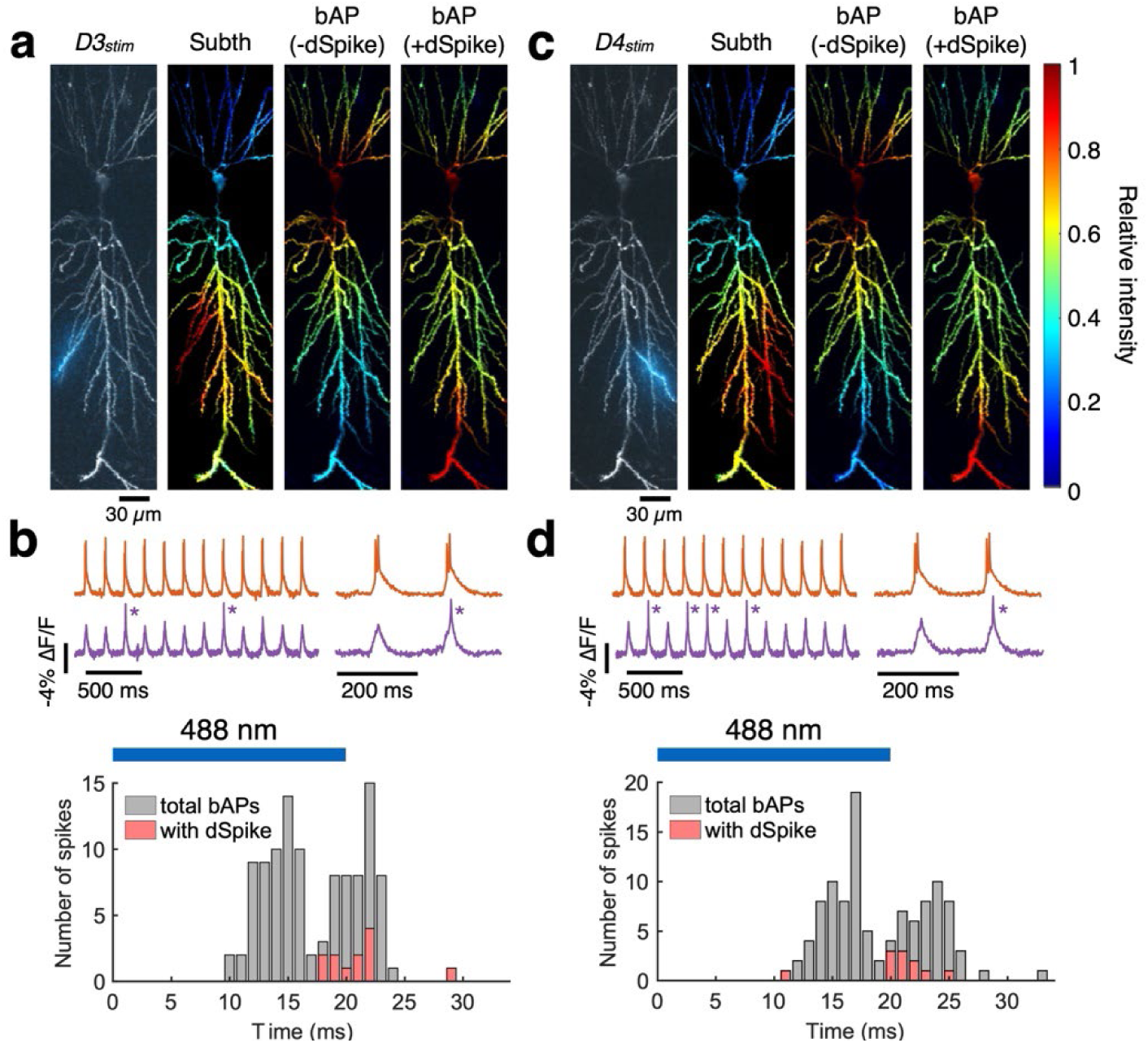
Additional examples of responses to dendrite-targeted stimuli. **a,** 2P structural image (gray) overlayed with fluorescence of eYFP (blue) showing the optogenetic stimulus (20 ms duration, 5 Hz). Normalized amplitude (ΔF/F_ref_) maps for subthreshold depolarization (stimulus-triggered average from 54 trials), back-propagating action potentials (bAPs) without dSpikes (-dSpikes; spike-triggered average from 80 spikes) and with dSpikes (+dSpikes; spike-triggered average from 12 spikes). **b,** Top: example traces at the soma (orange) and distal dendrites (> 300 µm; purple) showing cases with dSpike (indicated by asterisk). Bottom: counting all bAPs, and bAPs with dSpikes, as a function of time following optogenetic stimulus onset. Blue bar shows the timing of 488 nm stimulus (20 ms). **c-d,** Equivalent plots by stimulating a different dendrite (D4_stim_; 54 trials, 85 -dSpikes, and 11 +dSpikes). Related to Fig. 3.

**Fig. S9.**
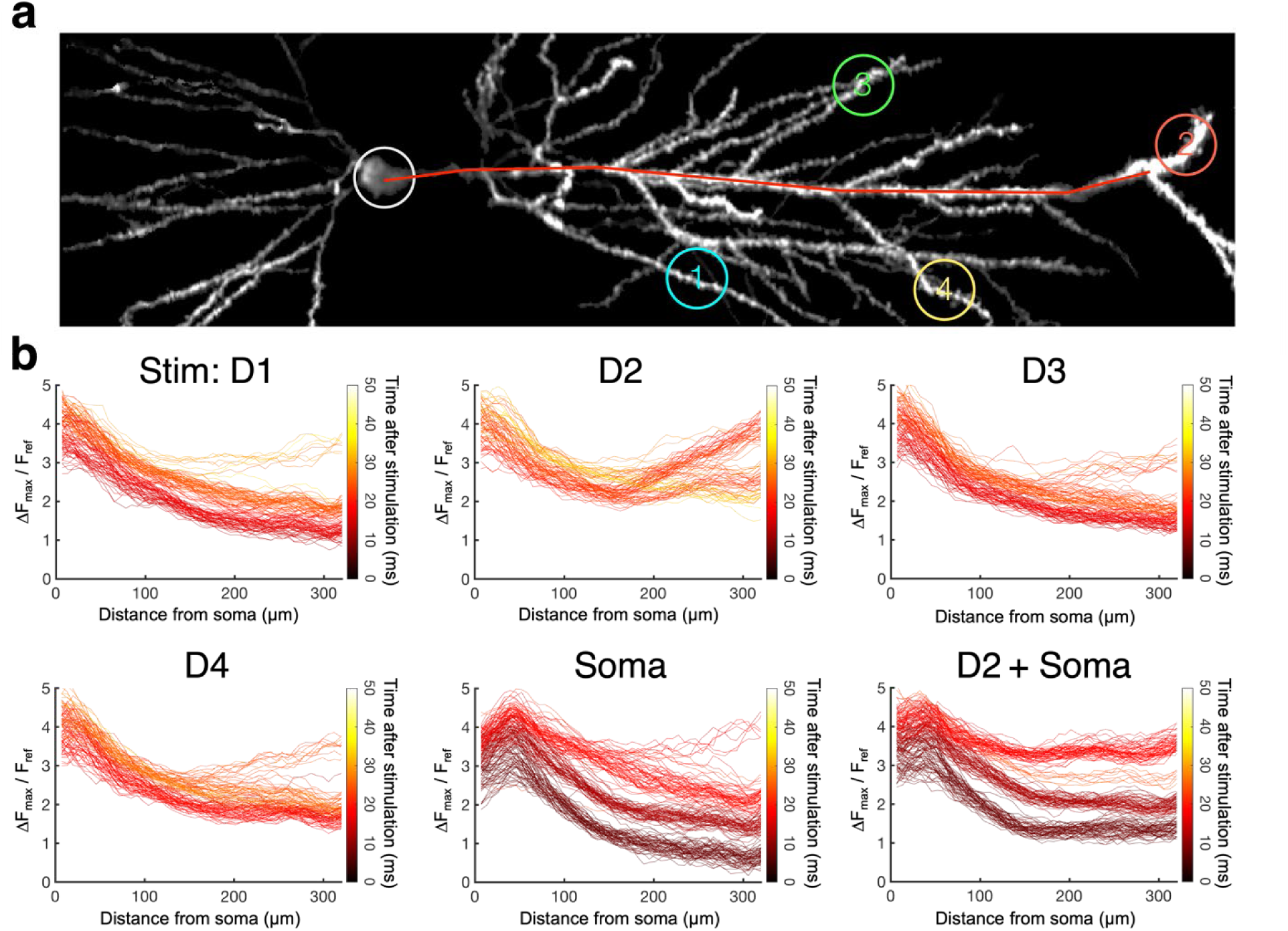
Spatial profiles for individual bAPs. **a,** 2P structural image (gray) showing stimulated branches indicated by numbers (stimuli: 20 ms duration at 5 Hz). **b,** Amplitude profiles for each bAP along the red line in **(a)**. Events that rise in the distal region indicate dSpikes. Plots were color-coded by the time after stimulus onset. Related to Fig. 2, Fig. 3, and Fig. S8.

**Fig. S10.**
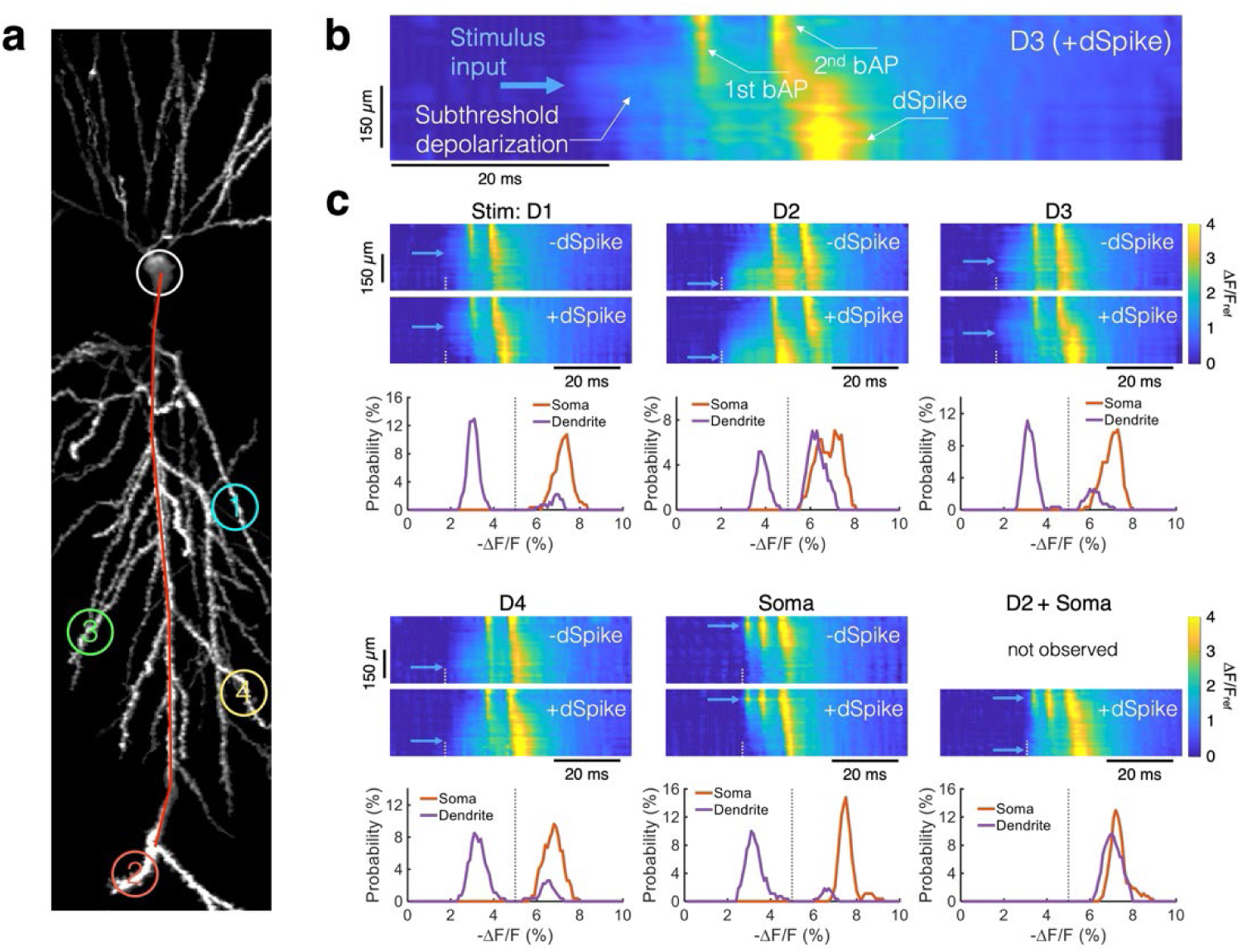
Quantifying propagation and amplitudes of bAPs and dSpikes. **a,** Data from Fig. S9. 2P structural image (gray) showing the neuron branches targeted for stimulation (20 ms duration at 5 Hz). **b,** Annotated example kymograph along the red line in **(a)**. Data from **(c)**. Blue arrowhead indicates place and time of stimulus onset. Subthreshold depolarization originated at the junction between the stimulated branch and the apical shaft and propagated diffusively in both the distal and proximal directions. The first bAP originated in the soma and back-propagated part way down the shaft, but then failed. The next bAP also originated in the soma and back-propagated until it triggered a dSpike. **c,** Top: kymographs comparing single-trial instances of bAPs with (+dSpike) and without (-dSpike) dendritic spikes. Blue arrowhead indicates place and time of stimulus onset. White dotted line indicates time of stimulus onset. Bottom: distribution of bAP amplitudes at the soma (orange) and at a distal dendrite (> 300 µm; purple). The amplitude in the soma had a unimodal distribution and the amplitude in the dendrites had a clearly bimodal distribution. Black dotted lines indicate threshold between -dSpike vs. +dSpike. Related to Fig. 2, Fig. 3, and Fig. S8.

**Fig. S11.**
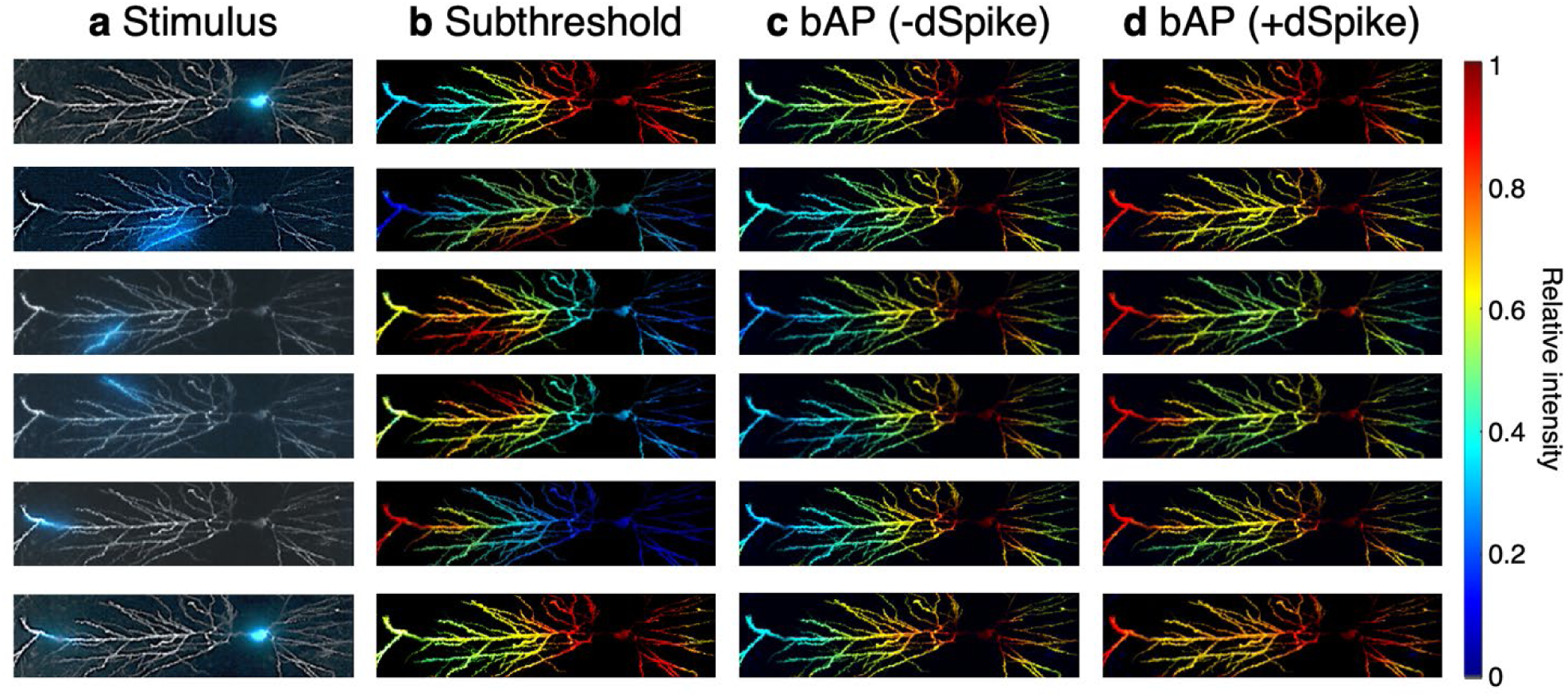
Comparison of spatial maps for different stimulus locations. **a,** 2P structural images (gray) overlayed with fluorescence of eYFP (blue) showing the optogenetic stimulus (20 ms duration at 5 Hz). Normalized amplitude (ΔF/F_ref_) maps for **b,** subthreshold depolarization, **c,** back-propagating action potentials (bAPs) without dSpikes (-dSpikes) and **d,** bAPs with dSpikes (+dSpikes). Data from Figs. 2, 3, and S8, arranged for side-by-side comparison.

**Fig. S12.**
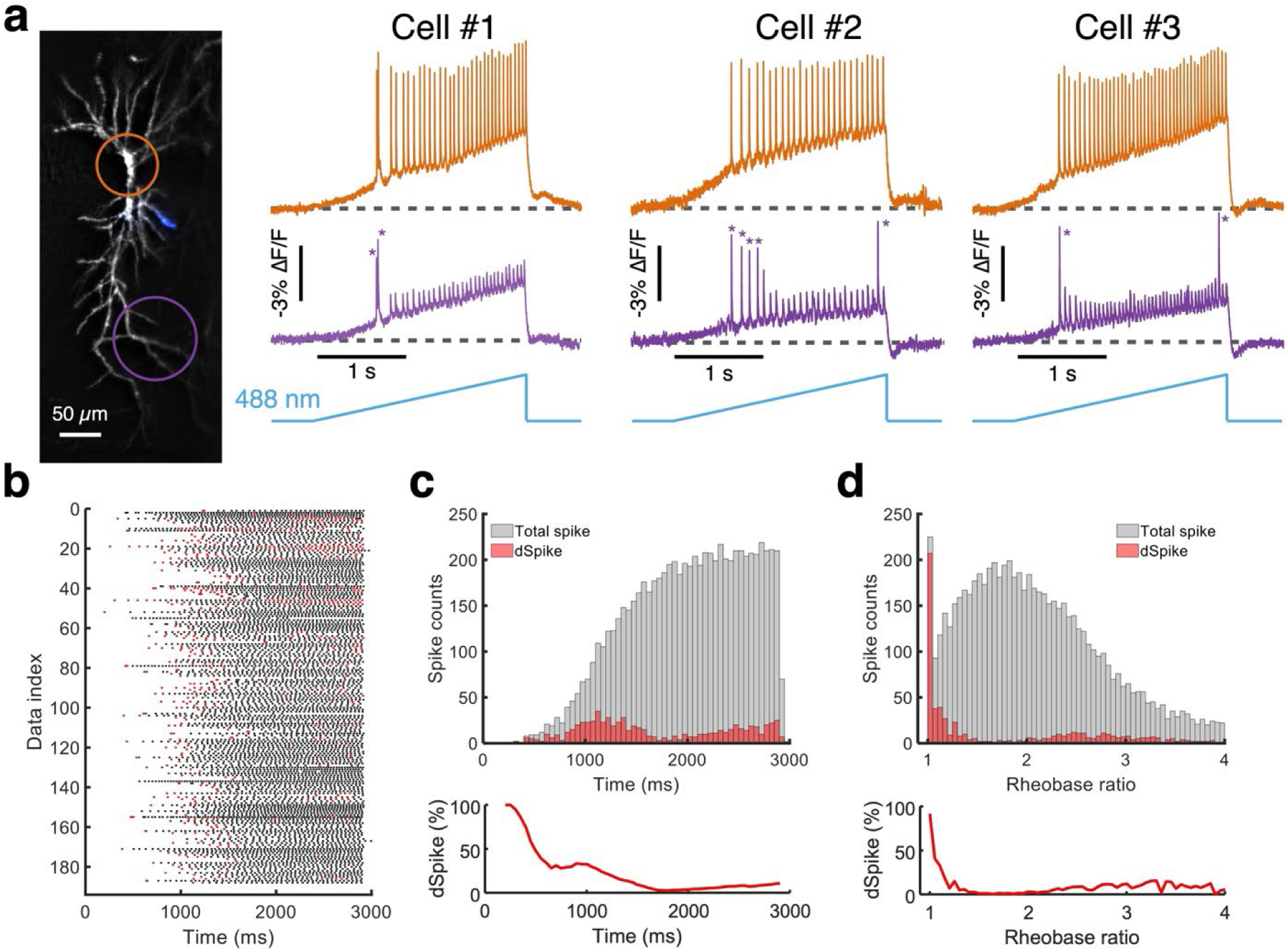
Production of dSpikes by optogenetic ramp stimulation. **a,** Left: HiLo structural image (gray) overlayed with fluorescence of eYFP (blue) showing the optogenetic stimulus (3 s linear ramp). Right: example traces from three cells at the soma (orange) and distal dendrites (> 300 µm; purple) showing cases with dSpike (indicated by asterisk). **b,** Spike raster plot showing the timing of all bAPs (black), and bAPs with dSpikes (red) following the optogenetic stimulation onset (n = 188 recordings, 56 cells, 44 animals). Optogenetic stimuli were targeted to the soma or proximal dendrites (< 150 µm from the soma). **c,** Top: Number of bAPs (gray) and bAPs with dSpikes (red), as a function of time after stimulus onset. Bottom: Percent of dSpikes among all bAPs. **d,** Same as **(c)**, with the time-axis scaled by time to reach optical rheobase (i.e. to induce the first spike). This scaling controls for variations in channelrhodopsin expression level and optogenetic stimulus strength. Under gradual ramp stimulation, the first bAP almost always evoked a dSpike (92% out of 225 bAPs). Related to Fig. 4.

**Fig. S13.**
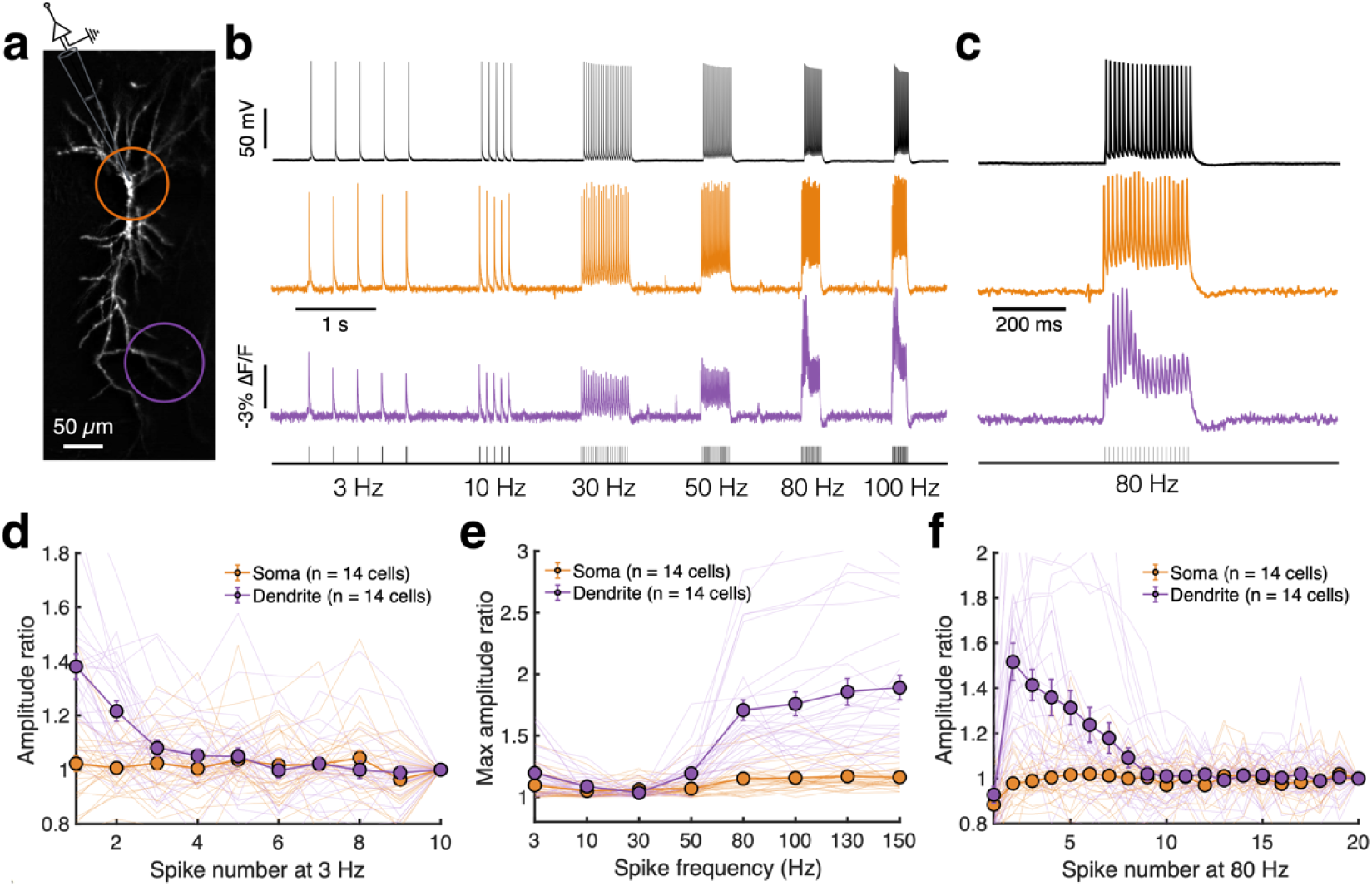
bAPs triggered by current injection also show a window for evoking dSpikes. **a,** HiLo image (gray) of a neuron stimulated via patch pipette at the soma. **b,** Recording of voltage at the soma via current clamp (black, 100 kHz sample rate) and fluorescence (orange, 1 kHz frame rate), and at a distal dendrite via fluorescence (> 300 µm; purple). Spikes were evoked by 2 nA current injection for 2 ms at frequencies ranging from 3 Hz to 100 Hz. **c,** Magnified view of responses at 80 Hz stimulus. **d,** Fluorescence amplitude (-ΔF/F) ratio relative to the last spike at 3 Hz (n = 29 recordings from 14 cells across 11 animals). The mean ± SEM and individual recordings (thin lines) are overlaid, showing responses at the soma (orange) and a distal dendrite (purple). The voltage at the distal dendrites gradually decreased, consistent with prior measurements^10^ and likely due to slow Na_V_ inactivation. **e,** Maximum spike amplitudes at each frequency, normalized to the minimum amplitude for each dataset. Somatic stimulation at 80 Hz and higher evoked dSpikes. **f,** Fluorescence amplitude of successive spikes, relative to the last spike (80 Hz stimulus). The dSpikes arose in a window after stimulus onset. Related to Fig. 4.

**Fig. S14.**
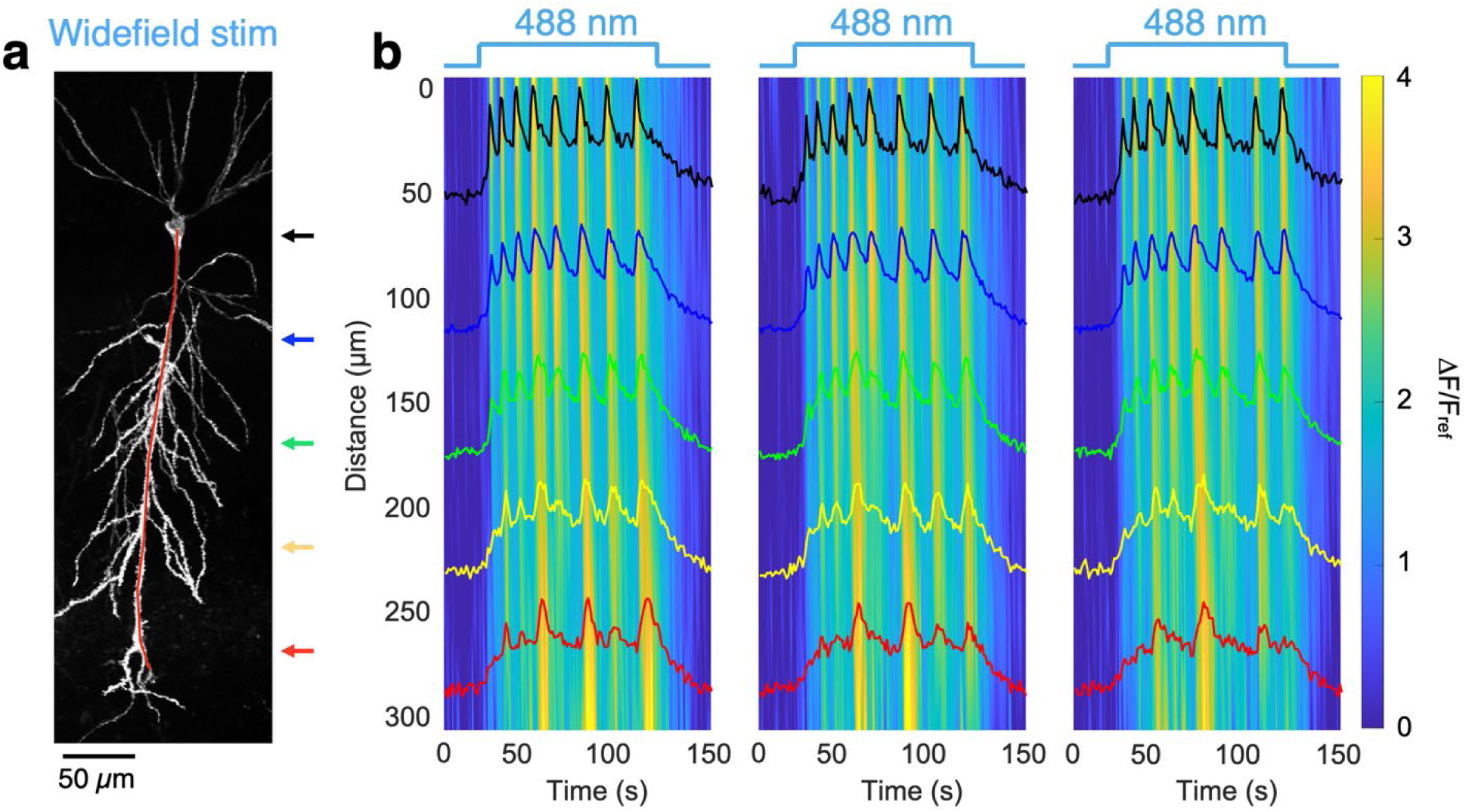
Stochastic back-propagation under strong drive. **a,** 2P structural image showing the measured neuron. The cell was exposed to wide-field optogenetic stimulation (100 ms duration at 5 Hz). Red line indicates the contour used to calculate kymographs. **b,** Successive responses to pulses of blue light. The spike train in the soma was regular, but the pattern of bAP propagation showed trial-to-trial variability, indicative of chaotic back-propagation. Related to Fig. 4.

**Fig. S15.**
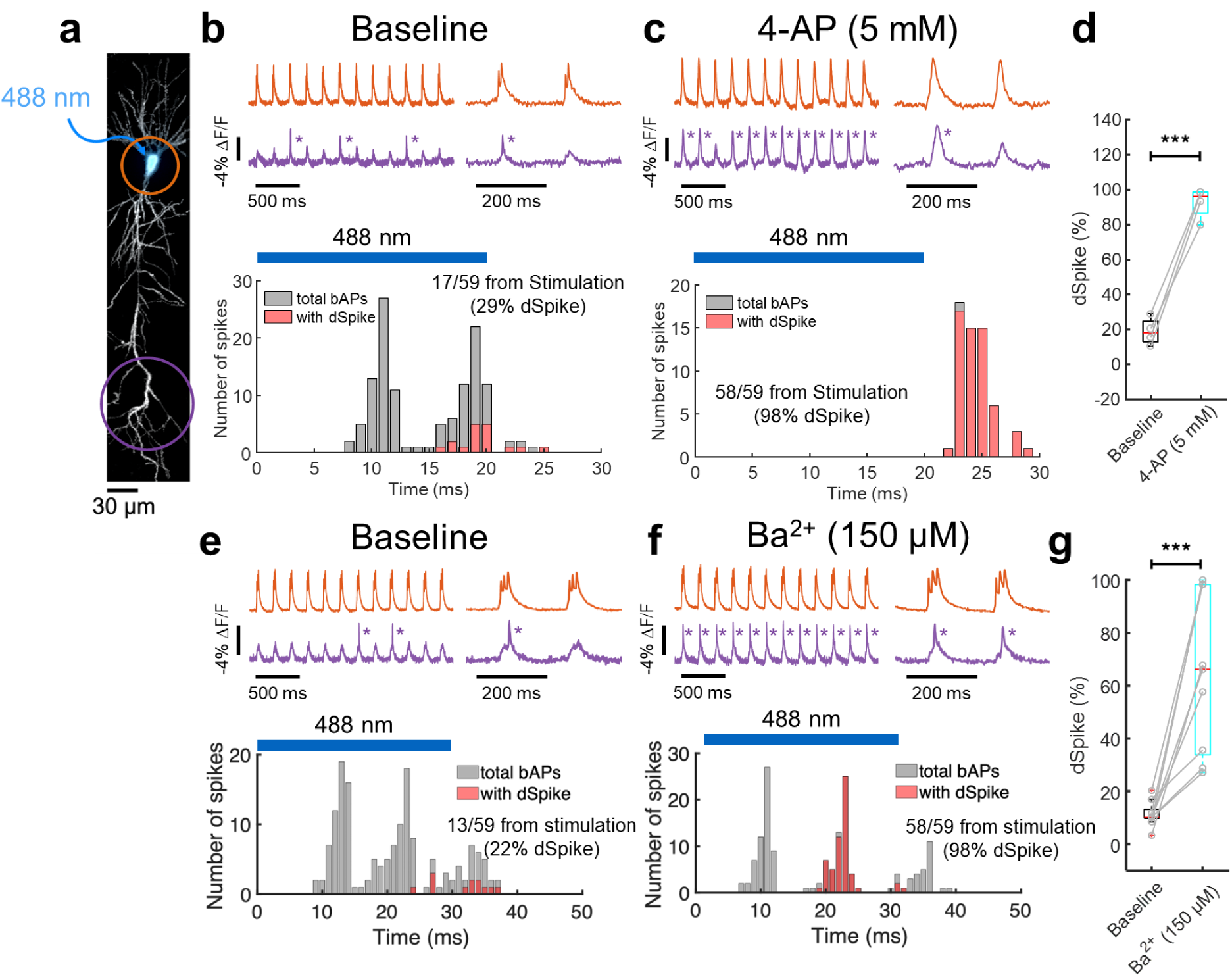
dSpike sensitivity to potassium channel blockers, 4-AP and BaCl_2_. **a,** 2P structural image (gray) overlayed with fluorescence of eYFP (blue) showing the optogenetic stimulus (20 or 30 ms duration at 5 Hz). **b,** Top: example traces at the soma (orange) and distal dendrites (> 300 µm; purple). DSpikes indicated by asterisks. Bottom: counting all bAPs, and bAPs with dSpikes, as a function of time following the optogenetic stimulation onset. Blue bar shows the timing of 488 nm. **c,** Corresponding analysis for the same cell in the presence of 4-AP (5 mM) in the bath. **d,** 4-AP significantly increased the dSpike probability compared to baseline (n = 4 cells from 3 animals). **e-g,** Equivalent experiments with BaCl_2_ (150 µM) application in the bath (n = 9 cells from 6 animals). Box plots show median, 25^th^ and 75^th^ percentile, extrema. \*\*\**p* < 0.001, paired Student’s t-test. Related to Fig. 4.

**Fig. S16.**
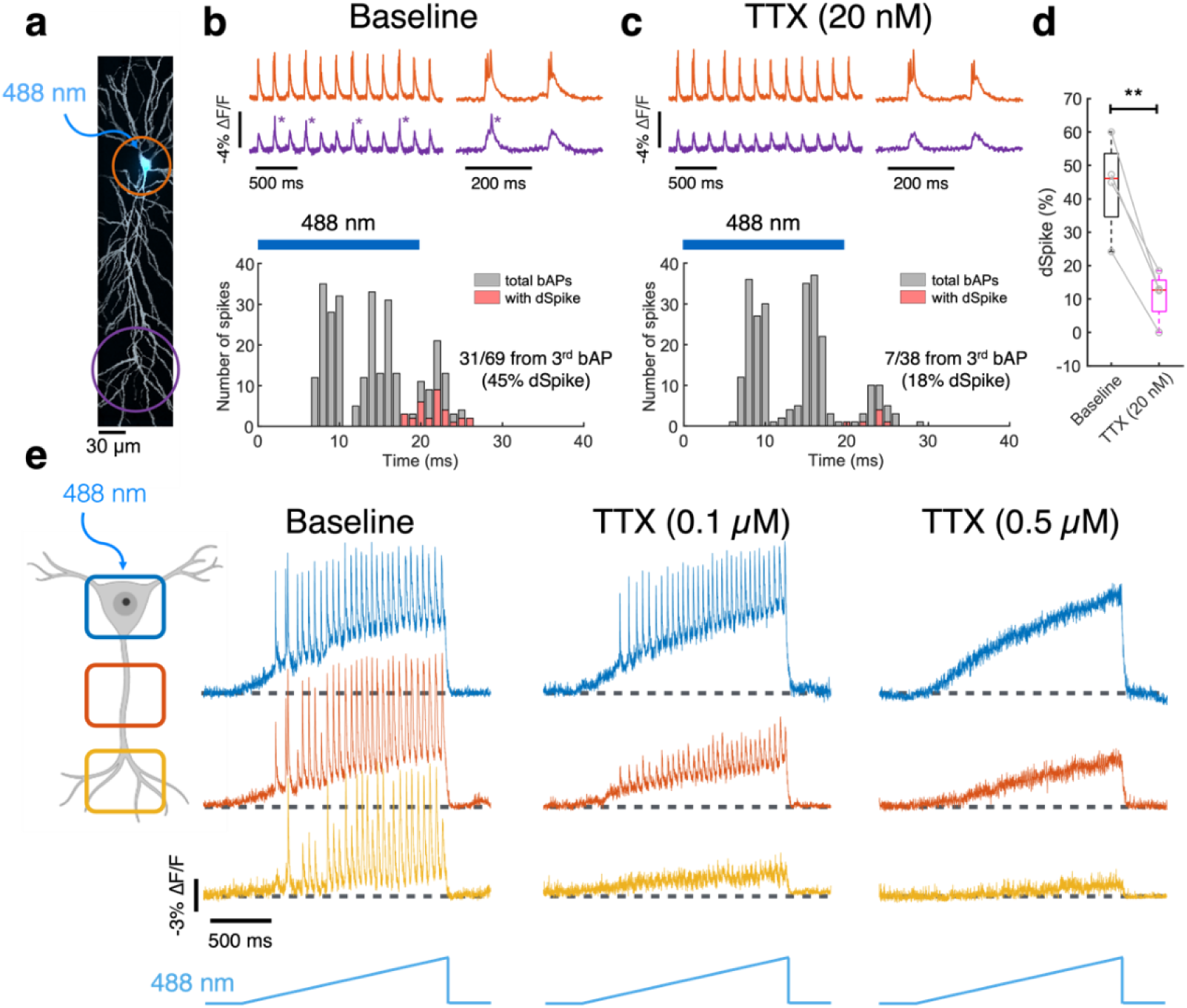
dSpike sensitivity to sodium channel blocker, TTX. **a,** 2P structural image (gray) overlayed with fluorescence of eYFP (blue) showing the optogenetic stimulus (20 ms duration at 5 Hz). **b,** Top: example traces at the soma (orange) and a distal dendrite (> 300 µm; purple). DSpikes indicated by asterisks. Bottom: counting all bAPs, and bAPs with dSpikes, as a function of time following the optogenetic stimulus onset. Blue bar shows the timing of the optogenetic stimulus (20 ms). **c,** Corresponding analysis in the presence of low concentration of TTX (20 nM). **d,** TTX significantly decreased the percentage of dSpikes vs. baseline (n = 4 cells from 3 animals). **e,** Example neuron showing responses at the soma (blue), proximal (orange; < 200 µm from soma), and distal (yellow; > 300 µm) dendrites. The soma was optogenetically stimulated with a linear ramp (2 s duration). The neuron was sequentially tested at 0.1 µM and 0.5 µM TTX in the bath using the same soma-targeted stimulus waveforms. At 0.1 µM, the soma continued to spike but dSpikes were suppressed. At 0.5 µM, spiking at the soma was suppressed. This cell’s soma, positioned deep in the slice (> 100 µm from the surface), showed reduced TTX sensitivity (typically soma spiking is suppressed by 0.1 µM of TTX). Box plots show median, 25^th^ and 75^th^ percentile, extrema. \*\**p* < 0.01, paired Student’s t-test. Related to Fig. 4.

**Fig. S17.**
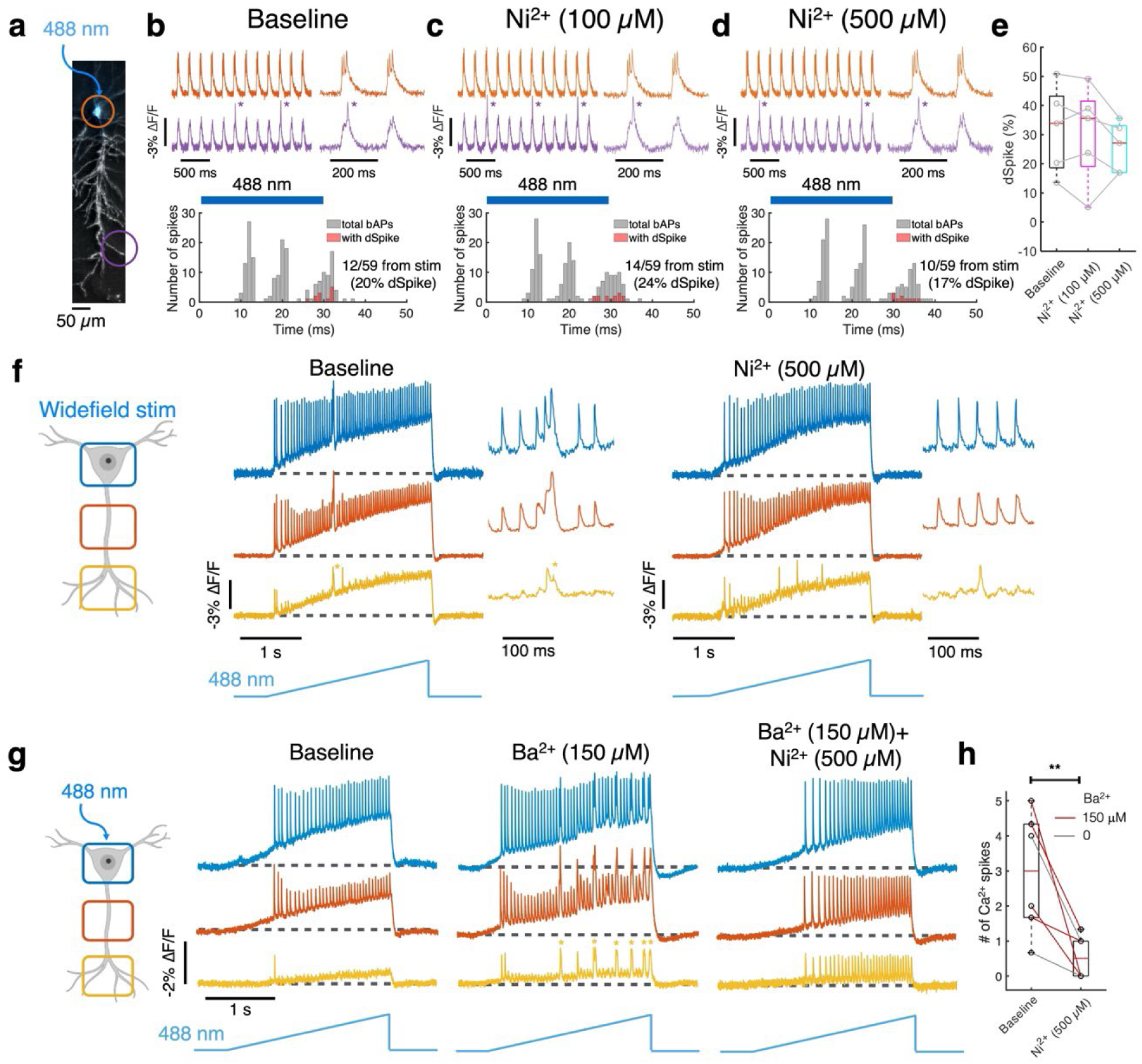
dSpike sensitivity to calcium channel blocker, Ni^2+^. **a,** HiLo image (gray) with soma-targeted optogenetic stimulation (blue; 30 ms duration at 5 Hz). **b,** Top: example traces at the soma (orange) and a distal dendrite (> 300 µm; purple). DSpikes indicated by asterisks. Bottom: counting all bAPs, and bAPs with dSpikes, as a function of time following the optogenetic stimulus onset. Blue bar shows the stimulus timing. **c-d,** Corresponding analysis with different concentrations of Ni^2+^ in the bath. **e,** Ni^2+^ did not affect the proportion of dSpikes vs. baseline (n = 5 cells from 4 animals). **f,** Example neuron showing responses at the soma (blue), proximal (orange; < 200 µm from soma), and distal (yellow; > 300 µm) dendrites. Widefield optogenetic stimulation with a linear ramp (3 s duration), was applied without, then with, 500 µM Ni^2+^. Putative dendritic Ca^2+^ spikes (depolarizations lasting > 20 ms) marked by asterisks. Ni^2+^ suppressed dendritic Ca^2+^ spikes but not short-duration dendritic Na^+^ spikes. **g,** Ba^2+^, a K_V_ channel blocker, increased the incidence of putative dendritic Ca^2+^ spikes under soma-targeted ramp stimulation. Ni^2+^ eliminated these Ca^2+^ spikes while preserving dendritic Na^+^ spikes. **h,** Ni^2+^ significantly decreased the frequency of putative Ca^2+^ spikes vs. baseline (the number of observations during a ramp stimulation; mean of three trials). These data are pooled from conditions **(f)** and **(g)** (n = 6 cells from 6 animals). Box plots show median, 25^th^ and 75^th^ percentile, extrema. \*\**p* < 0.01, paired Student’s t-test. Related to Fig. 4.

**Fig. S18.**
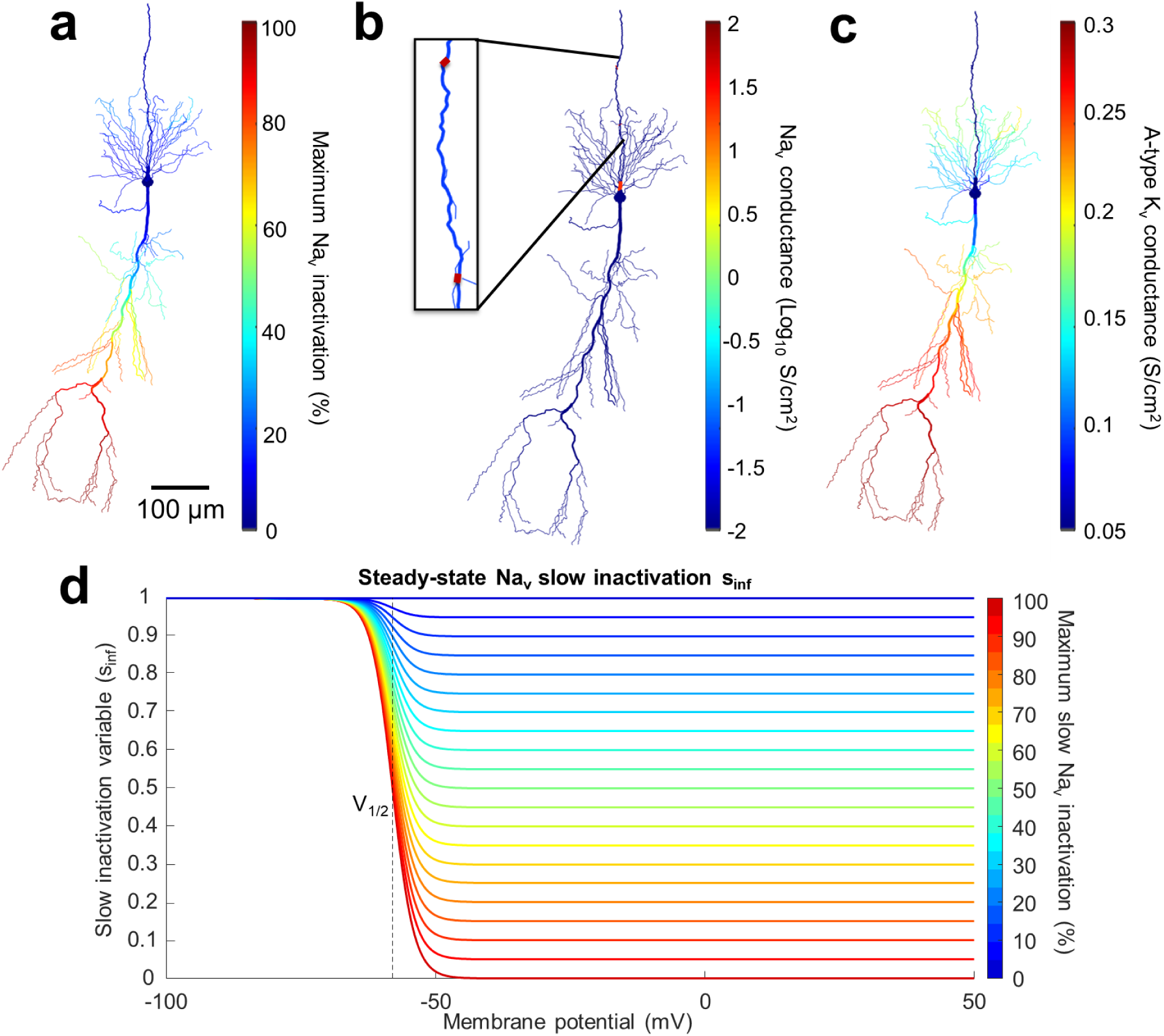
Parameters for Na_V_ and A-type K_V_ channels in CA1 model. **a,** Maximum Na_V_ inactivation (%) across the neuronal compartments. **b,** Available Na_V_ conductances distributed across the neuron (g_Na_). Note high density in node of Ranvier (30 S/cm^2^) and axon initial segment (15 S/cm^2^). **c,** Available A-type K_V_ conductances distributed across the neuron (g_KA_). **d,** Voltage-dependent changes of slow Na_V_ inactivation variables. Related to Fig. 4.

**Fig. S19.**
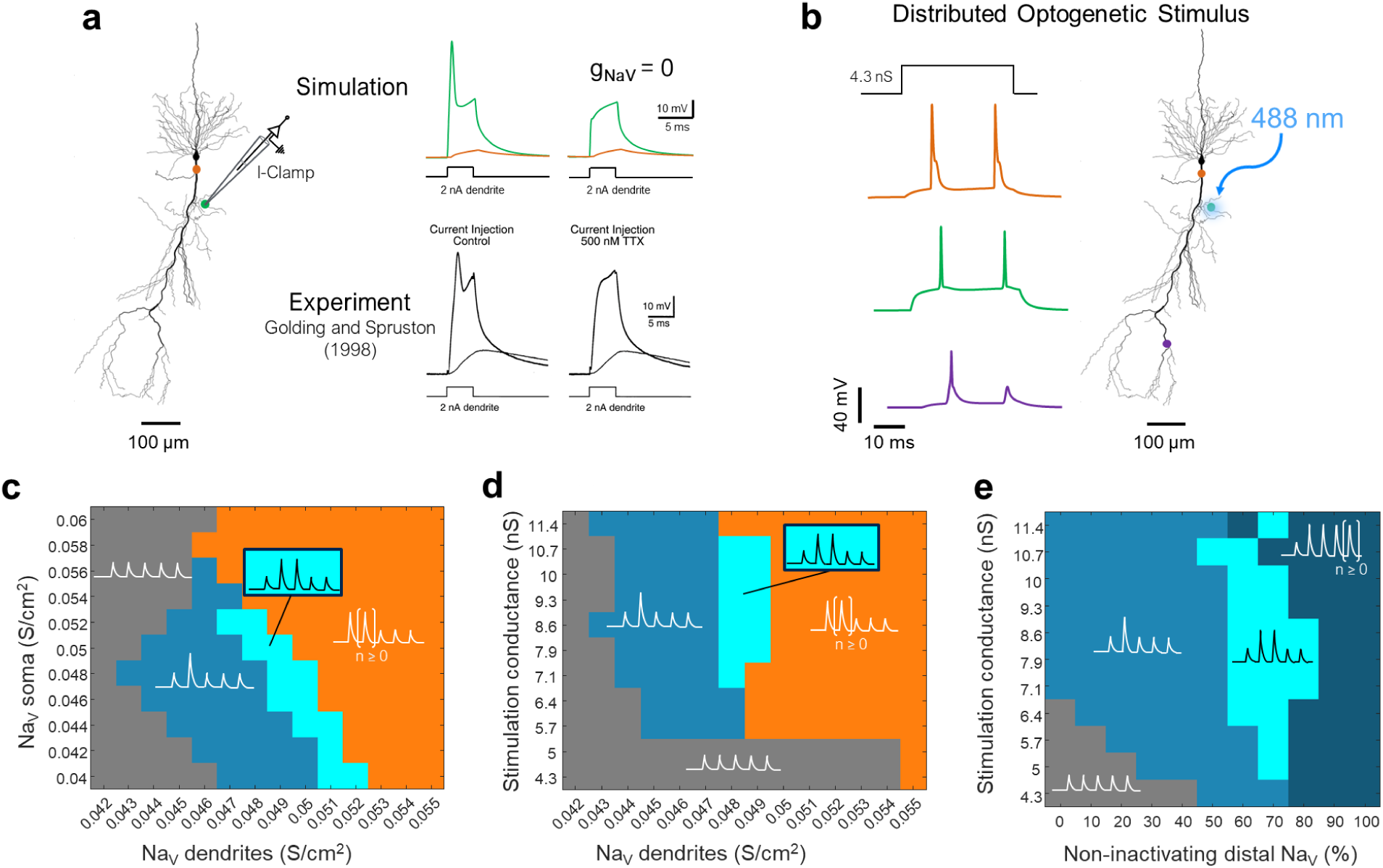
CA1 model validation and exploration of parameter space. **a,** Simulation (this work) and experimental data from Ref. ^38^ for patch clamp current injection (2 nA for 5 ms) into an oblique dendrite of a CA1 pyramidal neuron. In the simulation, voltage traces in the stimulated dendrite (green) and the soma (orange) are shown with (left) vs. without (right) dendritic Na_V_ channels. In the experiment, equivalent conditions are shown for the baseline (left) vs. the application of TTX (right). In both simulation and experiment, current injection evoked dendrite-localized sodium spikes. **b,** Simulated optogenetic stimulation over a 50 μm segment of the same dendritic branch and with the same model parameters as in **(a)**. The optogenetic stimulation evoked a spike at the soma first, which then back-propagated into the dendritic tree. The second evoked spike was attenuated in the distal dendrites due to slow Na_V_ inactivation. **c-e,** Phase diagrams of patterns of bAP amplitudes in distal dendrites as function of simulation parameters. In all cases, an optogenetic stimulus to the soma was modeled as stepwise turning on of a conductance with reversal potential 0 mV. Sample traces indicate spiking bAP motifs in a distal dendrite. Spike threshold for classification of a dSpike was set at –40 mV at 500 μm from soma. **c,** Dendritic bAP patterns for varying dendritic and somatic Na_V_ channel densities. Stimulus intensity was kept constant (d_1/2_ = 40 μm, z = 10 μm, g_Soma_= 0.7 mS/cm^2^, g_Distal_ = 0). Orange region covers any spiking pattern where the first bAP triggered a dSpike. **d**, Dendritic bAP patterns for varying stimulus strength (d_1/2_ = 40 μm, z = 10 μm, varying stimulus conductance) and dendritic Na_V_ density. Somatic Na_V_ at 50 mS/cm^2^. Orange region combines any spiking pattern where the first bAP successfully triggers a dSpike. **e,** Dendritic bAP patterns for varying stimulus strength (d_1/2_ = 40 μm, z = 10 μm, varying stimulus conductance) and fraction of non-inactivating distal Na_V_ channels. Somatic Na_V_ at 50 mS/cm^2^, dendritic Na_V_ at 47 mS/cm^2^. Dark teal region combines all patterns with three or more dSpikes. Related to Fig. 4.

**Fig. S20.**
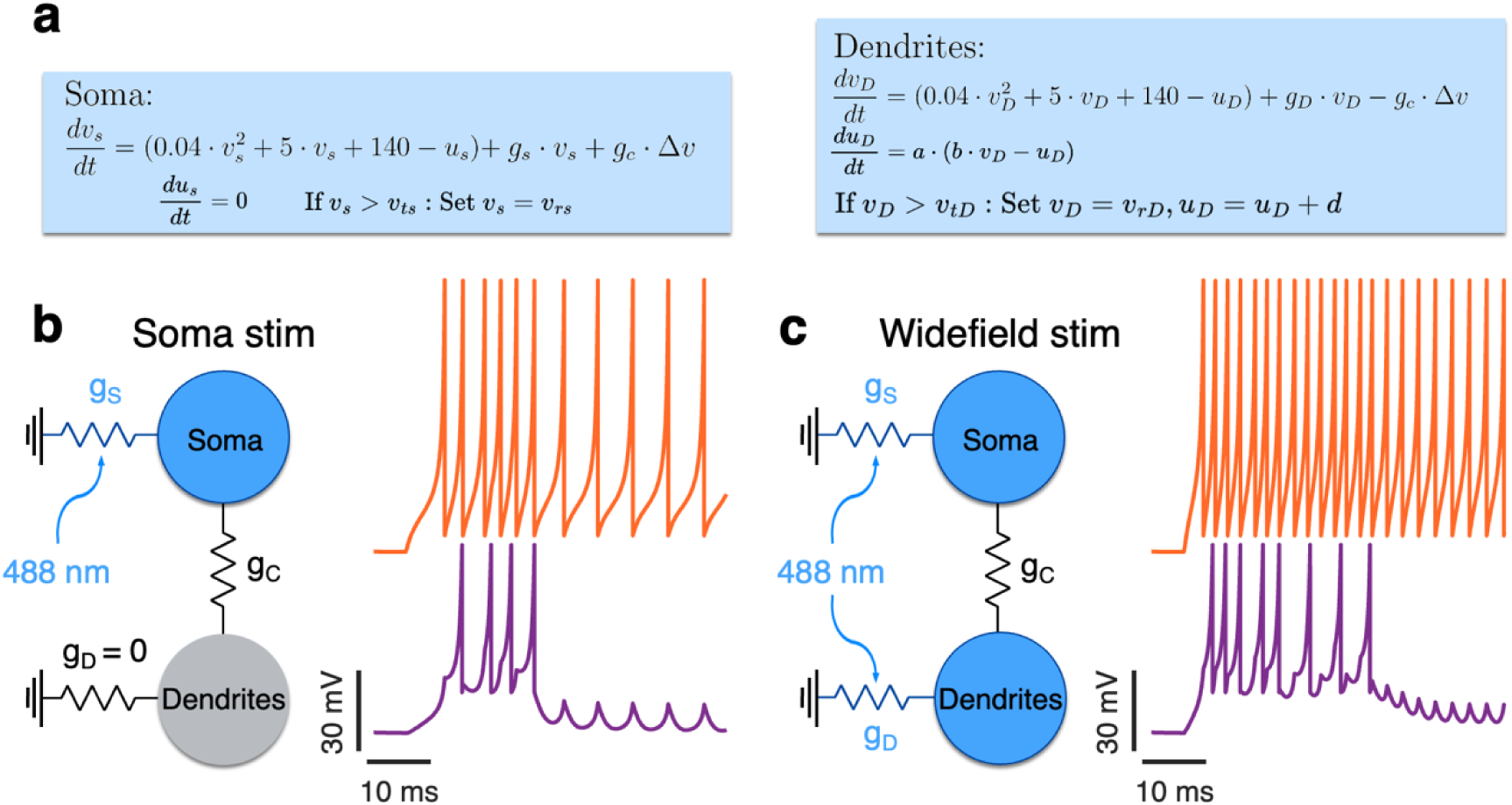
Two-compartment Izhikevich model reproduces the dynamics of dSpike successes and failures. **a,** Equations describing the two compartments. Soma without adaptation (*v_ts_* = 30, *v_rs_* = -55), dendrites with adaptation (*a* = 0.0025, *b* = 0.01, *d* = 1, *v_tD_* = 0, *v_rD_* = -55). Coupling strength *g_c_* = 0.325. **b,** Soma only stimulation with *g_s_* = 0.16. Soma *v_s_* trace in orange, dendritic *v_D_* trace in purple. **c,** Combined somatic and dendritic stimulation with *g_s_* = 0.30, *g_D_* = 0.05. Soma *v_s_* trace in orange, dendritic *v_D_* trace in purple. Related to Fig. 4.

**Fig. S21.**
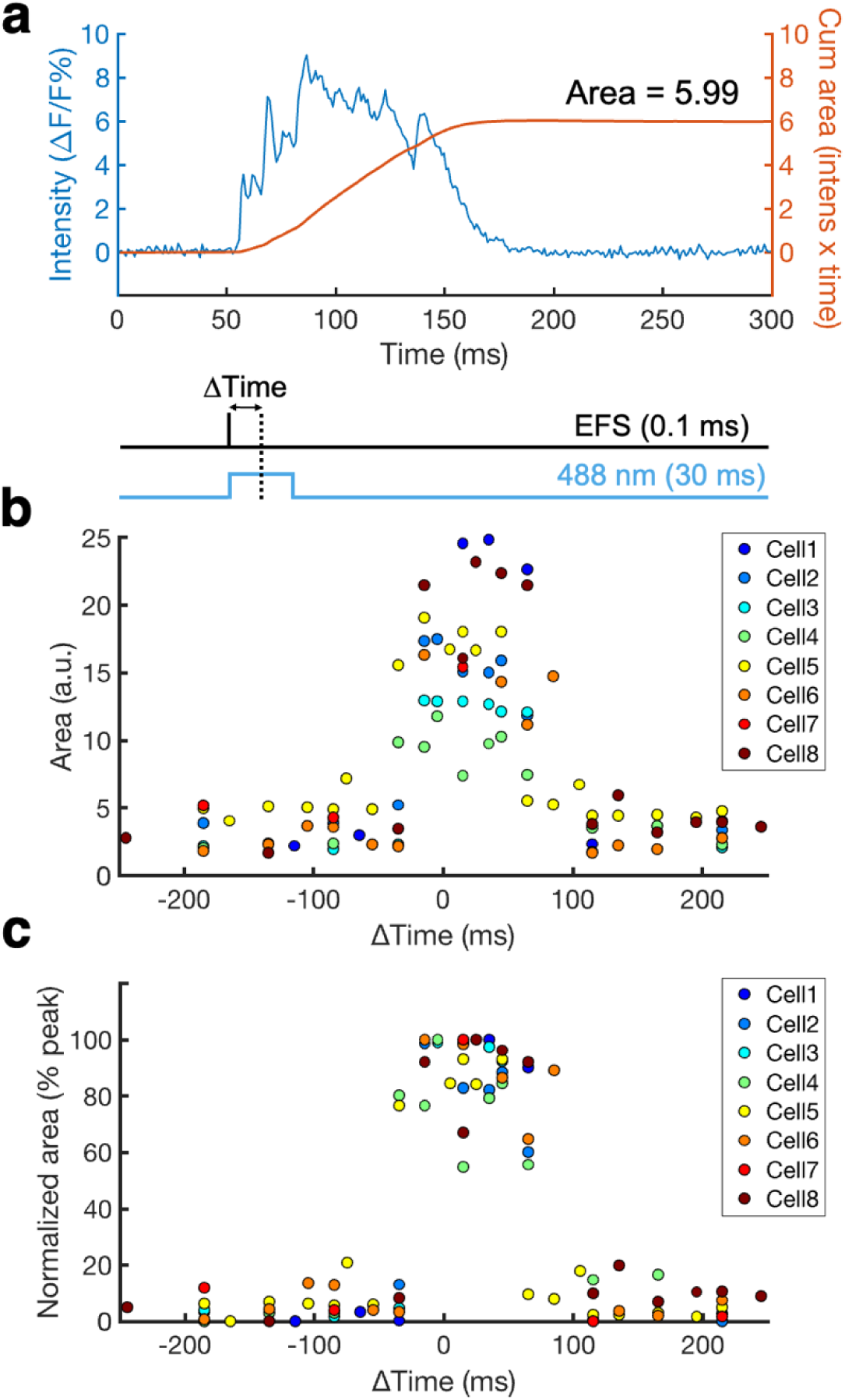
Estimating areas of plateau potentials. **a,** Example trace overlapped with cumulative area vs. time. **b,** Plateau areas vs. delay, ΔTime, between EFS (0.1 ms duration) and optogenetic stimulation (488 nm, 30 ms at the soma). Each color represents a different cell. **c,** Normalized area with each cell mapped to the range [0, 1]. Same data as in Fig. 5d.

**Fig. S22.**
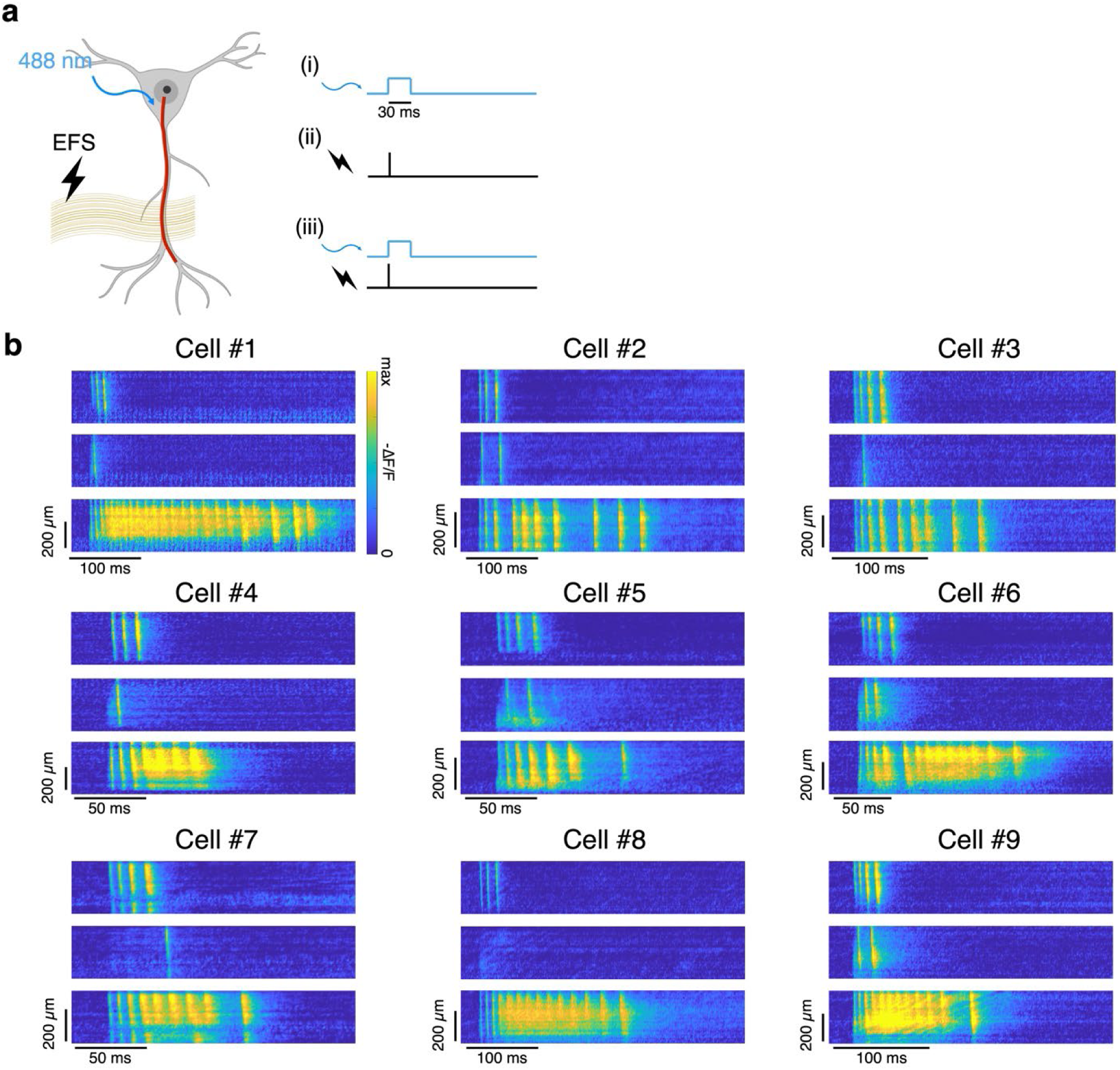
Kymographs of plateau potentials. **a,** Cells were stimulated with (i) optogenetic stimulus to the soma (30 ms), (ii) EFS targeting synapses onto distal synapses (0.1 ms), or (iii) combined optical and electrical stimuli. **b,** Gallery of response kymographs. In each kymograph, the soma is at the top and the distal apical dendrites are at the bottom. The kymographs show (top) optical, (middle) EFS, (bottom) combined stimuli.

**Fig. S23.**
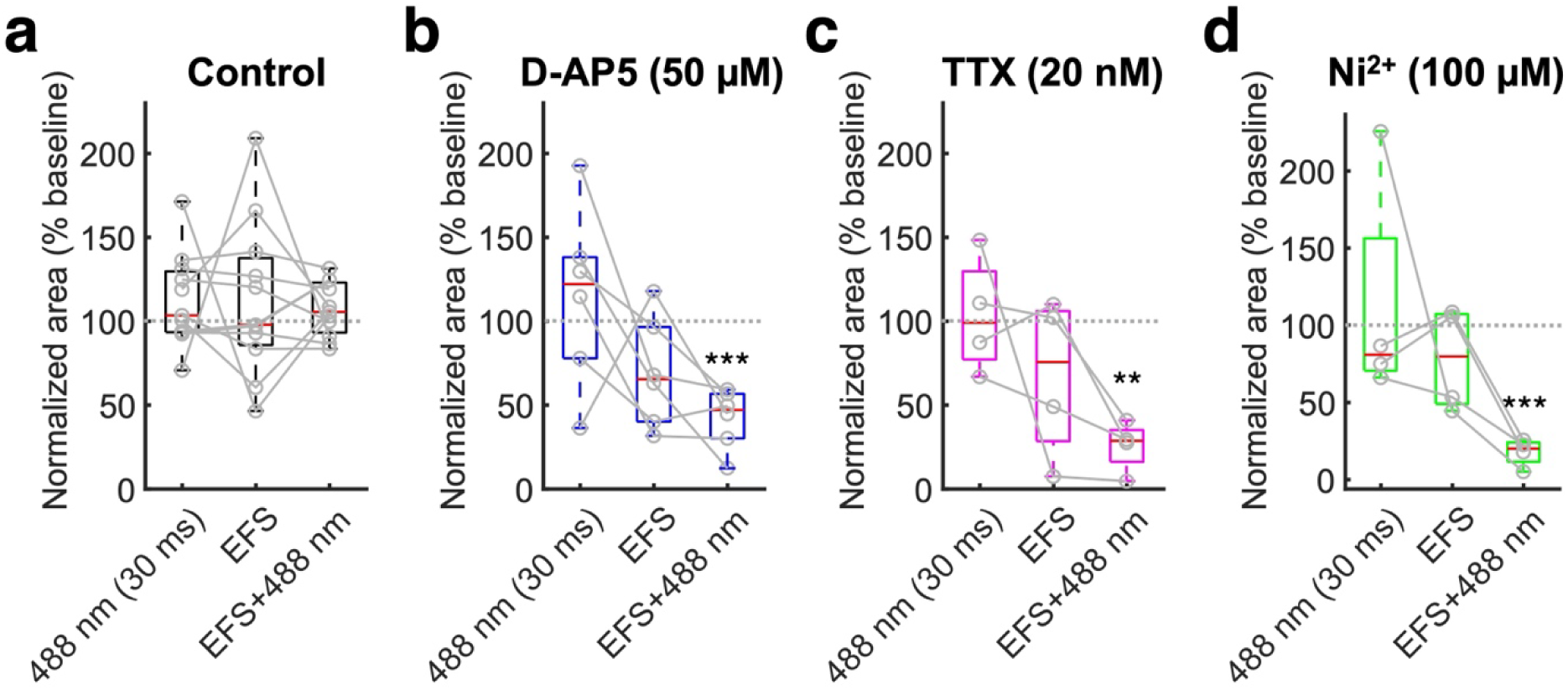
Effects of channel blockers on EFS and optogenetic signals. Effects of vehicle control (*n* = 11 cells from 8 animals), D-AP5 (NMDAR blocker, 50 µM; *n* = 6 cells from 5 animals), TTX (Na_V_ blocker, 20 nM, *n* = 4 cells from 3 animals), and Ni^2+^ (VGCC blocker, 100 µM, *n* = 4 cells from 4 animals) on responses to optogenetic stimulation alone (30 ms at the soma), electric-field stimulation alone (EFS; 0.1 ms), and combined stimulation. Optical voltage recordings were made at *d* > 200 μm from the soma. Box plots show median, 25^th^ and 75^th^ percentiles, and extrema. Each condition was compared to each baseline by paired Student’s t-test (\**p* < 0.05, \*\**p* < 0.01, and \*\*\**p* < 0.001 vs. baseline). Related to Fig. 5.

**Fig. S24.**
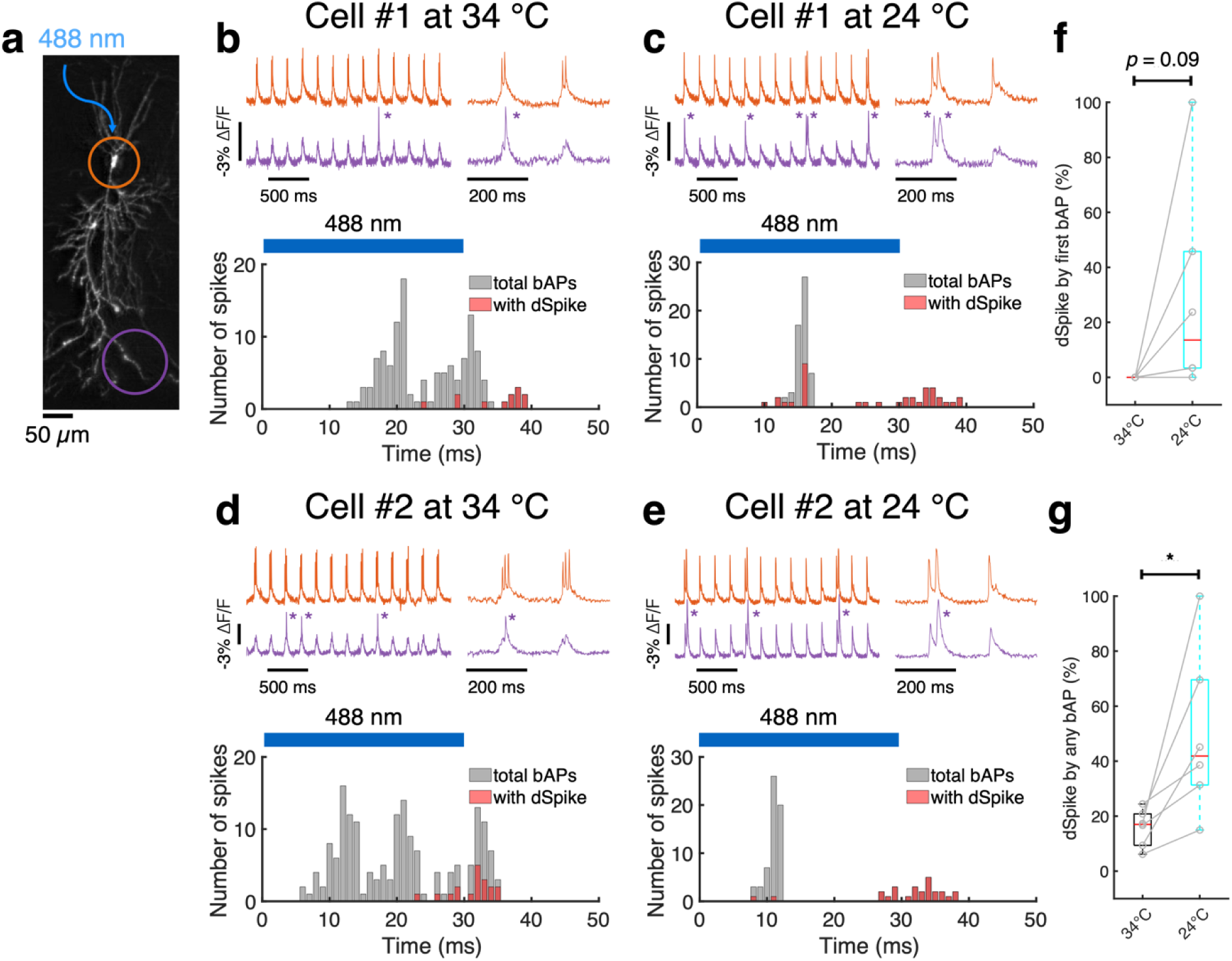
dSpike sensitivity to temperature. **a,** HiLo image (gray) with soma-targeted optogenetics stimulation (30 ms duration at 5 Hz). **b,** Top: example traces recorded at 34 °C at the soma (orange) and a distal dendrite (> 300 µm; purple). DSpikes indicated by asterisks. Bottom: counting all bAPs, and bAPs with dSpikes, as a function of time following the optogenetic stimulus onset. Blue bar shows the timing of the optogenetic stimulus (30 ms). **c,** Repeated measurement of the same cell at 24 °C. **d-e,** Same as **(b-c)**, recorded from another cell. **f,** At 24 °C, the first bAP after stimulus onset was more likely to evoke a dSpike than at 34 °C (n = 6 cells from 4 animals). **g,** Same as **(f)**, comparing across all bAPs (n = 6 cells from 4 animals). Box plots show median, 25^th^ and 75^th^ percentile, extrema. \**p* < 0.05, paired Student’s t-test.

## Notes

### Competing Interest Statement

The authors have declared no competing interest.

### Summary of Updates

Revised Introduction and Discussion. Added results mapping voltage dynamics onto 3D dendritic morphology. More detailed analysis of back-propagation.

