## Supplementary Izhikevich model for "Dendritic excitations govern back-propagation via a spike-rate accelerometer": Readme_Izhikevich.docx

\# Readme for 2-compartment Izhikevich type neuron model

\# Daniel Itkis,

\# Adam Cohen,

\# 6 June 2023, Cohen Lab (Harvard University)

\## Overview

This MATLAB script implements a 2-compartment Izhikevich neuron model to simulate backpropagating action potentials (bAPs) in a CA1 neuron stimulated by a channelrhodopsin conductance. The model contains a soma compartment and a dendrite compartment, which can be independently stimulated. The model is designed to show how different levels of conductance in the soma and dendrites impact the success-rate of dendritic bAPs driven by somatic spiking.

\## Parameters

The parameters of the Izhikevich model, such as `a`, `b`, `c`, `d`, can be modified to fit the desired conditions. The coupling between the soma and dendrites can be changed by modifying the `Coupling` variable. The variables are tuned to reproduce experimentally observed behavior.

Default Parameters & Comments:

a = 0.0025 Dendrites only (soma has no adaptation)

b = 0.01 Dendrites only (soma has no adaptation)

c = -55 Reset voltage for dendrites

cSoma = -65 Reset voltage for soma

d = 1 Dendrites only (soma has no adaptation)

vmax = 0 Spike threshold for dendrites

vmaxSoma = 30 Spike threshold for soma

rho = 1 Current asymmetry factor, can be used to model different sizes (and net conductances, capacitance) of the somatic and dendritic compartments

Additionally, the conductance of the soma and the dendrites can be altered using the `Conductance` and `ConductanceDendrites` variables, respectively.

\## Usage

To use the script, simply run it in a MATLAB environment. There are no required input arguments.

Suggested values for `Conductance` and `ConductanceDendrites`

Conductance = 0.16 and ConductanceDendrites = 0.0 for F S S S S F F F

Conductance = 0.3 and ConductanceDendrites = 0.05 for alternating success - failure motif

The output is a plot showing the u, v (membrane potential) in the soma and dendrites over time, with respect to the given conductance.

\## Modifying Input Current and Coupling

The input current to the neuron and the coupling between the soma and dendrites can be modified in the loop that simulates the Izhikevich neuron. This is achieved through the `gs` (Conductance at the soma) and `gd` (Conductance in the dendrites) variables, and the `Couple` variable (Coupling parameter between Soma and Dendrites).
