## Supplementary Izhikevich model for "Dendritic excitations govern back-propagation via a spike-rate accelerometer": Two_compartment_Izhikevich.m

```
% Two-compartment Izhikevich model of dendritic excitability in CA1
% pyramidal neurons
%
% Daniel Itkis,
% Adam Cohen,
% June 5, 2023

clear all;
close all;

%%
% Define parameters for the Izhikevich neuron model
a = 0.0025;
b = 0.01;
c = -55;
d = 1;

vmax = 0;
rho = 1;
cSoma = -65;

Conductance = 0.16; %0.16 or 0.3
ConductanceDendrites = 0.0; % 0 or 0.05

Coupling = 0.325;
nr = 1;

figure;
% Determine the input current and coupling
gs = Conductance; %Conductance at the soma
gd = ConductanceDendrites; %Conductance in the dendrites
Couple = Coupling; %Coupling parameter between Soma and Dendrites

% Initialize the membrane potential and recovery variable
v = -75;
v2 = -75;
u = -13;
u2 = -18;

% Define the time step and the number of steps to simulate
dt = 0.001;
steps = 500000;

% Initialize arrays to store the membrane potential and recovery variable
v_array = zeros(1, steps);
u_array = zeros(1, steps);
v2_array = zeros(1, steps);
u2_array = zeros(1, steps);

% Loop through the number of steps to simulate the Izhikevich neuron
for i = 1:steps
    % Update the membrane potential and recovery variable
    if i<steps/5
    g1 = 0;
    g2 = 0;
    elseif i<0.8*steps
    g1 = gs;
    g2 = gd;
    else
    g1 = 0;
    g2 = 0;
    end
    vold = v;
    v2old = v2;
    v = v + dt * (0.04 * v^2 + 5 * v + 140 - u - (g1*v) + (v2old - vold)*Couple/rho);
    u = u;
    v2 = v2 + dt * (0.04 * v2^2 + 5 * v2 + 140 - u2 - (g2*v2) - (v2old - vold)*Couple);
     u2 = u2 + dt * a * (b * v2 - u2);

    % Check for spike
    if v >= 30
        v = cSoma;
        %u = u + d;
    end

    if v2 >= vmax
        v2 = c;
        u2 = u2 + d;
    end

    % Store the membrane potential and recovery variable
    v_array(i) = v;
    u_array(i) = u;
    v2_array(i) = v2;
    u2_array(i) = u2;

end

plot(dt-100:dt:steps*dt-100,u2_array,'LineWidth',2,'Color',	"#EDB120");
hold on;	
plot(dt-100:dt:steps*dt-100,v_array,':','LineWidth',2,'Color','k');%,"#D95319");
plot(dt-100:dt:steps*dt-100,v2_array,'LineWidth',3,'Color','k');%"#0072BD");
xlim([-5.05 50.001]);
%xlim([0 steps*dt]);
ylim([-75; 40]);
hold off;        
xlabel('Time (rel. units)');
ylabel('u, v');
legend('u', 'v(Soma)', 'v(Dendrites)')
title(['gSoma=' num2str(gs) ', gDendrite=' num2str(gd)]);
```
