## Supplementary NEURON models for "Dendritic excitations govern back-propagation via a spike-rate accelerometer": FullCA1Model.py

```
from neuron import h, gui
import numpy as np
from scipy.optimize import fsolve
from math import sqrt, exp

class FullCA1Model:

    def __init__(self, channel_params, morphology_file, ka_cutoff):
        CheRiff = False

        # Load the morphology
        self.all = []
        self.load_morphology(morphology_file)

        self.add_passive()

        self.add_rxd_calcium()
        
        self.add_Nav_channels(*channel_params[0])
        self.add_Nav_inactivation(*channel_params[1])
        self.ka_cutoff = ka_cutoff
        self.add_Kap_channels(*channel_params[2])
        self.add_Kad_channels(*channel_params[3])
        self.add_Kdr_channels(*channel_params[4])

        if CheRiff == False:
            self.add_Chr_channels(channel_params[5])
        else:
            self.add_CheRiff_channels(channel_params[5])

        self.add_kca_channel(*channel_params[6])
        self.add_calH_channel(*channel_params[7])
        self.add_cal_channel(*channel_params[8])
        self.add_somacar_channel(*channel_params[9])
        self.add_nap_channel(*channel_params[10])
        self.add_mykca_channel(*channel_params[11])
        self.add_car_channel(*channel_params[12])
        self.add_car_mag_channel(*channel_params[13])
        self.add_cat_channel(*channel_params[14])
        self.add_hcn_channel(*channel_params[15])
        self.add_km_channel(*channel_params[16])
        self.set_axon_conductance()    
        self.geom_nseg(0.1, 100)


        self.times = None
        self.gmax_values = None
        self.time_vector = None
        self.gmax_vector = None
        self.clamps = [] 

    def load_morphology(self, morphology_file):
    
        if morphology_file == 'Jarsky_Gating_Morphology.nrn':
            # Load soma and dendrites
            h.load_file(morphology_file)
            h.load_file("Jarsky_Gating_axon.nrn")
            #self.add_Jarsky_morphology()
            for sec in h.allsec():
                self.all.append(sec)
            self.add_axon_morphology(self.all[0])

        elif morphology_file == 'Jarsky':
            # Load soma and dendrites
            #h.load_file(morphology_file)
            #h.load_file("Jarsky_Gating_axon.nrn")
            self.add_Jarsky_morphology()
            self.add_axon_morphology(self.all[0])
                        
        elif ".swc" in morphology_file:
            # Load hoc files for Import3d
            h.load_file('stdlib.hoc')
            h.load_file('import3d.hoc')
            # Initialize a new Import3d object
            morphology = h.Import3d_SWC_read()
            # Load your SWC file
            morphology.input(morphology_file)
            # Create Import3d_GUI object
            imprt_gui = h.Import3d_GUI(morphology, False)
            # Instantiate the morphology
            imprt_gui.instantiate(None) 
            for sec in h.allsec():
                self.all.append(sec)
            if 'axon' not in [sec.name() for sec in h.SectionList()]:
                 self.add_axon_morphology(self.all[0])

        else:
            print("Error: Not a valid morphology file?")

    def add_Nav_channels(self, g0, g1, d, z):
        self.add_channel('na3', 'gbar_na3', g0, g1, d, z)

    def add_Nav_inactivation(self, g0, g1, d, z):
        self.set_channel('na3', 'ar2_na3', g0, g1, d, z)

    def add_Kap_channels(self, g0, g1, d, z):
        self.add_channel('kap', 'gkabar_kap', g0, g1, d, z)
        soma_sec = next(h.allsec(), None)
        if soma_sec is None:
            print("Warning: No sections to iterate over. Are you sure the morphology has been loaded correctly?")
            return  # exit the method early
        for sec in self.all:
            for seg in sec:
                x = h.distance(soma_sec(0.5), seg)
                if x > self.ka_cutoff:
                    seg.gkabar_kap = 0

    def add_Kad_channels(self, g0, g1, d, z):
        self.add_channel('kad', 'gkabar_kad', g0, g1, d, z)
        soma_sec = next(h.allsec(), None)
        if soma_sec is None:
            print("Warning: No sections to iterate over. Are you sure the morphology has been loaded correctly?")
            return  # exit the method early
        for sec in self.all:
            for seg in sec:
                x = h.distance(soma_sec(0.5), seg)
                if x < self.ka_cutoff:
                    seg.gkabar_kad = 0

    def add_Kdr_channels(self, g0, g1, d, z):
        self.add_channel('kdr', 'gkdrbar_kdr', g0, g1, d, z)

    def add_Chr_channels(self, chrs):
        if not isinstance(chrs[0], list):  # If the input is just a list and not a list of lists
            chrs = [chrs]
        summation_flag = True if len(chrs) > 1 else False
        self.add_channel('chr', 'gexp_chr', 0, 0, 1, 1, 0, 0, summation=False)
        for chr_params in chrs:
            if len(chr_params) == 8:
                g0, g1, d, z, stim_start, stim_end, location, revpot = chr_params
            else:
                g0, g1, d, z, stim_start, stim_end = chr_params
                location = []
                revpot = 0

            self.add_channel('chr', 'gexp_chr', g0, g1, d, z, location, revpot, summation=summation_flag)
            self.set_channel('chr', 'del_chr', 0, 0, stim_start, stim_start)
            self.set_channel('chr', 'dur_chr', 0, 0, stim_end - stim_start, stim_end - stim_start)

    def add_CheRiff_channels(self, chrs):
        if not isinstance(chrs[0], list):  # If the input is just a list and not a list of lists
            chrs = [chrs]
        summation_flag = True if len(chrs) > 1 else False
        self.add_channel('cheriff', 'gmax_cheriff', 0, 0, 1, 1, 0, 0, summation=False)
        for chr_params in chrs:
            if len(chr_params) == 8:
                g0, g1, d, z, stim_start, stim_end, location, revpot = chr_params
            else:
                g0, g1, d, z, stim_start, stim_end = chr_params
                location = []
                revpot = 0

            self.add_channel('cheriff', 'gmax_cheriff', g0, g1, d, z, location, revpot, summation=summation_flag)

    # def add_Chr_channels(self, g0, g1, d, z, stim_start, stim_end, *args, **kwargs):
    #     self.add_channel('chr', 'gexp_chr', g0, g1, d, z, *args, **kwargs)
    #     self.set_channel('chr', 'del_chr', 0, 0, stim_start, stim_start)
    #     self.set_channel('chr', 'dur_chr', 0, 0, stim_end - stim_start, stim_end - stim_start)

    def add_kca_channel(self, g0, g1, d, z):
        self.add_channel('kca', 'gbar_kca', g0, g1, d, z)

    def add_calH_channel(self, g0, g1, d, z):
        self.add_channel('calH', 'gcalbar_calH', g0, g1, d, z)

    def add_cal_channel(self, g0, g1, d, z):
        self.add_channel('cal', 'gcalbar_cal', g0, g1, d, z)

    def add_somacar_channel(self, g0, g1, d, z):
        self.add_channel('somacar', 'gcabar_somacar', g0, g1, d, z)

    def add_nap_channel(self, g0, g1, d, z):
        self.add_channel('nap', 'gnabar_nap', g0, g1, d, z)

    def add_mykca_channel(self, g0, g1, d, z):
        self.add_channel('mykca', 'gkbar_mykca', g0, g1, d, z)

    def add_car_channel(self, g0, g1, d, z):
        self.add_channel('car', 'gcabar_car', g0, g1, d, z)

    def add_car_mag_channel(self, g0, g1, d, z):
        self.add_channel('car_mag', 'gmax_car_mag', g0, g1, d, z)

    def add_cat_channel(self, g0, g1, d, z):
        self.add_channel('cat', 'gcatbar_cat', g0, g1, d, z)

    def add_hcn_channel(self, g0, g1, d, z):
        self.add_channel('hd', 'ghdbar_hd', g0, g1, d, z)

    def add_km_channel(self, g0, g1, d, z):
        self.add_channel('km', 'gbar_km', g0, g1, d, z)

    def distance_soma(self, sec):
        soma_sec = next(h.allsec())
        return h.distance(sec(0.5), soma_sec(0.5))

    # def add_channel(self, channel_name, gvar, x1, x2, density_pre, density_start, density_end, density_post):
    #     soma_sec = next(h.allsec(), None)
    #     if soma_sec is None:
    #         print("Warning: No sections to iterate over. Are you sure the morphology has been loaded correctly?")
    #         return  # exit the method early        
        
    #     for sec in self.all:
    #         sec.insert(channel_name)
    #         # Use setattr to set the conductance attribute dynamically
    #         x = h.distance(sec(0.5),soma_sec(0.5))

    #         # Set the conductance based on the distance
    #         if x < x1:
    #             g = density_pre
    #         elif x1 <= x < x2:
    #             g = density_start + (density_end - density_start) * (x - x1) / (x2 - x1)
    #         else:
    #             g = density_post            

    #         not_dendsoma = 'hill' in sec.name() or 'iseg' in sec.name() or 'node' in sec.name()  or 'axon' in sec.name()
    #         if not_dendsoma:
    #             g = 0

    #         setattr(sec, gvar, g)

    def add_channel(self, channel_name, gvar, g0, g1, d, z, ref_sec_num=None, revpot = 0, summation=False):
        distance_mode = None
        if ref_sec_num is not None:
            ref_sec = self.all[ref_sec_num]
        else:
            soma_sec = next(h.allsec(), None)
            if soma_sec is None:
                print("Warning: No sections to iterate over. Are you sure the morphology has been loaded correctly?")
                return  # exit the method early
            ref_sec = h.cas()  

        for sec in self.all:
            sec.insert(channel_name)
            for seg in sec:
                # Determine distance x based on mode
                if distance_mode == '3d':
                    n3d = int(h.n3d()) 
                    x1, y1, z1 = [h.x3d(i) for i in range(n3d)], [h.y3d(i) for i in range(n3d)], [h.z3d(i) for i in range(n3d)]
                    x2 = sum([(1 - seg.x) * x1[i] + seg.x * x1[i + 1] for i in range(n3d - 1)])
                    y2 = sum([(1 - seg.x) * y1[i] + seg.x * y1[i + 1] for i in range(n3d - 1)])
                    z2 = sum([(1 - seg.x) * z1[i] + seg.x * z1[i + 1] for i in range(n3d - 1)])
                    x = sqrt((x1[0] - x2)**2 + (y1[0] - y2)**2 + (z1[0] - z2)**2)
                else:
                    x = h.distance(ref_sec(0.5), seg)

                if (d - x) / z < -100:
                    g = g1
                elif (d - x) / z > 100:
                    g = g0
                else:
                    g = g0 + (g1 - g0) / (1 + exp((d - x) / z))

                not_dend_or_soma = 'hill' in sec.name() or 'iseg' in sec.name() or 'node' in sec.name() or 'axon' in sec.name()
                if not_dend_or_soma:
                    g = 0

                if summation:
                    g_old = getattr(seg, gvar)
                    g_new = g + g_old                    
                    revpot_old = getattr(seg, "e_chr")
                    if g_new == 0:
                        revpot_new = 0
                    else:
                        revpot_new = (revpot * g + revpot_old * g_old) / g_new
                    setattr(seg, gvar, g_new) 
                    setattr(seg, "e_chr", revpot_new)
                    #print(str(seg) + " " + str(g_new) + " " + str(revpot_new))
                else:
                    setattr(seg, gvar, g)
                    if "chr" in channel_name:
                        setattr(seg, "e_chr", revpot)


    def set_channel(self, channel_name, gvar, g0, g1, d, z, ref_sec_num=None):

        if ref_sec_num is not None:
            ref_sec = self.all[ref_sec_num]
        else:
            soma_sec = next(h.allsec(), None)
            if soma_sec is None:
                print("Warning: No sections to iterate over. Are you sure the morphology has been loaded correctly?")
                return  # exit the method early
            ref_sec = h.cas()

        if channel_name != ('chr' or 'cheriff') and (gvar != 'del_chr' and gvar != 'dur_chr'):
            for sec in self.all:
                #sec.insert(channel_name)
                for seg in sec:
                    x = h.distance(soma_sec(0.5), seg)
                    g = g0 + (g1 - g0) / (1 + exp((d - x) / z))
                    
                    # Check if the section name indicates it is not a dendritic or soma section
                    not_dendsoma = 'hill' in sec.name() or 'iseg' in sec.name() or 'node' in sec.name() or 'axon' in sec.name()
                    if not_dendsoma:
                        g = 0
                    setattr(seg, gvar, g)
        else: 
            for sec in self.all:
                for seg in sec:
                    setattr(seg, gvar, d)


    # def set_channel(self, channel_name, gvar, x1, x2, density_pre, density_start, density_end, density_post):
    #     for sec in self.all:
    #         x = self.distance_soma(sec)
    #         # Set the conductance based on the distance
    #         if x < x1:
    #             g = density_pre
    #         elif x1 <= x < x2:
    #             g = density_start + (density_end - density_start) * (x - x1) / (x2 - x1)
    #         else:
    #             g = density_post            

    #         not_dendsoma = 'hill' in sec.name() or 'iseg' in sec.name() or 'node' in sec.name()  or 'axon' in sec.name()
    #         if not_dendsoma:
    #             g = 0

    #         setattr(sec, gvar, g)

    def add_passive(self):
        Vleak = -66  # Leak reversal in Cs+
        Rm = 40000  # Specific membrane resistivity in ohm-cm^2
        global_ra = 200.00  # Internal resistivity in ohm-cm
        Cm = 0.75  # Specific membrane capacitance in uF/cm^2
        spinefactor = 2.0
        spinelimit = 0 #300

        for sec in h.allsec():
            sec.insert('pas')
            sec.e_pas = Vleak
            if self.distance_soma(sec) <= spinelimit:
                sec.g_pas = 1 / Rm
                sec.cm = Cm
            else:
                sec.g_pas = spinefactor / Rm
                sec.cm = spinefactor * Cm
            sec.Ra = global_ra


    def add_rxd_calcium(self):
        for sec in h.allsec():
           # sec.insert('cad')
            sec.insert('cacum')

    def chr_vectorplay(self):
        freq = 15 # Hz
        amplitude = 0.02 # S/cm2
        offset = 1
        cond_vec = h.Vector()
        for t in np.arange(0, 1000 + h.dt, h.dt):
            cond = amplitude * (offset + h.sin(2 * h.PI * freq * (t-25) / 1000))
            cond_vec.append(cond)
        soma = self.all[0]
        cond_vec.play(soma.g_chr, h.dt)
        return cond, cond_vec, soma

    # def update_gmax(self):
    #     soma_sec = next(h.allsec(),None)
    #     t = h.t
    #     index = int(t/h.dt) 
    #     if 0 <= index < len(self.time_vector):
    #         new_gmax = self.gmax_vector.x[index]
    #         soma_sec.g_chr = new_gmax
    #         #rec_gmax.append(new_gmax) 
    # 
    # def prepare_simulation(self, times, gmax_values):
    #     self.time_vector = h.Vector()
    #     self.gmax_vector = h.Vector()
    #     self.rec_gmax = h.Vector()
    #     self.times = times
    #     self.gmax_values = gmax_values
    #     self.time_vector.from_python(list(times))
    #     self.gmax_vector.from_python(list(gmax_values))
    #     cvode = h.CVode()
    #     cvode.extra_scatter_gather(0, self.update_gmax)
        
    def prepare_simulation(self, times, gmax_values):
        self.time_vector = h.Vector()
        self.gmax_vectors = [h.Vector() for _ in self.all]
        self.rec_gmax = h.Vector()
        self.times = times
        # gmax_values should be a 2D list or array with shape (num_sections, num_timepoints)
        self.gmax_values = gmax_values
        self.time_vector.from_python(list(times))
        # Fill each gmax_vector from the corresponding row of gmax_values
        for vec, values in zip(self.gmax_vectors, gmax_values):
            vec.from_python(list(values))
        cvode = h.CVode()
        cvode.extra_scatter_gather(0, self.update_gmax)
    
    def update_gmax(self):
        self.allgmax = []
        for sec in h.allsec():
            self.allgmax.append(sec)
        t = h.t
        index = int(t/h.dt) 
        if 0 <= index < len(self.time_vector):
            for sec, gmax_vector in zip(self.allgmax, self.gmax_vectors):
                new_gmax = gmax_vector.x[index]
                sec.g_chr = new_gmax

    def prepare_simulation_cheriff(self, times, gmax_values):
        self.time_vector = h.Vector()
        self.gmax_vectors = [h.Vector() for _ in self.all]
        self.rec_gmax = h.Vector()
        self.times = times
        # gmax_values should be a 2D list or array with shape (num_sections, num_timepoints)
        self.gmax_values = gmax_values
        self.time_vector.from_python(list(times))
        # Fill each gmax_vector from the corresponding row of gmax_values
        for vec, values in zip(self.gmax_vectors, gmax_values):
            vec.from_python(list(values))
        cvode = h.CVode()
        cvode.extra_scatter_gather(0, self.update_gmax_cheriff)

    def update_gmax_cheriff(self):
        self.allgmax = []
        for sec in h.allsec():
            self.allgmax.append(sec)
        t = h.t
        index = int(t/h.dt) 
        if 0 <= index < len(self.time_vector):
            for sec, gmax_vector in zip(self.allgmax, self.gmax_vectors):
                new_gmax = gmax_vector.x[index]
                sec.lightintensity_cheriff = new_gmax

    def set_axon_conductance(self):
        gnabar = 0.05  # Sodium conductance
        gnode = 30.0  # Sodium conductance at a node
        gkdr = 0.040  # Delayed rectifier potassium conductance
        gkap = 0.048  # A-type potassium conductance
        Rn = 50  # Nodal resistivity
        Rm = 40000  # specific membrane resistivity in ohm-cm^2 
        Cmy = 0.04  # Capacitance in myelin

        for sec in h.allsec():
            
            if 'hill' in sec.name():
                sec.gbar_na3 = gnabar
                sec.gkdrbar_kdr = gkdr
                sec.gkabar_kap = gkap

            if 'iseg' in sec.name():
                sec.gbar_na3 = gnode/2#/10
                sec.gkdrbar_kdr = gkdr
                sec.gkabar_kap = gkap

            if 'node' in sec.name(): 
                sec.gbar_na3 = gnode
                sec.gkdrbar_kdr = gkdr
                sec.gkabar_kap = gkap * 0.2
                sec.g_pas = 1 / Rn

            if 'inode' in sec.name():            
                sec.gbar_na3 = gnabar
                sec.gkdrbar_kdr = gkdr
                sec.gkabar_kap = gkap * 0.2
                sec.cm = Cmy
                sec.g_pas = 1 / Rm
              
            if 'axon' in sec.name():
                sec.gbar_na3 = gnabar*1.4
                sec.ar2_na3 = 1
                sec.gkdrbar_kdr = gkdr
                sec.gkabar_kap = gkap * 0.2
                sec.cm = Cmy
                sec.g_pas = 1 / Rm

    def inverse_cdf(self,y,cdf):
        return fsolve(lambda x: cdf(x) - y, 0)

    def distribute_synapses(self, num_synapses, cdf, location, synapse_type):

        synapse_list = []
        synapse_positions = []  # Added this line

        # Initialize random number generator
        rng = np.random.default_rng()

        # Create a map of distances from soma to points 3um apart in each section
        distance_map = {}

        min_distance = 50       
        max_distance = 2000 

        if location == "basal":
            avoid = "apic"
        elif location == "apical":
            avoid = "dend"
        elif location == "distal":
            avoid = "dend"
            min_distance = 300
        elif location == "proximal":
            max_distance = 300
            avoid = "soma"
        else:
            avoid = "soma"
        
        for sec in self.all:
            not_dend = 'hill' in sec.name() or 'iseg' in sec.name() or 'node' in sec.name() or 'inode' in sec.name() or 'soma' in sec.name()  or 'axon' in sec.name() or avoid in sec.name()
            if not not_dend:
                
                # Distance from soma to center of section
                center_distance = self.distance_soma(sec)

                # Create points every 3um along section
                num_points = int(sec.L / 3) + 1
                for i in range(num_points):
                    location = i / max((num_points - 1),2)
                    point_distance = center_distance - (sec.L / 2) + (sec.L * location)
                    distance_map[sec, location] = point_distance

        for _ in range(num_synapses):
            # Draw random number and find corresponding distance
            rand_num = rng.random()
            target_distance = self.inverse_cdf(rand_num,cdf) * (max_distance - min_distance) + min_distance
            closest_distance = min(distance_map.values(), key=lambda x: abs(x - target_distance))

            # Find section and location with closest distance to target
            target_sec = [sec for sec, loc in distance_map.keys() if distance_map[sec, loc] == closest_distance][0]
            target_loc = [loc for sec, loc in distance_map.keys() if distance_map[sec, loc] == closest_distance][0]
            
            if target_loc == 1:
                target_loc -= 0.01
            elif target_loc == 0:
                target_loc += 0.01
                
            # Create synapse at target location
            if synapse_type == "ampa":
                syn = h.AmpaSyn(target_sec(target_loc))
                synapse_list.append(syn)
            elif synapse_type == "nmda":
                syn = h.NmdaSyn(target_sec(target_loc))
                synapse_list.append(syn)
            elif synapse_type == "ampa+nmda":
                syn_nmda = h.NmdaSyn(target_sec(target_loc))
                syn_ampa = h.AmpaSyn(target_sec(target_loc))
                synapse_list.append((syn_ampa, syn_nmda))
            elif synapse_type == "gabaa":
                syn = h.GABAa(target_sec(target_loc))
                synapse_list.append(syn)
            elif synapse_type == "gabab":
                syn = h.GABAb(target_sec(target_loc))
                synapse_list.append(syn)
            elif synapse_type == "exp2syn":
                syn = h.Exp2Syn(target_sec(target_loc))
                syn.e = 0
                syn.tau1 = 0.1
                syn.tau2 = 1.0
                synapse_list.append(syn)
            #elif synapse_type == "tsodyks-markram"
            #    syn = h.tsodyks_markram(target_sec(target_loc))
            else:
                print("Invalid synapse type!")
                

            # Get the 3D position of the synapse
            x = target_sec.x3d(int(target_loc * (target_sec.n3d() - 1)))
            y = target_sec.y3d(int(target_loc * (target_sec.n3d() - 1)))
            z = target_sec.z3d(int(target_loc * (target_sec.n3d() - 1)))
            synapse_positions.append((x, y, z))  

        return synapse_list, synapse_positions  # Return positions as well

    def generate_spike_train(self, start_time, end_time, avg_spike_rate, distribution_type):
        avg_spike_interval = 1000 / avg_spike_rate  # average time between spikes (ms)
        
        if distribution_type == 'regular':
            # For regularly spaced spikes, use arange to create an array of spike times
            spike_times = np.arange(start_time, end_time, avg_spike_interval)
            
        elif distribution_type == 'poisson':
            # For a Poisson process, inter-spike intervals are exponentially distributed
            num_spikes = int((end_time - start_time) * avg_spike_rate)
            ISIs = np.random.exponential(avg_spike_interval, num_spikes)
            spike_times = np.cumsum(ISIs) + start_time
    
        elif distribution_type == 'gamma':
            shape = 2  # arbitrary shape parameter for the gamma distribution
            scale = avg_spike_interval / shape  # scale parameter for the gamma distribution
            num_spikes = int((end_time - start_time) * avg_spike_rate)
            ISIs = np.random.gamma(shape, scale, num_spikes)
            spike_times = np.cumsum(ISIs) + start_time
    
        elif distribution_type == 'log-normal':
            # Correction. Needed to get correct avg. firing rate
            corr = 1.5; 
            # log normal distribution
            mu = np.log(avg_spike_interval/corr)  # log of the mean of the log-normal distribution
            sigma = 1.0  # standard deviation of the log-normal distribution
            num_spikes = int((end_time - start_time) * avg_spike_rate * corr)
            ISIs = np.random.lognormal(mu, sigma, num_spikes)
            spike_times = np.cumsum(ISIs) + start_time

        elif distribution_type == 'ramp':
            total_time = end_time - start_time
            num_spikes = int(avg_spike_rate * total_time / 1000)
            asym = 4 
            mode = start_time / asym + end_time * (1 - 1/asym)
            spike_times = np.random.triangular(start_time, mode, end_time, num_spikes)        
            spike_times = np.sort(spike_times)
        
        elif distribution_type == 'theta':
            # create the theta wave modulation
            theta_freq = 8  # theta frequency in Hz
            Amp = 0.1/theta_freq*1000  # Amplitude of theta wave in ms. 0.05 for 1:2 contrast, 0.1 for 1:5

            # Start with a base Poisson process
            num_spikes = int(((end_time - start_time) / 1000) * avg_spike_rate)*2
            ISIs = np.random.exponential(avg_spike_interval, num_spikes)
            #theta_wave = Amp *( offset + np.sin(2 * np.pi * theta_freq * spike_times / 1000))
            spike_times = np.cumsum(ISIs) + start_time - (end_time - start_time)/2
    
            # shift spike times according to the theta wave
            theta_wave = Amp * np.sin(2 * np.pi * theta_freq * spike_times / 1000)
            modulated_spike_times = spike_times + theta_wave
    
            # Ensure spikes times remain within the time interval
            modulated_spike_times = modulated_spike_times[(modulated_spike_times >= start_time) & (modulated_spike_times <= end_time)]
            spike_times = modulated_spike_times    
            
        else:
            raise ValueError('Invalid distribution type. Choose from "regular", "poisson", "gamma", "log-normal", "ramp" or "theta".')
        
        spike_times = spike_times[(spike_times  >= start_time) & (spike_times <= end_time)]
        return spike_times


    def distribute_spike_times(self, synapse_list, start_time, end_time, spike_rate, synapse_strength, distribution_type, ratio):
        # List to store NetStim and NetCon objects
        netstims = []
        netcons = []
        all_spike_times = []
    
        # Create a NetStim and NetCon for each synapse
        for synapse_entry in synapse_list:
            # If a tuple is found (ampa+nmda), unpack it
            if isinstance(synapse_entry, tuple):
                syn_ampa, syn_nmda = synapse_entry
                ampa_strength = synapse_strength 
                nmda_strength = synapse_strength * ratio
                synapses_weights = [(syn_ampa, ampa_strength), (syn_nmda, nmda_strength)]
            else:
                synapses_weights = [(synapse_entry, synapse_strength)]
    
            # Generate spike times
            spike_times = self.generate_spike_train(start_time, end_time, spike_rate, distribution_type)
    
            for synapse, weight in synapses_weights:
                for spike_time in spike_times:
                    stim = h.NetStim()
                    stim.number = 1
                    stim.start = spike_time
                
                    nc = h.NetCon(stim, synapse)
                    nc.weight[0] = weight  # Use the adjusted weight
                
                    netstims.append(stim)
                    netcons.append(nc)
                
            all_spike_times.append(spike_times)
    
        return netstims, netcons, all_spike_times


    def add_patch(self, loc, waveform, clamp_type):
        location = self.all[loc]
        if clamp_type.lower() == 'v':
            iclamp = h.VClamp(location(0.5))
        elif clamp_type.lower() == 'i':
            iclamp = h.IClamp(location(0.5))
        elif clamp_type.lower() == 'chr':
            print("This feature is not yet implemented.")
        else:
            raise ValueError("Invalid clamp type. Use 'v' for Voltage Clamp, 'i' for Current Clamp, or 'chr' for Local Conductance.")
        iclamp.dur = 1e9
        vec_waveform = h.Vector(waveform)
        vec_waveform.play(iclamp._ref_amp, h.dt)
        # iclamp.dur = 50
        # iclamp.amp = 2
        # iclamp.delay = 10
        recorded_amp = h.Vector()
        recorded_amp.record(iclamp._ref_amp)

        patch_objects = (iclamp, location, vec_waveform)
        self.clamps.append(patch_objects)

        return patch_objects, recorded_amp
    

    def add_axon_morphology(self, som):

        hill = h.Section(name='hill')

        hill.nseg = 11

        hill.pt3dclear()
        hill.pt3dadd(0.00, 0.00, 0.00, 4.04)
        hill.pt3dadd(-0.25, -1.00, 0.00, 3.73)
        hill.pt3dadd(-0.79, -2.04, 0.00, 3.44)
        hill.pt3dadd(-1.27, -3.02, 0.00, 3.11)
        hill.pt3dadd(-1.33, -3.96, 0.00, 2.76)
        hill.pt3dadd(-1.32, -4.84, 0.00, 2.40)
        hill.pt3dadd(-1.50, -5.80, 0.00, 2.11)
        hill.pt3dadd(-0.80, -6.82, 0.00, 1.78)
        hill.pt3dadd(-0.47, -7.92, 0.00, 1.39)
        hill.pt3dadd(-0.30, -8.98, 0.00, 1.05)
        hill.pt3dadd(-0.43, -10.04, 0.00, 1.05)
        hill.connect(som, 1)

        self.all.append(hill)
        
        iseg = h.Section(name='iseg')
        iseg.nseg = 11
        iseg.pt3dclear()
        iseg.pt3dadd(-0.53, -10.04, 0.00, 0.96)
        iseg.pt3dadd(0.43, -11.56, 0.00, 1.08)
        iseg.pt3dadd(0.07, -13.07, 0.00, 0.91)
        iseg.pt3dadd(-0.57, -14.58, 0.00, 1.16)
        iseg.pt3dadd(-0.60, -16.18, 0.00, 1.10)
        iseg.pt3dadd(-1.18, -17.72, 0.00, 0.94)
        iseg.pt3dadd(-1.03, -19.26, 0.00, 1.08)
        iseg.pt3dadd(-1.17, -20.88, 0.00, 0.89)
        iseg.pt3dadd(-1.40, -22.35, 0.00, 0.98)
        iseg.pt3dadd(-1.02, -23.93, 0.00, 0.95)
        iseg.pt3dadd(-0.90, -25.49, 0.00, 1.10)
        iseg.connect(hill, 1)
        self.all.append(iseg)
        
        inode1 = h.Section(name='inode1')
        inode1.nseg = 51
        inode1.pt3dclear()
        inode1.pt3dadd(-0.90, -25.49, 0.00, 1.40)
        inode1.pt3dadd(-1.48, -27.59, 0.00, 1.53)
        inode1.pt3dadd(0.40, -29.60, 0.00, 1.64)
        inode1.pt3dadd(1.63, -31.59, 0.00, 1.42)
        inode1.pt3dadd(1.43, -33.73, 0.00, 1.69)
        inode1.pt3dadd(1.16, -35.92, 0.00, 1.31)
        inode1.pt3dadd(1.97, -37.95, 0.00, 1.53)
        inode1.pt3dadd(2.76, -39.90, 0.00, 1.76)
        inode1.pt3dadd(2.19, -41.79, 0.00, 1.46)
        inode1.pt3dadd(1.61, -43.75, 0.00, 1.51)
        inode1.pt3dadd(-0.27, -45.70, 0.00, 1.48)
        inode1.pt3dadd(-0.25, -47.92, 0.00, 1.58)
        inode1.pt3dadd(-0.68, -50.02, 0.00, 1.49)
        inode1.pt3dadd(-0.57, -51.93, 0.00, 1.69)
        inode1.pt3dadd(-1.69, -53.77, 0.00, 1.54)
        inode1.pt3dadd(-1.84, -55.81, 0.00, 1.60)
        inode1.pt3dadd(-0.62, -57.77, 0.00, 1.52)
        inode1.pt3dadd(-1.57, -59.79, 0.00, 1.55)
        inode1.pt3dadd(-2.00, -61.64, 0.00, 1.51)
        inode1.pt3dadd(-1.36, -63.57, 0.00, 1.66)
        inode1.pt3dadd(-0.39, -65.44, 0.00, 1.56)
        inode1.pt3dadd(-0.46, -67.39, 0.00, 1.51)
        inode1.pt3dadd(-0.19, -69.27, 0.00, 1.60)
        inode1.pt3dadd(-2.07, -71.32, 0.00, 1.61)
        inode1.pt3dadd(-3.34, -73.17, 0.00, 1.44)
        inode1.pt3dadd(-4.58, -75.08, 0.00, 1.53)
        inode1.pt3dadd(-4.31, -76.99, 0.00, 1.45)
        inode1.pt3dadd(-4.00, -78.99, 0.00, 1.46)
        inode1.pt3dadd(-4.09, -80.92, 0.00, 1.53)
        inode1.pt3dadd(-4.80, -82.95, 0.00, 1.41)
        inode1.pt3dadd(-5.63, -84.88, 0.00, 1.57)
        inode1.pt3dadd(-6.75, -86.89, 0.00, 1.69)
        inode1.pt3dadd(-9.07, -88.99, 0.00, 1.54)
        inode1.pt3dadd(-8.50, -91.21, 0.00, 1.53)
        inode1.pt3dadd(-7.20, -93.23, 0.00, 1.46)
        inode1.pt3dadd(-6.14, -95.19, 0.00, 1.58)
        inode1.pt3dadd(-6.10, -97.05, 0.00, 1.50)
        inode1.pt3dadd(-5.64, -98.96, 0.00, 1.63)
        inode1.pt3dadd(-3.08, -100.98, 0.00, 1.36)
        inode1.pt3dadd(-1.48, -102.73, 0.00, 1.53)
        inode1.pt3dadd(-1.98, -104.46, 0.00, 1.57)
        inode1.pt3dadd(-1.71, -106.32, 0.00, 1.52)
        inode1.pt3dadd(-2.07, -108.41, 0.00, 1.57)
        inode1.pt3dadd(-0.39, -110.45, 0.00, 1.57)
        inode1.pt3dadd(-0.88, -112.45, 0.00, 1.38)
        inode1.pt3dadd(-0.44, -114.49, 0.00, 1.63)
        inode1.pt3dadd(-0.79, -116.49, 0.00, 1.53)
        inode1.pt3dadd(-0.93, -118.43, 0.00, 1.41)
        inode1.pt3dadd(-1.01, -120.40, 0.00, 1.46)
        inode1.pt3dadd(-0.70, -122.55, 0.00, 1.54)
        inode1.pt3dadd(-0.65, -124.54, 0.00, 1.52)
        inode1.connect(iseg, 1)
        self.all.append(inode1)

        
        node1 = h.Section(name='node1')
        node1.nseg = 2
        node1.pt3dclear()
        node1.pt3dadd(-0.65, -124.54, 0.00, 1.01)
        node1.pt3dadd(0.00, -125.70, 0.00, 1.09)
        node1.connect(inode1, 1)
        self.all.append(node1)
        
        inode2 = h.Section(name='inode2')
        inode2.nseg = 51
        inode2.pt3dclear()
        inode2.pt3dadd(0.00, -125.70, 0.00, 1.39)
        inode2.pt3dadd(0.00, -125.70, 0.00, 1.39)
        inode2.pt3dadd(-0.76, -127.73, 0.00, 1.38)
        inode2.pt3dadd(-1.17, -129.64, 0.00, 1.34)
        inode2.pt3dadd(-1.29, -131.74, 0.00, 1.47)
        inode2.pt3dadd(-1.47, -133.86, 0.00, 1.44)
        inode2.pt3dadd(-2.39, -135.72, 0.00, 1.63)
        inode2.pt3dadd(-1.94, -137.79, 0.00, 1.62)
        inode2.pt3dadd(-1.61, -139.84, 0.00, 1.50)
        inode2.pt3dadd(0.50, -141.68, 0.00, 1.39)
        inode2.pt3dadd(0.96, -143.67, 0.00, 1.49)
        inode2.pt3dadd(1.40, -145.81, 0.00, 1.46)
        inode2.pt3dadd(1.94, -147.70, 0.00, 1.27)
        inode2.pt3dadd(3.47, -149.59, 0.00, 1.63)
        inode2.pt3dadd(4.63, -151.39, 0.00, 1.46)
        inode2.pt3dadd(3.26, -153.33, 0.00, 1.43)
        inode2.pt3dadd(3.51, -155.53, 0.00, 1.47)
        inode2.pt3dadd(4.21, -157.74, 0.00, 1.51)
        inode2.pt3dadd(4.10, -159.71, 0.00, 1.66)
        inode2.pt3dadd(2.99, -161.48, 0.00, 1.48)
        inode2.pt3dadd(3.27, -163.50, 0.00, 1.41)
        inode2.pt3dadd(4.16, -165.33, 0.00, 1.35)
        inode2.pt3dadd(5.02, -167.32, 0.00, 1.68)
        inode2.pt3dadd(6.92, -169.16, 0.00, 1.38)
        inode2.pt3dadd(7.12, -171.03, 0.00, 1.41)
        inode2.pt3dadd(6.81, -173.20, 0.00, 1.60)
        inode2.pt3dadd(6.36, -175.37, 0.00, 1.38)
        inode2.pt3dadd(6.34, -177.22, 0.00, 1.39)
        inode2.pt3dadd(7.12, -179.33, 0.00, 1.49)
        inode2.pt3dadd(7.17, -181.38, 0.00, 1.55)
        inode2.pt3dadd(8.09, -183.45, 0.00, 1.53)
        inode2.pt3dadd(9.47, -185.63, 0.00, 1.52)
        inode2.pt3dadd(9.02, -187.74, 0.00, 1.59)
        inode2.pt3dadd(7.66, -189.86, 0.00, 1.42)
        inode2.pt3dadd(7.09, -191.95, 0.00, 1.43)
        inode2.pt3dadd(6.69, -193.89, 0.00, 1.50)
        inode2.pt3dadd(8.23, -195.90, 0.00, 1.35)
        inode2.pt3dadd(7.07, -197.93, 0.00, 1.57)
        inode2.pt3dadd(7.31, -199.87, 0.00, 1.53)
        inode2.pt3dadd(7.65, -201.78, 0.00, 1.37)
        inode2.pt3dadd(5.95, -203.68, 0.00, 1.32)
        inode2.pt3dadd(4.86, -205.61, 0.00, 1.56)
        inode2.pt3dadd(5.41, -207.54, 0.00, 1.55)
        inode2.pt3dadd(4.99, -209.48, 0.00, 1.57)
        inode2.pt3dadd(4.04, -211.26, 0.00, 1.46)
        inode2.pt3dadd(5.92, -213.18, 0.00, 1.38)
        inode2.pt3dadd(6.96, -214.97, 0.00, 1.44)
        inode2.pt3dadd(7.31, -217.01, 0.00, 1.48)
        inode2.pt3dadd(7.25, -219.02, 0.00, 1.53)
        inode2.pt3dadd(6.58, -221.06, 0.00, 1.47)
        inode2.pt3dadd(7.22, -223.13, 0.00, 1.38)
        inode2.pt3dadd(7.31, -225.06, 0.00, 1.65)
        inode2.connect(node1, 1)
        self.all.append(inode2)
        
        node2 = h.Section(name='node2')
        node2.nseg = 2
        node2.pt3dclear()
        node2.pt3dadd(7.31, -225.06, 0.00, 1.07)
        node2.pt3dadd(5.10, -225.83, 0.00, 1.00)
        node2.connect(inode2, 1)
        self.all.append(node2)
        
        inode3 = h.Section(name='inode3')
        inode3.nseg = 51
        inode3.pt3dclear()
        inode3.pt3dadd(5.10, -225.83, 0.00, 1.48)
        inode3.pt3dadd(5.10, -225.83, 0.00, 1.48)
        inode3.pt3dadd(4.58, -227.83, 0.00, 1.55)
        inode3.pt3dadd(4.16, -229.66, 0.00, 1.34)
        inode3.pt3dadd(4.17, -231.62, 0.00, 1.53)
        inode3.pt3dadd(4.64, -233.57, 0.00, 1.61)
        inode3.pt3dadd(2.81, -235.77, 0.00, 1.44)
        inode3.pt3dadd(1.03, -237.77, 0.00, 1.35)
        inode3.pt3dadd(0.58, -239.96, 0.00, 1.47)
        inode3.pt3dadd(-0.65, -242.05, 0.00, 1.65)
        inode3.pt3dadd(-0.57, -243.87, 0.00, 1.45)
        inode3.pt3dadd(-0.87, -245.85, 0.00, 1.56)
        inode3.pt3dadd(-2.21, -247.82, 0.00, 1.23)
        inode3.pt3dadd(-2.65, -249.79, 0.00, 1.53)
        inode3.pt3dadd(-2.80, -251.68, 0.00, 1.46)
        inode3.pt3dadd(-3.07, -253.62, 0.00, 1.40)
        inode3.pt3dadd(-3.47, -255.70, 0.00, 1.50)
        inode3.pt3dadd(-4.07, -257.73, 0.00, 1.43)
        inode3.pt3dadd(-5.49, -259.64, 0.00, 1.39)
        inode3.pt3dadd(-5.40, -261.68, 0.00, 1.41)
        inode3.pt3dadd(-5.61, -263.83, 0.00, 1.57)
        inode3.pt3dadd(-4.23, -265.91, 0.00, 1.47)
        inode3.pt3dadd(-3.79, -268.13, 0.00, 1.53)
        inode3.pt3dadd(-3.83, -270.25, 0.00, 1.52)
        inode3.pt3dadd(-3.56, -272.23, 0.00, 1.23)
        inode3.pt3dadd(-4.32, -274.27, 0.00, 1.59)
        inode3.pt3dadd(-3.86, -276.32, 0.00, 1.38)
        inode3.pt3dadd(-4.07, -278.36, 0.00, 1.49)
        inode3.pt3dadd(-4.13, -280.38, 0.00, 1.68)
        inode3.pt3dadd(-4.86, -282.47, 0.00, 1.67)
        inode3.pt3dadd(-4.31, -284.38, 0.00, 1.52)
        inode3.pt3dadd(-4.74, -286.23, 0.00, 1.54)
        inode3.pt3dadd(-4.25, -288.29, 0.00, 1.58)
        inode3.pt3dadd(-4.55, -290.46, 0.00, 1.41)
        inode3.pt3dadd(-4.09, -292.42, 0.00, 1.45)
        inode3.pt3dadd(-4.10, -294.46, 0.00, 1.55)
        inode3.pt3dadd(-3.96, -296.35, 0.00, 1.40)
        inode3.pt3dadd(-3.63, -298.28, 0.00, 1.47)
        inode3.pt3dadd(-2.97, -300.27, 0.00, 1.52)
        inode3.pt3dadd(-4.67, -302.39, 0.00, 1.31)
        inode3.pt3dadd(-3.15, -304.39, 0.00, 1.54)
        inode3.pt3dadd(-2.60, -306.29, 0.00, 1.64)
        inode3.pt3dadd(-1.61, -308.31, 0.00, 1.68)
        inode3.pt3dadd(-0.69, -310.36, 0.00, 1.56)
        inode3.pt3dadd(-2.15, -312.34, 0.00, 1.60)
        inode3.pt3dadd(-3.04, -314.45, 0.00, 1.56)
        inode3.pt3dadd(-3.50, -316.54, 0.00, 1.42)
        inode3.pt3dadd(-2.47, -318.50, 0.00, 1.48)
        inode3.pt3dadd(-1.99, -320.42, 0.00, 1.37)
        inode3.pt3dadd(-0.49, -322.23, 0.00, 1.49)
        inode3.pt3dadd(-0.19, -324.22, 0.00, 1.55)
        inode3.pt3dadd(-0.48, -326.23, 0.00, 1.56)
        inode3.connect(node2, 1)
        self.all.append(inode3)

    def add_Jarsky_morphology(self):
        soma = h.Section(name='soma')
        soma.nseg = 42
        soma.pt3dclear()
        soma.pt3dadd(0, 10.200000, 0.000000, 4.935061)
        soma.pt3dadd(0, 10.383960, 0.000000, 5.00)
        soma.pt3dadd(0, 9.824203, 0.000000, 5.034562)
        soma.pt3dadd(0, 9.073533, 0.000000, 5.134562)
        soma.pt3dadd(0, 8.413338, 0.000000, 5.971574)
        soma.pt3dadd(0, 7.096380, 0.000000, 10.381932)
        soma.pt3dadd(0, 6.398985, 0.000000, 10.929527)
        soma.pt3dadd(0, 5.582579, 0.000000, 11.707934)
        soma.pt3dadd(0, 4.945832, 0.000000, 12.534684)
        soma.pt3dadd(0, 4.214682, 0.000000, 13.118218)
        soma.pt3dadd(0, 3.519914, 0.000000, 13.704884)
        soma.pt3dadd(0, 2.638938, 0.000000, 14.477144)
        soma.pt3dadd(0, 1.854945, 0.000000, 14.707770)
        soma.pt3dadd(0, 1.008797, 0.000000, 15.347794)
        soma.pt3dadd(0, 0.224154, 0.000000, 14.957293)
        soma.pt3dadd(0, -0.503870, 0.000000, 15.471229)
        soma.pt3dadd(0, -1.162426, 0.000000, 14.750806)
        soma.pt3dadd(0, -1.926060, 0.000000, 14.787425)
        soma.pt3dadd(0, -2.705498, 0.000000, 14.152356)
        soma.pt3dadd(0, -3.370046, 0.000000, 13.472021)
        soma.pt3dadd(0, -4.099295, 0.000000, 12.055820)
        soma.pt3dadd(0, -4.733515, 0.000000, 11.312530)
        soma.pt3dadd(0, -5.484253, 0.000000, 10.915047)
        soma.pt3dadd(0, -6.261007, 0.000000, 10.231184)
        soma.pt3dadd(0, -7.062121, 0.000000, 8.872304)
        soma.pt3dadd(0, -7.865947, 0.000000, 7.839291)
        soma.pt3dadd(0, -8.659434, 0.000000, 7.836026)
        soma.pt3dadd(0, -9.447193, 0.000000, 6.741406)
        soma.pt3dadd(0, -10.207499, 0.000000, 6.422529)
        soma.pt3dadd(0, -10.962002, 0.000000, 6.200278)
        soma.pt3dadd(0, -11.694167, 0.000000, 5.373839)
        soma.pt3dadd(0, -12.436392, 0.000000, 4.809214)
        soma.pt3dadd(0, -13.095034, 0.000000, 3.895101)
        soma.pt3dadd(0, -13.769650, 0.000000, 4.600733)
        soma.pt3dadd(0, -14.120416, 0.000000, 3.000000)
        soma.pt3dadd(0, -14.871800, 0.000000, 3.0)
        soma.pt3dadd(0, -15.405928, 0.000000, 3.0)
        soma.pt3dadd(0, -15.670000, 0.000000, 3.0)
   
        dend = [None]*15

        dend[0] = h.Section(name='dend[0]')
        dend[0].connect(        soma(1), 0)
        dend[0].nseg = 1
        dend[0].pt3dclear()
        dend[0].pt3dadd(-0.72, -16.42, 0.00, 2.4)
        dend[0].pt3dadd(-0.72, -18.16, 0.00, 2.2)
        dend[0].pt3dadd(-0.72, -22.14, 0.00, 2.0)
        dend[0].pt3dadd(-0.72, -24.13, 0.00, 1.96)

        dend[1] = h.Section(name='dend[1]')
        dend[1].connect(        dend[0](1), 0)
        dend[1].nseg = 22
        dend[1].pt3dclear()
        dend[1].pt3dadd(-0.72, -24.13, 0.00, 0.77)
        dend[1].pt3dadd(-3.62, -25.87, 0.00, 0.77)
        dend[1].pt3dadd(-3.86, -25.87, 0.00, 0.77)
        dend[1].pt3dadd(-4.83, -27.36, 0.00, 0.77)
        dend[1].pt3dadd(-4.83, -28.36, 0.00, 0.77)
        dend[1].pt3dadd(-4.59, -28.86, 0.00, 0.77)
        dend[1].pt3dadd(-4.59, -29.35, 0.00, 0.77)
        dend[1].pt3dadd(-4.59, -30.60, 0.00, 0.77)
        dend[1].pt3dadd(-4.83, -32.84, -2.00, 0.77)
        dend[1].pt3dadd(-5.07, -34.83, -2.00, 0.77)
        dend[1].pt3dadd(-6.04, -37.31, -2.00, 0.77)
        dend[1].pt3dadd(-7.00, -39.80, -2.00, 0.77)
        dend[1].pt3dadd(-7.97, -42.54, -2.00, 0.77)
        dend[1].pt3dadd(-7.97, -42.79, -2.00, 0.77)
        dend[1].pt3dadd(-6.76, -43.78, -2.00, 0.77)
        dend[1].pt3dadd(-6.52, -44.28, -2.00, 0.77)
        dend[1].pt3dadd(-5.31, -45.77, -2.00, 0.77)
        dend[1].pt3dadd(-5.31, -46.02, -2.00, 0.61)
        dend[1].pt3dadd(-5.07, -47.76, -2.00, 0.61)
        dend[1].pt3dadd(-5.31, -48.01, -2.00, 0.61)
        dend[1].pt3dadd(-6.04, -49.50, -2.00, 0.61)
        dend[1].pt3dadd(-7.00, -51.24, -2.00, 0.61)
        dend[1].pt3dadd(-7.97, -52.99, -2.59, 0.61)
        dend[1].pt3dadd(-7.25, -54.98, -2.59, 0.61)
        dend[1].pt3dadd(-7.97, -57.71, -2.59, 0.61)
        dend[1].pt3dadd(-8.94, -59.70, -2.59, 0.61)
        dend[1].pt3dadd(-8.94, -61.69, -2.59, 0.61)
        dend[1].pt3dadd(-8.70, -66.17, 4.61, 0.61)
        dend[1].pt3dadd(-7.49, -69.15, 4.61, 0.61)
        dend[1].pt3dadd(-7.49, -69.40, 4.61, 0.61)
        dend[1].pt3dadd(-7.73, -71.39, 4.61, 0.61)
        dend[1].pt3dadd(-7.73, -72.64, 5.17, 0.61)
        dend[1].pt3dadd(-8.94, -75.12, 5.17, 0.61)
        dend[1].pt3dadd(-7.00, -78.36, 5.17, 0.61)
        dend[1].pt3dadd(-9.42, -79.10, 5.17, 0.61)
        dend[1].pt3dadd(-9.42, -79.35, 5.17, 0.61)
        dend[1].pt3dadd(-7.49, -83.08, 5.17, 0.61)
        dend[1].pt3dadd(-7.25, -83.08, 5.17, 0.61)
        dend[1].pt3dadd(-8.21, -86.82, 5.11, 0.61)
        dend[1].pt3dadd(-9.18, -89.55, 5.11, 0.61)
        dend[1].pt3dadd(-13.04, -95.27, -8.91, 0.61)
        dend[1].pt3dadd(-12.80, -95.27, -8.91, 0.61)
        dend[1].pt3dadd(-14.01, -100.25, -8.91, 0.61)
        dend[1].pt3dadd(-15.70, -103.48, -8.91, 0.61)
        dend[1].pt3dadd(-16.67, -106.72, -8.91, 0.61)
        dend[1].pt3dadd(-15.70, -111.44, -8.91, 0.61)
        dend[1].pt3dadd(-15.70, -115.42, -8.91, 0.61)
        dend[1].pt3dadd(-14.73, -120.15, -8.91, 0.61)
        dend[1].pt3dadd(-14.49, -120.65, -8.91, 0.61)
        dend[1].pt3dadd(-13.53, -121.89, -8.91, 0.61)
        dend[1].pt3dadd(-13.53, -122.14, -8.91, 0.61)
        dend[1].pt3dadd(-11.11, -123.13, -8.91, 0.61)
        dend[1].pt3dadd(-10.63, -123.38, -8.91, 0.61)
        dend[1].pt3dadd(-11.11, -126.62, -8.91, 0.61)
        dend[1].pt3dadd(-11.11, -127.36, -8.91, 0.61)
        dend[1].pt3dadd(-11.11, -130.10, -8.91, 0.61)
        dend[1].pt3dadd(-11.35, -130.60, -8.91, 0.61)
        dend[1].pt3dadd(-11.84, -134.08, -8.91, 0.43)
        dend[1].pt3dadd(-12.32, -134.58, -8.91, 0.43)
        dend[1].pt3dadd(-13.53, -137.06, -8.91, 0.43)
        dend[1].pt3dadd(-14.01, -137.81, -8.91, 0.43)
        dend[1].pt3dadd(-15.22, -139.30, -8.91, 0.43)
        dend[1].pt3dadd(-15.46, -139.55, -8.91, 0.43)
        dend[1].pt3dadd(-16.43, -142.04, -11.74, 0.43)
        dend[1].pt3dadd(-16.43, -144.03, -11.74, 0.43)
        dend[1].pt3dadd(-16.18, -144.78, -11.74, 0.43)
        dend[1].pt3dadd(-14.49, -146.52, -11.74, 0.43)
        dend[1].pt3dadd(-14.25, -146.77, -11.74, 0.43)
        dend[1].pt3dadd(-13.04, -148.76, -11.74, 0.43)
        dend[1].pt3dadd(-13.04, -149.01, -11.74, 0.43)
        dend[1].pt3dadd(-12.56, -155.22, -12.70, 0.43)
        dend[1].pt3dadd(-12.56, -154.98, -12.70, 0.43)
        dend[1].pt3dadd(-15.94, -158.21, -12.70, 0.43)
        dend[1].pt3dadd(-17.15, -159.20, -12.70, 0.43)
        dend[1].pt3dadd(-18.36, -159.20, -12.70, 0.43)
        dend[1].pt3dadd(-19.57, -158.96, -12.70, 0.43)
        dend[1].pt3dadd(-21.01, -158.21, -12.70, 0.43)
        dend[1].pt3dadd(-21.74, -155.22, -12.70, 0.43)
        dend[1].pt3dadd(-22.22, -154.98, -12.70, 0.43)
        dend[1].pt3dadd(-23.67, -154.23, -12.70, 0.43)
        dend[1].pt3dadd(-23.91, -154.23, -12.70, 0.43)
        dend[1].pt3dadd(-25.60, -153.48, -12.70, 0.43)
        dend[1].pt3dadd(-26.81, -152.99, -12.70, 0.43)
        dend[1].pt3dadd(-31.16, -151.74, -12.70, 0.43)
        dend[1].pt3dadd(-32.13, -150.00, -12.70, 0.43)
        dend[1].pt3dadd(-32.13, -149.25, -12.70, 0.43)
        dend[1].pt3dadd(-32.85, -148.01, -12.70, 0.43)
        dend[1].pt3dadd(-33.33, -148.01, -12.70, 0.43)
        dend[1].pt3dadd(-34.30, -146.27, -12.70, 0.43)

        dend[2] = h.Section(name='dend[2]')
        dend[2].connect(        dend[0](1), 0)
        dend[2].nseg = 2
        dend[2].pt3dclear()
        dend[2].pt3dadd(-0.72, -24.13, 0.00, 1.70)
        dend[2].pt3dadd(1.45, -26.12, -11.02, 1.70)
        dend[2].pt3dadd(1.45, -26.37, 3.48, 1.70)

        dend[3] = h.Section(name='dend[3]')
        dend[3].connect(        dend[1](1), 0)
        dend[3].nseg = 2
        dend[3].pt3dclear()
        dend[3].pt3dadd(1.45, -26.37, 3.48, 1.70)
        dend[3].pt3dadd(1.45, -28.36, -11.02, 1.70)
        dend[3].pt3dadd(1.45, -30.10, -11.02, 1.94)

        dend[4] = h.Section(name='dend[4]')
        dend[4].connect(        dend[3](1), 0)
        dend[4].nseg = 3
        dend[4].pt3dclear()
        dend[4].pt3dadd(1.45, -30.10, -11.02, 0.72)
        dend[4].pt3dadd(2.90, -32.09, -10.77, 0.72)
        dend[4].pt3dadd(3.38, -35.32, -10.77, 0.72)
        dend[4].pt3dadd(4.59, -38.06, -10.77, 0.72)
        dend[4].pt3dadd(4.83, -38.06, -10.77, 0.72)
        dend[4].pt3dadd(7.49, -40.30, -10.77, 0.72)
        dend[4].pt3dadd(7.73, -40.30, -10.77, 0.72)
        dend[4].pt3dadd(7.73, -41.79, -10.77, 0.72)
        dend[4].pt3dadd(11.35, -44.28, -10.77, 0.72)
        dend[4].pt3dadd(12.56, -46.02, -10.77, 0.72)
        dend[4].pt3dadd(12.80, -46.27, -10.77, 0.72)
        dend[4].pt3dadd(13.77, -48.26, -10.77, 0.72)

        dend[5] = h.Section(name='dend[5]')
        dend[5].connect(        dend[4](1), 0)
        dend[5].nseg = 16
        dend[5].pt3dclear()
        dend[5].pt3dadd(13.77, -48.26, -10.77, 0.43)
        dend[5].pt3dadd(12.80, -50.00, -5.03, 0.43)
        dend[5].pt3dadd(12.32, -52.49, -5.03, 0.43)
        dend[5].pt3dadd(12.08, -52.74, -5.03, 0.43)
        dend[5].pt3dadd(11.35, -55.97, -5.03, 0.43)
        dend[5].pt3dadd(11.35, -56.22, -5.03, 0.43)
        dend[5].pt3dadd(11.11, -58.96, -5.03, 0.43)
        dend[5].pt3dadd(11.11, -59.45, -5.03, 0.43)
        dend[5].pt3dadd(10.63, -62.94, -5.03, 0.43)
        dend[5].pt3dadd(10.14, -63.93, -5.03, 0.43)
        dend[5].pt3dadd(9.90, -65.67, -5.03, 0.43)
        dend[5].pt3dadd(9.66, -65.92, -5.03, 0.43)
        dend[5].pt3dadd(8.70, -67.66, -5.03, 0.43)
        dend[5].pt3dadd(8.45, -68.16, -5.03, 0.43)
        dend[5].pt3dadd(8.21, -69.40, -5.03, 0.43)
        dend[5].pt3dadd(8.45, -69.65, -5.03, 0.43)
        dend[5].pt3dadd(9.18, -71.14, -3.65, 0.43)
        dend[5].pt3dadd(10.63, -71.89, -3.65, 0.43)
        dend[5].pt3dadd(11.11, -71.89, -3.65, 0.43)
        dend[5].pt3dadd(11.35, -72.89, -3.65, 0.43)
        dend[5].pt3dadd(11.35, -73.13, -3.65, 0.43)
        dend[5].pt3dadd(10.39, -74.88, -3.65, 0.43)
        dend[5].pt3dadd(10.39, -75.12, -3.65, 0.43)
        dend[5].pt3dadd(9.66, -76.87, -3.65, 0.43)
        dend[5].pt3dadd(9.42, -78.11, -3.65, 0.43)
        dend[5].pt3dadd(9.42, -80.35, -3.65, 0.43)
        dend[5].pt3dadd(9.42, -80.60, -3.65, 0.43)
        dend[5].pt3dadd(9.18, -83.58, -3.65, 0.43)
        dend[5].pt3dadd(9.18, -84.83, -3.65, 0.43)
        dend[5].pt3dadd(8.94, -85.32, -3.65, 0.43)
        dend[5].pt3dadd(8.21, -87.56, -3.66, 0.43)
        dend[5].pt3dadd(8.21, -88.06, -3.66, 0.43)
        dend[5].pt3dadd(7.00, -90.30, -3.66, 0.43)
        dend[5].pt3dadd(7.00, -90.80, -3.66, 0.43)
        dend[5].pt3dadd(6.04, -93.28, -3.66, 0.43)
        dend[5].pt3dadd(5.80, -93.53, -3.66, 0.43)
        dend[5].pt3dadd(5.31, -94.78, -3.66, 0.43)
        dend[5].pt3dadd(5.31, -95.02, -3.66, 0.43)
        dend[5].pt3dadd(4.59, -96.02, -3.66, 0.43)
        dend[5].pt3dadd(4.35, -96.02, -3.66, 0.43)
        dend[5].pt3dadd(3.14, -97.02, -3.66, 0.43)
        dend[5].pt3dadd(2.90, -97.26, -3.66, 0.43)
        dend[5].pt3dadd(2.42, -98.76, -0.84, 0.43)
        dend[5].pt3dadd(2.90, -101.49, -0.84, 0.43)
        dend[5].pt3dadd(2.90, -103.48, -0.84, 0.43)
        dend[5].pt3dadd(1.21, -106.47, 4.44, 0.43)
        dend[5].pt3dadd(0.00, -107.96, 4.44, 0.43)
        dend[5].pt3dadd(-0.48, -108.21, 4.44, 0.43)
        dend[5].pt3dadd(-2.17, -109.70, 4.44, 0.43)
        dend[5].pt3dadd(-2.17, -109.95, 4.44, 0.43)
        dend[5].pt3dadd(-0.72, -111.19, 4.43, 0.43)
        dend[5].pt3dadd(-0.72, -111.44, 4.43, 0.43)
        dend[5].pt3dadd(0.00, -112.44, 4.43, 0.43)
        dend[5].pt3dadd(0.00, -112.94, 4.43, 0.43)
        dend[5].pt3dadd(0.00, -114.43, 4.43, 0.43)
        dend[5].pt3dadd(-3.14, -115.17, 11.54, 0.43)
        dend[5].pt3dadd(-7.73, -115.42, 11.53, 0.43)
        dend[5].pt3dadd(-9.18, -115.92, 11.49, 0.43)
        dend[5].pt3dadd(-9.42, -115.92, 11.48, 0.43)
        dend[5].pt3dadd(-11.84, -115.67, 11.37, 0.43)
        dend[5].pt3dadd(-11.84, -115.67, 11.36, 0.43)
        dend[5].pt3dadd(-14.98, -113.18, 11.18, 0.43)
        dend[5].pt3dadd(-15.22, -113.18, 11.15, 0.43)
        dend[5].pt3dadd(-15.22, -112.94, 7.28, 0.43)
        dend[5].pt3dadd(-19.81, -110.70, 7.28, 0.43)
        dend[5].pt3dadd(-21.98, -109.20, 7.28, 0.43)
        dend[5].pt3dadd(-24.64, -108.46, 7.28, 0.43)        

        dend[6] = h.Section(name='dend[6]')
        dend[6].connect(        dend[4](1), 0)
        dend[6].nseg = 12
        dend[6].pt3dclear()
        dend[6].pt3dadd(13.77, -48.26, -10.77, 0.43)
        dend[6].pt3dadd(15.94, -50.75, 2.15, 0.43)
        dend[6].pt3dadd(15.94, -50.50, 2.15, 0.43)
        dend[6].pt3dadd(16.91, -52.24, 2.15, 0.43)
        dend[6].pt3dadd(15.46, -54.23, 2.15, 0.43)
        dend[6].pt3dadd(15.46, -54.98, 2.15, 0.43)
        dend[6].pt3dadd(15.46, -56.97, 2.15, 0.43)
        dend[6].pt3dadd(15.70, -57.96, 2.15, 0.43)
        dend[6].pt3dadd(15.70, -58.96, 2.15, 0.43)
        dend[6].pt3dadd(15.70, -59.20, 2.15, 0.43)
        dend[6].pt3dadd(14.98, -61.19, 2.15, 0.43)
        dend[6].pt3dadd(14.25, -61.44, 2.15, 0.43)
        dend[6].pt3dadd(13.77, -62.19, 2.15, 0.43)
        dend[6].pt3dadd(13.53, -63.18, 2.15, 0.43)
        dend[6].pt3dadd(13.53, -64.43, 2.15, 0.43)
        dend[6].pt3dadd(13.77, -66.17, 2.15, 0.43)
        dend[6].pt3dadd(14.25, -67.91, 2.15, 0.43)
        dend[6].pt3dadd(15.22, -68.66, 2.15, 0.43)
        dend[6].pt3dadd(16.91, -69.65, 2.15, 0.43)
        dend[6].pt3dadd(17.39, -70.40, 2.15, 0.43)
        dend[6].pt3dadd(17.87, -71.39, 2.15, 0.43)
        dend[6].pt3dadd(17.87, -71.89, 2.15, 0.43)
        dend[6].pt3dadd(17.87, -72.89, 2.15, 0.43)
        dend[6].pt3dadd(17.87, -73.38, 2.15, 0.43)
        dend[6].pt3dadd(17.87, -74.13, 2.15, 0.43)
        dend[6].pt3dadd(17.87, -75.12, 2.15, 0.43)
        dend[6].pt3dadd(17.87, -75.87, 2.15, 0.43)
        dend[6].pt3dadd(17.63, -76.37, 2.15, 0.43)
        dend[6].pt3dadd(17.63, -77.36, 2.15, 0.43)
        dend[6].pt3dadd(17.63, -77.61, 2.15, 0.43)
        dend[6].pt3dadd(17.87, -79.35, 2.15, 0.43)
        dend[6].pt3dadd(18.36, -80.85, 2.15, 0.43)
        dend[6].pt3dadd(18.60, -81.59, 2.15, 0.43)
        dend[6].pt3dadd(22.46, -83.83, 6.67, 0.43)
        dend[6].pt3dadd(23.91, -86.32, 7.41, 0.43)
        dend[6].pt3dadd(24.64, -88.06, 10.01, 0.43)
        dend[6].pt3dadd(26.81, -89.05, 10.01, 0.43)
        dend[6].pt3dadd(26.81, -89.05, 10.00, 0.43)
        dend[6].pt3dadd(28.26, -91.04, 10.00, 0.43)
        dend[6].pt3dadd(29.71, -92.54, 10.00, 0.43)
        dend[6].pt3dadd(30.19, -92.54, 10.00, 0.43)
        dend[6].pt3dadd(31.16, -92.79, 10.00, 0.43)
        dend[6].pt3dadd(31.40, -93.03, 10.00, 0.43)
        dend[6].pt3dadd(32.13, -93.28, 10.00, 0.43)
        dend[6].pt3dadd(32.61, -94.28, 10.00, 0.43)
        dend[6].pt3dadd(34.30, -97.26, 10.00, 0.43)
        dend[6].pt3dadd(34.30, -97.51, 10.00, 0.43)
        dend[6].pt3dadd(35.51, -98.26, 10.00, 0.43)
        dend[6].pt3dadd(35.99, -98.51, 10.00, 0.43)
        dend[6].pt3dadd(38.89, -99.25, 10.00, 0.43)
        dend[6].pt3dadd(39.37, -99.50, 10.00, 0.43)
        dend[6].pt3dadd(41.55, -101.99, 10.40, 0.43)
        dend[6].pt3dadd(44.20, -101.74, 10.40, 0.43)
        dend[6].pt3dadd(44.44, -101.74, 10.40, 0.43)
        dend[6].pt3dadd(45.41, -101.00, 10.40, 0.43)
        dend[6].pt3dadd(46.14, -100.00, 10.40, 0.43)

        dend[7] = h.Section(name='dend[7]')
        dend[7].connect(        dend[6](1), 0)
        dend[7].nseg = 12
        dend[7].pt3dclear()
        dend[7].pt3dadd(1.45, -30.10, -11.02, 0.61)
        dend[7].pt3dadd(0.00, -32.34, -14.30, 0.61)
        dend[7].pt3dadd(-1.93, -33.08, -14.31, 0.61)
        dend[7].pt3dadd(-2.17, -33.08, -14.31, 0.61)
        dend[7].pt3dadd(-3.14, -34.08, -14.31, 0.61)
        dend[7].pt3dadd(-3.86, -36.32, -14.31, 0.61)
        dend[7].pt3dadd(-3.86, -36.82, -14.31, 0.61)
        dend[7].pt3dadd(-3.86, -38.06, -14.31, 0.61)
        dend[7].pt3dadd(-3.62, -38.81, -14.31, 0.61)
        dend[7].pt3dadd(-3.38, -41.04, -14.31, 0.61)
        dend[7].pt3dadd(-3.38, -41.29, -14.31, 0.61)
        dend[7].pt3dadd(-3.14, -42.29, -14.31, 0.61)
        dend[7].pt3dadd(-2.90, -43.03, -14.31, 0.61)
        dend[7].pt3dadd(-2.66, -44.53, -14.31, 0.61)
        dend[7].pt3dadd(-2.66, -45.27, -14.31, 0.61)
        dend[7].pt3dadd(-2.66, -46.77, -14.31, 0.61)
        dend[7].pt3dadd(-2.66, -47.01, -14.31, 0.61)
        dend[7].pt3dadd(-4.59, -50.75, -15.66, 0.61)
        dend[7].pt3dadd(-6.04, -53.48, -15.66, 0.61)
        dend[7].pt3dadd(-6.04, -53.73, -15.66, 0.61)
        dend[7].pt3dadd(-6.04, -55.22, -15.66, 0.61)
        dend[7].pt3dadd(-6.04, -55.47, -15.66, 0.61)
        dend[7].pt3dadd(-6.04, -58.21, -15.66, 0.61)
        dend[7].pt3dadd(-6.04, -59.45, -15.66, 0.61)
        dend[7].pt3dadd(-7.00, -61.44, -15.66, 0.61)
        dend[7].pt3dadd(-7.25, -61.69, -15.66, 0.61)
        dend[7].pt3dadd(-7.25, -63.18, -15.66, 0.61)
        dend[7].pt3dadd(-7.00, -64.68, -15.66, 0.61)
        dend[7].pt3dadd(-6.52, -65.92, -15.66, 0.61)
        dend[7].pt3dadd(-6.52, -66.17, -15.66, 0.61)
        dend[7].pt3dadd(-7.97, -67.41, -15.67, 0.61)
        dend[7].pt3dadd(-9.66, -68.91, -15.67, 0.61)
        dend[7].pt3dadd(-9.90, -69.65, -15.67, 0.61)
        dend[7].pt3dadd(-10.39, -72.14, -15.67, 0.61)
        dend[7].pt3dadd(-10.63, -72.64, -15.67, 0.61)
        dend[7].pt3dadd(-11.35, -74.38, -15.67, 0.61)
        dend[7].pt3dadd(-12.80, -72.89, -15.67, 0.61)
        dend[7].pt3dadd(-13.77, -72.39, -15.67, 0.61)
        dend[7].pt3dadd(-14.25, -72.39, -15.67, 0.61)
        dend[7].pt3dadd(-16.67, -72.89, -15.67, 0.61)
        dend[7].pt3dadd(-17.15, -73.38, -15.67, 0.61)
        dend[7].pt3dadd(-21.01, -78.11, -14.54, 0.61)
        dend[7].pt3dadd(-23.91, -78.36, -14.54, 0.61)
        dend[7].pt3dadd(-24.40, -78.36, -14.54, 0.61)
        dend[7].pt3dadd(-27.78, -79.85, -14.54, 0.61)
        dend[7].pt3dadd(-28.26, -80.10, -14.54, 0.61)
        dend[7].pt3dadd(-30.68, -81.34, -14.54, 0.61)
        dend[7].pt3dadd(-30.92, -81.59, -14.54, 0.61)
        dend[7].pt3dadd(-30.92, -82.09, -14.54, 0.61)
        dend[7].pt3dadd(-30.92, -82.59, -14.54, 0.61)
        dend[7].pt3dadd(-30.19, -85.57, -13.57, 0.61)
        dend[7].pt3dadd(-29.95, -85.57, -13.57, 0.61)
        dend[7].pt3dadd(-31.16, -88.56, -13.57, 0.61)
        dend[7].pt3dadd(-32.61, -88.81, -13.57, 0.61)
        dend[7].pt3dadd(-33.33, -90.55, -13.57, 0.61)
        dend[7].pt3dadd(-33.33, -91.04, -13.57, 0.61)
        dend[7].pt3dadd(-32.13, -91.54, -13.57, 0.61)
        dend[7].pt3dadd(-31.88, -92.04, -13.57, 0.61)
        dend[7].pt3dadd(-31.40, -94.53, -13.57, 0.61)
        dend[7].pt3dadd(-31.40, -95.27, -13.57, 0.61)
        dend[7].pt3dadd(-33.09, -96.27, -13.57, 0.61)
        dend[7].pt3dadd(-34.30, -96.52, -13.57, 0.61)
        dend[7].pt3dadd(-35.75, -96.77, -13.57, 0.61)
        dend[7].pt3dadd(-36.96, -97.26, -13.57, 0.61)
        dend[7].pt3dadd(-37.92, -97.76, -13.57, 0.61)
        dend[7].pt3dadd(-39.13, -98.01, -13.57, 0.61)
        dend[7].pt3dadd(-40.82, -98.01, -13.57, 0.61)

        dend[8] = h.Section(name='dend[8]')
        dend[8].connect(        dend[7](1), 0)
        dend[8].nseg = 17
        dend[8].pt3dclear()
        dend[8].pt3dadd(-40.82, -98.01, -13.57, 0.43)
        dend[8].pt3dadd(-43.00, -100.50, -13.58, 0.43)
        dend[8].pt3dadd(-44.69, -101.49, -13.58, 0.43)
        dend[8].pt3dadd(-45.17, -101.99, -13.58, 0.43)
        dend[8].pt3dadd(-45.89, -102.99, -13.58, 0.43)
        dend[8].pt3dadd(-45.89, -103.48, -13.58, 0.43)
        dend[8].pt3dadd(-46.14, -104.73, -13.58, 0.43)
        dend[8].pt3dadd(-46.62, -105.72, -13.58, 0.43)
        dend[8].pt3dadd(-46.86, -107.71, -13.58, 0.43)
        dend[8].pt3dadd(-46.86, -107.96, -13.58, 0.43)
        dend[8].pt3dadd(-48.07, -109.20, -13.58, 0.43)
        dend[8].pt3dadd(-49.03, -109.70, -13.58, 0.43)
        dend[8].pt3dadd(-52.17, -110.70, -13.58, 0.43)
        dend[8].pt3dadd(-52.42, -110.95, -13.58, 0.43)
        dend[8].pt3dadd(-55.56, -112.44, -10.17, 0.43)
        dend[8].pt3dadd(-57.97, -112.44, -10.17, 0.43)
        dend[8].pt3dadd(-58.21, -112.44, -10.17, 0.43)
        dend[8].pt3dadd(-60.14, -112.44, -10.17, 0.43)
        dend[8].pt3dadd(-60.63, -112.44, -10.17, 0.43)
        dend[8].pt3dadd(-62.80, -112.69, -10.17, 0.43)
        dend[8].pt3dadd(-64.49, -113.43, -10.17, 0.43)
        dend[8].pt3dadd(-64.98, -113.68, -10.17, 0.43)
        dend[8].pt3dadd(-66.18, -114.43, -10.17, 0.43)
        dend[8].pt3dadd(-66.67, -114.93, -10.17, 0.43)
        dend[8].pt3dadd(-66.91, -115.42, -10.17, 0.43)
        dend[8].pt3dadd(-66.91, -116.92, -10.17, 0.43)
        dend[8].pt3dadd(-67.15, -117.41, -10.17, 0.43)
        dend[8].pt3dadd(-68.12, -118.41, -10.17, 0.43)
        dend[8].pt3dadd(-68.12, -118.66, -10.17, 0.43)
        dend[8].pt3dadd(-69.32, -120.15, -10.18, 0.43)
        dend[8].pt3dadd(-70.05, -122.39, -15.90, 0.43)
        dend[8].pt3dadd(-72.46, -122.39, -15.90, 0.43)
        dend[8].pt3dadd(-72.71, -122.39, -15.90, 0.43)
        dend[8].pt3dadd(-74.88, -122.14, -15.96, 0.43)
        dend[8].pt3dadd(-74.88, -122.14, -16.04, 0.43)
        dend[8].pt3dadd(-76.33, -119.90, -16.39, 0.43)
        dend[8].pt3dadd(-76.33, -119.90, -16.41, 0.43)
        dend[8].pt3dadd(-78.26, -118.91, -16.73, 0.43)
        dend[8].pt3dadd(-80.19, -116.17, -19.00, 0.43)
        dend[8].pt3dadd(-80.19, -116.17, -18.98, 0.43)
        dend[8].pt3dadd(-81.40, -115.42, -18.97, 0.43)
        dend[8].pt3dadd(-81.64, -115.42, -18.96, 0.43)
        dend[8].pt3dadd(-84.54, -114.93, -18.96, 0.43)
        dend[8].pt3dadd(-85.51, -115.67, -18.96, 0.43)
        dend[8].pt3dadd(-85.75, -116.17, -18.96, 0.43)
        dend[8].pt3dadd(-85.99, -118.41, -18.96, 0.43)
        dend[8].pt3dadd(-85.99, -119.15, -18.96, 0.43)
        dend[8].pt3dadd(-86.47, -120.15, -18.96, 0.43)
        dend[8].pt3dadd(-86.72, -120.65, -18.96, 0.43)
        dend[8].pt3dadd(-86.72, -121.89, -18.96, 0.43)
        dend[8].pt3dadd(-86.72, -122.39, -18.96, 0.43)
        dend[8].pt3dadd(-86.47, -123.63, -18.96, 0.43)
        dend[8].pt3dadd(-85.75, -124.38, -18.96, 0.43)
        dend[8].pt3dadd(-85.02, -124.88, -18.96, 0.43)
        dend[8].pt3dadd(-84.78, -125.12, -18.96, 0.43)
        dend[8].pt3dadd(-84.78, -125.62, -18.96, 0.43)
        dend[8].pt3dadd(-85.27, -126.87, -18.96, 0.43)
        dend[8].pt3dadd(-85.99, -127.61, -18.96, 0.43)
        dend[8].pt3dadd(-86.96, -128.11, -18.96, 0.43)
        dend[8].pt3dadd(-87.92, -128.86, -18.96, 0.43)
        dend[8].pt3dadd(-88.65, -129.35, -18.96, 0.43)
        dend[8].pt3dadd(-88.89, -129.60, -18.96, 0.43)
        dend[8].pt3dadd(-88.89, -130.35, -18.96, 0.43)
        dend[8].pt3dadd(-88.89, -131.59, -18.96, 0.43)
        dend[8].pt3dadd(-88.65, -132.59, -18.96, 0.43)
        dend[8].pt3dadd(-87.68, -134.08, -19.91, 0.43)
        dend[8].pt3dadd(-88.89, -136.32, -19.91, 0.43)
        dend[8].pt3dadd(-89.61, -137.06, -19.91, 0.43)
        dend[8].pt3dadd(-90.82, -138.06, -19.91, 0.43)
        dend[8].pt3dadd(-91.79, -138.56, -19.91, 0.43)
        dend[8].pt3dadd(-92.03, -139.80, -19.91, 0.43)
        dend[8].pt3dadd(-92.27, -141.04, -19.91, 0.43)
        dend[8].pt3dadd(-92.27, -142.29, -19.91, 0.43)
        dend[8].pt3dadd(-92.51, -144.03, -19.91, 0.43)
        dend[8].pt3dadd(-92.75, -149.25, -16.44, 0.43)
        dend[8].pt3dadd(-92.51, -149.50, -16.44, 0.43)
        dend[8].pt3dadd(-91.55, -153.73, -16.44, 0.43)
        dend[8].pt3dadd(-91.06, -153.98, -16.44, 0.43)
        dend[8].pt3dadd(-90.34, -154.23, -16.44, 0.43)
        dend[8].pt3dadd(-89.37, -154.73, -16.44, 0.43)
        dend[8].pt3dadd(-88.16, -154.73, -16.44, 0.43)
        dend[8].pt3dadd(-87.68, -154.98, -16.44, 0.43)
        dend[8].pt3dadd(-86.47, -155.72, -16.44, 0.43)
        dend[8].pt3dadd(-85.51, -156.72, -16.44, 0.43)
        dend[8].pt3dadd(-84.06, -157.96, -16.44, 0.43)

        dend[9] = h.Section(name='dend[9]')
        dend[9].connect(        dend[7](1), 0)
        dend[9].nseg = 9
        dend[9].pt3dclear()
        dend[9].pt3dadd(-40.82, -98.01, -13.57, 0.43)
        dend[9].pt3dadd(-42.75, -97.02, -13.57, 0.43)
        dend[9].pt3dadd(-44.44, -96.77, -13.57, 0.43)
        dend[9].pt3dadd(-47.58, -97.51, -13.57, 0.43)
        dend[9].pt3dadd(-47.83, -99.01, -13.57, 0.43)
        dend[9].pt3dadd(-48.31, -100.00, -13.57, 0.43)
        dend[9].pt3dadd(-48.79, -100.00, -13.57, 0.43)
        dend[9].pt3dadd(-50.24, -100.75, -13.57, 0.43)
        dend[9].pt3dadd(-50.72, -101.00, -13.57, 0.43)
        dend[9].pt3dadd(-51.69, -101.00, -13.57, 0.43)
        dend[9].pt3dadd(-52.17, -101.00, -13.57, 0.43)
        dend[9].pt3dadd(-55.56, -100.00, -16.84, 0.43)
        dend[9].pt3dadd(-56.28, -100.00, -16.84, 0.43)
        dend[9].pt3dadd(-57.97, -100.25, -16.84, 0.43)
        dend[9].pt3dadd(-58.70, -101.00, -16.84, 0.43)
        dend[9].pt3dadd(-60.14, -101.99, -16.84, 0.43)
        dend[9].pt3dadd(-61.59, -101.99, -16.84, 0.43)
        dend[9].pt3dadd(-62.56, -101.99, -16.84, 0.43)
        dend[9].pt3dadd(-63.04, -101.99, -16.84, 0.43)
        dend[9].pt3dadd(-65.70, -102.24, -16.84, 0.43)
        dend[9].pt3dadd(-66.67, -102.74, -16.84, 0.43)
        dend[9].pt3dadd(-67.15, -103.98, -16.84, 0.43)
        dend[9].pt3dadd(-67.39, -104.23, -16.84, 0.43)
        dend[9].pt3dadd(-68.12, -104.98, -16.84, 0.43)
        dend[9].pt3dadd(-68.84, -105.72, -16.84, 0.43)
        dend[9].pt3dadd(-69.81, -105.97, -16.84, 0.43)
        dend[9].pt3dadd(-70.77, -105.97, -16.84, 0.43)
        dend[9].pt3dadd(-72.46, -106.22, -16.84, 0.43)
        dend[9].pt3dadd(-73.19, -106.72, -16.84, 0.43)
        dend[9].pt3dadd(-74.15, -107.46, -16.84, 0.43)
        dend[9].pt3dadd(-74.40, -107.96, -16.84, 0.43)
        dend[9].pt3dadd(-75.12, -109.45, -16.84, 0.43)
        dend[9].pt3dadd(-75.60, -109.95, -16.84, 0.43)
        dend[9].pt3dadd(-75.85, -110.70, -16.84, 0.43)
        dend[9].pt3dadd(-76.57, -110.95, -16.84, 0.43)
        dend[9].pt3dadd(-76.81, -110.95, -16.84, 0.43)
        dend[9].pt3dadd(-76.81, -111.19, -16.84, 0.43)
        dend[9].pt3dadd(-77.54, -111.19, -16.84, 0.43)
        dend[9].pt3dadd(-77.78, -111.44, -16.84, 0.43)
        dend[9].pt3dadd(-79.71, -111.94, -16.84, 0.43)
        dend[9].pt3dadd(-80.68, -117.16, -19.96, 0.43)
        dend[9].pt3dadd(-80.92, -117.16, -19.96, 0.43)
        dend[9].pt3dadd(-82.61, -118.66, -19.96, 0.43)
        dend[9].pt3dadd(-82.61, -118.91, -19.96, 0.43)
        dend[9].pt3dadd(-82.61, -120.15, -19.96, 0.43)
        dend[9].pt3dadd(-82.61, -121.14, -19.96, 0.43)
        dend[9].pt3dadd(-82.37, -122.89, -19.96, 0.43)
        dend[9].pt3dadd(-82.37, -123.38, -19.96, 0.43)
        dend[9].pt3dadd(-82.85, -124.13, -19.96, 0.43)
        dend[9].pt3dadd(-83.09, -124.38, -19.96, 0.43)

        dend[10] = h.Section(name='dend[10]')
        dend[10].connect(        dend[3](1), 0)
        dend[10].nseg = 6
        dend[10].pt3dclear()
        dend[10].pt3dadd(1.45, -26.37, 3.48, 0.48)
        dend[10].pt3dadd(3.38, -26.37, 3.48, 0.48)
        dend[10].pt3dadd(6.04, -31.59, 1.77, 0.48)
        dend[10].pt3dadd(7.97, -33.08, 1.78, 0.48)
        dend[10].pt3dadd(8.45, -33.33, 1.78, 0.48)
        dend[10].pt3dadd(9.90, -34.33, 1.78, 0.48)
        dend[10].pt3dadd(10.14, -34.33, 1.78, 0.48)
        dend[10].pt3dadd(14.25, -36.57, 1.78, 0.48)
        dend[10].pt3dadd(14.25, -36.82, 1.78, 0.48)
        dend[10].pt3dadd(17.15, -38.06, 1.78, 0.48)
        dend[10].pt3dadd(17.15, -38.31, 1.78, 0.48)
        dend[10].pt3dadd(23.43, -41.29, 2.53, 0.48)
        dend[10].pt3dadd(24.88, -44.53, 2.53, 0.48)
        dend[10].pt3dadd(25.12, -44.78, 2.53, 0.48)
        dend[10].pt3dadd(29.47, -48.26, -2.69, 0.48)
        dend[10].pt3dadd(30.19, -49.75, -2.69, 0.48)
        dend[10].pt3dadd(30.43, -49.75, -2.69, 0.48)

        dend[11] = h.Section(name='dend[11]')
        dend[11].connect(        dend[10](1), 0)
        dend[11].nseg = 7
        dend[11].pt3dclear()
        dend[11].pt3dadd(30.43, -49.75, -2.69, 0.48)
        dend[11].pt3dadd(37.20, -53.98, -2.69, 0.48)
        dend[11].pt3dadd(38.16, -57.21, -2.69, 0.48)
        dend[11].pt3dadd(38.41, -57.46, -2.69, 0.48)
        dend[11].pt3dadd(41.06, -60.70, -2.69, 0.48)
        dend[11].pt3dadd(41.06, -61.19, -2.69, 0.48)
        dend[11].pt3dadd(45.89, -64.93, -8.15, 0.48)
        dend[11].pt3dadd(49.28, -67.91, -8.15, 0.48)
        dend[11].pt3dadd(49.52, -67.91, -8.15, 0.48)
        dend[11].pt3dadd(53.86, -71.39, -8.41, 0.48)
        dend[11].pt3dadd(54.83, -74.88, -8.41, 0.48)
        dend[11].pt3dadd(54.83, -75.12, -8.41, 0.48)
        dend[11].pt3dadd(56.28, -76.62, -8.41, 0.48)
        dend[11].pt3dadd(57.00, -77.36, -8.41, 0.48)
        dend[11].pt3dadd(57.97, -79.35, -8.41, 0.48)
        dend[11].pt3dadd(58.21, -79.35, -8.41, 0.48)
        dend[11].pt3dadd(60.14, -82.59, -8.41, 0.48)

        dend[12] = h.Section(name='dend[12]')
        dend[12].connect(        dend[11](1), 0)
        dend[12].nseg = 7
        dend[12].pt3dclear()
        dend[12].pt3dadd(60.14, -82.59, -8.41, 0.43)
        dend[12].pt3dadd(61.84, -83.33, -15.23, 0.43)
        dend[12].pt3dadd(64.25, -82.59, -15.23, 0.43)
        dend[12].pt3dadd(66.91, -81.84, -15.23, 0.43)
        dend[12].pt3dadd(67.39, -82.09, -15.23, 0.43)
        dend[12].pt3dadd(68.12, -83.83, -15.23, 0.43)
        dend[12].pt3dadd(68.36, -84.08, -15.23, 0.43)
        dend[12].pt3dadd(70.77, -86.57, -15.23, 0.43)
        dend[12].pt3dadd(71.26, -86.57, -15.23, 0.43)
        dend[12].pt3dadd(72.95, -85.57, -15.23, 0.43)
        dend[12].pt3dadd(73.43, -85.57, -15.23, 0.43)
        dend[12].pt3dadd(78.50, -85.57, -15.23, 0.43)
        dend[12].pt3dadd(82.13, -85.57, -20.98, 0.43)
        dend[12].pt3dadd(83.57, -85.07, -20.98, 0.43)
        dend[12].pt3dadd(85.51, -85.07, -20.98, 0.43)
        dend[12].pt3dadd(85.75, -88.06, -20.97, 0.43)
        dend[12].pt3dadd(85.75, -88.56, -20.97, 0.43)
        dend[12].pt3dadd(89.37, -90.05, -20.97, 0.43)
        dend[12].pt3dadd(89.61, -89.80, -20.97, 0.43)
        dend[12].pt3dadd(91.55, -88.81, -20.97, 0.43)
        dend[12].pt3dadd(91.79, -88.81, -20.97, 0.43)

        dend[13] = h.Section(name='dend[13]')
        dend[13].connect(        dend[11](1), 0)
        dend[13].nseg = 11
        dend[13].pt3dclear()
        dend[13].pt3dadd(60.14, -82.59, -8.41, 0.43)
        dend[13].pt3dadd(59.18, -84.58, -8.41, 0.43)
        dend[13].pt3dadd(59.18, -84.83, -8.41, 0.43)
        dend[13].pt3dadd(59.90, -86.07, -8.41, 0.43)
        dend[13].pt3dadd(60.14, -86.32, -8.41, 0.43)
        dend[13].pt3dadd(62.80, -87.06, -8.41, 0.43)
        dend[13].pt3dadd(63.53, -87.06, -8.41, 0.43)
        dend[13].pt3dadd(64.73, -87.31, -8.41, 0.43)
        dend[13].pt3dadd(66.43, -88.06, -8.41, 0.43)
        dend[13].pt3dadd(68.36, -88.81, -8.41, 0.43)
        dend[13].pt3dadd(70.77, -89.80, -8.41, 0.43)
        dend[13].pt3dadd(75.12, -94.03, -8.41, 0.43)
        dend[13].pt3dadd(79.71, -98.26, -8.41, 0.43)
        dend[13].pt3dadd(79.95, -98.26, -8.41, 0.43)
        dend[13].pt3dadd(83.57, -102.24, -8.41, 0.43)
        dend[13].pt3dadd(83.57, -102.49, -8.41, 0.43)
        dend[13].pt3dadd(86.23, -108.21, -8.41, 0.43)
        dend[13].pt3dadd(88.41, -108.96, -8.41, 0.43)
        dend[13].pt3dadd(88.89, -109.20, -8.41, 0.43)
        dend[13].pt3dadd(89.37, -111.94, -8.41, 0.43)
        dend[13].pt3dadd(89.61, -112.19, -8.41, 0.43)
        dend[13].pt3dadd(89.86, -113.93, -8.41, 0.43)
        dend[13].pt3dadd(89.61, -115.42, -8.41, 0.43)
        dend[13].pt3dadd(88.89, -116.67, -8.41, 0.43)
        dend[13].pt3dadd(88.41, -117.41, -8.41, 0.43)
        dend[13].pt3dadd(90.10, -122.64, -13.92, 0.43)
        dend[13].pt3dadd(93.48, -124.13, -12.92, 0.43)
        dend[13].pt3dadd(93.72, -124.13, -12.92, 0.43)
        dend[13].pt3dadd(98.07, -124.63, -12.92, 0.43)
        dend[13].pt3dadd(98.55, -124.63, -12.92, 0.43)
        dend[13].pt3dadd(99.76, -124.63, -12.92, 0.43)
        dend[13].pt3dadd(100.00, -124.63, -12.92, 0.43)
        dend[13].pt3dadd(103.38, -124.38, -12.92, 0.43)
        dend[13].pt3dadd(103.86, -124.13, -12.92, 0.43)
        dend[13].pt3dadd(105.31, -123.38, -12.92, 0.43)

        dend[14] = h.Section(name='dend[14]')
        dend[14].connect(        dend[3](1), 0)
        dend[14].nseg = 6
        dend[14].pt3dclear()
        dend[14].pt3dadd(30.43, -49.75, -2.69, 0.43)
        dend[14].pt3dadd(29.47, -51.99, -11.83, 0.43)
        dend[14].pt3dadd(29.71, -55.72, -11.83, 0.43)
        dend[14].pt3dadd(29.71, -55.97, -11.83, 0.43)
        dend[14].pt3dadd(31.16, -57.71, -11.83, 0.43)
        dend[14].pt3dadd(31.16, -57.96, -11.83, 0.43)
        dend[14].pt3dadd(32.13, -60.20, -11.83, 0.43)
        dend[14].pt3dadd(34.06, -66.67, -15.57, 0.43)
        dend[14].pt3dadd(38.16, -67.41, -15.57, 0.43)
        dend[14].pt3dadd(38.41, -67.66, -15.57, 0.43)
        dend[14].pt3dadd(40.34, -72.89, -15.57, 0.43)
        dend[14].pt3dadd(40.82, -73.13, -15.57, 0.43)
        dend[14].pt3dadd(44.44, -74.38, -15.57, 0.43)
        dend[14].pt3dadd(46.62, -76.37, -15.57, 0.43)

        dend2 = [None]*31

        dend2[0] = h.Section(name='dend2[0]')
        dend2[0].connect(        soma(0.709232), 0)
        dend2[0].pt3dclear()
        dend2[0].pt3dadd(-5.80, -6.22, -0.01, 1.20)
        dend2[0].pt3dadd(-7.49, -6.97, -0.01, 1.20)
        dend2[0].pt3dadd(-10.14, -7.21, -0.01, 1.20)

        dend2[1] = h.Section(name='dend2[1]')
        dend2[1].connect(        dend2[0](1), 0)
        dend2[1].pt3dclear()
        dend2[1].pt3dadd(-10.14, -7.21, -0.01, 0.96)
        dend2[1].pt3dadd(-11.84, -5.22, -0.01, 0.96)
        dend2[1].pt3dadd(-15.22, -5.22, 4.55, 0.72)
        dend2[1].pt3dadd(-15.22, -5.22, 4.54, 0.72)
        dend2[1].pt3dadd(-15.94, -7.96, 4.54, 0.72)
        dend2[1].pt3dadd(-15.46, -9.95, 4.54, 0.72)
        dend2[1].pt3dadd(-15.22, -10.45, 4.54, 0.72)
        dend2[1].pt3dadd(-15.94, -13.18, 4.54, 0.72)
        dend2[1].pt3dadd(-16.43, -13.68, 4.54, 0.72)
        dend2[1].pt3dadd(-19.08, -13.43, 8.80, 0.72)
        dend2[1].pt3dadd(-20.29, -13.93, 8.80, 0.72)
        dend2[1].pt3dadd(-20.53, -14.18, 8.97, 0.72)
        dend2[1].pt3dadd(-21.01, -15.17, 9.04, 0.72)
        dend2[1].pt3dadd(-21.26, -15.42, 9.04, 0.72)
        dend2[1].pt3dadd(-27.54, -17.41, 14.93, 0.72)
        dend2[1].pt3dadd(-28.99, -19.15, 14.93, 0.72)
        dend2[1].pt3dadd(-29.23, -19.15, 14.93, 0.72)
        dend2[1].pt3dadd(-31.88, -19.40, 14.93, 0.72)
        dend2[1].pt3dadd(-32.85, -20.40, 14.93, 0.72)
        dend2[1].pt3dadd(-33.33, -20.90, 14.92, 0.72)
        dend2[1].pt3dadd(-33.57, -22.64, 14.92, 0.72)
        dend2[1].pt3dadd(-33.09, -26.12, 14.92, 0.72)
        dend2[1].pt3dadd(-33.09, -27.61, 14.92, 0.72)
        dend2[1].pt3dadd(-33.09, -27.86, 14.92, 0.72)
        dend2[1].pt3dadd(-33.09, -29.35, 14.92, 0.72)
        dend2[1].pt3dadd(-33.09, -29.60, 14.92, 0.72)
        dend2[1].pt3dadd(-32.37, -31.09, 14.92, 0.72)
        dend2[1].pt3dadd(-32.13, -31.09, 14.92, 0.72)

        dend2[2] = h.Section(name='dend2[2]')
        dend2[2].connect(        dend2[0](1), 0)
        dend2[2].pt3dclear()
        dend2[2].pt3dadd(-10.14, -7.21, -0.01, 1.20)
        dend2[2].pt3dadd(-9.90, -10.45, 0.22, 1.20)
        dend2[2].pt3dadd(-8.21, -11.19, 0.22, 1.20)

        dend2[3] = h.Section(name='dend2[3]')
        dend2[3].connect(        dend2[2](1), 0)
        dend2[3].pt3dclear()
        dend2[3].pt3dadd(-8.21, -11.19, 0.22, 0.77)
        dend2[3].pt3dadd(-9.42, -15.42, -2.21, 0.77)
        dend2[3].pt3dadd(-10.14, -17.41, -2.21, 0.77)
        dend2[3].pt3dadd(-10.14, -17.91, -2.21, 0.77)
        dend2[3].pt3dadd(-11.35, -19.15, -2.21, 0.77)
        dend2[3].pt3dadd(-11.84, -19.40, -2.21, 0.77)

        dend2[4] = h.Section(name='dend2[4]')
        dend2[4].connect(        dend2[3](1), 0)
        dend2[4].pt3dclear()
        dend2[4].pt3dadd(-11.84, -19.40, -2.21, 0.54)
        dend2[4].pt3dadd(-15.22, -19.65, -2.21, 0.54)
        dend2[4].pt3dadd(-15.46, -19.65, -2.21, 0.54)
        dend2[4].pt3dadd(-20.77, -18.66, -4.79, 0.54)
        dend2[4].pt3dadd(-22.46, -18.66, -4.79, 0.54)
        dend2[4].pt3dadd(-22.71, -18.91, -4.79, 0.54)
        dend2[4].pt3dadd(-24.15, -20.15, -4.79, 0.54)
        dend2[4].pt3dadd(-24.64, -20.65, -4.79, 0.54)
        dend2[4].pt3dadd(-26.33, -21.39, -4.79, 0.54)
        dend2[4].pt3dadd(-27.05, -21.64, -4.79, 0.54)
        dend2[4].pt3dadd(-28.02, -21.64, -4.79, 0.54)
        dend2[4].pt3dadd(-28.26, -21.64, -4.79, 0.54)

        dend2[5] = h.Section(name='dend2[5]')
        dend2[5].connect(        dend2[4](1), 0)
        dend2[5].pt3dclear()
        dend2[5].pt3dadd(-28.26, -21.64, -4.79, 0.43)
        dend2[5].pt3dadd(-32.13, -20.40, -8.10, 0.43)
        dend2[5].pt3dadd(-35.51, -18.91, -8.10, 0.43)
        dend2[5].pt3dadd(-38.65, -16.92, -8.10, 0.43)
        dend2[5].pt3dadd(-41.55, -15.92, -8.10, 0.43)
        dend2[5].pt3dadd(-44.69, -14.93, -8.10, 0.43)
        dend2[5].pt3dadd(-45.89, -14.93, -8.10, 0.43)
        dend2[5].pt3dadd(-47.83, -14.68, -5.22, 0.43)
        dend2[5].pt3dadd(-50.72, -15.17, -5.22, 0.43)
        dend2[5].pt3dadd(-51.21, -15.17, -5.22, 0.43)
        dend2[5].pt3dadd(-53.86, -15.17, -5.22, 0.43)
        dend2[5].pt3dadd(-54.59, -15.17, -5.22, 0.43)
        dend2[5].pt3dadd(-57.25, -15.42, -5.22, 0.43)
        dend2[5].pt3dadd(-57.73, -15.67, -5.22, 0.43)
        dend2[5].pt3dadd(-62.32, -17.91, -5.23, 0.43)
        dend2[5].pt3dadd(-63.04, -18.66, -5.23, 0.43)
        dend2[5].pt3dadd(-64.49, -19.90, -5.23, 0.43)
        dend2[5].pt3dadd(-64.98, -20.40, -5.23, 0.43)
        dend2[5].pt3dadd(-65.46, -21.89, -5.23, 0.43)
        dend2[5].pt3dadd(-66.18, -22.89, -5.23, 0.43)
        dend2[5].pt3dadd(-66.67, -24.13, -5.23, 0.43)
        dend2[5].pt3dadd(-66.91, -25.12, -5.23, 0.43)
        dend2[5].pt3dadd(-67.63, -27.11, -5.23, 0.43)
        dend2[5].pt3dadd(-67.87, -27.86, -5.23, 0.43)
        dend2[5].pt3dadd(-68.60, -29.35, -5.23, 0.43)
        dend2[5].pt3dadd(-69.08, -29.35, -5.23, 0.43)
        dend2[5].pt3dadd(-71.01, -30.60, -5.23, 0.43)
        dend2[5].pt3dadd(-71.50, -32.34, -5.23, 0.43)
        dend2[5].pt3dadd(-72.46, -34.83, -5.93, 0.43)
        dend2[5].pt3dadd(-72.22, -36.32, -5.93, 0.43)
        dend2[5].pt3dadd(-72.22, -36.57, -5.93, 0.43)
        dend2[5].pt3dadd(-73.19, -38.31, -5.93, 0.43)
        dend2[5].pt3dadd(-73.43, -38.56, -5.93, 0.43)
        dend2[5].pt3dadd(-75.12, -39.30, -5.93, 0.39)
        dend2[5].pt3dadd(-75.60, -40.05, -5.93, 0.39)
        dend2[5].pt3dadd(-77.78, -45.77, -9.05, 0.39)
        dend2[5].pt3dadd(-80.43, -48.01, -9.05, 0.39)
        dend2[5].pt3dadd(-82.37, -49.75, -9.05, 0.39)
        dend2[5].pt3dadd(-82.85, -50.25, -9.05, 0.39)
        dend2[5].pt3dadd(-84.54, -51.74, -9.05, 0.39)
        dend2[5].pt3dadd(-85.02, -51.99, -9.05, 0.39)
        dend2[5].pt3dadd(-86.23, -53.48, -9.05, 0.39)
        dend2[5].pt3dadd(-86.23, -54.23, -9.05, 0.39)
        dend2[5].pt3dadd(-86.23, -55.47, -9.05, 0.39)
        dend2[5].pt3dadd(-86.23, -56.97, -9.05, 0.39)
        dend2[5].pt3dadd(-86.23, -58.71, -9.05, 0.39)
        dend2[5].pt3dadd(-86.23, -59.45, -9.05, 0.39)
        dend2[5].pt3dadd(-86.47, -60.45, -9.05, 0.39)
        dend2[5].pt3dadd(-86.47, -61.69, -9.05, 0.39)
        dend2[5].pt3dadd(-86.47, -62.19, -9.05, 0.39)
        dend2[5].pt3dadd(-86.72, -63.43, -9.05, 0.39)
        dend2[5].pt3dadd(-87.20, -64.18, -9.05, 0.39)
        dend2[5].pt3dadd(-88.41, -65.17, -9.05, 0.39)
        dend2[5].pt3dadd(-88.65, -65.67, -9.05, 0.39)
        dend2[5].pt3dadd(-89.37, -66.17, -9.05, 0.39)
        dend2[5].pt3dadd(-90.34, -66.42, -9.05, 0.39)
        dend2[5].pt3dadd(-90.58, -66.67, -9.05, 0.39)

        dend2[6] = h.Section(name='dend2[6]')
        dend2[6].connect(        dend2[4](1), 0)
        dend2[6].pt3dclear()
        dend2[6].pt3dadd(-28.26, -21.64, -4.79, 0.43)
        dend2[6].pt3dadd(-30.19, -23.38, -4.79, 0.43)
        dend2[6].pt3dadd(-34.06, -25.37, -4.79, 0.43)
        dend2[6].pt3dadd(-37.20, -26.87, -4.79, 0.43)
        dend2[6].pt3dadd(-37.44, -26.87, -4.79, 0.43)
        dend2[6].pt3dadd(-40.10, -28.86, -4.79, 0.43)
        dend2[6].pt3dadd(-40.58, -29.10, -4.79, 0.43)
        dend2[6].pt3dadd(-44.69, -29.10, -4.79, 0.43)
        dend2[6].pt3dadd(-47.10, -29.10, -4.79, 0.43)
        dend2[6].pt3dadd(-47.83, -29.60, -4.79, 0.43)
        dend2[6].pt3dadd(-49.03, -30.85, -4.79, 0.43)
        dend2[6].pt3dadd(-49.03, -31.09, -4.79, 0.43)
        dend2[6].pt3dadd(-52.42, -32.09, -4.79, 0.43)
        dend2[6].pt3dadd(-52.66, -32.09, -4.79, 0.43)
        dend2[6].pt3dadd(-55.31, -32.59, -4.79, 0.43)
        dend2[6].pt3dadd(-56.04, -32.84, -4.79, 0.43)
        dend2[6].pt3dadd(-58.21, -33.08, -4.79, 0.43)
        dend2[6].pt3dadd(-58.45, -33.08, -4.79, 0.43)
        dend2[6].pt3dadd(-61.35, -34.58, -4.84, 0.43)
        dend2[6].pt3dadd(-61.35, -34.83, -4.84, 0.43)
        dend2[6].pt3dadd(-62.56, -37.56, -4.84, 0.43)
        dend2[6].pt3dadd(-63.04, -37.81, -4.84, 0.43)
        dend2[6].pt3dadd(-63.53, -39.05, -4.84, 0.43)
        dend2[6].pt3dadd(-64.01, -39.30, -4.84, 0.43)
        dend2[6].pt3dadd(-66.18, -40.80, -4.84, 0.43)
        dend2[6].pt3dadd(-67.15, -41.29, -4.84, 0.43)
        dend2[6].pt3dadd(-69.32, -42.04, -4.84, 0.43)
        dend2[6].pt3dadd(-69.81, -42.54, -4.84, 0.43)
        dend2[6].pt3dadd(-70.05, -44.78, -4.84, 0.43)
        dend2[6].pt3dadd(-70.05, -45.27, -4.84, 0.43)
        dend2[6].pt3dadd(-70.77, -46.52, -4.84, 0.43)
        dend2[6].pt3dadd(-71.74, -46.77, -4.84, 0.43)
        dend2[6].pt3dadd(-74.15, -46.77, -4.84, 0.43)
        dend2[6].pt3dadd(-74.40, -46.77, -4.84, 0.43)
        dend2[6].pt3dadd(-76.09, -48.51, -4.84, 0.43)
        dend2[6].pt3dadd(-76.57, -49.25, -4.84, 0.43)
        dend2[6].pt3dadd(-77.05, -51.49, -4.84, 0.43)
        dend2[6].pt3dadd(-77.05, -51.99, -4.84, 0.43)
        dend2[6].pt3dadd(-77.29, -53.48, -4.84, 0.43)
        dend2[6].pt3dadd(-77.54, -54.48, -4.84, 0.43)
        dend2[6].pt3dadd(-77.54, -54.98, -4.84, 0.43)
        dend2[6].pt3dadd(-77.54, -55.47, -4.84, 0.43)
        dend2[6].pt3dadd(-78.02, -56.72, -1.14, 0.43)
        dend2[6].pt3dadd(-78.50, -56.72, -1.14, 0.43)
        dend2[6].pt3dadd(-80.68, -56.47, -1.10, 0.43)
        dend2[6].pt3dadd(-81.16, -56.47, -1.08, 0.39)
        dend2[6].pt3dadd(-84.30, -55.47, -1.07, 0.39)
        dend2[6].pt3dadd(-84.54, -55.47, -1.07, 0.39)
        dend2[6].pt3dadd(-86.23, -54.98, -1.07, 0.39)
        dend2[6].pt3dadd(-86.96, -55.47, -1.06, 0.39)
        dend2[6].pt3dadd(-88.16, -56.47, -1.06, 0.39)
        dend2[6].pt3dadd(-88.65, -56.47, -1.06, 0.39)
        dend2[6].pt3dadd(-90.10, -56.47, -1.06, 0.39)
        dend2[6].pt3dadd(-90.58, -56.47, -1.06, 0.39)
        dend2[6].pt3dadd(-94.20, -53.98, -1.06, 0.39)
        dend2[6].pt3dadd(-94.93, -55.97, 1.37, 0.39)
        dend2[6].pt3dadd(-95.89, -58.46, 1.37, 0.39)
        dend2[6].pt3dadd(-96.38, -58.96, 1.37, 0.39)
        dend2[6].pt3dadd(-98.07, -60.20, 1.37, 0.39)
        dend2[6].pt3dadd(-99.03, -60.45, 1.37, 0.39)
        dend2[6].pt3dadd(-101.21, -60.70, 1.36, 0.39)
        dend2[6].pt3dadd(-102.17, -60.70, 1.36, 0.39)
        dend2[6].pt3dadd(-103.62, -61.94, 1.36, 0.39)
        dend2[6].pt3dadd(-104.11, -62.44, 1.36, 0.39)
        dend2[6].pt3dadd(-105.80, -62.44, 1.36, 0.39)
        dend2[6].pt3dadd(-106.76, -62.69, 1.36, 0.39)
        dend2[6].pt3dadd(-107.97, -63.93, 1.36, 0.39)
        dend2[6].pt3dadd(-108.70, -64.68, 1.36, 0.39)
        dend2[6].pt3dadd(-109.42, -65.67, 1.36, 0.39)
        dend2[6].pt3dadd(-110.14, -66.42, 1.36, 0.39)
        dend2[6].pt3dadd(-113.29, -66.92, 1.36, 0.39)
        dend2[6].pt3dadd(-113.53, -66.92, 1.36, 0.39)
        dend2[6].pt3dadd(-115.70, -67.41, 1.36, 0.39)
        dend2[6].pt3dadd(-116.43, -68.16, 1.36, 0.39)
        dend2[6].pt3dadd(-117.87, -69.40, 1.36, 0.39)
        dend2[6].pt3dadd(-118.36, -69.40, 1.36, 0.39)
        dend2[6].pt3dadd(-120.77, -70.65, 1.36, 0.39)
        dend2[6].pt3dadd(-121.01, -70.90, 1.36, 0.39)
        dend2[6].pt3dadd(-122.22, -72.39, 1.36, 0.39)
        dend2[6].pt3dadd(-122.71, -72.64, 1.36, 0.39)
        dend2[6].pt3dadd(-127.05, -71.14, 1.36, 0.39)
        dend2[6].pt3dadd(-128.99, -70.65, 1.36, 0.39)
        dend2[6].pt3dadd(-129.23, -70.40, 1.36, 0.39)
        dend2[6].pt3dadd(-130.68, -68.41, 1.36, 0.39)
        dend2[6].pt3dadd(-130.43, -62.19, 4.03, 0.39)
        dend2[6].pt3dadd(-131.88, -60.95, 4.11, 0.39)
        dend2[6].pt3dadd(-131.88, -60.70, 4.11, 0.39)
        dend2[6].pt3dadd(-133.33, -58.71, 4.33, 0.39)
        dend2[6].pt3dadd(-133.57, -58.46, 4.33, 0.39)
        dend2[6].pt3dadd(-134.78, -57.71, 4.73, 0.39)
        dend2[6].pt3dadd(-135.02, -57.71, 4.77, 0.39)
        dend2[6].pt3dadd(-135.99, -57.46, 4.77, 0.39)
        dend2[6].pt3dadd(-136.47, -57.46, 4.77, 0.39)
        dend2[6].pt3dadd(-137.44, -57.46, 4.77, 0.39)
        dend2[6].pt3dadd(-137.68, -57.71, 4.77, 0.39)

        dend2[7] = h.Section(name='dend2[7]')
        dend2[7].connect(        dend2[3](1), 0)
        dend2[7].pt3dclear()
        dend2[7].pt3dadd(-11.84, -19.40, -2.21, 0.54)
        dend2[7].pt3dadd(-14.98, -22.64, 2.25, 0.54)
        dend2[7].pt3dadd(-15.22, -24.63, 2.25, 0.54)
        dend2[7].pt3dadd(-15.94, -25.12, 2.25, 0.54)
        dend2[7].pt3dadd(-18.12, -26.87, 2.25, 0.54)
        dend2[7].pt3dadd(-18.36, -27.11, 2.25, 0.54)
        dend2[7].pt3dadd(-19.08, -27.86, 2.25, 0.54)

        dend2[8] = h.Section(name='dend2[8]')
        dend2[8].connect(        dend2[7](1), 0)
        dend2[8].pt3dclear()
        dend2[8].pt3dadd(-19.08, -27.86, 2.25, 0.48)
        dend2[8].pt3dadd(-24.64, -33.58, 2.24, 0.48)
        dend2[8].pt3dadd(-25.60, -35.32, 2.24, 0.48)
        dend2[8].pt3dadd(-25.60, -36.07, 2.24, 0.48)
        dend2[8].pt3dadd(-26.09, -38.31, 2.24, 0.48)
        dend2[8].pt3dadd(-26.33, -39.05, 2.24, 0.48)

        dend2[9] = h.Section(name='dend2[9]')
        dend2[9].connect(        dend2[8](1), 0)
        dend2[9].pt3dclear()
        dend2[9].pt3dadd(-26.33, -39.05, 2.24, 0.43)
        dend2[9].pt3dadd(-29.47, -39.55, -16.04, 0.43)
        dend2[9].pt3dadd(-29.71, -39.55, -16.04, 0.43)
        dend2[9].pt3dadd(-32.61, -41.29, -16.04, 0.43)
        dend2[9].pt3dadd(-32.85, -41.79, -16.05, 0.43)
        dend2[9].pt3dadd(-34.78, -43.78, -16.05, 0.43)
        dend2[9].pt3dadd(-35.27, -44.03, -16.05, 0.43)
        dend2[9].pt3dadd(-39.86, -45.02, -16.05, 0.43)
        dend2[9].pt3dadd(-40.10, -46.02, -16.05, 0.43)
        dend2[9].pt3dadd(-39.86, -49.01, -16.05, 0.43)
        dend2[9].pt3dadd(-39.86, -49.25, -16.05, 0.43)
        dend2[9].pt3dadd(-45.89, -50.75, -16.05, 0.43)
        dend2[9].pt3dadd(-46.14, -50.75, -16.05, 0.43)
        dend2[9].pt3dadd(-50.72, -51.99, -16.05, 0.43)
        dend2[9].pt3dadd(-51.21, -52.24, -16.05, 0.43)
        dend2[9].pt3dadd(-57.25, -49.01, -16.05, 0.43)
        dend2[9].pt3dadd(-63.53, -48.26, -16.05, 0.43)
        dend2[9].pt3dadd(-65.46, -47.26, -16.05, 0.43)
        dend2[9].pt3dadd(-65.70, -47.26, -16.05, 0.43)
        dend2[9].pt3dadd(-68.60, -47.01, -22.70, 0.43)
        dend2[9].pt3dadd(-68.84, -47.26, -22.71, 0.43)
        dend2[9].pt3dadd(-71.26, -47.26, -22.96, 0.43)
        dend2[9].pt3dadd(-71.26, -47.26, -22.98, 0.43)
        dend2[9].pt3dadd(-75.60, -47.26, -23.03, 0.43)
        dend2[9].pt3dadd(-75.60, -47.26, -23.11, 0.43)
        dend2[9].pt3dadd(-78.26, -48.01, -24.22, 0.43)
        dend2[9].pt3dadd(-78.26, -48.01, -24.23, 0.43)
        dend2[9].pt3dadd(-80.43, -48.76, -24.26, 0.43)
        dend2[9].pt3dadd(-80.68, -48.76, -24.33, 0.43)
        dend2[9].pt3dadd(-81.64, -48.76, -24.49, 0.43)
        dend2[9].pt3dadd(-82.13, -48.76, -24.49, 0.43)
        dend2[9].pt3dadd(-85.75, -48.76, -24.49, 0.43)
        dend2[9].pt3dadd(-86.23, -48.76, -24.49, 0.43)
        dend2[9].pt3dadd(-87.44, -50.00, -24.49, 0.39)
        dend2[9].pt3dadd(-87.68, -50.00, -24.49, 0.39)
        dend2[9].pt3dadd(-89.61, -49.25, -24.49, 0.39)
        dend2[9].pt3dadd(-89.86, -49.01, -24.49, 0.39)
        dend2[9].pt3dadd(-92.03, -47.76, -26.98, 0.39)
        dend2[9].pt3dadd(-92.27, -47.76, -26.99, 0.39)
        dend2[9].pt3dadd(-96.86, -46.27, -27.05, 0.39)
        dend2[9].pt3dadd(-96.86, -46.27, -27.08, 0.39)
        dend2[9].pt3dadd(-99.28, -46.52, -27.65, 0.39)
        dend2[9].pt3dadd(-99.28, -46.52, -27.68, 0.39)
        dend2[9].pt3dadd(-101.21, -47.01, -27.70, 0.39)
        dend2[9].pt3dadd(-105.07, -45.52, -27.70, 0.39)
        dend2[9].pt3dadd(-105.31, -45.52, -27.71, 0.39)
        dend2[9].pt3dadd(-108.21, -49.01, -30.50, 0.39)
        dend2[9].pt3dadd(-108.45, -49.01, -30.50, 0.39)
        dend2[9].pt3dadd(-113.04, -50.25, -30.56, 0.39)
        dend2[9].pt3dadd(-114.01, -50.50, -30.60, 0.39)
        dend2[9].pt3dadd(-115.46, -51.00, -30.77, 0.39)
        dend2[9].pt3dadd(-115.70, -51.00, -30.83, 0.39)
        dend2[9].pt3dadd(-117.39, -51.24, -30.84, 0.39)
        dend2[9].pt3dadd(-118.12, -51.49, -30.85, 0.39)
        dend2[9].pt3dadd(-119.32, -51.49, -30.86, 0.39)

        dend2[10] = h.Section(name='dend2[10]')
        dend2[10].connect(        dend2[8](1), 0)
        dend2[10].pt3dclear()
        dend2[10].pt3dadd(-26.33, -39.05, 2.24, 0.48)
        dend2[10].pt3dadd(-29.71, -43.78, 2.24, 0.48)
        dend2[10].pt3dadd(-31.64, -45.52, 2.24, 0.48)
        dend2[10].pt3dadd(-32.85, -47.51, 2.24, 0.48)
        dend2[10].pt3dadd(-34.54, -49.25, 2.24, 0.48)
        dend2[10].pt3dadd(-34.54, -50.00, 2.24, 0.48)
        dend2[10].pt3dadd(-35.02, -51.74, 2.24, 0.48)
        dend2[10].pt3dadd(-35.27, -52.49, 2.24, 0.48)
        dend2[10].pt3dadd(-35.27, -53.73, 2.24, 0.48)
        dend2[10].pt3dadd(-35.27, -54.48, 2.24, 0.48)
        dend2[10].pt3dadd(-37.68, -57.46, 1.55, 0.48)
        dend2[10].pt3dadd(-40.10, -58.96, 1.55, 0.48)
        dend2[10].pt3dadd(-40.58, -58.96, 1.55, 0.48)
        dend2[10].pt3dadd(-41.79, -59.20, -0.01, 0.48)

        dend2[11] = h.Section(name='dend2[11]')
        dend2[11].connect(        dend2[10](1), 0)
        dend2[11].pt3dclear()
        dend2[11].pt3dadd(-41.79, -59.20, -0.01, 0.48)
        dend2[11].pt3dadd(-43.96, -59.95, 1.55, 0.48)
        dend2[11].pt3dadd(-45.89, -61.94, 1.55, 0.48)
        dend2[11].pt3dadd(-46.14, -62.19, 1.55, 0.48)
        dend2[11].pt3dadd(-47.34, -63.68, 1.55, 0.48)
        dend2[11].pt3dadd(-47.34, -64.18, 1.55, 0.48)
        dend2[11].pt3dadd(-48.31, -65.67, 1.55, 0.48)
        dend2[11].pt3dadd(-48.55, -65.67, 1.55, 0.48)

        dend2[12] = h.Section(name='dend2[12]')
        dend2[12].connect(        dend2[11](1), 0)
        dend2[12].pt3dclear()
        dend2[12].pt3dadd(-48.55, -65.67, 1.55, 0.39)
        dend2[12].pt3dadd(-52.42, -68.91, -1.37, 0.39)
        dend2[12].pt3dadd(-52.42, -69.15, -1.37, 0.39)
        dend2[12].pt3dadd(-53.62, -71.89, -1.37, 0.39)
        dend2[12].pt3dadd(-53.62, -72.64, -1.37, 0.39)
        dend2[12].pt3dadd(-55.80, -75.12, -1.37, 0.39)
        dend2[12].pt3dadd(-56.52, -75.62, -1.37, 0.39)
        dend2[12].pt3dadd(-60.14, -77.11, 4.56, 0.39)
        dend2[12].pt3dadd(-60.39, -77.11, 4.56, 0.39)
        dend2[12].pt3dadd(-63.77, -77.36, 4.56, 0.39)
        dend2[12].pt3dadd(-63.77, -77.86, 4.56, 0.39)
        dend2[12].pt3dadd(-65.94, -79.60, 4.56, 0.39)
        dend2[12].pt3dadd(-66.18, -80.10, 4.56, 0.39)
        dend2[12].pt3dadd(-68.36, -81.84, 4.56, 0.39)
        dend2[12].pt3dadd(-68.60, -82.59, 4.56, 0.39)
        dend2[12].pt3dadd(-68.84, -84.08, 4.56, 0.39)
        dend2[12].pt3dadd(-68.84, -85.07, 4.56, 0.39)
        dend2[12].pt3dadd(-67.63, -86.32, 4.56, 0.39)
        dend2[12].pt3dadd(-66.91, -87.81, 5.15, 0.39)
        dend2[12].pt3dadd(-67.15, -90.30, 5.15, 0.39)
        dend2[12].pt3dadd(-69.81, -93.03, 7.80, 0.39)
        dend2[12].pt3dadd(-70.29, -96.52, 7.80, 0.39)
        dend2[12].pt3dadd(-71.74, -99.50, 7.80, 0.39)
        dend2[12].pt3dadd(-72.46, -100.25, 7.80, 0.39)
        dend2[12].pt3dadd(-73.91, -102.49, 7.80, 0.39)
        dend2[12].pt3dadd(-75.60, -103.23, 7.80, 0.39)
        dend2[12].pt3dadd(-76.33, -103.48, 7.80, 0.39)
        dend2[12].pt3dadd(-79.23, -104.48, 7.80, 0.39)
        dend2[12].pt3dadd(-79.71, -105.22, 7.80, 0.39)
        dend2[12].pt3dadd(-79.47, -107.21, 7.80, 0.39)
        dend2[12].pt3dadd(-79.47, -107.46, 7.80, 0.39)
        dend2[12].pt3dadd(-79.47, -109.20, 9.77, 0.39)
        dend2[12].pt3dadd(-82.13, -109.20, 9.80, 0.39)
        dend2[12].pt3dadd(-82.61, -109.20, 9.80, 0.39)
        dend2[12].pt3dadd(-84.54, -110.95, 9.80, 0.39)
        dend2[12].pt3dadd(-84.54, -111.44, 9.80, 0.39)
        dend2[12].pt3dadd(-84.54, -112.94, 9.80, 0.39)
        dend2[12].pt3dadd(-84.30, -113.43, 9.80, 0.39)
        dend2[12].pt3dadd(-81.40, -117.91, 10.29, 0.39)
        dend2[12].pt3dadd(-81.40, -117.91, 10.31, 0.39)
        dend2[12].pt3dadd(-79.71, -119.15, 10.31, 0.39)
        dend2[12].pt3dadd(-78.99, -123.63, 10.31, 0.39)
        dend2[12].pt3dadd(-78.74, -123.63, 10.31, 0.39)

        dend2[13] = h.Section(name='dend2[13]')
        dend2[13].connect(        dend2[11](1), 0)
        dend2[13].pt3dclear()
        dend2[13].pt3dadd(-48.55, -65.67, 1.55, 0.39)
        dend2[13].pt3dadd(-53.38, -68.16, 1.55, 0.39)
        dend2[13].pt3dadd(-53.86, -68.16, 1.55, 0.39)
        dend2[13].pt3dadd(-56.76, -69.40, 1.55, 0.39)
        dend2[13].pt3dadd(-57.25, -69.40, 1.55, 0.39)
        dend2[13].pt3dadd(-60.14, -72.14, 4.09, 0.39)
        dend2[13].pt3dadd(-60.63, -72.39, 4.09, 0.39)
        dend2[13].pt3dadd(-64.01, -74.13, 4.09, 0.39)
        dend2[13].pt3dadd(-64.73, -74.88, 4.09, 0.39)
        dend2[13].pt3dadd(-66.91, -76.87, 4.09, 0.39)
        dend2[13].pt3dadd(-67.39, -77.36, 4.09, 0.39)
        dend2[13].pt3dadd(-70.29, -80.60, 4.09, 0.39)
        dend2[13].pt3dadd(-71.74, -81.09, 4.09, 0.39)
        dend2[13].pt3dadd(-73.19, -82.34, 4.09, 0.39)
        dend2[13].pt3dadd(-73.19, -83.83, 4.09, 0.39)
        dend2[13].pt3dadd(-72.95, -87.31, -1.76, 0.39)
        dend2[13].pt3dadd(-73.67, -91.04, -1.76, 0.39)
        dend2[13].pt3dadd(-74.40, -93.28, -1.76, 0.39)
        dend2[13].pt3dadd(-75.36, -93.53, -1.76, 0.39)
        dend2[13].pt3dadd(-78.02, -94.03, -1.76, 0.39)
        dend2[13].pt3dadd(-79.23, -94.03, -1.76, 0.39)
        dend2[13].pt3dadd(-81.16, -92.04, -1.76, 0.39)
        dend2[13].pt3dadd(-81.16, -91.54, -8.32, 0.39)
        dend2[13].pt3dadd(-81.40, -91.54, -8.32, 0.39)
        dend2[13].pt3dadd(-84.06, -91.79, -8.31, 0.39)
        dend2[13].pt3dadd(-86.23, -93.28, -8.31, 0.39)
        dend2[13].pt3dadd(-86.47, -93.53, -8.31, 0.39)
        dend2[13].pt3dadd(-89.61, -94.78, -8.31, 0.39)
        dend2[13].pt3dadd(-90.10, -94.78, -8.31, 0.39)
        dend2[13].pt3dadd(-93.72, -95.52, -8.31, 0.39)
        dend2[13].pt3dadd(-93.96, -95.77, -8.31, 0.39)
        dend2[13].pt3dadd(-95.41, -97.02, -8.31, 0.39)
        dend2[13].pt3dadd(-96.14, -97.51, -8.31, 0.39)
        dend2[13].pt3dadd(-97.58, -98.26, -8.31, 0.39)
        dend2[13].pt3dadd(-98.55, -98.26, -8.31, 0.39)
        dend2[13].pt3dadd(-100.48, -97.26, -8.31, 0.39)
        dend2[13].pt3dadd(-102.17, -96.52, -8.31, 0.39)
        dend2[13].pt3dadd(-104.11, -93.28, -8.31, 0.39)
        dend2[13].pt3dadd(-104.59, -89.30, -11.71, 0.39)
        dend2[13].pt3dadd(-104.83, -86.07, -13.58, 0.39)
        dend2[13].pt3dadd(-104.83, -86.07, -13.63, 0.39)
        dend2[13].pt3dadd(-107.25, -83.08, -13.70, 0.39)

        dend2[14] = h.Section(name='dend2[14]')
        dend2[14].connect(        dend2[10](1), 0)
        dend2[14].pt3dclear()
        dend2[14].pt3dadd(-41.79, -59.20, -0.01, 0.43)
        dend2[14].pt3dadd(-42.03, -61.94, -3.00, 0.43)
        dend2[14].pt3dadd(-43.96, -64.18, -3.00, 0.43)
        dend2[14].pt3dadd(-44.20, -64.43, -3.00, 0.43)
        dend2[14].pt3dadd(-45.89, -66.92, -3.00, 0.43)
        dend2[14].pt3dadd(-46.14, -67.16, -3.00, 0.43)
        dend2[14].pt3dadd(-46.62, -69.15, -3.01, 0.43)
        dend2[14].pt3dadd(-46.86, -69.90, -3.01, 0.43)
        dend2[14].pt3dadd(-48.07, -71.14, -3.01, 0.43)
        dend2[14].pt3dadd(-48.55, -71.39, -3.01, 0.43)
        dend2[14].pt3dadd(-50.97, -72.14, -9.02, 0.43)
        dend2[14].pt3dadd(-51.21, -72.39, -9.02, 0.43)
        dend2[14].pt3dadd(-52.42, -74.38, -9.01, 0.43)
        dend2[14].pt3dadd(-52.66, -74.63, -9.01, 0.43)
        dend2[14].pt3dadd(-53.62, -76.12, -9.01, 0.43)
        dend2[14].pt3dadd(-54.35, -76.37, -9.01, 0.43)
        dend2[14].pt3dadd(-56.04, -77.11, -9.01, 0.43)
        dend2[14].pt3dadd(-56.76, -77.11, -9.01, 0.43)
        dend2[14].pt3dadd(-59.42, -77.86, -9.01, 0.43)
        dend2[14].pt3dadd(-59.90, -78.11, -9.01, 0.43)
        dend2[14].pt3dadd(-61.35, -79.60, -9.01, 0.43)
        dend2[14].pt3dadd(-61.59, -80.10, -9.01, 0.43)
        dend2[14].pt3dadd(-62.56, -80.85, -9.01, 0.43)
        dend2[14].pt3dadd(-63.04, -81.09, -9.01, 0.43)
        dend2[14].pt3dadd(-64.25, -84.08, -9.01, 0.43)
        dend2[14].pt3dadd(-64.49, -84.33, -9.01, 0.43)
        dend2[14].pt3dadd(-65.70, -87.06, -10.76, 0.43)
        dend2[14].pt3dadd(-67.63, -89.30, -10.76, 0.43)
        dend2[14].pt3dadd(-67.87, -89.30, -10.76, 0.43)
        dend2[14].pt3dadd(-71.26, -89.30, -10.76, 0.43)
        dend2[14].pt3dadd(-71.98, -89.55, -10.76, 0.43)
        dend2[14].pt3dadd(-74.64, -90.05, -10.76, 0.43)
        dend2[14].pt3dadd(-75.12, -90.30, -10.76, 0.43)
        dend2[14].pt3dadd(-77.29, -91.29, -10.76, 0.43)
        dend2[14].pt3dadd(-78.50, -91.29, -10.76, 0.43)
        dend2[14].pt3dadd(-80.92, -93.03, -10.76, 0.43)
        dend2[14].pt3dadd(-81.88, -94.28, -10.76, 0.43)
        dend2[14].pt3dadd(-82.85, -95.02, -10.76, 0.43)
        dend2[14].pt3dadd(-84.30, -95.27, -10.76, 0.43)
        dend2[14].pt3dadd(-86.72, -96.27, -10.76, 0.43)
        dend2[14].pt3dadd(-87.68, -97.51, -10.76, 0.43)
        dend2[14].pt3dadd(-88.41, -99.25, -10.76, 0.43)
        dend2[14].pt3dadd(-89.86, -100.75, -10.76, 0.43)
        dend2[14].pt3dadd(-94.44, -101.49, -8.99, 0.39)
        dend2[14].pt3dadd(-95.41, -104.23, -8.99, 0.39)
        dend2[14].pt3dadd(-95.89, -104.73, -8.99, 0.39)
        dend2[14].pt3dadd(-97.34, -105.72, -8.99, 0.39)
        dend2[14].pt3dadd(-98.31, -106.22, -8.99, 0.39)
        dend2[14].pt3dadd(-98.79, -107.46, -8.99, 0.39)
        dend2[14].pt3dadd(-99.28, -108.46, -8.99, 0.39)
        dend2[14].pt3dadd(-100.24, -110.45, -8.99, 0.39)
        dend2[14].pt3dadd(-100.97, -110.95, -8.99, 0.39)
        dend2[14].pt3dadd(-102.42, -111.94, -8.99, 0.39)
        dend2[14].pt3dadd(-102.90, -112.44, -8.99, 0.39)
        dend2[14].pt3dadd(-106.28, -115.17, -6.30, 0.39)
        dend2[14].pt3dadd(-106.52, -115.17, -6.30, 0.39)
        dend2[14].pt3dadd(-107.97, -115.42, -6.30, 0.39)
        dend2[14].pt3dadd(-108.45, -115.67, -6.30, 0.39)
        dend2[14].pt3dadd(-109.90, -117.16, -6.31, 0.39)
        dend2[14].pt3dadd(-110.39, -117.91, -6.31, 0.39)
        dend2[14].pt3dadd(-110.39, -119.15, -6.31, 0.39)
        dend2[14].pt3dadd(-110.39, -120.40, -6.31, 0.39)
        dend2[14].pt3dadd(-110.39, -121.39, -6.31, 0.39)
        dend2[14].pt3dadd(-110.39, -122.39, -6.31, 0.39)
        dend2[14].pt3dadd(-109.90, -123.38, -6.31, 0.39)
        dend2[14].pt3dadd(-110.14, -123.63, -6.31, 0.39)
        dend2[14].pt3dadd(-112.32, -124.38, -6.31, 0.39)
        dend2[14].pt3dadd(-113.29, -124.63, -6.31, 0.39)
        dend2[14].pt3dadd(-114.73, -125.62, -6.31, 0.39)
        dend2[14].pt3dadd(-114.98, -126.12, -6.31, 0.39)
        dend2[14].pt3dadd(-115.22, -127.86, -6.31, 0.39)
        dend2[14].pt3dadd(-116.67, -130.10, -6.32, 0.39)
        dend2[14].pt3dadd(-116.91, -132.34, -6.32, 0.39)

        dend2[15] = h.Section(name='dend2[15]')
        dend2[15].connect(        dend2[7](1), 0)
        dend2[15].pt3dclear()
        dend2[15].pt3dadd(-19.08, -27.86, 2.25, 0.77)
        dend2[15].pt3dadd(-18.11, -30.85, -3.10, 0.77)
        dend2[15].pt3dadd(-18.59, -32.35, -3.10, 0.77)
        dend2[15].pt3dadd(-19.32, -33.09, -3.10, 0.77)
        dend2[15].pt3dadd(-20.28, -34.83, -3.10, 0.77)
        dend2[15].pt3dadd(-19.32, -37.32, 4.32, 0.77)
        dend2[15].pt3dadd(-19.08, -37.32, 4.32, 0.77)
        dend2[15].pt3dadd(-17.39, -39.31, 4.32, 0.77)
        dend2[15].pt3dadd(-17.39, -39.81, 4.31, 0.77)
        dend2[15].pt3dadd(-18.84, -41.30, 4.31, 0.77)
        dend2[15].pt3dadd(-19.08, -41.55, 4.31, 0.77)

        dend2[16] = h.Section(name='dend2[16]')
        dend2[16].connect(        dend2[15](1), 0)
        dend2[16].pt3dclear()
        dend2[16].pt3dadd(-19.08, -41.55, 4.31, 0.43)
        dend2[16].pt3dadd(-20.28, -45.03, 2.71, 0.43)
        dend2[16].pt3dadd(-21.73, -48.27, 2.71, 0.43)
        dend2[16].pt3dadd(-23.67, -56.23, 9.41, 0.43)
        dend2[16].pt3dadd(-24.63, -58.47, 9.42, 0.43)
        dend2[16].pt3dadd(-26.56, -60.95, 9.43, 0.43)
        dend2[16].pt3dadd(-26.56, -61.20, 9.43, 0.43)

        dend2[17] = h.Section(name='dend2[17]')
        dend2[17].connect(        dend2[15](1), 0)
        dend2[17].pt3dclear()
        dend2[17].pt3dadd(-19.08, -41.55, 4.31, 0.43)
        dend2[17].pt3dadd(-22.46, -41.55, 4.85, 0.43)
        dend2[17].pt3dadd(-24.87, -42.05, 4.85, 0.43)
        dend2[17].pt3dadd(-25.12, -42.30, 4.85, 0.43)
        dend2[17].pt3dadd(-26.81, -44.54, 4.85, 0.43)
        dend2[17].pt3dadd(-26.81, -44.78, 4.85, 0.43)
        dend2[17].pt3dadd(-27.05, -46.53, 4.85, 0.43)
        dend2[17].pt3dadd(-27.29, -48.02, 4.85, 0.43)
        dend2[17].pt3dadd(-27.53, -51.50, 5.45, 0.43)
        dend2[17].pt3dadd(-28.98, -52.50, 5.45, 0.43)
        dend2[17].pt3dadd(-29.70, -52.99, 5.45, 0.43)
        dend2[17].pt3dadd(-30.67, -54.49, 5.45, 0.43)
        dend2[17].pt3dadd(-31.15, -54.98, 5.45, 0.43)
        dend2[17].pt3dadd(-32.60, -56.72, 5.45, 0.43)
        dend2[17].pt3dadd(-32.85, -56.97, 5.45, 0.43)
        dend2[17].pt3dadd(-33.33, -58.22, 5.45, 0.43)
        dend2[17].pt3dadd(-34.05, -59.46, 5.45, 0.43)
        dend2[17].pt3dadd(-34.29, -61.70, 5.45, 0.43)
        dend2[17].pt3dadd(-34.29, -61.95, 5.45, 0.43)
        dend2[17].pt3dadd(-35.02, -63.19, 5.45, 0.43)
        dend2[17].pt3dadd(-35.74, -64.44, 5.45, 0.43)
        dend2[17].pt3dadd(-36.47, -67.67, 5.45, 0.43)
        dend2[17].pt3dadd(-36.71, -68.91, 5.45, 0.43)
        dend2[17].pt3dadd(-36.71, -69.41, 5.45, 0.43)
        dend2[17].pt3dadd(-38.16, -70.66, 5.45, 0.43)
        dend2[17].pt3dadd(-38.88, -71.90, 5.45, 0.43)
        dend2[17].pt3dadd(-39.61, -72.40, 5.45, 0.43)
        dend2[17].pt3dadd(-41.30, -74.64, -3.09, 0.43)
        dend2[17].pt3dadd(-42.02, -76.63, -3.09, 0.43)
        dend2[17].pt3dadd(-41.30, -79.86, -0.65, 0.43)
        dend2[17].pt3dadd(-42.27, -81.35, -0.65, 0.43)
        dend2[17].pt3dadd(-43.23, -82.10, -0.65, 0.43)
        dend2[17].pt3dadd(-44.92, -84.09, -0.65, 0.43)
        dend2[17].pt3dadd(-45.41, -84.59, -0.65, 0.43)
        dend2[17].pt3dadd(-45.89, -85.83, -0.65, 0.43)
        dend2[17].pt3dadd(-46.61, -86.33, -0.65, 0.43)
        dend2[17].pt3dadd(-47.82, -87.82, -0.65, 0.43)
        dend2[17].pt3dadd(-48.30, -88.32, -0.65, 0.43)
        dend2[17].pt3dadd(-48.55, -90.56, -0.65, 0.43)
        dend2[17].pt3dadd(-48.55, -91.30, -0.65, 0.43)
        dend2[17].pt3dadd(-48.55, -92.55, -0.65, 0.43)
        dend2[17].pt3dadd(-48.79, -92.79, -0.65, 0.43)
        dend2[17].pt3dadd(-49.03, -93.54, -0.65, 0.43)
        dend2[17].pt3dadd(-49.51, -94.29, -0.65, 0.43)
        dend2[17].pt3dadd(-50.48, -96.03, -0.65, 0.43)
        dend2[17].pt3dadd(-50.72, -96.28, -0.65, 0.43)

        dend2[18] = h.Section(name='dend2[18]')
        dend2[18].connect(        dend2[17](1), 0)
        dend2[18].pt3dclear()
        dend2[18].pt3dadd(-50.72, -96.28, -0.65, 0.39)
        dend2[18].pt3dadd(-51.69, -99.01, -1.95, 0.39)
        dend2[18].pt3dadd(-53.38, -102.99, -1.95, 0.39)
        dend2[18].pt3dadd(-54.83, -105.73, -1.95, 0.39)
        dend2[18].pt3dadd(-55.07, -106.48, -1.95, 0.39)
        dend2[18].pt3dadd(-54.10, -108.22, -1.95, 0.39)
        dend2[18].pt3dadd(-53.86, -108.47, -1.95, 0.39)
        dend2[18].pt3dadd(-51.93, -112.69, -2.98, 0.39)
        dend2[18].pt3dadd(-52.65, -115.93, -2.98, 0.39)
        dend2[18].pt3dadd(-52.65, -116.18, -2.98, 0.39)
        dend2[18].pt3dadd(-54.10, -118.91, -2.98, 0.39)
        dend2[18].pt3dadd(-56.03, -119.91, -2.98, 0.39)
        dend2[18].pt3dadd(-58.45, -120.90, -2.98, 0.39)
        dend2[18].pt3dadd(-59.90, -121.40, -2.98, 0.39)
        dend2[18].pt3dadd(-60.62, -123.39, -2.98, 0.39)
        dend2[18].pt3dadd(-60.86, -123.64, -2.98, 0.39)
        dend2[18].pt3dadd(-62.56, -128.12, -2.98, 0.39)
        dend2[18].pt3dadd(-62.80, -128.12, -2.98, 0.39)
        dend2[18].pt3dadd(-64.49, -128.86, -2.98, 0.39)
        dend2[18].pt3dadd(-65.21, -130.11, -2.98, 0.39)
        dend2[18].pt3dadd(-66.42, -132.60, -2.98, 0.39)
        dend2[18].pt3dadd(-67.14, -132.84, -2.98, 0.39)
        dend2[18].pt3dadd(-70.53, -133.59, -2.98, 0.39)
        dend2[18].pt3dadd(-73.67, -136.82, -2.53, 0.39)
        dend2[18].pt3dadd(-75.12, -139.81, -2.53, 0.39)
        dend2[18].pt3dadd(-75.12, -140.31, -2.53, 0.39)
        dend2[18].pt3dadd(-75.12, -143.04, -2.53, 0.39)
        dend2[18].pt3dadd(-78.74, -144.78, -2.53, 0.39)
        dend2[18].pt3dadd(-79.22, -145.03, -2.54, 0.39)

        dend2[19] = h.Section(name='dend2[19]')
        dend2[19].connect(        dend2[17](1), 0)
        dend2[19].pt3dclear()
        dend2[19].pt3dadd(-50.72, -96.28, -0.65, 0.39)
        dend2[19].pt3dadd(-53.13, -100.26, -17.34, 0.39)
        dend2[19].pt3dadd(-55.31, -103.24, -17.34, 0.39)
        dend2[19].pt3dadd(-55.31, -104.24, -17.34, 0.39)
        dend2[19].pt3dadd(-56.03, -105.73, -17.34, 0.39)
        dend2[19].pt3dadd(-56.03, -106.72, -17.34, 0.39)
        dend2[19].pt3dadd(-56.76, -108.71, -17.34, 0.39)
        dend2[19].pt3dadd(-56.76, -108.96, -17.34, 0.39)
        dend2[19].pt3dadd(-58.45, -110.70, -17.34, 0.39)
        dend2[19].pt3dadd(-59.17, -110.95, -17.34, 0.39)
        dend2[19].pt3dadd(-60.38, -112.94, -17.34, 0.39)
        dend2[19].pt3dadd(-60.62, -115.68, -17.34, 0.39)
        dend2[19].pt3dadd(-60.62, -117.17, -16.45, 0.39)
        dend2[19].pt3dadd(-60.62, -117.42, -16.45, 0.39)
        dend2[19].pt3dadd(-61.35, -119.16, -16.45, 0.39)
        dend2[19].pt3dadd(-63.52, -120.66, -16.45, 0.39)
        dend2[19].pt3dadd(-64.73, -121.65, -16.45, 0.39)
        dend2[19].pt3dadd(-64.97, -122.15, -16.45, 0.39)
        dend2[19].pt3dadd(-65.21, -123.39, -16.45, 0.39)
        dend2[19].pt3dadd(-65.21, -124.88, -16.45, 0.39)
        dend2[19].pt3dadd(-65.21, -125.88, -16.45, 0.39)
        dend2[19].pt3dadd(-65.21, -126.63, -16.45, 0.39)
        dend2[19].pt3dadd(-64.97, -128.37, -16.45, 0.39)
        dend2[19].pt3dadd(-64.00, -130.36, -16.45, 0.39)
        dend2[19].pt3dadd(-63.28, -131.60, -16.45, 0.39)
        dend2[19].pt3dadd(-63.52, -132.35, -16.45, 0.39)
        dend2[19].pt3dadd(-64.00, -133.84, -16.45, 0.39)
        dend2[19].pt3dadd(-64.49, -134.59, -16.45, 0.39)
        dend2[19].pt3dadd(-65.21, -135.33, -16.45, 0.39)
        dend2[19].pt3dadd(-66.18, -136.58, -16.45, 0.39)
        dend2[19].pt3dadd(-66.42, -136.82, -16.45, 0.39)
        dend2[19].pt3dadd(-66.90, -137.57, -16.45, 0.39)
        dend2[19].pt3dadd(-67.14, -137.82, -16.45, 0.39)
        dend2[19].pt3dadd(-67.87, -138.32, -16.45, 0.39)
        dend2[19].pt3dadd(-70.77, -140.80, -26.08, 0.39)
        dend2[19].pt3dadd(-71.98, -144.78, -26.10, 0.39)
        dend2[19].pt3dadd(-72.22, -144.78, -26.10, 0.39)
        dend2[19].pt3dadd(-72.22, -146.28, -26.13, 0.39)
        dend2[19].pt3dadd(-72.22, -146.28, -26.17, 0.39)

        dend2[20] = h.Section(name='dend2[20]')
        dend2[20].connect(        dend2[2](1), 0)
        dend2[20].pt3dclear()
        dend2[20].pt3dadd(-8.21, -11.19, 0.22, 0.96)
        dend2[20].pt3dadd(-6.04, -15.92, -1.10, 0.96)
        dend2[20].pt3dadd(-5.07, -21.39, -1.76, 0.61)
        dend2[20].pt3dadd(-6.28, -24.38, -1.76, 0.61)
        dend2[20].pt3dadd(-6.76, -24.88, -1.76, 0.61)
        dend2[20].pt3dadd(-8.94, -27.36, -1.76, 0.61)
        dend2[20].pt3dadd(-9.18, -27.61, -1.76, 0.61)
        dend2[20].pt3dadd(-9.42, -29.10, -1.76, 0.61)
        dend2[20].pt3dadd(-9.42, -29.35, -1.76, 0.61)
        dend2[20].pt3dadd(-10.14, -35.57, -2.55, 0.61)
        dend2[20].pt3dadd(-9.66, -37.31, -2.55, 0.61)
        dend2[20].pt3dadd(-10.14, -39.05, -2.55, 0.61)
        dend2[20].pt3dadd(-10.39, -39.05, -2.55, 0.61)
        dend2[20].pt3dadd(-11.11, -40.55, -2.55, 0.61)
        dend2[20].pt3dadd(-11.59, -40.80, -2.55, 0.61)
        dend2[20].pt3dadd(-12.08, -42.29, -2.55, 0.61)
        dend2[20].pt3dadd(-12.56, -42.79, -2.55, 0.61)
        dend2[20].pt3dadd(-13.04, -43.28, -2.55, 0.61)
        dend2[20].pt3dadd(-13.29, -43.53, -2.55, 0.61)

        dend2[21] = h.Section(name='dend2[21]')
        dend2[21].connect(        dend2[20](1), 0)
        dend2[21].pt3dclear()
        dend2[21].pt3dadd(-13.29, -43.53, -2.55, 0.54)
        dend2[21].pt3dadd(-12.08, -47.51, -3.28, 0.54)
        dend2[21].pt3dadd(-13.53, -49.25, -3.28, 0.54)
        dend2[21].pt3dadd(-13.77, -49.50, -3.28, 0.54)
        dend2[21].pt3dadd(-14.49, -51.74, -3.28, 0.54)
        dend2[21].pt3dadd(-14.73, -52.49, -3.28, 0.54)
        dend2[21].pt3dadd(-14.25, -54.23, -3.28, 0.54)
        dend2[21].pt3dadd(-14.25, -55.22, -3.28, 0.54)
        dend2[21].pt3dadd(-13.04, -57.21, -3.28, 0.54)
        dend2[21].pt3dadd(-14.25, -62.44, 4.50, 0.54)
        dend2[21].pt3dadd(-16.18, -64.43, 4.50, 0.54)
        dend2[21].pt3dadd(-16.43, -64.68, 4.50, 0.54)
        dend2[21].pt3dadd(-16.67, -66.67, 4.50, 0.54)
        dend2[21].pt3dadd(-16.43, -66.67, 4.50, 0.54)
        dend2[21].pt3dadd(-16.43, -68.16, 4.49, 0.54)
        dend2[21].pt3dadd(-14.98, -72.14, 4.49, 0.54)
        dend2[21].pt3dadd(-14.73, -72.14, 4.49, 0.54)
        dend2[21].pt3dadd(-14.01, -75.37, 4.49, 0.54)
        dend2[21].pt3dadd(-14.25, -81.34, 0.63, 0.54)
        dend2[21].pt3dadd(-16.67, -84.58, 0.63, 0.54)
        dend2[21].pt3dadd(-14.49, -88.06, 0.63, 0.54)
        dend2[21].pt3dadd(-14.49, -88.31, 0.63, 0.54)
        dend2[21].pt3dadd(-11.59, -93.53, 1.59, 0.54)

        dend2[22] = h.Section(name='dend2[22]')
        dend2[22].connect(        dend2[21](1), 0)
        dend2[22].pt3dclear()
        dend2[22].pt3dadd(-11.59, -93.53, 1.59, 0.43)
        dend2[22].pt3dadd(-9.18, -98.01, 1.61, 0.43)
        dend2[22].pt3dadd(-7.00, -101.49, -1.55, 0.43)
        dend2[22].pt3dadd(-7.73, -103.73, -1.55, 0.43)
        dend2[22].pt3dadd(-7.97, -103.98, -1.55, 0.43)
        dend2[22].pt3dadd(-7.73, -105.97, -1.55, 0.43)
        dend2[22].pt3dadd(-6.76, -105.97, -1.55, 0.43)
        dend2[22].pt3dadd(-5.80, -107.21, -1.55, 0.43)
        dend2[22].pt3dadd(-5.31, -107.46, -1.55, 0.43)
        dend2[22].pt3dadd(-4.83, -108.71, -1.55, 0.43)
        dend2[22].pt3dadd(-4.59, -108.96, -1.55, 0.43)
        dend2[22].pt3dadd(-4.59, -110.20, -1.55, 0.43)
        dend2[22].pt3dadd(-4.59, -110.70, -1.55, 0.43)
        dend2[22].pt3dadd(-4.59, -114.43, -1.55, 0.43)
        dend2[22].pt3dadd(-4.59, -114.93, -1.55, 0.43)
        dend2[22].pt3dadd(-4.11, -116.42, -1.55, 0.43)
        dend2[22].pt3dadd(-3.86, -117.91, -1.55, 0.43)
        dend2[22].pt3dadd(-3.38, -120.65, -1.55, 0.43)
        dend2[22].pt3dadd(-3.86, -121.64, -1.55, 0.43)
        dend2[22].pt3dadd(-4.11, -123.13, -1.55, 0.43)
        dend2[22].pt3dadd(-4.35, -123.13, -1.55, 0.43)
        dend2[22].pt3dadd(-5.31, -125.12, -1.55, 0.43)
        dend2[22].pt3dadd(-5.31, -129.10, -1.55, 0.43)
        dend2[22].pt3dadd(-4.83, -132.09, -1.55, 0.43)
        dend2[22].pt3dadd(-7.00, -135.57, -1.55, 0.43)
        dend2[22].pt3dadd(-7.00, -135.82, -1.55, 0.43)
        dend2[22].pt3dadd(-8.70, -138.56, -1.55, 0.43)
        dend2[22].pt3dadd(-9.18, -139.30, -1.55, 0.43)
        dend2[22].pt3dadd(-11.11, -141.04, -1.55, 0.43)
        dend2[22].pt3dadd(-11.59, -141.29, -1.55, 0.43)
        dend2[22].pt3dadd(-11.35, -145.27, 0.59, 0.43)
        dend2[22].pt3dadd(-10.63, -148.76, 0.59, 0.43)
        dend2[22].pt3dadd(-10.63, -149.01, 0.59, 0.43)
        dend2[22].pt3dadd(-9.42, -149.50, 0.59, 0.43)
        dend2[22].pt3dadd(-8.70, -149.50, 0.59, 0.43)
        dend2[22].pt3dadd(-7.25, -149.75, 0.59, 0.43)
        dend2[22].pt3dadd(-7.00, -150.25, 0.59, 0.43)
        dend2[22].pt3dadd(-6.04, -152.24, 0.59, 0.43)
        dend2[22].pt3dadd(-6.04, -152.49, 0.59, 0.39)
        dend2[22].pt3dadd(-4.83, -153.98, 0.59, 0.39)
        dend2[22].pt3dadd(-4.83, -154.23, 0.59, 0.39)
        dend2[22].pt3dadd(-5.07, -158.46, -2.35, 0.39)
        dend2[22].pt3dadd(-5.07, -160.45, -2.35, 0.39)
        dend2[22].pt3dadd(-4.83, -160.70, -2.35, 0.39)
        dend2[22].pt3dadd(-2.66, -164.18, -2.35, 0.39)
        dend2[22].pt3dadd(-2.66, -164.68, -2.35, 0.39)
        dend2[22].pt3dadd(-1.21, -167.41, -2.35, 0.39)
        dend2[22].pt3dadd(-1.21, -167.91, -2.35, 0.39)
        dend2[22].pt3dadd(-0.00, -170.15, -2.35, 0.39)
        dend2[22].pt3dadd(0.24, -170.40, -2.35, 0.39)
        dend2[22].pt3dadd(0.97, -171.64, -2.35, 0.39)
        dend2[22].pt3dadd(0.72, -172.39, -2.35, 0.39)
        dend2[22].pt3dadd(-0.24, -174.13, -2.35, 0.39)
        dend2[22].pt3dadd(-0.48, -174.88, -2.35, 0.39)
        dend2[22].pt3dadd(-0.48, -175.62, -2.35, 0.39)
        dend2[22].pt3dadd(-0.72, -177.36, -2.35, 0.39)
        dend2[22].pt3dadd(-0.72, -177.36, -2.35, 0.39)

        dend2[23] = h.Section(name='dend2[23]')
        dend2[23].connect(        dend2[21](1), 0)
        dend2[23].pt3dclear()
        dend2[23].pt3dadd(-11.59, -93.53, 1.59, 0.39)
        dend2[23].pt3dadd(-10.87, -99.25, -4.67, 0.39)
        dend2[23].pt3dadd(-10.87, -102.49, -4.67, 0.39)
        dend2[23].pt3dadd(-10.39, -102.74, -4.67, 0.39)
        dend2[23].pt3dadd(-10.14, -105.22, -4.67, 0.39)
        dend2[23].pt3dadd(-10.63, -106.22, -4.67, 0.39)
        dend2[23].pt3dadd(-12.32, -108.21, -4.67, 0.39)
        dend2[23].pt3dadd(-12.80, -108.46, -4.67, 0.39)
        dend2[23].pt3dadd(-14.98, -108.71, -4.67, 0.39)
        dend2[23].pt3dadd(-16.43, -109.20, -4.67, 0.39)
        dend2[23].pt3dadd(-18.12, -109.95, -4.67, 0.39)
        dend2[23].pt3dadd(-19.08, -110.70, -4.67, 0.39)
        dend2[23].pt3dadd(-19.08, -111.69, -4.67, 0.39)
        dend2[23].pt3dadd(-19.81, -112.94, -4.67, 0.39)
        dend2[23].pt3dadd(-20.53, -114.93, -4.67, 0.39)
        dend2[23].pt3dadd(-20.53, -116.42, -4.67, 0.39)
        dend2[23].pt3dadd(-20.77, -118.66, -4.67, 0.39)
        dend2[23].pt3dadd(-21.26, -119.65, -4.67, 0.39)
        dend2[23].pt3dadd(-21.50, -121.14, -4.67, 0.39)
        dend2[23].pt3dadd(-21.98, -121.89, -4.67, 0.39)
        dend2[23].pt3dadd(-21.98, -124.13, -4.67, 0.39)
        dend2[23].pt3dadd(-21.98, -125.12, -4.67, 0.39)
        dend2[23].pt3dadd(-22.46, -127.61, -4.67, 0.39)
        dend2[23].pt3dadd(-22.95, -128.36, -4.67, 0.39)
        dend2[23].pt3dadd(-25.12, -131.09, -4.67, 0.39)
        dend2[23].pt3dadd(-25.85, -132.09, -4.67, 0.39)
        dend2[23].pt3dadd(-27.29, -132.34, -4.67, 0.39)
        dend2[23].pt3dadd(-28.50, -132.34, -4.67, 0.39)
        dend2[23].pt3dadd(-30.19, -131.84, -4.67, 0.39)
        dend2[23].pt3dadd(-31.40, -131.84, -4.67, 0.39)
        dend2[23].pt3dadd(-35.51, -129.85, -4.67, 0.39)
        dend2[23].pt3dadd(-36.47, -130.10, -4.67, 0.39)
        dend2[23].pt3dadd(-37.92, -130.85, -4.67, 0.39)
        dend2[23].pt3dadd(-39.61, -131.09, -4.67, 0.39)
        dend2[23].pt3dadd(-41.06, -130.10, -4.67, 0.39)
        dend2[23].pt3dadd(-44.69, -129.85, -4.67, 0.39)
        dend2[23].pt3dadd(-45.65, -130.35, -4.67, 0.39)
        dend2[23].pt3dadd(-45.89, -131.59, -4.67, 0.39)
        dend2[23].pt3dadd(-46.62, -132.59, -4.67, 0.39)
        dend2[23].pt3dadd(-47.58, -132.84, -4.67, 0.39)
        dend2[23].pt3dadd(-48.31, -133.08, -4.67, 0.39)

        dend2[24] = h.Section(name='dend2[24]')
        dend2[24].connect(        dend2[20](1), 0)
        dend2[24].pt3dclear()
        dend2[24].pt3dadd(-13.29, -43.53, -2.55, 0.54)
        dend2[24].pt3dadd(-16.18, -48.51, -0.99, 0.54)
        dend2[24].pt3dadd(-16.43, -48.51, -0.99, 0.54)
        dend2[24].pt3dadd(-17.39, -51.74, -0.99, 0.54)
        dend2[24].pt3dadd(-17.63, -51.99, -0.99, 0.54)

        dend2[25] = h.Section(name='dend2[25]')
        dend2[25].connect(        dend2[24](1), 0)
        dend2[25].pt3dclear()
        dend2[25].pt3dadd(-17.63, -51.99, -0.99, 0.43)
        dend2[25].pt3dadd(-19.57, -58.21, 0.08, 0.43)
        dend2[25].pt3dadd(-21.50, -61.69, 0.08, 0.43)
        dend2[25].pt3dadd(-21.74, -62.44, 0.08, 0.43)
        dend2[25].pt3dadd(-23.19, -65.17, 0.08, 0.43)
        dend2[25].pt3dadd(-23.43, -65.67, 0.08, 0.43)
        dend2[25].pt3dadd(-25.36, -69.90, 0.08, 0.43)
        dend2[25].pt3dadd(-25.36, -70.40, 0.08, 0.43)
        dend2[25].pt3dadd(-26.57, -72.39, 0.08, 0.43)
        dend2[25].pt3dadd(-26.81, -72.39, 0.08, 0.43)
        dend2[25].pt3dadd(-27.78, -74.63, -10.32, 0.43)
        dend2[25].pt3dadd(-27.78, -75.12, -10.32, 0.43)
        dend2[25].pt3dadd(-27.54, -76.37, -10.32, 0.43)
        dend2[25].pt3dadd(-27.29, -77.11, -10.32, 0.43)
        dend2[25].pt3dadd(-24.40, -84.33, -19.02, 0.43)
        dend2[25].pt3dadd(-25.85, -91.04, -19.02, 0.43)
        dend2[25].pt3dadd(-26.09, -96.02, -19.02, 0.43)
        dend2[25].pt3dadd(-26.57, -99.01, -19.02, 0.43)
        dend2[25].pt3dadd(-26.57, -99.50, -19.02, 0.43)
        dend2[25].pt3dadd(-26.81, -102.49, -19.02, 0.43)
        dend2[25].pt3dadd(-26.81, -102.74, -19.02, 0.43)
        dend2[25].pt3dadd(-27.05, -103.48, -19.02, 0.43)
        dend2[25].pt3dadd(-26.09, -109.45, -23.38, 0.43)
        dend2[25].pt3dadd(-25.12, -110.20, -23.38, 0.43)
        dend2[25].pt3dadd(-24.88, -110.45, -23.38, 0.43)
        dend2[25].pt3dadd(-24.64, -111.69, -23.38, 0.43)
        dend2[25].pt3dadd(-24.64, -111.94, -23.38, 0.43)
        dend2[25].pt3dadd(-25.85, -114.43, -23.38, 0.43)
        dend2[25].pt3dadd(-25.85, -114.68, -23.38, 0.43)
        dend2[25].pt3dadd(-26.33, -117.41, -23.38, 0.43)
        dend2[25].pt3dadd(-26.57, -118.16, -23.38, 0.43)
        dend2[25].pt3dadd(-26.57, -119.65, -23.38, 0.43)
        dend2[25].pt3dadd(-26.57, -120.40, -23.38, 0.43)
        dend2[25].pt3dadd(-25.85, -122.64, -23.38, 0.43)
        dend2[25].pt3dadd(-25.60, -122.64, -23.39, 0.43)
        dend2[25].pt3dadd(-24.64, -122.89, -23.39, 0.43)
        dend2[25].pt3dadd(-24.40, -122.89, -23.39, 0.43)
        dend2[25].pt3dadd(-16.18, -124.63, -23.35, 0.43)
        dend2[25].pt3dadd(-16.18, -124.63, -23.38, 0.43)
        dend2[25].pt3dadd(-7.73, -127.86, -30.27, 0.39)
        dend2[25].pt3dadd(1.45, -126.12, -31.52, 0.39)
        dend2[25].pt3dadd(1.45, -126.12, -31.54, 0.39)
        dend2[25].pt3dadd(3.62, -129.10, -31.73, 0.39)
        dend2[25].pt3dadd(3.62, -129.10, -31.84, 0.39)
        dend2[25].pt3dadd(7.73, -131.84, -33.25, 0.39)
        dend2[25].pt3dadd(8.21, -131.84, -33.25, 0.39)
        dend2[25].pt3dadd(11.11, -131.59, -33.25, 0.39)
        dend2[25].pt3dadd(11.59, -131.34, -33.25, 0.39)
        dend2[25].pt3dadd(13.04, -130.60, -33.25, 0.39)

        dend2[26] = h.Section(name='dend2[26]')
        dend2[26].connect(        dend2[24](1), 0)
        dend2[26].pt3dclear()
        dend2[26].pt3dadd(-17.63, -51.99, -0.99, 0.54)
        dend2[26].pt3dadd(-19.32, -52.99, -0.99, 0.54)
        dend2[26].pt3dadd(-20.53, -54.48, -0.99, 0.54)
        dend2[26].pt3dadd(-20.77, -54.48, -0.99, 0.54)
        dend2[26].pt3dadd(-25.85, -55.22, -9.73, 0.54)
        dend2[26].pt3dadd(-27.78, -58.46, -9.73, 0.54)
        dend2[26].pt3dadd(-28.02, -58.71, -9.73, 0.54)
        dend2[26].pt3dadd(-29.95, -60.95, -9.73, 0.54)

        dend2[27] = h.Section(name='dend2[27]')
        dend2[27].connect(        dend2[26](1), 0)
        dend2[27].pt3dclear()
        dend2[27].pt3dadd(-29.95, -60.95, -9.73, 0.39)
        dend2[27].pt3dadd(-31.16, -68.16, -13.51, 0.39)
        dend2[27].pt3dadd(-31.40, -68.16, -13.51, 0.39)
        dend2[27].pt3dadd(-31.88, -72.64, -13.51, 0.39)
        dend2[27].pt3dadd(-31.88, -73.13, -13.51, 0.39)
        dend2[27].pt3dadd(-33.09, -75.37, -13.51, 0.39)
        dend2[27].pt3dadd(-33.33, -76.12, -13.51, 0.39)
        dend2[27].pt3dadd(-34.54, -77.86, -13.51, 0.39)
        dend2[27].pt3dadd(-35.27, -79.60, -13.51, 0.39)
        dend2[27].pt3dadd(-35.75, -79.85, -13.51, 0.39)
        dend2[27].pt3dadd(-36.47, -80.60, -13.51, 0.39)
        dend2[27].pt3dadd(-37.92, -81.59, -13.51, 0.39)
        dend2[27].pt3dadd(-38.41, -81.84, -13.51, 0.39)
        dend2[27].pt3dadd(-40.34, -84.33, -17.74, 0.39)
        dend2[27].pt3dadd(-40.82, -84.33, -17.75, 0.39)
        dend2[27].pt3dadd(-42.51, -86.57, -17.75, 0.39)
        dend2[27].pt3dadd(-43.24, -87.06, -17.75, 0.39)
        dend2[27].pt3dadd(-44.20, -88.81, -17.75, 0.39)
        dend2[27].pt3dadd(-44.44, -89.05, -17.75, 0.39)
        dend2[27].pt3dadd(-47.34, -92.29, -17.75, 0.39)
        dend2[27].pt3dadd(-47.34, -93.03, -17.75, 0.39)
        dend2[27].pt3dadd(-51.93, -96.77, -17.75, 0.39)
        dend2[27].pt3dadd(-53.14, -97.51, -17.75, 0.39)
        dend2[27].pt3dadd(-54.11, -98.01, -17.75, 0.39)
        dend2[27].pt3dadd(-54.83, -98.51, -17.75, 0.39)

        dend2[28] = h.Section(name='dend2[28]')
        dend2[28].connect(        dend2[26](1), 0)
        dend2[28].pt3dclear()
        dend2[28].pt3dadd(-29.95, -60.95, -9.73, 0.48)
        dend2[28].pt3dadd(-31.64, -62.69, -5.89, 0.48)
        dend2[28].pt3dadd(-31.88, -62.69, -5.89, 0.48)
        dend2[28].pt3dadd(-35.02, -65.42, -5.89, 0.48)
        dend2[28].pt3dadd(-35.27, -65.42, -5.89, 0.48)
        dend2[28].pt3dadd(-36.47, -64.68, -5.88, 0.48)
        dend2[28].pt3dadd(-37.44, -64.68, -5.88, 0.48)
        dend2[28].pt3dadd(-39.61, -64.68, -5.88, 0.48)
        dend2[28].pt3dadd(-39.86, -64.93, -5.88, 0.48)
        dend2[28].pt3dadd(-40.82, -67.41, -5.88, 0.48)
        dend2[28].pt3dadd(-41.55, -68.16, -5.87, 0.48)
        dend2[28].pt3dadd(-43.48, -69.90, -5.87, 0.48)
        dend2[28].pt3dadd(-44.20, -70.65, -5.87, 0.48)
        dend2[28].pt3dadd(-45.17, -72.39, -5.87, 0.48)
        dend2[28].pt3dadd(-45.17, -73.38, -5.87, 0.48)

        dend2[29] = h.Section(name='dend2[29]')
        dend2[29].connect(        dend2[28](1), 0)
        dend2[29].pt3dclear()
        dend2[29].pt3dadd(-45.17, -73.38, -5.87, 0.39)
        dend2[29].pt3dadd(-39.13, -80.35, 4.09, 0.39)
        dend2[29].pt3dadd(-39.13, -84.58, 4.10, 0.39)
        dend2[29].pt3dadd(-39.13, -85.32, 4.10, 0.39)
        dend2[29].pt3dadd(-38.16, -90.80, 4.10, 0.39)
        dend2[29].pt3dadd(-37.92, -91.29, 4.10, 0.39)
        dend2[29].pt3dadd(-36.71, -92.54, 4.10, 0.39)
        dend2[29].pt3dadd(-36.47, -92.79, 4.10, 0.39)
        dend2[29].pt3dadd(-35.02, -97.26, 7.44, 0.39)
        dend2[29].pt3dadd(-35.27, -101.00, 7.46, 0.39)
        dend2[29].pt3dadd(-35.02, -102.74, 7.47, 0.39)
        dend2[29].pt3dadd(-35.02, -102.74, 7.48, 0.39)
        dend2[29].pt3dadd(-34.30, -104.73, 7.48, 0.39)
        dend2[29].pt3dadd(-33.82, -105.47, 7.49, 0.39)
        dend2[29].pt3dadd(-32.37, -106.47, 7.49, 0.39)
        dend2[29].pt3dadd(-32.13, -106.97, 7.50, 0.39)
        dend2[29].pt3dadd(-30.68, -108.46, 7.50, 0.39)
        dend2[29].pt3dadd(-30.68, -109.20, 7.50, 0.39)
        dend2[29].pt3dadd(-29.95, -110.95, 7.50, 0.39)
        dend2[29].pt3dadd(-29.95, -111.19, 7.50, 0.39)
        dend2[29].pt3dadd(-30.43, -112.94, 7.50, 0.39)
        dend2[29].pt3dadd(-30.68, -113.43, 7.50, 0.39)
        dend2[29].pt3dadd(-32.37, -114.18, 7.50, 0.39)
        dend2[29].pt3dadd(-33.09, -114.68, 7.51, 0.39)
        dend2[29].pt3dadd(-33.57, -115.17, 7.51, 0.39)
        dend2[29].pt3dadd(-34.30, -115.92, 7.51, 0.39)
        dend2[29].pt3dadd(-34.78, -116.67, 7.51, 0.39)
        dend2[29].pt3dadd(-35.27, -117.16, 7.51, 0.39)

        dend2[30] = h.Section(name='dend2[30]')
        dend2[30].connect(        dend2[28](1), 0)
        dend2[30].pt3dclear()
        dend2[30].pt3dadd(-45.17, -73.38, -5.87, 0.39)
        dend2[30].pt3dadd(-47.34, -79.85, -5.87, 0.39)
        dend2[30].pt3dadd(-47.83, -80.10, -5.87, 0.39)
        dend2[30].pt3dadd(-50.72, -82.34, -5.87, 0.39)
        dend2[30].pt3dadd(-51.21, -83.08, -5.87, 0.39)
        dend2[30].pt3dadd(-53.38, -85.32, -5.87, 0.39)
        dend2[30].pt3dadd(-53.86, -85.82, -5.87, 0.39)
        dend2[30].pt3dadd(-56.76, -86.57, -10.73, 0.39)
        dend2[30].pt3dadd(-56.76, -87.06, -10.73, 0.39)
        dend2[30].pt3dadd(-56.76, -91.79, -10.73, 0.39)
        dend2[30].pt3dadd(-57.00, -92.29, -10.73, 0.39)
        dend2[30].pt3dadd(-59.66, -95.52, -10.74, 0.39)
        dend2[30].pt3dadd(-59.90, -96.02, -10.74, 0.39)
        dend2[30].pt3dadd(-61.35, -97.76, -10.74, 0.39)
        dend2[30].pt3dadd(-61.59, -98.01, -10.74, 0.39)
        dend2[30].pt3dadd(-65.70, -99.50, -11.70, 0.39)
        dend2[30].pt3dadd(-65.94, -99.50, -11.70, 0.39)
        dend2[30].pt3dadd(-67.87, -100.75, -11.70, 0.39)
        dend2[30].pt3dadd(-68.36, -101.49, -11.70, 0.39)
        dend2[30].pt3dadd(-69.57, -103.23, -11.70, 0.39)
        dend2[30].pt3dadd(-69.81, -103.23, -11.70, 0.39)
        dend2[30].pt3dadd(-73.19, -104.48, -11.70, 0.39)
        dend2[30].pt3dadd(-73.67, -104.98, -11.70, 0.39)
        dend2[30].pt3dadd(-77.29, -105.97, -11.70, 0.39)
        dend2[30].pt3dadd(-82.13, -106.97, -17.40, 0.39)
        dend2[30].pt3dadd(-82.61, -107.46, -17.31, 0.39)
        dend2[30].pt3dadd(-85.27, -107.96, -17.17, 0.39)
        dend2[30].pt3dadd(-85.75, -108.46, -17.17, 0.39)
        dend2[30].pt3dadd(-85.99, -109.45, -17.15, 0.39)
        dend2[30].pt3dadd(-86.47, -109.45, -17.14, 0.39)

        dend3 = [None]*15

        dend3[0] = h.Section(name='dend3[0]')
        dend3[0].connect(        soma(0.443475), 0)
        dend3[0].pt3dclear()
        dend3[0].pt3dadd(8.21, -1.00, -8.20, 1.94)
        dend3[0].pt3dadd(9.18, -1.00, -8.20, 1.94)
        dend3[0].pt3dadd(10.39, -1.49, -8.20, 1.94)
        dend3[0].pt3dadd(10.87, -1.49, -8.20, 1.94)
        dend3[0].pt3dadd(12.08, -1.99, -8.20, 1.94)
        dend3[0].pt3dadd(12.32, -1.99, -8.20, 1.94)
        dend3[0].pt3dadd(12.80, -2.24, -8.20, 1.94)

        dend3[1] = h.Section(name='dend3[1]')
        dend3[1].connect(        dend3[0](1), 0)
        dend3[1].pt3dclear()
        dend3[1].pt3dadd(12.80, -2.24, -8.20, 0.77)
        dend3[1].pt3dadd(14.25, -6.47, -4.76, 0.77)
        dend3[1].pt3dadd(16.43, -6.97, -4.76, 0.77)
        dend3[1].pt3dadd(17.15, -6.97, -4.76, 0.77)
        dend3[1].pt3dadd(17.63, -8.71, -4.76, 0.77)
        dend3[1].pt3dadd(18.60, -8.71, -4.76, 0.77)

        dend3[2] = h.Section(name='dend3[2]')
        dend3[2].connect(        dend3[1](1), 0)
        dend3[2].pt3dclear()
        dend3[2].pt3dadd(18.60, -8.71, -4.76, 0.54)
        dend3[2].pt3dadd(18.84, -12.19, 4.21, 0.54)
        dend3[2].pt3dadd(16.67, -15.17, 4.21, 0.54)
        dend3[2].pt3dadd(17.15, -15.17, 4.21, 0.54)
        dend3[2].pt3dadd(17.39, -17.91, 4.21, 0.54)
        dend3[2].pt3dadd(17.63, -18.41, 4.21, 0.54)
        dend3[2].pt3dadd(18.84, -20.90, 4.21, 0.54)
        dend3[2].pt3dadd(19.08, -21.39, 4.21, 0.54)
        dend3[2].pt3dadd(20.29, -27.61, -5.02, 0.54)
        dend3[2].pt3dadd(21.74, -30.10, -5.03, 0.54)
        dend3[2].pt3dadd(22.71, -30.10, -5.03, 0.54)
        dend3[2].pt3dadd(24.64, -32.34, -5.03, 0.54)
        dend3[2].pt3dadd(24.88, -32.84, -5.03, 0.54)
        dend3[2].pt3dadd(25.36, -35.32, -5.03, 0.54)
        dend3[2].pt3dadd(25.60, -38.06, -5.03, 0.54)
        dend3[2].pt3dadd(26.57, -39.80, -5.03, 0.54)
        dend3[2].pt3dadd(26.81, -40.30, -5.03, 0.54)
        dend3[2].pt3dadd(28.50, -42.79, -5.03, 0.54)
        dend3[2].pt3dadd(28.74, -43.78, -5.03, 0.54)
        dend3[2].pt3dadd(30.19, -45.52, -5.03, 0.43)
        dend3[2].pt3dadd(30.19, -45.77, -5.03, 0.43)
        dend3[2].pt3dadd(30.92, -46.77, -5.03, 0.43)
        dend3[2].pt3dadd(33.09, -56.97, 7.34, 0.43)
        dend3[2].pt3dadd(34.30, -58.71, 7.34, 0.43)
        dend3[2].pt3dadd(34.54, -58.96, 7.34, 0.43)
        dend3[2].pt3dadd(35.99, -64.93, 7.34, 0.43)
        dend3[2].pt3dadd(39.37, -73.38, 13.47, 0.43)
        dend3[2].pt3dadd(42.27, -75.62, 13.47, 0.43)
        dend3[2].pt3dadd(42.27, -75.87, 13.47, 0.43)
        dend3[2].pt3dadd(43.24, -78.36, 13.47, 0.43)
        dend3[2].pt3dadd(43.24, -78.61, 13.48, 0.43)
        dend3[2].pt3dadd(43.48, -80.60, 13.50, 0.43)
        dend3[2].pt3dadd(43.24, -80.85, 13.50, 0.43)
        dend3[2].pt3dadd(43.72, -83.58, 13.51, 0.43)
        dend3[2].pt3dadd(43.96, -83.83, 13.51, 0.43)
        dend3[2].pt3dadd(44.69, -85.57, 13.51, 0.43)
        dend3[2].pt3dadd(44.69, -86.57, 13.51, 0.43)

        dend3[3] = h.Section(name='dend3[3]')
        dend3[3].connect(        dend3[1](1), 0)
        dend3[3].pt3dclear()
        dend3[3].pt3dadd(18.60, -8.71, -4.76, 0.54)
        dend3[3].pt3dadd(20.53, -9.45, -12.28, 0.54)
        dend3[3].pt3dadd(23.67, -9.95, -12.28, 0.54)
        dend3[3].pt3dadd(24.15, -10.20, -12.28, 0.54)
        dend3[3].pt3dadd(25.60, -10.70, -12.28, 0.54)
        dend3[3].pt3dadd(25.85, -11.19, -12.28, 0.54)
        dend3[3].pt3dadd(27.05, -13.43, -12.28, 0.54)
        dend3[3].pt3dadd(27.05, -13.68, -12.28, 0.54)

        dend3[4] = h.Section(name='dend3[4]')
        dend3[4].connect(        dend3[3](1), 0)
        dend3[4].pt3dclear()
        dend3[4].pt3dadd(27.05, -13.68, -12.28, 0.43)
        dend3[4].pt3dadd(28.99, -18.16, -19.38, 0.43)
        dend3[4].pt3dadd(28.99, -18.16, -19.37, 0.43)
        dend3[4].pt3dadd(30.68, -19.40, -19.37, 0.43)
        dend3[4].pt3dadd(31.40, -20.65, -19.37, 0.43)
        dend3[4].pt3dadd(32.37, -22.89, -19.37, 0.43)
        dend3[4].pt3dadd(32.61, -23.13, -19.37, 0.43)
        dend3[4].pt3dadd(35.02, -25.12, -19.37, 0.43)
        dend3[4].pt3dadd(35.51, -27.61, -19.37, 0.43)
        dend3[4].pt3dadd(35.99, -27.86, -19.37, 0.43)
        dend3[4].pt3dadd(40.34, -29.10, -19.63, 0.43)
        dend3[4].pt3dadd(40.58, -29.10, -19.63, 0.43)
        dend3[4].pt3dadd(45.41, -30.85, -19.63, 0.43)
        dend3[4].pt3dadd(46.62, -35.32, -19.63, 0.43)
        dend3[4].pt3dadd(46.62, -35.57, -19.63, 0.43)
        dend3[4].pt3dadd(48.31, -39.05, -23.24, 0.43)
        dend3[4].pt3dadd(50.97, -40.80, -23.24, 0.43)
        dend3[4].pt3dadd(51.45, -41.29, -23.24, 0.43)
        dend3[4].pt3dadd(53.38, -46.27, -23.24, 0.43)
        dend3[4].pt3dadd(54.11, -48.76, -23.23, 0.43)
        dend3[4].pt3dadd(54.35, -49.50, -23.23, 0.43)
        dend3[4].pt3dadd(56.04, -52.99, -23.23, 0.43)
        dend3[4].pt3dadd(56.52, -56.72, -23.23, 0.43)
        dend3[4].pt3dadd(57.25, -57.46, -23.23, 0.43)
        dend3[4].pt3dadd(57.73, -58.96, -23.23, 0.43)
        dend3[4].pt3dadd(60.87, -62.19, -23.23, 0.43)
        dend3[4].pt3dadd(61.59, -62.19, -23.23, 0.43)
        dend3[4].pt3dadd(62.56, -62.69, -23.23, 0.43)
        dend3[4].pt3dadd(62.80, -62.94, -23.23, 0.43)
        dend3[4].pt3dadd(64.98, -64.68, -23.23, 0.43)
        dend3[4].pt3dadd(65.70, -65.42, -23.23, 0.43)
        dend3[4].pt3dadd(71.74, -71.39, -23.23, 0.43)
        dend3[4].pt3dadd(73.91, -72.89, -23.23, 0.43)
        dend3[4].pt3dadd(74.40, -73.63, -23.23, 0.43)
        dend3[4].pt3dadd(76.09, -78.36, -23.23, 0.43)
        dend3[4].pt3dadd(76.09, -78.61, -23.23, 0.43)
        dend3[4].pt3dadd(81.40, -88.81, -23.82, 0.43)
        dend3[4].pt3dadd(84.06, -93.28, -23.82, 0.43)
        dend3[4].pt3dadd(84.54, -93.53, -23.82, 0.43)
        dend3[4].pt3dadd(88.41, -96.77, -23.82, 0.43)
        dend3[4].pt3dadd(88.41, -100.25, -26.05, 0.43)
        dend3[4].pt3dadd(88.41, -100.25, -26.07, 0.43)
        dend3[4].pt3dadd(86.23, -102.24, -26.13, 0.43)
        dend3[4].pt3dadd(86.96, -105.47, -26.13, 0.43)
        dend3[4].pt3dadd(87.20, -105.97, -26.13, 0.43)
        dend3[4].pt3dadd(88.65, -107.21, -26.13, 0.43)
        dend3[4].pt3dadd(86.96, -108.71, -26.13, 0.43)
        dend3[4].pt3dadd(85.75, -109.95, -26.13, 0.43)
        dend3[4].pt3dadd(85.27, -110.45, -26.13, 0.43)
        dend3[4].pt3dadd(84.06, -111.44, -26.13, 0.43)
        dend3[4].pt3dadd(83.33, -112.19, -26.13, 0.43)
        dend3[4].pt3dadd(83.09, -112.44, -26.13, 0.43)

        dend3[5] = h.Section(name='dend3[5]')
        dend3[5].connect(        dend3[3](1), 0)
        dend3[5].pt3dclear()
        dend3[5].pt3dadd(27.05, -13.68, -12.28, 0.54)
        dend3[5].pt3dadd(30.33, -15.12, -14.07, 0.54)
        dend3[5].pt3dadd(32.99, -17.61, -14.07, 0.54)
        dend3[5].pt3dadd(37.33, -17.61, -14.06, 0.54)

        dend3[6] = h.Section(name='dend3[6]')
        dend3[6].connect(        dend3[5](1), 0)
        dend3[6].pt3dclear()
        dend3[6].pt3dadd(37.33, -17.61, -14.06, 0.43)
        dend3[6].pt3dadd(44.34, -22.33, -17.46, 0.43)
        dend3[6].pt3dadd(46.27, -25.82, -17.46, 0.43)
        dend3[6].pt3dadd(51.10, -32.03, -17.45, 0.43)
        dend3[6].pt3dadd(51.34, -32.03, -17.45, 0.43)
        dend3[6].pt3dadd(53.76, -33.78, -17.45, 0.43)
        dend3[6].pt3dadd(54.24, -34.02, -17.44, 0.43)
        dend3[6].pt3dadd(60.28, -39.75, -14.56, 0.43)
        dend3[6].pt3dadd(63.90, -40.49, -14.56, 0.43)
        dend3[6].pt3dadd(70.43, -43.23, -12.71, 0.43)
        dend3[6].pt3dadd(70.43, -43.23, -12.70, 0.43)
        dend3[6].pt3dadd(71.39, -45.22, -12.70, 0.43)
        dend3[6].pt3dadd(72.60, -47.95, -12.70, 0.43)
        dend3[6].pt3dadd(75.26, -50.69, -12.70, 0.43)
        dend3[6].pt3dadd(75.74, -50.69, -12.70, 0.43)
        dend3[6].pt3dadd(77.43, -52.68, -12.70, 0.43)
        dend3[6].pt3dadd(77.91, -53.43, -12.70, 0.43)
        dend3[6].pt3dadd(79.36, -55.42, -12.70, 0.39)
        dend3[6].pt3dadd(80.33, -59.15, -12.70, 0.39)
        dend3[6].pt3dadd(82.99, -61.64, -12.70, 0.39)
        dend3[6].pt3dadd(83.95, -63.87, -12.70, 0.39)
        dend3[6].pt3dadd(84.19, -63.87, -12.70, 0.39)
        dend3[6].pt3dadd(88.78, -69.60, -12.61, 0.39)
        dend3[6].pt3dadd(89.51, -73.58, -12.62, 0.39)

        dend3[7] = h.Section(name='dend3[7]')
        dend3[7].connect(        dend3[5](1), 0)
        dend3[7].pt3dclear()
        dend3[7].pt3dadd(37.33, -17.61, -14.06, 0.43)
        dend3[7].pt3dadd(42.41, -16.86, -13.07, 0.43)
        dend3[7].pt3dadd(49.89, -15.86, -13.07, 0.43)
        dend3[7].pt3dadd(50.14, -15.86, -13.07, 0.43)
        dend3[7].pt3dadd(56.90, -13.63, -13.07, 0.43)
        dend3[7].pt3dadd(57.14, -13.63, -13.07, 0.43)
        dend3[7].pt3dadd(62.70, -13.38, -13.07, 0.43)
        dend3[7].pt3dadd(62.94, -13.38, -13.07, 0.43)
        dend3[7].pt3dadd(66.80, -11.64, -13.07, 0.43)
        dend3[7].pt3dadd(67.04, -11.64, -13.07, 0.43)
        dend3[7].pt3dadd(70.91, -11.64, -13.07, 0.43)
        dend3[7].pt3dadd(71.39, -11.64, -13.07, 0.43)
        dend3[7].pt3dadd(74.77, -11.88, -13.07, 0.43)
        dend3[7].pt3dadd(75.50, -11.88, -13.07, 0.43)
        dend3[7].pt3dadd(78.40, -10.89, -13.07, 0.43)
        dend3[7].pt3dadd(79.85, -10.89, -13.07, 0.43)
        dend3[7].pt3dadd(83.47, -10.89, -13.07, 0.43)
        dend3[7].pt3dadd(83.95, -10.89, -13.07, 0.43)
        dend3[7].pt3dadd(89.51, -9.89, -14.40, 0.43)
        dend3[7].pt3dadd(94.10, -8.65, -14.40, 0.43)
        dend3[7].pt3dadd(95.06, -8.40, -14.40, 0.43)
        dend3[7].pt3dadd(100.38, -8.90, -14.40, 0.43)
        dend3[7].pt3dadd(101.83, -8.90, -14.40, 0.43)
        dend3[7].pt3dadd(103.76, -11.64, -14.40, 0.43)
        dend3[7].pt3dadd(104.00, -11.64, -14.40, 0.43)
        dend3[7].pt3dadd(106.66, -11.64, -14.40, 0.43)
        dend3[7].pt3dadd(107.14, -11.64, -14.40, 0.43)
        dend3[7].pt3dadd(113.90, -11.14, -14.40, 0.43)
        dend3[7].pt3dadd(117.77, -10.89, -14.40, 0.43)
        dend3[7].pt3dadd(119.70, -10.64, -14.40, 0.43)
        dend3[7].pt3dadd(120.91, -10.39, -14.40, 0.43)
        dend3[7].pt3dadd(124.77, -9.65, -14.40, 0.43)
        dend3[7].pt3dadd(125.50, -9.40, -14.40, 0.43)
        dend3[7].pt3dadd(126.22, -9.40, -14.40, 0.43)
        dend3[7].pt3dadd(126.46, -9.15, -14.40, 0.43)
        dend3[7].pt3dadd(126.95, -9.15, -14.40, 0.43)
        dend3[7].pt3dadd(129.36, -8.65, -14.40, 0.43)

        dend3[8] = h.Section(name='dend3[8]')
        dend3[8].connect(        dend3[0](1), 0)
        dend3[8].pt3dclear()
        dend3[8].pt3dadd(12.80, -2.24, -8.20, 0.77)
        dend3[8].pt3dadd(15.59, -2.93, -8.97, 0.77)
        dend3[8].pt3dadd(17.53, -4.42, -8.97, 0.77)
        dend3[8].pt3dadd(19.22, -5.17, -8.97, 0.77)
        dend3[8].pt3dadd(22.84, -6.91, -8.97, 0.77)
        dend3[8].pt3dadd(23.32, -6.91, -8.97, 0.77)
        dend3[8].pt3dadd(24.77, -6.16, -8.97, 0.77)
        dend3[8].pt3dadd(25.01, -6.16, -8.97, 0.77)
        dend3[8].pt3dadd(26.46, -5.17, -8.97, 0.77)
        dend3[8].pt3dadd(27.19, -5.17, -8.97, 0.77)
        dend3[8].pt3dadd(27.67, -5.17, -8.97, 0.77)

        dend3[9] = h.Section(name='dend3[9]')
        dend3[9].connect(        dend3[8](1), 0)
        dend3[9].pt3dclear()
        dend3[9].pt3dadd(27.67, -5.17, -8.97, 0.54)
        dend3[9].pt3dadd(29.85, -6.41, -8.25, 0.54)
        dend3[9].pt3dadd(30.81, -7.41, -8.25, 0.54)
        dend3[9].pt3dadd(32.99, -7.16, -8.25, 0.54)
        dend3[9].pt3dadd(33.47, -7.16, -8.25, 0.54)

        dend3[10] = h.Section(name='dend3[10]')
        dend3[10].connect(        dend3[9](1), 0)
        dend3[10].pt3dclear()
        dend3[10].pt3dadd(33.47, -7.16, -8.25, 0.43)
        dend3[10].pt3dadd(32.50, -11.39, 2.39, 0.43)
        dend3[10].pt3dadd(35.40, -12.63, 2.39, 0.43)
        dend3[10].pt3dadd(35.64, -12.88, 2.39, 0.43)
        dend3[10].pt3dadd(41.92, -16.86, -30.16, 0.43)
        dend3[10].pt3dadd(48.44, -21.59, -30.16, 0.43)
        dend3[10].pt3dadd(48.69, -21.59, -30.16, 0.43)
        dend3[10].pt3dadd(51.83, -21.59, -30.16, 0.43)
        dend3[10].pt3dadd(51.83, -21.59, -30.17, 0.43)
        dend3[10].pt3dadd(54.97, -23.33, -30.17, 0.43)
        dend3[10].pt3dadd(55.45, -23.33, -30.19, 0.43)
        dend3[10].pt3dadd(58.83, -26.31, -30.26, 0.43)
        dend3[10].pt3dadd(60.04, -26.31, -30.30, 0.43)
        dend3[10].pt3dadd(62.94, -27.31, -30.48, 0.43)
        dend3[10].pt3dadd(63.42, -27.81, -30.50, 0.43)
        dend3[10].pt3dadd(78.64, -37.76, -21.09, 0.43)
        dend3[10].pt3dadd(78.64, -37.76, -21.08, 0.43)
        dend3[10].pt3dadd(85.16, -44.47, -21.07, 0.43)
        dend3[10].pt3dadd(92.16, -45.96, -21.07, 0.43)
        dend3[10].pt3dadd(93.37, -45.96, -21.07, 0.43)
        dend3[10].pt3dadd(96.51, -46.21, -21.07, 0.43)
        dend3[10].pt3dadd(97.24, -46.21, -21.07, 0.43)
        dend3[10].pt3dadd(100.86, -46.46, -21.07, 0.43)
        dend3[10].pt3dadd(101.83, -46.46, -21.07, 0.43)
        dend3[10].pt3dadd(103.52, -46.46, -21.07, 0.43)
        dend3[10].pt3dadd(103.76, -46.46, -21.07, 0.43)

        dend3[11] = h.Section(name='dend3[11]')
        dend3[11].connect(        dend3[9](1), 0)
        dend3[11].pt3dclear()
        dend3[11].pt3dadd(33.47, -7.16, -8.25, 0.43)
        dend3[11].pt3dadd(35.64, -6.91, -8.25, 0.43)
        dend3[11].pt3dadd(38.30, -7.41, -8.25, 0.43)
        dend3[11].pt3dadd(38.78, -7.66, -8.25, 0.43)
        dend3[11].pt3dadd(40.23, -7.90, -8.25, 0.43)
        dend3[11].pt3dadd(39.75, -10.14, -8.25, 0.43)
        dend3[11].pt3dadd(39.02, -10.89, -8.25, 0.43)
        dend3[11].pt3dadd(38.54, -11.14, -8.25, 0.43)
        dend3[11].pt3dadd(38.30, -12.13, -8.25, 0.43)
        dend3[11].pt3dadd(39.27, -13.38, -8.25, 0.43)
        dend3[11].pt3dadd(39.51, -13.63, -8.25, 0.43)
        dend3[11].pt3dadd(41.20, -14.12, -8.25, 0.43)
        dend3[11].pt3dadd(41.68, -14.62, -8.25, 0.43)
        dend3[11].pt3dadd(43.37, -15.86, -8.25, 0.43)
        dend3[11].pt3dadd(44.10, -16.36, -8.25, 0.43)
        dend3[11].pt3dadd(45.06, -16.61, -8.25, 0.43)
        dend3[11].pt3dadd(46.03, -17.11, -8.25, 0.43)
        dend3[11].pt3dadd(47.00, -17.85, -8.25, 0.43)
        dend3[11].pt3dadd(47.96, -18.10, -8.25, 0.43)
        dend3[11].pt3dadd(48.69, -18.10, -8.25, 0.43)
        dend3[11].pt3dadd(49.65, -18.60, -8.25, 0.43)
        dend3[11].pt3dadd(54.00, -19.35, -8.25, 0.43)
        dend3[11].pt3dadd(54.24, -19.60, -8.25, 0.43)
        dend3[11].pt3dadd(54.97, -19.60, -8.25, 0.43)
        dend3[11].pt3dadd(55.93, -19.84, -8.25, 0.43)
        dend3[11].pt3dadd(57.14, -20.09, -8.25, 0.43)
        dend3[11].pt3dadd(57.38, -20.09, -8.25, 0.43)
        dend3[11].pt3dadd(58.59, -20.59, -8.25, 0.43)
        dend3[11].pt3dadd(59.80, -21.09, -8.25, 0.43)
        dend3[11].pt3dadd(60.52, -21.34, -8.25, 0.43)
        dend3[11].pt3dadd(60.76, -21.34, -8.25, 0.43)
        dend3[11].pt3dadd(61.25, -21.83, -8.25, 0.43)
        dend3[11].pt3dadd(61.49, -21.83, -8.25, 0.43)
        dend3[11].pt3dadd(61.97, -23.08, -8.25, 0.43)
        dend3[11].pt3dadd(61.97, -23.83, -8.25, 0.43)
        dend3[11].pt3dadd(61.97, -25.32, -8.25, 0.43)
        dend3[11].pt3dadd(61.73, -27.56, -8.25, 0.43)
        dend3[11].pt3dadd(61.73, -28.55, -8.25, 0.43)
        dend3[11].pt3dadd(61.01, -30.04, -8.25, 0.39)
        dend3[11].pt3dadd(60.76, -30.29, -8.25, 0.39)
        dend3[11].pt3dadd(60.52, -31.29, -8.25, 0.39)
        dend3[11].pt3dadd(60.76, -31.29, -8.25, 0.39)
        dend3[11].pt3dadd(61.49, -31.79, -8.25, 0.39)
        dend3[11].pt3dadd(62.45, -32.78, -8.25, 0.39)
        dend3[11].pt3dadd(64.63, -35.02, -8.25, 0.39)
        dend3[11].pt3dadd(65.35, -36.26, -8.25, 0.39)
        dend3[11].pt3dadd(65.59, -36.76, -8.25, 0.39)
        dend3[11].pt3dadd(66.32, -37.76, -8.25, 0.39)
        dend3[11].pt3dadd(69.22, -45.22, -14.24, 0.39)
        dend3[11].pt3dadd(70.43, -46.96, -14.24, 0.39)
        dend3[11].pt3dadd(70.67, -47.46, -14.24, 0.39)
        dend3[11].pt3dadd(71.39, -50.69, -14.24, 0.39)
        dend3[11].pt3dadd(71.87, -51.19, -14.24, 0.39)
        dend3[11].pt3dadd(74.29, -53.43, -14.24, 0.39)
        dend3[11].pt3dadd(74.77, -53.92, -14.24, 0.39)
        dend3[11].pt3dadd(75.26, -55.17, -14.24, 0.39)
        dend3[11].pt3dadd(75.50, -55.67, -14.24, 0.39)
        dend3[11].pt3dadd(77.19, -58.15, -14.24, 0.39)
        dend3[11].pt3dadd(77.43, -59.15, -14.24, 0.39)
        dend3[11].pt3dadd(78.64, -60.39, -14.24, 0.39)
        dend3[11].pt3dadd(78.88, -61.14, -14.24, 0.39)
        dend3[11].pt3dadd(80.09, -63.63, -14.24, 0.39)
        dend3[11].pt3dadd(80.09, -64.12, -14.24, 0.39)
        dend3[11].pt3dadd(80.57, -65.12, -14.24, 0.39)
        dend3[11].pt3dadd(80.81, -65.62, -14.24, 0.39)
        dend3[11].pt3dadd(82.50, -66.36, -14.24, 0.39)
        dend3[11].pt3dadd(84.43, -68.35, -14.24, 0.39)
        dend3[11].pt3dadd(84.68, -68.60, -14.24, 0.39)
        dend3[11].pt3dadd(85.16, -69.60, -14.24, 0.39)
        dend3[11].pt3dadd(85.64, -70.59, -14.24, 0.39)
        dend3[11].pt3dadd(85.88, -71.59, -14.24, 0.39)
        dend3[11].pt3dadd(85.88, -72.58, -14.24, 0.39)
        dend3[11].pt3dadd(87.33, -74.07, -14.24, 0.39)
        dend3[11].pt3dadd(88.78, -76.81, -14.24, 0.39)
        dend3[11].pt3dadd(88.54, -79.30, -16.25, 0.39)
        dend3[11].pt3dadd(88.30, -81.54, -16.25, 0.39)
        dend3[11].pt3dadd(88.30, -82.03, -16.25, 0.39)
        dend3[11].pt3dadd(88.30, -83.03, -16.25, 0.39)
        dend3[11].pt3dadd(88.30, -83.28, -16.20, 0.39)
        dend3[11].pt3dadd(94.34, -86.26, -8.81, 0.39)
        dend3[11].pt3dadd(99.41, -88.75, -8.82, 0.39)
        dend3[11].pt3dadd(100.14, -88.75, -8.82, 0.39)
        dend3[11].pt3dadd(104.97, -89.00, -8.82, 0.39)
        dend3[11].pt3dadd(106.17, -89.00, -8.82, 0.39)
        dend3[11].pt3dadd(106.90, -89.50, -8.82, 0.39)

        dend3[12] = h.Section(name='dend3[12]')
        dend3[12].connect(        dend3[8](1), 0)
        dend3[12].pt3dclear()
        dend3[12].pt3dadd(27.67, -5.17, -8.97, 0.54)
        dend3[12].pt3dadd(34.92, -4.17, -4.34, 0.54)
        dend3[12].pt3dadd(37.09, -2.18, -4.34, 0.54)
        dend3[12].pt3dadd(39.27, -1.44, -4.34, 0.54)

        dend3[13] = h.Section(name='dend3[13]')
        dend3[13].connect(        dend3[12](1), 0)
        dend3[13].pt3dclear()
        dend3[13].pt3dadd(39.27, -1.44, -4.34, 0.43)
        dend3[13].pt3dadd(44.58, -4.67, 0.93, 0.43)
        dend3[13].pt3dadd(44.58, -4.67, 0.95, 0.43)
        dend3[13].pt3dadd(48.44, -5.67, 0.95, 0.43)
        dend3[13].pt3dadd(49.41, -5.91, 0.95, 0.43)
        dend3[13].pt3dadd(50.86, -6.66, 0.96, 0.43)
        dend3[13].pt3dadd(51.10, -7.90, 0.96, 0.43)
        dend3[13].pt3dadd(52.31, -8.40, 0.97, 0.43)
        dend3[13].pt3dadd(54.00, -8.90, 0.98, 0.43)
        dend3[13].pt3dadd(55.21, -9.15, 0.98, 0.43)
        dend3[13].pt3dadd(56.90, -9.15, 0.99, 0.43)
        dend3[13].pt3dadd(59.31, -9.65, 1.00, 0.43)
        dend3[13].pt3dadd(60.76, -9.89, 1.02, 0.43)
        dend3[13].pt3dadd(63.90, -8.65, 1.02, 0.43)
        dend3[13].pt3dadd(64.15, -8.65, 1.02, 0.43)
        dend3[13].pt3dadd(66.80, -9.89, 1.02, 0.43)
        dend3[13].pt3dadd(67.29, -10.64, 1.03, 0.43)
        dend3[13].pt3dadd(72.36, -18.60, 5.77, 0.43)
        dend3[13].pt3dadd(72.36, -18.60, 5.78, 0.43)
        dend3[13].pt3dadd(74.53, -22.33, 7.23, 0.43)
        dend3[13].pt3dadd(74.53, -22.33, 7.36, 0.43)
        dend3[13].pt3dadd(76.95, -25.57, 7.71, 0.43)
        dend3[13].pt3dadd(77.19, -25.82, 7.85, 0.43)
        dend3[13].pt3dadd(78.64, -27.31, 8.08, 0.43)
        dend3[13].pt3dadd(78.88, -27.56, 8.11, 0.43)
        dend3[13].pt3dadd(80.09, -29.30, 8.20, 0.43)
        dend3[13].pt3dadd(80.33, -29.80, 8.36, 0.43)
        dend3[13].pt3dadd(79.36, -32.78, 9.74, 0.43)
        dend3[13].pt3dadd(79.36, -32.78, 9.75, 0.43)
        dend3[13].pt3dadd(80.33, -35.27, 9.76, 0.43)
        dend3[13].pt3dadd(81.05, -35.77, 9.77, 0.43)
        dend3[13].pt3dadd(82.74, -37.51, 9.77, 0.43)
        dend3[13].pt3dadd(82.74, -37.51, 9.80, 0.43)
        dend3[13].pt3dadd(80.81, -41.98, 16.48, 0.43)
        dend3[13].pt3dadd(80.81, -41.98, 16.50, 0.43)
        dend3[13].pt3dadd(80.81, -43.48, 16.70, 0.43)
        dend3[13].pt3dadd(80.81, -43.73, 16.70, 0.43)
        dend3[13].pt3dadd(82.74, -45.72, 16.71, 0.39)
        dend3[13].pt3dadd(83.47, -45.96, 16.72, 0.39)
        dend3[13].pt3dadd(86.61, -46.96, 16.72, 0.39)
        dend3[13].pt3dadd(87.58, -46.96, 16.72, 0.39)
        dend3[13].pt3dadd(90.96, -47.71, 16.73, 0.39)
        dend3[13].pt3dadd(91.92, -47.71, 16.74, 0.39)
        dend3[13].pt3dadd(93.86, -47.95, 16.74, 0.39)
        dend3[13].pt3dadd(94.34, -48.20, 16.74, 0.39)
        dend3[13].pt3dadd(96.03, -48.95, 16.74, 0.39)
        dend3[13].pt3dadd(96.27, -48.95, 16.75, 0.39)
        dend3[13].pt3dadd(97.00, -49.45, 16.75, 0.39)
        dend3[13].pt3dadd(97.00, -50.69, 16.75, 0.39)
        dend3[13].pt3dadd(95.55, -51.44, 16.75, 0.39)
        dend3[13].pt3dadd(94.58, -52.43, 16.75, 0.39)
        dend3[13].pt3dadd(94.10, -53.18, 16.75, 0.39)
        dend3[13].pt3dadd(94.58, -55.67, 16.76, 0.39)
        dend3[13].pt3dadd(95.30, -56.16, 16.76, 0.39)
        dend3[13].pt3dadd(96.27, -57.16, 16.77, 0.39)
        dend3[13].pt3dadd(96.75, -57.41, 16.77, 0.39)
        dend3[13].pt3dadd(97.24, -57.90, 16.77, 0.39)
        dend3[13].pt3dadd(97.48, -58.15, 16.77, 0.39)
        dend3[13].pt3dadd(97.48, -59.40, 16.78, 0.39)
        dend3[13].pt3dadd(97.48, -59.40, 16.82, 0.39)

        dend3[14] = h.Section(name='dend3[14]')
        dend3[14].connect(        dend3[12](1), 0)
        dend3[14].pt3dclear()
        dend3[14].pt3dadd(39.27, -1.44, -4.34, 0.39)
        dend3[14].pt3dadd(44.34, 1.55, -2.33, 0.39)
        dend3[14].pt3dadd(49.65, 4.28, -2.33, 0.39)
        dend3[14].pt3dadd(54.24, 6.03, -2.33, 0.39)
        dend3[14].pt3dadd(55.45, 9.51, -2.33, 0.39)
        dend3[14].pt3dadd(56.90, 10.01, -2.33, 0.39)
        dend3[14].pt3dadd(57.62, 11.25, -2.33, 0.39)
        dend3[14].pt3dadd(58.11, 11.50, -2.33, 0.39)
        dend3[14].pt3dadd(59.56, 12.74, -2.33, 0.39)
        dend3[14].pt3dadd(59.56, 12.99, -2.33, 0.39)
        dend3[14].pt3dadd(61.25, 14.48, -2.33, 0.39)
        dend3[14].pt3dadd(61.25, 14.73, -2.33, 0.39)
        dend3[14].pt3dadd(62.94, 18.46, 8.02, 0.39)
        dend3[14].pt3dadd(62.70, 22.19, 8.02, 0.39)
        dend3[14].pt3dadd(64.15, 25.18, 8.10, 0.39)
        dend3[14].pt3dadd(64.63, 25.43, 8.11, 0.39)
        dend3[14].pt3dadd(65.35, 26.42, 8.11, 0.39)
        dend3[14].pt3dadd(66.56, 27.42, 8.12, 0.39)
        dend3[14].pt3dadd(68.01, 30.40, 8.19, 0.39)
        dend3[14].pt3dadd(68.01, 30.65, 8.19, 0.39)
        dend3[14].pt3dadd(68.98, 33.89, 8.20, 0.39)
        dend3[14].pt3dadd(69.46, 34.14, 8.22, 0.39)
        dend3[14].pt3dadd(71.15, 35.38, 8.22, 0.39)
        dend3[14].pt3dadd(71.87, 35.38, 8.22, 0.39)
        dend3[14].pt3dadd(71.87, 38.12, 8.23, 0.39)
        dend3[14].pt3dadd(72.12, 38.86, 8.24, 0.39)
        dend3[14].pt3dadd(72.12, 40.60, 8.24, 0.39)
        dend3[14].pt3dadd(72.60, 41.35, 8.24, 0.39)
        dend3[14].pt3dadd(72.84, 44.33, 8.24, 0.39)
        dend3[14].pt3dadd(73.08, 44.83, 8.24, 0.39)
        dend3[14].pt3dadd(75.74, 46.32, 8.25, 0.39)
        dend3[14].pt3dadd(76.22, 47.82, 8.25, 0.39)
        dend3[14].pt3dadd(76.22, 47.82, 8.26, 0.39)

        dend4 = [None]*13

        dend4[0] = h.Section(name='dend4[0]')
        dend4[0].connect(        soma(1), 0)
        dend4[0].pt3dclear()
        dend4[0].pt3dadd(2.07, -14.87, 10.07, 1.94)
        dend4[0].pt3dadd(2.07, -14.87, 10.06, 1.94)
        dend4[0].pt3dadd(4.72, -16.11, 10.06, 1.94)
        dend4[0].pt3dadd(4.97, -16.11, 10.06, 1.94)
        dend4[0].pt3dadd(8.35, -17.61, 10.06, 1.20)
        dend4[0].pt3dadd(10.04, -18.60, 10.06, 1.20)
        dend4[0].pt3dadd(10.52, -18.60, 10.06, 1.20)
        dend4[0].pt3dadd(12.21, -20.34, 2.26, 1.20)

        dend4[1] = h.Section(name='dend4[1]')
        dend4[1].connect(        dend4[0](1), 0)
        dend4[1].pt3dclear()
        dend4[1].pt3dadd(12.21, -20.34, 2.26, 0.48)
        dend4[1].pt3dadd(14.15, -23.33, 8.99, 0.48)
        dend4[1].pt3dadd(17.04, -21.83, 8.99, 0.48)
        dend4[1].pt3dadd(17.53, -21.59, 9.00, 0.48)
        dend4[1].pt3dadd(18.73, -21.34, 9.12, 0.48)
        dend4[1].pt3dadd(19.22, -21.09, 9.21, 0.48)
        dend4[1].pt3dadd(20.18, -22.83, 9.56, 0.48)
        dend4[1].pt3dadd(20.43, -22.83, 9.69, 0.48)
        dend4[1].pt3dadd(21.87, -23.33, 9.94, 0.48)
        dend4[1].pt3dadd(22.60, -23.33, 10.10, 0.48)
        dend4[1].pt3dadd(24.29, -23.33, 11.38, 0.48)
        dend4[1].pt3dadd(24.29, -23.33, 11.43, 0.48)
        dend4[1].pt3dadd(26.46, -23.83, 11.45, 0.48)
        dend4[1].pt3dadd(27.43, -22.58, 11.46, 0.48)
        dend4[1].pt3dadd(27.67, -22.58, 11.46, 0.48)

        dend4[2] = h.Section(name='dend4[2]')
        dend4[2].connect(        dend4[0](1), 0)
        dend4[2].pt3dclear()
        dend4[2].pt3dadd(12.21, -20.34, 2.26, 0.61)
        dend4[2].pt3dadd(13.66, -28.55, -3.91, 0.61)
        dend4[2].pt3dadd(14.87, -29.30, -3.91, 0.61)
        dend4[2].pt3dadd(15.35, -29.55, -3.91, 0.61)
        dend4[2].pt3dadd(17.77, -31.04, -3.91, 0.61)
        dend4[2].pt3dadd(18.25, -31.29, -3.91, 0.61)
        dend4[2].pt3dadd(19.94, -33.53, -3.90, 0.61)
        dend4[2].pt3dadd(20.18, -34.77, -3.90, 0.61)
        dend4[2].pt3dadd(20.43, -36.26, -3.90, 0.61)
        dend4[2].pt3dadd(20.43, -38.75, -3.90, 0.61)
        dend4[2].pt3dadd(20.43, -39.00, -3.90, 0.61)
        dend4[2].pt3dadd(20.43, -40.99, -3.90, 0.61)
        dend4[2].pt3dadd(20.18, -41.74, -3.90, 0.61)
        dend4[2].pt3dadd(20.18, -44.97, -3.90, 0.61)
        dend4[2].pt3dadd(20.18, -45.22, -3.90, 0.61)
        dend4[2].pt3dadd(19.46, -47.95, -3.93, 0.61)
        dend4[2].pt3dadd(19.46, -47.95, -3.94, 0.61)
        dend4[2].pt3dadd(18.98, -52.18, -1.55, 0.61)
        dend4[2].pt3dadd(18.49, -53.18, -1.55, 0.61)
        dend4[2].pt3dadd(18.49, -53.43, -1.55, 0.61)
        dend4[2].pt3dadd(17.77, -56.41, -1.55, 0.61)
        dend4[2].pt3dadd(18.01, -57.16, -1.55, 0.61)
        dend4[2].pt3dadd(18.98, -58.15, -1.55, 0.61)
        dend4[2].pt3dadd(19.94, -58.40, -1.55, 0.61)

        dend4[3] = h.Section(name='dend4[3]')
        dend4[3].connect(        dend4[2](1), 0)
        dend4[3].pt3dclear()
        dend4[3].pt3dadd(19.94, -58.40, -1.55, 0.54)
        dend4[3].pt3dadd(25.50, -60.64, -3.70, 0.54)
        dend4[3].pt3dadd(27.67, -59.40, -3.70, 0.54)
        dend4[3].pt3dadd(27.91, -59.40, -3.70, 0.54)
        dend4[3].pt3dadd(30.09, -58.90, -3.70, 0.54)
        dend4[3].pt3dadd(30.33, -58.90, -3.70, 0.54)
        dend4[3].pt3dadd(34.19, -57.90, -3.70, 0.54)
        dend4[3].pt3dadd(34.68, -57.90, -3.70, 0.54)
        dend4[3].pt3dadd(37.58, -57.66, -3.70, 0.54)
        dend4[3].pt3dadd(40.47, -57.16, -3.70, 0.54)
        dend4[3].pt3dadd(42.41, -57.16, -3.70, 0.54)
        dend4[3].pt3dadd(43.13, -57.16, -3.70, 0.43)
        dend4[3].pt3dadd(46.27, -57.41, -3.70, 0.43)
        dend4[3].pt3dadd(46.75, -57.41, -3.70, 0.43)
        dend4[3].pt3dadd(51.10, -57.66, -3.70, 0.43)
        dend4[3].pt3dadd(51.83, -57.66, -3.70, 0.43)
        dend4[3].pt3dadd(52.31, -57.66, -3.70, 0.43)
        dend4[3].pt3dadd(53.03, -57.66, -3.70, 0.43)
        dend4[3].pt3dadd(59.31, -59.89, 1.98, 0.43)
        dend4[3].pt3dadd(61.73, -59.89, 2.06, 0.43)
        dend4[3].pt3dadd(62.21, -60.14, 2.14, 0.43)
        dend4[3].pt3dadd(62.45, -61.39, 2.47, 0.43)
        dend4[3].pt3dadd(60.28, -66.36, 2.88, 0.43)
        dend4[3].pt3dadd(60.52, -66.86, 2.88, 0.43)
        dend4[3].pt3dadd(62.94, -68.10, 6.83, 0.43)
        dend4[3].pt3dadd(62.94, -68.10, 6.84, 0.43)
        dend4[3].pt3dadd(64.87, -68.85, 6.97, 0.43)
        dend4[3].pt3dadd(65.11, -68.85, 7.04, 0.43)
        dend4[3].pt3dadd(67.29, -67.36, 7.07, 0.43)
        dend4[3].pt3dadd(67.53, -67.36, 7.10, 0.43)
        dend4[3].pt3dadd(68.73, -66.61, 7.19, 0.43)
        dend4[3].pt3dadd(68.98, -66.61, 7.20, 0.43)
        dend4[3].pt3dadd(70.67, -66.61, 7.21, 0.43)
        dend4[3].pt3dadd(70.91, -66.61, 7.25, 0.43)
        dend4[3].pt3dadd(72.84, -67.36, 7.48, 0.43)
        dend4[3].pt3dadd(73.57, -67.61, 7.49, 0.39)
        dend4[3].pt3dadd(76.46, -68.10, 7.50, 0.39)
        dend4[3].pt3dadd(76.46, -68.10, 7.51, 0.39)
        dend4[3].pt3dadd(79.60, -69.35, 7.80, 0.39)
        dend4[3].pt3dadd(81.05, -71.09, 7.79, 0.39)
        dend4[3].pt3dadd(83.23, -70.34, 16.03, 0.39)
        dend4[3].pt3dadd(83.23, -70.34, 16.04, 0.39)
        dend4[3].pt3dadd(85.64, -72.33, 16.81, 0.39)
        dend4[3].pt3dadd(86.13, -72.33, 16.97, 0.39)
        dend4[3].pt3dadd(88.30, -72.83, 17.62, 0.39)
        dend4[3].pt3dadd(89.51, -73.08, 17.69, 0.39)
        dend4[3].pt3dadd(90.96, -73.83, 17.71, 0.39)
        dend4[3].pt3dadd(92.16, -79.05, 17.73, 0.39)

        dend4[4] = h.Section(name='dend4[4]')
        dend4[4].connect(        dend4[2](1), 0)
        dend4[4].pt3dclear()
        dend4[4].pt3dadd(19.94, -58.40, -1.55, 0.54)
        dend4[4].pt3dadd(23.32, -62.63, -5.32, 0.54)
        dend4[4].pt3dadd(25.98, -63.63, -5.32, 0.54)
        dend4[4].pt3dadd(26.71, -63.87, -5.32, 0.54)
        dend4[4].pt3dadd(27.91, -65.12, -10.57, 0.54)

        dend4[5] = h.Section(name='dend4[5]')
        dend4[5].connect(        dend4[4](1), 0)
        dend4[5].pt3dclear()
        dend4[5].pt3dadd(27.91, -65.12, -10.57, 0.48)
        dend4[5].pt3dadd(36.13, -70.84, -11.21, 0.48)
        dend4[5].pt3dadd(37.82, -74.82, -11.21, 0.48)
        dend4[5].pt3dadd(38.06, -74.82, -11.21, 0.48)
        dend4[5].pt3dadd(45.30, -83.03, -8.96, 0.48)
        dend4[5].pt3dadd(46.51, -87.76, -8.96, 0.48)
        dend4[5].pt3dadd(48.44, -89.75, -8.96, 0.48)
        dend4[5].pt3dadd(48.93, -89.99, -8.96, 0.48)
        dend4[5].pt3dadd(50.62, -91.98, -8.96, 0.48)
        dend4[5].pt3dadd(52.55, -93.73, -8.96, 0.48)
        dend4[5].pt3dadd(53.28, -94.22, -8.96, 0.48)
        dend4[5].pt3dadd(55.45, -95.72, -8.96, 0.48)
        dend4[5].pt3dadd(55.93, -95.96, -8.96, 0.48)
        dend4[5].pt3dadd(58.35, -97.46, -8.96, 0.48)
        dend4[5].pt3dadd(58.59, -97.71, -8.96, 0.48)
        dend4[5].pt3dadd(60.04, -101.44, -8.96, 0.48)
        dend4[5].pt3dadd(60.28, -101.44, -8.96, 0.48)
        dend4[5].pt3dadd(69.46, -109.40, -8.00, 0.48)
        dend4[5].pt3dadd(71.39, -112.88, -8.00, 0.48)
        dend4[5].pt3dadd(71.87, -112.88, -8.00, 0.48)
        dend4[5].pt3dadd(73.32, -113.63, -8.00, 0.48)
        dend4[5].pt3dadd(75.74, -117.61, -8.00, 0.48)
        dend4[5].pt3dadd(76.22, -120.84, -3.53, 0.48)
        dend4[5].pt3dadd(76.71, -123.08, -3.53, 0.48)

        dend4[6] = h.Section(name='dend4[6]')
        dend4[6].connect(        dend4[5](1), 0)
        dend4[6].pt3dclear()
        dend4[6].pt3dadd(76.71, -123.08, -3.53, 0.39)
        dend4[6].pt3dadd(78.15, -127.81, -3.53, 0.39)
        dend4[6].pt3dadd(79.12, -129.80, -3.53, 0.39)
        dend4[6].pt3dadd(79.12, -130.54, -3.53, 0.39)
        dend4[6].pt3dadd(79.60, -133.28, -3.53, 0.39)
        dend4[6].pt3dadd(79.85, -133.53, -3.54, 0.39)
        dend4[6].pt3dadd(80.33, -134.52, -3.54, 0.39)
        dend4[6].pt3dadd(80.33, -134.77, -3.54, 0.39)
        dend4[6].pt3dadd(78.88, -137.51, -4.57, 0.39)
        dend4[6].pt3dadd(79.85, -140.24, -4.57, 0.39)
        dend4[6].pt3dadd(79.85, -142.23, -4.57, 0.39)
        dend4[6].pt3dadd(78.64, -145.96, -4.58, 0.39)
        dend4[6].pt3dadd(78.40, -145.96, -4.58, 0.39)
        dend4[6].pt3dadd(76.95, -146.71, -4.58, 0.39)
        dend4[6].pt3dadd(74.77, -147.46, -4.58, 0.39)
        dend4[6].pt3dadd(74.29, -147.71, -4.58, 0.39)
        dend4[6].pt3dadd(73.81, -148.20, -4.58, 0.39)
        dend4[6].pt3dadd(72.84, -149.45, -4.58, 0.39)
        dend4[6].pt3dadd(72.60, -149.94, -4.58, 0.39)
        dend4[6].pt3dadd(72.60, -152.43, -4.58, 0.39)
        dend4[6].pt3dadd(72.60, -152.68, -4.58, 0.39)

        dend4[7] = h.Section(name='dend4[7]')
        dend4[7].connect(        dend4[5](1), 0)
        dend4[7].pt3dclear()
        dend4[7].pt3dadd(76.71, -123.08, -3.53, 0.39)
        dend4[7].pt3dadd(81.54, -128.55, 0.48, 0.39)
        dend4[7].pt3dadd(83.95, -129.30, 0.48, 0.39)
        dend4[7].pt3dadd(84.43, -129.30, 0.48, 0.39)
        dend4[7].pt3dadd(85.64, -133.78, 0.48, 0.39)
        dend4[7].pt3dadd(86.37, -135.52, 0.48, 0.39)
        dend4[7].pt3dadd(87.33, -136.01, 0.48, 0.39)
        dend4[7].pt3dadd(89.27, -136.51, 0.48, 0.39)
        dend4[7].pt3dadd(90.72, -137.51, 0.48, 0.39)
        dend4[7].pt3dadd(91.92, -140.74, 0.48, 0.39)
        dend4[7].pt3dadd(92.89, -141.98, 0.48, 0.39)
        dend4[7].pt3dadd(94.82, -143.23, 0.48, 0.39)
        dend4[7].pt3dadd(96.27, -144.47, 0.48, 0.39)
        dend4[7].pt3dadd(96.75, -145.72, 0.48, 0.39)
        dend4[7].pt3dadd(97.72, -146.71, 0.48, 0.39)
        dend4[7].pt3dadd(98.69, -148.95, 0.48, 0.39)
        dend4[7].pt3dadd(99.17, -149.45, 0.48, 0.39)

        dend4[8] = h.Section(name='dend4[8]')
        dend4[8].connect(        dend4[4](1), 0)
        dend4[8].pt3dclear()
        dend4[8].pt3dadd(27.91, -65.12, -10.57, 0.48)
        dend4[8].pt3dadd(28.15, -68.35, -15.30, 0.48)
        dend4[8].pt3dadd(28.64, -69.84, -15.30, 0.48)
        dend4[8].pt3dadd(28.64, -70.09, -15.30, 0.48)
        dend4[8].pt3dadd(29.12, -72.83, -15.30, 0.48)
        dend4[8].pt3dadd(29.36, -73.33, -15.30, 0.48)
        dend4[8].pt3dadd(30.09, -75.32, -15.30, 0.48)
        dend4[8].pt3dadd(30.09, -79.55, -20.21, 0.48)
        dend4[8].pt3dadd(29.85, -81.79, -20.21, 0.48)
        dend4[8].pt3dadd(29.60, -82.03, -20.21, 0.48)
        dend4[8].pt3dadd(30.33, -83.28, -20.21, 0.48)

        dend4[9] = h.Section(name='dend4[9]')
        dend4[9].connect(        dend4[8](1), 0)
        dend4[9].pt3dclear()
        dend4[9].pt3dadd(30.33, -83.28, -20.21, 0.43)
        dend4[9].pt3dadd(35.64, -83.53, -20.21, 0.43)
        dend4[9].pt3dadd(36.61, -87.51, -20.21, 0.43)
        dend4[9].pt3dadd(36.85, -87.51, -20.21, 0.43)
        dend4[9].pt3dadd(37.82, -88.50, -20.21, 0.43)
        dend4[9].pt3dadd(38.30, -89.50, -20.21, 0.43)
        dend4[9].pt3dadd(38.06, -91.24, -20.22, 0.43)

        dend4[10] = h.Section(name='dend4[10]')
        dend4[10].connect(        dend4[9](1), 0)
        dend4[10].pt3dclear()
        dend4[10].pt3dadd(38.06, -91.24, -20.22, 0.39)
        dend4[10].pt3dadd(36.37, -94.97, -28.98, 0.39)
        dend4[10].pt3dadd(36.13, -94.97, -28.98, 0.39)
        dend4[10].pt3dadd(35.64, -99.20, -28.98, 0.39)
        dend4[10].pt3dadd(35.88, -99.70, -28.98, 0.39)
        dend4[10].pt3dadd(36.37, -100.44, -28.99, 0.39)
        dend4[10].pt3dadd(36.37, -100.94, -28.99, 0.39)
        dend4[10].pt3dadd(34.92, -103.43, -28.99, 0.39)
        dend4[10].pt3dadd(34.68, -103.92, -28.99, 0.39)
        dend4[10].pt3dadd(34.19, -104.92, -28.99, 0.39)
        dend4[10].pt3dadd(33.95, -105.91, -28.99, 0.39)
        dend4[10].pt3dadd(33.95, -107.90, -28.99, 0.39)
        dend4[10].pt3dadd(34.43, -109.65, -28.99, 0.39)
        dend4[10].pt3dadd(34.68, -110.64, -28.99, 0.39)
        dend4[10].pt3dadd(34.68, -111.14, -28.99, 0.39)
        dend4[10].pt3dadd(34.43, -111.64, -28.99, 0.39)
        dend4[10].pt3dadd(34.19, -111.88, -28.99, 0.39)
        dend4[10].pt3dadd(34.19, -112.38, -34.72, 0.39)
        dend4[10].pt3dadd(32.02, -113.38, -34.72, 0.39)
        dend4[10].pt3dadd(32.02, -114.37, -34.72, 0.39)
        dend4[10].pt3dadd(32.02, -114.62, -34.72, 0.39)
        dend4[10].pt3dadd(32.26, -115.62, -34.72, 0.39)
        dend4[10].pt3dadd(32.26, -115.86, -34.72, 0.39)
        dend4[10].pt3dadd(33.71, -117.11, -34.72, 0.39)
        dend4[10].pt3dadd(33.95, -117.36, -34.72, 0.39)
        dend4[10].pt3dadd(33.95, -118.85, -34.72, 0.39)
        dend4[10].pt3dadd(33.71, -119.60, -34.72, 0.39)
        dend4[10].pt3dadd(36.61, -121.09, -34.72, 0.39)
        dend4[10].pt3dadd(36.85, -121.09, -34.72, 0.39)
        dend4[10].pt3dadd(39.51, -121.09, -34.72, 0.39)
        dend4[10].pt3dadd(39.99, -122.33, -34.72, 0.39)
        dend4[10].pt3dadd(39.99, -124.32, -34.72, 0.39)
        dend4[10].pt3dadd(40.23, -125.57, -34.72, 0.39)
        dend4[10].pt3dadd(40.23, -126.56, -34.72, 0.39)
        dend4[10].pt3dadd(40.23, -129.05, -34.72, 0.39)
        dend4[10].pt3dadd(40.23, -130.29, -34.72, 0.39)
        dend4[10].pt3dadd(40.47, -130.79, -34.72, 0.39)
        dend4[10].pt3dadd(40.47, -132.03, -34.72, 0.39)
        dend4[10].pt3dadd(39.75, -132.78, -34.72, 0.39)
        dend4[10].pt3dadd(38.54, -133.28, -34.72, 0.39)
        dend4[10].pt3dadd(35.16, -134.52, -34.72, 0.39)
        dend4[10].pt3dadd(35.16, -135.52, -34.72, 0.39)

        dend4[11] = h.Section(name='dend4[11]')
        dend4[11].connect(        dend4[9](1), 0)
        dend4[11].pt3dclear()
        dend4[11].pt3dadd(38.06, -91.24, -20.22, 0.39)
        dend4[11].pt3dadd(41.44, -93.73, -20.19, 0.39)
        dend4[11].pt3dadd(44.82, -96.96, -22.26, 0.39)
        dend4[11].pt3dadd(49.17, -97.71, -22.26, 0.39)
        dend4[11].pt3dadd(49.65, -97.71, -22.26, 0.39)
        dend4[11].pt3dadd(52.31, -98.70, -22.26, 0.39)
        dend4[11].pt3dadd(52.55, -98.70, -22.26, 0.39)
        dend4[11].pt3dadd(54.97, -101.69, -22.26, 0.39)
        dend4[11].pt3dadd(56.42, -103.43, -22.26, 0.39)
        dend4[11].pt3dadd(57.14, -104.42, -22.26, 0.39)
        dend4[11].pt3dadd(57.14, -106.41, -22.26, 0.39)
        dend4[11].pt3dadd(57.62, -107.90, -22.26, 0.39)
        dend4[11].pt3dadd(57.62, -109.15, -22.26, 0.39)
        dend4[11].pt3dadd(59.07, -113.63, -22.26, 0.39)
        dend4[11].pt3dadd(60.04, -115.12, -22.26, 0.39)
        dend4[11].pt3dadd(61.00, -116.11, -22.26, 0.39)
        dend4[11].pt3dadd(62.70, -118.35, -22.26, 0.39)
        dend4[11].pt3dadd(63.42, -119.10, -22.26, 0.39)
        dend4[11].pt3dadd(65.35, -120.34, -22.27, 0.39)
        dend4[11].pt3dadd(69.22, -120.84, -22.27, 0.39)
        dend4[11].pt3dadd(69.46, -120.84, -22.27, 0.39)
        dend4[11].pt3dadd(69.70, -120.84, -22.27, 0.39)
        dend4[11].pt3dadd(70.91, -127.06, -22.27, 0.39)
        dend4[11].pt3dadd(71.63, -127.31, -22.27, 0.39)
        dend4[11].pt3dadd(71.87, -127.81, -22.27, 0.39)
        dend4[11].pt3dadd(72.60, -132.28, -26.95, 0.39)
        dend4[11].pt3dadd(72.12, -135.52, -26.95, 0.39)
        dend4[11].pt3dadd(72.12, -136.01, -26.95, 0.39)
        dend4[11].pt3dadd(73.57, -136.51, -26.95, 0.39)
        dend4[11].pt3dadd(75.50, -137.51, -26.95, 0.39)
        dend4[11].pt3dadd(76.46, -138.00, -26.95, 0.39)
        dend4[11].pt3dadd(77.67, -140.49, -26.95, 0.39)
        dend4[11].pt3dadd(77.67, -141.98, -26.95, 0.39)
        dend4[11].pt3dadd(77.67, -142.48, -26.95, 0.39)
        dend4[11].pt3dadd(79.85, -142.98, -26.95, 0.39)
        dend4[11].pt3dadd(81.05, -143.23, -26.95, 0.39)
        dend4[11].pt3dadd(81.78, -143.48, -26.95, 0.39)
        dend4[11].pt3dadd(82.74, -143.97, -26.95, 0.39)
        dend4[11].pt3dadd(89.02, -146.21, -30.93, 0.39)
        dend4[11].pt3dadd(92.41, -147.21, -30.93, 0.39)
        dend4[11].pt3dadd(94.82, -148.45, -30.93, 0.39)
        dend4[11].pt3dadd(95.30, -148.45, -30.93, 0.39)
        dend4[11].pt3dadd(97.24, -149.45, -30.93, 0.39)
        dend4[11].pt3dadd(97.48, -149.45, -30.93, 0.39)
        dend4[11].pt3dadd(99.89, -153.18, -30.93, 0.39)
        dend4[11].pt3dadd(102.07, -153.43, -30.93, 0.39)
        dend4[11].pt3dadd(102.07, -152.93, -30.93, 0.39)
        dend4[11].pt3dadd(104.48, -150.69, -30.93, 0.39)
        dend4[11].pt3dadd(106.42, -148.70, -30.93, 0.39)
        dend4[11].pt3dadd(107.38, -148.20, -30.93, 0.39)
        dend4[11].pt3dadd(107.62, -147.71, -30.93, 0.39)
        dend4[11].pt3dadd(108.11, -144.72, -30.93, 0.39)
        dend4[11].pt3dadd(109.31, -143.97, -30.93, 0.39)
        dend4[11].pt3dadd(109.56, -143.48, -30.93, 0.39)

        dend4[12] = h.Section(name='dend4[12]')
        dend4[12].connect(        dend4[8](1), 0)
        dend4[12].pt3dclear()
        dend4[12].pt3dadd(30.33, -83.28, -20.21, 0.39)
        dend4[12].pt3dadd(30.81, -87.51, -20.21, 0.39)
        dend4[12].pt3dadd(29.12, -90.49, -16.08, 0.39)
        dend4[12].pt3dadd(27.67, -93.23, -16.08, 0.39)
        dend4[12].pt3dadd(27.43, -94.72, -16.08, 0.39)
        dend4[12].pt3dadd(27.43, -96.96, -16.08, 0.39)
        dend4[12].pt3dadd(27.43, -98.45, -16.08, 0.39)
        dend4[12].pt3dadd(27.43, -100.44, -16.08, 0.39)
        dend4[12].pt3dadd(27.67, -103.18, -16.08, 0.39)
        dend4[12].pt3dadd(27.67, -104.42, -16.07, 0.39)
        dend4[12].pt3dadd(27.91, -107.16, -16.06, 0.39)
        dend4[12].pt3dadd(27.91, -110.64, -16.04, 0.39)
        dend4[12].pt3dadd(28.15, -111.39, -16.04, 0.39)
        dend4[12].pt3dadd(27.91, -112.38, -16.04, 0.39)
        dend4[12].pt3dadd(27.91, -114.12, -16.04, 0.39)
        dend4[12].pt3dadd(28.88, -114.62, -16.04, 0.39)
        dend4[12].pt3dadd(30.81, -117.36, -16.04, 0.39)
        dend4[12].pt3dadd(33.23, -120.09, -16.04, 0.39)
        dend4[12].pt3dadd(35.16, -125.82, -13.02, 0.39)
        dend4[12].pt3dadd(35.40, -125.82, -13.02, 0.39)
        dend4[12].pt3dadd(37.58, -127.56, -13.02, 0.39)
        dend4[12].pt3dadd(38.78, -128.30, -13.02, 0.39)
        dend4[12].pt3dadd(39.51, -128.80, -13.02, 0.39)
        dend4[12].pt3dadd(39.75, -130.04, -13.02, 0.39)
        dend4[12].pt3dadd(38.06, -138.25, -18.36, 0.39)
        dend4[12].pt3dadd(38.30, -138.25, -18.36, 0.39)
        dend4[12].pt3dadd(40.23, -141.24, -18.35, 0.39)
        dend4[12].pt3dadd(40.47, -141.74, -18.35, 0.39)
        dend4[12].pt3dadd(40.72, -144.47, -18.35, 0.39)
        dend4[12].pt3dadd(40.23, -145.22, -18.35, 0.39)
        dend4[12].pt3dadd(39.75, -147.21, -18.35, 0.39)
        dend4[12].pt3dadd(39.51, -148.70, -18.35, 0.39)
        dend4[12].pt3dadd(39.51, -149.94, -18.35, 0.39)
        dend4[12].pt3dadd(39.51, -151.19, -18.35, 0.39)
        dend4[12].pt3dadd(39.51, -152.68, -18.35, 0.39)
        dend4[12].pt3dadd(39.51, -153.68, -18.35, 0.39)
        dend4[12].pt3dadd(39.51, -154.42, -18.35, 0.39)
        dend4[12].pt3dadd(39.02, -155.17, -18.35, 0.39)
        dend4[12].pt3dadd(38.54, -155.67, -18.35, 0.39)
        dend4[12].pt3dadd(37.58, -156.16, -18.35, 0.39)
        dend4[12].pt3dadd(36.61, -156.41, -18.35, 0.39)
        dend4[12].pt3dadd(33.95, -156.66, -18.35, 0.39)
        dend4[12].pt3dadd(33.71, -156.66, -18.35, 0.39)
        dend4[12].pt3dadd(32.26, -156.66, -18.39, 0.39)
        dend4[12].pt3dadd(31.05, -155.91, -20.44, 0.39)
        dend4[12].pt3dadd(28.15, -156.91, -20.44, 0.39)
        dend4[12].pt3dadd(27.67, -157.16, -20.44, 0.39)
        dend4[12].pt3dadd(26.71, -157.66, -20.44, 0.39)
        dend4[12].pt3dadd(25.74, -158.65, -20.44, 0.39)
        dend4[12].pt3dadd(24.77, -159.40, -20.44, 0.39)
        dend4[12].pt3dadd(23.32, -160.14, -20.44, 0.39)
        dend4[12].pt3dadd(22.60, -161.14, -20.44, 0.39)
        dend4[12].pt3dadd(21.87, -162.63, -20.44, 0.39)
        dend4[12].pt3dadd(21.39, -165.37, -20.44, 0.39)
        dend4[12].pt3dadd(20.91, -166.11, -20.44, 0.39)
        dend4[12].pt3dadd(20.18, -166.86, -20.44, 0.39)
        dend4[12].pt3dadd(18.73, -169.85, -20.44, 0.39)
        dend4[12].pt3dadd(18.01, -170.59, -20.44, 0.39)
        dend4[12].pt3dadd(17.77, -170.84, -20.44, 0.39)
        dend4[12].pt3dadd(17.29, -171.34, -20.44, 0.39)
        dend4[12].pt3dadd(16.80, -172.83, -20.44, 0.39)
        dend4[12].pt3dadd(17.04, -172.83, -20.44, 0.39)

        apic = [None]*79

        apic[0] = h.Section(name='apic[0]')
        apic[0].connect(        soma(0), 0)
        apic[0].pt3dclear()
        apic[0].pt3dadd(0, 10.20, 0.00, 4.0)
        apic[0].pt3dadd(0, 13.43, 0.00, 4.0)
        apic[0].pt3dadd(0, 16.17, 0.00, 3.8)
        apic[0].pt3dadd(-0.48, 19.90, 0.19, 3.8)
        apic[0].pt3dadd(0.00, 24.13, 0.20, 3.8)
        apic[0].pt3dadd(0.00, 27.36, 0.20, 3.8)
        apic[0].pt3dadd(0.00, 31.59, 0.20, 3.8)
        apic[0].pt3dadd(0.00, 34.83, 0.71, 3.8)
        apic[0].pt3dadd(-0.24, 37.56, 0.71, 3.8)
        
        apic[1] = h.Section(name='apic[1]')
        apic[1].connect(        apic[0](1), 0)
        apic[1].pt3dclear()
        apic[1].pt3dadd(-0.24, 37.56, 0.71, 0.96)
        apic[1].pt3dadd(3.14, 37.81, 0.04, 0.96)
        apic[1].pt3dadd(4.11, 42.54, 0.04, 0.96)
        apic[1].pt3dadd(5.31, 45.52, 0.04, 0.96)
        apic[1].pt3dadd(7.00, 47.76, 0.04, 0.96)
        apic[1].pt3dadd(7.49, 48.01, 0.04, 0.96)
        apic[1].pt3dadd(13.53, 51.49, -5.41, 0.54)
        apic[1].pt3dadd(14.73, 52.99, -5.44, 0.54)
        apic[1].pt3dadd(17.63, 52.99, -5.44, 0.54)
        apic[1].pt3dadd(18.12, 52.74, -5.44, 0.54)
        apic[1].pt3dadd(20.77, 51.99, -5.44, 0.54)
        apic[1].pt3dadd(21.01, 51.99, -5.44, 0.54)
        apic[1].pt3dadd(22.95, 52.49, -5.44, 0.54)
        apic[1].pt3dadd(23.43, 52.49, -5.44, 0.54)
        apic[1].pt3dadd(25.85, 51.99, -5.44, 0.54)
        apic[1].pt3dadd(26.09, 51.99, -5.44, 0.54)
        apic[1].pt3dadd(30.92, 49.01, -2.79, 0.54)
        apic[1].pt3dadd(33.82, 49.50, -2.79, 0.54)
        apic[1].pt3dadd(35.99, 50.25, -2.79, 0.54)
        apic[1].pt3dadd(36.23, 50.25, -2.79, 0.54)
        apic[1].pt3dadd(37.92, 51.00, -2.79, 0.54)
        apic[1].pt3dadd(38.41, 51.00, -2.79, 0.54)
        apic[1].pt3dadd(40.82, 51.49, -2.79, 0.54)
        apic[1].pt3dadd(41.55, 51.49, -2.79, 0.54)
        apic[1].pt3dadd(43.48, 51.24, -2.79, 0.54)
        apic[1].pt3dadd(43.96, 51.24, -2.79, 0.54)
        apic[1].pt3dadd(45.41, 53.73, -2.79, 0.54)
        apic[1].pt3dadd(45.89, 53.98, -2.79, 0.54)
        apic[1].pt3dadd(47.10, 53.98, -2.79, 0.54)
        apic[1].pt3dadd(47.83, 53.73, -2.79, 0.54)
        apic[1].pt3dadd(48.79, 52.49, -2.79, 0.54)
        apic[1].pt3dadd(49.76, 52.24, -2.79, 0.54)
        apic[1].pt3dadd(51.93, 52.24, -2.79, 0.54)
        apic[1].pt3dadd(54.35, 52.74, -2.82, 0.54)
        apic[1].pt3dadd(56.28, 53.73, 2.31, 0.43)
        apic[1].pt3dadd(57.00, 55.72, 2.33, 0.43)
        apic[1].pt3dadd(57.25, 55.72, 2.33, 0.43)
        apic[1].pt3dadd(58.45, 57.46, 2.35, 0.43)
        apic[1].pt3dadd(60.87, 56.72, 2.37, 0.43)
        apic[1].pt3dadd(61.11, 56.72, 2.37, 0.43)
        apic[1].pt3dadd(61.59, 56.47, 2.37, 0.43)
        apic[1].pt3dadd(61.84, 55.22, 2.41, 0.43)
        apic[1].pt3dadd(61.84, 55.22, 2.42, 0.43)
        apic[1].pt3dadd(62.80, 54.48, 2.71, 0.43)
        apic[1].pt3dadd(63.04, 54.48, 2.72, 0.43)
        apic[1].pt3dadd(64.01, 54.73, 2.77, 0.43)
        apic[1].pt3dadd(64.25, 54.73, 2.86, 0.43)
        apic[1].pt3dadd(65.70, 55.22, 3.86, 0.43)
        apic[1].pt3dadd(66.18, 55.47, 4.01, 0.43)
        apic[1].pt3dadd(68.12, 56.22, 5.04, 0.43)
        apic[1].pt3dadd(68.12, 56.22, 5.08, 0.43)
        apic[1].pt3dadd(69.32, 56.72, 5.08, 0.43)
        apic[1].pt3dadd(69.57, 56.72, 5.08, 0.43)
        apic[1].pt3dadd(71.01, 56.97, 5.14, 0.43)
        apic[1].pt3dadd(71.01, 56.97, 5.18, 0.43)
        apic[1].pt3dadd(72.46, 55.72, 5.37, 0.43)
        apic[1].pt3dadd(72.46, 55.72, 5.43, 0.43)
        apic[1].pt3dadd(75.36, 51.49, 7.57, 0.43)
        apic[1].pt3dadd(75.60, 51.49, 7.58, 0.43)
        apic[1].pt3dadd(78.02, 51.99, 7.61, 0.43)
        apic[1].pt3dadd(78.02, 52.24, 7.63, 0.43)
        apic[1].pt3dadd(82.37, 49.75, 11.60, 0.43)
        apic[1].pt3dadd(84.54, 48.26, 12.09, 0.43)
        apic[1].pt3dadd(84.54, 48.26, 12.20, 0.43)
        apic[1].pt3dadd(88.16, 46.27, 12.59, 0.43)
        apic[1].pt3dadd(88.16, 46.27, 12.64, 0.43)
        apic[1].pt3dadd(90.82, 45.52, 12.73, 0.43)
        apic[1].pt3dadd(91.06, 45.52, 12.74, 0.43)
        apic[1].pt3dadd(94.93, 43.28, 15.93, 0.43)
        apic[1].pt3dadd(97.10, 44.78, 16.15, 0.43)
        apic[1].pt3dadd(98.55, 44.53, 16.15, 0.39)
        apic[1].pt3dadd(98.79, 44.03, 16.15, 0.39)
        apic[1].pt3dadd(99.28, 43.03, 16.14, 0.39)
        apic[1].pt3dadd(98.55, 42.79, 16.14, 0.39)
        apic[1].pt3dadd(98.31, 41.79, 16.14, 0.39)
        apic[1].pt3dadd(96.86, 40.05, 16.14, 0.39)
        apic[1].pt3dadd(95.17, 39.30, 18.64, 0.39)
        apic[1].pt3dadd(95.17, 39.30, 19.32, 0.39)
        apic[1].pt3dadd(94.69, 37.81, 20.48, 0.39)
        apic[1].pt3dadd(94.69, 37.56, 20.68, 0.39)
        apic[1].pt3dadd(94.69, 36.57, 20.78, 0.39)
        apic[1].pt3dadd(95.17, 35.82, 20.81, 0.39)
        apic[1].pt3dadd(95.41, 35.32, 20.81, 0.39)
        apic[1].pt3dadd(96.38, 35.07, 20.82, 0.39)
        apic[1].pt3dadd(96.86, 35.07, 20.83, 0.39)
        apic[1].pt3dadd(97.10, 35.07, 20.85, 0.39)
        apic[1].pt3dadd(97.58, 34.33, 20.87, 0.39)
        
        apic[2] = h.Section(name='apic[2]')
        apic[2].connect(        apic[0](1), 0)
        apic[2].pt3dclear()
        apic[2].pt3dadd(-0.24, 37.56, 0.71, 3.8)
        apic[2].pt3dadd(-0.72, 40.80, 0.53, 3.8)
        apic[2].pt3dadd(-0.72, 43.78, 0.53, 3.8)
        apic[2].pt3dadd(-0.97, 43.78, 0.53, 3.8)
        apic[2].pt3dadd(-1.21, 47.26, 0.53, 3.8)
        apic[2].pt3dadd(-1.21, 50.50, 0.53, 3.8)
        apic[2].pt3dadd(-1.45, 53.73, 0.53, 3.8)
        apic[2].pt3dadd(-1.21, 57.21, 0.53, 3.8)
        apic[2].pt3dadd(-0.48, 61.19, 1.85, 3.8)
        apic[2].pt3dadd(0.72, 65.42, 1.85, 3.8)
        apic[2].pt3dadd(0.97, 69.65, 1.85, 3.8)
        apic[2].pt3dadd(0.97, 76.37, 1.85, 3.8)
        apic[2].pt3dadd(2.17, 83.08, 1.85, 3.8)
        apic[2].pt3dadd(2.17, 87.06, 1.85, 3.8)
        apic[2].pt3dadd(2.17, 87.31, 1.85, 3.8)
        apic[2].pt3dadd(2.66, 90.30, 1.85, 3.8)
        apic[2].pt3dadd(3.38, 94.03, 1.85, 4.00)
        
        apic[3] = h.Section(name='apic[3]')
        apic[3].connect(        apic[2](1), 0)
        apic[3].pt3dclear()
        apic[3].pt3dadd(3.38, 94.03, 1.85, 1.10)
        apic[3].pt3dadd(2.42, 96.77, 1.85, 1.10)
        apic[3].pt3dadd(2.17, 96.77, 1.85, 1.10)
        apic[3].pt3dadd(0.48, 97.26, 1.85, 1.10)
        apic[3].pt3dadd(-1.21, 98.76, 1.85, 0.96)
        apic[3].pt3dadd(-1.69, 100.75, 1.85, 0.96)
        apic[3].pt3dadd(-0.72, 103.23, 1.85, 0.96)
        apic[3].pt3dadd(-0.48, 105.97, 5.58, 0.96)
        apic[3].pt3dadd(1.21, 106.97, 6.29, 0.61)
        apic[3].pt3dadd(0.72, 109.20, 6.84, 0.61)
        apic[3].pt3dadd(2.42, 110.20, 6.84, 0.61)
        apic[3].pt3dadd(2.66, 110.45, 6.84, 0.61)
        
        apic[4] = h.Section(name='apic[4]')
        apic[4].connect(        apic[3](1), 0)
        apic[4].pt3dclear()
        apic[4].pt3dadd(2.66, 110.45, 6.84, 0.48)
        apic[4].pt3dadd(4.35, 112.44, 2.23, 0.48)
        apic[4].pt3dadd(7.49, 114.93, 2.23, 0.48)
        apic[4].pt3dadd(7.97, 114.93, 2.23, 0.48)
        apic[4].pt3dadd(8.94, 117.91, 2.23, 0.48)
        apic[4].pt3dadd(8.45, 118.41, 2.23, 0.48)
        apic[4].pt3dadd(8.45, 119.15, 2.23, 0.48)
        apic[4].pt3dadd(7.97, 119.40, 2.23, 0.48)
        apic[4].pt3dadd(7.49, 120.15, 2.23, 0.48)
        apic[4].pt3dadd(7.00, 120.65, 2.23, 0.48)
        apic[4].pt3dadd(6.76, 121.14, 2.23, 0.48)
        apic[4].pt3dadd(6.52, 121.14, 2.23, 0.48)
        apic[4].pt3dadd(5.80, 121.89, 2.23, 0.48)
        apic[4].pt3dadd(5.31, 122.39, 2.23, 0.48)
        apic[4].pt3dadd(5.07, 122.89, 2.23, 0.48)
        apic[4].pt3dadd(4.83, 123.38, 2.23, 0.48)
        apic[4].pt3dadd(4.59, 123.88, 2.23, 0.48)
        apic[4].pt3dadd(4.35, 125.37, 2.23, 0.48)
        apic[4].pt3dadd(4.35, 126.12, 2.23, 0.48)
        apic[4].pt3dadd(4.35, 128.11, 2.22, 0.48)
        apic[4].pt3dadd(4.35, 128.86, 2.22, 0.48)
        apic[4].pt3dadd(5.07, 129.10, 2.22, 0.48)
        apic[4].pt3dadd(5.80, 130.10, 2.22, 0.48)
        apic[4].pt3dadd(5.80, 133.33, 2.22, 0.48)
        apic[4].pt3dadd(5.56, 137.31, 2.22, 0.48)
        
        apic[5] = h.Section(name='apic[5]')
        apic[5].connect(        apic[3](1), 0)
        apic[5].pt3dclear()
        apic[5].pt3dadd(2.66, 110.45, 6.84, 0.48)
        apic[5].pt3dadd(2.42, 111.44, 2.21, 0.48)
        apic[5].pt3dadd(2.42, 111.44, 2.22, 0.48)
        apic[5].pt3dadd(2.17, 112.94, 2.23, 0.48)
        apic[5].pt3dadd(2.17, 112.94, -2.75, 0.48)
        apic[5].pt3dadd(1.93, 118.16, -2.75, 0.48)
        apic[5].pt3dadd(1.93, 119.65, -2.25, 0.48)
        apic[5].pt3dadd(1.93, 119.65, -2.24, 0.48)
        apic[5].pt3dadd(-0.97, 119.90, -0.24, 0.48)
        apic[5].pt3dadd(-0.97, 119.90, -0.12, 0.48)
        apic[5].pt3dadd(-2.66, 121.64, 0.59, 0.48)
        apic[5].pt3dadd(-2.66, 121.64, 0.91, 0.48)
        apic[5].pt3dadd(-4.59, 122.89, 1.56, 0.48)
        apic[5].pt3dadd(-4.83, 122.89, 2.13, 0.48)
        apic[5].pt3dadd(-5.56, 124.38, 2.67, 0.48)
        apic[5].pt3dadd(-5.56, 124.63, 2.93, 0.48)
        apic[5].pt3dadd(-5.80, 126.12, 3.33, 0.48)
        apic[5].pt3dadd(-5.80, 126.12, 3.61, 0.48)
        apic[5].pt3dadd(-5.80, 127.36, 4.16, 0.48)
        apic[5].pt3dadd(-5.80, 127.36, 4.35, 0.48)
        apic[5].pt3dadd(-7.49, 129.10, 9.18, 0.48)
        apic[5].pt3dadd(-9.90, 133.33, 11.43, 0.48)
        apic[5].pt3dadd(-9.90, 133.33, 11.66, 0.48)
        apic[5].pt3dadd(-12.08, 136.32, 12.25, 0.48)
        apic[5].pt3dadd(-12.32, 136.32, 12.25, 0.48)
        apic[5].pt3dadd(-13.04, 137.31, 12.25, 0.48)
        apic[5].pt3dadd(-13.04, 137.81, 12.25, 0.48)
        apic[5].pt3dadd(-13.04, 138.56, 12.25, 0.48)
        apic[5].pt3dadd(-13.04, 139.05, 12.25, 0.48)
        apic[5].pt3dadd(-13.04, 140.05, 12.25, 0.48)
        apic[5].pt3dadd(-13.29, 140.05, 12.25, 0.48)
        
        apic[6] = h.Section(name='apic[6]')
        apic[6].connect(        apic[2](1), 0)
        apic[6].pt3dclear()
        apic[6].pt3dadd(3.38, 94.03, 1.85, 3.62)
        apic[6].pt3dadd(4.59, 97.76, 6.45, 3.62)
        apic[6].pt3dadd(4.83, 101.00, 6.45, 3.62)
        apic[6].pt3dadd(4.83, 102.74, 6.45, 3.62)
        apic[6].pt3dadd(6.04, 105.97, 6.45, 3.62)
        apic[6].pt3dadd(7.73, 107.46, 6.45, 3.62)
        apic[6].pt3dadd(9.18, 109.70, 6.45, 3.62)
        apic[6].pt3dadd(10.14, 111.19, 6.45, 3.62)
        
        apic[7] = h.Section(name='apic[7]')
        apic[7].connect(        apic[6](1), 0)
        apic[7].pt3dclear()
        apic[7].pt3dadd(10.14, 111.19, 6.45, 1.20)
        apic[7].pt3dadd(7.25, 112.69, 11.70, 1.20)
        apic[7].pt3dadd(5.56, 113.68, 11.70, 1.20)
        apic[7].pt3dadd(5.31, 113.68, 11.70, 1.20)
        apic[7].pt3dadd(3.86, 114.43, 11.70, 1.20)
        apic[7].pt3dadd(3.62, 114.43, 11.70, 1.20)
        apic[7].pt3dadd(2.17, 115.17, 11.70, 1.20)
        apic[7].pt3dadd(1.45, 115.17, 11.70, 1.20)
        
        apic[8] = h.Section(name='apic[8]')
        apic[8].connect(        apic[7](1), 0)
        apic[8].pt3dclear()
        apic[8].pt3dadd(1.45, 115.17, 11.70, 0.72)
        apic[8].pt3dadd(-0.00, 117.66, 11.70, 0.72)
        apic[8].pt3dadd(-0.24, 117.66, 11.70, 0.72)
        apic[8].pt3dadd(-0.00, 120.15, 11.70, 0.72)
        apic[8].pt3dadd(1.21, 121.64, 11.70, 0.72)
        apic[8].pt3dadd(1.45, 121.89, 11.70, 0.72)
        apic[8].pt3dadd(2.17, 123.38, 11.70, 0.72)
        apic[8].pt3dadd(3.38, 125.12, 11.70, 0.72)
        apic[8].pt3dadd(3.38, 125.37, 11.70, 0.72)
        apic[8].pt3dadd(4.11, 126.12, 11.70, 0.72)
        apic[8].pt3dadd(4.59, 126.87, 11.70, 0.72)
        apic[8].pt3dadd(5.56, 127.86, 11.70, 0.72)
        apic[8].pt3dadd(7.00, 128.86, 11.70, 0.72)
        apic[8].pt3dadd(7.49, 129.35, 11.70, 0.72)
        apic[8].pt3dadd(7.73, 129.35, 11.70, 0.72)
        apic[8].pt3dadd(11.11, 131.84, 11.70, 0.72)
        apic[8].pt3dadd(11.35, 135.57, 9.82, 0.72)
        apic[8].pt3dadd(13.29, 136.57, 9.82, 0.72)
        apic[8].pt3dadd(14.25, 136.82, 9.82, 0.72)
        apic[8].pt3dadd(14.98, 140.55, 9.82, 0.72)
        apic[8].pt3dadd(14.98, 141.79, 9.82, 0.72)
        apic[8].pt3dadd(15.46, 142.29, 9.82, 0.72)
        apic[8].pt3dadd(16.18, 143.53, 9.82, 0.72)
        apic[8].pt3dadd(16.43, 144.03, 9.82, 0.72)
        apic[8].pt3dadd(16.67, 145.52, 9.82, 0.72)
        apic[8].pt3dadd(16.91, 146.02, 9.82, 0.72)
        apic[8].pt3dadd(17.63, 147.26, 9.82, 0.72)
        apic[8].pt3dadd(18.36, 148.26, 9.82, 0.72)
        apic[8].pt3dadd(18.60, 148.51, 9.82, 0.72)
        apic[8].pt3dadd(19.81, 149.75, 9.82, 0.72)
        apic[8].pt3dadd(20.77, 150.00, 9.82, 0.72)
        apic[8].pt3dadd(22.22, 150.00, 9.82, 0.72)
        apic[8].pt3dadd(23.91, 150.00, 9.82, 0.72)
        apic[8].pt3dadd(24.40, 150.00, 9.82, 0.72)
        apic[8].pt3dadd(25.36, 150.50, 9.82, 0.72)
        apic[8].pt3dadd(26.81, 151.24, 9.82, 0.72)
        apic[8].pt3dadd(27.29, 151.74, 9.82, 0.72)
        apic[8].pt3dadd(27.54, 154.23, 9.82, 0.72)
        apic[8].pt3dadd(27.54, 155.72, 9.82, 0.72)
        apic[8].pt3dadd(27.78, 157.46, 9.82, 0.72)
        apic[8].pt3dadd(28.02, 157.71, 9.82, 0.72)
        apic[8].pt3dadd(30.19, 158.96, 9.82, 0.72)
        apic[8].pt3dadd(34.06, 161.44, 7.99, 0.54)
        apic[8].pt3dadd(36.23, 164.18, 7.99, 0.54)
        apic[8].pt3dadd(39.61, 165.17, 7.99, 0.54)
        apic[8].pt3dadd(40.34, 165.17, 7.99, 0.54)
        apic[8].pt3dadd(43.48, 166.17, 7.99, 0.54)
        apic[8].pt3dadd(44.44, 166.67, 7.99, 0.54)
        apic[8].pt3dadd(45.17, 167.91, 7.99, 0.54)
        apic[8].pt3dadd(46.14, 168.16, 7.99, 0.54)
        apic[8].pt3dadd(47.10, 170.40, 7.99, 0.54)
        apic[8].pt3dadd(47.83, 171.64, 7.99, 0.54)
        apic[8].pt3dadd(48.79, 174.63, 7.99, 0.54)
        apic[8].pt3dadd(49.28, 176.87, 16.62, 0.54)
        apic[8].pt3dadd(49.28, 176.87, 16.67, 0.54)
        apic[8].pt3dadd(49.28, 182.34, 16.67, 0.54)
        apic[8].pt3dadd(50.24, 183.33, 16.67, 0.54)
        apic[8].pt3dadd(51.69, 184.08, 16.68, 0.54)
        apic[8].pt3dadd(51.93, 184.58, 16.68, 0.54)
        apic[8].pt3dadd(52.17, 185.82, 16.68, 0.54)
        apic[8].pt3dadd(54.11, 187.56, 16.69, 0.54)
        apic[8].pt3dadd(55.31, 189.80, 16.69, 0.54)
        apic[8].pt3dadd(55.56, 191.04, 16.69, 0.54)
        apic[8].pt3dadd(54.35, 196.77, 16.69, 0.54)
        apic[8].pt3dadd(54.11, 196.77, 16.69, 0.54)
        apic[8].pt3dadd(53.38, 199.25, 10.83, 0.54)
        apic[8].pt3dadd(53.38, 199.25, 10.78, 0.54)
        apic[8].pt3dadd(53.86, 202.74, 7.34, 0.54)
        apic[8].pt3dadd(53.86, 202.74, 7.27, 0.54)
        apic[8].pt3dadd(55.31, 204.73, 7.22, 0.54)
        apic[8].pt3dadd(55.56, 204.73, 7.16, 0.48)
        apic[8].pt3dadd(60.39, 209.20, 8.82, 0.48)
        apic[8].pt3dadd(60.39, 209.20, 8.81, 0.48)
        apic[8].pt3dadd(62.56, 210.70, 8.78, 0.48)
        apic[8].pt3dadd(63.53, 211.19, 8.77, 0.48)
        apic[8].pt3dadd(64.49, 211.44, 8.76, 0.48)
        apic[8].pt3dadd(64.73, 211.69, 8.76, 0.48)
        apic[8].pt3dadd(65.94, 216.42, 8.60, 0.48)
        apic[8].pt3dadd(62.32, 218.16, 12.01, 0.48)
        apic[8].pt3dadd(61.35, 221.89, 12.01, 0.48)
        apic[8].pt3dadd(61.35, 222.39, 12.01, 0.48)
        apic[8].pt3dadd(61.59, 224.38, 12.01, 0.48)
        apic[8].pt3dadd(62.32, 224.88, 12.02, 0.48)
        apic[8].pt3dadd(62.80, 225.62, 12.02, 0.48)
        apic[8].pt3dadd(63.04, 226.12, 12.02, 0.48)
        apic[8].pt3dadd(63.53, 226.87, 12.02, 0.48)
        apic[8].pt3dadd(64.01, 227.36, 12.02, 0.48)
        apic[8].pt3dadd(64.98, 228.86, 12.02, 0.48)
        apic[8].pt3dadd(65.22, 232.84, 12.04, 0.48)
        apic[8].pt3dadd(65.46, 234.08, 12.07, 0.48)
        apic[8].pt3dadd(66.67, 236.07, 12.08, 0.48)
        apic[8].pt3dadd(67.15, 236.32, 12.08, 0.48)
        apic[8].pt3dadd(69.08, 237.06, 12.08, 0.48)
        apic[8].pt3dadd(71.26, 238.06, 12.08, 0.48)
        apic[8].pt3dadd(71.74, 238.06, 12.08, 0.48)
        apic[8].pt3dadd(72.46, 238.31, 12.08, 0.48)
        apic[8].pt3dadd(74.64, 240.55, 16.25, 0.48)
        apic[8].pt3dadd(74.64, 240.55, 16.26, 0.48)
        apic[8].pt3dadd(76.81, 244.03, 16.79, 0.48)
        apic[8].pt3dadd(77.29, 244.28, 16.84, 0.48)
        apic[8].pt3dadd(78.26, 245.52, 17.19, 0.48)
        apic[8].pt3dadd(78.50, 245.77, 17.34, 0.48)
        apic[8].pt3dadd(78.74, 246.77, 17.70, 0.48)
        apic[8].pt3dadd(78.74, 247.26, 17.83, 0.48)
        apic[8].pt3dadd(79.23, 248.26, 17.88, 0.48)
        apic[8].pt3dadd(79.47, 248.76, 17.91, 0.48)
        apic[8].pt3dadd(80.19, 250.50, 17.92, 0.48)
        apic[8].pt3dadd(80.68, 251.49, 17.93, 0.48)
        apic[8].pt3dadd(81.64, 252.24, 17.93, 0.48)
        apic[8].pt3dadd(82.13, 252.99, 17.94, 0.48)
        apic[8].pt3dadd(82.85, 253.98, 17.94, 0.48)
        apic[8].pt3dadd(84.54, 256.47, 17.95, 0.48)
        apic[8].pt3dadd(85.51, 256.97, 17.95, 0.48)
        apic[8].pt3dadd(86.47, 258.71, 17.96, 0.48)
        
        apic[9] = h.Section(name='apic[9]')
        apic[9].connect(        apic[7](1), 0)
        apic[9].pt3dclear()
        apic[9].pt3dadd(1.45, 115.17, 11.70, 0.77)
        apic[9].pt3dadd(-0.24, 114.18, 8.33, 0.77)
        apic[9].pt3dadd(-2.42, 113.43, 8.33, 0.77)
        apic[9].pt3dadd(-4.35, 112.94, 8.33, 0.77)
        apic[9].pt3dadd(-5.80, 112.69, 8.33, 0.77)
        
        apic[10] = h.Section(name='apic[10]')
        apic[10].connect(        apic[9](1), 0)
        apic[10].pt3dclear()
        apic[10].pt3dadd(-5.80, 112.69, 8.33, 0.43)
        apic[10].pt3dadd(-6.52, 116.42, 11.62, 0.43)
        apic[10].pt3dadd(-6.52, 116.42, 11.63, 0.43)
        apic[10].pt3dadd(-7.49, 119.65, 11.65, 0.43)
        apic[10].pt3dadd(-7.49, 121.14, 11.66, 0.43)
        apic[10].pt3dadd(-8.94, 124.88, 11.92, 0.43)
        apic[10].pt3dadd(-9.42, 125.12, 11.96, 0.43)
        apic[10].pt3dadd(-10.63, 125.87, 12.27, 0.43)
        apic[10].pt3dadd(-10.63, 125.87, 12.29, 0.43)
        apic[10].pt3dadd(-11.59, 127.86, 12.40, 0.43)
        apic[10].pt3dadd(-11.84, 127.86, 12.40, 0.43)
        apic[10].pt3dadd(-13.77, 129.10, 12.42, 0.43)
        apic[10].pt3dadd(-14.49, 130.10, 12.42, 0.43)
        apic[10].pt3dadd(-14.49, 131.09, 12.42, 0.43)
        apic[10].pt3dadd(-15.22, 132.34, 12.42, 0.43)
        apic[10].pt3dadd(-15.70, 132.59, 12.42, 0.43)
        apic[10].pt3dadd(-18.12, 135.07, 12.07, 0.43)
        apic[10].pt3dadd(-18.12, 135.07, 12.06, 0.43)
        apic[10].pt3dadd(-19.08, 135.82, 12.06, 0.43)
        apic[10].pt3dadd(-20.05, 135.82, 12.06, 0.43)
        apic[10].pt3dadd(-22.22, 135.82, 12.06, 0.43)
        
        apic[11] = h.Section(name='apic[11]')
        apic[11].connect(        apic[9](1), 0)
        apic[11].pt3dclear()
        apic[11].pt3dadd(-5.80, 112.69, 8.33, 0.61)
        apic[11].pt3dadd(-5.80, 110.70, 8.31, 0.61)
        apic[11].pt3dadd(-6.04, 109.70, 8.31, 0.61)
        apic[11].pt3dadd(-7.00, 108.21, 8.31, 0.61)
        apic[11].pt3dadd(-7.25, 107.71, 8.31, 0.61)
        apic[11].pt3dadd(-7.73, 107.46, 8.31, 0.61)
        apic[11].pt3dadd(-7.97, 107.46, 8.31, 0.61)
        apic[11].pt3dadd(-8.21, 106.97, 8.31, 0.61)
        
        apic[12] = h.Section(name='apic[12]')
        apic[12].connect(        apic[11](1), 0)
        apic[12].pt3dclear()
        apic[12].pt3dadd(-8.21, 106.97, 8.31, 0.43)
        apic[12].pt3dadd(-9.90, 103.23, 6.08, 0.43)
        apic[12].pt3dadd(-11.84, 103.23, 6.08, 0.43)
        apic[12].pt3dadd(-12.56, 103.23, 6.08, 0.43)
        apic[12].pt3dadd(-16.91, 103.23, 6.08, 0.43)
        apic[12].pt3dadd(-17.15, 103.23, 6.08, 0.43)
        apic[12].pt3dadd(-18.36, 102.74, 6.08, 0.43)
        apic[12].pt3dadd(-19.08, 101.74, 6.08, 0.43)
        apic[12].pt3dadd(-19.81, 101.00, 6.08, 0.43)
        apic[12].pt3dadd(-20.29, 100.25, 6.08, 0.43)
        apic[12].pt3dadd(-21.01, 99.25, 6.10, 0.43)
        apic[12].pt3dadd(-21.01, 98.01, 6.10, 0.43)
        apic[12].pt3dadd(-21.50, 95.02, 9.16, 0.43)
        apic[12].pt3dadd(-24.40, 93.53, 9.17, 0.43)
        apic[12].pt3dadd(-24.64, 93.53, 9.17, 0.43)
        apic[12].pt3dadd(-26.09, 94.28, 9.18, 0.43)
        apic[12].pt3dadd(-28.26, 96.02, 9.18, 0.43)
        apic[12].pt3dadd(-31.16, 97.02, 9.18, 0.43)
        apic[12].pt3dadd(-35.27, 95.52, 9.19, 0.43)
        apic[12].pt3dadd(-37.20, 93.53, 9.19, 0.43)
        apic[12].pt3dadd(-37.44, 93.53, 9.20, 0.43)
        apic[12].pt3dadd(-40.82, 93.28, 13.41, 0.43)
        apic[12].pt3dadd(-42.03, 93.28, 13.42, 0.43)
        apic[12].pt3dadd(-42.27, 93.28, 13.44, 0.43)
        apic[12].pt3dadd(-43.48, 92.29, 13.78, 0.43)
        apic[12].pt3dadd(-43.96, 92.04, 13.78, 0.43)
        apic[12].pt3dadd(-45.65, 91.29, 14.78, 0.43)
        apic[12].pt3dadd(-45.65, 91.29, 15.04, 0.43)
        apic[12].pt3dadd(-46.86, 89.30, 15.37, 0.43)
        apic[12].pt3dadd(-47.58, 88.56, 15.39, 0.43)
        apic[12].pt3dadd(-48.31, 87.81, 15.44, 0.43)
        apic[12].pt3dadd(-49.28, 87.56, 15.55, 0.43)
        apic[12].pt3dadd(-49.52, 87.56, 15.58, 0.43)
        
        apic[13] = h.Section(name='apic[13]')
        apic[13].connect(        apic[11](1), 0)
        apic[13].pt3dclear()
        apic[13].pt3dadd(-8.21, 106.97, 8.31, 0.43)
        apic[13].pt3dadd(-10.87, 108.46, -1.11, 0.43)
        apic[13].pt3dadd(-14.25, 110.45, -1.11, 0.43)
        apic[13].pt3dadd(-14.49, 110.45, -1.11, 0.43)
        apic[13].pt3dadd(-16.18, 111.69, -1.11, 0.43)
        apic[13].pt3dadd(-17.15, 111.94, -1.11, 0.43)
        apic[13].pt3dadd(-19.57, 112.69, -1.11, 0.43)
        apic[13].pt3dadd(-21.26, 113.18, -1.11, 0.43)
        apic[13].pt3dadd(-23.19, 114.18, -1.11, 0.43)
        apic[13].pt3dadd(-23.43, 114.18, -1.11, 0.43)
        apic[13].pt3dadd(-24.15, 114.93, -1.11, 0.43)
        apic[13].pt3dadd(-24.40, 115.42, -1.11, 0.43)
        apic[13].pt3dadd(-26.09, 118.91, -1.11, 0.43)
        apic[13].pt3dadd(-26.81, 119.40, -1.11, 0.43)
        apic[13].pt3dadd(-28.50, 122.64, -1.11, 0.43)
        apic[13].pt3dadd(-28.50, 122.89, -1.11, 0.43)
        apic[13].pt3dadd(-30.19, 124.38, -1.11, 0.43)
        apic[13].pt3dadd(-34.54, 125.62, -1.11, 0.43)
        apic[13].pt3dadd(-37.44, 125.62, -1.11, 0.43)
        apic[13].pt3dadd(-41.06, 126.37, -1.11, 0.43)
        apic[13].pt3dadd(-42.75, 126.62, -1.11, 0.43)
        apic[13].pt3dadd(-43.00, 126.62, -1.11, 0.43)
        apic[13].pt3dadd(-44.69, 126.62, -1.11, 0.43)
        apic[13].pt3dadd(-44.93, 128.36, -1.11, 0.43)
        apic[13].pt3dadd(-46.62, 129.35, -0.25, 0.43)
        apic[13].pt3dadd(-46.62, 129.35, -0.26, 0.43)
        apic[13].pt3dadd(-52.42, 129.35, -0.26, 0.43)
        apic[13].pt3dadd(-54.35, 128.36, -0.26, 0.43)
        apic[13].pt3dadd(-55.07, 128.11, -0.26, 0.43)
        apic[13].pt3dadd(-58.45, 127.61, -0.27, 0.43)
        apic[13].pt3dadd(-58.94, 127.61, -0.28, 0.43)
        apic[13].pt3dadd(-61.84, 127.36, -0.29, 0.43)
        apic[13].pt3dadd(-62.08, 127.36, -0.30, 0.43)
        apic[13].pt3dadd(-64.98, 130.85, -0.30, 0.43)
        apic[13].pt3dadd(-65.22, 130.85, -0.30, 0.43)
        apic[13].pt3dadd(-68.60, 133.83, -0.30, 0.43)
        apic[13].pt3dadd(-71.50, 136.07, -0.30, 0.43)
        apic[13].pt3dadd(-72.22, 139.05, -0.30, 0.43)
        apic[13].pt3dadd(-72.22, 139.55, -0.30, 0.43)
        apic[13].pt3dadd(-71.98, 141.04, -0.30, 0.43)
        apic[13].pt3dadd(-71.98, 141.29, -0.30, 0.43)
        apic[13].pt3dadd(-71.74, 142.79, -0.30, 0.43)
        apic[13].pt3dadd(-71.74, 143.78, -0.30, 0.43)
        
        apic[14] = h.Section(name='apic[14]')
        apic[14].connect(        apic[6](1), 0)
        apic[14].pt3dclear()
        apic[14].pt3dadd(10.14, 111.19, 6.45, 3.47)
        apic[14].pt3dadd(11.35, 112.69, 11.55, 3.47)
        apic[14].pt3dadd(12.32, 114.43, 11.55, 3.47)
        
        apic[15] = h.Section(name='apic[15]')
        apic[15].connect(        apic[14](1), 0)
        apic[15].pt3dclear()
        apic[15].pt3dadd(12.32, 114.43, 11.55, 0.48)
        apic[15].pt3dadd(11.11, 116.67, 20.69, 0.48)
        apic[15].pt3dadd(11.11, 119.15, 20.69, 0.48)
        apic[15].pt3dadd(10.87, 120.40, 20.69, 0.48)
        apic[15].pt3dadd(10.87, 121.64, 20.68, 0.48)
        apic[15].pt3dadd(11.11, 122.14, 20.68, 0.48)
        apic[15].pt3dadd(11.59, 122.64, 20.68, 0.48)
        apic[15].pt3dadd(12.32, 122.64, 20.68, 0.48)
        apic[15].pt3dadd(13.29, 123.38, 20.68, 0.48)
        apic[15].pt3dadd(13.53, 125.12, 20.68, 0.48)
        apic[15].pt3dadd(13.53, 125.87, 20.68, 0.48)
        apic[15].pt3dadd(13.29, 127.11, 20.68, 0.48)
        apic[15].pt3dadd(12.56, 128.36, 20.68, 0.48)
        apic[15].pt3dadd(12.32, 128.86, 20.68, 0.48)
        
        apic[16] = h.Section(name='apic[16]')
        apic[16].connect(        apic[14](1), 0)
        apic[16].pt3dclear()
        apic[16].pt3dadd(12.32, 114.43, 11.55, 2.90)
        apic[16].pt3dadd(14.73, 116.92, 4.56, 2.90)
        apic[16].pt3dadd(16.43, 119.15, 4.56, 2.90)
        apic[16].pt3dadd(18.84, 123.13, 4.56, 2.90)
        apic[16].pt3dadd(19.57, 126.12, 4.56, 2.90)
        apic[16].pt3dadd(21.01, 129.60, 4.56, 2.90)
        apic[16].pt3dadd(22.46, 133.33, 8.97, 2.80)
        apic[16].pt3dadd(23.19, 136.32, 8.97, 2.80)
        
        apic[17] = h.Section(name='apic[17]')
        apic[17].connect(        apic[16](1), 0)
        apic[17].pt3dclear()
        apic[17].pt3dadd(23.19, 136.32, 8.97, 0.72)
        apic[17].pt3dadd(20.53, 135.07, 5.56, 0.72)
        apic[17].pt3dadd(19.08, 135.82, 5.56, 0.72)
        apic[17].pt3dadd(18.12, 137.56, 5.56, 0.72)
        apic[17].pt3dadd(18.12, 139.55, 5.56, 0.72)
        apic[17].pt3dadd(18.36, 139.80, 5.56, 0.72)
        apic[17].pt3dadd(19.57, 140.80, 5.56, 0.72)
        apic[17].pt3dadd(20.05, 140.80, 5.56, 0.72)
        apic[17].pt3dadd(24.40, 142.29, -6.80, 0.72)
        apic[17].pt3dadd(24.40, 142.29, -6.81, 0.72)
        apic[17].pt3dadd(23.91, 144.78, -7.48, 0.72)
        apic[17].pt3dadd(23.91, 144.78, -7.52, 0.72)
        apic[17].pt3dadd(23.91, 148.01, -8.06, 0.72)
        apic[17].pt3dadd(23.91, 148.01, -8.09, 0.72)
        apic[17].pt3dadd(24.64, 149.25, -8.18, 0.72)
        apic[17].pt3dadd(24.88, 149.25, -8.18, 0.72)
        apic[17].pt3dadd(27.05, 148.76, -6.85, 0.72)
        apic[17].pt3dadd(28.02, 148.76, -6.84, 0.72)
        apic[17].pt3dadd(29.95, 148.26, -6.83, 0.72)
        apic[17].pt3dadd(30.19, 148.26, -6.83, 0.72)
        apic[17].pt3dadd(30.92, 148.76, -6.82, 0.72)
        apic[17].pt3dadd(31.40, 148.76, -6.82, 0.72)
        apic[17].pt3dadd(31.88, 148.76, -6.82, 0.72)
        apic[17].pt3dadd(31.88, 148.26, -6.81, 0.72)
        apic[17].pt3dadd(31.88, 147.26, -6.81, 0.72)
        apic[17].pt3dadd(31.88, 146.27, -6.81, 0.72)
        apic[17].pt3dadd(31.88, 146.27, -6.81, 0.72)
        
        apic[18] = h.Section(name='apic[18]')
        apic[18].connect(        apic[17](1), 0)
        apic[18].pt3dclear()
        apic[18].pt3dadd(31.88, 146.27, -6.81, 0.43)
        apic[18].pt3dadd(31.88, 143.28, -9.93, 0.43)
        apic[18].pt3dadd(32.13, 143.28, -9.93, 0.43)
        apic[18].pt3dadd(33.82, 141.79, -9.93, 0.43)
        apic[18].pt3dadd(34.30, 141.79, -9.93, 0.43)
        apic[18].pt3dadd(35.99, 142.29, -9.93, 0.43)
        apic[18].pt3dadd(36.23, 142.29, -9.95, 0.43)
        apic[18].pt3dadd(36.96, 142.54, -9.96, 0.43)
        apic[18].pt3dadd(37.20, 142.79, -9.97, 0.43)
        apic[18].pt3dadd(37.92, 143.53, -9.97, 0.43)
        apic[18].pt3dadd(39.61, 144.03, -9.98, 0.43)
        apic[18].pt3dadd(39.86, 144.28, -9.98, 0.43)
        apic[18].pt3dadd(41.79, 144.28, -9.99, 0.43)
        apic[18].pt3dadd(42.27, 144.28, -9.99, 0.43)
        apic[18].pt3dadd(43.96, 142.79, -10.00, 0.43)
        apic[18].pt3dadd(44.20, 142.29, -10.00, 0.43)
        apic[18].pt3dadd(44.93, 141.79, -10.00, 0.43)
        apic[18].pt3dadd(45.89, 141.29, -10.00, 0.43)
        apic[18].pt3dadd(46.86, 140.05, -10.00, 0.43)
        apic[18].pt3dadd(48.79, 139.30, -13.17, 0.43)
        apic[18].pt3dadd(50.48, 142.04, -13.21, 0.43)
        apic[18].pt3dadd(51.21, 142.29, -13.22, 0.43)
        apic[18].pt3dadd(53.14, 142.79, -13.23, 0.43)
        apic[18].pt3dadd(53.62, 142.79, -13.23, 0.43)
        apic[18].pt3dadd(55.31, 142.29, -18.25, 0.43)
        apic[18].pt3dadd(55.31, 142.29, -18.29, 0.43)
        apic[18].pt3dadd(55.07, 138.56, -18.60, 0.43)
        apic[18].pt3dadd(55.07, 138.56, -18.61, 0.43)
        apic[18].pt3dadd(55.07, 136.32, -18.62, 0.43)
        apic[18].pt3dadd(55.07, 135.82, -18.65, 0.43)
        apic[18].pt3dadd(55.80, 134.83, -18.74, 0.43)
        apic[18].pt3dadd(56.28, 133.33, -18.87, 0.43)
        apic[18].pt3dadd(54.35, 131.34, -22.47, 0.43)
        apic[18].pt3dadd(54.11, 131.34, -22.49, 0.43)
        apic[18].pt3dadd(54.35, 130.35, -22.49, 0.43)
        apic[18].pt3dadd(55.07, 129.60, -22.49, 0.43)
        apic[18].pt3dadd(56.28, 129.10, -22.51, 0.43)
        apic[18].pt3dadd(56.76, 129.10, -22.52, 0.43)
        apic[18].pt3dadd(58.21, 129.10, -22.53, 0.43)
        apic[18].pt3dadd(59.42, 129.60, -22.53, 0.43)
        apic[18].pt3dadd(60.14, 129.85, -22.53, 0.43)
        apic[18].pt3dadd(60.39, 129.85, -22.54, 0.43)
        apic[18].pt3dadd(60.63, 130.10, -22.54, 0.43)
        apic[18].pt3dadd(60.87, 131.59, -22.54, 0.43)
        apic[18].pt3dadd(60.63, 131.84, -22.54, 0.43)
        apic[18].pt3dadd(60.63, 133.33, -22.54, 0.43)
        apic[18].pt3dadd(61.11, 133.58, -22.54, 0.43)
        apic[18].pt3dadd(62.56, 134.08, -22.54, 0.43)
        apic[18].pt3dadd(63.77, 134.08, -22.54, 0.43)
        apic[18].pt3dadd(68.36, 131.09, -22.54, 0.43)
        apic[18].pt3dadd(70.05, 130.10, -22.55, 0.43)
        apic[18].pt3dadd(72.22, 128.36, -31.04, 0.43)
        apic[18].pt3dadd(72.22, 128.61, -31.13, 0.43)
        apic[18].pt3dadd(75.12, 129.10, -31.39, 0.43)
        apic[18].pt3dadd(75.12, 128.86, -31.43, 0.43)
        apic[18].pt3dadd(76.33, 128.36, -31.44, 0.43)
        apic[18].pt3dadd(76.57, 128.36, -31.44, 0.43)
        apic[18].pt3dadd(76.81, 126.87, -31.30, 0.43)
        apic[18].pt3dadd(76.81, 126.87, -31.12, 0.43)
        apic[18].pt3dadd(75.85, 125.37, -32.66, 0.43)
        apic[18].pt3dadd(75.85, 125.12, -32.92, 0.43)
        apic[18].pt3dadd(74.40, 122.14, -34.38, 0.43)
        apic[18].pt3dadd(74.40, 122.14, -34.42, 0.43)
        apic[18].pt3dadd(72.46, 120.15, -34.60, 0.43)
        apic[18].pt3dadd(72.22, 119.40, -34.47, 0.43)
        apic[18].pt3dadd(71.74, 118.16, -34.47, 0.43)
        apic[18].pt3dadd(71.50, 117.41, -34.47, 0.43)
        apic[18].pt3dadd(70.05, 116.17, -34.47, 0.43)
        apic[18].pt3dadd(69.57, 113.93, -34.47, 0.43)
        apic[18].pt3dadd(68.84, 113.68, -34.45, 0.43)
        apic[18].pt3dadd(68.36, 113.43, -34.43, 0.43)
        apic[18].pt3dadd(68.12, 113.43, -34.43, 0.43)
        apic[18].pt3dadd(67.63, 113.43, -34.43, 0.43)
        apic[18].pt3dadd(64.49, 112.44, -36.79, 0.43)
        apic[18].pt3dadd(64.25, 112.44, -36.94, 0.43)
        apic[18].pt3dadd(62.32, 111.19, -37.02, 0.43)
        apic[18].pt3dadd(62.08, 110.95, -37.02, 0.43)
        apic[18].pt3dadd(61.35, 111.19, -37.02, 0.43)
        apic[18].pt3dadd(60.87, 111.69, -37.02, 0.43)
        apic[18].pt3dadd(60.39, 111.94, -37.02, 0.43)
        apic[18].pt3dadd(59.66, 111.94, -37.02, 0.43)
        apic[18].pt3dadd(58.70, 111.94, -37.02, 0.43)
        
        apic[19] = h.Section(name='apic[19]')
        apic[19].connect(        apic[17](1), 0)
        apic[19].pt3dclear()
        apic[19].pt3dadd(31.88, 146.27, -6.81, 0.43)
        apic[19].pt3dadd(30.68, 143.53, -11.00, 0.43)
        apic[19].pt3dadd(30.19, 138.81, -11.00, 0.43)
        apic[19].pt3dadd(30.43, 138.56, -11.00, 0.43)
        apic[19].pt3dadd(31.64, 137.31, -11.00, 0.43)
        apic[19].pt3dadd(32.37, 136.82, -11.00, 0.43)
        apic[19].pt3dadd(32.85, 134.83, -10.99, 0.43)
        apic[19].pt3dadd(32.85, 134.08, -10.99, 0.43)
        apic[19].pt3dadd(32.85, 132.59, -10.99, 0.43)
        apic[19].pt3dadd(32.61, 132.09, -10.99, 0.43)
        apic[19].pt3dadd(30.43, 131.59, -10.99, 0.43)
        apic[19].pt3dadd(30.19, 131.59, -10.99, 0.43)
        apic[19].pt3dadd(29.95, 130.85, -10.99, 0.43)
        apic[19].pt3dadd(29.71, 129.85, -10.99, 0.43)
        apic[19].pt3dadd(30.19, 128.61, -10.99, 0.43)
        apic[19].pt3dadd(29.95, 127.61, -10.99, 0.43)
        apic[19].pt3dadd(29.71, 126.62, -10.99, 0.43)
        apic[19].pt3dadd(29.47, 125.37, -10.98, 0.43)
        apic[19].pt3dadd(29.71, 125.12, -10.98, 0.43)
        apic[19].pt3dadd(29.95, 124.13, -10.98, 0.43)
        apic[19].pt3dadd(29.71, 123.38, -10.98, 0.43)
        apic[19].pt3dadd(29.23, 122.64, -10.98, 0.43)
        apic[19].pt3dadd(28.74, 121.89, -10.98, 0.43)
        apic[19].pt3dadd(28.74, 120.90, -11.03, 0.43)
        apic[19].pt3dadd(28.50, 120.15, -11.23, 0.43)
        apic[19].pt3dadd(26.81, 118.91, -11.30, 0.43)
        apic[19].pt3dadd(26.57, 118.91, -11.67, 0.43)
        apic[19].pt3dadd(25.85, 115.17, -15.65, 0.43)
        apic[19].pt3dadd(25.85, 114.93, -15.65, 0.43)
        apic[19].pt3dadd(26.09, 112.44, -15.65, 0.43)
        apic[19].pt3dadd(26.09, 111.19, -15.67, 0.43)
        apic[19].pt3dadd(25.85, 110.45, -15.67, 0.43)
        apic[19].pt3dadd(25.85, 109.45, -15.67, 0.43)
        apic[19].pt3dadd(25.12, 107.71, -18.42, 0.43)
        apic[19].pt3dadd(24.40, 105.72, -18.16, 0.43)
        apic[19].pt3dadd(24.15, 104.73, -18.01, 0.43)
        apic[19].pt3dadd(23.43, 101.24, -17.93, 0.43)
        apic[19].pt3dadd(23.43, 101.00, -17.93, 0.43)
        apic[19].pt3dadd(23.91, 99.50, -17.93, 0.43)
        apic[19].pt3dadd(24.40, 97.51, -17.93, 0.43)
        apic[19].pt3dadd(24.40, 97.01, -17.93, 0.43)
        apic[19].pt3dadd(24.40, 96.02, -17.93, 0.43)
        apic[19].pt3dadd(23.91, 94.78, -17.93, 0.43)
        apic[19].pt3dadd(23.43, 93.53, -17.93, 0.43)
        apic[19].pt3dadd(22.95, 92.79, -17.93, 0.43)
        apic[19].pt3dadd(22.71, 92.54, -17.93, 0.43)
        apic[19].pt3dadd(22.46, 92.29, -17.93, 0.43)
        apic[19].pt3dadd(22.22, 91.79, -17.93, 0.43)
        
        apic[20] = h.Section(name='apic[20]')
        apic[20].connect(        apic[16](1), 0)
        apic[20].pt3dclear()
        apic[20].pt3dadd(23.19, 136.32, 8.97, 3.14)
        apic[20].pt3dadd(22.46, 138.06, 6.84, 3.14)
        apic[20].pt3dadd(21.98, 140.80, 6.83, 3.14)
        apic[20].pt3dadd(21.74, 140.80, 6.83, 3.14)
        apic[20].pt3dadd(21.26, 142.29, 6.83, 3.14)
        apic[20].pt3dadd(21.01, 144.28, 6.83, 3.14)
        apic[20].pt3dadd(20.77, 144.53, 6.83, 3.14)
        apic[20].pt3dadd(20.77, 146.02, 6.83, 3.14)
        apic[20].pt3dadd(20.53, 146.27, 6.83, 3.14)
        
        apic[21] = h.Section(name='apic[21]')
        apic[21].connect(        apic[20](1), 0)
        apic[21].pt3dclear()
        apic[21].pt3dadd(20.53, 146.27, 6.83, 0.72)
        apic[21].pt3dadd(18.12, 145.52, 6.65, 0.72)
        apic[21].pt3dadd(16.43, 147.26, 6.65, 0.72)
        apic[21].pt3dadd(16.18, 147.26, 6.65, 0.72)
        apic[21].pt3dadd(15.22, 149.25, 6.65, 0.72)
        apic[21].pt3dadd(14.98, 149.25, 6.65, 0.72)
        apic[21].pt3dadd(13.77, 151.74, 1.92, 0.72)
        apic[21].pt3dadd(13.53, 151.74, 1.82, 0.72)
        apic[21].pt3dadd(12.08, 152.49, 1.38, 0.72)
        apic[21].pt3dadd(11.59, 152.74, 1.33, 0.72)
        apic[21].pt3dadd(10.14, 153.48, 1.11, 0.72)
        apic[21].pt3dadd(9.42, 153.73, 0.98, 0.72)
        apic[21].pt3dadd(8.21, 154.23, 0.94, 0.72)
        apic[21].pt3dadd(7.49, 154.48, 0.94, 0.72)
        apic[21].pt3dadd(6.28, 154.98, 0.94, 0.72)
        apic[21].pt3dadd(6.04, 155.22, 0.94, 0.72)
        apic[21].pt3dadd(5.80, 156.22, 0.94, 0.72)
        
        apic[22] = h.Section(name='apic[22]')
        apic[22].connect(        apic[21](1), 0)
        apic[22].pt3dclear()
        apic[22].pt3dadd(5.80, 156.22, 0.94, 0.39)
        apic[22].pt3dadd(3.38, 155.47, 0.35, 0.39)
        apic[22].pt3dadd(2.66, 154.23, 0.35, 0.39)
        apic[22].pt3dadd(2.17, 153.98, 0.35, 0.39)
        apic[22].pt3dadd(1.21, 152.74, 0.35, 0.39)
        apic[22].pt3dadd(1.21, 151.74, 0.35, 0.39)
        apic[22].pt3dadd(0.97, 149.75, 0.35, 0.39)
        apic[22].pt3dadd(0.72, 149.25, 0.35, 0.39)
        apic[22].pt3dadd(0.48, 148.51, 0.35, 0.39)
        apic[22].pt3dadd(0.97, 148.26, 0.35, 0.39)
        apic[22].pt3dadd(1.21, 148.01, 0.35, 0.39)
        apic[22].pt3dadd(1.93, 147.76, 0.35, 0.39)
        apic[22].pt3dadd(2.66, 148.01, 0.35, 0.39)
        apic[22].pt3dadd(2.90, 148.51, 0.35, 0.39)
        apic[22].pt3dadd(2.90, 149.25, 0.35, 0.39)
        apic[22].pt3dadd(3.14, 149.75, 0.35, 0.39)
        apic[22].pt3dadd(3.14, 150.25, 0.35, 0.39)
        apic[22].pt3dadd(3.14, 150.75, 0.35, 0.39)
        apic[22].pt3dadd(3.14, 151.49, 0.35, 0.39)
        apic[22].pt3dadd(2.66, 151.74, 0.35, 0.39)
        apic[22].pt3dadd(-0.24, 151.74, -6.96, 0.39)
        apic[22].pt3dadd(-3.62, 151.99, -10.66, 0.39)
        apic[22].pt3dadd(-4.83, 152.74, -10.66, 0.39)
        apic[22].pt3dadd(-5.07, 153.98, -10.66, 0.39)
        apic[22].pt3dadd(-4.83, 154.48, -10.66, 0.39)
        apic[22].pt3dadd(-4.11, 155.72, -10.66, 0.39)
        apic[22].pt3dadd(-3.14, 155.47, -10.66, 0.39)
        apic[22].pt3dadd(-1.93, 154.98, -10.66, 0.39)
        apic[22].pt3dadd(-1.45, 154.73, -10.66, 0.39)
        apic[22].pt3dadd(-0.97, 154.23, -10.66, 0.39)
        apic[22].pt3dadd(-2.17, 150.00, -11.57, 0.39)
        apic[22].pt3dadd(-2.17, 150.00, -11.58, 0.39)
        apic[22].pt3dadd(-2.90, 149.50, -14.40, 0.39)
        apic[22].pt3dadd(-3.14, 149.75, -14.63, 0.39)
        apic[22].pt3dadd(-5.56, 150.75, -15.16, 0.39)
        apic[22].pt3dadd(-5.80, 150.75, -15.16, 0.39)
        apic[22].pt3dadd(-7.25, 150.75, -15.16, 0.39)
        apic[22].pt3dadd(-7.49, 150.75, -15.16, 0.39)
        apic[22].pt3dadd(-8.45, 150.25, -15.16, 0.39)
        apic[22].pt3dadd(-8.70, 148.76, -15.16, 0.39)
        apic[22].pt3dadd(-8.45, 147.51, -15.16, 0.39)
        apic[22].pt3dadd(-7.25, 147.02, -15.16, 0.39)
        apic[22].pt3dadd(-6.76, 147.02, -15.16, 0.39)
        apic[22].pt3dadd(-6.76, 147.26, -20.56, 0.39)
        apic[22].pt3dadd(-6.76, 147.26, -20.60, 0.39)
        apic[22].pt3dadd(-7.97, 148.01, -20.63, 0.39)
        apic[22].pt3dadd(-9.66, 147.76, -20.63, 0.39)
        apic[22].pt3dadd(-10.14, 147.26, -20.63, 0.39)
        apic[22].pt3dadd(-10.63, 146.77, -20.62, 0.39)
        apic[22].pt3dadd(-12.32, 146.27, -20.62, 0.39)
        apic[22].pt3dadd(-12.56, 146.02, -20.62, 0.39)
        apic[22].pt3dadd(-13.04, 144.78, -20.62, 0.39)
        apic[22].pt3dadd(-13.77, 144.28, -20.62, 0.39)
        apic[22].pt3dadd(-14.98, 144.03, -20.49, 0.39)
        apic[22].pt3dadd(-14.98, 143.78, -20.49, 0.39)
        apic[22].pt3dadd(-16.43, 141.29, -27.28, 0.39)
        apic[22].pt3dadd(-17.39, 141.54, -27.28, 0.39)
        apic[22].pt3dadd(-17.39, 141.54, -27.26, 0.39)
        apic[22].pt3dadd(-17.63, 141.79, -27.15, 0.39)
        apic[22].pt3dadd(-18.36, 142.54, -27.15, 0.39)
        apic[22].pt3dadd(-18.60, 142.54, -27.15, 0.39)
        apic[22].pt3dadd(-21.26, 144.03, -27.15, 0.39)
        apic[22].pt3dadd(-21.50, 144.03, -27.15, 0.39)
        apic[22].pt3dadd(-23.19, 144.03, -27.15, 0.39)
        apic[22].pt3dadd(-23.67, 144.03, -27.15, 0.39)
        apic[22].pt3dadd(-27.05, 143.78, -27.15, 0.39)
        apic[22].pt3dadd(-27.54, 143.78, -27.15, 0.39)
        apic[22].pt3dadd(-30.68, 144.53, -32.82, 0.39)
        apic[22].pt3dadd(-32.85, 144.03, -33.00, 0.39)
        apic[22].pt3dadd(-32.85, 144.03, -33.05, 0.39)
        apic[22].pt3dadd(-34.78, 142.29, -33.67, 0.39)
        apic[22].pt3dadd(-34.78, 141.79, -33.78, 0.39)
        apic[22].pt3dadd(-36.47, 139.30, -33.91, 0.39)
        apic[22].pt3dadd(-36.47, 139.30, -33.93, 0.39)
        apic[22].pt3dadd(-39.86, 138.06, -38.01, 0.39)
        apic[22].pt3dadd(-39.86, 137.81, -38.33, 0.39)
        apic[22].pt3dadd(-41.79, 137.31, -39.71, 0.39)
        apic[22].pt3dadd(-42.03, 137.06, -39.90, 0.39)
        apic[22].pt3dadd(-46.38, 136.82, -39.99, 0.39)
        apic[22].pt3dadd(-46.62, 136.82, -40.00, 0.39)
        apic[22].pt3dadd(-49.03, 135.32, -40.01, 0.39)
        apic[22].pt3dadd(-50.00, 135.32, -40.01, 0.39)
        apic[22].pt3dadd(-50.72, 135.32, -40.02, 0.39)
        apic[22].pt3dadd(-51.21, 135.82, -40.02, 0.39)
        apic[22].pt3dadd(-51.69, 136.57, -40.02, 0.39)
        apic[22].pt3dadd(-51.93, 136.57, -40.02, 0.39)
        apic[22].pt3dadd(-54.35, 137.31, -40.03, 0.39)
        apic[22].pt3dadd(-55.31, 137.56, -40.05, 0.39)
        apic[22].pt3dadd(-55.31, 138.06, -40.05, 0.39)
        apic[22].pt3dadd(-55.56, 138.06, -40.05, 0.39)
        apic[22].pt3dadd(-56.76, 138.31, -40.05, 0.39)
        
        apic[23] = h.Section(name='apic[23]')
        apic[23].connect(        apic[21](1), 0)
        apic[23].pt3dclear()
        apic[23].pt3dadd(5.80, 156.22, 0.94, 0.54)
        apic[23].pt3dadd(6.04, 158.21, -3.16, 0.54)
        apic[23].pt3dadd(5.07, 159.20, -3.16, 0.54)
        apic[23].pt3dadd(5.07, 161.94, -3.16, 0.54)
        apic[23].pt3dadd(5.07, 162.19, -3.16, 0.54)
        apic[23].pt3dadd(3.38, 163.68, -3.16, 0.54)
        apic[23].pt3dadd(2.90, 163.68, -3.16, 0.54)
        apic[23].pt3dadd(1.93, 164.18, -3.16, 0.54)
        apic[23].pt3dadd(1.69, 164.18, -3.16, 0.54)
        apic[23].pt3dadd(0.97, 164.43, -3.16, 0.54)
        apic[23].pt3dadd(0.72, 164.68, -3.16, 0.54)
        
        apic[24] = h.Section(name='apic[24]')
        apic[24].connect(        apic[23](1), 0)
        apic[24].pt3dclear()
        apic[24].pt3dadd(0.72, 164.68, -3.16, 0.54)
        apic[24].pt3dadd(-1.69, 165.17, -5.47, 0.54)
        apic[24].pt3dadd(-4.59, 166.42, -5.47, 0.54)
        apic[24].pt3dadd(-5.80, 170.65, -5.47, 0.54)
        apic[24].pt3dadd(-7.25, 172.89, -5.47, 0.54)
        apic[24].pt3dadd(-7.73, 173.13, -5.47, 0.54)
        apic[24].pt3dadd(-8.70, 173.88, -5.47, 0.54)
        apic[24].pt3dadd(-9.18, 174.13, -5.47, 0.54)
        apic[24].pt3dadd(-10.87, 175.12, -5.47, 0.54)
        apic[24].pt3dadd(-11.35, 177.86, -5.47, 0.54)
        apic[24].pt3dadd(-12.56, 182.34, -5.47, 0.54)
        apic[24].pt3dadd(-17.63, 188.81, -5.31, 0.54)
        apic[24].pt3dadd(-20.05, 191.54, -5.31, 0.54)
        apic[24].pt3dadd(-20.29, 191.54, -5.31, 0.54)
        apic[24].pt3dadd(-21.50, 194.03, -5.31, 0.54)
        apic[24].pt3dadd(-23.43, 197.26, -5.31, 0.54)
        apic[24].pt3dadd(-25.12, 198.76, -5.31, 0.54)
        apic[24].pt3dadd(-25.85, 199.50, -5.32, 0.54)
        apic[24].pt3dadd(-27.54, 200.25, -4.08, 0.54)
        apic[24].pt3dadd(-28.74, 202.74, -4.08, 0.54)
        apic[24].pt3dadd(-28.50, 206.97, -4.08, 0.54)
        apic[24].pt3dadd(-28.26, 206.97, -4.08, 0.54)
        apic[24].pt3dadd(-27.29, 208.46, -4.08, 0.54)
        apic[24].pt3dadd(-26.09, 210.95, -4.08, 0.54)
        apic[24].pt3dadd(-24.88, 211.94, -4.08, 0.54)
        apic[24].pt3dadd(-24.64, 212.94, -4.08, 0.54)
        apic[24].pt3dadd(-24.40, 213.18, -4.08, 0.54)
        apic[24].pt3dadd(-24.64, 215.67, -4.08, 0.54)
        apic[24].pt3dadd(-25.85, 218.66, -4.08, 0.54)
        apic[24].pt3dadd(-25.85, 220.40, 3.08, 0.54)
        apic[24].pt3dadd(-27.54, 222.14, 3.08, 0.54)
        apic[24].pt3dadd(-28.26, 222.39, 3.08, 0.54)
        apic[24].pt3dadd(-29.23, 223.88, 3.08, 0.54)
        apic[24].pt3dadd(-28.99, 223.88, 3.08, 0.54)
        apic[24].pt3dadd(-27.54, 225.37, 3.08, 0.54)
        apic[24].pt3dadd(-27.54, 225.62, 3.08, 0.54)
        apic[24].pt3dadd(-27.05, 227.11, 3.08, 0.54)
        apic[24].pt3dadd(-27.78, 229.60, 3.08, 0.54)
        apic[24].pt3dadd(-27.78, 230.10, 3.08, 0.54)
        apic[24].pt3dadd(-27.78, 231.34, 3.08, 0.54)
        apic[24].pt3dadd(-27.78, 231.84, 3.08, 0.54)
        apic[24].pt3dadd(-28.26, 233.58, 3.08, 0.54)
        apic[24].pt3dadd(-28.50, 234.33, 3.08, 0.54)
        apic[24].pt3dadd(-28.74, 235.82, 3.08, 0.54)
        apic[24].pt3dadd(-28.02, 237.81, 3.08, 0.54)
        apic[24].pt3dadd(-29.47, 239.05, 5.52, 0.54)
        apic[24].pt3dadd(-30.92, 240.55, 5.51, 0.54)
        apic[24].pt3dadd(-31.16, 241.79, 5.51, 0.54)
        apic[24].pt3dadd(-31.16, 244.03, 5.51, 0.54)
        apic[24].pt3dadd(-31.16, 245.03, 5.51, 0.54)
        apic[24].pt3dadd(-30.92, 246.52, 5.51, 0.54)
        apic[24].pt3dadd(-33.09, 247.51, 5.51, 0.54)
        apic[24].pt3dadd(-33.57, 247.76, 5.51, 0.54)
        apic[24].pt3dadd(-34.06, 249.25, 5.51, 0.54)
        apic[24].pt3dadd(-34.06, 250.00, 5.51, 0.54)
        apic[24].pt3dadd(-34.06, 251.24, 5.51, 0.54)
        apic[24].pt3dadd(-34.06, 252.49, 5.51, 0.54)
        apic[24].pt3dadd(-33.57, 253.23, 5.51, 0.54)
        apic[24].pt3dadd(-33.33, 254.48, 5.51, 0.54)
        
        apic[25] = h.Section(name='apic[25]')
        apic[25].connect(        apic[23](1), 0)
        apic[25].pt3dclear()
        apic[25].pt3dadd(0.72, 164.68, -3.16, 0.48)
        apic[25].pt3dadd(0.24, 167.41, -3.16, 0.48)
        apic[25].pt3dadd(1.21, 170.15, -3.16, 0.48)
        apic[25].pt3dadd(-0.00, 171.14, -3.16, 0.48)
        apic[25].pt3dadd(-0.24, 171.39, -3.16, 0.48)
        apic[25].pt3dadd(-0.48, 173.63, -8.34, 0.48)
        apic[25].pt3dadd(-0.72, 176.37, -8.34, 0.48)
        apic[25].pt3dadd(-0.97, 176.62, -8.34, 0.48)
        apic[25].pt3dadd(-1.45, 178.86, -8.34, 0.48)
        apic[25].pt3dadd(-1.45, 179.10, -8.34, 0.48)
        apic[25].pt3dadd(-1.45, 180.60, -8.34, 0.48)
        apic[25].pt3dadd(-0.48, 182.84, -8.34, 0.48)
        apic[25].pt3dadd(-0.24, 183.33, -8.34, 0.48)
        apic[25].pt3dadd(0.97, 184.33, -8.34, 0.48)
        apic[25].pt3dadd(1.21, 184.58, -8.33, 0.48)
        apic[25].pt3dadd(1.45, 186.07, -8.33, 0.48)
        apic[25].pt3dadd(1.45, 186.82, -8.33, 0.48)
        apic[25].pt3dadd(1.45, 186.82, -8.33, 0.48)
        
        apic[26] = h.Section(name='apic[26]')
        apic[26].connect(        apic[25](1), 0)
        apic[26].pt3dclear()
        apic[26].pt3dadd(1.45, 186.82, -8.33, 0.43)
        apic[26].pt3dadd(-0.97, 188.06, -8.02, 0.43)
        apic[26].pt3dadd(-2.42, 192.04, -8.02, 0.43)
        apic[26].pt3dadd(-2.42, 192.79, -8.02, 0.43)
        apic[26].pt3dadd(-3.62, 195.03, -8.02, 0.43)
        apic[26].pt3dadd(-4.59, 198.51, -8.02, 0.43)
        apic[26].pt3dadd(-5.31, 203.48, -8.02, 0.43)
        apic[26].pt3dadd(-5.31, 203.98, -8.02, 0.43)
        apic[26].pt3dadd(-5.56, 207.46, -8.02, 0.43)
        apic[26].pt3dadd(-5.56, 207.71, -8.02, 0.43)
        apic[26].pt3dadd(-7.00, 210.45, -8.02, 0.43)
        apic[26].pt3dadd(-6.04, 214.43, -10.08, 0.43)
        apic[26].pt3dadd(-6.04, 214.43, -10.11, 0.43)
        apic[26].pt3dadd(-4.59, 216.17, -10.12, 0.43)
        apic[26].pt3dadd(-4.11, 216.92, -10.12, 0.43)
        apic[26].pt3dadd(-2.42, 219.65, -10.12, 0.43)
        apic[26].pt3dadd(-2.42, 220.90, -10.12, 0.43)
        apic[26].pt3dadd(-2.42, 223.88, -10.13, 0.43)
        apic[26].pt3dadd(-2.66, 223.88, -10.13, 0.43)
        apic[26].pt3dadd(-6.52, 227.11, -10.13, 0.43)
        apic[26].pt3dadd(-7.25, 227.86, -10.13, 0.43)
        apic[26].pt3dadd(-10.63, 229.10, -10.13, 0.43)
        apic[26].pt3dadd(-11.59, 229.10, -10.13, 0.43)
        apic[26].pt3dadd(-15.94, 229.10, -10.13, 0.43)
        apic[26].pt3dadd(-16.91, 229.10, -10.13, 0.43)
        apic[26].pt3dadd(-17.39, 229.35, -10.13, 0.43)
        apic[26].pt3dadd(-19.32, 228.61, -10.13, 0.43)
        apic[26].pt3dadd(-19.81, 226.87, -10.13, 0.43)
        apic[26].pt3dadd(-22.46, 228.11, -10.13, 0.43)
        apic[26].pt3dadd(-23.19, 228.36, -10.13, 0.43)
        apic[26].pt3dadd(-24.88, 226.37, -10.13, 0.43)
        apic[26].pt3dadd(-25.12, 226.37, -10.13, 0.43)
        apic[26].pt3dadd(-27.54, 226.37, -10.13, 0.43)
        apic[26].pt3dadd(-30.92, 227.86, -8.49, 0.43)
        apic[26].pt3dadd(-32.85, 228.86, -8.49, 0.43)
        apic[26].pt3dadd(-33.57, 229.35, -8.49, 0.43)
        apic[26].pt3dadd(-34.30, 230.10, -8.49, 0.43)
        apic[26].pt3dadd(-35.02, 230.60, -8.49, 0.43)
        apic[26].pt3dadd(-36.96, 231.59, -8.49, 0.43)
        apic[26].pt3dadd(-37.92, 231.59, -8.49, 0.43)
        apic[26].pt3dadd(-38.65, 231.59, -8.49, 0.43)
        apic[26].pt3dadd(-38.89, 231.59, -8.49, 0.43)
        
        apic[27] = h.Section(name='apic[27]')
        apic[27].connect(        apic[25](1), 0)
        apic[27].pt3dclear()
        apic[27].pt3dadd(1.45, 186.82, -8.33, 0.43)
        apic[27].pt3dadd(2.42, 189.30, -9.77, 0.43)
        apic[27].pt3dadd(0.48, 191.54, -9.77, 0.43)
        apic[27].pt3dadd(0.48, 192.29, -9.77, 0.43)
        apic[27].pt3dadd(0.48, 194.53, -9.77, 0.43)
        apic[27].pt3dadd(3.14, 195.52, -9.77, 0.43)
        apic[27].pt3dadd(4.11, 195.52, -9.77, 0.43)
        apic[27].pt3dadd(5.31, 195.52, -9.77, 0.43)
        apic[27].pt3dadd(6.04, 197.02, -9.77, 0.43)
        apic[27].pt3dadd(6.04, 197.76, -9.77, 0.43)
        apic[27].pt3dadd(6.04, 199.75, -9.77, 0.43)
        apic[27].pt3dadd(6.04, 200.25, -9.77, 0.43)
        apic[27].pt3dadd(6.52, 201.74, -9.77, 0.43)
        apic[27].pt3dadd(7.25, 201.99, -9.77, 0.43)
        apic[27].pt3dadd(7.25, 204.73, -9.77, 0.43)
        apic[27].pt3dadd(7.49, 204.98, -9.77, 0.43)
        apic[27].pt3dadd(8.21, 205.97, -5.17, 0.43)
        apic[27].pt3dadd(7.25, 208.46, -5.17, 0.43)
        apic[27].pt3dadd(7.25, 208.71, -5.17, 0.43)
        apic[27].pt3dadd(9.42, 208.96, -5.17, 0.43)
        apic[27].pt3dadd(9.90, 209.20, -5.17, 0.43)
        apic[27].pt3dadd(10.39, 211.69, -5.17, 0.43)
        apic[27].pt3dadd(10.87, 211.94, -5.17, 0.43)
        apic[27].pt3dadd(13.53, 215.17, -3.09, 0.43)
        apic[27].pt3dadd(16.18, 217.16, -3.09, 0.43)
        apic[27].pt3dadd(16.91, 217.16, -3.09, 0.43)
        apic[27].pt3dadd(19.57, 215.42, -3.09, 0.43)
        apic[27].pt3dadd(20.05, 215.17, -3.09, 0.43)
        apic[27].pt3dadd(22.46, 214.68, -3.09, 0.43)
        apic[27].pt3dadd(23.43, 214.68, -3.09, 0.43)
        apic[27].pt3dadd(24.64, 216.17, -3.09, 0.43)
        apic[27].pt3dadd(24.64, 216.67, -3.09, 0.43)
        apic[27].pt3dadd(24.64, 219.40, -3.09, 0.43)
        apic[27].pt3dadd(27.05, 220.90, -3.10, 0.43)
        
        apic[28] = h.Section(name='apic[28]')
        apic[28].connect(        apic[20](1), 0)
        apic[28].pt3dclear()
        apic[28].pt3dadd(20.53, 146.27, 6.83, 2.70)
        apic[28].pt3dadd(19.63, 148.37, 6.82, 2.70)
        apic[28].pt3dadd(18.90, 150.36, 6.82, 2.70)
        apic[28].pt3dadd(18.90, 153.09, 1.32, 2.70)
        apic[28].pt3dadd(18.90, 156.83, 1.32, 2.70)
        apic[28].pt3dadd(20.35, 161.30, 1.32, 2.70)
        apic[28].pt3dadd(22.04, 164.54, 1.32, 2.70)
        apic[28].pt3dadd(22.77, 165.03, 1.32, 2.70)
        apic[28].pt3dadd(24.22, 167.77, 1.32, 2.70)
        apic[28].pt3dadd(24.70, 167.77, 1.32, 2.70)
        apic[28].pt3dadd(25.91, 169.76, 7.56, 2.70)
        
        apic[29] = h.Section(name='apic[29]')
        apic[29].connect(        apic[28](1), 0)
        apic[29].pt3dclear()
        apic[29].pt3dadd(25.91, 169.76, 7.56, 0.72)
        apic[29].pt3dadd(24.46, 171.25, 7.55, 0.72)
        apic[29].pt3dadd(23.97, 172.99, 7.55, 0.72)
        apic[29].pt3dadd(23.97, 174.74, -5.05, 0.72)
        apic[29].pt3dadd(23.97, 174.99, -5.05, 0.72)
        apic[29].pt3dadd(23.97, 176.73, -5.05, 0.72)
        apic[29].pt3dadd(23.97, 176.98, -5.05, 0.72)
        apic[29].pt3dadd(23.97, 178.97, -5.07, 0.72)
        apic[29].pt3dadd(24.22, 179.21, -5.07, 0.72)
        apic[29].pt3dadd(25.18, 179.71, -5.08, 0.72)
        apic[29].pt3dadd(25.91, 182.70, -11.48, 0.54)
        apic[29].pt3dadd(27.84, 184.94, -17.43, 0.54)
        apic[29].pt3dadd(27.84, 184.94, -17.42, 0.54)
        apic[29].pt3dadd(27.84, 186.93, -17.42, 0.54)
        apic[29].pt3dadd(27.84, 187.42, -17.42, 0.54)
        apic[29].pt3dadd(27.60, 188.42, -17.46, 0.54)
        apic[29].pt3dadd(27.60, 188.42, -17.53, 0.54)
        apic[29].pt3dadd(27.84, 190.41, -26.20, 0.54)
        apic[29].pt3dadd(28.56, 192.15, -26.39, 0.54)
        apic[29].pt3dadd(28.56, 192.40, -26.56, 0.54)
        apic[29].pt3dadd(29.77, 193.64, -26.58, 0.54)
        apic[29].pt3dadd(30.01, 193.89, -26.58, 0.54)
        apic[29].pt3dadd(30.50, 194.39, -26.58, 0.54)
        apic[29].pt3dadd(30.74, 194.39, -26.58, 0.54)
        apic[29].pt3dadd(31.70, 196.38, -29.96, 0.54)
        
        apic[30] = h.Section(name='apic[30]')
        apic[30].connect(        apic[28](1), 0)
        apic[30].pt3dclear()
        apic[30].pt3dadd(25.91, 169.76, 7.56, 3.14)
        apic[30].pt3dadd(27.60, 172.25, 4.20, 3.14)
        apic[30].pt3dadd(27.60, 172.25, 4.21, 3.14)
        apic[30].pt3dadd(29.77, 174.49, 4.30, 3.50)
        
        apic[31] = h.Section(name='apic[31]')
        apic[31].connect(        apic[30](1), 0)
        apic[31].pt3dclear()
        apic[31].pt3dadd(29.77, 174.49, 4.30, 0.48)
        apic[31].pt3dadd(30.01, 170.51, 12.24, 0.48)
        apic[31].pt3dadd(29.77, 169.01, 12.24, 0.48)
        apic[31].pt3dadd(29.77, 168.77, 12.24, 0.48)
        apic[31].pt3dadd(29.77, 167.77, 12.24, 0.48)
        apic[31].pt3dadd(30.25, 166.28, 12.24, 0.48)
        apic[31].pt3dadd(30.74, 166.03, 12.24, 0.48)
        apic[31].pt3dadd(31.22, 165.03, 12.24, 0.48)
        apic[31].pt3dadd(31.22, 164.54, 12.25, 0.48)
        apic[31].pt3dadd(30.98, 163.79, 12.25, 0.48)
        apic[31].pt3dadd(30.74, 163.54, 12.25, 0.48)
        apic[31].pt3dadd(29.77, 163.04, 12.25, 0.48)
        apic[31].pt3dadd(28.81, 162.80, 12.25, 0.48)
        apic[31].pt3dadd(28.32, 162.80, 12.25, 0.48)
        apic[31].pt3dadd(27.60, 162.55, 12.25, 0.48)
        apic[31].pt3dadd(27.36, 162.30, 12.25, 0.48)
        apic[31].pt3dadd(26.39, 161.80, 12.25, 0.48)
        apic[31].pt3dadd(25.91, 161.30, 12.25, 0.48)
        apic[31].pt3dadd(25.42, 160.56, 12.25, 0.48)
        apic[31].pt3dadd(24.94, 159.31, 12.25, 0.48)
        apic[31].pt3dadd(24.70, 158.57, 12.25, 0.48)
        apic[31].pt3dadd(24.46, 156.83, 12.25, 0.48)
        apic[31].pt3dadd(23.97, 154.84, 12.25, 0.48)
        apic[31].pt3dadd(23.73, 154.34, 12.25, 0.48)
        apic[31].pt3dadd(23.49, 154.09, 12.25, 0.48)
        
        apic[32] = h.Section(name='apic[32]')
        apic[32].connect(        apic[30](1), 0)
        apic[32].pt3dclear()
        apic[32].pt3dadd(29.77, 174.49, 4.30, 3.50)
        apic[32].pt3dadd(31.70, 177.47, 4.15, 3.50)
        apic[32].pt3dadd(32.43, 178.97, 4.15, 3.50)
        
        apic[33] = h.Section(name='apic[33]')
        apic[33].connect(        apic[32](1), 0)
        apic[33].pt3dclear()
        apic[33].pt3dadd(32.43, 178.97, 4.15, 1.10)
        apic[33].pt3dadd(34.36, 172.00, 3.38, 1.10)
        apic[33].pt3dadd(36.29, 172.00, 3.38, 1.10)
        apic[33].pt3dadd(38.95, 172.50, 3.38, 1.10)
        apic[33].pt3dadd(39.19, 172.75, 3.38, 1.10)
        apic[33].pt3dadd(41.12, 173.74, 3.38, 1.10)
        apic[33].pt3dadd(41.37, 173.74, 3.38, 1.10)
        apic[33].pt3dadd(43.06, 175.23, 2.00, 1.10)
        apic[33].pt3dadd(43.30, 175.48, 2.00, 1.10)
        apic[33].pt3dadd(44.51, 176.98, 2.00, 1.10)
        apic[33].pt3dadd(45.23, 177.22, 2.00, 1.10)
        apic[33].pt3dadd(46.20, 177.97, 2.00, 1.10)
        apic[33].pt3dadd(46.44, 177.97, 2.00, 1.10)
        
        apic[34] = h.Section(name='apic[34]')
        apic[34].connect(        apic[33](1), 0)
        apic[34].pt3dclear()
        apic[34].pt3dadd(46.44, 177.97, 2.00, 0.54)
        apic[34].pt3dadd(49.10, 177.47, 2.00, 0.54)
        apic[34].pt3dadd(50.79, 177.47, 2.00, 0.54)
        apic[34].pt3dadd(51.27, 177.47, 2.00, 0.54)
        apic[34].pt3dadd(53.44, 178.47, 2.00, 0.54)
        apic[34].pt3dadd(53.93, 178.72, 2.00, 0.54)
        apic[34].pt3dadd(56.10, 180.71, 2.00, 0.54)
        apic[34].pt3dadd(56.10, 180.96, 2.00, 0.54)
        apic[34].pt3dadd(57.55, 181.20, 2.00, 0.54)
        apic[34].pt3dadd(58.27, 181.45, 2.00, 0.54)
        apic[34].pt3dadd(60.45, 178.97, -1.20, 0.54)
        apic[34].pt3dadd(62.62, 179.46, -1.20, 0.54)
        apic[34].pt3dadd(62.86, 179.46, -1.20, 0.54)
        apic[34].pt3dadd(63.35, 178.47, -1.20, 0.54)
        apic[34].pt3dadd(64.31, 178.47, -1.20, 0.54)
        apic[34].pt3dadd(65.04, 177.47, -1.20, 0.54)
        apic[34].pt3dadd(65.52, 176.73, -1.20, 0.54)
        apic[34].pt3dadd(66.97, 175.73, -1.20, 0.54)
        apic[34].pt3dadd(67.21, 174.99, -1.20, 0.54)
        apic[34].pt3dadd(67.69, 173.24, -1.20, 0.54)
        apic[34].pt3dadd(67.94, 172.99, -1.20, 0.54)
        apic[34].pt3dadd(69.14, 172.75, -1.19, 0.54)
        apic[34].pt3dadd(70.11, 172.50, -1.19, 0.54)
        apic[34].pt3dadd(70.83, 171.75, -1.19, 0.54)
        apic[34].pt3dadd(71.08, 170.51, -1.19, 0.54)
        apic[34].pt3dadd(71.32, 169.76, -1.19, 0.54)
        apic[34].pt3dadd(71.80, 169.76, -1.19, 0.54)
        apic[34].pt3dadd(73.01, 169.01, -1.19, 0.54)
        apic[34].pt3dadd(73.97, 168.52, -1.19, 0.54)
        apic[34].pt3dadd(75.18, 168.52, -1.19, 0.54)
        apic[34].pt3dadd(75.42, 168.52, -1.19, 0.54)
        apic[34].pt3dadd(76.39, 168.77, -1.19, 0.54)
        apic[34].pt3dadd(79.05, 168.77, -1.19, 0.54)
        apic[34].pt3dadd(79.77, 168.77, -1.19, 0.54)
        apic[34].pt3dadd(81.70, 167.27, -1.19, 0.54)
        apic[34].pt3dadd(82.43, 166.78, -1.19, 0.54)
        apic[34].pt3dadd(83.15, 164.54, -1.19, 0.54)
        apic[34].pt3dadd(82.91, 164.29, -1.19, 0.54)
        apic[34].pt3dadd(82.91, 164.04, -1.19, 0.54)
        apic[34].pt3dadd(82.67, 162.05, -1.19, 0.54)
        apic[34].pt3dadd(82.67, 161.55, -1.19, 0.54)
        apic[34].pt3dadd(83.40, 159.31, 4.79, 0.54)
        apic[34].pt3dadd(83.40, 159.31, 4.80, 0.54)
        apic[34].pt3dadd(86.54, 157.57, -1.80, 0.54)
        apic[34].pt3dadd(89.19, 155.58, -1.80, 0.54)
        apic[34].pt3dadd(89.43, 155.58, -1.80, 0.54)
        apic[34].pt3dadd(92.57, 155.08, -1.79, 0.54)
        apic[34].pt3dadd(92.82, 155.08, -1.79, 0.54)
        apic[34].pt3dadd(94.02, 154.84, -1.79, 0.54)
        apic[34].pt3dadd(94.02, 154.59, -1.79, 0.54)
        apic[34].pt3dadd(94.26, 153.34, -1.79, 0.54)
        apic[34].pt3dadd(94.26, 152.85, -1.79, 0.54)
        apic[34].pt3dadd(94.99, 149.61, -1.79, 0.54)
        apic[34].pt3dadd(94.99, 149.11, -1.78, 0.54)
        apic[34].pt3dadd(95.71, 146.63, -1.78, 0.54)
        apic[34].pt3dadd(95.96, 145.63, -1.78, 0.54)
        apic[34].pt3dadd(97.40, 144.64, -1.78, 0.54)
        apic[34].pt3dadd(97.89, 144.14, -1.78, 0.54)
        apic[34].pt3dadd(98.61, 143.14, -1.78, 0.54)
        apic[34].pt3dadd(98.61, 142.90, -1.77, 0.54)
        apic[34].pt3dadd(100.30, 137.67, -1.77, 0.54)
        apic[34].pt3dadd(101.03, 137.67, -1.77, 0.54)
        apic[34].pt3dadd(102.24, 135.93, -1.77, 0.54)
        apic[34].pt3dadd(102.96, 135.43, -1.77, 0.54)
        apic[34].pt3dadd(104.65, 134.44, -1.77, 0.54)
        apic[34].pt3dadd(104.89, 133.69, -1.77, 0.54)
        apic[34].pt3dadd(106.34, 130.71, -1.77, 0.54)
        apic[34].pt3dadd(107.55, 129.46, -1.77, 0.54)
        apic[34].pt3dadd(109.72, 128.22, -1.77, 0.54)
        apic[34].pt3dadd(110.93, 127.97, -1.77, 0.54)
        apic[34].pt3dadd(112.14, 126.73, -1.77, 0.54)
        apic[34].pt3dadd(112.86, 126.23, -1.77, 0.54)
        apic[34].pt3dadd(114.55, 123.74, -4.39, 0.54)
        apic[34].pt3dadd(116.49, 120.76, -4.37, 0.54)
        apic[34].pt3dadd(116.97, 119.76, -4.37, 0.54)
        apic[34].pt3dadd(117.94, 117.27, -4.37, 0.43)
        apic[34].pt3dadd(117.94, 116.28, -4.37, 0.43)
        apic[34].pt3dadd(118.90, 114.29, -4.37, 0.43)
        apic[34].pt3dadd(120.59, 112.80, -4.37, 0.43)
        apic[34].pt3dadd(123.49, 112.30, -4.37, 0.43)
        apic[34].pt3dadd(124.70, 112.05, -4.37, 0.43)
        apic[34].pt3dadd(126.87, 111.30, -4.37, 0.43)
        apic[34].pt3dadd(127.60, 111.05, -4.37, 0.43)
        apic[34].pt3dadd(128.81, 110.56, -4.37, 0.43)
        apic[34].pt3dadd(129.53, 107.57, -4.35, 0.43)
        apic[34].pt3dadd(130.01, 107.32, -4.35, 0.43)
        apic[34].pt3dadd(131.70, 105.33, -4.35, 0.43)
        apic[34].pt3dadd(132.43, 104.84, -4.35, 0.43)
        apic[34].pt3dadd(133.15, 103.59, -4.35, 0.43)
        apic[34].pt3dadd(133.40, 103.34, -4.35, 0.43)
        apic[34].pt3dadd(134.36, 102.35, -4.35, 0.43)
        apic[34].pt3dadd(135.81, 102.10, -4.35, 0.43)
        apic[34].pt3dadd(137.26, 101.60, -4.35, 0.43)
        apic[34].pt3dadd(138.95, 101.60, -4.35, 0.43)
        apic[34].pt3dadd(139.19, 101.60, -4.35, 0.43)
        apic[34].pt3dadd(140.16, 100.11, -4.35, 0.43)
        apic[34].pt3dadd(140.64, 100.11, -4.35, 0.43)
        apic[34].pt3dadd(141.61, 102.35, -4.35, 0.43)
        
        apic[35] = h.Section(name='apic[35]')
        apic[35].connect(        apic[33](1), 0)
        apic[35].pt3dclear()
        apic[35].pt3dadd(46.44, 177.97, 2.00, 0.54)
        apic[35].pt3dadd(47.65, 179.96, 7.31, 0.54)
        apic[35].pt3dadd(48.37, 182.70, 7.31, 0.54)
        apic[35].pt3dadd(49.82, 184.19, 7.31, 0.54)
        apic[35].pt3dadd(50.06, 184.44, 7.31, 0.54)
        apic[35].pt3dadd(51.03, 185.93, 7.31, 0.54)
        apic[35].pt3dadd(51.51, 188.42, 7.31, 0.54)
        apic[35].pt3dadd(51.75, 188.42, 7.31, 0.54)
        apic[35].pt3dadd(52.48, 189.16, 7.31, 0.54)
        apic[35].pt3dadd(52.72, 192.90, 7.31, 0.54)
        apic[35].pt3dadd(53.20, 192.90, 7.31, 0.54)
        apic[35].pt3dadd(53.44, 192.90, 7.31, 0.54)
        apic[35].pt3dadd(56.34, 196.13, 2.77, 0.54)
        apic[35].pt3dadd(56.34, 196.13, 2.76, 0.54)
        apic[35].pt3dadd(57.31, 197.87, 2.76, 0.54)
        apic[35].pt3dadd(57.55, 198.37, 2.76, 0.54)
        apic[35].pt3dadd(58.27, 200.36, 2.76, 0.54)
        apic[35].pt3dadd(58.52, 201.10, 2.76, 0.54)
        apic[35].pt3dadd(58.52, 202.85, 2.76, 0.54)
        apic[35].pt3dadd(58.52, 203.84, 2.76, 0.54)
        apic[35].pt3dadd(58.52, 204.34, 2.76, 0.54)
        apic[35].pt3dadd(58.27, 205.08, 2.76, 0.54)
        apic[35].pt3dadd(58.27, 205.33, 2.76, 0.54)
        apic[35].pt3dadd(58.76, 207.57, 5.79, 0.54)
        apic[35].pt3dadd(59.48, 211.55, 5.79, 0.54)
        apic[35].pt3dadd(59.48, 212.05, 5.79, 0.54)
        apic[35].pt3dadd(61.66, 212.80, 5.79, 0.54)
        apic[35].pt3dadd(63.11, 214.04, 5.79, 0.54)
        apic[35].pt3dadd(63.11, 214.29, 5.79, 0.54)
        apic[35].pt3dadd(63.59, 215.28, 5.79, 0.54)
        apic[35].pt3dadd(64.31, 219.26, 3.27, 0.54)
        apic[35].pt3dadd(66.73, 221.75, 3.27, 0.54)
        apic[35].pt3dadd(66.97, 221.75, 3.27, 0.54)
        apic[35].pt3dadd(66.97, 224.74, 3.27, 0.54)
        apic[35].pt3dadd(67.94, 226.23, 3.27, 0.54)
        apic[35].pt3dadd(69.14, 226.73, 3.27, 0.54)
        apic[35].pt3dadd(70.59, 227.47, 3.27, 0.54)
        apic[35].pt3dadd(71.32, 227.97, 3.27, 0.54)
        apic[35].pt3dadd(72.28, 229.46, 3.27, 0.54)
        apic[35].pt3dadd(73.25, 230.46, 3.27, 0.54)
        apic[35].pt3dadd(73.25, 231.95, 3.27, 0.54)
        apic[35].pt3dadd(73.25, 232.20, 3.27, 0.54)
        apic[35].pt3dadd(74.22, 234.69, 3.27, 0.54)
        apic[35].pt3dadd(74.22, 236.18, 3.27, 0.43)
        apic[35].pt3dadd(74.22, 237.92, 3.27, 0.43)
        apic[35].pt3dadd(73.25, 240.91, 6.65, 0.43)
        apic[35].pt3dadd(73.73, 243.64, 6.65, 0.43)
        apic[35].pt3dadd(73.97, 244.14, 6.65, 0.43)
        apic[35].pt3dadd(75.18, 246.13, 6.65, 0.43)
        apic[35].pt3dadd(75.91, 246.38, 6.65, 0.43)
        apic[35].pt3dadd(76.87, 247.37, 6.65, 0.43)
        apic[35].pt3dadd(77.60, 249.61, 6.65, 0.43)
        apic[35].pt3dadd(78.08, 250.11, 6.65, 0.43)
        apic[35].pt3dadd(79.05, 251.35, 6.65, 0.43)
        apic[35].pt3dadd(79.29, 251.85, 6.65, 0.43)
        apic[35].pt3dadd(80.01, 253.09, 6.65, 0.43)
        apic[35].pt3dadd(80.74, 254.34, 6.65, 0.43)
        apic[35].pt3dadd(80.74, 254.84, 6.65, 0.43)
        apic[35].pt3dadd(80.98, 256.83, 6.65, 0.43)
        apic[35].pt3dadd(80.98, 257.82, 6.65, 0.43)
        apic[35].pt3dadd(82.43, 260.31, 2.88, 0.43)
        apic[35].pt3dadd(83.64, 261.05, 2.89, 0.43)
        apic[35].pt3dadd(83.88, 261.30, 2.89, 0.43)
        apic[35].pt3dadd(85.09, 263.04, 2.89, 0.43)
        apic[35].pt3dadd(85.09, 266.78, 8.91, 0.43)
        apic[35].pt3dadd(85.09, 266.78, 8.90, 0.43)
        apic[35].pt3dadd(85.57, 268.02, 8.90, 0.43)
        apic[35].pt3dadd(86.05, 268.27, 8.90, 0.43)
        apic[35].pt3dadd(87.50, 269.01, 8.90, 0.43)
        apic[35].pt3dadd(87.74, 269.26, 8.90, 0.43)
        apic[35].pt3dadd(88.71, 271.01, 8.90, 0.43)
        apic[35].pt3dadd(90.40, 273.24, 11.89, 0.43)
        apic[35].pt3dadd(91.37, 275.23, 11.89, 0.43)
        apic[35].pt3dadd(91.12, 275.98, 11.89, 0.43)
        apic[35].pt3dadd(90.40, 277.72, 11.89, 0.43)
        apic[35].pt3dadd(90.16, 277.97, 11.89, 0.43)
        apic[35].pt3dadd(90.16, 280.96, 11.89, 0.43)
        apic[35].pt3dadd(90.16, 281.20, 11.89, 0.43)
        apic[35].pt3dadd(90.40, 282.20, 11.89, 0.43)
        apic[35].pt3dadd(90.64, 282.45, 11.89, 0.43)
        apic[35].pt3dadd(92.57, 283.94, 11.89, 0.43)
        apic[35].pt3dadd(93.06, 284.44, 11.89, 0.43)
        apic[35].pt3dadd(93.78, 285.43, 11.89, 0.43)
        apic[35].pt3dadd(94.02, 285.68, 11.89, 0.43)
        apic[35].pt3dadd(94.75, 286.43, 11.89, 0.43)
        apic[35].pt3dadd(95.23, 287.17, 11.89, 0.43)
        apic[35].pt3dadd(95.47, 287.67, 11.89, 0.43)
        apic[35].pt3dadd(95.71, 287.92, 11.89, 0.43)
        apic[35].pt3dadd(96.20, 289.41, 11.89, 0.43)
        apic[35].pt3dadd(95.71, 293.39, 9.67, 0.43)
        apic[35].pt3dadd(95.71, 293.89, 9.67, 0.43)
        apic[35].pt3dadd(96.44, 296.63, 9.67, 0.43)
        apic[35].pt3dadd(96.92, 298.12, 9.67, 0.43)
        apic[35].pt3dadd(97.16, 303.34, 9.67, 0.43)
        apic[35].pt3dadd(98.13, 305.33, 9.67, 0.43)
        apic[35].pt3dadd(99.34, 306.08, 9.67, 0.43)
        apic[35].pt3dadd(100.30, 306.83, 9.67, 0.43)
        apic[35].pt3dadd(100.54, 307.32, 9.67, 0.43)
        apic[35].pt3dadd(101.03, 310.56, 8.08, 0.43)
        apic[35].pt3dadd(100.79, 313.79, 8.09, 0.43)
        apic[35].pt3dadd(100.79, 315.28, 8.10, 0.39)
        apic[35].pt3dadd(101.03, 318.02, 8.10, 0.39)
        apic[35].pt3dadd(101.27, 318.27, 8.10, 0.39)
        apic[35].pt3dadd(103.44, 319.01, 8.10, 0.39)
        apic[35].pt3dadd(103.68, 319.26, 8.10, 0.39)
        apic[35].pt3dadd(104.89, 321.75, 8.10, 0.39)
        apic[35].pt3dadd(105.13, 322.75, 8.10, 0.39)
        apic[35].pt3dadd(105.38, 323.24, 8.10, 0.39)
        apic[35].pt3dadd(105.13, 326.23, 0.98, 0.39)
        apic[35].pt3dadd(107.31, 327.47, 0.98, 0.39)
        apic[35].pt3dadd(107.31, 327.72, 0.98, 0.39)
        apic[35].pt3dadd(108.03, 329.96, 0.98, 0.39)
        apic[35].pt3dadd(109.48, 333.44, 0.98, 0.39)
        apic[35].pt3dadd(110.69, 337.67, 0.98, 0.39)
        apic[35].pt3dadd(110.93, 337.67, 0.98, 0.39)
        apic[35].pt3dadd(112.62, 338.92, 0.98, 0.39)
        apic[35].pt3dadd(112.62, 339.16, 0.98, 0.39)
        apic[35].pt3dadd(111.41, 343.14, -2.65, 0.39)
        apic[35].pt3dadd(113.11, 345.63, -2.65, 0.39)
        apic[35].pt3dadd(113.59, 345.63, -2.65, 0.39)
        apic[35].pt3dadd(115.04, 346.38, -2.65, 0.39)
        apic[35].pt3dadd(116.00, 346.63, -2.65, 0.39)
        apic[35].pt3dadd(117.69, 346.88, -2.65, 0.39)
        apic[35].pt3dadd(117.94, 346.88, -2.65, 0.39)
        apic[35].pt3dadd(119.14, 347.37, -2.65, 0.39)
        apic[35].pt3dadd(119.63, 347.62, -2.65, 0.39)
        apic[35].pt3dadd(120.35, 348.12, -2.65, 0.39)
        apic[35].pt3dadd(120.35, 348.37, -2.65, 0.39)
        apic[35].pt3dadd(121.32, 350.11, -2.65, 0.39)
        apic[35].pt3dadd(121.32, 350.61, -2.65, 0.39)
        apic[35].pt3dadd(121.32, 351.60, -2.65, 0.39)
        apic[35].pt3dadd(121.32, 353.34, -2.65, 0.39)
        apic[35].pt3dadd(121.32, 354.34, -2.65, 0.39)
        
        apic[36] = h.Section(name='apic[36]')
        apic[36].connect(        apic[32](1), 0)
        apic[36].pt3dclear()
        apic[36].pt3dadd(32.43, 178.97, 4.15, 3.50)
        apic[36].pt3dadd(33.88, 182.70, 0.80, 3.50)
        apic[36].pt3dadd(34.60, 186.18, 0.81, 3.50)
        apic[36].pt3dadd(35.33, 189.66, 0.82, 3.50)
        apic[36].pt3dadd(35.33, 189.66, 0.83, 3.50)
        apic[36].pt3dadd(35.57, 193.64, 0.86, 3.50)
        apic[36].pt3dadd(35.57, 197.87, 0.86, 3.50)
        
        apic[37] = h.Section(name='apic[37]')
        apic[37].connect(        apic[36](1), 0)
        apic[37].pt3dclear()
        apic[37].pt3dadd(35.57, 197.87, 0.86, 1.20)
        apic[37].pt3dadd(33.64, 200.61, 4.29, 1.20)
        apic[37].pt3dadd(32.67, 204.34, 4.29, 1.20)
        apic[37].pt3dadd(32.67, 204.59, 4.29, 1.20)
        apic[37].pt3dadd(33.15, 210.06, 4.29, 1.20)
        apic[37].pt3dadd(33.88, 211.80, 4.29, 1.20)
        apic[37].pt3dadd(32.43, 216.03, 4.29, 1.20)
        apic[37].pt3dadd(31.95, 219.26, 4.28, 1.20)
        apic[37].pt3dadd(32.67, 226.23, -6.34, 1.20)
        apic[37].pt3dadd(32.91, 228.97, -6.34, 1.20)
        apic[37].pt3dadd(32.91, 229.21, -6.34, 1.20)
        apic[37].pt3dadd(33.40, 232.95, -6.34, 1.20)
        apic[37].pt3dadd(33.40, 232.95, -6.33, 1.20)
        apic[37].pt3dadd(33.40, 234.44, -6.33, 1.20)
        apic[37].pt3dadd(32.67, 235.68, -6.33, 1.20)
        apic[37].pt3dadd(31.70, 236.93, -2.44, 0.96)
        apic[37].pt3dadd(31.70, 237.17, -2.44, 0.96)
        apic[37].pt3dadd(31.70, 238.92, -2.44, 0.96)
        apic[37].pt3dadd(31.95, 242.15, -2.44, 0.96)
        apic[37].pt3dadd(31.46, 244.39, -2.44, 0.96)
        apic[37].pt3dadd(31.22, 244.89, -2.44, 0.96)
        apic[37].pt3dadd(30.50, 246.13, -2.44, 0.96)
        apic[37].pt3dadd(30.01, 247.37, -2.44, 0.96)
        apic[37].pt3dadd(29.77, 247.62, -2.44, 0.96)
        apic[37].pt3dadd(29.77, 247.87, -2.44, 0.96)
        apic[37].pt3dadd(27.84, 251.10, -4.67, 0.96)
        apic[37].pt3dadd(27.36, 253.34, -4.67, 0.96)
        apic[37].pt3dadd(27.36, 254.09, -4.67, 0.96)
        apic[37].pt3dadd(27.60, 257.07, -4.67, 0.96)
        apic[37].pt3dadd(26.63, 260.31, 2.66, 0.77)
        apic[37].pt3dadd(26.39, 263.04, 2.66, 0.77)
        apic[37].pt3dadd(25.91, 266.28, 2.66, 0.77)
        apic[37].pt3dadd(26.63, 269.01, 2.69, 0.77)
        apic[37].pt3dadd(25.42, 273.74, 2.72, 0.77)
        apic[37].pt3dadd(25.18, 273.74, 2.72, 0.77)
        apic[37].pt3dadd(23.25, 278.72, 6.85, 0.77)
        apic[37].pt3dadd(23.25, 279.21, 6.85, 0.77)
        apic[37].pt3dadd(23.97, 281.70, 6.85, 0.77)
        apic[37].pt3dadd(23.01, 283.94, 6.86, 0.77)
        apic[37].pt3dadd(23.01, 284.19, 6.86, 0.77)
        apic[37].pt3dadd(23.25, 287.17, 6.86, 0.77)
        apic[37].pt3dadd(24.22, 289.16, 6.85, 0.77)
        apic[37].pt3dadd(24.46, 290.91, 8.04, 0.54)
        apic[37].pt3dadd(23.49, 292.65, 8.04, 0.54)
        apic[37].pt3dadd(23.01, 292.90, 8.04, 0.54)
        apic[37].pt3dadd(22.28, 294.64, 8.04, 0.54)
        apic[37].pt3dadd(21.56, 297.12, 8.04, 0.54)
        apic[37].pt3dadd(21.56, 298.62, 8.04, 0.54)
        apic[37].pt3dadd(21.80, 299.86, 8.04, 0.54)
        apic[37].pt3dadd(22.77, 301.85, 8.04, 0.54)
        apic[37].pt3dadd(21.80, 303.34, 8.04, 0.54)
        apic[37].pt3dadd(21.56, 303.59, 8.04, 0.54)
        apic[37].pt3dadd(20.59, 307.32, 8.04, 0.54)
        apic[37].pt3dadd(20.35, 307.57, 8.04, 0.54)
        apic[37].pt3dadd(19.87, 309.56, 8.04, 0.54)
        apic[37].pt3dadd(19.87, 310.06, 8.04, 0.54)
        apic[37].pt3dadd(19.63, 312.30, 8.04, 0.54)
        apic[37].pt3dadd(19.14, 313.04, 8.04, 0.54)
        apic[37].pt3dadd(17.69, 314.79, 8.04, 0.54)
        apic[37].pt3dadd(17.45, 315.78, 8.04, 0.54)
        apic[37].pt3dadd(17.21, 318.27, 8.04, 0.54)
        apic[37].pt3dadd(17.21, 318.77, 8.04, 0.54)
        apic[37].pt3dadd(16.49, 320.51, 8.04, 0.54)
        apic[37].pt3dadd(16.49, 322.50, 8.04, 0.54)
        apic[37].pt3dadd(16.73, 323.24, 8.04, 0.54)
        apic[37].pt3dadd(17.69, 325.23, 11.62, 0.54)
        apic[37].pt3dadd(16.25, 328.97, 11.62, 0.54)
        apic[37].pt3dadd(15.76, 329.96, 11.62, 0.54)
        apic[37].pt3dadd(15.52, 331.20, 11.62, 0.54)
        apic[37].pt3dadd(15.52, 331.95, 11.62, 0.54)
        apic[37].pt3dadd(15.52, 334.19, 11.62, 0.39)
        apic[37].pt3dadd(16.00, 334.69, 11.62, 0.39)
        apic[37].pt3dadd(16.73, 336.18, 11.62, 0.39)
        apic[37].pt3dadd(17.21, 336.68, 11.62, 0.39)
        apic[37].pt3dadd(17.69, 339.16, 14.71, 0.39)
        apic[37].pt3dadd(17.69, 342.40, 14.73, 0.39)
        apic[37].pt3dadd(17.45, 345.13, 14.73, 0.39)
        apic[37].pt3dadd(16.97, 345.63, 14.73, 0.39)
        apic[37].pt3dadd(15.52, 346.13, 14.72, 0.39)
        apic[37].pt3dadd(14.07, 347.62, 14.71, 0.39)
        apic[37].pt3dadd(12.62, 348.12, 14.71, 0.39)
        apic[37].pt3dadd(12.38, 348.12, 14.71, 0.39)
        apic[37].pt3dadd(10.93, 350.61, 14.71, 0.39)
        apic[37].pt3dadd(10.69, 351.10, 14.71, 0.39)
        apic[37].pt3dadd(8.76, 355.58, 14.55, 0.39)
        apic[37].pt3dadd(8.76, 357.07, 14.55, 0.39)
        apic[37].pt3dadd(8.76, 358.32, 14.55, 0.39)
        apic[37].pt3dadd(8.76, 358.57, 14.55, 0.39)
        apic[37].pt3dadd(8.76, 359.81, 14.55, 0.39)
        
        apic[38] = h.Section(name='apic[38]')
        apic[38].connect(        apic[36](1), 0)
        apic[38].pt3dclear()
        apic[38].pt3dadd(35.57, 197.87, 0.86, 3.2)
        apic[38].pt3dadd(36.05, 201.10, 1.32, 3.2)
        apic[38].pt3dadd(36.05, 202.85, 1.32, 3.2)
        apic[38].pt3dadd(36.29, 204.84, 1.32, 3.2)
        apic[38].pt3dadd(36.29, 206.33, 1.32, 3.2)
        
        apic[39] = h.Section(name='apic[39]')
        apic[39].connect(        apic[38](1), 0)
        apic[39].pt3dclear()
        apic[39].pt3dadd(36.29, 206.33, 1.32, 1.00)
        apic[39].pt3dadd(36.54, 209.56, -5.27, 1.00)
        apic[39].pt3dadd(36.54, 210.06, -5.27, 1.00)
        apic[39].pt3dadd(36.78, 212.05, -5.27, 1.00)
        apic[39].pt3dadd(37.02, 212.55, -5.27, 1.00)
        apic[39].pt3dadd(37.74, 214.04, -5.38, 1.00)
        apic[39].pt3dadd(37.98, 214.29, -5.40, 1.00)
        apic[39].pt3dadd(38.47, 216.53, -5.74, 1.00)
        apic[39].pt3dadd(38.47, 216.53, -5.80, 1.00)
        apic[39].pt3dadd(39.19, 218.27, -5.95, 1.00)
        apic[39].pt3dadd(39.19, 218.27, -5.97, 1.00)
        apic[39].pt3dadd(41.37, 222.25, -6.40, 1.00)
        apic[39].pt3dadd(41.61, 222.50, -6.40, 1.00)
        apic[39].pt3dadd(42.33, 224.74, -6.40, 1.00)
        apic[39].pt3dadd(42.33, 224.74, -6.37, 1.00)
        apic[39].pt3dadd(42.82, 226.73, -1.43, 1.00)
        apic[39].pt3dadd(42.82, 229.21, -1.43, 1.00)
        apic[39].pt3dadd(43.06, 231.45, -1.43, 0.96)
        apic[39].pt3dadd(43.06, 231.70, -1.43, 0.96)
        apic[39].pt3dadd(43.06, 232.70, -1.43, 0.96)
        apic[39].pt3dadd(43.06, 233.69, -1.43, 0.96)
        apic[39].pt3dadd(43.06, 233.94, -1.43, 0.96)
        apic[39].pt3dadd(42.33, 236.18, -1.43, 0.96)
        apic[39].pt3dadd(42.09, 237.42, -1.43, 0.96)
        apic[39].pt3dadd(39.68, 239.41, -1.43, 0.96)
        apic[39].pt3dadd(39.19, 239.91, -1.43, 0.96)
        apic[39].pt3dadd(38.95, 240.16, -1.43, 0.96)
        apic[39].pt3dadd(37.74, 240.91, 0.91, 0.96)
        apic[39].pt3dadd(37.50, 241.15, 0.91, 0.96)
        apic[39].pt3dadd(35.81, 242.65, 0.91, 0.96)
        apic[39].pt3dadd(35.33, 246.88, 0.91, 0.96)
        apic[39].pt3dadd(35.09, 246.88, 0.91, 0.96)
        apic[39].pt3dadd(33.88, 249.36, 0.91, 0.96)
        apic[39].pt3dadd(33.15, 250.36, 0.91, 0.96)
        apic[39].pt3dadd(32.67, 251.60, 0.91, 0.96)
        apic[39].pt3dadd(31.95, 252.85, 0.91, 0.77)
        apic[39].pt3dadd(31.70, 254.59, -1.56, 0.77)
        apic[39].pt3dadd(31.22, 256.58, -1.56, 0.77)
        apic[39].pt3dadd(30.98, 256.58, -1.56, 0.77)
        apic[39].pt3dadd(29.05, 258.57, -1.56, 0.77)
        apic[39].pt3dadd(28.32, 260.81, -1.56, 0.77)
        apic[39].pt3dadd(28.32, 261.30, -1.56, 0.77)
        apic[39].pt3dadd(27.84, 263.29, -1.56, 0.77)
        apic[39].pt3dadd(27.84, 265.78, -1.56, 0.77)
        apic[39].pt3dadd(29.29, 268.27, -1.56, 0.77)
        apic[39].pt3dadd(29.29, 271.75, -4.83, 0.77)
        apic[39].pt3dadd(28.81, 274.24, -4.83, 0.77)
        apic[39].pt3dadd(27.84, 276.23, -5.53, 0.77)
        apic[39].pt3dadd(27.84, 276.23, -5.52, 0.77)
        apic[39].pt3dadd(27.36, 278.72, -5.54, 0.77)
        apic[39].pt3dadd(27.11, 278.97, -5.54, 0.77)
        apic[39].pt3dadd(26.87, 281.20, -5.54, 0.77)
        apic[39].pt3dadd(26.87, 281.45, -5.54, 0.77)
        apic[39].pt3dadd(28.32, 285.18, -5.54, 0.77)
        apic[39].pt3dadd(28.32, 285.43, -5.54, 0.77)
        apic[39].pt3dadd(28.56, 288.17, -5.54, 0.54)
        apic[39].pt3dadd(28.08, 290.66, -5.53, 0.54)
        apic[39].pt3dadd(27.11, 293.64, -4.32, 0.54)
        apic[39].pt3dadd(27.11, 293.89, -4.32, 0.54)
        apic[39].pt3dadd(26.63, 296.13, -4.32, 0.54)
        apic[39].pt3dadd(26.63, 296.63, -4.32, 0.54)
        apic[39].pt3dadd(25.91, 298.62, -4.32, 0.54)
        apic[39].pt3dadd(25.67, 299.11, -4.32, 0.54)
        apic[39].pt3dadd(26.15, 302.60, -4.32, 0.54)
        apic[39].pt3dadd(26.63, 303.34, -4.32, 0.54)
        apic[39].pt3dadd(27.60, 305.08, -4.32, 0.54)
        apic[39].pt3dadd(27.84, 305.08, -4.32, 0.54)
        apic[39].pt3dadd(28.08, 306.83, -4.33, 0.54)
        apic[39].pt3dadd(27.84, 308.82, -1.02, 0.54)
        apic[39].pt3dadd(27.84, 311.55, -1.02, 0.54)
        apic[39].pt3dadd(27.84, 311.80, -1.02, 0.54)
        apic[39].pt3dadd(29.05, 313.54, -1.02, 0.54)
        apic[39].pt3dadd(29.29, 313.79, -1.02, 0.54)
        apic[39].pt3dadd(30.01, 315.78, -1.02, 0.54)
        apic[39].pt3dadd(30.25, 316.03, -1.02, 0.54)
        apic[39].pt3dadd(30.25, 318.02, -1.02, 0.54)
        apic[39].pt3dadd(30.25, 318.52, -1.02, 0.54)
        apic[39].pt3dadd(29.29, 320.51, -2.28, 0.39)
        apic[39].pt3dadd(29.05, 320.51, -2.28, 0.39)
        apic[39].pt3dadd(28.08, 323.49, -2.28, 0.39)
        apic[39].pt3dadd(28.08, 323.99, -2.28, 0.39)
        apic[39].pt3dadd(27.60, 326.48, -2.28, 0.39)
        apic[39].pt3dadd(27.11, 327.72, -2.28, 0.39)
        apic[39].pt3dadd(27.36, 329.71, -2.28, 0.39)
        apic[39].pt3dadd(27.60, 329.96, -2.28, 0.39)
        apic[39].pt3dadd(29.53, 333.19, -2.28, 0.39)
        apic[39].pt3dadd(29.77, 334.19, -2.28, 0.39)
        apic[39].pt3dadd(29.77, 334.94, -2.28, 0.39)
        apic[39].pt3dadd(28.56, 337.67, -2.28, 0.39)
        apic[39].pt3dadd(27.60, 338.17, -2.28, 0.39)
        apic[39].pt3dadd(27.36, 338.42, -2.28, 0.39)
        apic[39].pt3dadd(26.63, 338.42, -2.28, 0.39)
        apic[39].pt3dadd(25.91, 338.42, -2.28, 0.39)
        apic[39].pt3dadd(25.42, 338.42, -2.28, 0.39)
        apic[39].pt3dadd(24.94, 338.42, -6.77, 0.39)
        apic[39].pt3dadd(24.94, 338.42, -6.81, 0.39)
        apic[39].pt3dadd(23.97, 341.15, -6.81, 0.39)
        apic[39].pt3dadd(22.04, 345.63, -6.81, 0.39)
        apic[39].pt3dadd(22.04, 346.38, -6.81, 0.39)
        apic[39].pt3dadd(22.04, 348.87, -6.81, 0.39)
        apic[39].pt3dadd(22.04, 349.36, -6.81, 0.39)
        apic[39].pt3dadd(22.28, 352.60, -6.81, 0.39)
        apic[39].pt3dadd(22.53, 353.59, -6.81, 0.39)
        apic[39].pt3dadd(22.77, 355.58, -6.81, 0.39)
        apic[39].pt3dadd(22.77, 355.83, -6.81, 0.39)
        apic[39].pt3dadd(21.32, 357.07, -6.81, 0.39)
        apic[39].pt3dadd(20.35, 357.07, -6.81, 0.39)
        apic[39].pt3dadd(17.94, 357.07, -6.81, 0.39)
        apic[39].pt3dadd(17.45, 357.07, -6.81, 0.39)
        apic[39].pt3dadd(15.04, 357.07, -6.81, 0.39)
        apic[39].pt3dadd(13.11, 359.31, -6.81, 0.39)
        apic[39].pt3dadd(11.90, 361.30, -6.81, 0.39)
        apic[39].pt3dadd(10.45, 363.29, -6.81, 0.39)
        apic[39].pt3dadd(9.48, 365.03, -7.05, 0.39)
        
        apic[40] = h.Section(name='apic[40]')
        apic[40].connect(        apic[38](1), 0)
        apic[40].pt3dclear()
        apic[40].pt3dadd(36.29, 206.33, 1.32, 3.14)
        apic[40].pt3dadd(38.47, 210.31, 2.97, 3.14)
        apic[40].pt3dadd(41.12, 215.28, 2.97, 3.14)
        apic[40].pt3dadd(43.06, 217.52, 2.97, 3.14)
        apic[40].pt3dadd(43.78, 219.51, 2.97, 3.14)
        
        apic[41] = h.Section(name='apic[41]')
        apic[41].connect(        apic[40](1), 0)
        apic[41].pt3dclear()
        apic[41].pt3dadd(43.78, 219.51, 2.97, 1.20)
        apic[41].pt3dadd(39.19, 222.75, 2.97, 1.20)
        apic[41].pt3dadd(34.60, 224.49, 2.97, 1.20)
        apic[41].pt3dadd(31.22, 227.47, 8.92, 0.77)
        apic[41].pt3dadd(29.53, 230.71, 8.92, 0.77)
        apic[41].pt3dadd(28.56, 233.44, 8.92, 0.77)
        apic[41].pt3dadd(28.56, 235.43, 8.92, 0.77)
        apic[41].pt3dadd(28.56, 237.67, 8.92, 0.77)
        apic[41].pt3dadd(28.81, 240.41, 8.92, 0.77)
        apic[41].pt3dadd(27.11, 242.15, -1.26, 0.77)
        apic[41].pt3dadd(24.70, 244.89, -1.26, 0.77)
        apic[41].pt3dadd(24.70, 245.13, -1.26, 0.77)
        apic[41].pt3dadd(23.73, 247.37, -1.26, 0.77)
        apic[41].pt3dadd(23.97, 249.36, -1.26, 0.77)
        apic[41].pt3dadd(24.22, 249.61, -1.26, 0.77)
        apic[41].pt3dadd(25.18, 252.10, -1.26, 0.77)
        apic[41].pt3dadd(25.18, 254.59, -1.27, 0.77)
        apic[41].pt3dadd(25.18, 256.58, -1.28, 0.77)
        apic[41].pt3dadd(25.18, 257.07, -1.28, 0.77)
        apic[41].pt3dadd(24.70, 261.05, -1.29, 0.77)
        apic[41].pt3dadd(24.46, 261.55, -1.29, 0.77)
        apic[41].pt3dadd(23.49, 263.04, -1.29, 0.77)
        apic[41].pt3dadd(23.49, 263.54, -1.29, 0.77)
        apic[41].pt3dadd(23.49, 266.28, -1.29, 0.77)
        apic[41].pt3dadd(23.73, 267.52, -1.29, 0.77)
        apic[41].pt3dadd(23.97, 268.27, -1.29, 0.77)
        apic[41].pt3dadd(23.73, 270.01, -3.47, 0.77)
        apic[41].pt3dadd(23.49, 272.00, -3.47, 0.77)
        apic[41].pt3dadd(23.25, 272.00, -3.47, 0.77)
        apic[41].pt3dadd(22.04, 273.74, -3.47, 0.61)
        apic[41].pt3dadd(21.80, 273.74, -3.47, 0.61)
        apic[41].pt3dadd(20.59, 274.99, -3.47, 0.61)
        apic[41].pt3dadd(20.11, 274.99, -3.47, 0.61)
        apic[41].pt3dadd(18.18, 275.98, -3.47, 0.61)
        apic[41].pt3dadd(17.69, 276.48, -3.47, 0.61)
        apic[41].pt3dadd(17.21, 277.47, -3.47, 0.61)
        apic[41].pt3dadd(16.73, 277.97, -3.47, 0.61)
        apic[41].pt3dadd(16.49, 278.97, -3.47, 0.61)
        apic[41].pt3dadd(16.49, 280.21, -3.47, 0.61)
        apic[41].pt3dadd(16.49, 281.45, -3.47, 0.61)
        apic[41].pt3dadd(16.49, 282.95, -3.46, 0.61)
        apic[41].pt3dadd(16.25, 283.94, -3.46, 0.61)
        
        apic[42] = h.Section(name='apic[42]')
        apic[42].connect(        apic[41](1), 0)
        apic[42].pt3dclear()
        apic[42].pt3dadd(16.25, 283.94, -3.46, 0.54)
        apic[42].pt3dadd(16.25, 285.93, -4.42, 0.54)
        apic[42].pt3dadd(16.49, 290.16, -4.42, 0.54)
        apic[42].pt3dadd(16.73, 292.40, -4.42, 0.54)
        apic[42].pt3dadd(16.73, 293.14, -4.42, 0.54)
        apic[42].pt3dadd(15.04, 294.89, -4.42, 0.54)
        apic[42].pt3dadd(15.04, 295.13, -4.42, 0.54)
        apic[42].pt3dadd(13.35, 296.88, -4.42, 0.54)
        apic[42].pt3dadd(13.11, 297.12, -4.42, 0.54)
        apic[42].pt3dadd(11.66, 298.87, -4.42, 0.54)
        apic[42].pt3dadd(11.66, 302.10, -4.42, 0.54)
        apic[42].pt3dadd(12.14, 302.35, -4.42, 0.54)
        apic[42].pt3dadd(10.93, 304.34, -4.42, 0.54)
        apic[42].pt3dadd(9.97, 304.34, -4.42, 0.54)
        apic[42].pt3dadd(8.76, 305.33, -4.42, 0.54)
        apic[42].pt3dadd(7.79, 305.83, -4.42, 0.54)
        apic[42].pt3dadd(6.82, 307.32, -4.42, 0.54)
        apic[42].pt3dadd(6.82, 310.06, -4.42, 0.54)
        apic[42].pt3dadd(6.82, 311.30, -4.42, 0.54)
        apic[42].pt3dadd(7.07, 313.29, -4.42, 0.54)
        apic[42].pt3dadd(7.07, 314.04, -4.42, 0.54)
        apic[42].pt3dadd(6.34, 317.02, -4.42, 0.54)
        apic[42].pt3dadd(6.58, 318.77, -4.42, 0.54)
        apic[42].pt3dadd(7.07, 319.26, -4.42, 0.54)
        apic[42].pt3dadd(8.27, 321.01, -3.57, 0.54)
        apic[42].pt3dadd(8.52, 322.75, 2.81, 0.54)
        apic[42].pt3dadd(8.52, 322.75, 2.82, 0.54)
        apic[42].pt3dadd(7.07, 328.22, 2.82, 0.54)
        apic[42].pt3dadd(6.82, 328.72, 2.82, 0.54)
        apic[42].pt3dadd(5.86, 330.96, 2.83, 0.54)
        apic[42].pt3dadd(5.13, 331.95, 2.83, 0.54)
        apic[42].pt3dadd(3.93, 334.19, 2.84, 0.54)
        apic[42].pt3dadd(3.68, 334.44, 2.84, 0.43)
        apic[42].pt3dadd(3.44, 337.67, 2.84, 0.43)
        apic[42].pt3dadd(3.44, 339.16, 2.84, 0.43)
        apic[42].pt3dadd(2.96, 341.40, 2.85, 0.43)
        apic[42].pt3dadd(2.96, 342.90, 2.85, 0.43)
        apic[42].pt3dadd(2.48, 346.88, 2.85, 0.43)
        apic[42].pt3dadd(2.48, 346.88, 2.86, 0.43)
        apic[42].pt3dadd(1.51, 349.61, 2.87, 0.43)
        apic[42].pt3dadd(1.51, 349.86, 2.87, 0.43)
        apic[42].pt3dadd(1.51, 352.35, 2.87, 0.43)
        apic[42].pt3dadd(1.51, 353.09, 2.87, 0.43)
        apic[42].pt3dadd(4.65, 355.58, 2.88, 0.43)
        apic[42].pt3dadd(4.89, 355.58, 2.88, 0.43)
        apic[42].pt3dadd(5.13, 357.82, 2.88, 0.43)
        apic[42].pt3dadd(5.62, 358.82, 2.88, 0.43)
        apic[42].pt3dadd(6.34, 360.31, 2.88, 0.43)
        apic[42].pt3dadd(6.34, 360.81, 2.88, 0.43)
        apic[42].pt3dadd(6.10, 363.54, 2.88, 0.43)
        apic[42].pt3dadd(5.86, 364.29, 2.88, 0.43)
        apic[42].pt3dadd(5.38, 368.52, -3.20, 0.43)
        
        apic[43] = h.Section(name='apic[43]')
        apic[43].connect(        apic[41](1), 0)
        apic[43].pt3dclear()
        apic[43].pt3dadd(16.25, 283.94, -3.46, 0.54)
        apic[43].pt3dadd(14.07, 285.43, -3.51, 0.54)
        apic[43].pt3dadd(13.59, 289.41, -3.52, 0.54)
        apic[43].pt3dadd(13.59, 289.66, -3.52, 0.54)
        apic[43].pt3dadd(14.07, 291.90, -3.52, 0.54)
        apic[43].pt3dadd(14.31, 292.65, -3.52, 0.54)
        apic[43].pt3dadd(15.04, 294.39, -3.52, 0.54)
        apic[43].pt3dadd(15.04, 296.63, -3.52, 0.54)
        apic[43].pt3dadd(14.31, 297.87, -3.52, 0.54)
        apic[43].pt3dadd(13.35, 300.36, -3.52, 0.54)
        apic[43].pt3dadd(13.35, 300.61, -3.52, 0.54)
        apic[43].pt3dadd(13.11, 302.10, -3.52, 0.54)
        apic[43].pt3dadd(12.86, 303.09, -3.52, 0.54)
        apic[43].pt3dadd(11.41, 305.33, -3.52, 0.54)
        apic[43].pt3dadd(11.17, 305.83, -3.52, 0.54)
        apic[43].pt3dadd(10.69, 307.32, -3.52, 0.54)
        apic[43].pt3dadd(10.69, 307.57, -3.52, 0.54)
        apic[43].pt3dadd(10.69, 309.06, -3.52, 0.54)
        apic[43].pt3dadd(10.69, 309.56, -3.52, 0.54)
        apic[43].pt3dadd(10.69, 312.80, -3.52, 0.54)
        apic[43].pt3dadd(10.45, 315.53, -3.52, 0.54)
        apic[43].pt3dadd(10.21, 317.02, -3.52, 0.54)
        apic[43].pt3dadd(10.21, 317.52, -3.52, 0.54)
        apic[43].pt3dadd(9.72, 318.52, -3.52, 0.54)
        apic[43].pt3dadd(9.72, 318.52, -3.51, 0.54)
        apic[43].pt3dadd(9.24, 320.26, -3.51, 0.54)
        apic[43].pt3dadd(9.00, 320.26, -3.51, 0.54)
        apic[43].pt3dadd(6.82, 322.00, -3.51, 0.54)
        apic[43].pt3dadd(6.58, 322.00, -3.51, 0.54)
        apic[43].pt3dadd(6.10, 322.50, -3.51, 0.54)
        apic[43].pt3dadd(6.10, 323.99, -3.51, 0.54)
        apic[43].pt3dadd(6.10, 324.99, -3.51, 0.54)
        apic[43].pt3dadd(4.17, 327.22, 0.62, 0.54)
        apic[43].pt3dadd(3.93, 327.47, 0.62, 0.54)
        apic[43].pt3dadd(2.48, 330.96, 0.62, 0.54)
        apic[43].pt3dadd(2.48, 331.20, 0.62, 0.54)
        apic[43].pt3dadd(1.75, 333.69, 0.62, 0.43)
        apic[43].pt3dadd(1.75, 333.94, 0.62, 0.43)
        apic[43].pt3dadd(1.75, 336.68, 0.62, 0.43)
        apic[43].pt3dadd(1.75, 337.42, 0.62, 0.43)
        apic[43].pt3dadd(1.75, 339.41, 0.62, 0.43)
        apic[43].pt3dadd(1.75, 339.91, 0.62, 0.43)
        apic[43].pt3dadd(1.03, 341.65, 0.62, 0.43)
        apic[43].pt3dadd(0.54, 342.15, 0.62, 0.43)
        apic[43].pt3dadd(-0.66, 344.64, 0.62, 0.43)
        apic[43].pt3dadd(-1.63, 346.38, 0.62, 0.43)
        apic[43].pt3dadd(-4.04, 350.61, 0.62, 0.43)
        apic[43].pt3dadd(-4.29, 351.10, 0.62, 0.43)
        apic[43].pt3dadd(-5.01, 352.35, 0.62, 0.43)
        apic[43].pt3dadd(-5.01, 352.60, 0.62, 0.43)
        apic[43].pt3dadd(-6.22, 353.84, 0.62, 0.43)
        apic[43].pt3dadd(-7.91, 356.33, 1.81, 0.43)
        apic[43].pt3dadd(-7.91, 356.33, 1.82, 0.43)
        apic[43].pt3dadd(-8.88, 359.81, 1.82, 0.43)
        apic[43].pt3dadd(-8.88, 363.29, 1.82, 0.43)
        apic[43].pt3dadd(-7.67, 365.53, 1.82, 0.43)
        apic[43].pt3dadd(-7.18, 366.28, 1.82, 0.43)
        apic[43].pt3dadd(-7.18, 368.02, 1.82, 0.43)
        apic[43].pt3dadd(-7.18, 368.77, 1.82, 0.43)
        apic[43].pt3dadd(-7.18, 369.51, 1.82, 0.43)
        apic[43].pt3dadd(-7.43, 372.25, 1.84, 0.43)
        apic[43].pt3dadd(-9.12, 374.24, 1.85, 0.43)
        apic[43].pt3dadd(-9.84, 375.98, 1.85, 0.43)
        
        apic[44] = h.Section(name='apic[44]')
        apic[44].connect(        apic[40](1), 0)
        apic[44].pt3dclear()
        apic[44].pt3dadd(43.78, 219.51, 2.97, 3.14)
        apic[44].pt3dadd(46.44, 223.24, 5.23, 3.14)
        apic[44].pt3dadd(48.13, 226.73, 5.23, 3.14)
        apic[44].pt3dadd(48.37, 226.98, 5.23, 3.14)
        apic[44].pt3dadd(49.82, 229.46, 5.23, 3.14)
        apic[44].pt3dadd(49.82, 229.71, 5.23, 3.14)
        apic[44].pt3dadd(51.03, 231.20, 5.23, 3.14)
        apic[44].pt3dadd(51.99, 234.19, 5.23, 3.14)
        apic[44].pt3dadd(52.72, 236.93, 9.82, 3.14)
        apic[44].pt3dadd(52.72, 237.17, 9.82, 2.70)
        apic[44].pt3dadd(52.48, 240.91, 9.82, 2.70)
        apic[44].pt3dadd(51.99, 243.64, 9.71, 2.70)
        apic[44].pt3dadd(51.51, 245.88, 9.71, 2.70)
        apic[44].pt3dadd(51.51, 247.62, 9.72, 2.70)
        
        apic[45] = h.Section(name='apic[45]')
        apic[45].connect(        apic[44](1), 0)
        apic[45].pt3dclear()
        apic[45].pt3dadd(51.51, 247.62, 9.72, 0.61)
        apic[45].pt3dadd(49.82, 249.11, 6.19, 0.61)
        apic[45].pt3dadd(48.85, 250.36, 6.04, 0.61)
        apic[45].pt3dadd(48.61, 250.61, 5.99, 0.61)
        apic[45].pt3dadd(47.89, 251.60, 5.83, 0.61)
        apic[45].pt3dadd(47.89, 251.60, 5.80, 0.61)
        apic[45].pt3dadd(47.89, 253.34, 5.25, 0.61)
        apic[45].pt3dadd(47.89, 253.59, 5.17, 0.61)
        apic[45].pt3dadd(49.34, 254.34, 4.75, 0.61)
        apic[45].pt3dadd(49.34, 254.34, 4.62, 0.61)
        apic[45].pt3dadd(49.82, 254.59, -0.24, 0.54)
        apic[45].pt3dadd(49.82, 254.59, -0.42, 0.54)
        apic[45].pt3dadd(51.51, 254.84, -0.41, 0.54)
        apic[45].pt3dadd(52.96, 255.33, -0.41, 0.54)
        apic[45].pt3dadd(54.41, 255.33, -0.41, 0.54)
        apic[45].pt3dadd(54.89, 255.33, -0.41, 0.54)
        apic[45].pt3dadd(55.86, 255.58, -0.41, 0.54)
        apic[45].pt3dadd(56.58, 255.83, -0.41, 0.54)
        apic[45].pt3dadd(57.55, 256.33, -0.41, 0.54)
        apic[45].pt3dadd(58.03, 256.58, -0.41, 0.54)
        apic[45].pt3dadd(58.27, 256.83, -0.41, 0.54)
        apic[45].pt3dadd(58.76, 257.07, -0.41, 0.54)
        apic[45].pt3dadd(59.72, 258.07, -0.41, 0.54)
        apic[45].pt3dadd(59.72, 258.32, -0.41, 0.54)
        apic[45].pt3dadd(59.72, 258.57, -0.41, 0.54)
        
        apic[46] = h.Section(name='apic[46]')
        apic[46].connect(        apic[45](1), 0)
        apic[46].pt3dclear()
        apic[46].pt3dadd(59.72, 258.57, -0.41, 0.43)
        apic[46].pt3dadd(61.41, 260.06, -3.80, 0.43)
        apic[46].pt3dadd(61.41, 263.29, -3.80, 0.43)
        apic[46].pt3dadd(62.86, 264.54, -3.80, 0.43)
        apic[46].pt3dadd(63.59, 264.79, -3.80, 0.43)
        apic[46].pt3dadd(64.31, 265.03, -3.80, 0.43)
        apic[46].pt3dadd(65.04, 265.28, -3.79, 0.43)
        apic[46].pt3dadd(65.76, 265.78, -3.79, 0.43)
        apic[46].pt3dadd(66.49, 268.52, -3.79, 0.43)
        apic[46].pt3dadd(67.69, 269.01, -3.79, 0.43)
        apic[46].pt3dadd(67.94, 269.26, -3.79, 0.43)
        apic[46].pt3dadd(68.90, 270.51, -3.79, 0.43)
        apic[46].pt3dadd(69.39, 270.76, -3.79, 0.43)
        apic[46].pt3dadd(70.11, 271.00, -3.79, 0.43)
        apic[46].pt3dadd(70.35, 271.25, -3.79, 0.43)
        apic[46].pt3dadd(71.08, 271.50, -3.79, 0.43)
        apic[46].pt3dadd(71.80, 271.75, -3.79, 0.43)
        apic[46].pt3dadd(73.01, 271.75, -3.79, 0.43)
        apic[46].pt3dadd(73.25, 271.75, -3.79, 0.43)
        apic[46].pt3dadd(74.46, 271.50, -3.79, 0.43)
        apic[46].pt3dadd(75.42, 271.50, -3.79, 0.43)
        apic[46].pt3dadd(76.63, 271.75, -3.79, 0.43)
        apic[46].pt3dadd(77.60, 272.25, -3.79, 0.43)
        apic[46].pt3dadd(78.32, 273.49, -3.79, 0.43)
        apic[46].pt3dadd(78.81, 273.99, -3.79, 0.43)
        apic[46].pt3dadd(79.29, 274.74, -3.79, 0.43)
        apic[46].pt3dadd(80.25, 276.73, -3.79, 0.43)
        apic[46].pt3dadd(80.74, 276.73, -3.79, 0.43)
        apic[46].pt3dadd(82.19, 276.73, -3.79, 0.43)
        apic[46].pt3dadd(83.64, 276.98, -3.79, 0.43)
        apic[46].pt3dadd(84.84, 276.98, -3.79, 0.43)
        apic[46].pt3dadd(86.05, 277.22, -3.79, 0.43)
        apic[46].pt3dadd(87.02, 277.47, -3.79, 0.43)
        apic[46].pt3dadd(88.95, 278.72, -3.79, 0.43)
        apic[46].pt3dadd(94.02, 282.45, -9.01, 0.43)
        apic[46].pt3dadd(94.51, 284.44, -9.01, 0.43)
        apic[46].pt3dadd(94.75, 284.69, -9.01, 0.43)
        apic[46].pt3dadd(96.92, 286.93, -9.01, 0.43)
        apic[46].pt3dadd(97.16, 287.17, -9.01, 0.43)
        apic[46].pt3dadd(99.34, 288.42, -9.01, 0.43)
        apic[46].pt3dadd(100.54, 289.66, -9.01, 0.43)
        apic[46].pt3dadd(101.51, 290.41, -9.01, 0.43)
        apic[46].pt3dadd(102.48, 292.15, -9.01, 0.43)
        apic[46].pt3dadd(103.44, 293.39, -9.01, 0.43)
        apic[46].pt3dadd(104.41, 294.39, -9.01, 0.43)
        apic[46].pt3dadd(106.34, 295.88, -9.01, 0.43)
        apic[46].pt3dadd(107.55, 296.88, -9.01, 0.43)
        apic[46].pt3dadd(107.79, 297.37, -9.01, 0.43)
        apic[46].pt3dadd(108.27, 297.62, -9.01, 0.43)
        apic[46].pt3dadd(109.00, 299.11, -9.01, 0.43)
        apic[46].pt3dadd(109.24, 299.11, -9.01, 0.43)
        apic[46].pt3dadd(110.21, 302.85, -9.01, 0.43)
        apic[46].pt3dadd(110.45, 303.09, -9.01, 0.43)
        apic[46].pt3dadd(112.14, 305.33, -9.01, 0.43)
        apic[46].pt3dadd(112.14, 305.83, -9.01, 0.43)
        apic[46].pt3dadd(112.14, 309.81, -9.01, 0.43)
        apic[46].pt3dadd(112.14, 310.31, -9.01, 0.43)
        apic[46].pt3dadd(112.14, 312.05, -9.01, 0.43)
        apic[46].pt3dadd(112.38, 312.05, -9.01, 0.43)
        apic[46].pt3dadd(113.11, 316.03, -11.62, 0.43)
        apic[46].pt3dadd(114.31, 317.77, -11.62, 0.43)
        apic[46].pt3dadd(114.31, 319.76, -11.62, 0.43)
        apic[46].pt3dadd(114.31, 320.01, -11.62, 0.43)
        apic[46].pt3dadd(114.07, 323.00, -11.62, 0.43)
        apic[46].pt3dadd(114.80, 324.49, -11.62, 0.39)
        apic[46].pt3dadd(115.04, 324.74, -11.62, 0.39)
        apic[46].pt3dadd(116.25, 326.48, -11.62, 0.39)
        apic[46].pt3dadd(117.21, 329.71, -10.53, 0.39)
        apic[46].pt3dadd(117.21, 329.96, -10.53, 0.39)
        apic[46].pt3dadd(117.94, 331.45, -10.54, 0.39)
        apic[46].pt3dadd(117.69, 331.45, -10.54, 0.39)
        apic[46].pt3dadd(116.97, 333.94, -7.75, 0.39)
        apic[46].pt3dadd(117.94, 336.68, -7.75, 0.39)
        apic[46].pt3dadd(118.66, 336.93, -7.75, 0.39)
        apic[46].pt3dadd(120.11, 337.42, -7.75, 0.39)
        apic[46].pt3dadd(118.90, 340.41, -7.75, 0.39)
        apic[46].pt3dadd(118.66, 342.40, -7.75, 0.39)
        apic[46].pt3dadd(120.11, 343.39, -7.75, 0.39)
        apic[46].pt3dadd(120.83, 343.64, -7.75, 0.39)
        apic[46].pt3dadd(121.56, 344.89, -7.75, 0.39)
        apic[46].pt3dadd(122.53, 345.38, -7.75, 0.39)
        apic[46].pt3dadd(122.77, 345.63, -7.75, 0.39)
        apic[46].pt3dadd(123.01, 345.63, -7.75, 0.39)
        apic[46].pt3dadd(122.53, 348.12, -13.04, 0.39)
        apic[46].pt3dadd(123.01, 349.86, -13.04, 0.39)
        apic[46].pt3dadd(123.25, 349.86, -13.04, 0.39)
        apic[46].pt3dadd(122.53, 352.10, -13.04, 0.39)
        apic[46].pt3dadd(122.53, 352.85, -13.04, 0.39)
        
        apic[47] = h.Section(name='apic[47]')
        apic[47].connect(        apic[45](1), 0)
        apic[47].pt3dclear()
        apic[47].pt3dadd(59.72, 258.57, -0.41, 0.54)
        apic[47].pt3dadd(58.76, 260.06, 0.48, 0.54)
        apic[47].pt3dadd(58.27, 261.55, 0.48, 0.54)
        apic[47].pt3dadd(58.27, 261.80, 0.48, 0.54)
        apic[47].pt3dadd(58.27, 263.29, 0.48, 0.54)
        apic[47].pt3dadd(58.52, 263.54, 0.48, 0.54)
        apic[47].pt3dadd(59.48, 264.79, 0.48, 0.54)
        apic[47].pt3dadd(59.97, 265.78, 0.48, 0.54)
        apic[47].pt3dadd(60.21, 266.03, 0.48, 0.54)
        apic[47].pt3dadd(60.45, 267.02, 0.48, 0.54)
        apic[47].pt3dadd(60.69, 267.77, 0.48, 0.54)
        apic[47].pt3dadd(60.93, 267.77, 0.48, 0.54)
        apic[47].pt3dadd(61.41, 269.01, 0.48, 0.54)
        apic[47].pt3dadd(61.41, 269.26, 0.48, 0.54)
        
        apic[48] = h.Section(name='apic[48]')
        apic[48].connect(        apic[47](1), 0)
        apic[48].pt3dclear()
        apic[48].pt3dadd(61.41, 269.26, 0.48, 0.48)
        apic[48].pt3dadd(59.00, 272.00, -4.90, 0.48)
        apic[48].pt3dadd(59.00, 274.74, -4.90, 0.48)
        apic[48].pt3dadd(59.00, 274.99, -4.90, 0.48)
        apic[48].pt3dadd(59.00, 276.98, -4.90, 0.48)
        apic[48].pt3dadd(59.00, 277.97, -4.90, 0.48)
        apic[48].pt3dadd(58.27, 278.97, -4.90, 0.48)
        apic[48].pt3dadd(56.82, 279.21, -4.90, 0.48)
        apic[48].pt3dadd(56.10, 279.46, -4.90, 0.48)
        apic[48].pt3dadd(54.89, 280.46, -4.90, 0.48)
        apic[48].pt3dadd(54.65, 280.96, -4.90, 0.48)
        apic[48].pt3dadd(55.38, 281.70, -4.90, 0.48)
        apic[48].pt3dadd(56.34, 281.95, -4.90, 0.48)
        apic[48].pt3dadd(57.55, 282.45, -4.90, 0.48)
        apic[48].pt3dadd(58.27, 282.95, -4.90, 0.48)
        apic[48].pt3dadd(58.76, 283.19, -4.90, 0.48)
        apic[48].pt3dadd(59.72, 284.44, -4.90, 0.48)
        apic[48].pt3dadd(59.97, 285.18, -4.90, 0.48)
        apic[48].pt3dadd(59.72, 286.93, -4.90, 0.48)
        apic[48].pt3dadd(59.24, 287.42, -4.89, 0.48)
        apic[48].pt3dadd(57.55, 288.92, -4.89, 0.48)
        apic[48].pt3dadd(57.07, 289.41, -4.89, 0.48)
        apic[48].pt3dadd(56.58, 290.16, -4.89, 0.48)
        apic[48].pt3dadd(56.10, 291.40, -4.89, 0.48)
        apic[48].pt3dadd(56.10, 292.15, -4.89, 0.48)
        apic[48].pt3dadd(57.07, 293.89, -4.89, 0.48)
        apic[48].pt3dadd(57.31, 294.14, -4.89, 0.48)
        apic[48].pt3dadd(57.31, 294.39, -4.89, 0.48)
        apic[48].pt3dadd(58.03, 296.63, -9.45, 0.48)
        apic[48].pt3dadd(57.79, 296.88, -9.45, 0.48)
        apic[48].pt3dadd(57.07, 298.12, -9.45, 0.48)
        apic[48].pt3dadd(57.55, 299.36, -9.45, 0.48)
        apic[48].pt3dadd(58.27, 299.86, -9.45, 0.48)
        apic[48].pt3dadd(58.52, 300.11, -9.45, 0.48)
        apic[48].pt3dadd(59.24, 301.35, -9.45, 0.48)
        apic[48].pt3dadd(59.48, 301.60, -9.45, 0.48)
        apic[48].pt3dadd(59.72, 302.10, -9.45, 0.48)
        apic[48].pt3dadd(59.72, 303.34, -9.45, 0.48)
        
        apic[49] = h.Section(name='apic[49]')
        apic[49].connect(        apic[48](1), 0)
        apic[49].pt3dclear()
        apic[49].pt3dadd(59.72, 303.34, -9.45, 0.39)
        apic[49].pt3dadd(58.27, 307.32, -9.45, 0.39)
        apic[49].pt3dadd(58.52, 310.31, -15.49, 0.39)
        apic[49].pt3dadd(58.52, 310.56, -15.49, 0.39)
        apic[49].pt3dadd(59.48, 316.78, -15.49, 0.39)
        apic[49].pt3dadd(59.72, 316.78, -15.49, 0.39)
        apic[49].pt3dadd(61.90, 314.54, -15.48, 0.39)
        
        apic[50] = h.Section(name='apic[50]')
        apic[50].connect(        apic[48](1), 0)
        apic[50].pt3dclear()
        apic[50].pt3dadd(59.72, 303.34, -9.45, 0.39)
        apic[50].pt3dadd(61.41, 307.82, -9.36, 0.39)
        apic[50].pt3dadd(61.41, 307.82, -9.35, 0.39)
        apic[50].pt3dadd(61.90, 310.56, -9.35, 0.39)
        apic[50].pt3dadd(61.90, 310.56, -9.34, 0.39)
        apic[50].pt3dadd(62.86, 311.55, -9.33, 0.39)
        apic[50].pt3dadd(63.83, 312.05, -9.33, 0.39)
        apic[50].pt3dadd(64.80, 313.54, -9.33, 0.39)
        apic[50].pt3dadd(65.28, 314.04, -9.33, 0.39)
        apic[50].pt3dadd(64.07, 316.78, -9.33, 0.39)
        apic[50].pt3dadd(64.07, 317.02, -9.33, 0.39)
        apic[50].pt3dadd(61.17, 319.26, -9.33, 0.39)
        apic[50].pt3dadd(60.45, 319.26, -9.33, 0.39)
        apic[50].pt3dadd(60.21, 321.25, -9.33, 0.39)
        apic[50].pt3dadd(59.97, 322.50, -16.57, 0.39)
        apic[50].pt3dadd(60.69, 324.49, -16.57, 0.39)
        apic[50].pt3dadd(59.97, 325.73, -19.71, 0.39)
        apic[50].pt3dadd(56.82, 326.98, -19.71, 0.39)
        apic[50].pt3dadd(56.58, 326.98, -19.71, 0.39)
        apic[50].pt3dadd(56.10, 328.47, -21.95, 0.39)
        apic[50].pt3dadd(56.10, 328.47, -21.97, 0.39)
        apic[50].pt3dadd(57.31, 329.71, -22.00, 0.39)
        apic[50].pt3dadd(58.76, 330.96, -23.28, 0.39)
        apic[50].pt3dadd(59.00, 333.19, -23.26, 0.39)
        apic[50].pt3dadd(59.00, 333.19, -23.28, 0.39)
        apic[50].pt3dadd(57.79, 335.18, -23.28, 0.39)
        apic[50].pt3dadd(57.31, 335.18, -23.28, 0.39)
        apic[50].pt3dadd(55.13, 336.43, -23.29, 0.39)
        apic[50].pt3dadd(54.89, 336.43, -23.29, 0.39)
        apic[50].pt3dadd(54.89, 338.42, -26.90, 0.39)
        apic[50].pt3dadd(54.89, 338.42, -26.92, 0.39)
        apic[50].pt3dadd(55.86, 339.91, -27.07, 0.39)
        apic[50].pt3dadd(58.76, 341.65, -27.09, 0.39)
        apic[50].pt3dadd(57.07, 343.14, -27.09, 0.39)
        apic[50].pt3dadd(55.62, 343.64, -27.09, 0.39)
        apic[50].pt3dadd(55.13, 343.89, -27.31, 0.39)
        apic[50].pt3dadd(54.89, 344.14, -27.40, 0.39)
        apic[50].pt3dadd(53.20, 346.38, -28.06, 0.39)
        apic[50].pt3dadd(53.20, 346.38, -28.11, 0.39)
        apic[50].pt3dadd(52.72, 348.87, -28.16, 0.39)
        apic[50].pt3dadd(52.72, 350.86, -28.16, 0.39)
        apic[50].pt3dadd(52.72, 351.10, -28.16, 0.39)
        apic[50].pt3dadd(53.44, 353.34, -28.16, 0.39)
        apic[50].pt3dadd(55.38, 354.84, -28.84, 0.39)
        apic[50].pt3dadd(55.38, 354.84, -28.89, 0.39)
        apic[50].pt3dadd(55.38, 358.32, -29.68, 0.39)
        apic[50].pt3dadd(55.38, 358.32, -29.73, 0.39)
        apic[50].pt3dadd(55.86, 361.55, -30.22, 0.39)
        apic[50].pt3dadd(55.86, 361.80, -30.30, 0.39)
        apic[50].pt3dadd(55.86, 364.04, -30.94, 0.39)
        apic[50].pt3dadd(54.89, 364.29, -30.97, 0.39)
        apic[50].pt3dadd(53.44, 365.03, -30.98, 0.39)
        apic[50].pt3dadd(52.72, 365.78, -31.04, 0.39)
        apic[50].pt3dadd(51.99, 368.52, -31.39, 0.39)
        apic[50].pt3dadd(51.99, 368.52, -31.42, 0.39)
        apic[50].pt3dadd(51.99, 369.26, -31.43, 0.39)
        apic[50].pt3dadd(51.99, 369.26, -31.44, 0.39)
        
        apic[51] = h.Section(name='apic[51]')
        apic[51].connect(        apic[47](1), 0)
        apic[51].pt3dclear()
        apic[51].pt3dadd(61.41, 269.26, 0.48, 0.39)
        apic[51].pt3dadd(62.38, 271.00, -3.33, 0.39)
        apic[51].pt3dadd(62.38, 271.00, -3.34, 0.39)
        apic[51].pt3dadd(63.59, 272.99, -3.35, 0.39)
        apic[51].pt3dadd(63.35, 273.99, -3.36, 0.39)
        apic[51].pt3dadd(62.62, 275.23, -3.36, 0.39)
        apic[51].pt3dadd(62.38, 276.48, -3.37, 0.39)
        apic[51].pt3dadd(62.38, 277.97, -3.38, 0.39)
        apic[51].pt3dadd(62.62, 278.72, -3.39, 0.39)
        apic[51].pt3dadd(63.59, 279.21, -3.40, 0.39)
        apic[51].pt3dadd(64.07, 279.71, -3.40, 0.39)
        apic[51].pt3dadd(65.04, 281.45, -3.40, 0.39)
        apic[51].pt3dadd(66.00, 281.95, -3.40, 0.39)
        apic[51].pt3dadd(67.21, 282.70, -3.40, 0.39)
        apic[51].pt3dadd(67.69, 284.94, -3.40, 0.39)
        apic[51].pt3dadd(67.69, 286.43, -3.40, 0.39)
        apic[51].pt3dadd(67.69, 287.92, -3.40, 0.39)
        apic[51].pt3dadd(67.69, 288.17, -3.40, 0.39)
        apic[51].pt3dadd(67.21, 290.66, -3.40, 0.39)
        apic[51].pt3dadd(66.49, 290.91, -3.40, 0.39)
        apic[51].pt3dadd(65.52, 291.90, -3.40, 0.39)
        apic[51].pt3dadd(65.28, 292.15, -3.40, 0.39)
        apic[51].pt3dadd(65.28, 293.64, -3.41, 0.39)
        apic[51].pt3dadd(65.28, 294.14, -3.41, 0.39)
        apic[51].pt3dadd(65.28, 296.88, -3.41, 0.39)
        apic[51].pt3dadd(65.52, 297.12, -3.41, 0.39)
        apic[51].pt3dadd(65.76, 298.62, -3.41, 0.39)
        apic[51].pt3dadd(65.76, 298.87, -3.41, 0.39)
        apic[51].pt3dadd(65.04, 301.60, -7.85, 0.39)
        apic[51].pt3dadd(65.04, 301.60, -7.96, 0.39)
        apic[51].pt3dadd(66.73, 303.34, -8.05, 0.39)
        apic[51].pt3dadd(66.97, 303.59, -8.08, 0.39)
        apic[51].pt3dadd(67.94, 305.33, -8.13, 0.39)
        apic[51].pt3dadd(68.18, 305.83, -8.13, 0.39)
        apic[51].pt3dadd(68.18, 307.57, -8.14, 0.39)
        apic[51].pt3dadd(67.94, 307.57, -8.14, 0.39)
        apic[51].pt3dadd(66.25, 307.57, -13.07, 0.39)
        apic[51].pt3dadd(65.28, 308.82, -13.07, 0.39)
        
        apic[52] = h.Section(name='apic[52]')
        apic[52].connect(        apic[44](1), 0)
        apic[52].pt3dclear()
        apic[52].pt3dadd(51.51, 247.62, 9.72, 2.70)
        apic[52].pt3dadd(52.06, 250.61, 12.55, 2.70)
        apic[52].pt3dadd(52.54, 254.10, 12.55, 2.70)
        apic[52].pt3dadd(52.30, 254.59, 12.55, 2.70)
        apic[52].pt3dadd(52.79, 257.83, 12.55, 2.70)
        apic[52].pt3dadd(53.99, 260.56, 12.55, 2.70)
        apic[52].pt3dadd(55.44, 264.05, 12.55, 2.70)
        apic[52].pt3dadd(55.44, 264.30, 12.55, 2.70)
        apic[52].pt3dadd(56.89, 267.03, 12.55, 2.70)
        apic[52].pt3dadd(58.83, 268.28, 12.55, 2.70)
        apic[52].pt3dadd(60.76, 270.76, 12.55, 2.70)
        apic[52].pt3dadd(63.41, 273.00, 12.55, 2.70)
        apic[52].pt3dadd(63.66, 273.00, 12.55, 2.80)
        apic[52].pt3dadd(65.83, 274.49, 12.55, 2.80)
        apic[52].pt3dadd(66.07, 274.49, 12.55, 2.80)
        apic[52].pt3dadd(67.76, 275.49, 12.55, 2.80)
        apic[52].pt3dadd(70.18, 277.23, 13.10, 3.10)
        apic[52].pt3dadd(71.14, 280.46, 13.10, 3.10)
        apic[52].pt3dadd(72.11, 282.95, 13.10, 3.10)
        apic[52].pt3dadd(72.83, 286.19, 13.10, 3.10)
        
        apic[53] = h.Section(name='apic[53]')
        apic[53].connect(        apic[52](1), 0)
        apic[53].pt3dclear()
        apic[53].pt3dadd(72.83, 286.19, 13.10, 0.86)
        apic[53].pt3dadd(75.49, 288.42, 3.86, 0.86)
        apic[53].pt3dadd(75.97, 289.92, 3.86, 0.86)
        apic[53].pt3dadd(75.97, 290.17, 3.86, 0.86)
        apic[53].pt3dadd(77.18, 291.41, 3.86, 0.86)
        apic[53].pt3dadd(78.15, 291.66, 3.86, 0.86)
        apic[53].pt3dadd(79.60, 292.40, 3.86, 0.86)
        apic[53].pt3dadd(80.08, 292.65, 3.86, 0.86)
        apic[53].pt3dadd(81.29, 293.40, 3.86, 0.86)
        apic[53].pt3dadd(81.53, 294.15, 3.86, 0.86)
        apic[53].pt3dadd(81.53, 295.14, 3.86, 0.86)
        apic[53].pt3dadd(81.77, 297.88, 3.86, 0.86)
        apic[53].pt3dadd(82.98, 299.12, 3.86, 0.86)
        apic[53].pt3dadd(83.22, 299.37, 3.86, 0.86)
        apic[53].pt3dadd(84.91, 300.12, 3.86, 0.86)
        apic[53].pt3dadd(85.88, 301.36, 3.86, 0.86)
        apic[53].pt3dadd(85.88, 301.86, 3.86, 0.72)
        apic[53].pt3dadd(87.09, 302.85, 3.86, 0.72)
        apic[53].pt3dadd(88.54, 305.59, 9.12, 0.72)
        apic[53].pt3dadd(88.54, 305.84, 9.18, 0.72)
        apic[53].pt3dadd(88.54, 308.33, 9.21, 0.72)
        apic[53].pt3dadd(88.54, 308.57, 9.22, 0.72)
        apic[53].pt3dadd(90.71, 309.82, 9.24, 0.72)
        apic[53].pt3dadd(91.19, 310.07, 9.24, 0.72)
        apic[53].pt3dadd(94.09, 311.56, 6.08, 0.72)
        apic[53].pt3dadd(94.09, 311.81, 6.08, 0.72)
        apic[53].pt3dadd(95.78, 312.55, 6.08, 0.72)
        apic[53].pt3dadd(96.99, 313.30, 6.07, 0.72)
        apic[53].pt3dadd(97.71, 313.80, 6.07, 0.72)
        apic[53].pt3dadd(98.20, 314.05, 6.07, 0.72)
        apic[53].pt3dadd(99.16, 314.30, 6.07, 0.72)
        apic[53].pt3dadd(101.58, 314.05, 6.07, 0.72)
        apic[53].pt3dadd(101.82, 314.05, 6.07, 0.72)
        apic[53].pt3dadd(103.03, 314.54, 6.07, 0.72)
        apic[53].pt3dadd(103.75, 314.79, 6.07, 0.72)
        apic[53].pt3dadd(104.48, 315.29, 6.07, 0.72)
        apic[53].pt3dadd(105.44, 315.79, 6.07, 0.72)
        apic[53].pt3dadd(106.17, 316.04, 6.07, 0.72)
        apic[53].pt3dadd(106.65, 316.53, 6.07, 0.72)
        apic[53].pt3dadd(106.89, 316.53, 6.07, 0.72)
        apic[53].pt3dadd(108.34, 317.28, 6.07, 0.72)
        apic[53].pt3dadd(108.58, 317.53, 6.07, 0.72)
        apic[53].pt3dadd(110.03, 318.52, 6.07, 0.72)
        apic[53].pt3dadd(110.27, 318.77, 6.07, 0.72)
        apic[53].pt3dadd(111.48, 320.02, 6.07, 0.72)
        apic[53].pt3dadd(112.69, 320.51, 2.09, 0.72)
        apic[53].pt3dadd(112.93, 320.51, 2.09, 0.72)
        apic[53].pt3dadd(115.11, 319.77, 2.09, 0.72)
        apic[53].pt3dadd(115.59, 319.77, 2.09, 0.72)
        apic[53].pt3dadd(118.00, 320.76, 2.09, 0.72)
        apic[53].pt3dadd(118.49, 322.50, 2.09, 0.72)
        apic[53].pt3dadd(119.21, 323.00, 2.09, 0.72)
        apic[53].pt3dadd(120.66, 323.50, 2.09, 0.72)
        apic[53].pt3dadd(121.39, 323.75, 2.09, 0.72)
        apic[53].pt3dadd(122.83, 324.25, 2.09, 0.72)
        apic[53].pt3dadd(124.04, 324.74, 2.09, 0.72)
        apic[53].pt3dadd(125.49, 325.24, 2.09, 0.55)
        apic[53].pt3dadd(127.18, 325.99, 2.09, 0.55)
        apic[53].pt3dadd(127.67, 326.24, 2.09, 0.55)
        apic[53].pt3dadd(128.63, 326.48, 2.09, 0.55)
        apic[53].pt3dadd(129.12, 327.48, 0.14, 0.55)
        apic[53].pt3dadd(129.12, 328.23, 0.14, 0.55)
        apic[53].pt3dadd(129.36, 328.97, 0.14, 0.55)
        apic[53].pt3dadd(130.08, 328.97, 0.14, 0.55)
        apic[53].pt3dadd(132.01, 329.47, 0.14, 0.55)
        apic[53].pt3dadd(133.70, 329.47, 0.14, 0.55)
        apic[53].pt3dadd(133.95, 329.47, 0.14, 0.55)
        apic[53].pt3dadd(135.15, 329.47, 0.14, 0.55)
        apic[53].pt3dadd(135.15, 329.72, 0.14, 0.55)
        apic[53].pt3dadd(135.88, 330.96, 0.14, 0.55)
        apic[53].pt3dadd(135.88, 331.96, 0.14, 0.55)
        apic[53].pt3dadd(136.36, 332.45, 0.14, 0.55)
        apic[53].pt3dadd(137.09, 333.45, 0.14, 0.55)
        apic[53].pt3dadd(139.26, 333.95, -0.79, 0.55)
        apic[53].pt3dadd(141.68, 334.20, -0.79, 0.55)
        apic[53].pt3dadd(141.92, 334.20, -0.79, 0.55)
        apic[53].pt3dadd(143.61, 334.69, -0.79, 0.55)
        apic[53].pt3dadd(144.09, 334.94, -0.79, 0.55)
        apic[53].pt3dadd(145.06, 335.69, -0.79, 0.55)
        apic[53].pt3dadd(146.02, 336.68, -0.79, 0.55)
        apic[53].pt3dadd(146.02, 336.93, -0.79, 0.55)
        apic[53].pt3dadd(148.44, 338.67, 0.58, 0.55)
        apic[53].pt3dadd(150.61, 338.67, 0.57, 0.55)
        apic[53].pt3dadd(150.61, 338.92, 0.57, 0.55)
        apic[53].pt3dadd(153.51, 340.17, 0.56, 0.55)
        apic[53].pt3dadd(154.24, 341.41, 0.55, 0.55)
        apic[53].pt3dadd(154.48, 341.41, 0.55, 0.55)
        apic[53].pt3dadd(156.65, 339.92, 0.55, 0.55)
        apic[53].pt3dadd(156.89, 339.92, 0.55, 0.55)
        apic[53].pt3dadd(158.58, 339.67, -2.67, 0.55)
        apic[53].pt3dadd(158.58, 339.67, -2.68, 0.55)
        apic[53].pt3dadd(160.03, 341.16, -2.72, 0.55)
        apic[53].pt3dadd(161.24, 342.40, -3.49, 0.55)
        apic[53].pt3dadd(161.24, 342.40, -3.58, 0.55)
        apic[53].pt3dadd(161.72, 342.90, -5.62, 0.55)
        apic[53].pt3dadd(163.66, 343.65, -5.61, 0.55)
        apic[53].pt3dadd(163.90, 343.65, -5.61, 0.55)
        apic[53].pt3dadd(166.07, 344.15, -5.61, 0.55)
        apic[53].pt3dadd(166.31, 344.39, -5.61, 0.55)
        apic[53].pt3dadd(168.25, 345.14, -5.74, 0.55)
        apic[53].pt3dadd(170.90, 345.39, -5.74, 0.48)
        apic[53].pt3dadd(172.35, 346.88, -5.74, 0.48)
        apic[53].pt3dadd(173.08, 347.13, -5.74, 0.48)
        apic[53].pt3dadd(174.28, 347.13, -5.74, 0.48)
        apic[53].pt3dadd(175.01, 346.63, -5.74, 0.48)
        apic[53].pt3dadd(176.46, 346.38, -5.74, 0.48)
        apic[53].pt3dadd(177.18, 346.38, -5.74, 0.48)
        apic[53].pt3dadd(177.91, 346.38, -5.74, 0.48)
        apic[53].pt3dadd(178.87, 346.38, -5.74, 0.48)
        apic[53].pt3dadd(179.12, 346.88, -5.74, 0.48)
        apic[53].pt3dadd(179.84, 348.13, -5.74, 0.48)
        apic[53].pt3dadd(181.05, 348.87, -5.81, 0.48)
        apic[53].pt3dadd(181.53, 348.87, -5.81, 0.48)
        apic[53].pt3dadd(181.77, 349.12, -5.82, 0.48)
        apic[53].pt3dadd(183.22, 348.62, -9.17, 0.48)
        apic[53].pt3dadd(183.70, 348.62, -9.16, 0.48)
        apic[53].pt3dadd(185.15, 349.37, -9.16, 0.48)
        apic[53].pt3dadd(186.12, 349.87, -9.16, 0.48)
        apic[53].pt3dadd(186.84, 350.12, -9.16, 0.48)
        apic[53].pt3dadd(188.29, 351.36, -9.15, 0.48)
        apic[53].pt3dadd(188.78, 351.86, -9.15, 0.48)
        apic[53].pt3dadd(190.23, 355.84, -9.15, 0.48)
        apic[53].pt3dadd(190.23, 355.84, -9.15, 0.48)
        
        apic[54] = h.Section(name='apic[54]')
        apic[54].connect(        apic[52](1), 0)
        apic[54].pt3dclear()
        apic[54].pt3dadd(72.83, 286.19, 13.10, 2.70)
        apic[54].pt3dadd(72.83, 288.42, 13.84, 2.70)
        apic[54].pt3dadd(72.83, 288.67, 13.84, 2.70)
        apic[54].pt3dadd(72.83, 290.17, 13.84, 2.70)
        apic[54].pt3dadd(72.83, 290.41, 13.84, 2.70)
        apic[54].pt3dadd(72.83, 292.16, 13.84, 2.70)
        apic[54].pt3dadd(72.83, 292.40, 13.84, 2.70)
        apic[54].pt3dadd(72.83, 294.15, 13.84, 2.70)
        apic[54].pt3dadd(72.83, 294.39, 13.84, 2.70)
        apic[54].pt3dadd(72.83, 295.39, 13.84, 2.70)
        
        apic[55] = h.Section(name='apic[55]')
        apic[55].connect(        apic[54](1), 0)
        apic[55].pt3dclear()
        apic[55].pt3dadd(72.83, 295.39, 13.84, 0.86)
        apic[55].pt3dadd(75.01, 295.89, 7.61, 0.86)
        apic[55].pt3dadd(76.94, 296.88, 7.61, 0.86)
        apic[55].pt3dadd(77.18, 296.88, 7.61, 0.86)
        apic[55].pt3dadd(78.63, 298.37, 7.61, 0.86)
        apic[55].pt3dadd(79.36, 301.11, 7.61, 0.72)
        apic[55].pt3dadd(79.84, 302.35, 7.61, 0.72)
        apic[55].pt3dadd(80.32, 302.60, 7.61, 0.72)
        apic[55].pt3dadd(81.77, 302.85, 7.61, 0.72)
        apic[55].pt3dadd(82.01, 302.85, 7.61, 0.72)
        apic[55].pt3dadd(83.46, 304.35, 7.61, 0.72)
        apic[55].pt3dadd(83.70, 304.59, 7.61, 0.72)
        apic[55].pt3dadd(85.15, 305.84, 7.61, 0.72)
        apic[55].pt3dadd(86.36, 306.09, 7.61, 0.72)
        apic[55].pt3dadd(87.09, 306.34, 7.61, 0.72)
        apic[55].pt3dadd(87.33, 306.34, 7.61, 0.72)
        apic[55].pt3dadd(89.26, 303.35, 4.75, 0.72)
        apic[55].pt3dadd(89.50, 303.60, 4.75, 0.72)
        apic[55].pt3dadd(91.43, 303.35, 4.75, 0.72)
        apic[55].pt3dadd(91.68, 303.35, 4.75, 0.72)
        apic[55].pt3dadd(93.12, 304.84, 4.75, 0.72)
        apic[55].pt3dadd(94.82, 305.59, 4.75, 0.72)
        apic[55].pt3dadd(95.06, 305.84, 4.75, 0.72)
        apic[55].pt3dadd(96.75, 306.58, 4.75, 0.72)
        apic[55].pt3dadd(98.68, 307.33, 4.75, 0.72)
        apic[55].pt3dadd(99.16, 307.58, 4.75, 0.72)
        apic[55].pt3dadd(101.58, 307.58, 4.00, 0.72)
        apic[55].pt3dadd(101.58, 307.58, 4.03, 0.72)
        apic[55].pt3dadd(103.27, 308.33, 4.06, 0.72)
        apic[55].pt3dadd(103.51, 308.57, 4.06, 0.72)
        apic[55].pt3dadd(103.99, 309.32, 4.06, 0.72)
        apic[55].pt3dadd(104.24, 309.57, 4.07, 0.72)
        apic[55].pt3dadd(104.96, 310.81, 4.07, 0.72)
        apic[55].pt3dadd(105.20, 310.81, 4.07, 0.72)
        apic[55].pt3dadd(106.65, 311.56, 4.07, 0.72)
        apic[55].pt3dadd(107.13, 311.56, 4.07, 0.72)
        apic[55].pt3dadd(110.52, 310.56, 4.06, 0.48)
        apic[55].pt3dadd(111.48, 310.56, 4.06, 0.48)
        apic[55].pt3dadd(112.93, 311.31, 4.06, 0.48)
        apic[55].pt3dadd(113.17, 311.56, 4.06, 0.48)
        apic[55].pt3dadd(113.90, 314.05, 4.06, 0.48)
        apic[55].pt3dadd(114.62, 314.54, 4.06, 0.48)
        apic[55].pt3dadd(115.11, 315.04, 4.06, 0.48)
        apic[55].pt3dadd(116.31, 315.54, 4.06, 0.48)
        apic[55].pt3dadd(119.94, 315.04, -1.60, 0.48)
        apic[55].pt3dadd(120.18, 315.04, -1.60, 0.48)
        apic[55].pt3dadd(121.87, 315.79, -1.60, 0.48)
        apic[55].pt3dadd(122.35, 316.04, -1.60, 0.48)
        apic[55].pt3dadd(123.80, 318.03, -1.60, 0.48)
        apic[55].pt3dadd(124.04, 318.52, -1.60, 0.48)
        apic[55].pt3dadd(124.77, 319.02, -1.60, 0.48)
        apic[55].pt3dadd(125.01, 319.27, -1.60, 0.48)
        apic[55].pt3dadd(126.46, 320.02, -1.60, 0.48)
        apic[55].pt3dadd(129.36, 319.27, -1.60, 0.48)
        apic[55].pt3dadd(130.08, 318.03, -1.60, 0.48)
        apic[55].pt3dadd(131.29, 317.78, -1.60, 0.48)
        apic[55].pt3dadd(131.53, 317.78, -1.60, 0.48)
        apic[55].pt3dadd(132.26, 317.78, -1.60, 0.48)
        apic[55].pt3dadd(132.74, 318.28, -1.60, 0.48)
        apic[55].pt3dadd(133.46, 319.52, -1.61, 0.48)
        apic[55].pt3dadd(133.95, 320.02, -1.61, 0.48)
        apic[55].pt3dadd(134.91, 320.02, -1.61, 0.48)
        apic[55].pt3dadd(137.33, 320.02, -1.61, 0.48)
        apic[55].pt3dadd(137.57, 320.02, -1.61, 0.48)
        apic[55].pt3dadd(139.02, 320.02, -1.61, 0.48)
        apic[55].pt3dadd(140.23, 320.02, -1.61, 0.48)
        apic[55].pt3dadd(142.40, 319.77, -3.59, 0.48)
        apic[55].pt3dadd(144.57, 322.01, -3.59, 0.48)
        apic[55].pt3dadd(144.57, 322.50, -3.59, 0.48)
        apic[55].pt3dadd(145.06, 324.00, -3.59, 0.48)
        apic[55].pt3dadd(145.54, 324.25, -3.59, 0.48)
        apic[55].pt3dadd(146.51, 326.73, -5.53, 0.48)
        apic[55].pt3dadd(146.51, 326.73, -5.55, 0.48)
        apic[55].pt3dadd(147.47, 328.23, -5.99, 0.48)
        apic[55].pt3dadd(147.47, 328.23, -6.03, 0.48)
        apic[55].pt3dadd(148.92, 328.97, -6.03, 0.48)
        apic[55].pt3dadd(149.16, 328.97, -6.03, 0.48)
        apic[55].pt3dadd(150.37, 329.72, -6.23, 0.48)
        apic[55].pt3dadd(151.10, 329.47, -6.23, 0.48)
        apic[55].pt3dadd(151.34, 329.22, -6.23, 0.48)
        apic[55].pt3dadd(151.58, 327.23, -6.23, 0.48)
        apic[55].pt3dadd(151.58, 326.98, -6.23, 0.48)
        apic[55].pt3dadd(153.27, 326.98, -6.25, 0.48)
        apic[55].pt3dadd(155.44, 327.98, -6.25, 0.48)
        apic[55].pt3dadd(155.69, 328.23, -6.24, 0.48)
        apic[55].pt3dadd(157.13, 328.97, -6.24, 0.48)
        apic[55].pt3dadd(159.55, 330.22, -9.58, 0.48)
        apic[55].pt3dadd(161.97, 331.21, -9.58, 0.48)
        apic[55].pt3dadd(162.45, 331.46, -9.58, 0.48)
        apic[55].pt3dadd(162.69, 331.71, -9.58, 0.48)
        apic[55].pt3dadd(162.93, 331.71, -9.58, 0.48)
        apic[55].pt3dadd(163.66, 332.70, -9.59, 0.48)
        apic[55].pt3dadd(163.90, 332.95, -9.59, 0.48)
        apic[55].pt3dadd(165.59, 333.95, -9.59, 0.48)
        apic[55].pt3dadd(166.80, 333.95, -9.60, 0.48)
        apic[55].pt3dadd(167.28, 333.95, -9.60, 0.48)
        apic[55].pt3dadd(168.73, 333.20, -9.60, 0.48)
        apic[55].pt3dadd(169.21, 333.20, -9.60, 0.48)
        apic[55].pt3dadd(171.87, 334.94, -9.60, 0.48)
        apic[55].pt3dadd(172.11, 334.94, -9.60, 0.48)
        apic[55].pt3dadd(172.59, 335.94, -9.61, 0.48)
        apic[55].pt3dadd(173.08, 336.19, -9.61, 0.48)
        apic[55].pt3dadd(174.77, 336.68, -10.71, 0.48)
        apic[55].pt3dadd(174.77, 336.93, -10.72, 0.48)
        apic[55].pt3dadd(175.98, 339.42, -10.72, 0.48)
        apic[55].pt3dadd(176.22, 339.92, -10.72, 0.48)
        apic[55].pt3dadd(176.46, 340.41, -10.72, 0.48)
        apic[55].pt3dadd(176.94, 340.66, -10.73, 0.48)
        apic[55].pt3dadd(178.87, 340.66, -10.73, 0.48)
        apic[55].pt3dadd(179.12, 340.17, -10.73, 0.39)
        apic[55].pt3dadd(180.08, 339.17, -10.73, 0.39)
        apic[55].pt3dadd(181.53, 338.92, -10.73, 0.39)
        apic[55].pt3dadd(182.01, 338.42, -10.73, 0.39)
        apic[55].pt3dadd(182.98, 338.42, -10.75, 0.39)
        apic[55].pt3dadd(184.19, 338.42, -13.32, 0.39)
        apic[55].pt3dadd(184.19, 338.42, -13.35, 0.39)
        apic[55].pt3dadd(185.88, 339.42, -13.35, 0.39)
        apic[55].pt3dadd(185.88, 339.67, -13.36, 0.39)
        apic[55].pt3dadd(186.60, 340.41, -13.36, 0.39)
        apic[55].pt3dadd(187.09, 341.91, -13.36, 0.39)
        apic[55].pt3dadd(188.05, 344.39, -14.01, 0.39)
        apic[55].pt3dadd(188.29, 344.64, -14.01, 0.39)
        apic[55].pt3dadd(188.54, 345.39, -14.01, 0.39)
        apic[55].pt3dadd(189.02, 345.39, -14.01, 0.39)
        apic[55].pt3dadd(189.74, 345.89, -14.01, 0.39)
        apic[55].pt3dadd(190.47, 346.38, -14.01, 0.39)
        apic[55].pt3dadd(190.95, 346.63, -14.01, 0.39)
        apic[55].pt3dadd(191.19, 346.88, -14.01, 0.39)
        apic[55].pt3dadd(191.68, 347.38, -14.01, 0.39)
        apic[55].pt3dadd(193.12, 348.62, -14.01, 0.39)
        apic[55].pt3dadd(193.85, 349.12, -14.01, 0.39)
        apic[55].pt3dadd(195.30, 349.62, -14.01, 0.39)
        apic[55].pt3dadd(195.54, 349.87, -14.01, 0.39)
        apic[55].pt3dadd(195.78, 349.87, -14.01, 0.39)
        
        apic[56] = h.Section(name='apic[56]')
        apic[56].connect(        apic[54](1), 0)
        apic[56].pt3dclear()
        apic[56].pt3dadd(72.83, 295.39, 13.84, 2.18)
        apic[56].pt3dadd(72.11, 298.62, 15.22, 2.18)
        apic[56].pt3dadd(71.87, 300.36, 15.22, 2.18)
        apic[56].pt3dadd(72.11, 303.10, 15.22, 2.18)
        apic[56].pt3dadd(72.35, 303.60, 15.22, 2.18)
        apic[56].pt3dadd(72.83, 308.33, 15.22, 2.18)
        apic[56].pt3dadd(73.56, 310.81, 15.22, 2.18)
        apic[56].pt3dadd(74.28, 313.05, 9.43, 2.70)
        apic[56].pt3dadd(74.04, 313.30, 9.43, 2.70)
        apic[56].pt3dadd(74.53, 316.78, 9.43, 2.70)
        apic[56].pt3dadd(75.01, 317.28, 9.43, 2.70)
        apic[56].pt3dadd(75.25, 319.02, 9.43, 2.70)
        apic[56].pt3dadd(75.49, 319.52, 9.44, 2.70)
        apic[56].pt3dadd(76.22, 321.26, 9.44, 2.70)
        apic[56].pt3dadd(76.70, 322.26, 9.45, 2.70)
        apic[56].pt3dadd(78.15, 324.25, 9.45, 2.70)
        apic[56].pt3dadd(80.08, 326.24, 9.45, 2.70)
        apic[56].pt3dadd(81.05, 327.73, 9.45, 2.70)
        apic[56].pt3dadd(82.26, 328.97, 9.45, 2.70)
        apic[56].pt3dadd(82.74, 329.22, 9.46, 2.70)
        apic[56].pt3dadd(84.19, 331.46, 9.46, 2.70)
        apic[56].pt3dadd(84.19, 332.45, 9.46, 2.70)
        apic[56].pt3dadd(84.19, 334.20, 9.46, 2.70)
        apic[56].pt3dadd(84.19, 334.94, 9.46, 2.70)
        apic[56].pt3dadd(84.19, 337.18, 9.46, 2.70)
        apic[56].pt3dadd(83.95, 337.93, 9.48, 2.70)
        apic[56].pt3dadd(83.46, 339.67, 9.48, 2.70)
        apic[56].pt3dadd(83.46, 341.91, 9.48, 2.70)
        apic[56].pt3dadd(83.22, 345.39, 9.48, 2.70)
        apic[56].pt3dadd(83.22, 348.62, 9.48, 2.60)
        apic[56].pt3dadd(83.22, 349.12, 9.48, 2.60)
        apic[56].pt3dadd(84.91, 351.86, 9.48, 2.60)
        apic[56].pt3dadd(85.88, 353.35, 9.48, 2.60)
        apic[56].pt3dadd(86.36, 353.85, 9.48, 2.60)
        apic[56].pt3dadd(87.09, 355.09, 9.48, 2.60)
        apic[56].pt3dadd(88.05, 356.83, 9.48, 2.60)
        apic[56].pt3dadd(88.78, 358.33, 9.48, 2.60)
        apic[56].pt3dadd(89.02, 358.57, 9.48, 2.60)
        apic[56].pt3dadd(90.23, 360.07, 9.48, 2.60)
        apic[56].pt3dadd(90.47, 360.32, 9.48, 2.60)
        apic[56].pt3dadd(92.88, 362.31, 9.48, 2.60)
        
        apic[57] = h.Section(name='apic[57]')
        apic[57].connect(        apic[56](1), 0)
        apic[57].pt3dclear()
        apic[57].pt3dadd(92.88, 362.31, 9.48, 0.48)
        apic[57].pt3dadd(92.40, 365.29, 15.33, 0.48)
        apic[57].pt3dadd(92.40, 365.29, 15.34, 0.48)
        apic[57].pt3dadd(92.64, 367.03, 15.34, 0.48)
        apic[57].pt3dadd(92.64, 367.78, 15.34, 0.48)
        apic[57].pt3dadd(92.88, 368.77, 15.34, 0.48)
        apic[57].pt3dadd(92.16, 369.27, 15.34, 0.48)
        apic[57].pt3dadd(91.43, 369.52, 15.34, 0.48)
        apic[57].pt3dadd(90.71, 370.02, 15.34, 0.48)
        apic[57].pt3dadd(90.71, 370.76, 15.34, 0.48)
        apic[57].pt3dadd(90.71, 371.01, 15.34, 0.48)
        apic[57].pt3dadd(90.71, 372.26, 15.34, 0.48)
        apic[57].pt3dadd(90.71, 373.75, 15.34, 0.48)
        apic[57].pt3dadd(90.71, 374.25, 15.34, 0.48)
        apic[57].pt3dadd(90.47, 375.24, 15.34, 0.48)
        apic[57].pt3dadd(89.98, 375.49, 15.36, 0.48)
        apic[57].pt3dadd(89.26, 375.99, 15.37, 0.48)
        apic[57].pt3dadd(88.29, 376.98, 15.38, 0.48)
        apic[57].pt3dadd(87.81, 377.73, 15.38, 0.48)
        apic[57].pt3dadd(87.57, 377.73, 15.39, 0.48)
        apic[57].pt3dadd(86.36, 378.47, 15.47, 0.48)
        apic[57].pt3dadd(86.12, 378.47, 15.47, 0.48)
        apic[57].pt3dadd(85.40, 378.47, 15.48, 0.48)
        apic[57].pt3dadd(84.67, 378.47, 15.48, 0.48)
        apic[57].pt3dadd(84.19, 378.47, 15.48, 0.48)
        apic[57].pt3dadd(83.70, 378.72, 15.49, 0.48)
        apic[57].pt3dadd(83.22, 379.22, 15.49, 0.48)
        apic[57].pt3dadd(82.74, 381.46, 15.50, 0.48)
        apic[57].pt3dadd(82.26, 381.96, 15.50, 0.48)
        apic[57].pt3dadd(81.77, 383.95, 19.43, 0.48)
        apic[57].pt3dadd(81.77, 383.95, 19.26, 0.48)
        apic[57].pt3dadd(81.05, 384.94, 18.62, 0.48)
        apic[57].pt3dadd(80.08, 385.44, 18.38, 0.48)
        apic[57].pt3dadd(79.12, 385.69, 18.18, 0.48)
        apic[57].pt3dadd(77.42, 385.94, 18.09, 0.48)
        apic[57].pt3dadd(76.22, 386.43, 18.05, 0.48)
        apic[57].pt3dadd(76.22, 389.42, 18.05, 0.48)
        apic[57].pt3dadd(76.22, 389.42, 18.05, 0.48)
        
        apic[58] = h.Section(name='apic[58]')
        apic[58].connect(        apic[56](1), 0)
        apic[58].pt3dclear()
        apic[58].pt3dadd(92.88, 362.31, 9.48, 3.5)
        apic[58].pt3dadd(94.57, 363.30, 9.48, 3.5)
        apic[58].pt3dadd(95.54, 365.29, 9.48, 3.5)
        apic[58].pt3dadd(95.78, 365.29, 9.48, 3.5)
        apic[58].pt3dadd(96.99, 367.03, 9.48, 3.5)
        apic[58].pt3dadd(98.92, 368.77, 9.48, 3.5)
        
        apic[59] = h.Section(name='apic[59]')
        apic[59].connect(        apic[58](1), 0)
        apic[59].pt3dclear()
        apic[59].pt3dadd(98.92, 368.77, 9.48, 1.40)
        apic[59].pt3dadd(101.58, 368.52, 9.47, 1.40)
        apic[59].pt3dadd(102.55, 369.77, 9.47, 1.40)
        apic[59].pt3dadd(103.75, 370.76, 9.47, 1.40)
        apic[59].pt3dadd(103.99, 371.01, 9.47, 1.40)
        apic[59].pt3dadd(104.72, 371.76, 9.47, 1.40)
        apic[59].pt3dadd(106.89, 372.75, 8.45, 1.20)
        apic[59].pt3dadd(109.31, 374.00, 8.45, 1.20)
        apic[59].pt3dadd(109.55, 374.25, 8.45, 1.20)
        apic[59].pt3dadd(110.76, 375.49, 8.45, 1.20)
        apic[59].pt3dadd(111.48, 375.49, 8.45, 1.20)
        apic[59].pt3dadd(114.14, 375.99, 8.45, 1.20)
        apic[59].pt3dadd(114.62, 376.24, 8.45, 1.20)
        apic[59].pt3dadd(116.07, 376.48, 8.45, 1.20)
        apic[59].pt3dadd(116.55, 376.24, 8.45, 1.20)
        apic[59].pt3dadd(119.21, 374.99, 8.45, 1.20)
        apic[59].pt3dadd(119.45, 374.99, 8.45, 1.20)
        apic[59].pt3dadd(122.83, 374.25, 8.45, 1.20)
        apic[59].pt3dadd(122.83, 374.00, 8.45, 1.20)
        apic[59].pt3dadd(124.28, 373.00, 8.45, 1.20)
        apic[59].pt3dadd(124.77, 372.75, 8.45, 1.20)
        apic[59].pt3dadd(125.25, 372.26, 8.45, 1.20)
        apic[59].pt3dadd(125.73, 372.01, 8.45, 1.20)
        apic[59].pt3dadd(127.91, 371.26, 8.45, 1.20)
        apic[59].pt3dadd(128.39, 371.26, 8.45, 1.20)
        apic[59].pt3dadd(130.56, 371.26, 8.45, 1.20)
        apic[59].pt3dadd(132.01, 372.01, 8.45, 1.20)
        apic[59].pt3dadd(132.50, 372.26, 8.45, 1.20)
        apic[59].pt3dadd(134.43, 372.75, 8.45, 1.20)
        apic[59].pt3dadd(135.88, 373.00, 8.45, 1.20)
        apic[59].pt3dadd(136.84, 373.25, 8.45, 1.20)
        apic[59].pt3dadd(137.09, 373.25, 8.45, 1.20)
        apic[59].pt3dadd(140.23, 373.25, 8.45, 1.20)
        apic[59].pt3dadd(140.47, 373.25, 8.45, 1.20)
        apic[59].pt3dadd(142.40, 373.25, 8.45, 1.20)
        apic[59].pt3dadd(142.88, 373.25, 8.45, 1.20)
        apic[59].pt3dadd(144.57, 373.25, 8.45, 1.20)
        apic[59].pt3dadd(145.06, 373.25, 8.45, 1.10)
        apic[59].pt3dadd(145.54, 374.99, 8.45, 1.10)
        apic[59].pt3dadd(146.26, 378.23, 8.45, 1.10)
        apic[59].pt3dadd(148.68, 378.72, 8.45, 1.10)
        apic[59].pt3dadd(149.65, 378.72, 8.45, 1.10)
        apic[59].pt3dadd(151.34, 378.72, 8.45, 1.10)
        apic[59].pt3dadd(152.79, 379.22, 8.45, 1.10)
        apic[59].pt3dadd(154.24, 379.72, 8.45, 1.10)
        apic[59].pt3dadd(154.48, 379.97, 8.45, 1.10)
        apic[59].pt3dadd(156.41, 380.46, 8.45, 1.10)
        apic[59].pt3dadd(159.31, 380.96, 6.09, 1.10)
        apic[59].pt3dadd(160.76, 381.71, 6.09, 1.10)
        apic[59].pt3dadd(161.00, 381.71, 6.09, 1.10)
        apic[59].pt3dadd(163.17, 382.95, 6.09, 1.10)
        apic[59].pt3dadd(163.66, 383.20, 6.09, 1.10)
        apic[59].pt3dadd(165.59, 383.45, 6.09, 1.10)
        apic[59].pt3dadd(165.83, 383.45, 6.09, 1.10)
        apic[59].pt3dadd(168.25, 383.20, 6.09, 1.10)
        apic[59].pt3dadd(168.73, 383.20, 6.09, 1.10)
        apic[59].pt3dadd(170.90, 382.95, 6.09, 1.10)
        apic[59].pt3dadd(171.39, 382.95, 6.09, 1.10)
        apic[59].pt3dadd(173.32, 382.45, 6.09, 1.10)
        apic[59].pt3dadd(173.80, 382.45, 6.09, 1.10)
        apic[59].pt3dadd(175.25, 382.45, 6.09, 1.10)
        apic[59].pt3dadd(175.73, 382.45, 6.09, 1.10)
        apic[59].pt3dadd(177.42, 382.45, 6.09, 1.10)
        apic[59].pt3dadd(177.42, 382.70, 6.09, 1.10)
        apic[59].pt3dadd(178.87, 383.45, 6.09, 1.10)
        apic[59].pt3dadd(180.32, 384.94, 6.09, 1.10)
        apic[59].pt3dadd(180.56, 385.19, 6.09, 1.10)
        apic[59].pt3dadd(180.81, 385.44, 6.09, 1.10)
        apic[59].pt3dadd(182.01, 386.19, 6.09, 1.10)
        apic[59].pt3dadd(182.01, 386.43, 6.09, 1.10)
        apic[59].pt3dadd(183.70, 386.93, 6.09, 1.10)
        apic[59].pt3dadd(183.95, 387.18, 6.09, 1.10)
        apic[59].pt3dadd(185.64, 387.68, 6.09, 1.10)
        apic[59].pt3dadd(186.36, 387.68, 6.09, 1.10)
        apic[59].pt3dadd(187.09, 388.67, 6.09, 1.10)
        apic[59].pt3dadd(188.05, 390.17, 6.09, 1.10)
        apic[59].pt3dadd(188.05, 391.16, 6.09, 1.10)
        apic[59].pt3dadd(187.33, 391.91, 6.09, 1.10)
        
        apic[60] = h.Section(name='apic[60]')
        apic[60].connect(        apic[59](1), 0)
        apic[60].pt3dclear()
        apic[60].pt3dadd(187.33, 391.91, 6.09, 0.86)
        apic[60].pt3dadd(189.26, 393.40, 1.09, 0.86)
        apic[60].pt3dadd(191.19, 394.15, 1.09, 0.86)
        apic[60].pt3dadd(191.43, 394.39, 1.09, 0.86)
        apic[60].pt3dadd(193.61, 395.14, 1.09, 0.86)
        apic[60].pt3dadd(194.57, 395.64, 1.09, 0.86)
        apic[60].pt3dadd(195.78, 396.38, 1.09, 0.86)
        apic[60].pt3dadd(196.26, 396.38, 1.09, 0.86)
        apic[60].pt3dadd(195.54, 397.13, 1.08, 0.86)
        apic[60].pt3dadd(195.54, 397.38, 1.08, 0.86)
        apic[60].pt3dadd(194.82, 398.87, 1.08, 0.86)
        apic[60].pt3dadd(194.57, 399.12, 1.08, 0.86)
        apic[60].pt3dadd(194.33, 400.86, 1.08, 0.86)
        apic[60].pt3dadd(195.54, 401.36, 1.08, 0.86)
        apic[60].pt3dadd(195.78, 401.36, 1.08, 0.86)
        apic[60].pt3dadd(196.75, 400.37, 1.09, 0.77)
        apic[60].pt3dadd(196.99, 400.37, 1.09, 0.77)
        apic[60].pt3dadd(198.68, 401.61, -0.94, 0.77)
        apic[60].pt3dadd(199.89, 403.60, -0.98, 0.77)
        apic[60].pt3dadd(200.13, 403.60, -0.98, 0.77)
        apic[60].pt3dadd(201.34, 402.85, -0.98, 0.77)
        apic[60].pt3dadd(201.58, 402.60, -0.98, 0.77)
        apic[60].pt3dadd(203.03, 404.10, -1.03, 0.77)
        apic[60].pt3dadd(203.51, 404.10, -1.03, 0.77)
        apic[60].pt3dadd(205.69, 406.58, -1.03, 0.77)
        apic[60].pt3dadd(205.44, 408.08, -1.04, 0.77)
        apic[60].pt3dadd(205.20, 408.33, -1.04, 0.77)
        apic[60].pt3dadd(204.72, 408.82, -1.04, 0.77)
        apic[60].pt3dadd(204.48, 409.82, -1.04, 0.77)
        apic[60].pt3dadd(204.48, 410.56, -1.04, 0.77)
        apic[60].pt3dadd(203.99, 411.56, -1.04, 0.77)
        apic[60].pt3dadd(204.96, 412.31, -1.04, 0.77)
        apic[60].pt3dadd(206.17, 412.80, -1.04, 0.77)
        apic[60].pt3dadd(206.17, 413.05, -1.04, 0.77)
        apic[60].pt3dadd(207.62, 414.54, -1.06, 0.77)
        apic[60].pt3dadd(208.34, 415.04, -1.08, 0.77)
        apic[60].pt3dadd(208.58, 415.79, -1.10, 0.77)
        apic[60].pt3dadd(209.31, 416.53, -1.12, 0.77)
        apic[60].pt3dadd(209.55, 417.28, -1.12, 0.77)
        apic[60].pt3dadd(209.79, 417.53, -1.13, 0.77)
        apic[60].pt3dadd(211.00, 418.03, -1.13, 0.77)
        apic[60].pt3dadd(211.72, 418.28, -1.13, 0.77)
        apic[60].pt3dadd(213.90, 417.53, -4.75, 0.77)
        apic[60].pt3dadd(213.90, 417.53, -4.76, 0.77)
        apic[60].pt3dadd(215.59, 418.28, -4.77, 0.77)
        apic[60].pt3dadd(216.80, 418.77, -4.78, 0.77)
        apic[60].pt3dadd(217.28, 419.02, -4.78, 0.77)
        apic[60].pt3dadd(217.28, 419.27, -4.81, 0.77)
        apic[60].pt3dadd(217.04, 420.51, -5.85, 0.54)
        apic[60].pt3dadd(217.04, 420.76, -5.85, 0.54)
        apic[60].pt3dadd(216.80, 422.01, -5.86, 0.54)
        apic[60].pt3dadd(216.80, 422.50, -5.86, 0.54)
        apic[60].pt3dadd(217.04, 423.50, -5.86, 0.54)
        apic[60].pt3dadd(217.52, 424.00, -5.86, 0.54)
        apic[60].pt3dadd(218.73, 424.74, -5.86, 0.54)
        apic[60].pt3dadd(219.69, 424.99, -5.86, 0.54)
        apic[60].pt3dadd(220.90, 425.49, -5.86, 0.54)
        apic[60].pt3dadd(221.14, 425.49, -5.86, 0.54)
        apic[60].pt3dadd(221.87, 425.74, -5.86, 0.54)
        apic[60].pt3dadd(222.59, 425.99, -5.86, 0.54)
        apic[60].pt3dadd(223.32, 426.24, -5.86, 0.54)
        apic[60].pt3dadd(223.56, 426.48, -5.86, 0.54)
        apic[60].pt3dadd(224.28, 427.98, -8.42, 0.54)
        apic[60].pt3dadd(225.73, 429.72, -8.44, 0.54)
        apic[60].pt3dadd(226.22, 430.22, -8.44, 0.54)
        apic[60].pt3dadd(227.18, 430.71, -8.44, 0.54)
        apic[60].pt3dadd(228.15, 431.21, -8.44, 0.54)
        apic[60].pt3dadd(228.63, 431.46, -8.44, 0.54)
        apic[60].pt3dadd(228.87, 431.71, -8.44, 0.54)
        apic[60].pt3dadd(228.87, 432.70, -8.45, 0.54)
        apic[60].pt3dadd(228.87, 433.20, -8.49, 0.54)
        apic[60].pt3dadd(228.87, 433.95, -8.55, 0.54)
        apic[60].pt3dadd(228.87, 433.95, -8.58, 0.54)
        apic[60].pt3dadd(229.12, 436.68, -8.66, 0.54)
        apic[60].pt3dadd(229.12, 436.93, -8.68, 0.54)
        apic[60].pt3dadd(229.36, 437.43, -8.68, 0.54)
        apic[60].pt3dadd(230.56, 438.42, -8.70, 0.54)
        apic[60].pt3dadd(231.53, 439.17, -8.70, 0.54)
        apic[60].pt3dadd(232.01, 439.17, -8.72, 0.54)
        apic[60].pt3dadd(232.26, 439.42, -8.76, 0.54)
        apic[60].pt3dadd(235.64, 441.66, -11.17, 0.54)
        apic[60].pt3dadd(236.84, 442.16, -11.18, 0.54)
        apic[60].pt3dadd(237.57, 442.40, -11.18, 0.54)
        apic[60].pt3dadd(239.26, 442.90, -11.19, 0.54)
        apic[60].pt3dadd(239.50, 443.15, -11.19, 0.54)
        apic[60].pt3dadd(239.74, 445.39, -11.19, 0.54)
        apic[60].pt3dadd(239.74, 445.39, -11.20, 0.54)
        apic[60].pt3dadd(238.29, 446.63, -13.03, 0.54)
        apic[60].pt3dadd(238.29, 448.13, -13.03, 0.54)
        apic[60].pt3dadd(238.29, 448.13, -13.04, 0.54)
        apic[60].pt3dadd(239.02, 448.87, -13.05, 0.54)
        apic[60].pt3dadd(241.92, 449.87, -17.59, 0.54)
        apic[60].pt3dadd(242.64, 453.60, -17.63, 0.54)
        apic[60].pt3dadd(242.64, 453.60, -17.64, 0.54)
        apic[60].pt3dadd(243.37, 455.59, -17.64, 0.54)
        apic[60].pt3dadd(243.37, 456.83, -17.64, 0.54)
        apic[60].pt3dadd(243.37, 457.83, -17.65, 0.54)
        apic[60].pt3dadd(243.37, 458.57, -17.65, 0.54)
        apic[60].pt3dadd(243.85, 461.56, -17.65, 0.54)
        apic[60].pt3dadd(243.85, 461.81, -17.65, 0.54)
        apic[60].pt3dadd(244.33, 462.55, -17.65, 0.54)
        apic[60].pt3dadd(245.78, 463.55, -17.65, 0.54)
        apic[60].pt3dadd(246.75, 463.80, -17.65, 0.54)
        apic[60].pt3dadd(247.96, 463.55, -17.65, 0.54)
        apic[60].pt3dadd(248.92, 463.55, -17.65, 0.54)
        apic[60].pt3dadd(249.41, 463.55, -17.65, 0.54)
        apic[60].pt3dadd(250.13, 463.80, -17.65, 0.54)
        apic[60].pt3dadd(250.61, 465.54, -17.65, 0.54)
        apic[60].pt3dadd(250.85, 469.02, -17.65, 0.54)
        apic[60].pt3dadd(251.10, 470.02, -17.65, 0.54)
        apic[60].pt3dadd(251.58, 471.01, -17.65, 0.54)
        apic[60].pt3dadd(252.30, 471.76, -17.65, 0.54)
        apic[60].pt3dadd(252.55, 472.01, -17.65, 0.54)
        apic[60].pt3dadd(253.03, 472.01, -17.65, 0.54)
        apic[60].pt3dadd(253.51, 472.26, -17.65, 0.54)
        apic[60].pt3dadd(254.24, 472.26, -17.65, 0.54)
        apic[60].pt3dadd(256.17, 472.75, -17.65, 0.54)
        apic[60].pt3dadd(257.38, 473.75, -17.65, 0.43)
        apic[60].pt3dadd(258.10, 474.74, -22.25, 0.43)
        apic[60].pt3dadd(259.55, 478.23, -22.26, 0.43)
        apic[60].pt3dadd(259.79, 478.23, -22.26, 0.43)
        apic[60].pt3dadd(260.03, 479.47, -22.27, 0.43)
        apic[60].pt3dadd(260.27, 480.46, -22.29, 0.43)
        apic[60].pt3dadd(260.76, 481.46, -22.29, 0.43)
        apic[60].pt3dadd(261.00, 481.96, -22.29, 0.43)
        apic[60].pt3dadd(261.24, 482.45, -22.29, 0.43)
        apic[60].pt3dadd(261.72, 482.21, -22.30, 0.43)
        apic[60].pt3dadd(262.45, 481.96, -22.30, 0.43)
        apic[60].pt3dadd(266.80, 481.21, -22.32, 0.43)
        apic[60].pt3dadd(267.04, 481.21, -22.32, 0.43)
        apic[60].pt3dadd(267.28, 481.21, -22.32, 0.43)
        apic[60].pt3dadd(268.00, 482.70, -22.33, 0.43)
        apic[60].pt3dadd(268.00, 483.45, -22.33, 0.43)
        apic[60].pt3dadd(268.00, 483.95, -22.33, 0.43)
        apic[60].pt3dadd(268.00, 484.20, -22.33, 0.43)
        apic[60].pt3dadd(268.00, 485.19, -22.33, 0.43)
        apic[60].pt3dadd(268.25, 485.44, -22.33, 0.43)
        apic[60].pt3dadd(268.97, 486.19, -22.33, 0.43)
        apic[60].pt3dadd(269.94, 486.68, -22.34, 0.43)
        apic[60].pt3dadd(270.42, 486.68, -22.34, 0.43)
        apic[60].pt3dadd(274.77, 488.18, -22.34, 0.43)
        apic[60].pt3dadd(274.28, 491.41, -27.61, 0.43)
        apic[60].pt3dadd(275.73, 494.89, -27.62, 0.43)
        apic[60].pt3dadd(275.98, 497.13, -27.63, 0.43)
        apic[60].pt3dadd(275.98, 497.38, -27.63, 0.43)
        apic[60].pt3dadd(277.42, 499.87, -31.96, 0.43)
        apic[60].pt3dadd(277.67, 500.12, -31.96, 0.43)
        apic[60].pt3dadd(276.70, 503.85, -31.96, 0.43)
        apic[60].pt3dadd(279.12, 503.85, -31.96, 0.43)
        apic[60].pt3dadd(281.05, 503.60, -35.32, 0.43)
        apic[60].pt3dadd(280.56, 506.34, -35.35, 0.43)
        apic[60].pt3dadd(280.56, 506.34, -35.36, 0.43)
        apic[60].pt3dadd(280.56, 508.08, -35.47, 0.43)
        apic[60].pt3dadd(281.53, 508.33, -35.65, 0.43)
        apic[60].pt3dadd(283.46, 509.57, -35.74, 0.43)
        apic[60].pt3dadd(284.43, 510.32, -35.80, 0.43)
        apic[60].pt3dadd(285.15, 510.56, -35.84, 0.43)
        apic[60].pt3dadd(285.15, 510.56, -35.85, 0.43)
        apic[60].pt3dadd(285.40, 511.56, -35.93, 0.43)
        apic[60].pt3dadd(285.40, 511.81, -36.05, 0.43)
        apic[60].pt3dadd(283.95, 514.54, -36.66, 0.43)
        apic[60].pt3dadd(283.95, 514.79, -36.83, 0.43)
        apic[60].pt3dadd(283.95, 517.28, -37.28, 0.43)
        apic[60].pt3dadd(284.67, 518.03, -37.34, 0.43)
        apic[60].pt3dadd(285.40, 518.52, -37.81, 0.39)
        apic[60].pt3dadd(285.64, 518.52, -37.97, 0.39)
        apic[60].pt3dadd(286.84, 519.02, -38.04, 0.39)
        apic[60].pt3dadd(287.57, 519.27, -38.09, 0.39)
        apic[60].pt3dadd(289.50, 519.52, -38.17, 0.39)
        apic[60].pt3dadd(289.74, 519.52, -38.19, 0.39)
        apic[60].pt3dadd(291.19, 520.51, -42.56, 0.39)
        apic[60].pt3dadd(291.19, 520.51, -42.57, 0.39)
        apic[60].pt3dadd(291.92, 524.00, -44.01, 0.39)
        apic[60].pt3dadd(292.40, 524.25, -44.01, 0.39)
        apic[60].pt3dadd(292.88, 524.99, -44.00, 0.39)
        apic[60].pt3dadd(293.61, 525.49, -44.00, 0.39)
        apic[60].pt3dadd(294.33, 527.48, -44.79, 0.39)
        apic[60].pt3dadd(294.57, 527.73, -45.12, 0.39)
        apic[60].pt3dadd(294.82, 528.97, -45.61, 0.39)
        apic[60].pt3dadd(294.82, 528.97, -45.64, 0.39)
        apic[60].pt3dadd(295.78, 529.97, -45.65, 0.39)
        apic[60].pt3dadd(296.75, 531.21, -45.66, 0.39)
        apic[60].pt3dadd(297.23, 531.71, -45.66, 0.39)
        apic[60].pt3dadd(297.47, 532.21, -45.66, 0.39)
        apic[60].pt3dadd(297.71, 532.45, -45.66, 0.39)
        
        apic[61] = h.Section(name='apic[61]')
        apic[61].connect(        apic[59](1), 0)
        apic[61].pt3dclear()
        apic[61].pt3dadd(187.33, 391.91, 6.09, 0.96)
        apic[61].pt3dadd(187.81, 397.63, 7.80, 0.96)
        apic[61].pt3dadd(187.09, 400.61, 7.80, 0.96)
        apic[61].pt3dadd(187.09, 401.11, 7.80, 0.96)
        apic[61].pt3dadd(186.12, 404.35, 7.80, 0.96)
        apic[61].pt3dadd(186.12, 404.59, 7.80, 0.96)
        apic[61].pt3dadd(184.91, 407.83, 7.80, 0.96)
        apic[61].pt3dadd(184.67, 408.57, 7.80, 0.96)
        apic[61].pt3dadd(183.70, 410.56, 7.80, 0.96)
        apic[61].pt3dadd(182.74, 411.81, 7.80, 0.96)
        apic[61].pt3dadd(180.56, 412.55, 7.80, 0.96)
        apic[61].pt3dadd(180.32, 412.55, 7.80, 0.96)
        apic[61].pt3dadd(177.91, 413.80, 7.80, 0.96)
        apic[61].pt3dadd(177.18, 416.53, 4.31, 0.96)
        apic[61].pt3dadd(177.18, 419.02, 4.31, 0.96)
        apic[61].pt3dadd(177.18, 419.52, 4.31, 0.96)
        apic[61].pt3dadd(176.46, 421.01, 4.31, 0.96)
        apic[61].pt3dadd(175.98, 421.26, 4.31, 0.96)
        apic[61].pt3dadd(175.25, 422.50, 4.31, 0.96)
        apic[61].pt3dadd(175.25, 423.00, 4.31, 0.96)
        apic[61].pt3dadd(174.28, 425.24, 4.31, 0.96)
        apic[61].pt3dadd(174.28, 425.49, 4.31, 0.96)
        apic[61].pt3dadd(173.80, 426.48, 4.31, 0.96)
        apic[61].pt3dadd(173.80, 426.73, 4.31, 0.96)
        apic[61].pt3dadd(175.01, 428.47, 4.31, 0.96)
        apic[61].pt3dadd(175.25, 428.97, 4.31, 0.96)
        apic[61].pt3dadd(175.98, 430.46, 4.31, 0.96)
        apic[61].pt3dadd(177.42, 431.46, 4.31, 1.00)
        apic[61].pt3dadd(177.91, 432.21, 4.31, 1.00)
        apic[61].pt3dadd(179.60, 433.20, 4.31, 1.00)
        apic[61].pt3dadd(180.81, 433.45, 4.31, 1.00)
        apic[61].pt3dadd(182.26, 433.95, 4.31, 1.00)
        apic[61].pt3dadd(183.70, 434.94, 4.31, 1.00)
        apic[61].pt3dadd(184.43, 435.19, 4.31, 1.00)
        apic[61].pt3dadd(184.91, 435.44, 4.31, 1.00)
        apic[61].pt3dadd(189.02, 437.93, -0.48, 1.00)
        apic[61].pt3dadd(189.02, 437.93, -0.49, 1.00)
        apic[61].pt3dadd(188.54, 440.41, -2.54, 1.00)
        apic[61].pt3dadd(188.54, 440.41, -2.59, 1.00)
        apic[61].pt3dadd(188.29, 442.90, -3.41, 1.00)
        apic[61].pt3dadd(188.78, 443.65, -3.54, 1.00)
        apic[61].pt3dadd(189.50, 445.39, -3.78, 1.00)
        apic[61].pt3dadd(189.74, 445.39, -3.78, 1.00)
        apic[61].pt3dadd(191.92, 448.38, -5.07, 1.00)
        apic[61].pt3dadd(191.92, 448.38, -5.20, 1.00)
        apic[61].pt3dadd(194.82, 451.11, -6.58, 1.00)
        apic[61].pt3dadd(194.82, 451.11, -6.70, 1.00)
        apic[61].pt3dadd(195.54, 452.11, -6.71, 1.10)
        apic[61].pt3dadd(196.02, 452.85, -6.71, 1.10)
        apic[61].pt3dadd(196.51, 454.35, -6.71, 1.20)
        apic[61].pt3dadd(197.23, 455.59, -8.17, 1.20)
        
        apic[62] = h.Section(name='apic[62]')
        apic[62].connect(        apic[61](1), 0)
        apic[62].pt3dclear()
        apic[62].pt3dadd(197.23, 455.59, -8.17, 0.72)
        apic[62].pt3dadd(199.40, 456.09, -6.28, 0.72)
        apic[62].pt3dadd(200.13, 457.58, -6.28, 0.72)
        apic[62].pt3dadd(200.13, 460.07, -6.28, 0.72)
        apic[62].pt3dadd(199.89, 462.06, -6.28, 0.72)
        apic[62].pt3dadd(200.37, 462.31, -6.28, 0.72)
        apic[62].pt3dadd(201.34, 462.55, -6.28, 0.72)
        apic[62].pt3dadd(202.06, 462.80, -6.28, 0.72)
        apic[62].pt3dadd(203.51, 463.55, -6.28, 0.72)
        apic[62].pt3dadd(203.75, 463.55, -6.28, 0.72)
        apic[62].pt3dadd(206.41, 464.05, -2.74, 0.54)
        apic[62].pt3dadd(206.41, 464.05, -2.73, 0.54)
        apic[62].pt3dadd(208.58, 463.80, -2.72, 0.54)
        apic[62].pt3dadd(209.07, 464.05, -2.72, 0.54)
        apic[62].pt3dadd(210.52, 464.54, -2.72, 0.54)
        apic[62].pt3dadd(210.76, 464.79, -2.72, 0.54)
        apic[62].pt3dadd(211.72, 466.53, -2.67, 0.54)
        apic[62].pt3dadd(211.72, 466.78, -2.67, 0.54)
        apic[62].pt3dadd(211.97, 469.02, -2.67, 0.54)
        apic[62].pt3dadd(212.21, 469.27, -2.66, 0.54)
        apic[62].pt3dadd(212.45, 471.01, -2.20, 0.54)
        apic[62].pt3dadd(212.69, 471.26, -2.11, 0.54)
        apic[62].pt3dadd(212.69, 472.26, -1.36, 0.54)
        apic[62].pt3dadd(212.93, 474.74, -1.23, 0.54)
        apic[62].pt3dadd(212.93, 474.74, -1.22, 0.54)
        apic[62].pt3dadd(212.93, 476.24, -1.22, 0.54)
        apic[62].pt3dadd(212.93, 476.73, -1.22, 0.54)
        apic[62].pt3dadd(212.93, 478.47, -1.21, 0.54)
        apic[62].pt3dadd(212.93, 478.72, -1.17, 0.54)
        apic[62].pt3dadd(213.41, 479.22, -1.16, 0.54)
        apic[62].pt3dadd(214.14, 479.47, -1.16, 0.54)
        apic[62].pt3dadd(214.62, 479.72, -1.16, 0.54)
        apic[62].pt3dadd(215.59, 480.22, -1.16, 0.54)
        apic[62].pt3dadd(216.55, 480.71, -1.16, 0.54)
        apic[62].pt3dadd(216.80, 480.71, -1.14, 0.54)
        apic[62].pt3dadd(219.45, 482.21, -0.59, 0.54)
        apic[62].pt3dadd(220.66, 483.20, -0.59, 0.54)
        apic[62].pt3dadd(221.14, 483.70, -0.59, 0.54)
        apic[62].pt3dadd(221.63, 484.20, -0.57, 0.54)
        apic[62].pt3dadd(222.35, 484.69, -0.57, 0.54)
        apic[62].pt3dadd(222.59, 485.19, -0.57, 0.54)
        apic[62].pt3dadd(223.08, 485.69, -0.57, 0.43)
        apic[62].pt3dadd(223.80, 486.19, -0.57, 0.43)
        apic[62].pt3dadd(223.80, 486.43, -0.57, 0.43)
        apic[62].pt3dadd(224.04, 486.93, -0.56, 0.43)
        apic[62].pt3dadd(224.28, 486.93, -0.56, 0.43)
        apic[62].pt3dadd(224.77, 487.93, -0.56, 0.43)
        apic[62].pt3dadd(225.01, 490.17, -0.56, 0.43)
        apic[62].pt3dadd(225.01, 490.41, -0.56, 0.43)
        apic[62].pt3dadd(225.01, 491.41, -0.56, 0.43)
        apic[62].pt3dadd(225.01, 491.91, -0.56, 0.43)
        apic[62].pt3dadd(225.01, 493.15, -0.56, 0.43)
        apic[62].pt3dadd(224.77, 494.64, -0.56, 0.43)
        apic[62].pt3dadd(224.77, 494.89, -0.56, 0.43)
        apic[62].pt3dadd(225.25, 496.39, 3.24, 0.43)
        apic[62].pt3dadd(225.25, 496.88, 3.24, 0.43)
        apic[62].pt3dadd(226.22, 497.88, 3.24, 0.43)
        apic[62].pt3dadd(226.46, 498.13, 3.25, 0.43)
        apic[62].pt3dadd(226.94, 498.38, 3.32, 0.43)
        apic[62].pt3dadd(227.18, 499.12, 3.35, 0.43)
        apic[62].pt3dadd(227.18, 499.62, 3.42, 0.43)
        apic[62].pt3dadd(227.18, 501.61, 3.55, 0.43)
        apic[62].pt3dadd(227.18, 501.61, 3.56, 0.43)
        apic[62].pt3dadd(227.18, 503.60, 3.57, 0.43)
        apic[62].pt3dadd(226.94, 504.84, 3.59, 0.43)
        apic[62].pt3dadd(226.94, 504.84, 3.65, 0.43)
        apic[62].pt3dadd(227.42, 506.34, 4.24, 0.43)
        apic[62].pt3dadd(227.42, 506.34, 4.31, 0.43)
        apic[62].pt3dadd(227.91, 507.58, 4.34, 0.43)
        apic[62].pt3dadd(228.87, 508.33, 3.02, 0.43)
        apic[62].pt3dadd(227.42, 510.81, 3.02, 0.43)
        apic[62].pt3dadd(226.94, 514.05, 3.01, 0.43)
        apic[62].pt3dadd(226.94, 514.54, 3.01, 0.43)
        apic[62].pt3dadd(227.18, 515.04, 3.01, 0.43)
        apic[62].pt3dadd(227.42, 515.79, 3.01, 0.43)
        apic[62].pt3dadd(227.91, 517.03, 3.01, 0.43)
        apic[62].pt3dadd(227.91, 517.78, 3.01, 0.43)
        apic[62].pt3dadd(227.91, 519.77, 3.01, 0.43)
        apic[62].pt3dadd(227.91, 520.27, 3.01, 0.43)
        apic[62].pt3dadd(226.70, 522.26, 3.01, 0.43)
        apic[62].pt3dadd(226.46, 522.75, 3.01, 0.43)
        apic[62].pt3dadd(225.98, 524.00, 3.01, 0.43)
        apic[62].pt3dadd(225.98, 524.00, 3.02, 0.43)
        apic[62].pt3dadd(223.56, 526.48, 9.61, 0.43)
        apic[62].pt3dadd(223.56, 526.48, 9.62, 0.43)
        apic[62].pt3dadd(222.11, 530.96, 11.05, 0.43)
        apic[62].pt3dadd(222.11, 530.96, 11.07, 0.43)
        apic[62].pt3dadd(222.11, 533.20, 11.31, 0.43)
        apic[62].pt3dadd(220.18, 534.20, 11.33, 0.43)
        apic[62].pt3dadd(219.94, 534.44, 11.33, 0.43)
        apic[62].pt3dadd(218.97, 536.43, 11.34, 0.43)
        apic[62].pt3dadd(218.97, 537.18, 11.34, 0.43)
        apic[62].pt3dadd(218.97, 538.18, 11.34, 0.43)
        apic[62].pt3dadd(219.21, 539.17, 11.34, 0.43)
        apic[62].pt3dadd(219.45, 539.67, 11.34, 0.43)
        apic[62].pt3dadd(219.45, 540.66, 11.34, 0.43)
        apic[62].pt3dadd(218.73, 542.40, 11.34, 0.43)
        apic[62].pt3dadd(218.49, 542.65, 11.34, 0.43)
        apic[62].pt3dadd(218.00, 545.39, 11.33, 0.43)
        apic[62].pt3dadd(217.04, 546.39, 11.33, 0.43)
        apic[62].pt3dadd(217.04, 546.39, 11.30, 0.43)
        apic[62].pt3dadd(217.04, 546.39, 7.74, 0.43)
        apic[62].pt3dadd(215.83, 550.61, 7.72, 0.43)
        apic[62].pt3dadd(214.62, 553.85, 7.72, 0.43)
        apic[62].pt3dadd(214.62, 554.35, 7.72, 0.43)
        apic[62].pt3dadd(214.62, 555.59, 7.72, 0.43)
        apic[62].pt3dadd(214.62, 557.33, 7.72, 0.43)
        apic[62].pt3dadd(213.90, 558.08, 7.72, 0.43)
        apic[62].pt3dadd(212.45, 559.32, 7.72, 0.43)
        apic[62].pt3dadd(210.27, 563.80, 6.11, 0.43)
        apic[62].pt3dadd(209.07, 566.04, 6.11, 0.43)
        apic[62].pt3dadd(206.89, 569.27, 10.28, 0.43)
        apic[62].pt3dadd(206.65, 572.01, 10.28, 0.43)
        apic[62].pt3dadd(205.69, 574.25, 10.28, 0.43)
        apic[62].pt3dadd(205.44, 576.48, 10.29, 0.43)
        apic[62].pt3dadd(205.44, 576.73, 10.29, 0.43)
        apic[62].pt3dadd(204.72, 577.73, 10.29, 0.43)
        apic[62].pt3dadd(204.72, 577.98, 10.29, 0.43)
        apic[62].pt3dadd(203.51, 581.46, 10.29, 0.43)
        apic[62].pt3dadd(203.51, 581.71, 10.29, 0.43)
        apic[62].pt3dadd(203.75, 583.45, 10.29, 0.43)
        apic[62].pt3dadd(203.99, 584.20, 10.29, 0.43)
        apic[62].pt3dadd(204.24, 585.19, 10.29, 0.43)
        apic[62].pt3dadd(204.24, 585.94, 10.29, 0.43)
        apic[62].pt3dadd(203.27, 586.68, 10.29, 0.43)
        apic[62].pt3dadd(202.55, 586.93, 10.29, 0.43)
        apic[62].pt3dadd(202.30, 586.93, 10.29, 0.43)
        apic[62].pt3dadd(201.34, 586.68, 10.29, 0.43)
        apic[62].pt3dadd(200.13, 585.94, 10.29, 0.43)
        apic[62].pt3dadd(199.65, 585.44, 10.29, 0.43)
        apic[62].pt3dadd(197.96, 584.94, 10.29, 0.43)
        apic[62].pt3dadd(197.47, 584.94, 10.29, 0.43)
        apic[62].pt3dadd(194.09, 584.94, 10.30, 0.43)
        apic[62].pt3dadd(193.37, 584.94, 10.30, 0.43)
        apic[62].pt3dadd(191.43, 583.20, 13.61, 0.43)
        apic[62].pt3dadd(189.50, 583.45, 13.61, 0.43)
        apic[62].pt3dadd(189.02, 583.70, 13.66, 0.43)
        apic[62].pt3dadd(186.84, 584.44, 13.74, 0.43)
        apic[62].pt3dadd(186.84, 584.69, 13.85, 0.43)
        apic[62].pt3dadd(185.15, 586.19, 15.00, 0.39)
        apic[62].pt3dadd(185.15, 586.19, 15.01, 0.39)
        apic[62].pt3dadd(185.15, 588.18, 23.25, 0.39)
        apic[62].pt3dadd(184.91, 588.18, 23.25, 0.39)
        apic[62].pt3dadd(182.01, 589.17, 23.22, 0.39)
        apic[62].pt3dadd(181.53, 588.67, 23.16, 0.39)
        apic[62].pt3dadd(180.56, 588.67, 23.04, 0.39)
        apic[62].pt3dadd(180.08, 588.42, 23.03, 0.39)
        apic[62].pt3dadd(177.91, 587.43, 23.02, 0.39)
        apic[62].pt3dadd(177.91, 587.43, 23.01, 0.39)
        apic[62].pt3dadd(177.67, 587.43, 21.85, 0.39)
        apic[62].pt3dadd(177.67, 587.43, 21.60, 0.39)
        apic[62].pt3dadd(176.22, 587.43, 20.84, 0.39)
        apic[62].pt3dadd(175.25, 588.18, 20.84, 0.39)
        apic[62].pt3dadd(174.28, 588.18, 20.84, 0.39)
        apic[62].pt3dadd(172.35, 588.18, 20.84, 0.39)
        apic[62].pt3dadd(172.11, 588.18, 20.84, 0.39)
        apic[62].pt3dadd(171.39, 587.68, 20.84, 0.39)
        apic[62].pt3dadd(171.39, 587.43, 20.84, 0.39)
        apic[62].pt3dadd(169.94, 587.43, 20.84, 0.39)
        apic[62].pt3dadd(169.21, 587.43, 20.84, 0.39)
        apic[62].pt3dadd(168.97, 587.43, 20.84, 0.39)
        
        apic[63] = h.Section(name='apic[63]')
        apic[63].connect(        apic[61](1), 0)
        apic[63].pt3dclear()
        apic[63].pt3dadd(197.23, 455.59, -8.17, 0.86)
        apic[63].pt3dadd(197.23, 457.08, -6.44, 0.86)
        apic[63].pt3dadd(197.71, 459.82, -6.44, 0.86)
        apic[63].pt3dadd(197.23, 461.81, -6.44, 0.86)
        apic[63].pt3dadd(196.75, 463.80, -6.44, 0.86)
        apic[63].pt3dadd(196.26, 464.79, -6.44, 0.86)
        apic[63].pt3dadd(196.26, 465.29, -6.44, 0.86)
        apic[63].pt3dadd(196.75, 467.53, -6.44, 0.86)
        apic[63].pt3dadd(196.99, 468.28, -6.44, 0.86)
        apic[63].pt3dadd(197.96, 470.02, -6.44, 0.86)
        apic[63].pt3dadd(197.96, 470.27, -6.44, 0.86)
        apic[63].pt3dadd(199.65, 472.26, -6.03, 0.86)
        apic[63].pt3dadd(200.61, 473.75, -6.03, 0.86)
        apic[63].pt3dadd(201.34, 474.49, -6.03, 0.86)
        apic[63].pt3dadd(201.58, 474.74, -6.03, 0.86)
        apic[63].pt3dadd(203.27, 475.24, -6.03, 0.86)
        apic[63].pt3dadd(205.20, 476.24, -6.06, 0.86)
        apic[63].pt3dadd(207.62, 479.22, -6.19, 0.72)
        apic[63].pt3dadd(208.34, 479.97, -6.19, 0.72)
        apic[63].pt3dadd(208.58, 481.96, -6.19, 0.72)
        apic[63].pt3dadd(208.58, 482.70, -6.19, 0.72)
        apic[63].pt3dadd(208.58, 483.95, -6.19, 0.72)
        apic[63].pt3dadd(208.58, 484.69, -6.19, 0.72)
        apic[63].pt3dadd(208.10, 485.69, -6.19, 0.72)
        apic[63].pt3dadd(207.86, 487.43, -6.19, 0.72)
        apic[63].pt3dadd(207.62, 487.43, -6.19, 0.72)
        apic[63].pt3dadd(207.13, 489.42, -6.19, 0.72)
        apic[63].pt3dadd(207.13, 489.67, -6.19, 0.72)
        apic[63].pt3dadd(206.89, 491.16, -6.19, 0.72)
        apic[63].pt3dadd(206.65, 491.91, -6.19, 0.72)
        apic[63].pt3dadd(205.93, 494.15, -6.19, 0.72)
        apic[63].pt3dadd(205.69, 494.64, -6.19, 0.72)
        apic[63].pt3dadd(204.96, 496.39, -6.19, 0.72)
        apic[63].pt3dadd(204.72, 496.63, -6.19, 0.72)
        apic[63].pt3dadd(204.24, 498.38, -6.19, 0.72)
        apic[63].pt3dadd(204.24, 500.12, -6.19, 0.72)
        apic[63].pt3dadd(204.48, 500.86, -6.19, 0.72)
        apic[63].pt3dadd(204.96, 502.11, -6.19, 0.72)
        apic[63].pt3dadd(205.93, 503.60, -6.19, 0.72)
        apic[63].pt3dadd(206.41, 504.10, -6.19, 0.72)
        apic[63].pt3dadd(207.13, 504.84, -6.19, 0.72)
        apic[63].pt3dadd(207.86, 505.59, -6.19, 0.72)
        apic[63].pt3dadd(209.31, 507.58, -6.19, 0.72)
        apic[63].pt3dadd(209.31, 508.08, -6.19, 0.72)
        apic[63].pt3dadd(209.55, 508.57, -6.19, 0.72)
        apic[63].pt3dadd(211.00, 511.56, 0.30, 0.72)
        apic[63].pt3dadd(212.21, 514.30, 0.30, 0.72)
        apic[63].pt3dadd(212.21, 514.30, 0.29, 0.72)
        apic[63].pt3dadd(213.41, 515.79, 0.29, 0.72)
        
        apic[64] = h.Section(name='apic[64]')
        apic[64].connect(        apic[63](1), 0)
        apic[64].pt3dclear()
        apic[64].pt3dadd(213.41, 515.79, 0.29, 0.61)
        apic[64].pt3dadd(214.86, 515.29, -7.60, 0.61)
        apic[64].pt3dadd(216.31, 515.29, -7.60, 0.61)
        apic[64].pt3dadd(216.55, 515.29, -7.60, 0.61)
        apic[64].pt3dadd(216.55, 517.03, -7.66, 0.61)
        apic[64].pt3dadd(216.55, 517.28, -7.67, 0.61)
        apic[64].pt3dadd(215.35, 520.27, -7.74, 0.61)
        apic[64].pt3dadd(215.35, 520.27, -7.80, 0.61)
        apic[64].pt3dadd(212.45, 522.26, -8.45, 0.61)
        apic[64].pt3dadd(212.45, 522.26, -8.61, 0.61)
        apic[64].pt3dadd(211.48, 525.49, -8.73, 0.61)
        apic[64].pt3dadd(211.48, 525.49, -8.75, 0.61)
        apic[64].pt3dadd(210.76, 528.72, -8.76, 0.61)
        apic[64].pt3dadd(210.52, 528.72, -8.76, 0.61)
        apic[64].pt3dadd(207.62, 529.72, -8.76, 0.61)
        apic[64].pt3dadd(207.13, 529.72, -8.76, 0.61)
        apic[64].pt3dadd(203.75, 532.70, -8.76, 0.61)
        apic[64].pt3dadd(203.03, 536.19, -8.76, 0.61)
        apic[64].pt3dadd(202.30, 538.67, -8.84, 0.61)
        apic[64].pt3dadd(202.06, 538.67, -8.85, 0.61)
        apic[64].pt3dadd(201.82, 540.91, -15.03, 0.43)
        apic[64].pt3dadd(201.82, 541.16, -15.03, 0.43)
        apic[64].pt3dadd(202.06, 543.15, -15.03, 0.43)
        apic[64].pt3dadd(202.06, 544.15, -15.03, 0.43)
        apic[64].pt3dadd(202.06, 544.40, -15.03, 0.43)
        apic[64].pt3dadd(201.58, 544.89, -15.03, 0.43)
        apic[64].pt3dadd(200.37, 545.64, -15.03, 0.43)
        apic[64].pt3dadd(198.68, 545.39, -15.03, 0.43)
        apic[64].pt3dadd(197.23, 545.64, -15.03, 0.43)
        apic[64].pt3dadd(196.99, 545.89, -15.03, 0.43)
        apic[64].pt3dadd(196.75, 546.88, -15.03, 0.43)
        apic[64].pt3dadd(197.96, 548.62, -15.03, 0.43)
        apic[64].pt3dadd(198.68, 549.62, -15.03, 0.43)
        apic[64].pt3dadd(197.71, 552.85, -16.77, 0.43)
        apic[64].pt3dadd(197.71, 552.85, -16.80, 0.43)
        apic[64].pt3dadd(196.02, 555.34, -16.81, 0.43)
        apic[64].pt3dadd(195.54, 556.83, -16.81, 0.43)
        apic[64].pt3dadd(195.30, 557.33, -16.81, 0.43)
        apic[64].pt3dadd(194.57, 560.07, -16.81, 0.43)
        apic[64].pt3dadd(193.61, 562.06, -16.81, 0.43)
        apic[64].pt3dadd(193.12, 562.55, -16.81, 0.43)
        apic[64].pt3dadd(192.40, 564.05, -16.81, 0.43)
        apic[64].pt3dadd(192.16, 564.30, -16.81, 0.43)
        apic[64].pt3dadd(191.43, 565.54, -16.81, 0.43)
        apic[64].pt3dadd(190.47, 567.53, -16.81, 0.43)
        apic[64].pt3dadd(189.98, 567.78, -16.81, 0.43)
        apic[64].pt3dadd(189.02, 567.78, -16.81, 0.43)
        apic[64].pt3dadd(186.84, 567.53, -16.81, 0.43)
        apic[64].pt3dadd(186.36, 567.28, -16.81, 0.43)
        apic[64].pt3dadd(185.64, 567.28, -16.81, 0.43)
        apic[64].pt3dadd(185.64, 567.53, -16.81, 0.43)
        apic[64].pt3dadd(184.91, 568.52, -16.81, 0.43)
        apic[64].pt3dadd(184.43, 569.52, -16.81, 0.43)
        apic[64].pt3dadd(182.01, 568.52, -16.84, 0.43)
        apic[64].pt3dadd(181.05, 567.53, -18.66, 0.43)
        apic[64].pt3dadd(181.05, 567.53, -18.70, 0.43)
        apic[64].pt3dadd(180.32, 568.28, -18.70, 0.39)
        apic[64].pt3dadd(178.15, 569.02, -18.70, 0.39)
        apic[64].pt3dadd(178.15, 569.27, -18.70, 0.39)
        apic[64].pt3dadd(178.63, 570.76, -23.98, 0.39)
        apic[64].pt3dadd(178.63, 570.76, -23.99, 0.39)
        apic[64].pt3dadd(177.91, 571.76, -23.99, 0.39)
        apic[64].pt3dadd(177.18, 571.26, -23.99, 0.39)
        apic[64].pt3dadd(176.46, 571.01, -23.99, 0.39)
        apic[64].pt3dadd(175.25, 571.01, -23.99, 0.39)
        apic[64].pt3dadd(174.53, 571.01, -23.99, 0.39)
        apic[64].pt3dadd(174.04, 571.01, -23.99, 0.39)
        
        apic[65] = h.Section(name='apic[65]')
        apic[65].connect(        apic[63](1), 0)
        apic[65].pt3dclear()
        apic[65].pt3dadd(213.41, 515.79, 0.29, 0.43)
        apic[65].pt3dadd(212.45, 517.03, -7.49, 0.43)
        apic[65].pt3dadd(213.41, 519.02, -7.49, 0.43)
        apic[65].pt3dadd(215.35, 520.27, -7.49, 0.43)
        apic[65].pt3dadd(216.07, 521.51, -7.49, 0.43)
        apic[65].pt3dadd(216.31, 522.01, -7.49, 0.43)
        apic[65].pt3dadd(216.80, 522.26, -7.49, 0.43)
        apic[65].pt3dadd(217.76, 522.50, -7.49, 0.43)
        apic[65].pt3dadd(218.97, 522.50, -7.49, 0.43)
        apic[65].pt3dadd(219.69, 522.50, -7.49, 0.43)
        apic[65].pt3dadd(220.18, 522.50, -7.49, 0.43)
        apic[65].pt3dadd(221.39, 522.50, -7.49, 0.43)
        apic[65].pt3dadd(222.11, 522.75, -7.49, 0.43)
        apic[65].pt3dadd(222.83, 523.00, -7.49, 0.43)
        apic[65].pt3dadd(224.04, 523.75, -3.60, 0.43)
        apic[65].pt3dadd(223.56, 525.74, -2.87, 0.43)
        apic[65].pt3dadd(223.56, 525.74, -2.83, 0.43)
        apic[65].pt3dadd(221.63, 527.23, -2.18, 0.43)
        apic[65].pt3dadd(221.63, 527.23, -2.14, 0.43)
        apic[65].pt3dadd(221.39, 529.22, -2.11, 0.43)
        apic[65].pt3dadd(221.39, 530.71, -2.10, 0.43)
        apic[65].pt3dadd(222.11, 531.71, -2.09, 0.43)
        apic[65].pt3dadd(223.32, 532.21, -2.08, 0.43)
        apic[65].pt3dadd(225.98, 532.45, -2.07, 0.43)
        apic[65].pt3dadd(226.22, 532.45, -2.07, 0.43)
        apic[65].pt3dadd(227.18, 532.70, -2.07, 0.43)
        apic[65].pt3dadd(227.67, 532.95, -2.07, 0.43)
        apic[65].pt3dadd(227.67, 534.69, 1.57, 0.43)
        apic[65].pt3dadd(227.67, 534.69, 1.66, 0.43)
        apic[65].pt3dadd(227.18, 535.69, 2.26, 0.43)
        apic[65].pt3dadd(227.18, 535.94, 2.49, 0.43)
        apic[65].pt3dadd(226.46, 537.18, 2.89, 0.43)
        apic[65].pt3dadd(226.22, 537.43, 2.97, 0.43)
        apic[65].pt3dadd(225.98, 539.67, 3.17, 0.43)
        apic[65].pt3dadd(225.98, 540.17, 3.24, 0.43)
        apic[65].pt3dadd(225.98, 541.41, 3.29, 0.43)
        apic[65].pt3dadd(225.98, 541.41, 3.31, 0.43)
        apic[65].pt3dadd(226.94, 542.16, 3.32, 0.43)
        apic[65].pt3dadd(227.42, 542.16, 3.33, 0.43)
        apic[65].pt3dadd(229.12, 543.40, 3.37, 0.43)
        apic[65].pt3dadd(229.60, 543.65, 3.37, 0.43)
        apic[65].pt3dadd(230.32, 543.90, 3.37, 0.43)
        apic[65].pt3dadd(230.32, 545.14, 3.38, 0.39)
        apic[65].pt3dadd(230.32, 545.64, 3.38, 0.39)
        apic[65].pt3dadd(229.60, 546.88, 3.38, 0.39)
        apic[65].pt3dadd(229.36, 547.13, 3.38, 0.39)
        apic[65].pt3dadd(229.36, 547.38, 3.39, 0.39)
        apic[65].pt3dadd(228.63, 547.88, 3.43, 0.39)
        apic[65].pt3dadd(228.39, 548.13, 3.44, 0.39)
        apic[65].pt3dadd(227.18, 548.13, 3.45, 0.39)
        apic[65].pt3dadd(226.22, 548.13, 3.45, 0.39)
        apic[65].pt3dadd(225.98, 548.38, 3.45, 0.39)
        apic[65].pt3dadd(225.98, 548.38, 3.83, 0.39)
        apic[65].pt3dadd(225.25, 549.87, 4.39, 0.39)
        apic[65].pt3dadd(225.25, 549.87, 4.42, 0.39)
        apic[65].pt3dadd(226.22, 551.36, 6.16, 0.39)
        apic[65].pt3dadd(226.22, 551.36, 6.38, 0.39)
        apic[65].pt3dadd(227.91, 551.86, 6.29, 0.39)
        apic[65].pt3dadd(227.91, 551.86, 6.35, 0.39)
        apic[65].pt3dadd(228.87, 553.10, 7.18, 0.39)
        apic[65].pt3dadd(228.87, 553.10, 7.36, 0.39)
        apic[65].pt3dadd(230.08, 554.59, 9.90, 0.39)
        apic[65].pt3dadd(230.08, 557.08, 11.75, 0.39)
        apic[65].pt3dadd(230.08, 557.33, 11.74, 0.39)
        apic[65].pt3dadd(231.77, 557.58, 13.18, 0.39)
        apic[65].pt3dadd(231.77, 557.33, 13.18, 0.39)
        apic[65].pt3dadd(233.22, 557.33, 13.24, 0.39)
        apic[65].pt3dadd(233.22, 557.58, 13.25, 0.39)
        apic[65].pt3dadd(232.74, 559.57, 13.27, 0.39)
        apic[65].pt3dadd(232.50, 559.82, 13.33, 0.39)
        apic[65].pt3dadd(231.77, 560.07, 13.38, 0.39)
        apic[65].pt3dadd(231.77, 560.07, 13.42, 0.39)
        apic[65].pt3dadd(230.32, 560.07, 13.44, 0.39)
        apic[65].pt3dadd(229.84, 560.32, 13.45, 0.39)
        apic[65].pt3dadd(229.36, 560.81, 13.45, 0.39)
        apic[65].pt3dadd(230.56, 562.06, 14.64, 0.39)
        apic[65].pt3dadd(230.81, 562.06, 14.72, 0.39)
        apic[65].pt3dadd(232.01, 562.55, 14.59, 0.39)
        apic[65].pt3dadd(232.26, 562.80, 14.64, 0.39)
        apic[65].pt3dadd(234.43, 564.05, 19.18, 0.39)
        apic[65].pt3dadd(234.43, 564.05, 19.20, 0.39)
        apic[65].pt3dadd(235.15, 565.29, 19.33, 0.39)
        apic[65].pt3dadd(235.15, 565.29, 19.59, 0.39)
        apic[65].pt3dadd(235.15, 566.53, 20.14, 0.39)
        apic[65].pt3dadd(235.15, 567.03, 20.15, 0.39)
        apic[65].pt3dadd(234.91, 567.78, 20.13, 0.39)
        apic[65].pt3dadd(234.67, 568.03, 20.34, 0.39)
        apic[65].pt3dadd(234.19, 568.77, 20.88, 0.39)
        apic[65].pt3dadd(233.70, 569.02, 21.08, 0.39)
        apic[65].pt3dadd(232.26, 569.02, 21.82, 0.39)
        apic[65].pt3dadd(231.77, 569.02, 21.90, 0.39)
        apic[65].pt3dadd(231.05, 569.02, 21.94, 0.39)
        apic[65].pt3dadd(230.81, 569.02, 21.94, 0.39)
        
        apic[66] = h.Section(name='apic[66]')
        apic[66].connect(        apic[58](1), 0)
        apic[66].pt3dclear()
        apic[66].pt3dadd(98.92, 368.77, 9.48, 2.70)
        apic[66].pt3dadd(100.61, 370.51, 11.42, 2.70)
        apic[66].pt3dadd(101.82, 372.26, 11.42, 2.70)
        apic[66].pt3dadd(102.06, 372.26, 11.42, 2.70)
        apic[66].pt3dadd(103.51, 373.50, 11.42, 2.70)
        apic[66].pt3dadd(105.69, 375.24, 11.42, 2.70)
        apic[66].pt3dadd(107.62, 376.24, 11.42, 2.70)
        apic[66].pt3dadd(108.10, 376.48, 11.42, 2.70)
        apic[66].pt3dadd(109.79, 377.23, 11.42, 2.70)
        apic[66].pt3dadd(111.24, 378.47, 11.42, 2.70)
        apic[66].pt3dadd(111.72, 378.72, 11.42, 2.70)
        apic[66].pt3dadd(113.41, 379.72, 11.42, 2.70)
        apic[66].pt3dadd(114.38, 381.71, 7.91, 2.44)
        apic[66].pt3dadd(114.14, 383.45, 7.91, 2.44)
        apic[66].pt3dadd(114.14, 383.95, 7.91, 2.44)
        apic[66].pt3dadd(114.38, 384.69, 7.91, 2.44)
        apic[66].pt3dadd(114.14, 387.43, 7.91, 2.44)
        apic[66].pt3dadd(114.14, 388.42, 7.91, 2.44)
        
        apic[67] = h.Section(name='apic[67]')
        apic[67].connect(        apic[66](1), 0)
        apic[67].pt3dclear()
        apic[67].pt3dadd(114.14, 388.42, 7.91, 0.48)
        apic[67].pt3dadd(116.80, 388.42, 12.99, 0.48)
        apic[67].pt3dadd(117.28, 390.17, 12.99, 0.48)
        apic[67].pt3dadd(119.21, 391.91, 12.99, 0.48)
        apic[67].pt3dadd(119.94, 392.65, 12.99, 0.48)
        apic[67].pt3dadd(120.42, 392.90, 12.99, 0.48)
        apic[67].pt3dadd(121.63, 393.40, 12.99, 0.48)
        apic[67].pt3dadd(123.08, 393.40, 12.99, 0.48)
        apic[67].pt3dadd(124.04, 393.15, 12.99, 0.48)
        apic[67].pt3dadd(125.01, 393.15, 12.99, 0.48)
        apic[67].pt3dadd(125.73, 393.40, 12.99, 0.48)
        apic[67].pt3dadd(126.70, 398.13, 12.99, 0.48)
        apic[67].pt3dadd(127.67, 398.13, 12.99, 0.48)
        apic[67].pt3dadd(128.63, 398.62, 12.99, 0.48)
        apic[67].pt3dadd(129.36, 398.62, 12.99, 0.48)
        apic[67].pt3dadd(129.60, 398.87, 13.76, 0.48)
        apic[67].pt3dadd(129.84, 398.87, 14.33, 0.48)
        apic[67].pt3dadd(130.08, 399.12, 14.48, 0.48)
        apic[67].pt3dadd(131.53, 399.62, 14.72, 0.48)
        apic[67].pt3dadd(132.01, 399.87, 14.85, 0.48)
        apic[67].pt3dadd(134.43, 399.87, 15.13, 0.48)
        apic[67].pt3dadd(134.91, 399.87, 15.23, 0.48)
        apic[67].pt3dadd(135.64, 400.12, 15.46, 0.48)
        apic[67].pt3dadd(135.88, 400.12, 15.71, 0.48)
        apic[67].pt3dadd(136.84, 400.12, 15.83, 0.48)
        apic[67].pt3dadd(139.02, 400.61, 16.44, 0.48)
        apic[67].pt3dadd(139.98, 400.86, 16.50, 0.48)
        apic[67].pt3dadd(140.95, 400.86, 16.70, 0.48)
        
        apic[68] = h.Section(name='apic[68]')
        apic[68].connect(        apic[66](1), 0)
        apic[68].pt3dclear()
        apic[68].pt3dadd(114.14, 388.42, 7.91, 2.20)
        apic[68].pt3dadd(112.93, 389.42, 15.82, 2.20)
        apic[68].pt3dadd(111.48, 391.66, 15.81, 2.20)
        apic[68].pt3dadd(110.03, 393.65, 15.81, 2.20)
        apic[68].pt3dadd(106.89, 395.64, 15.80, 2.20)
        apic[68].pt3dadd(106.41, 396.14, 15.79, 2.20)
        apic[68].pt3dadd(104.48, 397.88, 15.78, 1.90)
        apic[68].pt3dadd(102.79, 400.86, 15.78, 1.90)
        apic[68].pt3dadd(102.30, 403.60, 15.78, 1.90)
        apic[68].pt3dadd(101.10, 405.34, 15.78, 1.90)
        apic[68].pt3dadd(100.61, 405.59, 15.78, 1.90)
        apic[68].pt3dadd(98.20, 406.58, 15.78, 1.90)
        apic[68].pt3dadd(97.96, 406.58, 15.78, 1.90)
        apic[68].pt3dadd(96.02, 407.58, 15.78, 1.90)
        apic[68].pt3dadd(95.54, 407.83, 15.78, 1.90)
        apic[68].pt3dadd(93.85, 409.32, 15.78, 1.90)
        apic[68].pt3dadd(93.12, 410.56, 15.78, 1.90)
        apic[68].pt3dadd(93.12, 410.81, 15.78, 1.90)
        apic[68].pt3dadd(92.16, 411.81, 15.78, 1.70)
        apic[68].pt3dadd(91.92, 412.31, 15.78, 1.70)
        apic[68].pt3dadd(89.50, 416.78, 15.78, 1.70)
        apic[68].pt3dadd(88.29, 418.03, 15.78, 1.70)
        apic[68].pt3dadd(88.05, 418.77, 15.78, 1.70)
        apic[68].pt3dadd(88.54, 424.25, 15.78, 1.70)
        apic[68].pt3dadd(88.29, 426.48, 15.78, 1.70)
        apic[68].pt3dadd(89.98, 429.22, 7.16, 1.70)
        apic[68].pt3dadd(91.19, 430.71, 7.16, 1.70)
        apic[68].pt3dadd(92.64, 434.44, 7.16, 1.70)
        apic[68].pt3dadd(92.40, 437.43, 7.16, 1.70)
        apic[68].pt3dadd(92.16, 437.93, 7.16, 1.70)
        apic[68].pt3dadd(90.95, 440.17, 7.16, 1.70)
        apic[68].pt3dadd(90.95, 440.66, 7.16, 1.70)
        apic[68].pt3dadd(89.74, 444.15, 7.51, 1.70)
        apic[68].pt3dadd(88.54, 446.63, 7.51, 1.70)
        apic[68].pt3dadd(88.54, 446.88, 7.51, 1.70)
        apic[68].pt3dadd(87.57, 449.12, 7.51, 1.70)
        apic[68].pt3dadd(87.57, 449.37, 7.51, 1.70)
        apic[68].pt3dadd(86.60, 451.86, 7.51, 1.70)
        apic[68].pt3dadd(86.60, 452.36, 7.51, 1.70)
        apic[68].pt3dadd(85.15, 455.09, 7.51, 2.20)
        apic[68].pt3dadd(83.70, 458.08, 7.51, 2.20)
        apic[68].pt3dadd(82.26, 459.82, 7.51, 2.20)
        apic[68].pt3dadd(80.56, 461.56, 7.51, 2.20)
        apic[68].pt3dadd(80.32, 462.31, 7.51, 2.20)
        apic[68].pt3dadd(80.56, 463.55, 7.51, 2.20)
        
        apic[69] = h.Section(name='apic[69]')
        apic[69].connect(        apic[68](1), 0)
        apic[69].pt3dclear()
        apic[69].pt3dadd(80.56, 463.55, 7.51, 1.40)
        apic[69].pt3dadd(81.53, 466.53, 8.20, 1.40)
        apic[69].pt3dadd(82.50, 468.52, 8.20, 1.40)
        apic[69].pt3dadd(82.74, 468.52, 8.20, 1.40)
        apic[69].pt3dadd(84.19, 469.27, 8.20, 1.40)
        apic[69].pt3dadd(85.88, 469.52, 8.20, 1.40)
        apic[69].pt3dadd(88.05, 470.02, 4.27, 1.40)
        apic[69].pt3dadd(88.29, 472.50, 4.27, 1.40)
        apic[69].pt3dadd(88.54, 472.50, 4.27, 1.40)
        apic[69].pt3dadd(89.02, 473.50, 4.27, 1.40)
        apic[69].pt3dadd(89.50, 473.75, 4.27, 1.40)
        apic[69].pt3dadd(90.47, 474.74, 4.27, 1.40)
        apic[69].pt3dadd(91.19, 475.49, 4.27, 1.40)
        apic[69].pt3dadd(91.43, 476.48, 4.27, 1.40)
        apic[69].pt3dadd(91.43, 476.73, 4.27, 1.40)
        apic[69].pt3dadd(92.40, 477.48, 4.27, 1.10)
        apic[69].pt3dadd(92.64, 477.73, 4.27, 1.10)
        apic[69].pt3dadd(93.61, 478.47, 4.27, 1.10)
        apic[69].pt3dadd(93.85, 478.47, 4.27, 1.10)
        apic[69].pt3dadd(95.78, 480.96, 4.92, 1.10)
        apic[69].pt3dadd(96.51, 482.21, 4.92, 1.10)
        apic[69].pt3dadd(97.23, 483.45, 4.91, 1.10)
        apic[69].pt3dadd(97.71, 484.20, 5.16, 1.10)
        apic[69].pt3dadd(97.71, 484.44, 5.16, 1.10)
        apic[69].pt3dadd(97.71, 485.44, 5.16, 1.10)
        apic[69].pt3dadd(96.02, 485.44, 5.16, 1.10)
        apic[69].pt3dadd(96.02, 485.69, 5.16, 1.10)
        apic[69].pt3dadd(95.54, 487.68, 5.16, 1.10)
        apic[69].pt3dadd(94.33, 489.17, 5.16, 1.10)
        apic[69].pt3dadd(94.09, 490.91, 5.16, 1.10)
        apic[69].pt3dadd(94.09, 492.65, 5.16, 1.10)
        apic[69].pt3dadd(94.09, 493.15, 5.16, 1.10)
        apic[69].pt3dadd(94.33, 494.15, 5.16, 1.10)
        apic[69].pt3dadd(94.33, 495.64, 5.16, 1.10)
        apic[69].pt3dadd(94.33, 495.89, 5.16, 1.10)
        apic[69].pt3dadd(95.06, 497.13, 5.16, 1.10)
        apic[69].pt3dadd(95.30, 497.38, 5.16, 1.10)
        apic[69].pt3dadd(96.02, 498.13, 5.16, 1.10)
        apic[69].pt3dadd(96.51, 499.12, 5.16, 1.10)
        apic[69].pt3dadd(96.75, 499.12, 5.15, 1.10)
        apic[69].pt3dadd(97.23, 500.12, 5.15, 1.10)
        apic[69].pt3dadd(97.47, 500.37, 5.14, 1.10)
        apic[69].pt3dadd(97.96, 503.10, 1.73, 0.77)
        apic[69].pt3dadd(98.20, 503.85, 1.73, 0.77)
        apic[69].pt3dadd(98.44, 504.35, 1.73, 0.77)
        apic[69].pt3dadd(98.92, 505.34, 1.73, 0.77)
        apic[69].pt3dadd(99.89, 506.34, 1.73, 0.77)
        apic[69].pt3dadd(100.37, 507.08, 1.73, 0.77)
        apic[69].pt3dadd(100.85, 509.07, 1.73, 0.77)
        apic[69].pt3dadd(101.34, 511.31, 0.40, 0.77)
        apic[69].pt3dadd(101.82, 512.06, 0.40, 0.77)
        apic[69].pt3dadd(102.06, 512.31, 0.40, 0.77)
        apic[69].pt3dadd(102.79, 513.05, 0.40, 0.77)
        apic[69].pt3dadd(103.27, 513.80, 0.40, 0.77)
        
        apic[70] = h.Section(name='apic[70]')
        apic[70].connect(        apic[69](1), 0)
        apic[70].pt3dclear()
        apic[70].pt3dadd(103.27, 513.80, 0.40, 0.54)
        apic[70].pt3dadd(103.99, 515.54, -9.55, 0.54)
        apic[70].pt3dadd(104.24, 515.54, -9.56, 0.54)
        apic[70].pt3dadd(105.69, 516.04, -9.57, 0.54)
        apic[70].pt3dadd(106.17, 516.53, -9.59, 0.54)
        apic[70].pt3dadd(104.24, 517.78, -9.59, 0.54)
        apic[70].pt3dadd(103.51, 518.03, -9.61, 0.54)
        apic[70].pt3dadd(103.51, 518.52, -9.63, 0.54)
        apic[70].pt3dadd(103.51, 519.02, -9.63, 0.54)
        apic[70].pt3dadd(103.75, 519.27, -9.63, 0.54)
        apic[70].pt3dadd(104.72, 519.77, -9.63, 0.54)
        apic[70].pt3dadd(104.96, 520.02, -9.63, 0.54)
        apic[70].pt3dadd(105.93, 520.51, -9.64, 0.54)
        apic[70].pt3dadd(107.13, 520.76, -9.64, 0.54)
        apic[70].pt3dadd(107.86, 521.26, -9.64, 0.54)
        apic[70].pt3dadd(107.86, 522.50, -9.64, 0.54)
        apic[70].pt3dadd(107.86, 522.50, -9.66, 0.54)
        apic[70].pt3dadd(106.89, 524.25, -12.46, 0.54)
        apic[70].pt3dadd(106.89, 524.25, -12.52, 0.54)
        apic[70].pt3dadd(106.17, 525.74, -12.56, 0.54)
        apic[70].pt3dadd(105.93, 525.74, -12.99, 0.54)
        apic[70].pt3dadd(105.44, 527.23, -13.03, 0.54)
        apic[70].pt3dadd(105.44, 527.23, -13.04, 0.54)
        apic[70].pt3dadd(106.17, 528.23, -13.04, 0.54)
        apic[70].pt3dadd(106.65, 528.72, -13.05, 0.54)
        apic[70].pt3dadd(107.13, 529.97, -13.05, 0.54)
        apic[70].pt3dadd(107.62, 530.46, -13.05, 0.54)
        apic[70].pt3dadd(108.34, 531.46, -13.05, 0.54)
        apic[70].pt3dadd(108.34, 531.71, -13.05, 0.54)
        apic[70].pt3dadd(108.83, 532.45, -13.05, 0.54)
        apic[70].pt3dadd(109.07, 532.95, -13.05, 0.54)
        apic[70].pt3dadd(109.07, 533.45, -13.05, 0.43)
        apic[70].pt3dadd(108.83, 535.44, -13.05, 0.43)
        apic[70].pt3dadd(108.34, 536.43, -18.64, 0.43)
        apic[70].pt3dadd(109.31, 536.93, -19.20, 0.43)
        apic[70].pt3dadd(110.03, 537.18, -19.20, 0.43)
        apic[70].pt3dadd(112.69, 538.18, -19.20, 0.43)
        apic[70].pt3dadd(111.48, 541.16, -19.21, 0.43)
        apic[70].pt3dadd(111.48, 541.16, -19.23, 0.43)
        apic[70].pt3dadd(111.72, 542.40, -25.27, 0.43)
        apic[70].pt3dadd(113.17, 543.15, -25.27, 0.43)
        apic[70].pt3dadd(114.14, 544.15, -25.27, 0.43)
        apic[70].pt3dadd(113.90, 545.39, -25.27, 0.43)
        apic[70].pt3dadd(113.90, 545.64, -25.27, 0.43)
        apic[70].pt3dadd(114.38, 547.88, -25.46, 0.43)
        apic[70].pt3dadd(114.86, 548.13, -25.56, 0.43)
        apic[70].pt3dadd(117.04, 547.13, -27.43, 0.43)
        apic[70].pt3dadd(117.52, 547.13, -27.46, 0.43)
        apic[70].pt3dadd(119.45, 547.88, -29.02, 0.43)
        apic[70].pt3dadd(119.45, 547.88, -29.19, 0.43)
        apic[70].pt3dadd(119.69, 550.12, -29.68, 0.43)
        apic[70].pt3dadd(119.69, 550.37, -30.10, 0.43)
        apic[70].pt3dadd(119.69, 551.61, -30.19, 0.43)
        apic[70].pt3dadd(118.73, 552.85, -30.18, 0.43)
        apic[70].pt3dadd(118.25, 553.35, -30.18, 0.43)
        apic[70].pt3dadd(118.00, 554.35, -30.18, 0.43)
        apic[70].pt3dadd(118.97, 555.59, -30.18, 0.43)
        apic[70].pt3dadd(119.21, 555.59, -30.18, 0.43)
        apic[70].pt3dadd(119.94, 556.34, -30.18, 0.43)
        apic[70].pt3dadd(121.14, 557.83, -30.89, 0.43)
        apic[70].pt3dadd(121.14, 558.82, -35.34, 0.43)
        apic[70].pt3dadd(119.21, 559.57, -35.34, 0.43)
        apic[70].pt3dadd(119.21, 559.82, -35.34, 0.43)
        apic[70].pt3dadd(118.97, 562.31, -35.39, 0.43)
        apic[70].pt3dadd(119.21, 562.31, -35.39, 0.43)
        apic[70].pt3dadd(119.21, 564.30, -35.39, 0.43)
        apic[70].pt3dadd(119.21, 564.79, -35.39, 0.43)
        apic[70].pt3dadd(119.21, 565.79, -35.39, 0.43)
        apic[70].pt3dadd(119.94, 567.28, -35.39, 0.43)
        apic[70].pt3dadd(120.18, 567.53, -35.39, 0.43)
        apic[70].pt3dadd(120.42, 568.03, -35.39, 0.43)
        apic[70].pt3dadd(121.39, 568.77, -35.39, 0.43)
        apic[70].pt3dadd(121.87, 569.52, -35.39, 0.43)
        apic[70].pt3dadd(122.35, 569.77, -35.39, 0.43)
        apic[70].pt3dadd(124.77, 572.01, -33.50, 0.43)
        apic[70].pt3dadd(125.25, 572.01, -33.50, 0.43)
        apic[70].pt3dadd(126.46, 573.50, -33.51, 0.43)
        apic[70].pt3dadd(126.46, 573.75, -33.52, 0.43)
        apic[70].pt3dadd(125.73, 574.99, -33.53, 0.43)
        apic[70].pt3dadd(125.49, 575.24, -33.53, 0.39)
        apic[70].pt3dadd(124.77, 576.98, -33.57, 0.39)
        apic[70].pt3dadd(124.28, 577.48, -33.59, 0.39)
        apic[70].pt3dadd(123.08, 578.97, -33.83, 0.39)
        apic[70].pt3dadd(123.08, 578.97, -33.87, 0.39)
        apic[70].pt3dadd(122.11, 580.96, -40.61, 0.39)
        apic[70].pt3dadd(122.11, 580.96, -40.68, 0.39)
        apic[70].pt3dadd(120.66, 582.95, -43.43, 0.39)
        apic[70].pt3dadd(120.66, 582.95, -43.64, 0.39)
        apic[70].pt3dadd(120.42, 583.70, -44.47, 0.39)
        apic[70].pt3dadd(120.18, 583.70, -44.79, 0.39)
        apic[70].pt3dadd(119.21, 584.94, -45.67, 0.39)
        apic[70].pt3dadd(119.21, 585.19, -45.81, 0.39)
        apic[70].pt3dadd(118.97, 586.68, -45.94, 0.39)
        apic[70].pt3dadd(118.73, 586.68, -45.96, 0.39)
        apic[70].pt3dadd(117.76, 587.18, -46.23, 0.39)
        apic[70].pt3dadd(117.76, 587.18, -46.26, 0.39)
        apic[70].pt3dadd(115.11, 586.68, -46.37, 0.39)
        apic[70].pt3dadd(115.35, 586.68, -46.38, 0.39)
        apic[70].pt3dadd(116.31, 589.17, -46.82, 0.39)
        apic[70].pt3dadd(116.31, 589.67, -47.08, 0.39)
        apic[70].pt3dadd(116.31, 590.41, -47.57, 0.39)
        apic[70].pt3dadd(116.07, 590.41, -47.69, 0.39)
        apic[70].pt3dadd(115.35, 590.41, -47.97, 0.39)
        apic[70].pt3dadd(115.35, 590.41, -48.17, 0.39)
        apic[70].pt3dadd(110.76, 589.92, -53.76, 0.39)
        apic[70].pt3dadd(110.27, 589.92, -53.78, 0.39)
        apic[70].pt3dadd(109.07, 594.40, -53.87, 0.39)
        apic[70].pt3dadd(109.07, 594.40, -53.88, 0.39)
        apic[70].pt3dadd(108.34, 594.40, -53.89, 0.39)
        apic[70].pt3dadd(107.38, 595.89, -54.38, 0.39)
        apic[70].pt3dadd(107.38, 595.89, -54.44, 0.39)
        apic[70].pt3dadd(106.65, 597.13, -54.53, 0.39)
        apic[70].pt3dadd(106.41, 597.38, -54.55, 0.39)
        
        apic[71] = h.Section(name='apic[71]')
        apic[71].connect(        apic[69](1), 0)
        apic[71].pt3dclear()
        apic[71].pt3dadd(103.27, 513.80, 0.40, 0.61)
        apic[71].pt3dadd(103.27, 517.28, 3.97, 0.61)
        apic[71].pt3dadd(103.27, 521.01, 3.98, 0.61)
        apic[71].pt3dadd(103.27, 522.75, 4.01, 0.61)
        apic[71].pt3dadd(102.55, 525.49, 0.65, 0.61)
        apic[71].pt3dadd(102.55, 525.49, 0.64, 0.61)
        apic[71].pt3dadd(102.55, 527.48, 0.59, 0.61)
        apic[71].pt3dadd(102.55, 527.48, 0.56, 0.61)
        apic[71].pt3dadd(102.55, 528.97, 0.30, 0.61)
        apic[71].pt3dadd(102.55, 529.22, 0.24, 0.61)
        apic[71].pt3dadd(102.55, 530.71, 0.22, 0.61)
        apic[71].pt3dadd(103.03, 531.46, 0.21, 0.61)
        apic[71].pt3dadd(103.51, 532.45, 0.20, 0.61)
        apic[71].pt3dadd(103.51, 532.45, 0.19, 0.61)
        apic[71].pt3dadd(103.75, 533.70, 0.18, 0.61)
        apic[71].pt3dadd(103.99, 533.95, 0.18, 0.61)
        apic[71].pt3dadd(103.99, 534.94, 0.18, 0.61)
        
        apic[72] = h.Section(name='apic[72]')
        apic[72].connect(        apic[71](1), 0)
        apic[72].pt3dclear()
        apic[72].pt3dadd(103.99, 534.94, 0.18, 0.54)
        apic[72].pt3dadd(104.72, 538.92, 0.13, 0.54)
        apic[72].pt3dadd(105.93, 540.17, 0.13, 0.54)
        apic[72].pt3dadd(106.17, 540.17, 0.13, 0.54)
        apic[72].pt3dadd(107.38, 541.66, 0.13, 0.54)
        apic[72].pt3dadd(108.10, 542.16, 0.13, 0.54)
        apic[72].pt3dadd(109.55, 543.15, 0.13, 0.54)
        apic[72].pt3dadd(110.03, 543.40, 0.13, 0.54)
        apic[72].pt3dadd(111.00, 544.39, 0.13, 0.54)
        apic[72].pt3dadd(111.24, 544.89, 0.13, 0.54)
        apic[72].pt3dadd(111.72, 546.88, 0.13, 0.54)
        apic[72].pt3dadd(111.72, 548.62, 0.13, 0.54)
        apic[72].pt3dadd(111.72, 550.37, 0.13, 0.54)
        apic[72].pt3dadd(111.72, 552.36, 0.13, 0.54)
        apic[72].pt3dadd(111.72, 553.10, 0.13, 0.54)
        apic[72].pt3dadd(111.72, 555.09, 0.13, 0.54)
        apic[72].pt3dadd(111.72, 556.09, 0.13, 0.54)
        apic[72].pt3dadd(112.45, 557.58, 0.13, 0.54)
        apic[72].pt3dadd(112.69, 557.58, 0.12, 0.54)
        apic[72].pt3dadd(114.38, 558.08, 0.12, 0.54)
        apic[72].pt3dadd(118.00, 558.33, 0.12, 0.54)
        apic[72].pt3dadd(119.69, 558.33, 0.12, 0.54)
        apic[72].pt3dadd(120.42, 558.33, 0.12, 0.54)
        apic[72].pt3dadd(120.66, 558.33, 0.12, 0.54)
        apic[72].pt3dadd(121.87, 558.57, 0.12, 0.54)
        apic[72].pt3dadd(122.35, 558.82, 0.12, 0.54)
        apic[72].pt3dadd(123.08, 558.82, 0.12, 0.54)
        apic[72].pt3dadd(123.32, 559.32, 0.12, 0.54)
        apic[72].pt3dadd(124.04, 560.07, 0.12, 0.54)
        apic[72].pt3dadd(125.25, 560.81, 0.12, 0.54)
        apic[72].pt3dadd(125.97, 561.81, 0.12, 0.54)
        apic[72].pt3dadd(126.70, 563.55, 3.21, 0.54)
        apic[72].pt3dadd(126.70, 565.54, 3.21, 0.54)
        apic[72].pt3dadd(126.70, 566.29, 3.21, 0.54)
        apic[72].pt3dadd(126.22, 568.28, 3.20, 0.54)
        apic[72].pt3dadd(126.22, 570.27, 3.20, 0.54)
        apic[72].pt3dadd(125.97, 571.26, 3.20, 0.54)
        apic[72].pt3dadd(127.91, 571.51, 3.20, 0.54)
        apic[72].pt3dadd(128.87, 571.51, 3.20, 0.54)
        apic[72].pt3dadd(130.56, 572.75, 3.28, 0.54)
        apic[72].pt3dadd(130.81, 574.99, 3.28, 0.54)
        apic[72].pt3dadd(130.81, 575.49, 3.28, 0.54)
        apic[72].pt3dadd(132.26, 577.98, 10.70, 0.54)
        apic[72].pt3dadd(134.43, 578.97, 10.69, 0.54)
        apic[72].pt3dadd(134.91, 579.22, 10.69, 0.54)
        apic[72].pt3dadd(136.12, 580.96, 10.69, 0.54)
        apic[72].pt3dadd(136.36, 581.21, 10.69, 0.54)
        apic[72].pt3dadd(136.84, 582.95, 10.69, 0.54)
        apic[72].pt3dadd(136.84, 583.45, 10.69, 0.54)
        apic[72].pt3dadd(136.84, 585.19, 10.69, 0.54)
        apic[72].pt3dadd(136.84, 585.44, 10.69, 0.54)
        apic[72].pt3dadd(136.84, 587.18, 10.69, 0.54)
        apic[72].pt3dadd(136.84, 587.68, 10.68, 0.54)
        apic[72].pt3dadd(137.81, 590.41, 10.68, 0.54)
        apic[72].pt3dadd(139.02, 591.16, 10.68, 0.54)
        apic[72].pt3dadd(139.98, 591.16, 10.68, 0.54)
        apic[72].pt3dadd(140.95, 591.16, 10.68, 0.43)
        apic[72].pt3dadd(141.68, 589.92, 10.68, 0.43)
        apic[72].pt3dadd(142.64, 589.42, 10.68, 0.43)
        apic[72].pt3dadd(143.61, 589.42, 10.68, 0.43)
        apic[72].pt3dadd(145.06, 589.67, 10.68, 0.43)
        apic[72].pt3dadd(145.78, 589.92, 10.68, 0.43)
        apic[72].pt3dadd(146.26, 589.92, 10.68, 0.43)
        apic[72].pt3dadd(148.68, 587.18, 10.68, 0.43)
        apic[72].pt3dadd(153.03, 585.69, 6.05, 0.43)
        apic[72].pt3dadd(153.03, 585.69, 6.07, 0.43)
        apic[72].pt3dadd(155.93, 584.20, 6.09, 0.43)
        apic[72].pt3dadd(156.41, 584.20, 6.09, 0.43)
        apic[72].pt3dadd(158.83, 582.95, 6.09, 0.43)
        apic[72].pt3dadd(159.07, 582.95, 6.09, 0.43)
        apic[72].pt3dadd(160.27, 582.21, 6.09, 0.43)
        apic[72].pt3dadd(160.52, 582.21, 6.09, 0.43)
        apic[72].pt3dadd(164.14, 584.20, 6.09, 0.39)
        apic[72].pt3dadd(164.86, 585.44, 6.09, 0.39)
        apic[72].pt3dadd(165.59, 585.44, 6.09, 0.39)
        apic[72].pt3dadd(168.73, 585.19, 6.09, 0.39)
        apic[72].pt3dadd(169.45, 584.69, 6.09, 0.39)
        apic[72].pt3dadd(170.66, 584.69, 6.09, 0.39)
        apic[72].pt3dadd(171.39, 584.69, 6.09, 0.39)
        apic[72].pt3dadd(173.32, 584.69, 6.09, 0.39)
        apic[72].pt3dadd(174.28, 584.69, 6.09, 0.39)
        apic[72].pt3dadd(175.49, 585.44, 6.09, 0.39)
        apic[72].pt3dadd(176.22, 586.93, 6.09, 0.39)
        apic[72].pt3dadd(176.70, 587.43, 6.09, 0.39)
        apic[72].pt3dadd(177.42, 589.42, 6.09, 0.39)
        apic[72].pt3dadd(178.39, 590.17, 6.09, 0.39)
        apic[72].pt3dadd(179.36, 590.41, 6.09, 0.39)
        apic[72].pt3dadd(179.60, 590.41, 6.09, 0.39)
        apic[72].pt3dadd(180.56, 590.41, 6.09, 0.39)
        apic[72].pt3dadd(181.53, 590.41, 6.09, 0.39)
        
        apic[73] = h.Section(name='apic[73]')
        apic[73].connect(        apic[71](1), 0)
        apic[73].pt3dclear()
        apic[73].pt3dadd(103.99, 534.94, 0.18, 0.43)
        apic[73].pt3dadd(104.48, 540.91, -6.73, 0.43)
        apic[73].pt3dadd(105.93, 543.40, -6.73, 0.43)
        apic[73].pt3dadd(105.93, 545.39, -6.73, 0.43)
        apic[73].pt3dadd(105.69, 545.64, -6.73, 0.43)
        apic[73].pt3dadd(105.44, 546.14, -6.73, 0.43)
        apic[73].pt3dadd(105.20, 546.63, -6.73, 0.43)
        apic[73].pt3dadd(105.20, 547.38, -6.73, 0.43)
        apic[73].pt3dadd(105.20, 548.13, -6.73, 0.43)
        apic[73].pt3dadd(105.69, 549.12, -6.73, 0.43)
        apic[73].pt3dadd(105.93, 549.62, -6.73, 0.43)
        apic[73].pt3dadd(106.17, 550.37, -6.73, 0.43)
        apic[73].pt3dadd(106.41, 550.86, -6.73, 0.43)
        apic[73].pt3dadd(107.13, 552.60, -6.73, 0.43)
        apic[73].pt3dadd(107.13, 553.10, -6.73, 0.43)
        apic[73].pt3dadd(106.89, 554.35, -6.73, 0.43)
        apic[73].pt3dadd(106.65, 554.59, -6.73, 0.43)
        apic[73].pt3dadd(106.89, 556.09, -9.45, 0.43)
        apic[73].pt3dadd(105.44, 558.33, -9.45, 0.43)
        apic[73].pt3dadd(104.96, 558.82, -9.45, 0.43)
        apic[73].pt3dadd(103.75, 560.56, -9.45, 0.43)
        apic[73].pt3dadd(103.75, 561.06, -9.45, 0.43)
        apic[73].pt3dadd(103.99, 563.55, -9.45, 0.43)
        apic[73].pt3dadd(105.93, 563.55, -9.45, 0.43)
        apic[73].pt3dadd(106.17, 563.05, -9.45, 0.43)
        apic[73].pt3dadd(106.89, 562.55, -9.45, 0.43)
        apic[73].pt3dadd(107.38, 562.31, -9.45, 0.43)
        apic[73].pt3dadd(108.34, 562.06, -9.45, 0.43)
        apic[73].pt3dadd(109.07, 562.06, -9.45, 0.43)
        apic[73].pt3dadd(109.55, 562.31, -9.45, 0.43)
        apic[73].pt3dadd(110.03, 562.80, -9.45, 0.43)
        apic[73].pt3dadd(110.27, 564.05, -9.45, 0.43)
        apic[73].pt3dadd(111.00, 565.79, -13.62, 0.43)
        apic[73].pt3dadd(111.48, 565.79, -13.62, 0.43)
        apic[73].pt3dadd(113.66, 566.29, -13.62, 0.43)
        apic[73].pt3dadd(113.66, 567.78, -13.62, 0.43)
        apic[73].pt3dadd(112.93, 569.77, -17.75, 0.43)
        apic[73].pt3dadd(113.17, 571.01, -17.75, 0.43)
        apic[73].pt3dadd(113.90, 571.01, -17.75, 0.43)
        apic[73].pt3dadd(116.07, 571.26, -17.75, 0.43)
        apic[73].pt3dadd(116.55, 571.26, -17.74, 0.43)
        apic[73].pt3dadd(118.49, 571.76, -17.74, 0.43)
        apic[73].pt3dadd(118.97, 572.01, -17.74, 0.43)
        apic[73].pt3dadd(119.69, 572.75, -24.88, 0.43)
        apic[73].pt3dadd(119.69, 574.00, -24.88, 0.43)
        apic[73].pt3dadd(119.69, 574.25, -24.88, 0.43)
        apic[73].pt3dadd(120.18, 574.99, -24.88, 0.43)
        apic[73].pt3dadd(121.14, 575.74, -24.88, 0.43)
        apic[73].pt3dadd(121.63, 576.73, -24.88, 0.43)
        apic[73].pt3dadd(122.11, 576.98, -24.88, 0.43)
        apic[73].pt3dadd(123.08, 577.48, -24.88, 0.43)
        apic[73].pt3dadd(123.80, 577.73, -24.88, 0.43)
        apic[73].pt3dadd(124.28, 578.47, -24.88, 0.43)
        apic[73].pt3dadd(124.77, 578.72, -24.88, 0.43)
        apic[73].pt3dadd(125.25, 580.22, -24.88, 0.43)
        apic[73].pt3dadd(125.49, 580.46, -24.88, 0.43)
        apic[73].pt3dadd(125.73, 581.46, -24.88, 0.43)
        apic[73].pt3dadd(125.73, 581.71, -24.88, 0.43)
        apic[73].pt3dadd(125.73, 582.45, -24.88, 0.43)
        apic[73].pt3dadd(125.73, 583.20, -24.88, 0.43)
        apic[73].pt3dadd(125.73, 584.94, -24.88, 0.43)
        apic[73].pt3dadd(127.67, 585.94, -24.88, 0.43)
        apic[73].pt3dadd(128.63, 586.43, -24.88, 0.43)
        apic[73].pt3dadd(131.29, 586.93, -28.82, 0.43)
        apic[73].pt3dadd(130.56, 590.17, -28.82, 0.43)
        apic[73].pt3dadd(131.29, 591.66, -28.82, 0.43)
        apic[73].pt3dadd(131.77, 591.91, -28.82, 0.43)
        apic[73].pt3dadd(132.26, 593.90, -28.82, 0.43)
        
        apic[74] = h.Section(name='apic[74]')
        apic[74].connect(        apic[68](1), 0)
        apic[74].pt3dclear()
        apic[74].pt3dadd(80.56, 463.55, 7.51, 0.72)
        apic[74].pt3dadd(78.87, 464.54, 13.27, 0.72)
        apic[74].pt3dadd(77.42, 466.53, 13.27, 0.72)
        apic[74].pt3dadd(77.18, 468.52, 13.27, 0.72)
        apic[74].pt3dadd(76.46, 470.27, 13.27, 0.72)
        apic[74].pt3dadd(76.22, 470.27, 13.27, 0.72)
        apic[74].pt3dadd(75.01, 471.51, 13.27, 0.72)
        apic[74].pt3dadd(74.53, 473.25, 13.27, 0.72)
        apic[74].pt3dadd(74.04, 474.25, 13.27, 0.72)
        apic[74].pt3dadd(74.04, 474.49, 13.27, 0.72)
        apic[74].pt3dadd(73.80, 476.48, 13.26, 0.72)
        apic[74].pt3dadd(73.80, 478.97, 13.26, 0.72)
        apic[74].pt3dadd(73.56, 479.47, 13.26, 0.72)
        apic[74].pt3dadd(73.56, 479.72, 13.26, 0.72)
        
        apic[75] = h.Section(name='apic[75]')
        apic[75].connect(        apic[74](1), 0)
        apic[75].pt3dclear()
        apic[75].pt3dadd(73.56, 479.72, 13.26, 0.43)
        apic[75].pt3dadd(72.59, 482.21, 12.05, 0.43)
        apic[75].pt3dadd(71.87, 483.45, 12.05, 0.43)
        apic[75].pt3dadd(72.83, 484.94, 12.05, 0.43)
        apic[75].pt3dadd(72.83, 485.19, 12.05, 0.43)
        apic[75].pt3dadd(72.83, 487.18, 12.05, 0.43)
        apic[75].pt3dadd(72.35, 487.18, 12.05, 0.43)
        apic[75].pt3dadd(70.90, 487.93, 12.05, 0.43)
        apic[75].pt3dadd(70.66, 488.18, 12.05, 0.43)
        apic[75].pt3dadd(69.69, 488.92, 12.05, 0.43)
        apic[75].pt3dadd(69.45, 489.42, 12.05, 0.43)
        apic[75].pt3dadd(69.21, 491.41, 12.05, 0.43)
        apic[75].pt3dadd(70.18, 492.16, 15.28, 0.43)
        apic[75].pt3dadd(67.76, 492.90, 15.28, 0.43)
        apic[75].pt3dadd(67.76, 494.40, 15.28, 0.43)
        apic[75].pt3dadd(67.76, 494.64, 15.28, 0.43)
        apic[75].pt3dadd(68.49, 495.64, 15.28, 0.43)
        apic[75].pt3dadd(68.97, 496.39, 15.28, 0.43)
        apic[75].pt3dadd(69.45, 497.63, 15.33, 0.43)
        apic[75].pt3dadd(69.45, 498.38, 15.34, 0.43)
        apic[75].pt3dadd(69.45, 499.37, 15.50, 0.43)
        apic[75].pt3dadd(69.21, 499.62, 15.52, 0.43)
        apic[75].pt3dadd(68.49, 500.61, 17.03, 0.43)
        apic[75].pt3dadd(67.04, 502.11, 17.24, 0.43)
        apic[75].pt3dadd(67.04, 502.11, 17.33, 0.43)
        apic[75].pt3dadd(66.80, 503.35, 20.08, 0.43)
        apic[75].pt3dadd(66.55, 503.85, 20.09, 0.43)
        apic[75].pt3dadd(66.31, 504.84, 20.09, 0.43)
        apic[75].pt3dadd(65.35, 506.34, 20.09, 0.43)
        apic[75].pt3dadd(65.11, 506.58, 20.09, 0.43)
        apic[75].pt3dadd(64.14, 508.08, 20.09, 0.43)
        apic[75].pt3dadd(63.90, 508.33, 20.09, 0.43)
        
        apic[76] = h.Section(name='apic[76]')
        apic[76].connect(        apic[74](1), 0)
        apic[76].pt3dclear()
        apic[76].pt3dadd(73.56, 479.72, 13.26, 0.72)
        apic[76].pt3dadd(75.73, 485.69, 10.23, 0.72)
        apic[76].pt3dadd(77.42, 487.18, 10.23, 0.72)
        apic[76].pt3dadd(77.67, 487.18, 10.23, 0.72)
        apic[76].pt3dadd(78.39, 488.92, 10.23, 0.72)
        apic[76].pt3dadd(79.36, 490.17, 10.23, 0.72)
        apic[76].pt3dadd(80.08, 490.91, 10.23, 0.72)
        apic[76].pt3dadd(80.32, 491.41, 10.23, 0.72)
        apic[76].pt3dadd(82.50, 492.16, 10.23, 0.72)
        apic[76].pt3dadd(83.22, 492.65, 10.23, 0.72)
        apic[76].pt3dadd(83.95, 495.39, 10.23, 0.72)
        apic[76].pt3dadd(83.95, 498.87, 10.23, 0.72)
        apic[76].pt3dadd(83.70, 499.37, 10.23, 0.72)
        apic[76].pt3dadd(82.98, 500.37, 9.88, 0.72)
        apic[76].pt3dadd(82.74, 500.61, 9.88, 0.72)
        apic[76].pt3dadd(82.98, 502.36, 9.88, 0.72)
        apic[76].pt3dadd(82.98, 503.10, 9.88, 0.72)
        apic[76].pt3dadd(82.98, 504.35, 9.88, 0.72)
        apic[76].pt3dadd(82.26, 505.09, 9.88, 0.72)
        apic[76].pt3dadd(79.84, 506.09, 6.43, 0.72)
        apic[76].pt3dadd(78.87, 508.08, 6.43, 0.72)
        apic[76].pt3dadd(78.63, 508.33, 6.43, 0.72)
        apic[76].pt3dadd(77.91, 510.32, 6.43, 0.72)
        apic[76].pt3dadd(77.91, 510.56, 6.43, 0.72)
        apic[76].pt3dadd(78.15, 513.55, 6.43, 0.72)
        apic[76].pt3dadd(78.63, 514.30, 6.43, 0.72)
        apic[76].pt3dadd(78.63, 516.78, 6.43, 0.72)
        apic[76].pt3dadd(78.39, 519.27, 1.08, 0.72)
        apic[76].pt3dadd(76.70, 521.76, 1.08, 0.72)
        apic[76].pt3dadd(75.97, 524.25, 1.08, 0.72)
        apic[76].pt3dadd(75.97, 526.24, 1.08, 0.72)
        apic[76].pt3dadd(76.46, 527.23, 1.08, 0.61)
        apic[76].pt3dadd(76.70, 529.22, 1.08, 0.61)
        apic[76].pt3dadd(76.22, 529.97, 1.08, 0.61)
        apic[76].pt3dadd(74.28, 531.71, 1.08, 0.61)
        apic[76].pt3dadd(74.04, 531.71, 1.08, 0.61)
        apic[76].pt3dadd(72.59, 532.70, 1.08, 0.61)
        apic[76].pt3dadd(71.39, 533.70, 3.44, 0.61)
        apic[76].pt3dadd(71.39, 536.19, 3.44, 0.61)
        apic[76].pt3dadd(71.87, 536.68, 3.44, 0.61)
        apic[76].pt3dadd(72.11, 538.92, 3.44, 0.61)
        apic[76].pt3dadd(72.11, 539.42, 3.44, 0.61)
        apic[76].pt3dadd(71.63, 541.91, 3.44, 0.61)
        apic[76].pt3dadd(71.39, 542.65, 3.44, 0.61)
        apic[76].pt3dadd(69.45, 544.89, 3.44, 0.61)
        apic[76].pt3dadd(69.21, 545.14, 3.44, 0.61)
        apic[76].pt3dadd(68.73, 546.39, 3.44, 0.61)
        apic[76].pt3dadd(68.49, 546.88, 3.44, 0.61)
        apic[76].pt3dadd(68.25, 548.38, 3.44, 0.61)
        apic[76].pt3dadd(68.00, 548.62, 3.44, 0.61)
        apic[76].pt3dadd(66.80, 549.37, 3.44, 0.61)
        apic[76].pt3dadd(66.31, 549.62, 3.44, 0.61)
        apic[76].pt3dadd(64.86, 549.87, 3.44, 0.61)
        
        apic[77] = h.Section(name='apic[77]')
        apic[77].connect(        apic[76](1), 0)
        apic[77].pt3dclear()
        apic[77].pt3dadd(64.86, 549.87, 3.44, 0.43)
        apic[77].pt3dadd(63.17, 550.12, -4.20, 0.43)
        apic[77].pt3dadd(63.66, 553.85, -4.20, 0.43)
        apic[77].pt3dadd(64.62, 555.09, -4.20, 0.43)
        apic[77].pt3dadd(64.86, 555.09, -4.20, 0.43)
        apic[77].pt3dadd(66.55, 555.59, -4.20, 0.43)
        apic[77].pt3dadd(67.76, 555.84, -4.20, 0.43)
        apic[77].pt3dadd(68.00, 556.09, -4.20, 0.43)
        apic[77].pt3dadd(68.25, 556.09, -4.20, 0.43)
        apic[77].pt3dadd(69.21, 559.07, -4.20, 0.43)
        apic[77].pt3dadd(71.39, 559.57, -4.20, 0.43)
        apic[77].pt3dadd(72.35, 559.57, -4.20, 0.43)
        apic[77].pt3dadd(73.32, 559.82, -4.20, 0.43)
        apic[77].pt3dadd(74.28, 560.07, -4.20, 0.43)
        apic[77].pt3dadd(74.77, 560.32, -4.20, 0.43)
        apic[77].pt3dadd(75.73, 561.56, -4.20, 0.43)
        apic[77].pt3dadd(75.73, 562.06, -4.20, 0.43)
        apic[77].pt3dadd(75.73, 563.05, -4.20, 0.43)
        apic[77].pt3dadd(75.73, 563.80, -4.20, 0.43)
        apic[77].pt3dadd(75.73, 565.04, -4.20, 0.43)
        apic[77].pt3dadd(77.42, 567.53, -7.87, 0.43)
        apic[77].pt3dadd(77.42, 567.53, -7.90, 0.43)
        apic[77].pt3dadd(76.94, 569.77, -7.91, 0.43)
        apic[77].pt3dadd(76.46, 570.76, -7.91, 0.43)
        apic[77].pt3dadd(76.46, 572.26, -7.91, 0.43)
        apic[77].pt3dadd(76.46, 572.75, -7.91, 0.43)
        apic[77].pt3dadd(77.42, 573.75, -7.91, 0.43)
        apic[77].pt3dadd(77.91, 573.75, -7.91, 0.43)
        apic[77].pt3dadd(78.39, 574.25, -7.91, 0.43)
        apic[77].pt3dadd(80.08, 575.74, -11.20, 0.43)
        apic[77].pt3dadd(80.32, 577.73, -11.20, 0.43)
        apic[77].pt3dadd(80.32, 578.23, -11.21, 0.43)
        apic[77].pt3dadd(80.32, 579.47, -11.21, 0.43)
        apic[77].pt3dadd(80.08, 580.46, -11.21, 0.43)
        apic[77].pt3dadd(80.08, 581.21, -11.21, 0.43)
        apic[77].pt3dadd(80.08, 581.46, -11.21, 0.43)
        apic[77].pt3dadd(80.32, 582.70, -11.21, 0.43)
        apic[77].pt3dadd(80.32, 583.20, -11.21, 0.43)
        apic[77].pt3dadd(80.56, 584.44, -11.21, 0.43)
        apic[77].pt3dadd(80.56, 584.94, -11.21, 0.43)
        apic[77].pt3dadd(80.56, 585.94, -11.21, 0.43)
        apic[77].pt3dadd(81.29, 586.68, -11.22, 0.43)
        apic[77].pt3dadd(82.01, 587.43, -11.22, 0.43)
        apic[77].pt3dadd(82.26, 587.68, -11.22, 0.43)
        apic[77].pt3dadd(82.74, 588.42, -11.22, 0.43)
        apic[77].pt3dadd(82.98, 588.92, -11.22, 0.39)
        apic[77].pt3dadd(83.46, 591.16, -11.22, 0.39)
        apic[77].pt3dadd(83.95, 591.41, -11.22, 0.39)
        apic[77].pt3dadd(85.15, 591.91, -11.22, 0.39)
        apic[77].pt3dadd(86.36, 592.65, -11.22, 0.39)
        apic[77].pt3dadd(85.88, 596.39, -14.92, 0.39)
        apic[77].pt3dadd(85.88, 598.62, -14.92, 0.39)
        apic[77].pt3dadd(86.60, 599.37, -14.92, 0.39)
        apic[77].pt3dadd(87.81, 600.37, -14.92, 0.39)
        apic[77].pt3dadd(88.78, 601.36, -14.93, 0.39)
        apic[77].pt3dadd(89.26, 602.11, -14.93, 0.39)
        apic[77].pt3dadd(89.74, 602.36, -14.93, 0.39)
        apic[77].pt3dadd(90.47, 603.10, -14.93, 0.39)
        apic[77].pt3dadd(91.68, 603.60, -14.94, 0.39)
        apic[77].pt3dadd(92.16, 603.85, -14.94, 0.39)
        apic[77].pt3dadd(92.88, 604.59, -14.94, 0.39)
        apic[77].pt3dadd(93.61, 605.59, -14.95, 0.39)
        apic[77].pt3dadd(93.61, 607.08, -14.95, 0.39)
        apic[77].pt3dadd(93.61, 607.58, -14.95, 0.39)
        apic[77].pt3dadd(93.61, 608.33, -14.95, 0.39)
        apic[77].pt3dadd(93.37, 608.82, -14.95, 0.39)
        apic[77].pt3dadd(92.88, 609.57, -14.95, 0.39)
        apic[77].pt3dadd(92.16, 609.82, -14.95, 0.39)
        
        apic[78] = h.Section(name='apic[78]')
        apic[78].connect(        apic[76](1), 0)
        apic[78].pt3dclear()
        apic[78].pt3dadd(64.86, 549.87, 3.44, 0.43)
        apic[78].pt3dadd(67.76, 551.11, 3.44, 0.43)
        apic[78].pt3dadd(69.94, 553.10, 3.44, 0.43)
        apic[78].pt3dadd(70.42, 554.10, -13.65, 0.43)
        apic[78].pt3dadd(70.42, 554.10, -13.74, 0.43)
        apic[78].pt3dadd(70.66, 556.09, -14.00, 0.43)
        apic[78].pt3dadd(70.66, 556.09, -14.14, 0.43)
        apic[78].pt3dadd(70.66, 557.33, -14.44, 0.43)
        apic[78].pt3dadd(70.90, 557.33, -14.64, 0.43)
        apic[78].pt3dadd(71.39, 559.32, -15.28, 0.43)
        apic[78].pt3dadd(71.39, 559.32, -15.33, 0.43)
        apic[78].pt3dadd(71.39, 561.31, -15.48, 0.43)
        apic[78].pt3dadd(71.39, 561.81, -15.51, 0.43)
        apic[78].pt3dadd(71.39, 563.55, -15.52, 0.43)
        apic[78].pt3dadd(71.39, 564.05, -15.53, 0.43)
        apic[78].pt3dadd(71.87, 565.04, -15.59, 0.43)
        apic[78].pt3dadd(72.35, 565.79, -15.60, 0.43)
        apic[78].pt3dadd(73.08, 566.78, -15.60, 0.43)
        apic[78].pt3dadd(73.56, 567.28, -15.60, 0.43)
        apic[78].pt3dadd(73.56, 568.28, -15.60, 0.43)
        apic[78].pt3dadd(73.56, 568.52, -15.60, 0.43)
        apic[78].pt3dadd(73.56, 569.52, -15.60, 0.43)
        apic[78].pt3dadd(73.56, 569.52, -15.71, 0.43)
        apic[78].pt3dadd(73.56, 570.51, -16.31, 0.43)
        apic[78].pt3dadd(73.56, 571.26, -16.65, 0.43)
        apic[78].pt3dadd(73.56, 572.75, -17.22, 0.43)
        apic[78].pt3dadd(73.56, 572.75, -17.47, 0.43)
        apic[78].pt3dadd(74.77, 573.50, -17.51, 0.43)
        apic[78].pt3dadd(75.25, 573.75, -17.52, 0.43)
        apic[78].pt3dadd(75.73, 574.74, -17.52, 0.43)
        apic[78].pt3dadd(74.04, 576.48, -17.52, 0.43)
        apic[78].pt3dadd(73.80, 576.73, -17.52, 0.43)
        apic[78].pt3dadd(73.32, 577.23, -17.52, 0.43)
        apic[78].pt3dadd(73.32, 578.23, -17.52, 0.43)
        apic[78].pt3dadd(75.01, 579.72, -17.52, 0.43)
        apic[78].pt3dadd(75.25, 580.46, -17.52, 0.43)
        apic[78].pt3dadd(75.25, 581.46, -17.52, 0.43)
        apic[78].pt3dadd(76.70, 582.95, -17.52, 0.43)
        apic[78].pt3dadd(76.94, 583.20, -17.52, 0.43)
        apic[78].pt3dadd(77.67, 584.69, -17.52, 0.43)
        apic[78].pt3dadd(78.15, 585.69, -20.21, 0.43)
        apic[78].pt3dadd(79.12, 586.68, -20.21, 0.43)
        apic[78].pt3dadd(79.12, 586.93, -20.21, 0.43)
        apic[78].pt3dadd(79.36, 587.43, -20.22, 0.43)
        apic[78].pt3dadd(79.60, 588.42, -20.22, 0.43)
        apic[78].pt3dadd(78.87, 589.92, -20.23, 0.43)
        apic[78].pt3dadd(78.63, 589.92, -20.23, 0.43)
        apic[78].pt3dadd(77.42, 590.42, -20.24, 0.43)
        apic[78].pt3dadd(77.42, 590.91, -20.25, 0.43)
        apic[78].pt3dadd(77.18, 591.66, -20.26, 0.43)
        apic[78].pt3dadd(77.18, 592.90, -20.26, 0.43)
        apic[78].pt3dadd(77.42, 593.40, -20.26, 0.43)
        apic[78].pt3dadd(77.91, 593.65, -20.27, 0.43)
        apic[78].pt3dadd(79.12, 594.15, -20.27, 0.43)
        apic[78].pt3dadd(79.12, 594.40, -20.27, 0.43)
        apic[78].pt3dadd(79.12, 594.40, -20.21, 0.43)
        apic[78].pt3dadd(80.08, 595.64, -23.51, 0.39)
        apic[78].pt3dadd(80.08, 597.38, -23.50, 0.39)
        apic[78].pt3dadd(79.60, 599.62, -23.50, 0.39)
        apic[78].pt3dadd(79.60, 600.12, -23.50, 0.39)
        apic[78].pt3dadd(79.60, 601.11, -23.50, 0.39)
        apic[78].pt3dadd(79.60, 601.36, -23.50, 0.39)
        apic[78].pt3dadd(79.84, 601.86, -23.50, 0.39)
        apic[78].pt3dadd(81.05, 603.85, -23.50, 0.39)
        apic[78].pt3dadd(81.05, 606.09, -23.50, 0.39)
        apic[78].pt3dadd(81.05, 607.08, -23.50, 0.39)
        apic[78].pt3dadd(81.05, 607.33, -23.50, 0.39)
        apic[78].pt3dadd(81.77, 608.08, -23.50, 0.39)
        apic[78].pt3dadd(82.01, 608.82, -23.50, 0.39)
        
        for sec in h.allsec():
            self.all.append(sec)

    def delete_cell(self):
        for sec in h.allsec():
            h.delete_section(sec=sec)
        del self.times
        del self.gmax_values 
        del self.time_vector 
        del self.gmax_vector
        del self.all
        del self

    def geom_nseg(self, d_lambda, freq):
        for sec in h.allsec():
            sec.nseg = max(int((sec.L / (d_lambda * self.lambda_f(sec, freq)) + 0.9) / 2) * 2 + 1, sec.nseg)

    def lambda_f(self, sec, freq):
        if sec.n3d() < 2:
            return 1e5 * sqrt(sec.diam / (4 * np.pi * freq * sec.Ra * sec.cm))
        
        x1 = sec.arc3d(0)
        d1 = sec.diam3d(0)
        lam = 0
        for i in range(1, int(sec.n3d())):
            x2 = sec.arc3d(i)
            d2 = sec.diam3d(i)
            lam += (x2 - x1) / sqrt(d1 + d2)
            x1, d1 = x2, d2
        
        lam *= sqrt(2) * 1e-5 * sqrt(4 * np.pi * freq * sec.Ra * sec.cm)
        return sec.L / lam
```
