## Supplementary NEURON models for "Dendritic excitations govern back-propagation via a spike-rate accelerometer": main.py

```
import numpy as np
from run_simulation import run_simulation
from run_morphology import run_morphology
from delete_neuron import delete_all_sections
from plot_results import plot_results
from plot_results import kymograph
from run_simulation import reshape_v
import matplotlib.pyplot as plt
import json
import os
from parameters_list import setup_parameters

# File to load morphology from
morphology_file = "Jarsky"

dt = 0.025  # ms

# Channels parameters
Nav_inactivation = [1, 0.6, 300, 100]
Kdr_params = [0.04, 0.04, 0, 1]

Nap_params = [1e-9, 1e-9, 300, 100]
Kca_params = [1e-9, 1e-9, 550, 100]
Km_params = [1e-9, 1e-9, 550, 100]
Mykca_params = [1e-9, 1e-9, 550, 100]

CalH_params = [1e-9, 1e-9, 300, 100]
Car_mag_params = [1e-9, 1e-9, 550, 100]
Somacar_params = [1e-9, 1e-9, 50, 30]
Cat_params = [1e-9, 1e-9, 200, 100]
Hcn_params = [1e-9, 1e-9, 200, 100]
# Channel Parameters: [g0, g1, d, z]
# where g = g0 + (g1 - g0) / (1 + exp((d - x) / z))

#sampling = "sections"
sampling = "segments"


for run_nr in [0, 1]:

    [duration, stim_start, stim_end, Chr_params, synapse_params, sim_params, outputs, waveform, waveform_patch, patch_params, Nav_params, Kad_params, Kap_params, Cal_params, Car_params] = setup_parameters(run_nr)

    outputs = ["v", "g_chr", "gexp_chr", "s_na3", "h_na3", "gbar_na3", "l_kad", "gkabar_kad", "ina_na3", "inavalue_nap", "ik_kdr", "ik_kad", "ik_kap", "ikvalue_km", "ivalue_hd", "i_chr", "cai", "iCa_cat", "icavalue_car", "icavalue_cal", "icavalue_calH", "ikvalue_mykca"]
    outputs = ["v", "g_chr", "gexp_chr"]

    channel_params = [Nav_params, Nav_inactivation, Kap_params, Kad_params, Kdr_params, Chr_params, Kca_params, CalH_params, Cal_params, Somacar_params, Nap_params, Mykca_params, Car_params, Car_mag_params, Cat_params, Hcn_params, Km_params]

    # Run simulation
    morpho = run_morphology(channel_params, morphology_file, sampling)
    delete_all_sections()
    d = morpho[1]
    p = morpho[2]
    path = morpho[3]
    distance_matrix = [[float(x) for x in item] for item in d]

    result = run_simulation(sim_params, channel_params, synapse_params, waveform, outputs, morphology_file, patch_params, distance_matrix, sampling)
    delete_all_sections()

    # Get results
    time = [float(x) for x in result[0]]
    section_names = [x for x in morpho[0]]
    result_vars = result[1]
    positions = [[float(x) for x in item] for item in p]
    waveform = [x if stim_start <= t <= stim_end else 0 for x, t in zip(waveform, time)]


    synapse_nr = 0
    for param in synapse_params:
        synapse_nr += int(param[0])

    synapse_pos = []
    spike_times = []
    spike_times_frames = []
    for idx in range(synapse_nr):
        synapse_pos.append([float(x) for x in result[2][idx]])
        spike_times.append([float(x) for x in result[3][idx]])
        spike_times_frames.append([float(x) for x in result[4][idx]])

    if 'synapse_pos' not in locals():
        synapse_pos = []
    if 'spike_times' not in locals():
        spike_times = []
        spike_times_frames = []

    allSpikeTimes = [item for sublist in spike_times for item in sublist]

    varNames = outputs
    for varName in varNames:
        varCell = result_vars[varName]
        varMatrix = []
        for item in varCell:
            varMatrix.extend([float(x) for x in item])
        locals()[varName] = varMatrix

    for output in outputs:
        variable = locals()[output]
        reshaped_variable = reshape_v(variable, distance_matrix)
        locals()[output] = reshaped_variable

    # Plot results
    plot_results(time, v, distance_matrix, duration, section_names, False)
    [distances, voltages] = kymograph(time,v,distance_matrix,path)

    # Animate results
    data_file = "data.json"
    data_syn_file = "data_syn.json"
    #animation_path = ""
    animation_path = "none"
    if animation_path != "none":
        # Combine the strings for the file names and paths
        combined_data_file = os.path.join(animation_path, data_file)
        combined_data_syn_file = os.path.join(animation_path, data_syn_file)
        # If the files exist, delete them
        if os.path.isfile(combined_data_file):
            os.remove(combined_data_file)
        if os.path.isfile(combined_data_syn_file):
            os.remove(combined_data_syn_file)

        # Define data and write to file
        v_short = np.array(v).astype(float).tolist()  # Convert to list
        data = {'positions': positions, 'var_plot': v_short}  # No need to convert positions to list
        with open(combined_data_file, 'w') as file:
            json.dump(data, file)

        # Prepare spike times and synapse positions
        spike_times_cell = [list(spike_times_frame) for spike_times_frame in spike_times_frames] # Converting inner lists if needed
        data_syn = {'spike_times': spike_times_cell, 'synapse_pos': list(synapse_pos)}  # Convert synapse_pos to list if needed
        with open(combined_data_syn_file, 'w') as file_syn:
            json.dump(data_syn, file_syn)

    # Create the figure
    fig, ax1 = plt.subplots()
    fig.patch.set_facecolor('none')
    ax1.set_xlabel('Time (ms)')
    ax1.set_ylabel('Net Synaptic Stimulation (Hz)', color=(240/255, 240/255, 240/255))
    if len(allSpikeTimes) == 0:
        ax1.plot(0, 0)
        ax1.set_ylim([0, 10])
    else:
        _, bins, _ = ax1.hist(allSpikeTimes, bins=range(0, int(duration), 1), facecolor=(240/255, 240/255, 240/255), edgecolor=(240/255, 240/255, 240/255))
        ax1.set_ylim([0, max(np.histogram(allSpikeTimes, bins=bins)[0])])

    ax2 = ax1.twinx()
    if max(waveform) == 0:
        waveform_norm = waveform
    else:
        waveform_norm = waveform / max(waveform)
    ax2.plot(time, waveform_norm, linewidth=2, color=(0, 0.705, 0.896))
    ax2.set_ylabel('Optogenetic Stimulation', rotation=-90, labelpad=15)
    if max(waveform) == 0:
        ax2.set_ylim([0, 1.05])
    else:
        ax2.set_ylim([0, max(waveform_norm)*1.05])
    for ax in [ax1, ax2]:
        ax.spines['bottom'].set_color((240/255, 240/255, 240/255))
        ax.spines['top'].set_color((240/255, 240/255, 240/255))
        ax.xaxis.label.set_color((240/255, 240/255, 240/255))
        ax.yaxis.label.set_color((240/255, 240/255, 240/255))
        ax.tick_params(axis='x', colors=(240/255, 240/255, 240/255))
        ax.tick_params(axis='y', colors=(240/255, 240/255, 240/255))
        ax.xaxis.label.set_size(14)
        ax.yaxis.label.set_size(14)
        ax.tick_params(axis='both', which='major', labelsize=14)
    plt.xlim([0, duration])
    plt.tight_layout()

    if animation_path != "none":
        plt.savefig(f"{animation_path}/waveform.png", transparent=True)
    plt.show()
```
