## Supplementary NEURON models for "Dendritic excitations govern back-propagation via a spike-rate accelerometer": ampa_forti.mod

```
COMMENT

Forti etal's (1997) AMPA synapse with two decay time constants based on the 
exp2syn.mod mechanism from the NEURON distribution

The value of 0.002 for the fraction of Ca flowing at low membrane
potentials is derived from the measurements of Spruston &al (1995)

Forti & al. 1997 "Loose-patch recordings of single quanta at
  individual hippocampal synapses" Nature 388, 874-878

Spruston & al. 1995 "Dendritic glutamate receptor channels in rat
  hippocampal CA3 and CA1 neurons" J. Physiol. 482, 325-352

ENDCOMMENT

NEURON {
	  POINT_PROCESS AmpaSyn
	  RANGE  e, g, i, ica
	  NONSPECIFIC_CURRENT i
    USEION ca READ cai,cao WRITE ica
    GLOBAL total, taurise, taufast, tauslow, taurise_exp, taufast_exp, tauslow_exp, afast, aslow, normfac, T_exp, fracca
}

INCLUDE "units.inc"

PARAMETER {
	  taurise_exp =.2 (ms) <1e-9,1e9>       : rise
	  taufast_exp = 0.61 (ms) <1e-9,1e9>    : fast decay
	  tauslow_exp = 2.55 (ms) <1e-9,1e9>    : slow decay
    afast = 0.75 <0,1>
	  e=0	(mV)
    fracca= 0.002        : fraction of current that is ca ions
    z = 2
    celsius = 22	(degC)
    T_exp = 22    (degC)
    q10 = 3
}

ASSIGNED {
    v       (mV)
    i       (nA)
    ica	    (nA) 	
    g       (uS)
    aslow 
    total   (uS)
    cai     (mM)
    cao     (mM)
    taurise (ms)
    taufast (ms)
    tauslow (ms)
    normfac 
}

BREAKPOINT {
	  SOLVE state METHOD cnexp
    g = (B + C - A) 
    i =   g * (1-fracca) * (v - e)
    ica = g * fracca     * ghkg(v,cai,cao,z)
}

INCLUDE "triexpsyn.inc"

INCLUDE "ghk.inc"
```
