## Supplementary NEURON models for "Dendritic excitations govern back-propagation via a spike-rate accelerometer": cacum.mod

```
COMMENT
	calcium accumulation into a volume of area*depth next to the
	membrane with a decay (time constant tau) to resting level
	given by the global calcium variable cai0_ca_ion
ENDCOMMENT

NEURON {
	SUFFIX cacum
	USEION ca READ ica WRITE cai
	RANGE depth, tau, cai0
}

UNITS {
	(mM) = (milli/liter)
	(mA) = (milliamp)
	F = (faraday) (coulombs)
}

PARAMETER {
	depth = 1 (nm)	: assume volume = area*depth
	tau = 10 (ms)
	cai0 = 50e-6 (mM)	: Requires explicit use in INITIAL
			: block for it to take precedence over cai0_ca_ion
			: Do not forget to initialize in hoc if different
			: from this default.
}

ASSIGNED {
	ica (mA/cm2)
}

STATE {
	cai (mM)
}

INITIAL {
	cai = cai0
}

BREAKPOINT {
	SOLVE integrate METHOD derivimplicit
}

DERIVATIVE integrate {
	cai' = -ica/depth/F/2 * (1e7) + (cai0 - cai)/tau
}
```
