## Supplementary NEURON models for "Dendritic excitations govern back-propagation via a spike-rate accelerometer": cad.mod

```
TITLE decay of internal calcium concentration
:
: Internal calcium concentration due to calcium currents and pump.
: Differential equations.
:
: Simple model of ATPase pump with 3 kinetic constants (Destexhe 92)
:     Cai + P <-> CaP -> Cao + P  (k1,k2,k3)
: A Michaelis-Menten approximation is assumed, which reduces the complexity
: of the system to 2 parameters: 
:       kt = <tot enzyme concentration> * k3  -> TIME CONSTANT OF THE PUMP
:	kd = k2/k1 (dissociation constant)    -> EQUILIBRIUM CALCIUM VALUE
: The values of these parameters are chosen assuming a high affinity of 
: the pump to calcium and a low transport capacity (cfr. Blaustein, 
: TINS, 11: 438, 1988, and references therein).  
:
: Units checked using "modlunit" -> factor 10000 needed in ca entry
:
: VERSION OF PUMP + DECAY (decay can be viewed as simplified buffering)
:
: All variables are range variables
:
:
: This mechanism was published in:  Destexhe, A. Babloyantz, A. and 
: Sejnowski, TJ.  Ionic mechanisms for intrinsic slow oscillations in
: thalamic relay neurons. Biophys. J. 65: 1538-1552, 1993)
:
: Written by Alain Destexhe, Salk Institute, Nov 12, 1992
:
: This file was modified by Yiota Poirazi on April 18, 2001 to account for the sharp
: Ca++ spike repolarization observed in: Golding, N. Jung H-Y., Mickus T. and Spruston N
: "Dendritic Calcium Spike Initiation and Repolarization are controlled by distinct potassium channel
: subtypes in CA1 pyramidal neurons". J. of Neuroscience 19(20) 8789-8798, 1999.
:
:  factor 10000 is replaced by 10000/18 needed in ca entry
:  taur --rate of calcium removal-- is replaced by taur*7 (7 times faster) 


INDEPENDENT {t FROM 0 TO 1 WITH 1 (ms)}

NEURON {
	SUFFIX cad
	USEION ca READ ica, cai WRITE cai	
        RANGE ca
	GLOBAL depth,cainf,taur
}

UNITS {
	(molar) = (1/liter)			: moles do not appear in units
	(mM)	= (millimolar)
	(um)	= (micron)
	(mA)	= (milliamp)
	(msM)	= (ms mM)
	FARADAY = (faraday) (coulomb)
}


PARAMETER {
	depth	= .1	(um)		: depth of shell
	taur	= 200	(ms)		: rate of calcium removal
	cainf	= 100e-6(mM)
	cai		(mM)
}

STATE {
	ca		(mM) 
}

INITIAL {
	ca = cainf
}

ASSIGNED {
	ica		(mA/cm2)
	drive_channel	(mM/ms)
}
	
BREAKPOINT {
	SOLVE state METHOD euler
}

DERIVATIVE state { 

	drive_channel =  - (10000) * ica / (2 * FARADAY * depth)
	if (drive_channel <= 0.) { drive_channel = 0.  }   : cannot pump inward 
         
	:ca' = drive_channel + (cainf-ca)/taur
        ca' = drive_channel/18 + (cainf -ca)/taur*7
	cai = ca
}
```
