## Supplementary NEURON models for "Dendritic excitations govern back-propagation via a spike-rate accelerometer": cagk.mod

```
TITLE CaGk
: Calcium activated mAHP K channel.
: From Moczydlowski and Latorre (1983) J. Gen. Physiol. 82

UNITS {
	(molar) = (1/liter)
	(mV) =	(millivolt)
	(mA) =	(milliamp)
	(mM) =	(millimolar)
	FARADAY = (faraday)  (kilocoulombs)
	R = (k-mole) (joule/degC)
}

NEURON {
	SUFFIX mykca
	USEION ca READ cai
	USEION k READ ek WRITE ik
	RANGE gkbar, g, gmax, ikvalue
	GLOBAL oinf, tau
}

PARAMETER {
	celsius = 20	(degC)
	v		(mV)
	ek		(mV)
	gkbar = 0.01	(mho/cm2)	: Maximum Permeability
	cai = 1e-3	(mM)
	d1 = 0.84
	d2 = 1.0
	k1 = 0.18	(mM)
	k2 = 0.011	(mM)
	bbar = 0.28	(/ms)
	abar = 0.48	(/ms)
	ikvalue (mA/cm2)
}
COMMENT
the preceding two numbers were switched on 8/19/92 in response to a bug
report by Bartlett Mel. In the paper the kinetic scheme is
C <-> CCa (K1)
CCa <-> OCa (beta2,alpha2)
OCa <-> OCa2 (K4)
In this model abar = beta2 and bbar = alpha2 and K4 comes from d2 and k2
I was forcing things into a nomenclature where alpha is the rate from
closed to open. Unfortunately I didn't switch the numbers.
ENDCOMMENT

ASSIGNED {
	ik		(mA/cm2)
	oinf
	tau		(ms)
  g     (mho/cm2)
  gmax  (mho/cm2)
}

STATE {	o }		: fraction of open channels

BREAKPOINT {
	  SOLVE state METHOD cnexp
    g = gkbar*o
	  ik = g*(v - ek)
	  ikvalue = ik
    if (g > gmax ) {
        gmax = g
    }
}

DERIVATIVE state {
	rate(v, cai)
	o' = (oinf - o)/tau
}

INITIAL {
	rate(v, cai)
	o = oinf
  gmax = 0
}

FUNCTION alp(v (mV), ca (mM)) (1/ms) { :callable from hoc
	alp = abar/(1 + exp1(k1,d1,v)/ca)
}

FUNCTION bet(v (mV), ca (mM)) (1/ms) { :callable from hoc
	bet = bbar/(1 + ca/exp1(k2,d2,v))
}

FUNCTION exp1(k (mM), d, v (mV)) (mM) { :callable from hoc
	exp1 = k*exp(-2*d*FARADAY*v/R/(273.15 + celsius))
}

PROCEDURE rate(v (mV), ca (mM)) { :callable from hoc
	LOCAL a
	a = alp(v,ca)
	tau = 1/(a + bet(v, ca))
	oinf = a*tau
}
```
