## Supplementary NEURON models for "Dendritic excitations govern back-propagation via a spike-rate accelerometer": cal.mod

```
TITLE L-type calcium channel with low threshold for activation
: used in somatic and proximal dendritic regions 
: it calculates I_Ca using channel permeability instead of conductance

UNITS {
	  (mA) = (milliamp)
	  (mV) = (millivolt)
    (molar) = (1/liter)
    (mM) = (millimolar)
    FARADAY = (faraday) (coulomb)
    R = (k-mole) (joule/degC)
}

PARAMETER {		:parameters that can be entered when function is called in cell-setup 
	  v             (mV)
	  celsius = 34	(degC)
	  gcalbar = 0   (mho/cm2)   : initialized conductance
	  ki  = 0.001   (mM)  
	  cai = 5.e-5   (mM)        : initial internal Ca++ concentration
	  cao = 2       (mM)        : initial external Ca++ concentration
    tfa = 5                   : time constant scaling factor
    eca = 140     (mV)        : Ca++ reversal potential
    icavalue (mA/cm2)
}

NEURON {
	  SUFFIX cal
	  USEION ca READ cai,cao WRITE ica
    RANGE gcalbar, gmax, gcal, minf, taum, icavalue
}

STATE {	m }                      : unknown parameter to be solved in the DEs 

ASSIGNED {                       : parameters needed to solve DE
	  ica   (mA/cm2)
    gcal  (mho/cm2)
    gmax  (mho/cm2) 
    minf
    taum  (ms)
}

INITIAL {                        : initialize the following parameter using rates()
    rates(v)
    m = minf
	  gcal = gcalbar*m*h2(cai)
}

BREAKPOINT {
	  SOLVE states METHOD cnexp
	  gcal = gcalbar*m*h2(cai) : maximum channel permeability
	  ica = gcal*ghk(v,cai,cao): calcium current induced by this channel
      icavalue = ica
    if (gcal > gmax) {
        gmax = gcal
    }
}

FUNCTION h2(cai(mM)) {
	  h2 = ki/(ki+cai)
}

FUNCTION ghk(v(mV), ci(mM), co(mM)) (mV) {
    LOCAL nu,f
    f = KTF(celsius)/2
    nu = v/f
    ghk=-f*(1. - (ci/co)*exp(nu))*efun(nu)
}

FUNCTION KTF(celsius (degC)) (mV) { : temperature-dependent adjustment factor
    KTF = ((25.(mV)/293.15(degC))*(celsius + 273.15(degC)))
}

FUNCTION efun(z) {
	  if (fabs(z) < 1e-4) {
		    efun = 1 - z/2
	  }else{
		    efun = z/(exp(z) - 1)
	  }
}

FUNCTION alpm(v (mV)) (/ms) {
	  alpm = 0.055(/ms/mV)*(-27.01(mV) - v)/(exp((-27.01(mV)-v)/3.8(mV)) - 1)
}


FUNCTION betm(v (mV)) (/ms) {
    betm =0.94(/ms)*exp((-63.01(mV)-v)/17(mV))
}

:if state_cagk is called from hoc, garbage or segmentation violation will
:result because range variables won't have correct pointer.  This is because
:only BREAKPOINT sets up the correct pointers to range variables.
DERIVATIVE states {     : exact when v held constant; integrates over dt step
    rates(v)
    m' = (minf - m)/taum
}

PROCEDURE rates(v (mV)) { :callable from hoc
    LOCAL a
    TABLE taum, minf FROM -150 TO 150 WITH 300 
    a = alpm(v)
    taum = 1/(tfa*(a+betm(v))) : estimation of activation tau
    minf = a/(a+betm(v))       : estimation of activation steady state value
}
```
