## Supplementary NEURON models for "Dendritic excitations govern back-propagation via a spike-rate accelerometer": calH.mod

```
TITLE Ca L-type channel with high treshold of activation
: inserted in distal dendrites to account for distally
: restricted initiation of Ca++ spikes
: uses channel conductance (not permeability)
: written by Yiota Poirazi, 1/8/00

NEURON {
	  SUFFIX calH
	  USEION ca READ eca WRITE ica
    RANGE gcalbar, m, h, g, gmax, icavalue
	  RANGE inf, tau
}

UNITS {
	  (mA) = (milliamp)
	  (mV) = (millivolt)
}

PARAMETER {          : parameters that can be entered when function is called in cell-setup
    v             (mV)
    celsius = 34	(degC)
    gcalbar = 0   (mho/cm2) : initialized conductance
	  eca = 140     (mV)      : Ca++ reversal potential
	  icavalue (mA/cm2)
}

STATE {	m h }                     : unknown activation and inactivation parameters to be solved in the DEs  

ASSIGNED {                        : parameters needed to solve DE
	  ica    (mA/cm2)
    inf[2]
	  tau[2] (ms)
    g      (mho/cm2)
    gmax   (mho/cm2)
}

BREAKPOINT {
	  SOLVE states METHOD cnexp
    g = gcalbar*m*m*m*h
	  ica = g*(v - eca)
	  icavalue = ica       
    if (g > gmax) {
        gmax = g
    }
}

INITIAL {
	  mhn(v)
    m = inf[0]
    h = inf[1]
	  ica = gcalbar*m*m*m*h*(v - eca) : initial Ca++ current value
}

DERIVATIVE states {	: exact when v held constant
	  mhn(v)
	  m' = (inf[0] - m)/tau[0]
	  h' = (inf[1] - h)/tau[1]
}

FUNCTION varss(v (mV), i) {
	  if (i==0) { 
        varss = 1 / (1 + exp((v+37(mV))/(-1(mV))))  : Ca activation 
	  }
	  else if (i==1) { 
        varss = 1 / (1 + exp((v+41(mV))/(0.5(mV)))) : Ca inactivation 
	  }
}

FUNCTION vartau(v (mV), i) (ms) {
	  if (i==0) {
        vartau = 3.6(ms)  : activation variable time constant
    }
	  else if (i==1) {
        :           vartau = 25   : inactivation variable time constant
        vartau = 29(ms)   : inactivation variable time constant
    }
}	

PROCEDURE mhn(v (mV)) {LOCAL a, b :rest = -70
    TABLE inf, tau DEPEND celsius FROM -100 TO 100 WITH 200
	  FROM i=0 TO 1 {
		    tau[i] = vartau(v,i)
		    inf[i] = varss(v,i)
	  }
}
```
