## Supplementary NEURON models for "Dendritic excitations govern back-propagation via a spike-rate accelerometer": car_mag.mod

```
TITLE Ca R-type channel with medium threshold for activation

COMMENT 
  Kinetics taken from Jeffrey C Magee and Daniel Johnston (1995)
  "Characterization of single voltage-gated Na+ and Ca2+ channels in
  apical dendrites of rat CA1 pyramidal neurons", J. Physiol. 487(1):
  67-90.
ENDCOMMENT

NEURON {
    SUFFIX car_mag
    USEION ca READ cai, cao WRITE ica
    RANGE gmax, m, h
    RANGE minf, hinf, taum, tauh
    GLOBAL q10, taum_exp, z
}

INCLUDE "units.inc"

PARAMETER {   
    gmax = 0      (S/cm2) <0,1e9> 
    q10  = 3  
    taum_exp = 0.92  (ms)            : experimentally-measured taum
    z = 2                         : valency of Ca ions
}  

STATE {	mO mC hO hC }    

ASSIGNED {               : parameters needed to solve DE
    v       (mV)
    celsius (degC)
    cai     (mM)
    cao     (mM)
	  ica     (mA/cm2)
    minf
    hinf
	  taum    (ms)
    tauh    (ms)
}

BREAKPOINT {
    SOLVE kin METHOD sparse
	  ica = gmax*mO*mO*mO*hO*ghkg(v,cai,cao,z)
}

INITIAL { 
    taum = q10^(-(celsius-22(degC))/10(degC))*taum_exp
    tauh = q10^(-(celsius-22(degC))/10(degC))*53(ms)
    SOLVE kin STEADYSTATE sparse    
    ica = gmax*mO*mO*mO*hO*ghkg(v,cai,cao,z)
}

KINETIC kin {
    minf = 1/(1+exp(-(v- 3(mV))/8.3(mV)))
    hinf = 1/(1+exp( (v+39(mV))/9.2(mV)))
    ~ mC <-> mO (minf/taum, (1-minf)/taum)
    ~ hC <-> hO (hinf/tauh, (1-hinf)/tauh)
    CONSERVE mC + mO = 1
    CONSERVE hC + hO = 1
}

INCLUDE "ghk.inc"
```
