## Supplementary NEURON models for "Dendritic excitations govern back-propagation via a spike-rate accelerometer": cat.mod

```
TITLE t-type calcium channel with high threshold for activation
: used in somatic and dendritic regions 
: it calculates I_Ca using channel permeability instead of conductance

UNITS {
	  (mA) = (milliamp)
	  (mV) = (millivolt)
    (molar) = (1/liter)
    (mM) = (millimolar)
    FARADAY = (faraday) (coulomb)
    R = (k-mole) (joule/degC)
}


PARAMETER {           :parameters that can be entered when function is called in cell-setup 
	  v             (mV)
    tBase = 23.5  (degC)
	  celsius = 22  (degC)
	  gcatbar = 0   (mho/cm2)  : initialized conductance
	  ki = 0.001    (mM)
	  cai = 5.e-5   (mM)       : initial internal Ca++ concentration
	  cao = 2       (mM)       : initial external Ca++ concentration
    tfa = 1                  : activation time constant scaling factor
    tfi = 0.68               : inactivation time constant scaling factor
    eca = 140                : Ca++ reversal potential
}

NEURON {
	  SUFFIX cat
	  USEION ca READ cai,cao 
    USEION Ca WRITE iCa VALENCE 2
    : The T-current does not activate calcium-dependent currents.
    : The construction with dummy ion Ca prevents the updating of the 
    : internal calcium concentration. 
    RANGE gcatbar, hinf, minf, taum, tauh, iCa, gmax
}

STATE {	m h }  : unknown activation and inactivation parameters to be solved in the DEs 

ASSIGNED {     : parameters needed to solve DE
	  iCa  (mA/cm2)
    gcat (mho/cm2) 
    gmax (mho/cm2) 
    minf
    hinf
    taum (ms)
    tauh (ms)
}

INITIAL {
    :        tadj = 3^((celsius-tBase)/10)   : assume Q10 of 3
    rates(v)
    m = minf
    h = hinf
	  gcat = gcatbar*m*m*h*h2(cai)
    gmax = gcat
}

BREAKPOINT {
	  SOLVE states METHOD cnexp
	  gcat = gcatbar*m*m*h*h2(cai) : maximum channel permeability
	  iCa = gcat*ghk(v,cai,cao)    : dummy calcium current induced by this channel
    if (gcat > gmax) {
        gmax = gcat
    }
}

FUNCTION h2(cai(mM)) {
	  h2 = ki/(ki+cai)
}

FUNCTION ghk(v(mV), ci(mM), co(mM)) (mV) { LOCAL nu,f
    f = KTF(celsius)/2
    nu = v/f
    ghk=-f*(1. - (ci/co)*exp(nu))*efun(nu)
}

FUNCTION KTF(celsius (degC)) (mV) {   : temperature-dependent adjustment factor
    KTF = ((25(mV)/293.15(degC))*(celsius + 273.15(degC)))
}

FUNCTION efun(z) {
	  if (fabs(z) < 1e-4) {
		    efun = 1 - z/2
	  }else{
		    efun = z/(exp(z) - 1)
	  }
}

FUNCTION alph(v(mV)) (/ms) {
	  alph = 1.6e-4(/ms)*exp(-(v+57(mV))/19(mV))
}

FUNCTION beth(v(mV)) (/ms) {
	  beth = 1(/ms)/(exp((-v+15(mV))/10(mV))+1.0)
}

FUNCTION alpm(v(mV)) (/ms) {
	  alpm = 0.1967(/ms)*(-1.0(/mV)*v+19.88)/(exp((-1.0*v+19.88(mV))/10.0(mV))-1.0)
}

FUNCTION betm(v(mV)) (/ms) {
	  betm = 0.046(/ms)*exp(-v/22.73(mV))
}

:if state_cagk is called from hoc, garbage or segmentation violation will
:result because range variables won't have correct pointer.  This is because
: only BREAKPOINT sets up the correct pointers to range variables.
DERIVATIVE states {     : exact when v held constant; integrates over dt step
    rates(v)
    m' = (minf - m)/taum
    h' = (hinf - h)/tauh
}

PROCEDURE rates(v (mV)) { :callable from hoc
    LOCAL a
    TABLE taum, minf, tauh, hinf FROM -150 TO 150 WITH 300
    a = alpm(v)
    taum = 1/(tfa*(a + betm(v))) : estimation of activation tau
    minf =  a/(a+betm(v))        : estimation of activation steady state
    a = alph(v)
    tauh = 1/(tfi*(a + beth(v))) : estimation of inactivation tau
    hinf = a/(a+beth(v))         : estimation of inactivation steady state
}
```
