## Supplementary NEURON models for "Dendritic excitations govern back-propagation via a spike-rate accelerometer": gabaa.mod

```
TITLE minimal model of GABAa receptors

COMMENT
-----------------------------------------------------------------------------

	Minimal kinetic model for GABA-A receptors
	==========================================

  Model of Destexhe, Mainen & Sejnowski, 1994:

	(closed) + T <-> (open)

  The simplest kinetics are considered for the binding of transmitter (T)
  to open postsynaptic receptors.   The corresponding equations are in
  similar form as the Hodgkin-Huxley model:

	dr/dt = alpha * [T] * (1-r) - beta * r

	I = gmax * [open] * (V-Erev)

  where [T] is the transmitter concentration and r is the fraction of 
  receptors in the open form.

  If the time course of transmitter occurs as a pulse of fixed duration,
  then this first-order model can be solved analytically, leading to a very
  fast mechanism for simulating synaptic currents, since no differential
  equation must be solved (see Destexhe, Mainen & Sejnowski, 1994).

-----------------------------------------------------------------------------

  Based on voltage-clamp recordings of GABAA receptor-mediated currents in rat
  hippocampal slices (Otis and Mody, Neuroscience 49: 13-32, 1992), this model
  was fit directly to experimental recordings in order to obtain the optimal
  values for the parameters (see Destexhe, Mainen and Sejnowski, 1996).

-----------------------------------------------------------------------------

  This mod file includes a mechanism to describe the time course of transmitter
  on the receptors.  The time course is approximated here as a brief pulse
  triggered when the presynaptic compartment produces an action potential.
  The pointer "pre" represents the voltage of the presynaptic compartment and
  must be connected to the appropriate variable in oc.

-----------------------------------------------------------------------------

  See details in:

  Destexhe, A., Mainen, Z.F. and Sejnowski, T.J.  An efficient method for
  computing synaptic conductances based on a kinetic model of receptor binding
  Neural Computation 6: 10-14, 1994.  

  Destexhe, A., Mainen, Z.F. and Sejnowski, T.J.  Kinetic models of 
  synaptic transmission.  In: Methods in Neuronal Modeling (2nd edition; 
  edited by Koch, C. and Segev, I.), MIT press, Cambridge, 1996.


  Written by Alain Destexhe, Laval University, 1995

-----------------------------------------------------------------------------

  Modified by D. Itkis, Cohen Lab, Harvard University, 2023.
  Replaced prethreshold with a netcom that takes presynaptic spike trains.
  Changed gmax to g, g to g_inst for compatibility with other synapse types.
  
-----------------------------------------------------------------------------
ENDCOMMENT


INDEPENDENT {t FROM 0 TO 1 WITH 1 (ms)}

NEURON {
	POINT_PROCESS GABAa
	RANGE C, R, R0, R1, g_inst, g, lastrelease
	NONSPECIFIC_CURRENT i
	GLOBAL Cmax, Cdur, Alpha, Beta, Erev, Deadtime, Rinf, Rtau
}

UNITS {
	(nA) = (nanoamp)
	(mV) = (millivolt)
	(umho) = (micromho)
	(mM) = (milli/liter)
}

PARAMETER {
	Cmax	= 1	(mM)		: max transmitter concentration
	Cdur	= 1	(ms)		: transmitter duration (rising phase)
	Alpha	= 5	(/ms mM)	: forward (binding) rate
	Beta	= 0.18	(/ms)		: backward (unbinding) rate
	Erev	= -80	(mV)		: reversal potential
	Deadtime = 1	(ms)		: mimimum time between release events
	g		(umho)		: maximum conductance
}

ASSIGNED {
	v		(mV)		: postsynaptic voltage
	i 		(nA)		: current = g*(v - Erev)
	g_inst 		(umho)		: conductance
	C		(mM)		: transmitter concentration
	R				: fraction of open channels
	R0				: open channels at start of release
	R1				: open channels at end of release
	Rinf				: steady state channels open
	Rtau		(ms)		: time constant of channel binding
	lastrelease	(ms)		: time of last spike
}

INITIAL {
	R = 0
	C = 0
	Rinf = Cmax*Alpha / (Cmax*Alpha + Beta)
	Rtau = 1 / ((Alpha * Cmax) + Beta)
	lastrelease = -1000
}

BREAKPOINT {
	SOLVE release
	g_inst = g * R
	i = g_inst*(v - Erev)
}

PROCEDURE release() { LOCAL q

	q = ((t - lastrelease) - Cdur)		: time since last release ended

	if (q < 0) {			: still releasing?
		: do nothing
	} else if (C == Cmax) {			: in dead time after release
		R1 = R
		C = 0.
	}

	if (C > 0) {				: transmitter being released?
	   R = Rinf + (R0 - Rinf) * exptable (- (t - lastrelease) / Rtau)		
	} else {				: no release occurring
  	   R = R1 * exptable (- Beta * (t - (lastrelease + Cdur)))
	}

	VERBATIM
	return 0;
	ENDVERBATIM
}

NET_RECEIVE(weight (uS)) {
	g = weight
	if (t - lastrelease > Deadtime) {
		C = Cmax
		R0 = R
		lastrelease = t
	}
}

FUNCTION exptable(x) { 
	TABLE  FROM -10 TO 10 WITH 2000

	if ((x > -10) && (x < 10)) {
		exptable = exp(x)
	} else {
		exptable = 0.
	}
}
```
