## Supplementary NEURON models for "Dendritic excitations govern back-propagation via a spike-rate accelerometer": gabab.mod

```
TITLE minimal model of GABAB receptors

COMMENT
-----------------------------------------------------------------------------

	Minimal kinetic model for GABA-B receptors
	==========================================

	Minimal model of GABAB currents including nonlinear stimulus 
	dependency (fundamental to take into account for GABAB receptors).


	Features:

  	  - peak at 100 ms; time course fit from experimental PSC
	  - NONLINEAR SUMMATION (psc is much stronger with bursts)
	    due to cooperativity of G-protein binding on K+ channels

	Approximations:

	  - single binding site on receptor	
	  - model of alpha G-protein activation (direct) of K+ channel
	  - G-protein dynamics is second-order; simplified as follows:
		- saturating receptor
		- no desensitization
		- Michaelis-Menten of receptor for G-protein production
		- "resting" G-protein is in excess
		- Quasi-stat of intermediate enzymatic forms
	  - binding on K+ channel is fast


	Kinetic Equations:

	  dR/dt = K1 * T * (1-R) - K2 * R

	  dG/dt = K3 * R - K4 * G

	  R : activated receptor
	  T : transmitter
	  G : activated G-protein
	  K1,K2,K3,K4 = kinetic rate cst

  n activated G-protein bind to a K+ channel:

	n G + C <-> O		(Alpha,Beta)

  If the binding is fast, the fraction of open channels is given by:

	O = G^n / ( G^n + KD )

  where KD = Beta / Alpha is the dissociation constant

-----------------------------------------------------------------------------

  Based on voltage-clamp recordings of GABAB receptor-mediated currents in rat
  hippocampal slices (Otis et al, J. Physiol. 463: 391-407, 1993), this model 
  was fit directly to experimental recordings in order to obtain the optimal
  values for the parameters (see Destexhe and Sejnowski, 1995).

-----------------------------------------------------------------------------

  This mod file includes a mechanism to describe the time course of transmitter
  on the receptors.  The time course is approximated here as a brief pulse
  triggered when the presynaptic compartment produces an action potential.
  The pointer "pre" represents the voltage of the presynaptic compartment and
  must be connected to the appropriate variable in oc.

-----------------------------------------------------------------------------

  See details in:

  Destexhe, A. and Sejnowski, T.J.  G-protein activation kinetics and
  spill-over of GABA may account for differences between inhibitory responses
  in the hippocampus and thalamus.  Proc. Natl. Acad. Sci. USA  92:
  9515-9519, 1995.

  See also: 

  Destexhe, A., Mainen, Z.F. and Sejnowski, T.J.  Kinetic models of 
  synaptic transmission.  In: Methods in Neuronal Modeling (2nd edition; 
  edited by Koch, C. and Segev, I.), MIT press, Cambridge, 1996.


  Written by Alain Destexhe, Laval University, 1995

-----------------------------------------------------------------------------

  Modified by D. Itkis, Cohen Lab, Harvard University, 2023.
  Replaced prethreshold with a netcom that takes presynaptic spike trains.
  Changed gmax to g, g to g_inst for compatibility with other synapse types.

-----------------------------------------------------------------------------
ENDCOMMENT


INDEPENDENT {t FROM 0 TO 1 WITH 1 (ms)}

NEURON {
	POINT_PROCESS GABAb
	RANGE C, R, G, g_inst, g, lastrelease
	NONSPECIFIC_CURRENT i
	GLOBAL Cmax, Cdur, Deadtime, K1, K2, K3, K4, KD, Erev
}
UNITS {
	(nA) = (nanoamp)
	(mV) = (millivolt)
	(umho) = (micromho)
	(mM) = (milli/liter)
}

PARAMETER {
	Cmax	= 1	(mM)		: max transmitter concentration
	Cdur	= 1	(ms)		: transmitter duration (rising phase)
	Deadtime = 1	(ms)		: mimimum time between release events
	K1	= 0.09	(/ms mM)	: forward binding rate to receptor
	K2	= 0.0012 (/ms)		: backward (unbinding) rate of receptor
	K3	= 0.18 (/ms)		: rate of G-protein production
	K4	= 0.034 (/ms)		: rate of G-protein decay
	KD	= 100			: dissociation constant of K+ channel
	n	= 4			: nb of binding sites of G-protein on K+
	Erev	= -95	(mV)		: reversal potential (E_K)
	g		(umho)		: maximum conductance
}

ASSIGNED {
	v		(mV)		: postsynaptic voltage
	i 		(nA)		: current = g*(v - Erev)
	g_inst 		(umho)		: conductance
	C		(mM)		: transmitter concentration
	Gn
	lastrelease	(ms)		: time of last spike
}

STATE {
	R				: fraction of activated receptor
	G				: fraction of activated G-protein
}

INITIAL {
	C = 0
	lastrelease = -1000
	R = 0
	G = 0
}

BREAKPOINT {
	SOLVE bindkin METHOD euler
	Gn = G^n
	g_inst = g * Gn / (Gn+KD)
	i = g_inst*(v - Erev)
}

DERIVATIVE bindkin {
	release()		: evaluate the variable C
	R' = K1 * C * (1-R) - K2 * R
	G' = K3 * R - K4 * G
}

PROCEDURE release() { LOCAL q
	q = ((t - lastrelease) - Cdur)		: time since last release ended
	if (q < 0) {				: still releasing?
		: do nothing
	} else if (C == Cmax) {			: in dead time after release
		C = 0.
	}
}

NET_RECEIVE(weight (uS)) {
	g = weight
	if (t - lastrelease > Deadtime) {
		C = Cmax
		lastrelease = t
	}
}
```
