## Supplementary NEURON models for "Dendritic excitations govern back-propagation via a spike-rate accelerometer": hd.mod

```
TITLE I-h channel from Magee 1998 for distal dendrites
: https://modeldb.science/156120?tab=2

UNITS {
	(mA) = (milliamp)
	(mV) = (millivolt)

}

PARAMETER {
	v 		           (mV)
  ehd = -30      		   (mV)        
	celsius 	       (degC)
	ghdbar=1e-4 	   (Siemens/cm2)
  vhalfl=-81       (mV)
	kl=-7
  vhalft=-75   	   (mV)
  a0t=0.011        (/ms)
  zetat=2.2        (1)
  gmt=.4   	       (1)
	q10=4.5
	qtl=1
        ivalue = 0 (mA/cm2)
}


NEURON {
	SUFFIX hd
	NONSPECIFIC_CURRENT i
        RANGE ghdbar, vhalfl, ivalue
        GLOBAL linf,taul
}

STATE {
        l
}

ASSIGNED {
	i (mA/cm2)
        linf      
        taul
        ghd
}

INITIAL {
	rate(v)
	l=linf
}


BREAKPOINT {
	SOLVE states METHOD cnexp
	ghd = ghdbar*l
	i = ghd*(v-ehd)
        ivalue = i

}


FUNCTION alpt(v(mV)) {
  alpt = exp(0.0378*zetat*(v-vhalft)) 
}

FUNCTION bett(v(mV)) {
  bett = exp(0.0378*zetat*gmt*(v-vhalft)) 
}

DERIVATIVE states {     : exact when v held constant; integrates over dt step
        rate(v)
        l' =  (linf - l)/taul
}

PROCEDURE rate(v (mV)) { :callable from hoc
        LOCAL a,qt
        qt=q10^((celsius-33)/10)
        a = alpt(v)
        linf = 1/(1 + exp(-(v-vhalfl)/kl))
:       linf = 1/(1+ alpl(v))
        taul = bett(v)/(qtl*qt*a0t*(1+a))
}
```
