## Supplementary NEURON models for "Dendritic excitations govern back-propagation via a spike-rate accelerometer": kca.mod

```
TITLE Slow Ca-dependent potassium current
:
:   Ca++ dependent K+ current IC responsible for slow AHP
:   Differential equations
:
:   Model based on a first order kinetic scheme
:
:       + n cai <->     (alpha,beta)
:
:   Following this model, the activation fct will be half-activated at 
:   a concentration of Cai = (beta/alpha)^(1/n) = cac (parameter)
:
:   The mod file is here written for the case n=2 (2 binding sites)
:   ---------------------------------------------
:
:   This current models the "slow" IK[Ca] (IAHP): 
:      - potassium current
:      - activated by intracellular calcium
:      - NOT voltage dependent
:
:   A minimal value for the time constant has been added
:
:   Ref: Destexhe et al., J. Neurophysiology 72: 803-818, 1994.
:   See also: http://www.cnl.salk.edu/~alain , http://cns.fmed.ulaval.ca
:   modifications by Yiota Poirazi 2001
:   taumin = 0.5 ms instead of 0.1 ms	

NEURON {
    SUFFIX kca
    USEION k READ ek WRITE ik
    USEION ca READ cai
    RANGE gk, gbar, m_inf, tau_m, gmax
    GLOBAL beta, cac
}

UNITS {
    (mA) = (milliamp)
    (mV) = (millivolt)
    (molar) = (1/liter)
    (mM) = (millimolar)
}

PARAMETER {
    v                (mV)
    celsius = 36     (degC)
    ek      = -80    (mV)
    cai     = 2.4e-5 (mM)           : initial [Ca]i
    gbar    = 0.01   (mho/cm2)
    beta    = 0.03   (1/ms)          : backward rate constant
    cac     = 0.025  (mM)            : middle point of activation fct
    taumin  = 0.5    (ms)            : minimal value of the time cst
}

STATE {m}        : activation variable to be solved in the DEs       

ASSIGNED {       : parameters needed to solve DE 
    ik      (mA/cm2)
    m_inf
    tau_m   (ms)
    tadj
    gk      (mho/cm2)
    gmax    (mho/cm2)
}
BREAKPOINT { 
    SOLVE states METHOD derivimplicit
    gk = gbar*m*m*m     : maximum channel conductance
    ik = gk*(v - ek)    : potassium current induced by this channel
    if (gk > gmax) {
        gmax = gk
    }
}

DERIVATIVE states { 
    evaluate_fct(v,cai)
    m' = (m_inf - m) / tau_m
}

INITIAL {
    :
    :  activation kinetics are assumed to be at 22 deg. C
    :  Q10 is assumed to be 3
    :
    tadj = 3 ^ ((celsius-22.0(degC))/10(degC)) : temperature-dependent adjastment factor
    evaluate_fct(v,cai)
    m = m_inf
    gk = gbar*m*m*m
    gmax = gk
}

PROCEDURE evaluate_fct(v(mV),cai(mM)) {  LOCAL car
    car = (cai/cac)^2
    m_inf = car / ( 1 + car )      : activation steady state value
    tau_m =  1 / beta / (1 + car) / tadj
    if(tau_m < taumin) { tau_m = taumin }   : activation min value of time cst
}
```
