## Supplementary NEURON models for "Dendritic excitations govern back-propagation via a spike-rate accelerometer": km.mod

```
COMMENT
km.mod
Potassium channel, Hodgkin-Huxley style kinetics
Based on I-M (muscarinic K channel)
Slow, noninactivating
Author: Zach Mainen, Salk Institute, 1995,

ENDCOMMENT

NEURON {
	  SUFFIX km
	  USEION k READ ek WRITE ik
	  RANGE n, gk, gbar, gmax, ikvalue
	  RANGE ninf, ntau
	  GLOBAL Ra, Rb
	  GLOBAL q10, temp, tadj, vmin, vmax
}

UNITS {
	  (mA) = (milliamp)
	  (mV) = (millivolt)
	  (pS) = (picosiemens)
	  (um) = (micron)
} 

PARAMETER {
	  v 		        (mV)
	  gbar = 10   	(pS/um2)              : 0.03 mho/cm2
	  tha  = -30	  (mV)                  : v 1/2 for inf
	  qa   = 9	    (mV)                  : inf slope		
	  Ra   = 0.001	(/ms/mV)              : max act rate  (slow)
	  Rb   = 0.001	(/ms/mV)              : max deact rate  (slow)
	  celsius		    (degC)
	  temp = 23	    (degC)                : original temp 	
	  q10  = 2.3                          : temperature sensitivity
	  vmin = -120	  (mV)
	  vmax = 100	  (mV)
	  ikvalue = 0    (mA/cm2)
} 


ASSIGNED {
	  a		 (/ms)
	  b		 (/ms)
	  ik 	 (mA/cm2)
	  gk	 (pS/um2)
	  ek   (mV)
	  ninf
	  ntau (ms)	
	  tadj
    gmax (mho/cm2)
}


STATE { n }

INITIAL { 
	  trates(v)
	  n = ninf
    gk = tadj*gbar*n
    gmax = (1e-4) * gk
}

BREAKPOINT {
    SOLVE states METHOD cnexp
	  gk = tadj*gbar*n
	  ik = (1e-4) * gk * (v - ek)
	  ikvalue = ik
    if (((1e-4) * gk) > gmax) {
        gmax = (1e-4) * gk
    }
} 

LOCAL nexp

DERIVATIVE states {   
    trates(v)     
    n' =  (ninf-n)/ntau*tadj
}

PROCEDURE trates(v (mV)) {  :Computes rate and other constants at current v.
    :Call once from HOC to initialize inf at resting v.
    TABLE ninf, ntau
	  DEPEND celsius, temp, Ra, Rb, tha, qa
	  FROM vmin TO vmax WITH 199
    
	  rates(v): not consistently executed from here if usetable_hh == 1
    tadj = q10^((celsius - temp)/10(degC))  :temperature adjastment
}


PROCEDURE rates(v (mV)) {  :Computes rate and other constants at current v.
    :Call once from HOC to initialize inf at resting v.
    
    a =  Ra * (v - tha) / (1 - exp(-(v - tha)/qa))
    b = -Rb * (v - tha) / (1 - exp((v - tha)/qa))
    ntau = 1/(a+b)
	  ninf = a*ntau
}
```
