## Supplementary NEURON models for "Dendritic excitations govern back-propagation via a spike-rate accelerometer": nap.mod

```
TITLE  Na persistent channel
: used in distal oblique dendrites to assist Ca spike initiation  
: a typo in the exponential function was pointed out by Michele Migliore and
: corrected by Yiota Poirazi on December 4th, 2003
NEURON {
	  SUFFIX nap
	  USEION na READ ena WRITE ina
    RANGE  gnabar,vhalf, K, g, gmax, inavalue
}

UNITS {
	  (mA) = (milliamp)
	  (mV) = (millivolt)
}

PARAMETER { : parameters that can be entered when function is called in cell-setup 
	  v               (mV)
    ena = 50        (mV) : Na reversal potential  (reset in cell-setup.hoc)
	  K = 1           (mV)              : slope of steady state variable
    :	gnabar = 0.001e-2 (mho/cm2) : suggested conductance, 1 percent of the transient Na current
	  gnabar = 0      (mho/cm2)          : initialized conductance
	  vhalf  = -51.90 (mV)                : half potential
    inavalue = 0   (mA/cm2)            : reference current as range variable
}	

STATE { n }

ASSIGNED {
	  ina  (mA/cm2)
    g    (mho/cm2)
    gmax (mho/cm2)
}

INITIAL {
    :	SOLVE states not used
    gmax = 0
}

BREAKPOINT {
    states(v)
    g = gnabar*n*n*n
	  ina = g*(v-ena)
      inavalue = ina
    if (g > gmax) {
        gmax = g
    }
}

PROCEDURE states(v (mV)) {     : exact when v held constant; integrates over dt step
    :        n = 1 / (1 + (exp(vhalf - v))/K) : steady state value !!!typo in the exponential function!!!
    TABLE n DEPEND vhalf, K FROM -150 TO 150 WITH 300     
    n = 1 / (1 + exp((vhalf - v)/K)) : steady state value
}
```
