## Supplementary NEURON models for "Dendritic excitations govern back-propagation via a spike-rate accelerometer": nmda_andr.mod

```
COMMENT

AMPA synapse with two decay time constants based on the exp2syn.mod
mechanism from the NEURON distribution.

We use the time constants measured by Andrasfalvy & Magee (2001) from
patches excised from different parts of the stratum radiatum at
22\degC.  These are in broad agreement with the time constants measured
by Spruston &al (1995), epecially when the age-dependent change in
receptor properties (Hestrin &al., 1992) is taken into account.
Although they are considerably slower than the time constants measured
using whole-cell recordings from hippocampal cells (Hestrin & al., 1990),
this might be accounted for by the more depolarised potentials used in
these recordings.  We corrected the time constants recorded at 22\degC
to 34\degC using a \Qten of 3 (Hestrin & al., 1990).  This gave a
rise time of 1.7\ms and a dual exponential decay comprising a fast
component fast time constant of 67\ms and a slow component with a time
constant of 428\ms in the ratio 0.61 to 0.39. 

References

B K Andr\'asfalvy & J C Magee 2001 "Distance-Dependent Increase in
  AMPA Receptor Number in the Dendrites of Adult Hippocampal CA1
  Pyramidal Neurons" J. Neurosci. 21, 9151-9159

S Hestrin & al. 1990 "Mechanisms Generating the time course of dual
  component excitatory synaptic currents recorded in hippocampal
  slices" Neuron 5, 247-253

S Hestrin 1992 "Developmental regulation of NMDA receptor-mediated
  currents at a central synapse" Nature 357, 686-689

N Spruston & al. 1995 "Dendritic glutamate receptor channels in rat
  hippocampal CA3 and CA1 neurons" J. Physiol. 482, 325-352

ENDCOMMENT

NEURON {
    POINT_PROCESS NmdaSyn
    RANGE  e, g, i, b, ica
    NONSPECIFIC_CURRENT i
    USEION ca READ cai,cao WRITE ica
    GLOBAL total, mg, q10, taurise, taufast, tauslow, taurise_exp, taufast_exp, tauslow_exp, afast, aslow, normfac, T_exp, K0, delta, fracca
}

INCLUDE "units.inc"

PARAMETER {
    : Time constants and afast from Andrasfalvy & Magee 01
    taurise_exp =    6.46 (ms) <1e-9,1e9>    : rise      
    taufast_exp =  252.5  (ms) <1e-9,1e9>    : fast decay
    tauslow_exp = 1660    (ms) <1e-9,1e9>    : slow decay
    afast = 0.61 <0,1>
    e=0	(mV)
    mg	= 1    (mM)		: external magnesium concentration
    fracca= 0.13        : fraction of current that is ca ions; Srupuston &al 95
    z = 2
    celsius = 22	(degC)
    T_exp = 22    (degC)
    q10 = 3       : Hestrin 90
    K0 = 4.1 (mM) : From Spruston &al 95
    delta = 0.8   : From Spruston &al 95
}

ASSIGNED {
    v       (mV)
    i       (nA)
    ica	    (nA) 	
    g       (uS)
    aslow 
    total   (uS)
    cai     (mM)
    cao     (mM)
    taurise (ms)
    taufast (ms)
    tauslow (ms)
    normfac 
    b
}

BREAKPOINT {
    SOLVE state METHOD cnexp
    b = mgblock(v)		: b is the block by magnesium at this voltage
    g = (B + C - A) * b
    i =   g * (1-fracca) * (v - e)
    ica = g * fracca     * ghkg(v,cai,cao,z)
}

INCLUDE "triexpsyn.inc"

INCLUDE "ghk.inc"

FUNCTION mgblock(v(mV)) {
    TABLE 
    DEPEND mg, K0, delta, celsius
    FROM -140 TO 80 WITH 1000
    : From Spruston &al 95
    mgblock = 1/(1+(mg/K0)*exp(-delta*z*FARADAY*v*(0.001)/R/(celsius+273)))
}
```
