## Supplementary NEURON models for "Dendritic excitations govern back-propagation via a spike-rate accelerometer": somacar.mod

```
TITLE Ca R-type channel with medium threshold for activation
: used in somatic regions. It has lower threshold for activation/inactivation
: and slower activation time constant
: than the same mechanism in dendritic regions
: uses channel conductance (not permeability)
: written by Yiota Poirazi on 3/12/01

NEURON {
	  SUFFIX somacar
	  USEION ca READ eca WRITE ica
    RANGE gcabar, m, h, g, gmax
	  RANGE inf, tau
}

UNITS {
	  (mA) = (milliamp)
	  (mV) = (millivolt)
}

PARAMETER {      : parameters that can be entered when function is called in cell-setup
    v               (mV)
 	  celsius = 34	  (degC)
    gcabar = 0      (mho/cm2) : initialized conductance
	  eca = 140       (mV)      : Ca++ reversal potential
}

STATE {	m h }   : unknown activation and inactivation parameters to be solved in the DEs

ASSIGNED {               : parameters needed to solve DE
	  ica    (mA/cm2)
    inf[2]
	  tau[2] (ms)
    g      (mho/cm2)
    gmax   (mho/cm2)
}

BREAKPOINT {
	  SOLVE states METHOD cnexp
    g = gcabar*m*m*m*h
	  ica = g*(v - eca)
    if (g > gmax) {
        gmax = g
    }
}

INITIAL {
    mhn(v)
    m = inf[0]
    h = inf[1]
    g = gcabar*m*m*m*h
    ica = g*(v - eca) : initial Ca++ current value
    gmax = g
}

DERIVATIVE states {
	  mhn(v)
	  m' = (inf[0] - m)/tau[0]
	  h' = (inf[1] - h)/tau[1]
}	

FUNCTION varss(v (mV), i) {
	  if (i==0) {
	      varss = 1 / (1 + exp((v+60(mV))/(-3(mV)))) :Ca activation
	  }
	  else if (i==1) {
        varss = 1/ (1 + exp((v+62(mV))/(1(mV))))   :Ca inactivation
	  }
}

FUNCTION vartau(v (mV), i) (ms) {
	  if (i==0) {
        vartau = 100  : activation variable time constant
    }
	  else if (i==1) {
        vartau = 5    : inactivation variable time constant
    }
}	

PROCEDURE mhn(v (mV)) {LOCAL a, b :rest = -70
	  TABLE inf, tau DEPEND celsius FROM -100 TO 100 WITH 200
	  FROM i=0 TO 1 {
		    tau[i] = vartau(v,i) 
		    inf[i] = varss(v,i)
	  }
}
```
