## Supplementary NEURON models for "Dendritic excitations govern back-propagation via a spike-rate accelerometer": cheriff.mod

```
TITLE CheRiff model of light-activated current 
: based on All-optical electrophysiology in mammalian neurons using engineered microbial
: rhodopsins, Nature Methods, 11, pages 825–833 (2014)
: Added by D. Itkis 2023.

NEURON {
    SUFFIX cheriff
    NONSPECIFIC_CURRENT i
    RANGE i, g, e, gmax, ton, toff, tdes, ipeak, isteady_state, lightintensity
}

PARAMETER {
    gmax = 1e-2 (siemens/cm2)
    e = 0 (millivolt)
    ton = 4.5 (ms) : 4.5 ms for 500 mW/cm2
    toff = 16 (ms)
    tdes = 400 (ms)
    ipeak = 2.0 
    isteady_state = 1.33 
    lightintensity = 0 : Half activation intensity relative
}

STATE {
    off peak steady_state
}

ASSIGNED {
    v (millivolt)
    i (milliamp/cm2)
    g (siemens/cm2)
    kon (1/ms)
    koff (1/ms)
    kdes (1/ms)
}

INITIAL {
    off = 1
    peak = 0
    steady_state = 0
    kon = 1/ton         : On-rate constant
    koff = 1/toff        : Off-rate constant
    kdes = 1/tdes         : Desensitization rate constant
}

BREAKPOINT { 
    g = (ipeak * peak + isteady_state * steady_state) * gmax
    i = (ipeak * peak + isteady_state * steady_state) * gmax * (v - e)
}

KINETIC state {
    ~ off <-> peak (kon*lightintensity, kdes)
    ~ peak <-> steady_state (kdes, koff)
    CONSERVE off + peak + steady_state = 1
}
```
