## Supplementary NEURON models for "Dendritic excitations govern back-propagation via a spike-rate accelerometer": chr.mod

```
TITLE Distributed conductance stimulus as
: model for CheRiff channelrhodopsin 
: Added by D. Itkis 2023.

NEURON {
 SUFFIX chr
 NONSPECIFIC_CURRENT i
 RANGE i, e, g, gexp, del, dur
}
PARAMETER {
del = 5 (ms)
dur = 75 (ms)
g = 0.00 (siemens/cm2)
e = 0 (millivolt)
gexp = 1
}
ASSIGNED {
 i (milliamp/cm2)
 v (millivolt)
}
BREAKPOINT { 
at_time(del)
at_time(del+dur)
if (t < del + dur && t > del) {
i = (v-e)*g*gexp
} else {
i = 0
}
}
```
