## Supplementary NEURON models for "Dendritic excitations govern back-propagation via a spike-rate accelerometer": gclamp.mod

```
TITLE Point-like conductance clamp
: Added by D. Itkis 2023.

NEURON {
POINT_PROCESS gclamp
NONSPECIFIC_CURRENT i
RANGE del, dur, g, i, e 
}
UNITS { (nA) = (nanoamp) }
PARAMETER {
del = 5(ms)
dur = 20 (ms)
g = 0.001 (1/megaohm)
e = 0 (millivolt)
}
ASSIGNED { 
   i (nA)
   v (millivolt) }
INITIAL { i = 0 }
BREAKPOINT {
at_time(del)
at_time(del+dur)
if (t < del + dur && t > del) {
i = (v-e)*g
} else {
i = 0
}
}
```
