## Supplementary NEURON models for "Dendritic excitations govern back-propagation via a spike-rate accelerometer": kadist.mod

```
TITLE K-A channel from Klee Ficker and Heinemann
: modified to account for Dax A Current ----------
: M.Migliore Jun 1997

UNITS {
	(mA) = (milliamp)
	(mV) = (millivolt)
	(mS) = (millisiemens)

}

INDEPENDENT {t FROM 0 TO 1 WITH 1 (ms)}

PARAMETER {
        dt (ms)
	v (mV)
        ek = -90 (mV)              : must be explicitely def. in hoc
	celsius = 24	(degC)
	gkabar=.008 (mho/cm2)
        vhalfn=-1   (mV)
        vhalfl=-56   (mV)
        a0l=0.05      (/ms)
        a0n=.1    (/ms)
        zetan=-1.8    (1)
        zetal=3    (1)
        gmn=0.39   (1)
        gml=1   (1)
	lmin=2  (ms)
	nmin=0.1  (ms)
	pw=-1    (1)
	tq=-40 (mV)
	qq=5  (mV)
	q10=5 
	qtl=1
        nscale=1
        lscale=1
}


NEURON {
	SUFFIX kad
	USEION k READ ek WRITE ik
        RANGE gkabar,gka,ik
        RANGE ninf,linf,taul,taun
        GLOBAL lmin,nscale,lscale
}

STATE {
	n
        l
}

ASSIGNED {
	ik (mA/cm2)
        ninf
        linf      
        taul   (ms)
        taun   (ms)
        gka    (mho/cm2)
        qt     
}

INITIAL {
        rates(v)
        n=ninf
        l=linf
        gka = gkabar*n*l
	ik = gka*(v-ek)
}        

BREAKPOINT {
	SOLVE states METHOD cnexp
	gka = gkabar*n*l
	ik = gka*(v-ek)
}

DERIVATIVE states {
        rates(v)
        n' = (ninf-n)/taun
        l' = (linf-l)/taul
}

FUNCTION alpn(v(mV)) {
LOCAL zeta
  zeta=zetan+pw/(1+exp((v-tq)/qq))
  alpn = exp(1.e-3*zeta*(v-vhalfn)*9.648e4 (degC/mV)/(8.315*(273.16+celsius))) 
}

FUNCTION betn(v(mV)) {
LOCAL zeta
  zeta=zetan+pw/(1+exp((v-tq)/qq))
  betn = exp(1.e-3*zeta*gmn*(v-vhalfn)*9.648e4 (degC/mV)/(8.315*(273.16+celsius))) 
}

FUNCTION alpl(v(mV)) {
  alpl = exp(1.e-3*zetal*(v-vhalfl)*9.648e4 (degC/mV)/(8.315*(273.16+celsius))) 
}

FUNCTION betl(v(mV)) {
  betl = exp(1.e-3*zetal*gml*(v-vhalfl)*9.648e4 (degC/mV)/(8.315*(273.16+celsius))) 
}
LOCAL facn,facl

:if state_borgka is called from hoc, garbage or segmentation violation will
:result because range variables won't have correct pointer.  This is because
: only BREAKPOINT sets up the correct pointers to range variables.
:PROCEDURE states() {     : exact when v held constant; integrates over dt step
:        rates(v)
:        n = n + facn*(ninf - n)
:        l = l + facl*(linf - l)
:        VERBATIM
:        return 0;
:        ENDVERBATIM
:}

PROCEDURE rates(v (mV)) { :callable from hoc
        LOCAL a,qt
        qt=q10^((celsius-24)/10 (degC))
        a = alpn(v)
        ninf = 1/(1 + a)
        taun = betn(v)/(qt*a0n*(1+a))
        taun = taun/nscale
	if (taun<nmin) {taun=nmin}
        facn = (1 - exp(-dt/taun))
        a = alpl(v)
        linf = 1/(1+ a)
	taul = 0.26(ms/mV)*(v+50)/qtl
        taul = taul/lscale
	if (taul<lmin/qtl) {taul=lmin/qtl}
        facl = (1 - exp(-dt/taul))
}
```
