## Supplementary NEURON models for "Dendritic excitations govern back-propagation via a spike-rate accelerometer": kdrca1.mod

```
TITLE K-DR channel
: from Klee Ficker and Heinemann
: modified to account for Dax et al.
: M.Migliore 1997

UNITS {
	(mA) = (milliamp)
	(mV) = (millivolt)

}

INDEPENDENT {t FROM 0 TO 1 WITH 1 (ms)}

PARAMETER {
        dt (ms)
	v (mV)
        ek = -90	(mV)	: must be explicitely def. in hoc
	celsius = 24	(degC)
	gkdrbar=.003 (mho/cm2)
        ikmax = 0.3 (mA/cm2)
        vhalfn=13   (mV)
        a0n=0.02      (/ms)
        zetan=-3    (1)
        gmn=0.7  (1)
	nmax=2  (ms)
	q10=1
        nscale=1
}


NEURON {
	SUFFIX kdr
	USEION k READ ek WRITE ik
        RANGE gkdr,gkdrbar,ik
	RANGE ninf,taun
        GLOBAL nscale
}

STATE {
	n
}

ASSIGNED {
	ik (mA/cm2)
        ninf
        gkdr (mho/cm2)
        taun (ms)
}

INITIAL {
        rates(v)
        n=ninf
	gkdr = gkdrbar*n
	ik = gkdr*(v-ek)
}        

BREAKPOINT {
	SOLVE states METHOD cnexp
	gkdr = gkdrbar*n
	ik = gkdr*(v-ek)
:       if (ik>ikmax) { ik=ikmax }
}

DERIVATIVE states {
        rates(v)
        n' = (ninf-n)/taun
}

FUNCTION alpn(v(mV)) {
  alpn = exp(1.e-3*zetan*(v-vhalfn)*9.648e4(degC/mV)/(8.315*(273.16+celsius))) 
}

FUNCTION betn(v(mV)) {
  betn = exp(1.e-3*zetan*gmn*(v-vhalfn)*9.648e4(degC/mV)/(8.315*(273.16+celsius))) 
}

LOCAL facn

:if state_borgka is called from hoc, garbage or segmentation violation will
:result because range variables won't have correct pointer.  This is because
: only BREAKPOINT sets up the correct pointers to range variables.
:PROCEDURE states() {     : exact when v held constant; integrates over dt step
:        rates(v)
:        n = n + facn*(ninf - n)
:        VERBATIM
:        return 0;
:        ENDVERBATIM
:}

PROCEDURE rates(v (mV)) { :callable from hoc
        LOCAL a,qt
        qt=q10^((celsius-24)/10(degC))
        a = alpn(v)
        ninf = 1/(1+a)
        taun = betn(v)/(qt*a0n*(1+a))
	if (taun<nmax) {taun=nmax}
        taun=taun/nscale
        facn = (1 - exp(-dt/taun))
}
```
