## Supplementary NEURON models for "Dendritic excitations govern back-propagation via a spike-rate accelerometer": na3.mod

```
TITLE na3
: Na current modified by D.Itkis 3/23
: from Jeff M.
:  ---------- modified -------M.Migliore may97

INDEPENDENT {t FROM 0 TO 1 WITH 1 (ms)}

NEURON {
	SUFFIX na3
	USEION na READ ena WRITE ina
	RANGE  gbar, ar2, ina, thegna, smin
	RANGE  minf, hinf, mtau, htau, sinf, taus, tauss
        GLOBAL qinf, thinf, mscale, hscale, sscale
}

PARAMETER {
	gbar = 0.010   	(mho/cm2)	
								
	tha  =  -30	(mV)		: v 1/2 for act	
	qa   = 7.2	(mV)		: act slope (4.5)		
	Ra   = 0.4	(/ms)		: open (v)		
	Rb   = 0.124 	(/ms)		: close (v)		

	thi1  = -45	(mV)		: v 1/2 for inact 	
	thi2  = -45 	(mV)		: v 1/2 for inact 	
	qd   = 1.5	(mV)	        : inact tau slope
	qg   = 1.5      (mV)
	mmin=0.02	(ms)
	hmin=0.5	(ms)		
	q10=2
	Rg   = 0.01 	(/ms)		: inact recov (v) 	
	Rd   = .03 	(/ms)		: inact (v)	
	qq   = 10        (mV)
	tq   = -55      (mV)

	thinf  = -50 	(mV)		: inact inf slope	
	qinf  = 4 	(mV)		: inact inf slope 

        vhalfs=-60	(mV)		: slow inact.
        a0s=0.0003	(/ms)		: a0s=b0s
:        a0s=0.00003	(/ms)		: a0s=b0s
        zetas=12	(1)
        gms=0.2		(1)
        smin=10		(ms)
        vvh=-58		(mV) 
        vvs=2		(mV)
        ar2=1		(1)		: 1=no inact., 0=max inact.
	ena = 55	(mV)            : must be explicitly def. in hoc

	v 		(mV)
	dt		(ms)
	celsius=24	(degC)

        mscale = 1
        hscale = 1
        sscale = 1
}


UNITS {
	(mA) = (milliamp)
	(mV) = (millivolt)
	(pS) = (picosiemens)
	(um) = (micron)
} 

ASSIGNED {
	ina 		(mA/cm2)
	thegna		(mho/cm2)
	minf 		hinf 		
	mtau (ms)	htau (ms) 	
	sinf      	taus (ms)
        tauss (s)
}
 

STATE { m h s}

INITIAL {
        trates(v,ar2)
        m=minf
        h=hinf
        s=sinf
        thegna = gbar*m*m*m*h*s
	ina = thegna * (v - ena)
} 

BREAKPOINT {
        SOLVE states METHOD cnexp
        thegna = gbar*m*m*m*h*s
	ina = thegna * (v - ena)
} 

DERIVATIVE states {
         trates(v,ar2)
         m' = (minf - m)/mtau
         h' = (hinf - h)/htau
         s' = (sinf - s)/taus
}

FUNCTION alpv(v(mV),vh(mV)) {
         alpv = 1/(1+exp((v-vvh)/vvs))
}

:FUNCTION alpv(v(mV),vh(mV)) {    :used in "s" activation system infinity calculation, added by D Itkis 3/27/23, instead of sinf = c + a2*(1-c)
:    alpv = (1+ar2*exp((v-vh)/vvs))/(1+exp((v-vh)/vvs))
:}

        
FUNCTION alps(v(mV)) {  
  alps = exp(1.e-3*zetas*(v-vhalfs)*9.648e4(degC/mV)/(8.315*(273.16+celsius)))
}

FUNCTION bets(v(mV)) {
  bets = exp(1.e-3*zetas*gms*(v-vhalfs)*9.648e4(degC/mV)/(8.315*(273.16+celsius)))
}

LOCAL mexp, hexp, sexp

:PROCEDURE states() {   
:        trates(v,ar2)      
:        m = m + mexp*(minf-m)
:        h = h + hexp*(hinf-h)
:        s = s + sexp*(sinf - s)
:        VERBATIM
:        return 0;
:        ENDVERBATIM
:}

PROCEDURE trates(vm(mV),a2) {  
        LOCAL  a, b, c, qt
        qt=q10^((celsius-24)/10 (degC))
	a = trap0(vm,tha,Ra,qa)
	b = trap0(-vm,-tha,Rb,qa)
	mtau = 1(ms)/(a+b)/qt
        if (mtau<mmin) {mtau=mmin}
        mtau=mtau/mscale
	minf = a/(a+b)
        mexp = 1 - exp(-dt/mtau)

	a = trap0(vm,thi1,Rd,qd)
	b = trap0(-vm,-thi2,Rg,qg)
	htau =  1(ms)/(a+b)/qt
        if (htau<hmin) {htau=hmin}
        htau=htau/hscale
	hinf = 1/(1+exp((vm-thinf)/qinf))
        hexp = 1 - exp(-dt/htau)
   	c=alpv(vm,vhalfs)  
        sinf = c+a2*(1-c)
        taus = bets(vm)/(a0s*(1+alps(vm)))
        if (taus<smin) {taus=smin}
        taus=taus/sscale
        sexp = (1 - exp(-dt/taus))
        tauss = (0.001)*taus
}

FUNCTION trap0(v(mV),th(mV),a(/ms),q(mV)) {
	if (fabs((v-th)*1(/mV)) > 1e-6) {
	        trap0 = 1(ms/mV)*a * (v - th) / (1 - exp(-(v - th)/q))
	} else {
	        trap0 = 1(ms/mV)*a * q
 	}
}	

        
```
