## Supplementary NEURON models for "Dendritic excitations govern back-propagation via a spike-rate accelerometer": Plot_Phase_Diagrams.m

```
% Results of parameter scans for bAP propagation profile, Fig. S20 b-d


% NaV density vs NaV density, fixed stim profile
Nav1 = [0.04, 0.042, 0.044, 0.046, 0.048, 0.05, 0.052, 0.054, 0.056, 0.058, 0.06]; % S/cm2 NaV density soma
Nav2 = [0.042, 0.043, 0.044, 0.045, 0.046, 0.047, 0.048, 0.049, 0.05, 0.051, 0.052, 0.053, 0.054, 0.055]; % S/cm2 NaV density dendrites 
% 
        % duration = 600/6  # ms
        % stim_start = 50/10  # ms
        % stim_end = duration-50/10  # ms
        % Chr_params = [[1, 0, 40, 10, stim_start, stim_end, 0, 0]]
        % synapse_params = []
        % outputs = ["v", "g_chr", "gexp_chr", "s_na3", "ar2_na3"]
        % stimmax = 7e-4  # Light intensity
        % sim_params = [duration, dt, 10, 190, 9, 0]
        % waveform = generate_waveform(sim_params, stimmax, stim_start, stim_end, "step")
        % sim_params[2] = 99 # ms pulse duration for patch clamp pulses
        % sim_params[3] = 99 # ms times between pulses for patch clamp pulses
        % sim_params[4] = 99 # number of pulses for patch clamp pulses
        % sim_params[5] = 0 # baseline current/voltage for patch clamp pulses (nA or mV)
        % waveform_patch = generate_waveform(sim_params, 0, stim_start, stim_end, "pulses")
        % patch_params = [0, waveform_patch, "i"]
        % Nav s Nav_inactivation = [1, 0.5, 300, 100]
        % Kad_params = [-0.2, 0.3, 0, 150]

    vals =[  42      43      44      45      46      47      48      49      50      51      52      53      54      55];
results = [000000, 000000, 000000, 000000, 000000, 010000, 010000, 010000, 010000, 011000, 011000, 110000, 111000, 111000;
           000000, 000000, 000000, 000000, 010000, 010000, 010000, 010000, 010000, 011000, 110000, 111000, 111000, 111100;
           000000, 000000, 000000, 010000, 010000, 010000, 010000, 010000, 011000, 011000, 111000, 111000, 111100, 111100;
           000000, 000000, 010000, 010000, 010000, 010000, 010000, 011000, 011000, 110000, 111000, 111000, 111000, 111100;
           000000, 010000, 010000, 010000, 010000, 010000, 010000, 011000, 011000, 110000, 111000, 111000, 111100, 111100;
           000000, 000000, 010000, 010000, 010000, 010000, 011000, 011000, 110000, 110000, 110000, 110000, 111000, 111000;
           000000, 000000, 000000, 010000, 010000, 011000, 011000, 110000, 110000, 110000, 110000, 110000, 111000, 111000;
           000000, 000000, 000000, 000000, 010000, 010000, 100000, 100000, 110000, 110000, 110000, 110000, 111000, 111000;
           000000, 000000, 000000, 000000, 010000, 100000, 100000, 100000, 100000, 110000, 110000, 110000, 111000, 111000;
           000000, 000000, 000000, 000000, 100000, 100000, 100000, 100000, 100000, 110000, 110000, 110000, 111000, 111000;
           000000, 000000, 000000, 100000, 000000, 010000, 010000, 010000, 010000, 110000, 110000, 110000, 111000, 111000];

figure; 

colormap_vals = [
    0.5, 0.5, 0.5; % gray
    0, 0, 1;  % blue
    0, 1, 1;  % cyan
    0, 1, 0;  % green
    1, 1, 0;  % yellow
    1, 0, 1;  % magenta
    1, 0, 0;  % red
];

colormap(colormap_vals);

binary_results = [
    0, 10000, 11000, 100000, 110000, 111000, 111100
];

binary_to_idx = containers.Map(binary_results, 1:numel(binary_results));

results_idx = arrayfun(@(x) binary_to_idx(x), results);

imagesc(results_idx);
axis equal tight;

yticks(1:length(Nav1));
yticklabels(Nav1);
xticks(1:length(Nav2));
xticklabels(Nav2);

ylabel('Na_V Soma (S/cm^2)');
xlabel('Na_V Dendrites (S/cm^2)');

hold on;
h = zeros(length(binary_results), 1);
for i = 1:length(binary_results)
    h(i) = plot(NaN,NaN,'s','MarkerFaceColor', colormap_vals(i,:), 'MarkerEdgeColor', colormap_vals(i,:));
end
hold off;

results_str = ["FFFFFF", "FSFFFF", "FSSFFF", "SFFFFF", "SSFFFF", "SSSFFF", "SSSSFF"];
% Create the legend
lgd = legend(h, results_str, 'Location', 'northeastoutside');
title(lgd, 'Results');

%%
clear all;
% ri-06 cell parameter stability
% NaV density vs NaV density, fixed stim profile. Low somatic NaV
Nav1 = 0.05; % S/cm2 somatic NaV density
stim_max = [3e-4 3.5e-4 4e-4 4.5e-4 5e-4 5.5e-4 6e-4 6.5e-4 7e-4 7.5e-4 8e-4]; % S/cm2
I_total = [300 350 400 450 500 550 600 650 700 750 800 850 900 950 1000]/70; % nS net stimulus
Nav2 = [0.042, 0.043, 0.044, 0.045, 0.046, 0.047, 0.048, 0.049, 0.05, 0.051, 0.052, 0.053, 0.054, 0.055]; % S/cm2 dendritic NaV density
% 
        % duration = 600/6  # ms
        % stim_start = 50/10  # ms
        % stim_end = duration-50/10  # ms
        % Chr_params = [[1, 0, 40, 10, stim_start, stim_end, 0, 0]]
        % synapse_params = []
        % outputs = ["v", "g_chr", "gexp_chr", "s_na3", "ar2_na3"]
        % sim_params = [duration, dt, 10, 190, 9, 0]
        % waveform = generate_waveform(sim_params, stimmax, stim_start, stim_end, "step")
        % sim_params[2] = 99 # ms pulse duration for patch clamp pulses
        % sim_params[3] = 99 # ms times between pulses for patch clamp pulses
        % sim_params[4] = 99 # number of pulses for patch clamp pulses
        % sim_params[5] = 0 # baseline current/voltage for patch clamp pulses (nA or mV)
        % waveform_patch = generate_waveform(sim_params, 0, stim_start, stim_end, "pulses")
        % patch_params = [0, waveform_patch, "i"]
        % Nav s Nav_inactivation = [1, 0.5, 300, 100]
        % Kad_params = [-0.2, 0.3, 0, 150]

    vals =[  42      43      44      45      46      47      48      49      50      51      52      53      54      55];
results = [000000, 000000, 000000, 000000, 000000, 000000, 000000, 000000, 000000, 000000, 000000, 000000, 000000, 100000;
           000000, 000000, 000000, 000000, 000000, 000000, 000000, 000000, 000000, 000000, 000000, 000000, 000000, 100000;
           000000, 000000, 000000, 010000, 010000, 010000, 010000, 110000, 110000, 110000, 110000, 110000, 110000, 111000;
           000000, 000000, 000000, 010000, 010000, 010000, 010000, 110000, 110000, 110000, 110000, 110000, 110000, 111000;
           000000, 000000, 010000, 010000, 010000, 010000, 011000, 100000, 110000, 110000, 110000, 110000, 111000, 111000;
           000000, 000000, 010000, 010000, 010000, 010000, 011000, 011000, 110000, 110000, 110000, 110000, 111000, 111000;
           000000, 010000, 010000, 010000, 010000, 010000, 011000, 011000, 110000, 110000, 110000, 110000, 111000, 111000;
           000000, 000000, 010000, 010000, 010000, 010000, 011000, 011000, 110000, 110000, 110000, 111000, 111000, 111000;
           000000, 000000, 010000, 010000, 010000, 010000, 011000, 011000, 110000, 110000, 110000, 110000, 111000, 111000;
           000000, 000000, 010000, 010000, 010000, 010000, 011000, 011000, 110100, 110000, 111000, 111000, 111000, 111100;
           000000, 010000, 010000, 010000, 010101, 010000, 011010, 011010, 110101, 110101, 111000, 111001, 111001, 111101;
           ];

figure;

% Ensure the colormap order matches binary_results
colormap_vals = [
    0.5, 0.5, 0.5; % gray for 000000
    0, 0, 1;  % blue for 010000
    0, 1, 1;  % cyan for 011000
    0, 1, 0;  % green for 011100
    1, 1, 0;  % yellow for 011110
    1, 0, 1;  % magenta for 011111
    1, 0, 0;  % red for alternating
];
colormap(colormap_vals);

binary_results = [0, 10000, 11000, 100000, 110000, 111000, 111100];
binary_to_idx = containers.Map(binary_results, 1:numel(binary_results));

% Add mapping for "alternating"
binary_to_idx(-1) = 7; % Map "alternating" to the red color

% Initialize results_idx matrix with the same size as results
results_idx = zeros(size(results));

% Populate results_idx
for i = 1:size(results, 1)
    for j = 1:size(results, 2)
        if isKey(binary_to_idx, results(i,j))
            results_idx(i,j) = binary_to_idx(results(i,j));
        else
            results_idx(i,j) = binary_to_idx(-1); 
        end
    end
end

imagesc(results_idx);
axis equal tight;

yticks(1:length(stim_max));
yticklabels(I_total);
xticks(1:length(Nav2));
xticklabels(Nav2);

ylabel('Net equivalent stimulation current (nS)');
xlabel('Na_V Dendrites (S/cm^2)');

hold on;
h = zeros(length(binary_results), 1);
for i = 1:length(binary_results)
    h(i) = plot(NaN,NaN,'s','MarkerFaceColor', colormap_vals(i,:), 'MarkerEdgeColor', colormap_vals(i,:));
end
hold off;

results_str = ["FFFFFF", "FSFFFF", "FSSFFF", "SFFFFF", "SSFFFF", "SSSFFF", "alternating"];
% Create the legend
lgd = legend(h, results_str, 'Location', 'northeastoutside');
title(lgd, 'Results');

%%
clear all;
% ri-06 cell parameter stability
% NaV inactivation vs. Stim strength
Nav1 = 0.05; % S/cm2 somatic NaV density
Nav2 = 0.047; % S/cm2 dendritic NaV density
stim_max = [3e-4 3.5e-4 4e-4 4.5e-4 5e-4 5.5e-4 6e-4 6.5e-4 7e-4 7.5e-4 8e-4]; % S/cm2
I_total = [300 350 400 450 500 550 600 650 700 750 800 850 900 950 1000]/70; % nS net stimulus
inact = [0, 0.1, 0.2, 0.3, 0.4, 0.5, 0.6, 0.7, 0.8, 0.9, 1]; % ar2

% 
        % duration = 600/6  # ms
        % stim_start = 50/10  # ms
        % stim_end = duration-50/10  # ms
        % Chr_params = [[1, 0, 40, 10, stim_start, stim_end, 0, 0]]
        % synapse_params = []
        % outputs = ["v", "g_chr", "gexp_chr", "s_na3", "ar2_na3"]
        % sim_params = [duration, dt, 10, 190, 9, 0]
        % waveform = generate_waveform(sim_params, stimmax, stim_start, stim_end, "step")
        % sim_params[2] = 99 # ms pulse duration for patch clamp pulses
        % sim_params[3] = 99 # ms times between pulses for patch clamp pulses
        % sim_params[4] = 99 # number of pulses for patch clamp pulses
        % sim_params[5] = 0 # baseline current/voltage for patch clamp pulses (nA or mV)
        % waveform_patch = generate_waveform(sim_params, 0, stim_start, stim_end, "pulses")
        % patch_params = [0, waveform_patch, "i"]
        % Nav s Nav_inactivation = [1, 0.5, 300, 100]
        % Kad_params = [-0.2, 0.3, 0, 150]

    vals =[  0       1        2      3      4        5       6       7       8       9       1.0 ];
results = [000000, 000000, 000000, 000000, 000000, 010000, 010000, 010000, 011100, 011111, 011111;
           000000, 000000, 000000, 010000, 010000, 010000, 010000, 011000, 011100, 011100, 011111;
           000000, 000000, 010000, 010000, 010000, 010000, 010000, 011000, 011010, 011100, 011111;
           000000, 010000, 010000, 010000, 010000, 010000, 011000, 011000, 011101, 011100, 011111;
           010000, 010000, 010000, 010000, 010000, 010000, 011000, 011000, 011000, 011100, 011111;
           010000, 010000, 010000, 010000, 010000, 010000, 011000, 011000, 011000, 011110, 011111;
           010000, 010000, 010000, 010000, 010000, 010000, 011000, 011000, 011000, 011110, 011111;
           010000, 010000, 010000, 010000, 010000, 010000, 011000, 011000, 011100, 011111, 011111;
           010000, 010000, 010000, 010000, 010000, 010000, 011000, 011000, 011101, 011111, 011111;
           010000, 010000, 010000, 010000, 010000, 011000, 011000, 011001, 011101, 011111, 011111;
           010000, 010000, 010000, 010000, 010000, 010000, 011010, 011000, 011100, 011101, 011111;                   
           ];

figure;

% Ensure the colormap order matches binary_results
colormap_vals = [
    0.5, 0.5, 0.5; % gray for 000000
    0, 0, 1;  % blue for 010000
    0, 1, 1;  % cyan for 011000
    0, 1, 0;  % green for 011100
    1, 1, 0;  % yellow for 011110
    1, 0, 1;  % magenta for 011111
    1, 0, 0;  % red for alternating
];
colormap(colormap_vals);

% Map using integers directly
binary_results = [0, 10000, 11000, 11100, 11110, 11111, -1]; % -1 for "alternating"
binary_to_idx = containers.Map(binary_results, 1:numel(binary_results));

% Initialize results_idx matrix with the same size as results
results_idx = zeros(size(results));

% Populate results_idx
for i = 1:size(results, 1)
    for j = 1:size(results, 2)
        if isKey(binary_to_idx, results(i,j))
            results_idx(i,j) = binary_to_idx(results(i,j));
        else
            results_idx(i,j) = binary_to_idx(-1); % For "alternating"
        end
    end
end

imagesc(results_idx);
%colorbar;
axis equal tight;

yticks(1:length(stim_max));
yticklabels(I_total);
xticks(1:length(inact));
xticklabels(100*inact);

ylabel('Net stimulation conductance (nS)');
xlabel('Percent non-inactivating distal Na_V channels (%)');

hold on;
h = zeros(length(binary_results), 1);
for i = 1:length(binary_results)
    h(i) = plot(NaN,NaN,'s','MarkerFaceColor', colormap_vals(i,:), 'MarkerEdgeColor', colormap_vals(i,:));
end
hold off;

results_str = ["FFFFFF", "FSFFFF", "FSSFFF", "FSSSFF", "FSSSSF", "FSSSSS", "alternating"];
% Create the legend
lgd = legend(h, results_str, 'Location', 'northeastoutside');
title(lgd, 'Results');

function idx = safe_lookup(x)
    if isKey(binary_to_idx, x)
        idx = binary_to_idx(x);
    else
        idx = binary_to_idx(-1); % For "alternating"
    end
end
```
