## Supplementary NEURON models for "Dendritic excitations govern back-propagation via a spike-rate accelerometer": parameters_list.py

```
from run_simulation import generate_waveform

def setup_parameters(run_nr):
    runs = ["Patch pulses soma", "Optopatch pulses soma", "Patch step soma", "Optopatch step soma", 
            "Patch pulses oblique", "Optopatch pulses oblique", "Patch pulses apical", "Optopatch pulses apical", 
            "Optopatch pulses soma+apical", "Patch pulses basal", "Optopatch pulses basal", "Optopatch pulses widefield", 
            "Optopatch step widefield", "Optopatch pulses+TTX soma", "Optopatch pulses+4-AP soma", "Optopatch ramp+TTX soma", 
            "Optopatch+EFS DeltaT soma+apical", "Optopatch+EFS Ni soma+apical", "Optopatch+EFS D-AP5 soma+apical", "Optopatch+EFS TTX soma+apical"]
    
    print("Starting run: ", runs[run_nr])

    dt = 0.025  # ms

    Nav_params = [0.05, 0.048, 300, 100]
    Kad_params = [-0.2, 0.3, 0, 150]
    Kap_params = [-0.2, 0.3, 0, 150]
    Cal_params = [1e-9, 1e-9, 550, 100]
    Car_params = [1e-9, 1e-9, 300, 100]
    #Car_params = [1e-2, 3e-2, 300, 100]

    #Patch pulses soma 
    if run_nr == 0:
        duration = 1100  # ms
        stim_start = 50  # ms
        stim_end = duration-48 # ms
        Chr_params = [[1, 0, 30, 10, stim_start, stim_end, 0, 0]]
        synapse_params = []
        outputs = ["v", "g_chr", "gexp_chr"]
        stimmax = 0  # Light intensity
        sim_params = [duration, dt, 99, 99, 99, 0]
        waveform = generate_waveform(sim_params, stimmax, stim_start, stim_end, "pulses")
        sim_params[2] = 2 # ms pulse duration for patch clamp pulses
        sim_params[3] = 98 # ms times between pulses for patch clamp pulses
        sim_params[4] = 10 # number of pulses for patch clamp pulses
        sim_params[5] = 0 # baseline current/voltage for patch clamp pulses (nA or mV)
        waveform_patch = generate_waveform(sim_params, 2, stim_start, stim_end, "pulses")
        patch_params = [0, waveform_patch, "i"]
    
    #Optopatch pulses soma 
    if run_nr == 1:
        duration = 2000  # ms
        stim_start = 50 # ms
        stim_end = duration-48  # ms
        Chr_params = [[1, 0, 30, 10, stim_start, stim_end, 0, 0]]
        synapse_params = []
        outputs = ["v", "g_chr", "gexp_chr"]
        stimmax = 8e-4  # Light intensity
        sim_params = [duration, dt, 20, 180, 10, 0] # 1c 10 ms, 2g 20ms stim
        waveform = generate_waveform(sim_params, stimmax, stim_start, stim_end, "pulses")
        sim_params[2] = 99 # ms pulse duration for patch clamp pulses
        sim_params[3] = 99 # ms times between pulses for patch clamp pulses
        sim_params[4] = 99 # number of pulses for patch clamp pulses
        sim_params[5] = 0 # baseline current/voltage for patch clamp pulses (nA or mV)
        waveform_patch = generate_waveform(sim_params, 0, stim_start, stim_end, "pulses")
        patch_params = [0, waveform_patch, "i"]

    #Patch step soma 
    # Compare to Optopatch step soma. 
    if run_nr == 2:
        duration = 85  # ms
        stim_start = 25  # ms
        stim_end = duration-10  # ms
        Chr_params = [[1, 0, 30, 10, stim_start, stim_end, 0, 0]]
        synapse_params = []
        outputs = ["v", "g_chr", "gexp_chr"]
        stimmax = 0  # Light intensity
        sim_params = [duration, dt, 99, 99, 99, 0]
        waveform = generate_waveform(sim_params, stimmax, stim_start, stim_end, "pulses")
        sim_params[2] = 2 # ms pulse duration for patch clamp pulses
        sim_params[3] = 98 # ms times between pulses for patch clamp pulses
        sim_params[4] = 10 # number of pulses for patch clamp pulses
        sim_params[5] = 0 # baseline current/voltage for patch clamp pulses (nA or mV)
        waveform_patch = generate_waveform(sim_params, 0.5, stim_start, stim_end, "step")
        patch_params = [0, waveform_patch, "i"]

    #Optopatch step soma
    if run_nr == 3:
        duration = 85  # ms
        stim_start = 25  # ms
        stim_end = duration-10  # ms
        Chr_params = [[1, 1e-15, 40, 10, stim_start, stim_end, 0, 0]]
        #Cheriff_params = [[1, 1e-15, 40, 10, stim_start, stim_end, 0, 0]]
        synapse_params = []
        outputs = ["v", "g_chr", "gexp_chr"]
        stimmax = 4.9e-4  # Light intensity mW/cm^2
        sim_params = [duration, dt, 10, 190, 9, 0]
        waveform = generate_waveform(sim_params, stimmax, stim_start, stim_end, "step")
        sim_params[2] = 10 # ms pulse duration for patch clamp pulses
        sim_params[3] = 100 # ms times between pulses for patch clamp pulses
        sim_params[4] = 2 # number of pulses for patch clamp pulses
        sim_params[5] = 0 # baseline current/voltage for patch clamp pulses (nA or mV)
        waveform_patch = generate_waveform(sim_params, 0, stim_start, stim_end, "pulses")
        patch_params = [0, waveform_patch, "i"]

    #Patch pulses oblique 
    # Use section e.g. 94, 98.
    if run_nr == 4:
        duration = 150#2500/8  # ms
        stim_start = 50  # ms
        stim_end = duration-50/10  # ms
        Chr_params = [[1, 0, 30, 1, stim_start, stim_end, 0, 0]]
        synapse_params = []
        outputs = ["v", "g_chr", "gexp_chr"]
        stimmax = 0  # Light intensity
        sim_params = [duration, dt, 2, 20, 50, 0]
        waveform = generate_waveform(sim_params, stimmax, stim_start, stim_end, "pulses")
        sim_params[2] = 90#2*10*5 # ms pulse duration for patch clamp pulses
        sim_params[3] = 198 # ms times between pulses for patch clamp pulses
        sim_params[4] = 10 # number of pulses for patch clamp pulses
        sim_params[5] = 0 # baseline current/voltage for patch clamp pulses (nA or mV)
        waveform_patch = generate_waveform(sim_params, 1.5, stim_start, stim_end, "pulses")
        patch_params = [94, waveform_patch, "i"]

    #Optopatch pulses oblique 
    # Use section e.g. 94, 98.
    if run_nr == 5:
        duration = 150#2500  # ms
        stim_start = 50 # ms
        stim_end = duration-50/10  # ms
        Chr_params = [[1, 0, 60, 10, stim_start, stim_end, 94, 0]]
        synapse_params = []
        outputs = ["v", "g_chr", "gexp_chr"]
        stimmax = 2.5e-3  # Light intensity
        sim_params = [duration, dt, 20, 180, 45, 0]
        waveform = generate_waveform(sim_params, stimmax, stim_start, stim_end, "pulses")
        sim_params[2] = 99 # ms pulse duration for patch clamp pulses
        sim_params[3] = 99 # ms times between pulses for patch clamp pulses
        sim_params[4] = 99 # number of pulses for patch clamp pulses
        sim_params[5] = 0 # baseline current/voltage for patch clamp pulses (nA or mV)
        waveform_patch = generate_waveform(sim_params, 0, 0, 300, "pulses")
        patch_params = [0, waveform_patch, "i"]

    #Patch pulses apical 
    if run_nr == 6:
        duration = 2500  # ms
        stim_start = 50  # ms
        stim_end = duration-50  # ms
        Chr_params = [[1, 0, 30, 10, stim_start, stim_end, 0, 0]]
        synapse_params = []
        outputs = ["v", "g_chr", "gexp_chr"]
        stimmax = 0  # Light intensity
        sim_params = [duration, dt, 99, 99, 99, 0]
        waveform = generate_waveform(sim_params, stimmax, stim_start, stim_end, "pulses")
        sim_params[2] = 2 # ms pulse duration for patch clamp pulses
        sim_params[3] = 198 # ms times between pulses for patch clamp pulses
        sim_params[4] = 20 # number of pulses for patch clamp pulses
        sim_params[5] = 0 # baseline current/voltage for patch clamp pulses (nA or mV)
        waveform_patch = generate_waveform(sim_params, 1, stim_start, stim_end, "pulses")
        patch_params = [144, waveform_patch, "i"]

    #Optopatch pulses apical 
    if run_nr == 7:
        duration = 2500  # ms
        stim_start = 50  # ms
        stim_end = duration-50  # ms
        Chr_params = [[1, 0, 30, 10, stim_start, stim_end, 144, 0]]
        synapse_params = []
        outputs = ["v", "g_chr", "gexp_chr"]
        stimmax = 1e-4  # Light intensity
        sim_params = [duration, dt, 20, 180, 20, 0]
        waveform = generate_waveform(sim_params, stimmax, stim_start, stim_end, "step")
        sim_params[2] = 99 # ms pulse duration for patch clamp pulses
        sim_params[3] = 99 # ms times between pulses for patch clamp pulses
        sim_params[4] = 99 # number of pulses for patch clamp pulses
        sim_params[5] = 0 # baseline current/voltage for patch clamp pulses (nA or mV)
        waveform_patch = generate_waveform(sim_params, 0, 0, 300, "pulses")
        patch_params = [0, waveform_patch, "i"]

    #Optopatch pulses soma+apical - Figure 2h
    if run_nr == 8:
        duration = 2500  # ms
        stim_start = 50  # ms
        stim_end = duration-50  # ms
        Chr_params = [[1, 0, 30, 10, stim_start, stim_end, 0, 0],[1, 0, 30, 10, stim_start, stim_end, 144, 0]]
        synapse_params = []
        outputs = ["v", "g_chr", "gexp_chr", "gkabar_kad"]
        stimmax = 8e-4  # Light intensity
        sim_params = [duration, dt, 20, 180, 50, 0]
        waveform = generate_waveform(sim_params, stimmax, stim_start, stim_end, "full")
        sim_params[2] = 99 # ms pulse duration for patch clamp pulses
        sim_params[3] = 99 # ms times between pulses for patch clamp pulses
        sim_params[4] = 99 # number of pulses for patch clamp pulses
        sim_params[5] = 0 # baseline current/voltage for patch clamp pulses (nA or mV)
        waveform_patch = generate_waveform(sim_params, 0, 0, 300, "pulses")
        patch_params = [0, waveform_patch, "i"]

    #Patch pulses basal  
    # Use section e.g. 25, 50 or 74.
    # For separate branches: 15, 46, 70, 83
    if run_nr == 9:
        duration = 2500 # ms
        stim_start = 50  # ms
        stim_end = duration-50  # ms
        Chr_params = [1, 0, 30, 10, stim_start, stim_end, 0, 0]
        synapse_params = []
        outputs = ["v", "g_chr", "gexp_chr", "gkabar_kad"]
        stimmax = 0  # Light intensity
        sim_params = [duration, dt, 99, 99, 99, 0]
        waveform = generate_waveform(sim_params, stimmax, stim_start, stim_end, "pulses")
        sim_params[2] = 2 # ms pulse duration for patch clamp pulses
        sim_params[3] = 198 # ms times between pulses for patch clamp pulses
        sim_params[4] = 20 # number of pulses for patch clamp pulses
        sim_params[5] = 0 # baseline current/voltage for patch clamp pulses (nA or mV)
        waveform_patch = generate_waveform(sim_params, 1, stim_start, stim_end, "pulses")
        patch_params = [25, waveform_patch, "i"]

    #Optopatch pulses basal
    # Use section e.g. 25, 50 or 74.
    # For separate branches: 15, 46, 70, 83
    if run_nr == 10:
        duration = 2500  # ms
        stim_start = 50  # ms
        stim_end = duration-50  # ms
        Chr_params = [1, 0, 30, 10, stim_start, stim_end, 34, 0]
        synapse_params = []
        outputs = ["v", "g_chr", "gexp_chr", "gkabar_kad"]
        stimmax = 9e-4  # Light intensity
        sim_params = [duration, dt, stim_start, stim_end, 25, 0]
        waveform = generate_waveform(sim_params, stimmax, stim_start, stim_end, "pulses")
        sim_params[2] = 99 # ms pulse duration for patch clamp pulses
        sim_params[3] = 99 # ms times between pulses for patch clamp pulses
        sim_params[4] = 99 # number of pulses for patch clamp pulses
        sim_params[5] = 0 # baseline current/voltage for patch clamp pulses (nA or mV)
        waveform_patch = generate_waveform(sim_params, 0, 0, 300, "pulses")
        patch_params = [0, waveform_patch, "i"]

    #Optopatch step oblique - Figure 3a
    #Use section e.g. 94, 98.
    if run_nr == 11:
        duration = 600/12  # ms
        stim_start = 50/10  # ms
        stim_end = duration-50/10 # ms
        Chr_params = [1, 0, 30, 10, stim_start, stim_end, 98, 0]
        synapse_params = []
        outputs = ["v", "g_chr", "gexp_chr"]
        stimmax = 1e-3  # Light intensity
        sim_params = [duration, dt, 99, 99, 99, 0]
        waveform = generate_waveform(sim_params, stimmax, stim_start, stim_end, "step")
        sim_params[2] = 99 # ms pulse duration for patch clamp pulses
        sim_params[3] = 99 # ms times between pulses for patch clamp pulses
        sim_params[4] = 99 # number of pulses for patch clamp pulses
        sim_params[5] = 0 # baseline current/voltage for patch clamp pulses (nA or mV)
        waveform_patch = generate_waveform(sim_params, 0, 0, 300, "pulses")
        patch_params = [0, waveform_patch, "i"]

    #Optopatch step widefield 
    if run_nr == 12:
        duration = 700/7  # ms
        stim_start = 50/10  # ms
        stim_end = duration-100/20  # ms
        Chr_params = [1, 0.1, 200, 100, stim_start, stim_end, 0, 0]
        synapse_params = []
        outputs = ["v", "g_chr", "gexp_chr"]
        stimmax = 3e-4  # Light intensity
        sim_params = [duration, dt, 2, 20, 50, 0]
        waveform = generate_waveform(sim_params, stimmax, stim_start, stim_end, "step")
        sim_params[2] = 99 # ms pulse duration for patch clamp pulses
        sim_params[3] = 99 # ms times between pulses for patch clamp pulses
        sim_params[4] = 99 # number of pulses for patch clamp pulses
        sim_params[5] = 0 # baseline current/voltage for patch clamp pulses (nA or mV)
        waveform_patch = generate_waveform(sim_params, 0, 0, 300, "pulses")
        patch_params = [0, waveform_patch, "i"]

    #Optopatch pulses+TTX soma
    if run_nr == 13:
        duration = 2500  # ms
        stim_start = 50  # ms
        stim_end = duration-50  # ms
        Chr_params = [[1, 0, 30, 10, stim_start, stim_end, 0, 0]]
        synapse_params = []
        outputs = ["v", "g_chr", "gexp_chr"]
        stimmax = 2e-3  # Light intensity
        sim_params = [duration, dt, 20, 180, 50, 0]
        waveform = generate_waveform(sim_params, stimmax, stim_start, stim_end, "pulses")
        sim_params[2] = 99 # ms pulse duration for patch clamp pulses
        sim_params[3] = 99 # ms times between pulses for patch clamp pulses
        sim_params[4] = 99 # number of pulses for patch clamp pulses
        sim_params[5] = 0 # baseline current/voltage for patch clamp pulses (nA or mV)
        waveform_patch = generate_waveform(sim_params, 0, 0, 300, "pulses")
        patch_params = [0, waveform_patch, "i"]
        Nav_params = [0.04/5, 0.02/5, 250, 200]

    #Optopatch pulses+4-AP soma 
    # 5 mM 4-AP bath application
    if run_nr == 14:
        duration = 2500  # ms
        stim_start = 50  # ms
        stim_end = duration-50  # ms
        Chr_params = [1, 0, 30, 10, stim_start, stim_end, 0, 0]
        synapse_params = []
        outputs = ["v", "g_chr", "gexp_chr"]
        stimmax = 1e-4  # Light intensity
        sim_params = [duration, dt, 20, 180, 50, 0]
        waveform = generate_waveform(sim_params, stimmax, stim_start, stim_end, "pulses")
        sim_params[2] = 99 # ms pulse duration for patch clamp pulses
        sim_params[3] = 99 # ms times between pulses for patch clamp pulses
        sim_params[4] = 99 # number of pulses for patch clamp pulses
        sim_params[5] = 0 # baseline current/voltage for patch clamp pulses (nA or mV)
        waveform_patch = generate_waveform(sim_params, 0, 0, 300, "pulses")
        patch_params = [0, waveform_patch, "i"]
        Kad_params = [-0.3/2, 0.3/2, 0, 150]

    #Optopatch ramp+TTX soma - Figure S8
    # Baseline, 100 nM and 500 nM TTX bath application
    if run_nr == 15:
        duration = 2500  # ms
        stim_start = 100  # ms
        stim_end = duration-100  # ms
        Chr_params = [1, 0, 30, 10, stim_start, stim_end, 0, 0]
        synapse_params = []
        outputs = ["v", "g_chr", "gexp_chr"]
        stimmax = 1e-3  # Light intensity
        sim_params = [duration, dt, 0, 0, 0, 0]
        waveform = generate_waveform(sim_params, stimmax, stim_start, stim_end, "ramp")
        sim_params[2] = 99 # ms pulse duration for patch clamp pulses
        sim_params[3] = 99 # ms times between pulses for patch clamp pulses
        sim_params[4] = 99 # number of pulses for patch clamp pulses
        sim_params[5] = 0 # baseline current/voltage for patch clamp pulses (nA or mV)
        waveform_patch = generate_waveform(sim_params, 0, 0, 300, "pulses")
        patch_params = [0, waveform_patch, "i"]
        Nav_params = [0.04, 0.02, 250, 200]

    #Optopatch+EFS DeltaT soma+apical 
    # Time delays relative to middle of light pulse
    # 8 combinations in total:
    # 30 ms light pulse w/ time delay -85ms (EFS 85 ms after middle of light pulse), 
    # -15ms, +45ms and +115ms
    # 15 ms light pulse w/ time delay -95ms, -25ms, +35ms and +105ms 
    # AMPA + NMDA broken, only AMPA usable for EFS
    if run_nr == 16:
        duration = 600  # ms
        stim_start = 50  # ms
        stim_end = 80 # ms
        Chr_params = [1, 0, 40, 10, stim_start, stim_end, 0, 0]
        synapse_params = [[1500, 65+15, 200, 0.1, 0.00033, "regular", "distal", "ampa+nmda", 2]]
        outputs = ["v", "g_chr", "gexp_chr"]
        stimmax = 6e-4  # Light intensity
        sim_params = [duration, dt, 0, 0, 0, 0]
        waveform = generate_waveform(sim_params, stimmax, stim_start, stim_end, "step")
        sim_params[2] = 99 # ms pulse duration for patch clamp pulses
        sim_params[3] = 99 # ms times between pulses for patch clamp pulses
        sim_params[4] = 99 # number of pulses for patch clamp pulses
        sim_params[5] = 0 # baseline current/voltage for patch clamp pulses (nA or mV)
        waveform_patch = generate_waveform(sim_params, 0, 0, 300, "pulses")
        patch_params = [0, waveform_patch, "i"]

    #Optopatch+EFS Ni soma+apical 
    # 100 uM Ni2+ blocks Ca channels 
    if run_nr == 17:
        duration = 500  # ms
        stim_start = 50  # ms
        stim_end = 80 # ms
        Chr_params = [1, 0, 30, 10, stim_start, stim_end, 0, 0]
        synapse_params = [[1000, 65-15, 200, 0.1, 0.0005, "regular", "apical", "ampa+nmda"]]
        outputs = ["v", "g_chr", "gexp_chr"]
        stimmax = 5e-4  # Light intensity
        sim_params = [duration, dt, 0, 0, 0, 0]
        waveform = generate_waveform(sim_params, stimmax, stim_start, stim_end, "step")
        sim_params[2] = 99 # ms pulse duration for patch clamp pulses
        sim_params[3] = 99 # ms times between pulses for patch clamp pulses
        sim_params[4] = 99 # number of pulses for patch clamp pulses
        sim_params[5] = 0 # baseline current/voltage for patch clamp pulses (nA or mV)
        waveform_patch = generate_waveform(sim_params, 0, 0, 300, "pulses")
        patch_params = [0, waveform_patch, "i"]
        Cal_params = [0, 0, 550, 100]
        Car_params = [0, 0, 550, 100]

    #Optopatch+EFS NMDA soma+apical 
    # 50 uM D-AP5 blocks NMDA channels
    if run_nr == 18:
        duration = 500  # ms
        stim_start = 50  # ms
        stim_end = 80 # ms
        Chr_params = [1, 0, 30, 10, stim_start, stim_end, 0, 0]
        synapse_params = [[1000, 65-15, 200, 0.1, 0.0005, "regular", "apical", "ampa"]]
        outputs = ["v", "g_chr", "gexp_chr"]
        stimmax = 5e-4  # Light intensity
        sim_params = [duration, dt, 0, 0, 0, 0]
        waveform = generate_waveform(sim_params, stimmax, stim_start, stim_end, "step")
        sim_params[2] = 99 # ms pulse duration for patch clamp pulses
        sim_params[3] = 99 # ms times between pulses for patch clamp pulses
        sim_params[4] = 99 # number of pulses for patch clamp pulses
        sim_params[5] = 0 # baseline current/voltage for patch clamp pulses (nA or mV)
        waveform_patch = generate_waveform(sim_params, 0, 0, 300, "pulses")
        patch_params = [0, waveform_patch, "i"]

    #Optopatch+EFS TTX soma+apical 
    # 20 nM TTX blocks ~80% of NaV channels
    if run_nr == 19:
        duration = 500  # ms
        stim_start = 50  # ms
        stim_end = 80 # ms
        Chr_params = [1, 0, 30, 10, stim_start, stim_end, 0, 0]
        synapse_params = [[1000, 65-15, 200, 0.1, 0.0005, "regular", "apical", "ampa+nmda",0.0]]
        outputs = ["v", "g_chr", "gexp_chr"]
        stimmax = 5e-4  # Light intensity
        sim_params = [duration, dt, 0, 0, 0, 0]
        waveform = generate_waveform(sim_params, stimmax, stim_start, stim_end, "step")
        sim_params[2] = 99 # ms pulse duration for patch clamp pulses
        sim_params[3] = 99 # ms times between pulses for patch clamp pulses
        sim_params[4] = 1 # number of pulses for patch clamp pulses
        sim_params[5] = 0 # baseline current/voltage for patch clamp pulses (nA or mV)
        waveform_patch = generate_waveform(sim_params, 0, stim_start, stim_end, "full")
        patch_params = [0, waveform_patch, "i"]
        Nav_params = [0.05/5, 0.05/5, 250, 200]

    print(str(duration) + " ms")

    return duration, stim_start, stim_end, Chr_params, synapse_params, sim_params, outputs, waveform, waveform_patch, patch_params, Nav_params, Kad_params, Kap_params, Cal_params, Car_params
```
