## Supplementary NEURON models for "Dendritic excitations govern back-propagation via a spike-rate accelerometer": Plot_Voltage_Traces.m

```
clear all;
close all;
% Define the paths to the data files
data_file = 'C:\Users\danie\Documents\GitHub\neuron-visualizer\online-visualizer\data.json';
data_syn_file = 'C:\Users\danie\Documents\GitHub\neuron-visualizer\online-visualizer\data_syn.json';

% Read the data from the JSON files
data = jsondecode(fileread(data_file));
data_syn = jsondecode(fileread(data_syn_file));

% Extract voltage data
voltage = data.var_plot;

figure;
for i = 640:1:700%length(voltage(:,1))
% Plot voltage(1,:) and voltage(98,:)
%subplot(2,1,1);
plot(voltage(i,:));
hold off;
title(i);
pause(1);
end

figure;
plot(0:0.025:length(voltage(1,:))*0.025-0.025,voltage(1,:), 'LineWidth', 2);
xlabel("Time (ms)");
ylabel("Membrane Potential (mV)");
xlim([0 40]);
ylim([-80 40]);
figure;
plot(0:0.025:length(voltage(1,:))*0.025-0.025,voltage(660,:), 'LineWidth', 2);  %921
xlabel("Time (ms)");
ylabel("Membrane Potential (mV)");
xlim([0 40]);
ylim([-80 0]);

figure;
plot(0:0.025:length(voltage(1,:))*0.025-0.025,voltage(660:2:670,:)', 'LineWidth', 2);
xlabel("Time (ms)");
ylabel("Membrane Potential (mV)");
```
