## Supplementary NEURON models for "Dendritic excitations govern back-propagation via a spike-rate accelerometer": Plot_Voltage_Traces.m

```
clear all;
close all;
% Define the paths to the data files
data_file = 'C:\Users\danie\Documents\GitHub\neuron-visualizer\online-visualizer\data.json';
data_syn_file = 'C:\Users\danie\Documents\GitHub\neuron-visualizer\online-visualizer\data_syn.json';

% Read the data from the JSON files
data = jsondecode(fileread(data_file));
data_syn = jsondecode(fileread(data_syn_file));

% Extract voltage data
voltage = data.var_plot;

% figure;
% for i = 1:10:length(voltage(:,1))
% % Plot voltage(1,:) and voltage(98,:)
% %subplot(2,1,1);
% plot(voltage(i,:));
% hold off;
% title(i);
% pause(0.5);
% end

figure;
plot(0:0.025:length(voltage(1,:))*0.025-0.025,voltage(1,:), 'LineWidth', 2);
xlabel("Time (ms)");
ylabel("Membrane Potential (mV)");
figure;
plot(0:0.025:length(voltage(1,:))*0.025-0.025,voltage(919,:), 'LineWidth', 2);  %921
xlabel("Time (ms)");
ylabel("Membrane Potential (mV)");

figure;
plot(0:0.025:length(voltage(1,:))*0.025-0.025,voltage(680:700,:)', 'LineWidth', 2);
xlabel("Time (ms)");
ylabel("Membrane Potential (mV)");
```
