## Supplementary NEURON models for "Dendritic excitations govern back-propagation via a spike-rate accelerometer": parameters_list.py

```
from run_simulation import generate_waveform

def setup_parameters(run_nr):
    runs = ["Patch pulse oblique", "Optopatch pulse oblique"]
    
    print("Starting run: ", runs[run_nr])

    dt = 0.025  # ms

    Nav_params = [0.045, 0.045, 300, 100]
    Kad_params = [-0.2, 0.3, 0, 150]
    Kap_params = [-0.2, 0.3, 0, 150]
    Cal_params = [1e-9, 1e-9, 550, 100]
    Car_params = [1e-9, 1e-9, 300, 100]

    #Patch pulses oblique 
    # This results in a single patch current pulse of 5ms duration.
    # Use section e.g. 94, 98.
    if run_nr == 0:
        duration = 60 # ms
        stim_start = 15  # ms
        stim_end = duration-10  # ms
        Chr_params = [[1, 0, 30, 1, stim_start, stim_end, 0, 0]]
        synapse_params = []
        outputs = ["v", "g_chr", "gexp_chr"]
        stimmax = 0  # Light intensity
        sim_params = [duration, dt, 2, 20, 50, 0]
        waveform = generate_waveform(sim_params, stimmax, stim_start, stim_end, "pulses")
        sim_params[2] = 5 # ms pulse duration for patch clamp pulses
        sim_params[3] = 198 # ms times between pulses for patch clamp pulses
        sim_params[4] = 10 # number of pulses for patch clamp pulses
        sim_params[5] = 0 # baseline current/voltage for patch clamp pulses (nA or mV)
        waveform_patch = generate_waveform(sim_params, 2, stim_start, stim_end, "pulses")
        patch_params = [98, waveform_patch, "i"] # 98 is the section number

    #Optopatch pulses oblique instantaneous 
    # This results in a step of conductance of 45ms duration.
    # Use section e.g. 94, 98.
    if run_nr == 1:
        duration = 80  # ms
        stim_start = 15 # ms
        stim_end = duration-20  # ms
        Chr_params = [[1, 0, 50, 10, stim_start, stim_end, 98, 0]] # 98 is the section number
        synapse_params = []
        outputs = ["v", "g_chr", "gexp_chr"]
        stimmax = 1e-3  # Light intensity
        sim_params = [duration, dt, 30, 180, 45, 0]
        waveform = generate_waveform(sim_params, stimmax, stim_start, stim_end, "step")
        sim_params[2] = 99 # ms pulse duration for patch clamp pulses
        sim_params[3] = 99 # ms times between pulses for patch clamp pulses
        sim_params[4] = 99 # number of pulses for patch clamp pulses
        sim_params[5] = 0 # baseline current/voltage for patch clamp pulses (nA or mV)
        waveform_patch = generate_waveform(sim_params, 0, 0, 300, "pulses")
        patch_params = [0, waveform_patch, "i"]

    print(str(duration) + " ms")

    return duration, stim_start, stim_end, Chr_params, synapse_params, sim_params, outputs, waveform, waveform_patch, patch_params, Nav_params, Kad_params, Kap_params, Cal_params, Car_params
```
