## Supplementary NEURON models for "Dendritic excitations govern back-propagation via a spike-rate accelerometer": Plot_Channel_Distributions.m

```
clear all; close all;

%% Import Geometry Data
% Import Segment Labels
load('Labels.mat');

% Import Distance Matrix
load('Distance.mat');
D = distance;

% Import Segment Labels
load('PositionsN.mat');
NC = [0, cumsum(Nsec)];

% Import Segment Positions
load('PositionsXYZ.mat');


%% NaV densities in S/cm2
NaV_densities = ones(1,length(D(:,1)))*0.05;
NaV_densities(156) = 15;
NaV_densities(157) = 30;
NaV_densities(158) = 30;

%% NaV slow inactivation 
gprox = 1;
gdistal = 0;
d12 = 300;
z = 100;

for i = 1:length(D(1,:))
    NaV_inactivation(i) = gprox + (gdistal - gprox)/(1 + exp((D(1,i)-d12)/z));
end

%% A-type K density
gprox = 0.3;
gdistal = -0.2;
d12 = 0;
z = 150;

for i = 1:length(D(1,:))
    AK_density(i) = gprox + (gdistal - gprox)/(1 + exp((D(1,i)-d12)/z));
end

%%  Plot Neuron LUT Colormap Snap Detailed Soma

% dyn = log(NaV_densities);
%Vrange = [log10(0.05); log10(100)];

dyn = AK_density;
Vrange = [0.05; 0.3];

% dyn = NaV_inactivation;
% Vrange = [0; 1];

red = [zeros(1,95) (linspace(0,1,160-95)) ones(1,224-161) (linspace(1,0.5,258-225))];
blue = [(linspace(0.5+1/63,1,257-225)) ones(1,224-161) (linspace(1,0,160-95)) zeros(1,96)];
green = [zeros(1,31) (linspace(0,1,65)) ones(1,224-161) (linspace(1,0,160-95)) zeros(1,32)];
colorMap = [red; green; blue]';


fh = figure();
fh.WindowState = 'maximized';

kymo = [1];
    for i = 1:161
        clear Cmap;
        if ismember(i,[155,156,157,158,159,160,161])
        Cval = (max(min(dyn(1),Vrange(2)),Vrange(1))-Vrange(1))/(Vrange(2)-Vrange(1));
        else
        Cval = (max(min(dyn(i),Vrange(2)),Vrange(1))-Vrange(1))/(Vrange(2)-Vrange(1));
        end
        Cmap(1,:) = red(round(255*Cval+1))*255*ones(1,Nsec(i));
        Cmap(2,:) = green(round(255*Cval+1))*255*ones(1,Nsec(i));
        Cmap(3,:) = blue(round(255*Cval+1))*255*ones(1,Nsec(i));
        Cmap(4,:) = 1;
        Cmap = uint8(Cmap);
        if ismember(i,kymo)
            for j =NC(i)+1:NC(i+1)+1
            if j == 1
            bonuspoint = [0	10.1999998092651	0	4];
            p = plot3([(Psec(j,1)+bonuspoint(1))/2 (Psec(j,1)+Psec(j+1,1))/2],[(Psec(j,2)+bonuspoint(2))/2 (Psec(j,2)+Psec(j+1,2))/2],[(Psec(j,3)+bonuspoint(3))/2 (Psec(j,3)+Psec(j+1,3))/2],'Color',[0 0 0.5]);
            p.LineWidth = (Psec(j,4)+bonuspoint(4))/2;
            else
            p = plot3([(Psec(j,1)+Psec(j-1,1))/2 (Psec(j,1)+Psec(j+1,1))/2],[(Psec(j,2)+Psec(j-1,2))/2 (Psec(j,2)+Psec(j+1,2))/2],[(Psec(j,3)+Psec(j-1,3))/2 (Psec(j,3)+Psec(j+1,3))/2],'Color', [0 0 0.5]);
            p.LineWidth = Psec(j,4);
            end
            hold on;
            end
        else
            p = plot3(Psec(NC(i)+1:NC(i+1),1),Psec(NC(i)+1:NC(i+1),2),Psec(NC(i)+1:NC(i+1),3),'k-');
            if i == 156  
            p.LineWidth = Psec(NC(i)+1,4)*5;
            elseif i == 157  
            p.LineWidth = Psec(NC(i)+1,4)*5;
            elseif i == 158  
            p.LineWidth = Psec(NC(i)+1,4)*5;
            else
            p.LineWidth = Psec(NC(i)+1,4);
            end
            drawnow
            set(p.Edge, 'ColorBinding','interpolated', 'ColorData',Cmap);
            hold on;
        end
        hold on;
    end
    a = colorbar;
    a.FontSize = 21;
    a.Title.Rotation = 270.0;
    a.Title.FontSize = 21;
    a.Title.Position = [60 225 0];
    colormap(colorMap);
    caxis(Vrange);
    view(90,90);

    xlabel('Distance from Soma (µm)');
    ylabel('Distance from Soma (µm)');
    set(gca,'FontSize',21);
    pbaspect([450, 1200, 70]);
    pause(0.1);
    hold off;

clear a dyn1 dyn2 dyn3 p Cval blue red green dt VrangeR VrangeG VrangeB ts tim B G R t v fh i frame colorMap Cmap allnoaxon noaxon Vrange dyn;
```
