## Supplementary NEURON models for "Dendritic excitations govern back-propagation via a spike-rate accelerometer": plot_results.py

```
import numpy as np
import matplotlib.pyplot as plt

def plot_results(time, v, distance_matrix, duration, section_names, axon):
    plt.figure()

    AXON_SUBSTRINGS = ["inode", "node", "axon", "hill", "iseg"]
    if axon == False:
        # Filter out axon sections based on their names
        v = [v[i] for i in range(len(section_names)) if not any(sub in section_names[i] for sub in AXON_SUBSTRINGS)]
        distances_from_soma = [distance_matrix[0][i] for i in range(len(section_names)) if not any(sub in section_names[i] for sub in AXON_SUBSTRINGS)]

    # Pair each row of v with its corresponding distance
    v_with_distances = zip(v, distances_from_soma)

    # Sort the pairs by distance
    sorted_v_with_distances = sorted(v_with_distances, key=lambda x: x[1])

    # Unzip to get the sorted v values and distances
    sorted_v, _ = zip(*sorted_v_with_distances)

    cmap = plt.get_cmap('jet', len(sorted_v))
    for idx, values in enumerate(sorted_v):
        plt.plot(time[0:len(sorted_v[0])], values, color=cmap(idx))
    plt.ylabel("Voltage (mV)")
    plt.xlabel("Time (ms)")
    plt.xlim([0, duration])
    #plt.xlim([25, 50])
    plt.show()

def kymograph(time, v, distance_matrix, kymomod):
    
    Dmod = np.array(distance_matrix[0])
    volmod = np.array(v)

    #kymomod = [0, 1, 75, 77, 81, 89, 91, 95, 103, 105, 107, 111, 113, 115, 119, 127, 129, 131, 133, 141, 143, 144, 145]

    selected_Dmod = [Dmod[i] for i in kymomod]

    max_distance = int(np.max(selected_Dmod))
    vol_kymo = np.zeros((max_distance, len(time)))

    # Sort distances
    D_sorted, D_order = np.sort(Dmod[kymomod]), np.argsort(Dmod[kymomod])

    # Construct the kymograph
    for nr_idx in range(1, len(D_order)):
        nr = D_order[nr_idx]
        for t in range(len(time)):
            start_idx = int(Dmod[kymomod[nr - 1]])
            end_idx = int(Dmod[kymomod[nr]]) if nr_idx != len(D_order) - 1 else max_distance
            vol_kymo[start_idx:end_idx, t] = np.linspace(volmod[kymomod[nr - 1], t], volmod[kymomod[nr], t], end_idx - start_idx)

    # Plot the kymograph
    plt.figure()
    posit = np.arange(0, max(D_sorted) - 1)
    plt.imshow(vol_kymo, aspect='auto', extent=[time[0], time[-1], posit[-1], posit[0]]) # Note the reversed order of posit
    plt.colorbar(label='Membrane Potential (mV)')
    plt.ylabel("Distance to soma (μm)")
    plt.xlabel("Time (ms)")
    plt.title("Voltage kymograph")
    plt.ylim(max_distance, 0) 
    plt.grid(False)
    #plt.xlim([0, duration])
    #plt.xlim([20, 60])

    plt.show()

    distances = list(range(max_distance))
    voltages = [vol_kymo[i, :].tolist() for i in range(max_distance)]

    # Plot individual voltage traces
    plt.figure()
    cmap = plt.get_cmap('jet', len(kymomod)) # Define the colormap
    for i, idx in enumerate(kymomod):
        color = cmap(i) # Get the color for the current trace
        plt.plot(time, volmod[idx, :], color=color, label=f"Distance {Dmod[idx]}")
    plt.xlabel("Time (ms)")
    #plt.xlim([0, duration])
    #plt.xlim([20, 60])
    plt.ylabel("Membrane Potential (mV)")
    plt.show()

    return distances, voltages

def kymograph_stim(time, v, distance_matrix,kymomod):
    
    Dmod = np.array(distance_matrix[0])
    volmod = np.array(v)

    #kymomod = [0, 1, 75, 77, 81, 89, 91, 95, 103, 105, 107, 111, 113, 115, 119, 127, 129, 131, 133, 141, 143, 144, 145]

    selected_Dmod = [Dmod[i] for i in kymomod]

    max_distance = int(np.max(selected_Dmod))
    vol_kymo = np.zeros((max_distance, len(time)))

    # Sort distances
    D_sorted, D_order = np.sort(Dmod[kymomod]), np.argsort(Dmod[kymomod])

    # Construct the kymograph
    for nr_idx in range(1, len(D_order)):
        nr = D_order[nr_idx]
        for t in range(len(time)):
            start_idx = int(Dmod[kymomod[nr - 1]])
            end_idx = int(Dmod[kymomod[nr]]) if nr_idx != len(D_order) - 1 else max_distance
            vol_kymo[start_idx:end_idx, t] = np.linspace(volmod[kymomod[nr - 1], t], volmod[kymomod[nr], t], end_idx - start_idx)

    # Plot the kymograph
    plt.figure()
    posit = np.arange(0, max(D_sorted) - 1)
    plt.imshow(vol_kymo, aspect='auto', extent=[time[0], time[-1], posit[-1], posit[0]]) # Note the reversed order of posit
    plt.colorbar(label='Channelrhodopsin Conductance (S/cm2)')
    plt.ylabel("Distance to soma (μm)")
    plt.xlabel("Time (ms)")
    plt.ylim(max_distance, 0) 
    plt.grid(False)
    plt.title("Optogenetic stimulation pattern")
    plt.show()

    distances = list(range(max_distance))
    voltages = [vol_kymo[i, :].tolist() for i in range(max_distance)]

    return distances, voltages
```
