## Supplementary NEURON models for "Dendritic excitations govern back-propagation via a spike-rate accelerometer": Readme_NEURON.docx

**Dendritic voltage imaging maps the biophysical basis of plateau potentials in the hippocampus**

\# Readme for NEURON model of bAPs in CA1 pyramidal cells

\# Daniel Itkis,

\# Adam Cohen,

\# 3 March 2024, Cohen Lab (Harvard University)

\## Description

This repository contains code for simulating backpropagating action potentials (bAPs) in CA1 pyramidal cells using the NEURON simulation environment. This model uses biophysically realistic properties of a CA1 pyramidal neuron and is derived from a model by Jarsky & Spruston^1^ (ModelDB accession number: 116084), with updates of ion channel parameters, tuning of spatial distributions, adding slow Na_V_ inactivation, and inclusion of channelrhodopsin for optogenetic stimulation.

The ion channels in the model are:

Slowly inactivating Voltage-gated sodium (Nav), Delayed -rectifier potassium (Kdr) and two types of A-type potassium (Kap proximal, Kad distal) channels. Channelrhodopsin stimulation is implemented using a time-varying distributed conductance with 0 mV reversal potential (ChR).

Detailed equations for the dynamics of the channels can be found in their respective .mod files:

na3.mod, kdrca1.mod, kap.mod, kad.mod, chr.mod

\## Overview of the role of each type of channel in the model:

- Nav Channels (Nav): These are voltage-gated sodium channels, which drive the upstroke of action potentials. Slow inactivation of Nav in the distal dendrites causes bAP failures during sustained stimulation.
- Kap, Kad, and Kdr Channels (Kap, Kad, Kdr): These are voltage-gated potassium channels.

Kap and Kad channels are A-type potassium channels, which are fast-acting and contribute to the transient outward current. They are partially engaged at resting potential and rapidly inactivate during depolarization, either from distal inputs or from failed bAPs.

- Kdr channels are delayed rectifier potassium channels that are needed for repolarization after an AP and contribute to the neuron's ability to fire repetitively.
- Channelrhodopsin Channels (ChR): This is a model of the light-gated ion channel, CheRiff. The model allows for varying the conductance densities of ChR channels to model different illumination patterns and intensities.

\## File List

The following files should be included in the repository to reproduce the simulations:

1. **Ion channel mod files:** The NMODL files that define the dynamics of the Na_V_, K_V_, K_DR_, and ChR channels. Other ion channels such as additional types of Na_V_ and K_V_, VGCC and various synapses (NMODL in directory ChannelsUnused) can be added to the model by setting their densities in the model.

2. **Morphology file:** The file that details the morphology of the CA1 pyramidal neuron. “Jarsky” uses morphology from the original Jarsky & Spruston model. Replacing Morphology file in main.py with a different name and placing the appropriate .swc file in the same directory will run the simulation with the new morphology.

3. **Python run_simulation.py:** The Python script that sets the parameters, runs the simulation, and returns the results.

4. **Python run_morphology.py:** Complementary file that returns morphology data about the loaded neuron for various plots in Matlab.

5. **Python FullCA1Model.py:** The Python Class definition that loads the morphology, distributes channels and sets model properties.

\## How to Use

1. Install Python (3.10) and NEURON (8.2). (Optional) Install Matlab (2023b) to plot simulation results.

2. (Optional) Before running the script, replace 'path_to_your_python_script' with the actual path to the Python script on your system.

3. Compile the .mod files using NEURON's 'nrnivmodl' or 'mknrndll' command.

4. To run the simulation, execute the main.py python script. The script initializes parameters and includes a for loop to run batch simulations. Parameters that vary from run to run such as simulation duration, stimulation start and end times, and ion channel densities are contained in parameter_list.py. It then calls Python functions in run_simulation.py, run_morphology.py and other helper functions to run the simulation and retrieve the morphology of the model neuron.

The simulation results are processed and visualized in MATLAB.

MATLAB scripts to generate various plots are included in
“.\Plot Channel Distributions\Figure_S19a-d\Plot_Channel_Distributions.m”
“.\Oblique Patch Clamp and Optogenetic Stimulus\Figures\Figure_S20a_PatchStimulusShort\ Plot_Voltage_Traces.m”

“.\Oblique Patch Clamp and Optogenetic Stimulus\Figures\Figure_S20b_OptopatchStimulus\ Plot_Voltage_Traces.m”

“.\Model Robustness\Figures\Figure_S20b-d\Plot_Phase_Diagrams.m”

\## Parameters

Parameters for different ion channels and their densities can be modified to fit specific experimental conditions.

The parameters are structured as follows: {gsoma, gdistal, d1/2, z}

The default ion channel parameters are fine tuned to reproduce experimental results.

Channelrhodopsin (Chr) parameters can be modified with a stimulation start (stim_start) and end time (stim_end).

These parameters {gsoma, gdistal, d1/2, z} specify how the density of ion channels changes over the spatial domain of a neuron section.

- gsoma sets the channel density in S/cm^2^ at d 🡪 -∞. For channelrhodopsin stimulation, the reference position for d = 0 (typically section 0, i.e. the soma) can be selected in the Chr_params[0][6] in parameters_list.py.
- gdistal sets the channel density in S/cm^2^ at at d 🡪 ∞.
- d1/2 sets the distance in μm at which the channel density is halfway between gsoma and gdistal.
- z sets the length constant in μm for decay of g(d) towards either gsoma or gdistal.

This parameter structure models ion channel densities as continuous and smooth sigmoid distributions.

1.	Jarsky, T., Roxin, A., Kath, W. L. & Spruston, N. Conditional dendritic spike propagation following distal synaptic activation of hippocampal CA1 pyramidal neurons. *Nature Neuroscience* **8**, 1667–1676 (2005).
