## Supplementary NEURON models for "Dendritic excitations govern back-propagation via a spike-rate accelerometer": run_morphology.py

```
import numpy as np
from neuron import h, gui
from FullCA1Model import FullCA1Model

def run_morphology(channel_params, morphology_file, sampling):
    # Create a model using the provided channel parameters
    model = FullCA1Model(channel_params, morphology_file)

    if sampling == "sections":
        positions = []
        distance_matrix = [
        np.array([h.distance(section(0.5), second_section(0.5)) for second_section in h.allsec()])
        for section in h.allsec()
        ]
        section_count = sum(1 for _ in h.allsec())
        section_nums = [0] * section_count
        positions = []
        for index, section in enumerate(h.allsec()):
            section_nums[index] = section.n3d()
            for k in range(section_nums[index]):
                position_info = [
                    section.x3d(k),
                    section.y3d(k),
                    section.z3d(k),
                    section.diam3d(k),
                    index,
                    index,
                ]
                positions.append(position_info)
        section_names = []  
        for section in h.allsec():
            section_names.append(section.name())

    elif sampling == "segments":
        positions = []
        distance_matrix = []
        section_names = []

        # Initialize segment index
        segment_index = 0

        # Loop through each section to populate 'positions' and 'section_names'
        seg_global = 0
        for index, section in enumerate(h.allsec()):
            n3d = section.n3d()
            nseg = section.nseg
            section_length = section.L
            
            # Loop through n3d and find arc3d for each point
            seg_max = 0
            for i in range(n3d):
                arc3d_val = section.arc3d(i)
                
                # Assign it to a segment based on arc3d
                seg_idx = int(arc3d_val / section_length * nseg) 
                seg_max = max(seg_max, seg_idx)
                
                position_info = [
                    section.x3d(i),
                    section.y3d(i),
                    section.z3d(i),
                    section.diam3d(i),
                    index,
                    seg_global + seg_idx,
                ]
                positions.append(position_info)
            
            seg_global = seg_global + seg_max 

        # Populate 'section_names'
        for section in h.allsec():
            for seg in section:
                section_names.append(f"{section.name()}({seg.x})")

        # Initialize distance_matrix as an empty list
        distance_matrix = []

        # Loop through all sections and segments to populate 'distance_matrix'
        for section1 in h.allsec():
            for seg1 in section1:
                row = []
                for section2 in h.allsec():
                    for seg2 in section2:
                        d = h.distance(seg1, seg2)  # Replace with your actual distance function
                        row.append(d)
                distance_matrix.append(row)

        # Convert distance_matrix to NumPy array for easier manipulation
        distance_matrix = np.array(distance_matrix)
        # Convert distance_matrix to NumPy array for easier manipulation
        distance_matrix = np.array(distance_matrix)
  
    else: 
        # Initialize distance_matrix as an empty list
        distance_matrix = []
        # Initialize a list to store the points that should be sampled
        sampling_points = []
        # Initialize a list to store the 3D positions of the sampling points
        sampling_positions = []
        # Start at the soma (assuming soma is the first section)
        soma_sec = next(h.allsec())
        sampling_points.append((soma_sec, 0.5))
        closest_idx = nearest_idx(soma_sec, 0.5)
        sampling_positions.append([soma_sec.x3d(closest_idx), soma_sec.y3d(closest_idx), soma_sec.z3d(closest_idx)])
        added_points = set()
        added_points.add((soma_sec, 0.5))

        while sampling_points:
            sec1, loc1 = sampling_points.pop(0)

            internal_points = []
            if (loc1*sec1.L - sampling) > 0:
                new_point = (sec1, loc1-sampling/sec1.L)
                internal_points.append(new_point)
            if (loc1*sec1.L + sampling) < sec1.L:
                new_point = (sec1, loc1+sampling/sec1.L)
                internal_points.append(new_point)

            for point in internal_points:
                point_sec, point_loc = point
                if (point not in added_points):
                        # Check if this point is closer than 'sampling'/10 um to any existing point
                        too_close = False
                        for existing_point in added_points:
                            sec_existing, loc_existing = existing_point
                            distance_to_existing = h.distance(point_sec(point_loc), sec_existing(loc_existing))
                            
                            if distance_to_existing < sampling/10:
                                too_close = True
                                break
                        
                        if not too_close:
                            added_points.add(point)
                            sampling_points.append(point)
                            
                            # Record the 3D position of this sampling point
                            closest_idx = nearest_idx(sec2, loc2)
                            x, y, z = sec2.x3d(closest_idx), sec2.y3d(closest_idx), sec2.z3d(closest_idx)
                            sampling_positions.append([x, y, z])

            for sec2 in h.allsec():
                distance_to_start = h.distance(sec1(loc1), sec2(0))
                distance_to_end = h.distance(sec1(loc1), sec2(1))
                
                if (distance_to_start-sampling)*(distance_to_end-sampling) < 0:
                    
                    # Interpolate to find the exact location within the section
                    loc2 = (sampling - distance_to_start) / (distance_to_end - distance_to_start)
                    # Create a tuple to represent this point
                    point = (sec2, loc2)
                    
                    # Check if this point has already been added
                    if point not in added_points:
                        # Check if this point is closer than 'sampling' um to any existing point
                        too_close = False
                        for existing_point in added_points:
                            sec_existing, loc_existing = existing_point
                            distance_to_existing = h.distance(sec2(loc2), sec_existing(loc_existing))
                            
                            if 0 < distance_to_existing < sampling:
                                too_close = True
                                break
                        
                        if not too_close:
                            added_points.add(point)
                            sampling_points.append(point)
                            
                            # Record the 3D position of this sampling point
                            closest_idx = nearest_idx(sec2, loc2)
                            x, y, z = sec2.x3d(closest_idx), sec2.y3d(closest_idx), sec2.z3d(closest_idx)
                            sampling_positions.append([x, y, z])        

        # Initialize an empty list to store the distance matrix
        distance_matrix = []

        # Loop through each pair of points in added_points
        for point1 in added_points:
            sec1, loc1 = point1
            distance_row = []
            for point2 in added_points:
                sec2, loc2 = point2
                distance = h.distance(sec1(loc1), sec2(loc2))
                distance_row.append(distance)
            distance_matrix.append(distance_row)

        # Convert the list of lists to a NumPy array for easier manipulation
        distance_matrix = np.array(distance_matrix)

        # Collect 3D position information
        section_count = sum(1 for _ in h.allsec())
        section_nums = [0] * section_count
        positions = []
        sampling_indices = {}  # To store the nearest sampling point index for each section
        for index, section in enumerate(h.allsec()):
            section_nums[index] = section.n3d()
            # Create points every 'sampling' um along section
            num_points = int(section.L / sampling) + 1
            for i in range(num_points):
                location = i / max((num_points - 1), 1)
                
                # Find the nearest 3D point to this location
                min_distance = float('inf')
                nearest_k = 0
                for k in range(section_nums[index]):
                    x, y, z = section.x3d(k), section.y3d(k), section.z3d(k)
                    distance = ((x - section.x3d(0))**2 + (y - section.y3d(0))**2 + (z - section.z3d(0))**2)**0.5
                    if abs(distance - section.L * location) < min_distance:
                        min_distance = abs(distance - section.L * location)
                        nearest_k = k
                
                # Record the nearest sampling point index
                sampling_indices[section, location] = nearest_k
                
                position_info = [
                    section.x3d(nearest_k),
                    section.y3d(nearest_k),
                    section.z3d(nearest_k),
                    section.diam3d(nearest_k),
                    index,
                    sampling_indices[section, location],
                ]
                positions.append(position_info)

    if sampling == "sections":
        path = longest_path_from_soma_sections(distance_matrix,section_names)
    elif sampling == "segments":
        path = longest_path_from_soma_segments(distance_matrix,section_names)
    else:
        path = longest_path_from_soma_two_pass(distance_matrix,section_names,sampling)

    return section_names, distance_matrix, positions, path

def longest_path_from_soma_sections(distance_matrix, section_names):
    n = len(distance_matrix)
    AXON_SUBSTRINGS = ["inode", "node", "axon", "hill", "iseg"]

    def is_axon(section_index):
        """Check if a section is an axon based on its name."""
        return any(sub in section_names[section_index] for sub in AXON_SUBSTRINGS)

    # 1. Find the farthest section from the soma
    farthest_section = -1
    max_distance = -1
    for i in range(1, n):  # Start from 1 to exclude soma itself
        if not is_axon(i) and distance_matrix[0][i] > max_distance:
            max_distance = distance_matrix[0][i]
            farthest_section = i

    path = [farthest_section]
    current_section = farthest_section

    while current_section != 0:  # Loop until we reach the soma
        # 2. Find the neighbors of the current section
        neighbors = [(i, distance_matrix[current_section][i]) for i in range(n) 
                     if distance_matrix[current_section][i] > 0 and i != current_section and not is_axon(i)]
        
        # Sort neighbors based on distance from current section
        neighbors.sort(key=lambda x: x[1])

        added_to_path = False

        # 3. & 4. Check each neighbor
        for (neighbor, distance) in neighbors:
            if abs(distance_matrix[0][neighbor] + distance_matrix[neighbor][current_section] - distance_matrix[0][current_section]) < 1e-9:
                path.append(neighbor)
                current_section = neighbor
                added_to_path = True
                break

        # If we didn't find a valid neighbor, backtrack
        if not added_to_path:
            path.pop()
            current_section = path[-1]

    # 5. Repeat until you reach the soma
    path.reverse()
    return path

def longest_path_from_soma_segments(distance_matrix, section_names):
    n = len(distance_matrix)
    AXON_SUBSTRINGS = ["inode", "node", "axon", "hill", "iseg"]

    def is_axon(segment_index):
        """Check if a segment is an axon based on its name."""
        return any(sub in section_names[segment_index] for sub in AXON_SUBSTRINGS)

    # 1. Find the farthest segment from the soma
    farthest_segment = -1
    max_distance = -1
    for i in range(1, n):  # Start from 1 to exclude soma itself
        if not is_axon(i) and distance_matrix[0][i] > max_distance:
            max_distance = distance_matrix[0][i]
            farthest_segment = i

    path = [farthest_segment]
    current_segment = farthest_segment

    while current_segment != 0:  # Loop until we reach the soma
        # 2. Find the neighbors of the current segment
        neighbors = [(i, distance_matrix[current_segment][i]) for i in range(n) 
                     if distance_matrix[current_segment][i] > 0 and i != current_segment and not is_axon(i)]
        
        # Sort neighbors based on distance from current segment
        neighbors.sort(key=lambda x: x[1])

        added_to_path = False

        # 3. & 4. Check each neighbor
        for (neighbor, distance) in neighbors:
            if abs(distance_matrix[0][neighbor] + distance_matrix[neighbor][current_segment] - distance_matrix[0][current_segment]) < 1e-9:
                path.append(neighbor)
                current_segment = neighbor
                added_to_path = True
                break

        # If we didn't find a valid neighbor, backtrack
        if not added_to_path:
            path.pop()
            current_segment = path[-1]

    # 5. Repeat until you reach the soma
    path.reverse()
    return path


def longest_path_from_soma_two_pass(distance_matrix, section_names, sampling):
    n = len(distance_matrix[0])
    AXON_SUBSTRINGS = ["inode", "node", "axon", "hill", "iseg"]

    def is_axon(section_index):
        """Check if a section is an axon based on its name."""
        return any(sub in section_names[section_index] for sub in AXON_SUBSTRINGS)

    # Find the farthest section from the soma using the original method
    farthest_section = -1
    max_distance = -1
    for i in range(1, n):  # Start from 1 to exclude soma itself
        if not is_axon(i) and distance_matrix[0][i] > max_distance:
            max_distance = distance_matrix[0][i]
            farthest_section = i

    # Find the furthest point from the soma using the new method
    furthest_point_idx = np.argmax(distance_matrix[0, :])

    # Choose the furthest point based on both methods
    current_point_idx = furthest_point_idx if furthest_point_idx > farthest_section else farthest_section

    # Initialize the longest path with the furthest point
    longest_path = [current_point_idx]

    # Loop to build the longest path
    while current_point_idx != 0:
        # Calculate the desired next distance
        desired_next_distance = distance_matrix[0, current_point_idx] - sampling
        
        # Find the point that is closest to the desired next distance and closer to the soma
        next_point_idx = np.argmin(np.abs(distance_matrix[0, :current_point_idx] - desired_next_distance))
        
        # Add this point to the longest path
        longest_path.append(next_point_idx)
        
        # Update the current point
        current_point_idx = next_point_idx

    # Reverse the path to start from the soma
    longest_path.reverse()
    return longest_path


def all_paths_from_soma_to_terminals(distance_matrix):
    n = len(distance_matrix)
    visited = [False] * n
    all_paths = []

    def is_terminal(section):
        """Check if a section is terminal."""
        for i in range(n):
            if not visited[i] and distance_matrix[section][i] > 0:
                return False
        return True

    def dfs(curr_section, curr_path):
        nonlocal all_paths
        visited[curr_section] = True
        curr_path.append(curr_section)

        if is_terminal(curr_section):
            all_paths.append(curr_path.copy())
        else:
            for i in range(n):
                if not visited[i] and distance_matrix[curr_section][i] > 0:
                    dfs(i, curr_path.copy())

        visited[curr_section] = False

    dfs(0, [])

    all_paths.sort(key=len, reverse=True)
    return all_paths

def nearest_idx(section, loc):
    min_distance = float('inf')
    nearest_k = 0
    n3d = section.n3d()
    
    for k in range(n3d):
        arc_distance = section.arc3d(k)
        distance = abs(arc_distance - loc)
        
        if distance < min_distance:
            min_distance = distance
            nearest_k = k
    
    return nearest_k
```
