## Supplementary NEURON models for "Dendritic excitations govern back-propagation via a spike-rate accelerometer": run_simulation.py

```
from neuron import h, gui
from FullCA1Model import FullCA1Model
import numpy as np
from scipy.optimize import fsolve

def uniform_cdf(x):
    return x
    
def sqrt_cdf(x):
    return x**1/2

def run_simulation(sim_params, channel_params, synapse_params_list, waveform, record_vars, morphology_file, patch_params, distance_matrix, sampling):

    ka_cutoff = 100 # switch from proximal to distal A-type K at 100 um
    model = FullCA1Model(channel_params, morphology_file, ka_cutoff)

    # Simulation parameters
    h.tstop = sim_params[0]
    h.dt = sim_params[1]
    h.celsius = 35 # Temperature in celsius

    # Initialize as empty lists
    all_synapses = []
    all_syn_positions = []
    all_netstims = []
    all_netcons = []
    all_spike_times = []

    for synapse_params in synapse_params_list:
        synapse_nr = int(round(synapse_params[0]))
        start_time = synapse_params[1]
        end_time = synapse_params[2]
        spike_rate = synapse_params[3]
        synapse_strength = synapse_params[4]
        distribution_type = synapse_params[5]
        location = synapse_params[6]
        synapse_type = synapse_params[7]
        try:
            ratio = synapse_params[8] 
        except IndexError: 
            ratio = 0.1
            
        [synapses, syn_positions] = model.distribute_synapses(synapse_nr, uniform_cdf, location, synapse_type)
        [netstims, netcons, spike_times] = model.distribute_spike_times(synapses, start_time, end_time, spike_rate, synapse_strength, distribution_type, ratio)

        # Append to the lists
        all_synapses.extend(synapses)
        all_syn_positions.extend(syn_positions)
        all_netstims.extend(netstims)
        all_netcons.extend(netcons)
        all_spike_times.extend(spike_times)

    # Initialize dictionaries to hold recording vectors for each variable
    rec_t = h.Vector()
    rec_t.record(h._ref_t)
    rec_vars = {var: [] for var in record_vars}

    # Iterate over all sections
    if sampling == "sections":
        for sec in h.allsec():
            for var in record_vars:
                vec = h.Vector()
                try:
                    vec.record(getattr(sec(0.5), '_ref_' + var))
                except AttributeError:
                    print(f"Error: {var} does not exist in {sec.name()}")
                    continue
                rec_vars[var].append(vec)
    elif sampling == "segments":
        for sec in h.allsec():
            for seg in sec:
                for var in record_vars:
                    vec = h.Vector()
                    try:
                        vec.record(getattr(seg, '_ref_' + var))
                    except AttributeError:
                        print(f"Error: {var} does not exist in {sec.name()}")
                        continue
                    rec_vars[var].append(vec)


    # Add Vectorplay stimulation 
    #stims, stimulation, somsec = model.chr_vectorplay()
    #stimulation_out = np.array(stimulation)
    
    num_sections = sum(1 for _ in h.allsec())
    times = np.arange(0, h.tstop + h.dt, h.dt)          
    gmax_values = waveform.tolist()
    gmax_values_2D = np.tile(gmax_values, (num_sections, 1))
    model.prepare_simulation(times, gmax_values_2D)    
    [clamp, recorded_amp] = model.add_patch(patch_params[0], patch_params[1], patch_params[2])
    recorded_curr = h.Vector()
    recorded_curr.record(clamp[0]._ref_i)
 
    total_g_chr = 0.0

    # Iterate over all sections
    for sec in h.allsec():
        # Iterate over all segments in the current section
        for seg in sec:
            segment_conductance = seg.gexp_chr * h.area(seg.x) / 1e8 # S
            total_g_chr += segment_conductance
    total_g_chr = total_g_chr * max(waveform)

    print(f"Total conductance g_ChR: {round(total_g_chr*1e9,2)} nS")
    print(f"Total peak current I_ChR: {round(total_g_chr*70*1e9,2)} pA")
    
    # Initialize and run the simulation
    h.finitialize(-73)
    h.v_init = -73 # mV
    h.run()

    # Convert recorded data to NumPy arrays
    time = np.array(rec_t)
    record_arrays = {var: [np.array(vec) for vec in vec_list] 
                     for var, vec_list in rec_vars.items()}
    
    all_spike_times_frames = [[round(time / h.dt) for time in inner_list] for inner_list in all_spike_times]
        
    return time, record_arrays, all_syn_positions, all_spike_times, all_spike_times_frames

def reshape_v(v, distance_matrix):
    n = len(distance_matrix)
    length_of_subarray = len(v) // n
    reshaped_v = [v[i * length_of_subarray: (i + 1) * length_of_subarray] for i in range(n)]
    return reshaped_v

def generate_waveform(sim_params, stimmax, stim_start, stim_end, protocol):
    if protocol == "full": # Pulses then ramp
        # Stimulation parameters
        stim_duration = stim_end - stim_start
        scale = stim_duration / 260
        times = np.arange(0, sim_params[0] + sim_params[1], sim_params[1])
        t_down = 20 * scale  # ms
        t_up = 20 * scale  # ms
        t_ramp = 75 * scale  # ms

        n_start = int(stim_start / sim_params[1])
        n_down = int(t_down / sim_params[1])
        n_up = int(t_up / sim_params[1])
        n_ramp = int(t_ramp / sim_params[1])
        zeros_start = np.zeros(n_start)
        zeros_down = np.zeros(n_down)
        ramp_up_1 = np.ones(n_up) * stimmax / 3
        ramp_up_2 = np.ones(n_up) * stimmax * 2 / 3
        ramp_up_3 = np.ones(n_up) * stimmax
        ramp = np.linspace(0, stimmax, n_ramp)

        # Combine them to form the complete waveform
        waveform = np.concatenate([zeros_start, zeros_down, ramp_up_1, zeros_down, ramp_up_2, zeros_down, ramp_up_3, np.tile(zeros_down, 2), ramp, zeros_down])
    
    elif protocol == "step":
        times = np.arange(0, sim_params[0] + sim_params[1], sim_params[1])
        waveform = np.ones(len(times)) * sim_params[5] 
        waveform[int(stim_start / sim_params[1]):int(stim_end / sim_params[1])] = stimmax

    elif protocol == "ramp":
        times = np.arange(0, sim_params[0] + sim_params[1], sim_params[1])
        waveform = np.ones(len(times)) * sim_params[5]
        waveform[int(stim_start / sim_params[1]):int(stim_end / sim_params[1])] = np.linspace(sim_params[5], sim_params[5]+stimmax, round((stim_end - stim_start) / sim_params[1]))

    elif protocol == "pulses":
        baseline = sim_params[5] 
        pulse_dur = sim_params[2] 
        pulse_nr = sim_params[4]
        pulse_interval = sim_params[3] 

        times = np.arange(0, sim_params[0] + sim_params[1], sim_params[1])
        waveform = np.ones(len(times))*baseline

        for pulse in range(pulse_nr):
            start_idx = int(pulse * pulse_interval / sim_params[1])
            end_idx = int(start_idx + pulse_dur / sim_params[1])
            if end_idx < len(waveform): 
                waveform[start_idx+int(stim_start / sim_params[1]):end_idx+int(stim_start / sim_params[1])] = waveform[start_idx+int(stim_start / sim_params[1]):end_idx+int(stim_start / sim_params[1])] + stimmax

    if len(waveform) < len(times):
        waveform = np.concatenate([waveform, np.zeros(len(times) - len(waveform))])
    elif len(waveform) > len(times):
        waveform = waveform[:len(times)]
        
    return waveform
```
