## Supplementary NEURON models for "Dendritic excitations govern back-propagation via a spike-rate accelerometer": v_amplification_ratio.py

```
from mpl_toolkits.mplot3d import Axes3D
from scipy.stats import linregress
import numpy as np
import matplotlib.pyplot as plt
import random
from scipy.stats import gaussian_kde

def analyze_neuron_bAP(voltage, s_na3,h_na3, gbar_na3, l_kad, gkabar_kad, distance_matrix):
    def find_closest_neighbors_towards_soma(distance_matrix):
        closest_neighbors = [-1]  # -1 for the soma itself as it doesn't have a neighbor towards the soma
        for i in range(1, len(distance_matrix)):
            min_distance = float('inf')
            closest_neighbor = -1
            for j in range(len(distance_matrix)):
                if distance_matrix[i][j] < min_distance and distance_matrix[0][j] < distance_matrix[0][i]:
                    min_distance = distance_matrix[i][j]
                    closest_neighbor = j
            closest_neighbors.append(closest_neighbor)
        return closest_neighbors

    closest_neighbors = find_closest_neighbors_towards_soma(distance_matrix)
    # for i, section in enumerate(closest_neighbors):
    #     print(f"Section {i} has closest neighbor {section}.")

    equilibration_time = int(25/0.025)  # Assuming the equilibration time is 25 ms
    #stim_end_time = 4000#int(70/0.025)  # Assuming the stimulus ends at 70 ms

    t_max_list = [[] for _ in range(len(closest_neighbors))]
    t_init_list = [[] for _ in range(len(closest_neighbors))]

    for i, section in enumerate(closest_neighbors):
        if section == -1:  # Skip the soma
            continue
        voltage_series = voltage[section]
        
        for j in range(equilibration_time, len(voltage_series) - 1):  # Start from equilibration_time
            if voltage_series[j] > voltage_series[j-1] and voltage_series[j] > voltage_series[j+1]:
                t_max = j
                t_max_list[i].append(t_max)
                
                if len(t_max_list[i]) > 1:
                    prev_t_max = t_max_list[i][-2]
                    min_voltage_idx = np.argmin(voltage_series[prev_t_max:t_max]) + prev_t_max
                else:
                    min_voltage_idx = np.argmin(voltage_series[equilibration_time:t_max]) + equilibration_time  # Start from equilibration_time
                
                t_init_list[i].append(min_voltage_idx)

 
    amplification_list = []
    ratio_list = []
    ratio2_list = []
    voltage_amplitude_list = []
    
    # New lists for debugging
    debug_sections = []#random.sample(range(1, len(closest_neighbors)), len(closest_neighbors)//50)
    
    for i in range(len(closest_neighbors) - 1):
        parent_section = closest_neighbors[i]
        daughter_section = closest_neighbors[i+1]

        if parent_section == -1 or daughter_section == -1 or gkabar_kad[daughter_section][1] == 0:  # Skip if either is the soma
            continue

        min_length = min(len(t_max_list[parent_section]), len(t_max_list[daughter_section]))

        for j in range(min_length):
            parent_t_max = t_max_list[parent_section][j]
            daughter_t_max = t_max_list[daughter_section][j]

            parent_t_init = t_init_list[parent_section][j]
            daughter_t_init = t_init_list[daughter_section][j]

            voltage_amplitude = voltage[parent_section][parent_t_max] - voltage[parent_section][parent_t_init]

            # Check the condition to ignore certain spikes
            if voltage_amplitude < 10:
                continue

            # Debugging plots
            if parent_section in debug_sections:
                plt.figure()
                plt.plot(voltage[parent_section], label='Voltage')
                plt.scatter([parent_t_max], [voltage[parent_section][parent_t_max]], color='red')  # maxima
                plt.scatter([parent_t_init], [voltage[parent_section][parent_t_init]], color='green')  # minima
                plt.legend()
                plt.title(f"Section {parent_section}")
                plt.show()

            print(voltage[parent_section][parent_t_max] - voltage[parent_section][parent_t_init])
            amplification = ((voltage[daughter_section][daughter_t_max] - voltage[daughter_section][daughter_t_init]) - (voltage[parent_section][parent_t_max] - voltage[parent_section][parent_t_init])) / \
                            (voltage[parent_section][parent_t_max] - voltage[parent_section][parent_t_init])

            min_l_kad = min(l_kad[daughter_section][daughter_t_init:daughter_t_max+1])
            print(daughter_section)
            ratio = (s_na3[daughter_section][daughter_t_init] * h_na3[daughter_section][daughter_t_init] * gbar_na3[daughter_section][1]) - (min_l_kad * gkabar_kad[daughter_section][1])
            ratio2 = (s_na3[daughter_section][daughter_t_init]*gbar_na3[daughter_section][1]) - (min_l_kad * gkabar_kad[daughter_section][1])
            #ratio = (s_na3[daughter_section][daughter_t_init]) / l_kad[daughter_section][daughter_t_init]
            #ratio = (s_na3[daughter_section][daughter_t_init]*h_na3[daughter_section][daughter_t_init]) / l_kad[daughter_section][daughter_t_init]
            ratio_list.append(ratio)
            ratio2_list.append(ratio2)
            amplification_list.append(amplification)
            voltage_amplitude_list.append(voltage_amplitude)


    plt.figure(figsize=(10, 6))
    plt.scatter(ratio_list, amplification_list, c='blue', marker='o', s=10)
    plt.xlabel('(g_na * s_na * h_na) - (g_ka * l_ka) at t_init')
    plt.ylabel('Amplification or Decrement')
    plt.title('Amplification/Decrement vs nash/kad Ratio')
    plt.grid(True)
    plt.show()

    plt.figure(figsize=(10, 6))
    plt.hist2d(ratio_list, amplification_list, bins=(500, 50), cmap='Blues')
    plt.colorbar(label='Density')
    plt.xlabel('(g_na * s_na * h_na) - (g_ka * l_ka) at t_init')
    plt.ylabel('Amplification or Decrement')
    plt.title('Amplification/Decrement vs nash/kad Ratio')
    plt.grid(True)
    plt.show()

    # Calculate the point density
    xy = np.vstack([ratio2_list, amplification_list])
    z = gaussian_kde(xy)(xy)

    # Create grid
    x_min, x_max = np.min(ratio2_list), np.max(ratio2_list)
    y_min, y_max = np.min(amplification_list), np.max(amplification_list)
    x_grid, y_grid = np.mgrid[x_min:x_max:100j, y_min:y_max:100j]
    positions = np.vstack([x_grid.ravel(), y_grid.ravel()])
    values = gaussian_kde(xy)(positions).reshape(x_grid.shape)

    # Create the KDE plot
    plt.figure(figsize=(10, 6))
    plt.contourf(x_grid, y_grid, values, levels=50, cmap='Blues')
    plt.colorbar(label='Density')
    plt.xlabel('(g_na * s_na * h_na) - (g_ka * l_ka) at t_init')
    plt.ylabel('Amplification or Decrement')
    plt.title('Amplification/Decrement vs Na(sh)/KA Ratio')
    plt.grid(True)
    plt.show()


    fig = plt.figure(figsize=(10, 6))
    ax = fig.add_subplot(111, projection='3d')

    ax.scatter(ratio_list, voltage_amplitude_list, amplification_list, c='blue', marker='o')

    ax.set_xlabel('(g_na * s_na * h_na) - (g_ka * l_ka) at t_init')
    ax.set_ylabel('Voltage Amplitude at Parent')
    ax.set_zlabel('Amplification or Decrement')

    plt.title('3D Plot of Amplification/Decrement vs nash/kad Ratio and Voltage Amplitude')
    plt.grid(True)
    plt.show()

    fig = plt.figure(figsize=(10, 6))
    ax = fig.add_subplot(111, projection='3d')

    ax.scatter(ratio2_list, voltage_amplitude_list, amplification_list, c='blue', marker='o')

    ax.set_xlabel('(g_na * s_na) / (g_ka * l_ka) at t_init')
    ax.set_ylabel('Voltage Amplitude at Parent')
    ax.set_zlabel('Amplification or Decrement')

    plt.title('3D Plot of Amplification/Decrement vs nas/kad Ratio and Voltage Amplitude')
    plt.grid(True)
    plt.show()


def analyze_neuron_bAP_2(voltage, path, equilibration_time, distances, dt):
    # Initialize lists to store the maxima, minima, and their time points
    spike_maxima_minima_pairs = []
    current_maximum_time = {}

    # Iterate only along the longest path
    previous_index = path[0]
    for section_index in path:
        voltage_series = voltage[section_index]
        previous_minimum = []
        previous_minimum_time = []
        current_maximum_time = []
        section_distance = distances[previous_index][section_index]
        previous_index = section_index

        # Start time for the first maximum search is the equilibration time
        start_time = equilibration_time

        # Differentiate between first path and subsequent paths
        if section_index == path[0]:
            previous_maximum_time = start_time
            maxima_ref = []
            # Find spikes above -10mV threshold for path[0]
            for time_point in range(int(round(start_time/dt)), len(voltage_series)):  # Start after equilibration_time
                if voltage_series[time_point] > -10 and voltage_series[time_point] > voltage_series[time_point - 1] and voltage_series[time_point] > voltage_series[time_point + 1]:
                    current_maximum = voltage_series[time_point]
                    current_maximum_time = time_point
                    previous_minimum = min(voltage_series[previous_maximum_time:time_point])
                    previous_minimum_time = voltage_series[previous_maximum_time:time_point].index(previous_minimum) + previous_maximum_time
                    spike_maxima_minima_pairs.extend([
                        section_index,
                        section_distance,
                        current_maximum_time * dt,
                        current_maximum,
                        previous_minimum_time * dt,
                        previous_minimum
                    ])
                    previous_maximum_time = time_point
                    maxima_ref.append(time_point)
            print(len(maxima_ref))
            maxima_ref.append(len(voltage_series))

                    
        else:
            # For subsequent paths, find global maxima in the specified intervals
            previous_maximum_time = int(round(start_time/dt))
            for i in range(len(maxima_ref) - 1):
                # Search within a window offset by 5 from the reference maximum points
                search_start = maxima_ref[i] - 5
                search_end = maxima_ref[i+1] - 5
                current_maximum = max(voltage_series[search_start:search_end])
                current_maximum_time = voltage_series[search_start:search_end].index(current_maximum) + search_start
                previous_minimum = min(voltage_series[previous_maximum_time:current_maximum_time])
                previous_minimum_time = voltage_series[previous_maximum_time:current_maximum_time].index(previous_minimum) + previous_maximum_time                
                # Validate the current maximum by checking neighboring points
                if (current_maximum_time - 1 >= 0 and current_maximum_time + 1 < len(voltage_series)
                        and voltage_series[current_maximum_time] > voltage_series[current_maximum_time - 1]
                        and voltage_series[current_maximum_time] > voltage_series[current_maximum_time + 1]):
                    # If validation passes, find the previous minimum
                    spike_maxima_minima_pairs.extend([
                        section_index,
                        section_distance,
                        current_maximum_time * dt,
                        current_maximum,
                        previous_minimum_time * dt,
                        previous_minimum
                    ])
                else:
                    # If validation fails, set the voltage values for the maximum and minimum to 0
                    spike_maxima_minima_pairs.extend([
                        section_index,
                        section_distance,
                        current_maximum_time * dt,
                        0,  # Set maximum to 0
                        previous_minimum_time * dt,
                        0   # Set minimum to 0
                    ])
                # Update the previous maximum time to the current
                previous_maximum_time = current_maximum_time

    # Plot voltage traces for sections specified in path
    for section_index in path:
        plt.figure()
        plt.plot([i * dt for i in range(len(voltage[section_index]))], voltage[section_index], label='Voltage Trace')
        # Extract maxima and minima for the current section
        section_pairs = [spike_maxima_minima_pairs[i:i + 6] for i in range(0, len(spike_maxima_minima_pairs), 6) if spike_maxima_minima_pairs[i] == section_index]
        for pair in section_pairs:
            plt.scatter(pair[2], pair[3], color='red')  # maxima
            if len(pair) > 4:
                plt.scatter(pair[4], pair[5], color='green')  # minima
        plt.title(f"Voltage Trace for Section {section_index}")
        plt.xlabel('Time (ms)')
        plt.ylabel('Voltage (mV)')
        plt.legend()
        plt.show()

    return spike_maxima_minima_pairs
```
